# Worldwide tracing of mutations and the evolutionary dynamics of SARS-CoV-2

**DOI:** 10.1101/2020.08.07.242263

**Authors:** Zhong-Yin Zhou, Hang Liu, Yue-Dong Zhang, Yin-Qiao Wu, Min-Sheng Peng, Aimin Li, David M. Irwin, Haipeng Li, Jian Lu, Yiming Bao, Xuemei Lu, Di Liu, Ya-Ping Zhang

**Affiliations:** State Key Laboratory of Genetic Resources and Evolution, Kunming Institute of Zoology, Chinese Academy of Sciences, Kunming, Yunnan, 650223, China; Kunming College of Life Science, University of Chinese Academy of Sciences, Kunming, Yunnan, 650204, China; Shaanxi Key Laboratory for Network Computing and Security Technology, School of Computer Science and Engineering, Xi’an University of Technology, Xi’an, Shaanxi, 710048, China; Department of Laboratory Medicine and Pathobiology, University of Toronto, Toronto, Canada; CAS Key Laboratory of Computational Biology, CAS-MPG Partner Institute for Computational Biology, Shanghai Institute of Nutrition and Health, University of Chinese Academy of Sciences, Chinese Academy of Sciences, China; Center for Excellence in Animal Evolution and Genetics, Chinese Academy of Sciences, Kunming, 650223, China; State Key Laboratory of Protein and Plant Gene Research, Center for Bioinformatics, School of Life Sciences, Peking University, Beijing 100871, China; National Genomics Data Center, Beijing Institute of Genomics, Chinese Academy of Sciences and China National Center for Bioinformation, Beijing 100101, China; CAS Key Laboratory of Special Pathogens, Wuhan Institute of Virology, Center for Biosafety Mega-Science, Chinese Academy of Sciences, Wuhan, China

## Abstract

Understanding the mutational and evolutionary dynamics of SARS-CoV-2 is essential for treating COVID-19 and the development of a vaccine. Here, we analyzed publicly available 15,818 assembled SARS-CoV-2 genome sequences, along with 2,350 raw sequence datasets sampled worldwide. We investigated the distribution of inter-host single nucleotide polymorphisms (inter-host SNPs) and intra-host single nucleotide variations (iSNVs). Mutations have been observed at 35.6% (10,649/29,903) of the bases in the genome. The substitution rate in some protein coding regions is higher than the average in SARS-CoV-2 viruses, and the high substitution rate in some regions might be driven to escape immune recognition by diversifying selection. Both recurrent mutations and human-to-human transmission are mechanisms that generate fitness advantageous mutations. Furthermore, the frequency of three mutations (S protein, F400L; ORF3a protein, T164I; and ORF1a protein, Q6383H) has gradual increased over time on lineages, which provides new clues for the early detection of fitness advantageous mutations. Our study provides theoretical support for vaccine development and the optimization of treatment for COVID-19. We call researchers to submit raw sequence data to public databases.

---

SARS-CoV-2 is an RNA virus that causes the coronavirus disease 2019 (COVID-19). This disease has spread worldwide, and as of June 26, 2020, has infected more than nine million humans^1^. The whole-genome sequence of SARS-CoV-2 is 96.2% similar to that of a bat SARS-related coronavirus (RaTG13) and is 79% similar to human SARS-CoV^2^. Recent studies have indicated that SARS-CoV-2 is more easily transmitted from person to person than SARS-CoV^3,4^. SARS-CoV-2 has also gradually accumulated new mutations that may make it more suitable to the human host^5,6^.

With the development of next-generation sequencing, it has become possible to conduct large-scale studies to detect inter-host single nucleotide polymorphisms (inter-host SNPs) and intra-host single nucleotide variations (iSNVs) for infectious diseases, such as Ebola virus that can uncover essential information concerning their transmission and evolution^7,8^. Thus, tracing inter-host SNPs and iSNVs, and revealing the evolutionary dynamics of SARS-CoV-2 worldwide is a priority.

To trace mutations in SARS-CoV-2, we first identified similarities and differences in SNPs from assembled genome sequences and publicly available raw reads. We collected 15,818 high-quality SARS-CoV-2 genome sequences from the publicly accessible GISAID, CNGBdb, GenBank, GWH^9^ and NMDC databases on May 27, 2020. All of these sequences were aligned against the reference genome (Wuhan-Hu-1, GenBank NC_045512.2)^10^ using MUSCLE^11^, revealing 7,700 inter-host SNP sites (4,672 nonsynonymous, 2,570 synonymous, 98 stop gain, 7 stop loss, and 433 noncoding) (Extended Data Table 1). Publicly available raw reads were extracted from 1,364 clinical SARS-CoV-2 samples from Australia, 109 samples from India, 400 samples from the United Kingdom, 112 samples from China^12^, and 365 samples from the United States that were collected between January and May 2020 (Extended Data Table 2). After quality control steps, 5,705 iSNVs (3,796 nonsynonymous, 1,572 synonymous, 261 stop gain, 4 stop loss and 134 noncoding) (Extended Data Table 3) were identified from these samples, with only 48.3% (2,756/5,705) of them shared with the SNPs from the inter-host SNP data (Extended Data Fig.1). Combining these two SNPs dataset revealed that 35.6% (10,649/29,903) of the bases in the SARS-CoV-2 genome sequence had been mutated at least once Extended Data Fig.1). Because only 2,350 samples with raw reads have been analyzed, the number of iSNV is much underestimated.

We then investigated the distribution of inter-host SNPs and iSNVs in each open reading frame (ORF) and found similar trends in the SNP numbers and SNP numbers normalized by the length between inter-host SNPs and iSNVs (Fig. 1, a-d). We then focused on the S-protein receptor binding domain (RBD) as it is being used for monoclonal antibody isolation and vaccine design^13,14^. Several mutations were found in this region, including 68 inter-host nonsynonymous and 65 intra-host nonsynonymous mutations (Fig. 1, a and c). An excess of C-to-T, A-to-G and T-to-C mutations were observed in both the inter-host SNPs and iSNVs, which may be the result of the RNA editing signature of APOBEC deaminases and ADAR enzymes (Fig. 1e). Furthermore, we analyzed the distributions of the numbers of nonsynonymous and synonymous mutations (Fig. 1f). Some regions of the SARS-CoV-2 genome have greater numbers of nonsynonymous mutations than the average level (Fig. 1f). The high numbers of nonsynonymous mutations found at the ends of the SARS-CoV-2 genome might stem from errors of genome assembly. The higher numbers of nonsynonymous mutations for some regions might be due to higher substitution rates and increased recombination. To address this, we estimated the substitution rates using BEAST^15^ a set of assembled genomes obtained from patients from Shanghai. A substitution rate of 1.20×10^−3^ mutations per site per year (95% highest posterior density interval, 6.14×10^−4^ to 1.87×10^−3^ mutations per site per year) for these SARS-CoV-2 genomes was estimated (Extended Data Table 4), which is similar to the previous estimate of 9.7× 10^−4^ mutations per site per year^16^. However, higher substitution rates were found for some protein-coding regions (such as genome positions 21,563-22,700 and 24,300-25,384 in the S protein), of approximately 2.95× 10^−3^ to 3.94× 10^−3^ mutations per site per year (Fig. 1f and Extended Data Table 4). When the sequences were tested for evidence of recombination using the Recombination Detection Program (RDP)^17^, none was detected in the genomes from the Shanghai patients. These results suggest that some regions of the SARS-CoV-2 genome are experiencing faster rates of evolution.

**Fig. 1.**
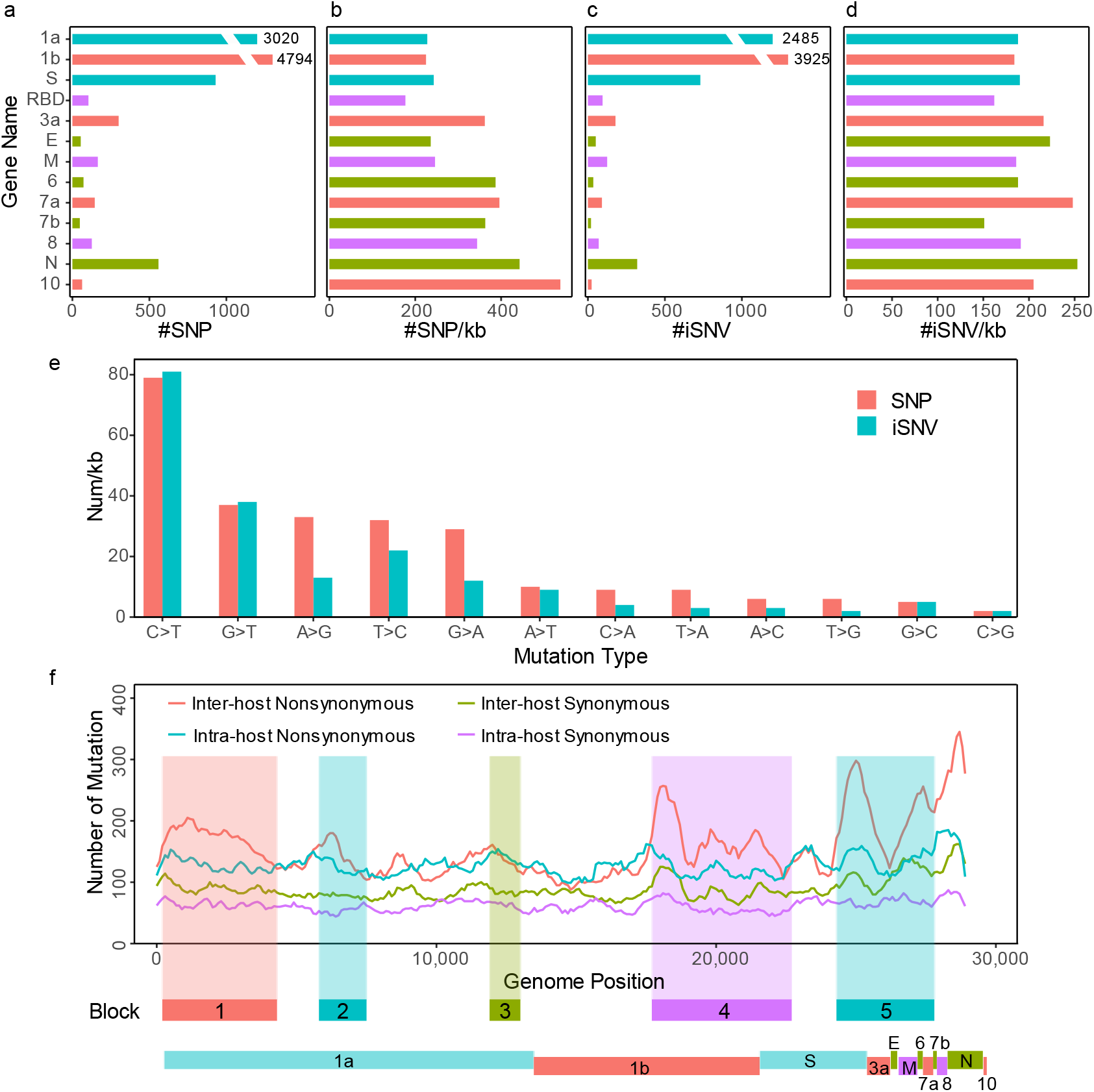
Distributions of inter-host SNPs and iSNVs in the SARS-CoV-2 viral genome. **a**. Total number of inter-host SNPs in each ORF and RBD region. **b**. The number of inter-host SNPs in each kb of each ORF and RBD region. **c**. Total number of iSNVs in each ORF and RBD region. **d**. The number of iSNVs for each ORF and RBD region normalized by kb. **e**. The numbers of each SNP type normalized by kb for inter-host SNPs and iSNVs. **f.** Distribution of nonsynonymous and synonymous mutations in inter-host and intra-host SNPs. Number of mutations were calculated using a sliding window with a size of 1000bp and a step size of 100bp. The blocks are regions having more protein gene mutations than the average level.

To determine whether diversifying selection was occurring in SARS-CoV-2, we first calculated π, the nucleotide diversity of the viral populations using pairwise comparisons for each gene from the inter-host sequences. We can evaluate the level of natural selection acting upon the sequences by comparing the frequencies of the synonymous (πS) and nonsynonymous (πN) polymorphisms^18^. πS>πN is indicative of purifying selection removing deleterious mutations, πS<πN indicates diversifying selection, and πS=πN suggests neutrality. For most genes, the inter-host sequence data shows that πN is less than πS, implying that purifying selection plays a central role in the evolution of these SARS-CoV-2 genes (Extended Data Table 5). For ORF3a and ORF8, πN was greater than πS, although the standard error of the πN values is high (Extended Data Table 5), which prevents us from concluding that diversifying selection is occurring in those two genes. Previous studies have indicated that evolution of the mucin-like domain of the Ebola virus glycoprotein might have been driven by diversifying selection imposed by antibodies^7^. To further investigate regions of the SARS-CoV-2 genome whose evolution might have been driven by diversifying selection, we calculated πN and πS using a sliding window of 30 codons and a step size of 3 codons^18^. Using this sliding window analysis, we found signals for diversifying selection in small regions of the ORF1ab, S, ORF3a, and ORF8 genes (Fig. 2a). πN is greater than πS in one region (Genome position: 23,318-23,488, S protein: 584-641) in the SD2 domain of the S protein (27 nonsynonymous and 16 synonymous within this region) (Fig. 2a). This region, including the fitness advantage pandemic mutation D614G^5^, is in the surface-exposed viral protein of the SARS-CoV virus and might be a primary target for antibodies. Thus, antibodies might be the driving force for the diversifying selection seen in this region. Further studies are needed to test the enrichment of mutations within the B cell epitopes within this region.

**Fig. 2.**
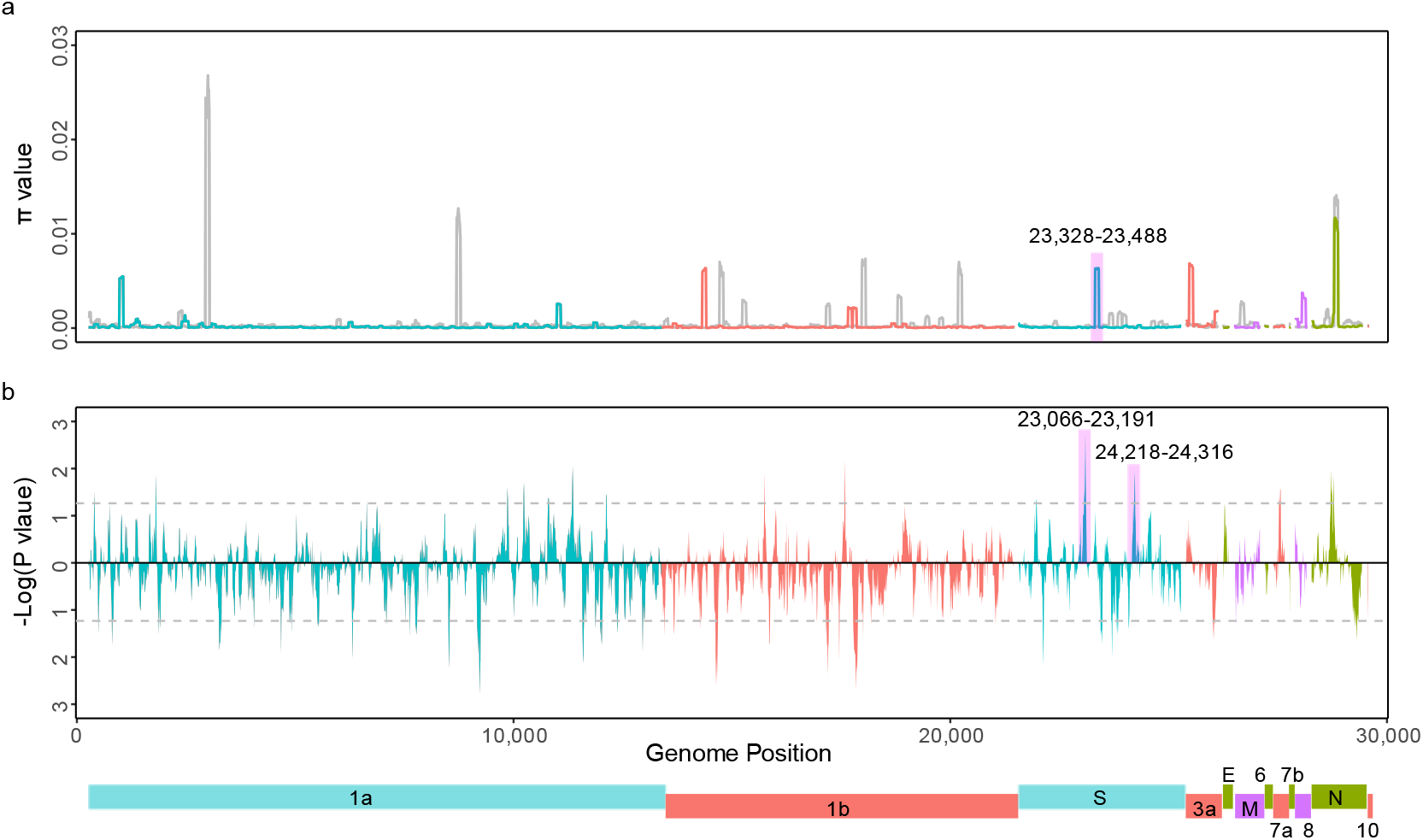
Analysis of natural selection of the SARS-CoV-2 virus ORFs. **a**. πN (colored lines) and πS (grey lines) were estimated for the SARS-CoV-2 virus using a 30 amino acids window and 3 amino acids step. Different ORFs are distinguished by different colors for πN. **b**. The Kolmogorov-Smirnov test was performed between the nonsynonymous and synonymous iSNVs mutated allele frequencies (MuAFs). The upper segment indicates that the MuAF values of the nonsynonymous mutation is higher than the synonymous mutation, while in the lower segment the nonsynonymous mutations are lower than the synonymous mutations. The dashed lines represent P<0.05.

The frequency of iSNVs can also be used to reveal selection signals^8,19^. We analyzed the mutated allele frequency (MuAF) using the Wuhan-Hu-1 genome as a reference. We performed the MuAF analysis using a sliding window of 30 codons and a step size of 3 codons. A small region (Genome position: 23,066-23,191, S protein: 502-543) of the RBD and SD1 domains of the S protein has a nonsynonymous frequency that is significantly higher than the synonymous frequency (there are 13 nonsynonymous and 7 synonymous mutations in this region) (Kolmogorov-Smirnov test, P<0.05) (Fig. 2b). Another region (Genome position: 24,218-24,316, S protein: 886-918) in the S protein CR and HR1 domains also has a significantly higher nonsynonymous frequency than the synonymous frequency (there are 9 nonsynonymous and 4 synonymous mutations in this region) (Fig. 2b) and is overlapping with substitution mutation rate region (block 5 in Extended Data Table 4; genome position: 24,300-27,800). Diversifying selection on these two regions might also be driven by an intra-host escape from antibody recognition.

To study the mechanisms for the generation and transmission of fitness advantage mutations, we first investigated the number of iSNVs over time and noticed that the numbers of iSNVs was generally steady with few fluctuations (Fig. 3a). Thus, we concluded that there was no increase in diversity within intra-host genomes over time. iSNVs provide valuable information about human-to-human transmission and are useful for characterizing human-to-human transmission chains. In the 2,187 worldwide cases, which spanned 5 months of the COVID-19 epidemic, 1,988 samples shared at least one iSNVs (Fig. 3b). The shared iSNVs can be explained by superinfection, recurring mutation or human-to-human transmission^7^. We can rule out superinfection, as 2,949 shared iSNVs were detected only in raw reads and were not identified in the assembled sequences. Additionally, the fitness advantage mutation (G to A change) at position 23,403 (spike protein, D614G, Wuhan-Hu-1, GenBank NC_045512.2)^5^ is at an intermediate frequency (~60% in four samples) in the Australia population, and belong to G clade (Extended Data Fig.2). If superinfection were common in COVID-19 patients, we would expect to see near equal distributions of iSNVs containing individuals in both the A and G clades. From the above results, we conclude that superinfection is not an important reason for the existence of iSNVs. Another possible reason for shared iSNVs is recurring mutation and human-to-human transmission. We can see the shared iSNVs at low frequencies in different countries or regions and at different times. Thus, we cannot exclude the influence of recurring mutation. Importantly, all of the shared iSNVs are unlikely to be the result of only recurring mutations as 23,403 variants were found at >90% frequency in 34 different samples from the United Kingdom, the United States, and Australia. The high frequency of the shared iSNVs might arise from human-to-human transmission and/or adaption. In summary, we can conclude that recurrent mutations and human-to-human transmission could lead to the observed iSNV pattern seen in COVID-19 patients.

**Fig. 3.**
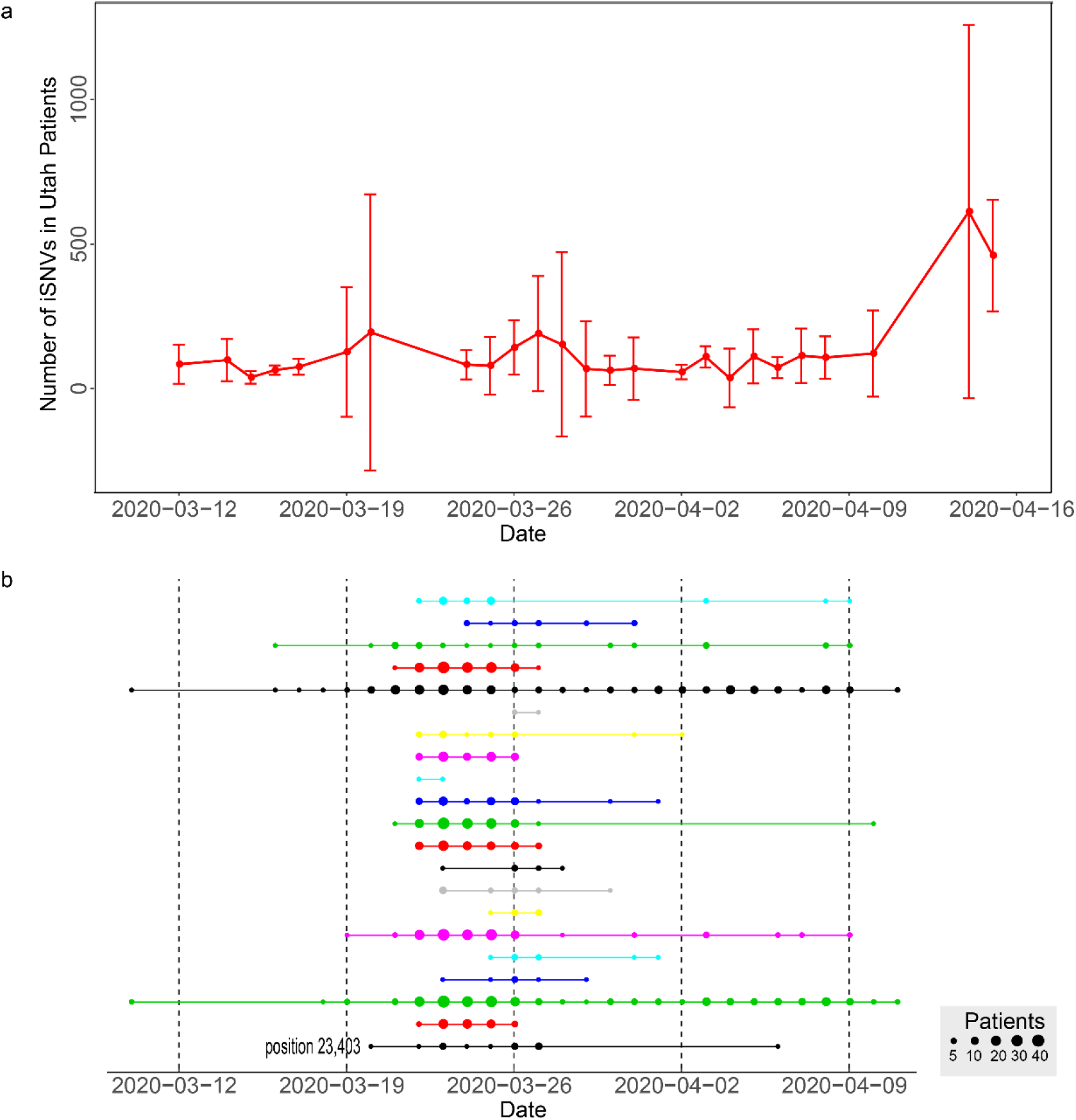
Distribution and sharing of iSNVs in SARS-CoV-2 virus. **a**. Time distribution of iSNVs in the Utah COVID-19 patients. Numbers indicate the number of iSNVs in all the patients of the same day. Error bars represent the 95% confidence interval. **b**. Evidence of human-to-human sharing in the Victoria (Australia) iSNVs. SNPs shared in two or more patients are displayed as a row of connected points. Each row represents a randomly selected SNP from the shared iSNVs. Position 23,403 is an example of a shared fitness advantageous mutation.

Population genetics theory points out that an allele with a fitness advantage will increase in frequency faster than alleles without a fitness advantage. Thus, if a mutation has a selective advantage, then the frequency of this mutation will gradually increase over time in a transmission chain (Fig. 4a). To detect possible fitness advantage mutations in SARS-CoV-2, we analyzed 343 iSNVs in genomes from Shanghai, China, requiring that the iSNVs existed for at least three days and from two samples for each day. Furthermore, we constructed a median-joining haplotype network (Fig. 4b) using PopART (http://popart.otago.ac.nz) and a phylogenetic tree (Extended Data Fig.3) using IQ-TREE for the Shanghai samples^20^. We then calculated the allele frequency and plotted the population-level frequency dynamic with isolate sampling times for the above iSNVs for each lineage. We found two shared iSNVs (mutation type) in the red cluster and two shared iSNVs in the black cluster that had gradual frequency increases over time (Fig. 4c and Extended Data Fig.4a). The T to C mutation (position 22,760, S protein) causing the F400L amino acid substitution might be a fitness advantage mutation for SARS-CoV-2 in these patients, while another synonymous mutation (T to C, position 22,762) in the same codon is likely hitchhiking. Another pair of mutations (C to T, position 25,883, ORF3a protein, T164I and A to T, position 19413, ORF1a protein, Q6383H) might also provide a fitness advantage within these SARS-CoV-2 patients. Genetic drift could also lead to increases in iSNV frequency. Whereas genetic drift may affect the whole genome, some iSNVs may not follow similar patterns and thus, likely were driven by purifying or neutral selection (Fig. 4d and Extended Data Fig.4b). However, we cannot completely rule out the influence of genetic drift. These mutations, to date, are not yet highly prevalent in COVID-19 patients. We urged researchers to focus on these types of mutations and upload raw data of SARS-CoV-2 genomes to public databases for the monitoring of iSNV frequencies.

**Fig. 4.**
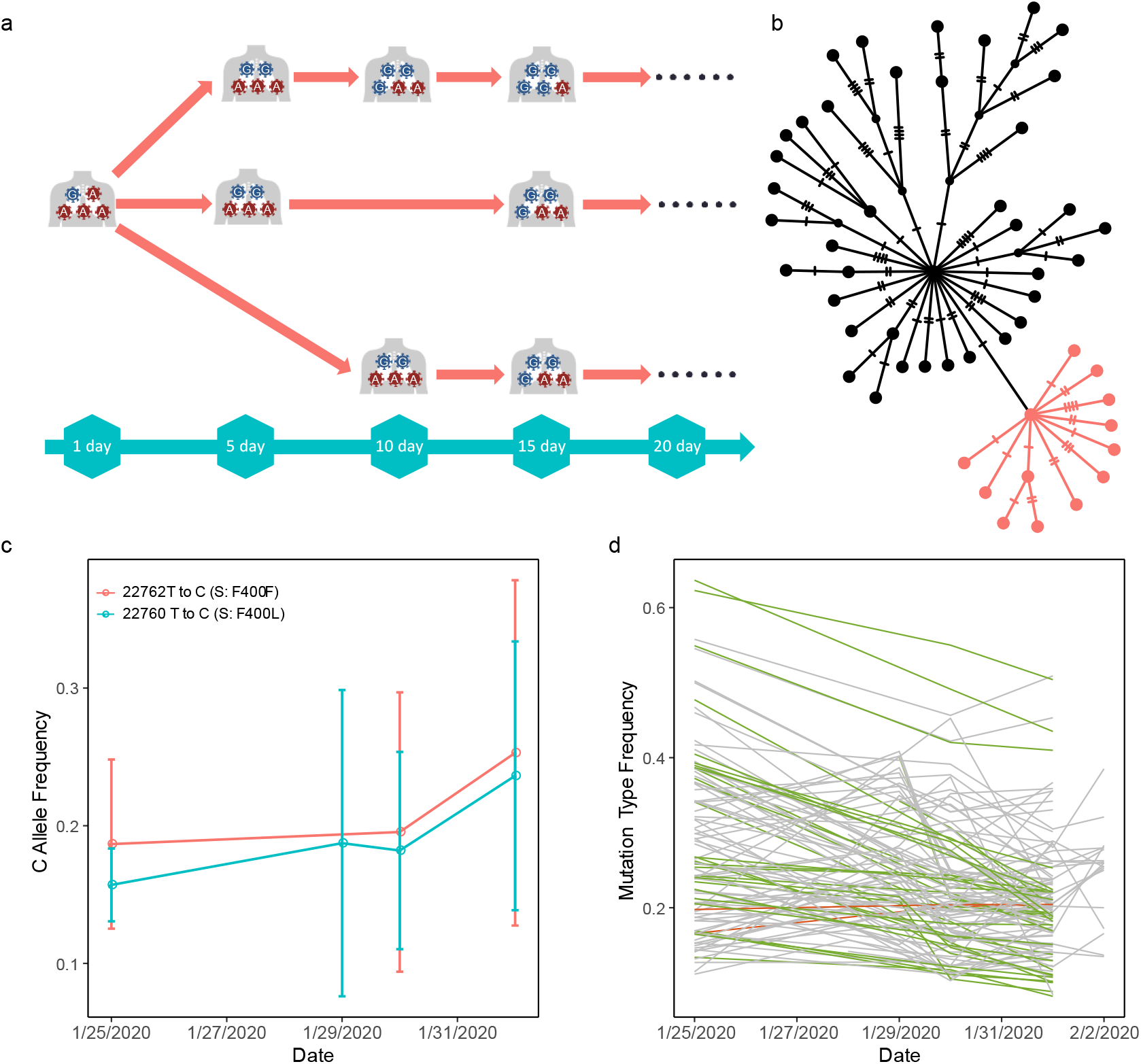
Early prediction of fitness advantageous mutations for SARS-CoV-2 virus using iSNVs. **a**. Model of the human-to-human transmission of an iSNVs. G mutation is the supposed fitness advantageous mutation. **b**. Median-joining haplotype network constructed from the Shanghai SARS-CoV-2 genomes. Red colored dots represent a viral haplotype, which included samples used for the fitness advantageous mutation analysis. The clustered samples were assumed to be in the same transmission chain. **c**. Example of the early prediction of a fitness advantageous mutation. Mutation type frequency was calculated as the ratio of the number of C reads to the sum of the numbers of the T and C reads. Date was the sampling day. **d**. iSNVs dynamics for SARS-CoV-2 from the Shanghai patients (from a cluster marked in red in B). These iSNVs represent mutations that might be driven by neutral and purifying selection. Mutation types were inferring using the Wuhan-Hu-1 genome as the reference genome. The red color represents the mutation types frequency gradually increase over time. The green color indicates decrease of mutation types frequency over time. Grey color represents the fluctuation of mutation types frequency.

In summary, we found that to May 27, at least 35.6% of the bases of the SARS-CoV-2 genome harbor one or more mutation and that evolution of mutations in the spike protein gene might have been driven by diversifying selection to increase diversity to escape immune system recognition. Furthermore, we found that recurring mutations and human-to-human transmission are the reasons for the prevalence of the fitness advantage mutations in SARS-CoV-2. Through an analysis of shared iSNVs in SARS-CoV-2 genomes from patients from Shanghai, we found three alleles that might be fitness advantage mutations for SARS-CoV-2. This study provides theoretical support for vaccine development and optimizing treatment. We also strongly recommend that researchers submit raw sequence data and epidemiological information of SARS-CoV-2 isolates to public databases, which will improve the detection of fitness advantage mutations of SARS-CoV-2.

## Methods

### SARS-CoV-2 data

15,818 high-quality assembled SARS-CoV-2 sequences were acquired from the GISAID (https://www.gisaid.org/), CNGBdb (https://db.cngb.org/), GenBank (https://www.ncbi.nlm.nih.gov/genbank/), GWH (https://bigd.big.ac.cn/gwh/), and NMDC (http://nmdc.cn/nCoV) databases (on May 27, 2020). These sequences were sampled from December 2019 to May 2020 and distributed across 77 countries.

We downloaded raw sequence datasets for SARS-CoV-2 that included 1,160 samples from Australia (SRA accession number PRJNA613958), 400 from United Kingdom (SRA accession number PRJEB37886), 281 from United States (SRA accession number PRJNA614995), and 112 from China (SRA accession number PRJNA627662). All data used in this study are acknowledged and listed in Extended Data Table 2.

### Assembled sequence analysis

All 15,817 assembled sequences were aligned against the SARS-CoV-2 reference genome (Wuhan-Hu-1, GenBank NC_045512.2) using MUSCLE version 3.8.31^11^. Subsequently, SNPs were called from the aligned sequences. SNPs were annotated to protein-coding genes and intergenic regions using ANNOVAR^21^.

Assembled sequences from Shanghai and Australia were downloaded from the GISAID database (https://www.gisaid.org/), which have corresponding raw sequence data in the SRA database (accession number PRJNA627662 and PRJNA613958), were aligned using MUSCLE version 3.8.31^11^. Maximum likelihood phylogenetic trees were constructed using IQ-TREE version 2.0.5^20^. Median-joining haplotype networks were built using PopART version 1.7 (http://popart.otago.ac.nz/). Phylogenetic trees and haplotype networks were used for further analyses.

### iSNVs analysis

To obtain high-confidence iSNVs, we first trimmed low-quality bases from the raw reads using Trimmomatic 0.39 with default parameters^22^. The passed reads were mapped to the SARS-CoV-2 reference genome using BWA mem version 0.7.17 with default parameters^23^. GATK MarkDuplicates version 4.1.3.0 was used to mark duplicate reads^24,25^. The iSNVs were detected using bcftools mpileup version 1.10.2 with : mapping quality >20, base quality >20, mapping quality >50^26^. iSNVs were then filtered using bcftools filter version 1.10.2 with ‘QUAL<10 || DP<15’ parameters for data from India, United Kingdom, and China, and ‘QUAL<10 || DP<30’ for data from the United States and Australia^26^. We used different parameters due to their differences in sequencing depth. To remove errors from the sequencing and mapping data, only iSNVs that were covered by at least five reads were used for further analyses.

### Substitution rates

Substitution rates for the Shanghai samples were assessed using the Bayesian Markov chain Monte Carlo (MCMC) implemented in BEAST v1.8.4^15^. We performed the analysis for 100 million generations, sampling every 10,000 steps as done in a previous study^27^.

### Natural selection analysis

We calculated the nonsynonymous (πN) and synonymous (πS) nucleotide diversity using SNPGeine with a script using the aligned sequences^18^. To intuitively display the potential natural selection regions in the SARS-CoV-2 genome, we further carried out sliding window analyses (window size, 30 codons; step size, 3 codons) of the πN and πS.

Mutated allele frequency (MuAF) of the iSNVs were calculated using a sliding window (window size, 30 codons; step size, 3 codons) with a sequencing error rate of 0.001 and minor allele frequency of 0.001, and other parameters as used in a previous study^19^.

### iSNVs dynamics analysis

To prevent a mapping bias caused by polymorphisms, the iSNVs in the SARS-CoV-2 reference genome were replaced with ‘N’ for each sample. We used BWA mem 0.7.17 with default parameters to re-map the reads^23^.

Following the haplotype network and phylogenetic tree in the Shanghai samples, we calculated the allele frequency and plotted the frequency with collection time for each lineage.

## Acknowledgments

This work was supported by the Chinese Academy of Sciences, National Nature Science Foundation of China (91331101), and the National Key Research and Development Project (No. 2020YFC0847000).

## Author contributions

Y.P.Z. and Z.Y.Z. designed the project. H.L. and Z.Y.Z. performed the data analysis. Z.Y.Z. wrote the primary manuscript. Z.Y.Z., H.L., A.L., Y.D.Z. and Y.Q.W drew the figures. Y.B. provided the SARS-CoV-2 sequence alignment. M.S.P., D.M.I., H.L., J.L., X.L., D.L. and Y.P.Z. revised the manuscript.

## Additional information

## Extended data table legends

**Extended Data Table 1**. SNPs identified in the assembled SARS-CoV-2 sequences.

**Extended Data Table 2**. Acknowledgement of the sharing of SARS-CoV-2 raw genome sequences in NCBI SRA database used in this study.

**Extended Data Table 3**. Annotations of the SNPs identified from the raw reads generated from next generation sequencing of SARS-CoV-2 samples.

**Extended Data Table 4**. Estimates of the substitution rate for the SARS-CoV-2 genome and the regions with high numbers of mutations in protein genes. HPD: 95% highest posterior density interval.

**Extended Data Table 5**. Mean nonsynonymous and synonymous nucleotide diversity of each ORF

**Extended Data Fig.1.**
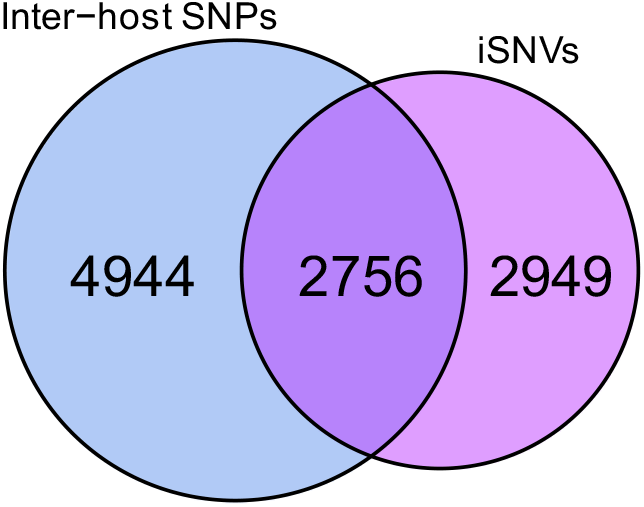
Overlapping of SNPs identified in the inter-host and the intra-host.

**Extended Data Fig.2.**
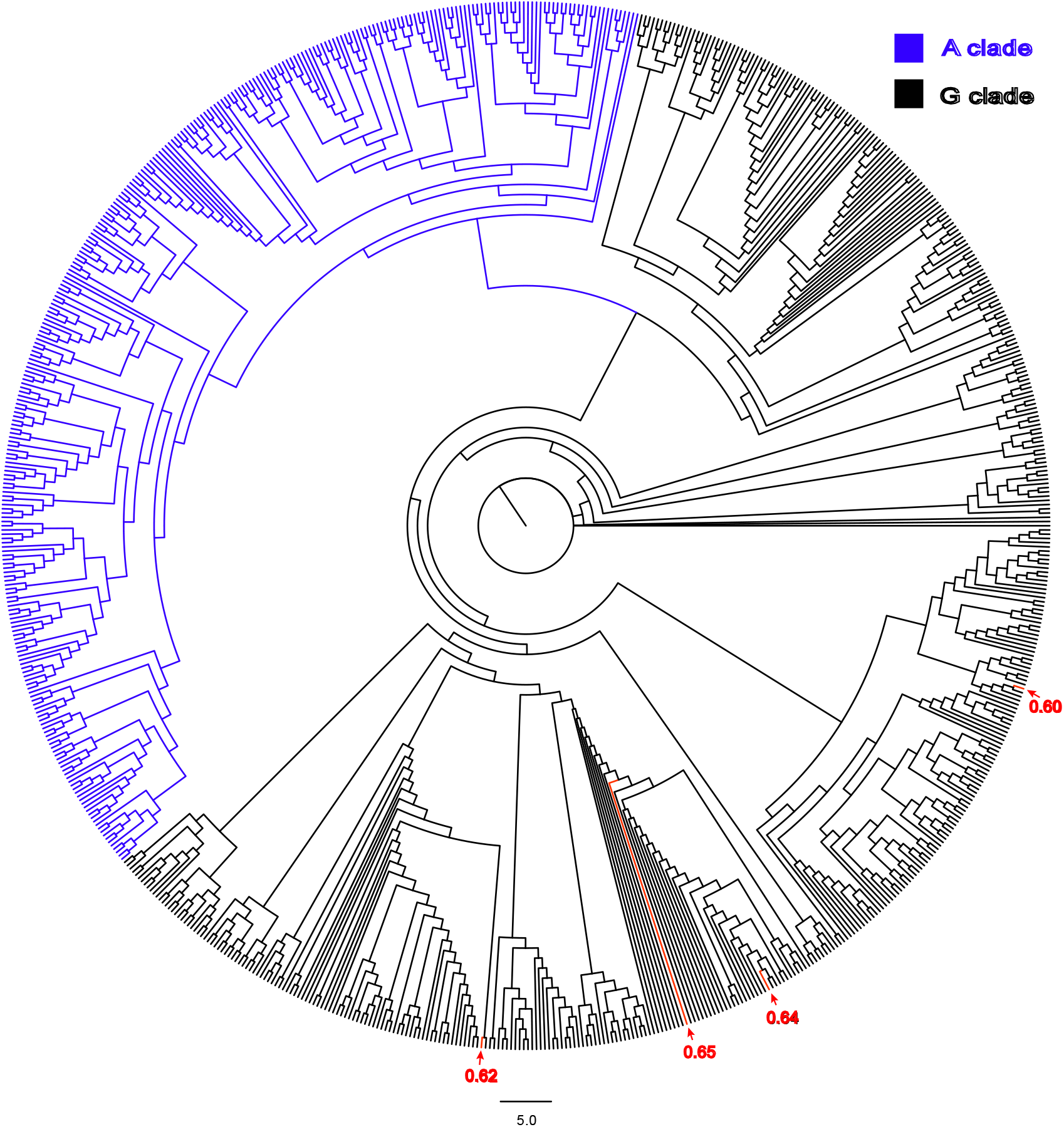
Phylogenetic placement of intermediate frequency (~60%) SNP at position 23,403. The frequency of the G mutation type was calculated as the ratio of number of G reads to the sum of numbers of A and G reads. Samples related to the intermediate frequency are marked in red.

**Extended Data Fig.3.**
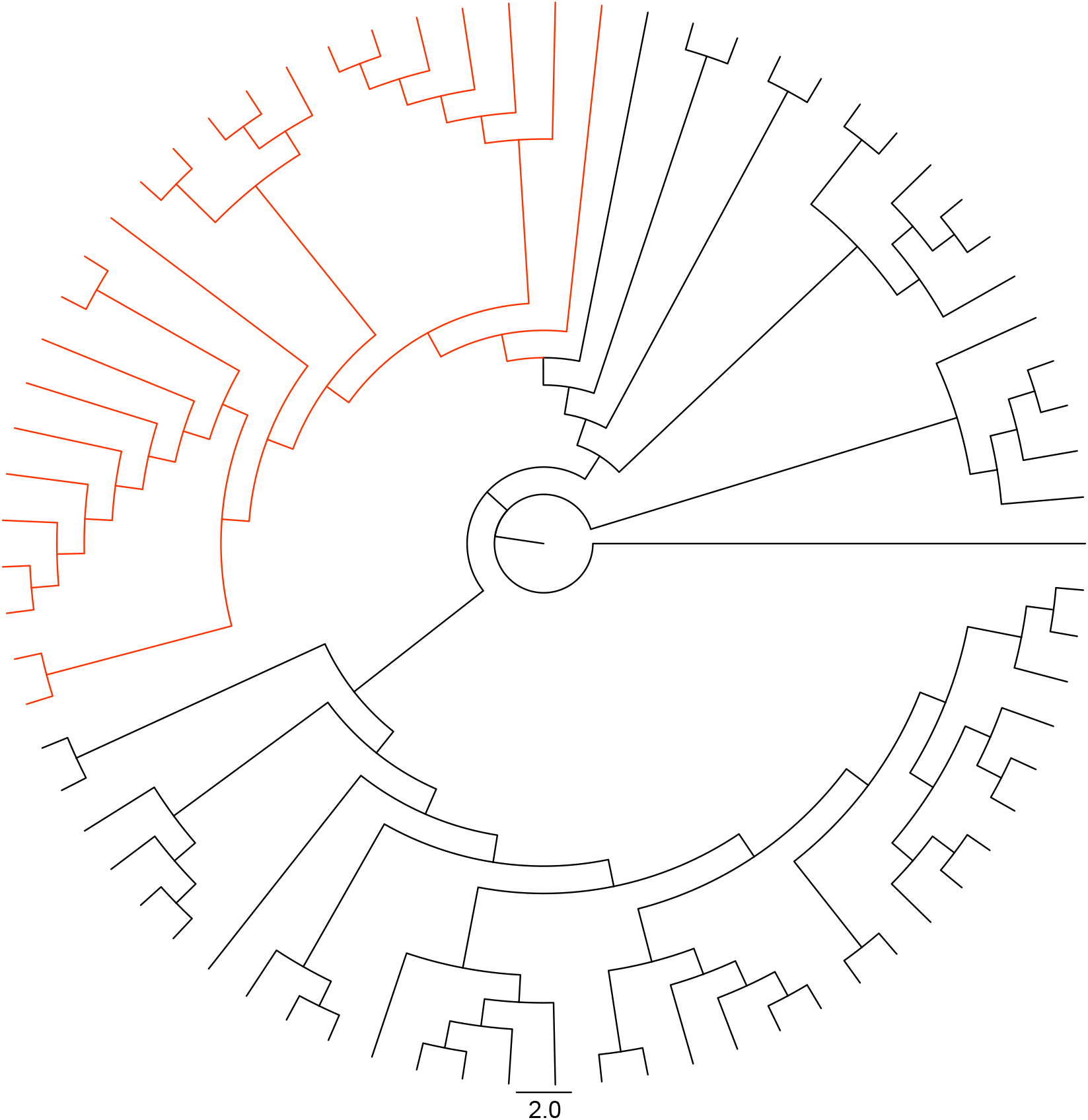
Phylogenetic placement of the Figure 3B viral haplotype (marked in red). Clustered samples were assumed to be in the same transmission chain.

**Extended Data Fig.4.**
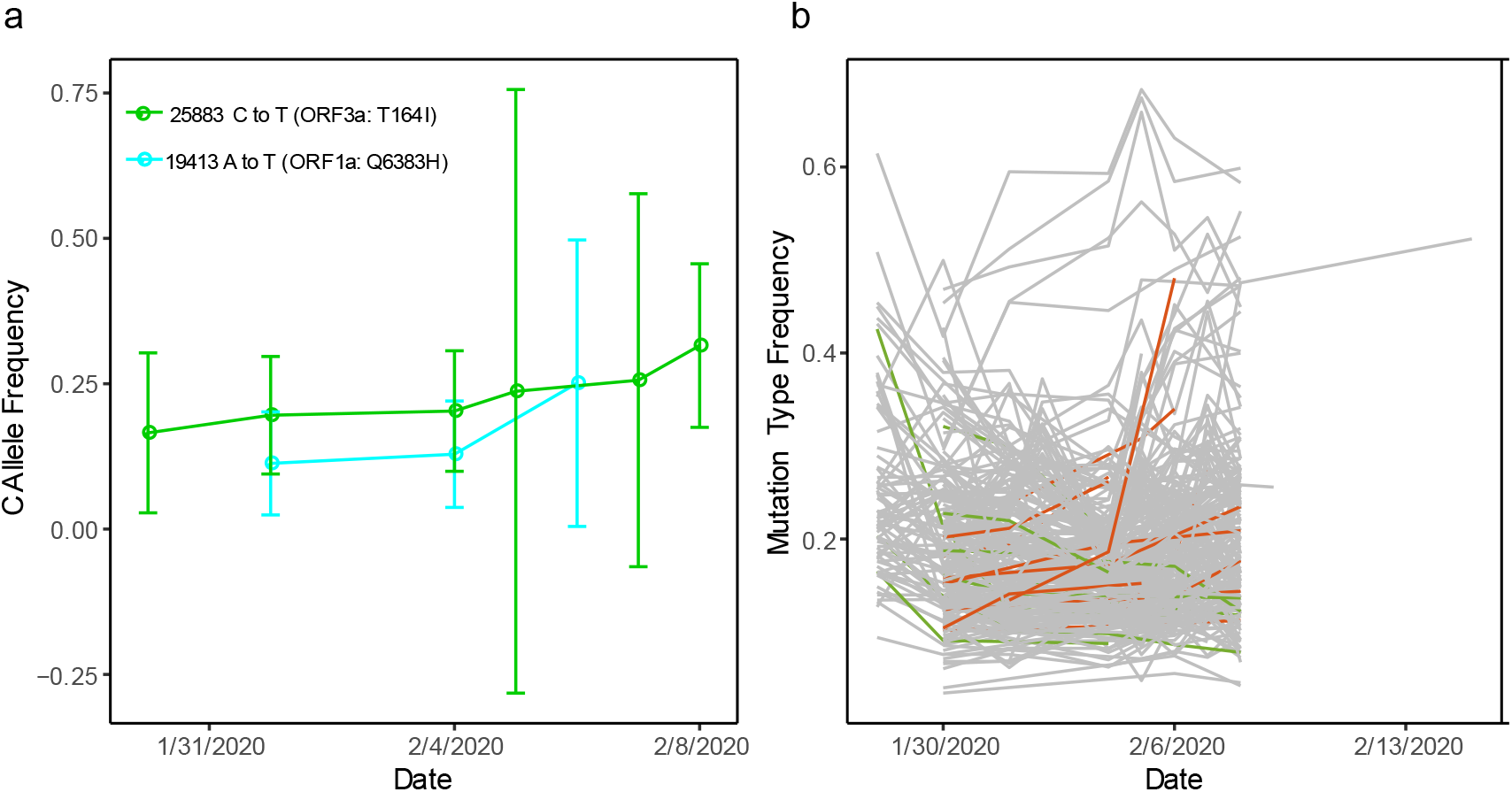
Example of other candidate fitness advantageous mutations. **a.** Two candidate fitness advantageous mutations identified in the Shanghai patients. Mutation types were inferring using the Wuhan-Hu-1 genome as the reference genome. Mutation type frequency was calculated as the ratio of number of mutation type reads to the sum of numbers of wild- and mutation-type reads. Date was the sampling day. **b.** iSNVs dynamics for the Shanghai patients (from the cluster labeled in black in Fig. 4B). These iSNVs may be driven by neutral and purifying selection.

## Main References

1 Organization, W. H. WHO Coronavirus disease(COVID-19) Situation Report, https://www.who.int/emergencies/diseases/novel-coronavirus-2019/situation-reports/ (2020).

2 Zhou, P. et al. A pneumonia outbreak associated with a new coronavirus of probable bat origin. Nature 579, 270–273, https://doi.org/10.1038/s41586-020-2012-7 (2020).

3 Chan, J. F. et al. A familial cluster of pneumonia associated with the 2019 novel coronavirus indicating person-to-person transmission: a study of a family cluster. Lancet (London, England) 395, 514–523, https://doi.org/10.1016/s0140-6736(20)30154-9 (2020).

4 Wrapp, D. et al. Cryo-EM structure of the 2019-nCoV spike in the prefusion conformation. Science (New York, N.Y.) 367, 1260–1263, https://doi.org/10.1126/science.abb2507 (2020).

5 Korber, B. et al. Tracking changes in SARS-CoV-2 Spike: evidence that D614G increases infectivity of the COVID-19 virus. Cell (2020).

6 Tang, X. et al. On the origin and continuing evolution of SARS-CoV-2. National science review (2020).

7 Park, D. J. et al. Ebola Virus Epidemiology, Transmission, and Evolution during Seven Months in Sierra Leone. Cell 161, 1516–1526, https://doi.org/10.1016/j.cell.2015.06.007 (2015).

8 Ni, M. et al. Intra-host dynamics of Ebola virus during 2014. Nature microbiology 1, 16151, https://doi.org/10.1038/nmicrobiol.2016.151 (2016).

9 Zhao, W. M. et al. The 2019 novel coronavirus resource. Yi chuan = Hereditas 42, 212–221, https://doi.org/10.16288/j.yczz.20-030 (2020).

10 Wu, F. et al. A new coronavirus associated with human respiratory disease in China. Nature 579, 265–269, https://doi.org/10.1038/s41586-020-2008-3 (2020).

11 Edgar, R. C. MUSCLE: multiple sequence alignment with high accuracy and high throughput. Nucleic acids research 32, 1792–1797, https://doi.org/10.1093/nar/gkh340 (2004).

12 Zhang, X. et al. Viral and host factors related to the clinical outcome of COVID-19. Nature, https://doi.org/10.1038/s41586-020-2355-0 (2020).

13 Wu, Y. et al. A noncompeting pair of human neutralizing antibodies block COVID-19 virus binding to its receptor ACE2. Science (New York, N.Y.) 368, 1274–1278, https://doi.org/10.1126/science.abc2241 (2020).

14 Dai, L. et al. A Universal Design of Betacoronavirus Vaccines against COVID-19, MERS, and SARS. Cell, https://doi.org/10.1016/j.cell.2020.06.035 (2020).

15 Suchard, M. A. et al. Bayesian phylogenetic and phylodynamic data integration using BEAST 1.10. Virus evolution 4, vey016, https://doi.org/10.1093/ve/vey016 (2018).

16 van Dorp, L. et al. No evidence for increased transmissibility from recurrent mutations in SARS-CoV-2. bioRxiv, https://doi.org/doi.org/10.1101/2020.05.21.108506 (2020).

17 Martin, D. P., Murrell, B., Golden, M., Khoosal, A. & Muhire, B. RDP4: Detection and analysis of recombination patterns in virus genomes. Virus evolution 1, vev003, https://doi.org/10.1093/ve/vev003 (2015).

18 Nelson, C. W., Moncla, L. H. & Hughes, A. L. SNPGenie: estimating evolutionary parameters to detect natural selection using pooled next-generation sequencing data. Bioinformatics (Oxford, England) 31, 3709–3711, https://doi.org/10.1093/bioinformatics/btv449 (2015).

19 Li, Y. et al. Resequencing of 200 human exomes identifies an excess of low-frequency non-synonymous coding variants. Nature genetics 42, 969–972, https://doi.org/10.1038/ng.680 (2010).

20 Minh, B. Q. et al. IQ-TREE 2: New Models and Efficient Methods for Phylogenetic Inference in the Genomic Era. Molecular biology and evolution 37, 1530–1534, https://doi.org/10.1093/molbev/msaa015 (2020).

## Methods References

5 Korber, B. et al. Tracking changes in SARS-CoV-2 Spike: evidence that D614G increases infectivity of the COVID-19 virus. Cell, https://doi.org/doi:10.1016/j.cell.2020.06.043 (2020).

6 Tang, X. et al. On the origin and continuing evolution of SARS-CoV-2. National science review, https://doi.org/doi.org/10.1093/nsr/nwaa036 (2020).

21 Wang, K., Li, M. & Hakonarson, H. ANNOVAR: functional annotation of genetic variants from high-throughput sequencing data. Nucleic acids research 38, e164, https://doi.org/10.1093/nar/gkq603 (2010).

22 Bolger, A. M., Lohse, M. & Usadel, B. Trimmomatic: a flexible trimmer for Illumina sequence data. Bioinformatics (Oxford, England) 30, 2114–2120, https://doi.org/10.1093/bioinformatics/btu170 (2014).

23 Li, H. & Durbin, R. Fast and accurate short read alignment with Burrows-Wheeler transform. Bioinformatics (Oxford, England) 25, 1754–1760, https://doi.org/10.1093/bioinformatics/btp324 (2009).

24 McKenna, A. et al. The Genome Analysis Toolkit: a MapReduce framework for analyzing next-generation DNA sequencing data. Genome research 20, 1297–1303, https://doi.org/10.1101/gr.107524.110 (2010).

25 DePristo, M. A. et al. A framework for variation discovery and genotyping using next-generation DNA sequencing data. Nature genetics 43, 491–498, https://doi.org/10.1038/ng.806 (2011).

26 Li, H. A statistical framework for SNP calling, mutation discovery, association mapping and population genetical parameter estimation from sequencing data. Bioinformatics (Oxford, England) 27, 2987–2993, https://doi.org/10.1093/bioinformatics/btr509 (2011).

27 Tong, Y.-G. et al. Genetic diversity and evolutionary dynamics of Ebola virus in Sierra Leone. Nature 524, 93–96, https://doi.org/ doi: 10.1038/nature14490 (2015).

