## Extended Data Table for "Worldwide tracing of mutations and the evolutionary dynamics of SARS-CoV-2": Extended Data Table 1.pdf

**Extended Data Table 1. SNPs identified in the assembled SARS-CoV-2 sequences.**

| Genome Position | Ref | Alt1 | Alt1 frequency in worldwide | Alt2 | Alt2 frequency in worldwide | Alt3 | Alt3 frequency in worldwide | Annotation | |
|---|---|---|---|---|---|---|---|---|---|
| 29838 | C | A | 8.23E-01 | G | 1.81E-03 | T | 1.57E-03 | three_prime_UTR | |
| 241 | C | T | 7.09E-01 | | | | | five_prime_UTR | |
| 23403 | A | G | 7.07E-01 | | | | | S | nonsynonymous_D614G |
| 3037 | C | T | 7.04E-01 | | | | | ORF1ab | synonymous_F924F |
| 14408 | C | T | 7.04E-01 | | | | | ORF1ab | nonsynonymous_P4715L |
| 29880 | A | G | 5.91E-01 | T | 5.89E-03 | C | 3.75E-03 | three_prime_UTR | |
| 29841 | G | A | 5.80E-01 | T | 1.62E-03 | C | 6.47E-04 | three_prime_UTR | |
| 1 | A | T | 5.36E-01 | C | 5.47E-03 | G | 3.65E-03 | five_prime_UTR | |
| 29903 | A | G | 5.35E-01 | C | 4.79E-03 | T | 3.19E-03 | three_prime_UTR | |
| 29882 | A | G | 5.03E-01 | C | 2.85E-01 | T | 2.71E-03 | three_prime_UTR | |
| 29898 | A | T | 5.39E-03 | T | 5.39E-03 | | | three_prime_UTR | |
| 29879 | A | T | 4.88E-01 | G | 1.01E-01 | C | 1.59E-03 | three_prime_UTR | |
| 29902 | A | G | 4.83E-01 | T | 4.29E-03 | C | 1.43E-03 | three_prime_UTR | |
| 29883 | A | G | 4.56E-01 | C | 2.16E-03 | T | 1.62E-03 | three_prime_UTR | |
| 29893 | A | G | 4.34E-01 | T | 2.99E-03 | C | 1.50E-03 | three_prime_UTR | |
| 3 | T | A | 4.33E-01 | G | 4.58E-03 | C | 3.20E-03 | five_prime_UTR | |
| 29864 | G | A | 4.25E-01 | T | 2.15E-03 | T | 2.15E-03 | three_prime_UTR | |
| 29865 | A | G | 4.17E-01 | C | 3.94E-03 | T | 8.75E-04 | three_prime_UTR | |
| 4 | A | T | 4.00E-01 | G | 6.70E-03 | C | 3.94E-03 | five_prime_UTR | |
| 29886 | A | G | 3.81E-01 | T | 8.22E-03 | C | 1.10E-03 | three_prime_UTR | |
| 6 | A | C | 3.70E-01 | T | 5.35E-03 | G | 1.78E-03 | five_prime_UTR | |
| 7 | G | C | 3.58E-01 | T | 3.74E-03 | A | 1.02E-03 | five_prime_UTR | |
| 29900 | A | T | 3.51E-01 | G | 6.76E-03 | C | 3.86E-03 | three_prime_UTR | |
| 8 | G | T | 3.51E-01 | A | 4.22E-03 | C | 3.25E-03 | five_prime_UTR | |
| 29899 | A | T | 3.49E-01 | G | 2.89E-03 | C | 1.92E-03 | three_prime_UTR | |
| 10 | T | C | 3.46E-01 | A | 4.64E-03 | G | 3.41E-03 | five_prime_UTR | |

| | | | | | | | | | |
|---|---|---|---|---|---|---|---|---|---|
| 29896 | A | G | 3.39E-01 | T | 2.52E-01 | C | 3.21E-03 | three_prime_UTR | |
| 25563 | G | T | 3.11E-01 | C | 1.27E-04 | | | ORF3a | nonsynonymous_Q57H; nonsynonymous_Q57H |
| 29842 | T | G | 2.62E-01 | A | 1.85E-01 | C | 4.28E-04 | three_prime_UTR | |
| 29901 | A | T | 2.62E-01 | C | 3.09E-03 | G | 2.06E-03 | three_prime_UTR | |
| 1059 | C | T | 2.55E-01 | | | | | ORF1ab | nonsynonymous_T265I |
| 29846 | T | A | 2.51E-01 | C | 1.67E-02 | G | 5.77E-04 | three_prime_UTR | |
| 29847 | T | A | 2.50E-01 | G | 5.79E-04 | | | three_prime_UTR | |
| 29848 | T | A | 2.48E-01 | G | 1.75E-03 | | | three_prime_UTR | |
| 29845 | T | A | 2.47E-01 | C | 4.00E-03 | G | 1.72E-03 | three_prime_UTR | |
| 29843 | G | A | 2.46E-01 | C | 1.93E-02 | T | 2.27E-03 | three_prime_UTR | |
| 29837 | C | A | 2.32E-01 | G | 6.35E-03 | T | 2.12E-03 | three_prime_UTR | |
| 28881 | G | A | 1.99E-01 | T | 3.80E-04 | | | N | nonsynonymous_R203K; nonsynonymous_R203M |
| 28882 | G | A | 1.99E-01 | T | 2.53E-04 | | | N | synonymous_R203R; nonsynonymous_R203S |
| 28883 | G | C | 1.99E-01 | A | 1.90E-04 | | | N | nonsynonymous_G204R; nonsynonymous_G204R |
| 29836 | C | A | 1.96E-01 | G | 7.00E-03 | T | 2.47E-03 | three_prime_UTR | |
| 29887 | A | G | 1.61E-01 | T | 5.72E-03 | C | 3.05E-03 | three_prime_UTR | |
| 29891 | A | C | 1.29E-01 | G | 4.34E-03 | T | 3.91E-03 | three_prime_UTR | |
| 29889 | A | G | 1.28E-01 | C | 1.23E-02 | T | 1.32E-03 | three_prime_UTR | |
| 28144 | T | C | 1.22E-01 | | | | | ORF8 | nonsynonymous_L84S |
| 8782 | C | T | 1.21E-01 | | | | | ORF1ab | synonymous_S2839S |
| 29888 | A | T | 1.19E-01 | C | 5.85E-02 | G | 4.18E-02 | three_prime_UTR | |
| 29885 | A | G | 1.05E-01 | T | 3.23E-02 | C | 5.52E-04 | three_prime_UTR | |
| 11083 | G | T | 9.79E-02 | A | 1.27E-04 | C | 6.34E-05 | ORF1ab | nonsynonymous_L3606F; synonymous_L3606L; nonsynonymous_L3606F |
| 18060 | C | T | 7.67E-02 | | | | | ORF1ab | synonymous_L5932L |

| | | | | | | | | | |
|---|---|---|---|---|---|---|---|---|---|
| 17858 | A | G | 7.58E-02 | | | | | ORF1ab | nonsynonymous_Y5865C |
| 17747 | C | T | 7.45E-02 | | | | | ORF1ab | nonsynonymous_P5828L |
| 14805 | C | T | 6.59E-02 | | | | | ORF1ab | synonymous_Y4847Y |
| 26144 | G | T | 6.23E-02 | | | | | ORF3a | nonsynonymous_G251V |
| 29840 | T | A | 6.00E-02 | G | 1.30E-03 | C | 1.30E-04 | three_prime_UTR | |
| 36 | C | T | 5.20E-02 | G | 3.64E-04 | | | five_prime_UTR | |
| 20268 | A | G | 4.94E-02 | | | | | ORF1ab | synonymous_L6668L |
| 29890 | A | G | 4.27E-02 | T | 9.58E-03 | C | 4.36E-04 | three_prime_UTR | |
| 29872 | A | G | 4.00E-02 | T | 7.99E-03 | C | 2.28E-03 | three_prime_UTR | |
| 18877 | C | T | 3.33E-02 | | | | | ORF1ab | synonymous_L6205L |
| 35 | A | T | 3.18E-02 | G | 1.23E-04 | G | 1.23E-04 | five_prime_UTR | |
| 27964 | C | T | 3.09E-02 | | | | | ORF8 | nonsynonymous_S24L |
| 29877 | A | T | 2.59E-02 | G | 7.76E-03 | C | 1.29E-03 | three_prime_UTR | |
| 15324 | C | T | 2.56E-02 | | | | | ORF1ab | synonymous_N5020N |
| 17247 | T | C | 2.49E-02 | | | | | ORF1ab | synonymous_R5661R |
| 2558 | C | T | 2.30E-02 | | | | | ORF1ab | nonsynonymous_P765S |
| 34 | A | T | 2.16E-02 | C | 1.86E-03 | G | 1.24E-04 | five_prime_UTR | |
| 29875 | A | T | 2.14E-02 | G | 1.76E-02 | C | 1.13E-02 | three_prime_UTR | |
| 2480 | A | G | 2.13E-02 | | | | | ORF1ab | nonsynonymous_I739V |
| 29873 | A | G | 2.13E-02 | C | 1.30E-02 | T | 3.54E-03 | three_prime_UTR | |
| 29870 | C | A | 2.12E-02 | T | 1.24E-02 | G | 8.85E-03 | three_prime_UTR | |
| 29861 | G | A | 2.08E-02 | T | 3.33E-03 | C | 2.08E-03 | three_prime_UTR | |
| 29862 | G | A | 1.98E-02 | T | 2.11E-03 | C | 8.43E-04 | three_prime_UTR | |
| 28854 | C | T | 1.98E-02 | | | | | N | nonsynonymous_S194L |
| 29844 | A | T | 1.89E-02 | G | 2.86E-03 | C | 1.72E-03 | three_prime_UTR | |
| 11916 | C | T | 1.87E-02 | | | | | ORF1ab | nonsynonymous_S3884L |
| 29553 | G | A | 1.85E-02 | | | | | intergenic | |
| 29851 | T | A | 1.84E-02 | G | 9.75E-03 | C | 1.62E-03 | three_prime_UTR | |
| 2416 | C | T | 1.78E-02 | | | | | ORF1ab | synonymous_Y717Y |
| 27046 | C | T | 1.76E-02 | | | | | M | nonsynonymous_T175M |

| | | | | | | | | | |
|---|---|---|---|---|---|---|---|---|---|
| 29850 | A | G | 1.75E-02 | T | 3.03E-03 | C | 2.28E-03 | three_prime_UTR | |
| 37 | C | A | 1.73E-02 | T | 2.92E-03 | G | 1.22E-04 | five_prime_UTR | |
| 25429 | G | T | 1.64E-02 | A | 6.32E-05 | | | ORF3a | nonsynonymous_V13L; nonsynonymous_V13I |
| 28311 | C | T | 1.56E-02 | G | 1.90E-04 | | | N | nonsynonymous_P13L; nonsynonymous_P13R |
| 29876 | A | C | 1.53E-02 | G | 1.41E-02 | T | 7.67E-03 | three_prime_UTR | |
| 13730 | C | T | 1.53E-02 | | | | | ORF1ab | nonsynonymous_A4489V |
| 23731 | C | T | 1.46E-02 | | | | | S | synonymous_T723T |
| 10097 | G | A | 1.42E-02 | | | | | ORF1ab | nonsynonymous_G3278S |
| 6312 | C | A | 1.35E-02 | T | 1.90E-04 | T | 1.90E-04 | ORF1ab | nonsynonymous_T2016K; nonsynonymous_T2016I; nonsynonymous_T2016I |
| 29874 | A | T | 1.34E-02 | T | 1.34E-02 | C | 1.22E-03 | three_prime_UTR | |
| 29540 | G | A | 1.33E-02 | T | 5.06E-04 | | | intergenic | |
| 23929 | C | T | 1.33E-02 | | | | | S | synonymous_Y789Y |
| 18998 | C | T | 1.32E-02 | | | | | ORF1ab | nonsynonymous_A6245V |
| 29897 | A | T | 1.32E-02 | G | 5.26E-03 | G | 5.26E-03 | three_prime_UTR | |
| 24034 | C | T | 1.30E-02 | | | | | S | synonymous_N824N |
| 1440 | G | A | 1.29E-02 | | | | | ORF1ab | nonsynonymous_G392D |
| 29868 | G | A | 1.24E-02 | C | 8.88E-03 | T | 5.33E-03 | three_prime_UTR | |
| 26735 | C | T | 1.23E-02 | | | | | M | synonymous_Y71Y |
| 29869 | A | T | 1.22E-02 | G | 6.99E-03 | C | 8.73E-04 | three_prime_UTR | |
| 28077 | G | C | 1.22E-02 | T | 2.34E-03 | | | ORF8 | nonsynonymous_V62L; nonsynonymous_V62L |
| 2891 | G | A | 1.22E-02 | | | | | ORF1ab | nonsynonymous_A876T |
| 29700 | A | G | 1.20E-02 | | | | | three_prime_UTR | |
| 26729 | T | C | 1.20E-02 | | | | | M | synonymous_A69A |
| 3177 | C | T | 1.18E-02 | A | 1.90E-04 | | | ORF1ab | nonsynonymous_P971L; nonsynonymous_P971H |

| | | | | | | | | | |
|---|---|---|---|---|---|---|---|---|---|
| 313 | C | T | 1.17E-02 | | | | | ORF1ab | synonymous_L16L |
| 29878 | A | T | 1.16E-02 | C | 6.87E-03 | G | 3.17E-03 | three_prime_UTR | |
| 490 | T | A | 1.15E-02 | | | | | ORF1ab | nonsynonymous_D75E |
| 18736 | T | C | 1.11E-02 | A | 6.32E-05 | | | ORF1ab | nonsynonymous_F6158L; nonsynonymous_F6158I |
| 1397 | G | A | 1.09E-02 | | | | | ORF1ab | nonsynonymous_V378I |
| 29742 | G | T | 1.07E-02 | A | 6.90E-03 | | | three_prime_UTR | |
| 28688 | T | C | 1.05E-02 | | | | | N | synonymous_L139L |
| 29881 | A | G | 1.01E-02 | T | 2.19E-03 | C | 1.92E-03 | three_prime_UTR | |
| 19839 | T | C | 9.55E-03 | | | | | ORF1ab | synonymous_N6525N |
| 26530 | A | G | 9.43E-03 | | | | | M | nonsynonymous_D3G |
| 29853 | G | A | 9.41E-03 | T | 9.25E-04 | C | 4.63E-04 | three_prime_UTR | |
| 12 | A | T | 8.94E-03 | G | 2.56E-03 | G | 2.56E-03 | five_prime_UTR | |
| 28657 | C | T | 8.92E-03 | | | | | N | synonymous_D128D |
| 28863 | C | T | 8.54E-03 | | | | | N | nonsynonymous_S197L |
| 29711 | G | T | 8.29E-03 | C | 6.38E-05 | | | three_prime_UTR | |
| 9477 | T | A | 8.28E-03 | | | | | ORF1ab | nonsynonymous_F3071Y |
| 29734 | G | C | 8.17E-03 | T | 3.19E-04 | A | 1.28E-04 | three_prime_UTR | |
| 25979 | G | T | 8.16E-03 | | | | | ORF3a | nonsynonymous_G196V |
| 29855 | T | A | 8.12E-03 | C | 2.07E-03 | G | 7.96E-04 | three_prime_UTR | |
| 29867 | T | A | 7.99E-03 | C | 2.66E-03 | G | 8.88E-04 | three_prime_UTR | |
| 31 | A | T | 7.92E-03 | G | 6.38E-04 | G | 6.38E-04 | five_prime_UTR | |
| 29854 | C | A | 7.78E-03 | G | 1.71E-03 | T | 9.33E-04 | three_prime_UTR | |
| 29858 | T | A | 7.72E-03 | C | 2.63E-03 | G | 9.27E-04 | three_prime_UTR | |
| 13536 | C | T | 7.72E-03 | | | | | ORF1ab | synonymous_Y4424Y |
| 24368 | G | T | 7.65E-03 | C | 6.32E-05 | | | S | nonsynonymous_D936Y; nonsynonymous_D936H |
| 514 | T | C | 7.54E-03 | | | | | ORF1ab | synonymous_H83H |
| 28836 | C | T | 7.53E-03 | | | | | N | nonsynonymous_S188L |
| 13862 | C | T | 7.33E-03 | | | | | ORF1ab | nonsynonymous_T4533I |

| | | | | | | | | | |
|---|---|---|---|---|---|---|---|---|---|
| 16289 | C | T | 7.27E-03 | | | | | ORF1ab | nonsynonymous_A5342V |
| 1302 | C | T | 7.27E-03 | | | | | ORF1ab | nonsynonymous_T346I |
| 11 | T | G | 7.24E-03 | C | 2.72E-03 | A | 1.81E-03 | five_prime_UTR | |
| 24694 | A | T | 6.96E-03 | | | | | S | synonymous_G1044G |
| 4002 | C | T | 6.89E-03 | | | | | ORF1ab | nonsynonymous_T1246I |
| 29863 | A | T | 6.64E-03 | G | 2.21E-03 | | | three_prime_UTR | |
| 29856 | T | A | 6.59E-03 | C | 1.12E-03 | G | 8.03E-04 | three_prime_UTR | |
| 187 | A | G | 6.58E-03 | | | | | five_prime_UTR | |
| 6040 | C | T | 6.58E-03 | | | | | ORF1ab | synonymous_F1925F |
| 29859 | T | A | 6.27E-03 | C | 5.49E-03 | G | 9.41E-04 | three_prime_UTR | |
| 21575 | C | T | 6.27E-03 | | | | | S | nonsynonymous_L5F |
| 5 | A | T | 6.19E-03 | G | 1.82E-03 | C | 1.46E-03 | five_prime_UTR | |
| 19524 | C | T | 6.14E-03 | | | | | ORF1ab | synonymous_L6420L |
| 29857 | C | A | 6.14E-03 | T | 2.61E-03 | G | 2.30E-03 | three_prime_UTR | |
| 28878 | G | A | 5.88E-03 | C | 3.16E-04 | T | 1.26E-04 | N | nonsynonymous_S202N; nonsynonymous_S202T; nonsynonymous_S202I |
| 24862 | A | G | 5.75E-03 | | | | | S | synonymous_T1100T |
| 29866 | A | T | 5.74E-03 | G | 4.41E-03 | C | 3.53E-03 | three_prime_UTR | |
| 25350 | C | T | 5.63E-03 | | | | | S | nonsynonymous_P1263L |
| 10323 | A | G | 5.63E-03 | | | | | ORF1ab | nonsynonymous_K3353R |
| 29871 | A | G | 5.46E-03 | T | 2.19E-03 | T | 2.19E-03 | three_prime_UTR | |
| 10798 | C | A | 5.44E-03 | | | | | ORF1ab | nonsynonymous_D3511E |
| 833 | T | C | 5.37E-03 | | | | | ORF1ab | nonsynonymous_F190L |
| 14786 | C | T | 5.31E-03 | | | | | ORF1ab | nonsynonymous_A4841V |
| 379 | C | A | 4.99E-03 | | | | | ORF1ab | synonymous_V38V |
| 28580 | G | T | 4.87E-03 | A | 2.53E-04 | | | N | nonsynonymous_D103Y; nonsynonymous_D103N |
| 29816 | T | A | 4.84E-03 | G | 6.82E-05 | | | three_prime_UTR | |

| | | | | | | | | | |
|---|---|---|---|---|---|---|---|---|---|
| 28851 | G | T | 4.81E-03 | C | 3.79E-04 | | | N | nonsynonymous_S193I; nonsynonymous_S193T |
| 2 | T | A | 4.76E-03 | C | 3.71E-03 | G | 3.18E-03 | five_prime_UTR | |
| 10319 | C | T | 4.74E-03 | | | | | ORF1ab | nonsynonymous_L3352F |
| 29817 | T | A | 4.71E-03 | C | 2.05E-04 | G | 1.36E-04 | three_prime_UTR | |
| 25688 | C | T | 4.68E-03 | A | 6.96E-04 | | | ORF3a | nonsynonymous_A99V; nonsynonymous_A99D |
| 29894 | A | T | 4.67E-03 | T | 4.67E-03 | C | 1.56E-03 | three_prime_UTR | |
| 18 | T | G | 4.67E-03 | A | 1.23E-03 | | | five_prime_UTR | |
| 26088 | C | T | 4.62E-03 | | | | | ORF3a | synonymous_I232I |
| 6310 | C | A | 4.55E-03 | T | 1.20E-03 | | | ORF1ab | nonsynonymous_S2015R; synonymous_S2015S |
| 28896 | C | G | 4.49E-03 | | | | | N | nonsynonymous_A208G |
| 12478 | G | A | 4.49E-03 | | | | | ORF1ab | nonsynonymous_M4071I |
| 29892 | A | G | 4.46E-03 | G | 4.46E-03 | T | 2.98E-03 | three_prime_UTR | |
| 27635 | C | T | 4.31E-03 | | | | | ORF7a | nonsynonymous_S81L |
| 11320 | T | C | 4.30E-03 | | | | | ORF1ab | synonymous_Y3685Y |
| 29095 | C | T | 4.24E-03 | | | | | N | synonymous_F274F |
| 20 | C | G | 4.21E-03 | T | 7.65E-04 | A | 1.91E-04 | five_prime_UTR | |
| 19 | C | G | 4.17E-03 | T | 2.70E-03 | A | 1.47E-03 | five_prime_UTR | |
| 1917 | C | T | 4.17E-03 | | | | | ORF1ab | nonsynonymous_T551I |
| 13 | T | A | 4.08E-03 | C | 3.67E-03 | G | 2.04E-03 | five_prime_UTR | |
| 19170 | C | T | 4.05E-03 | A | 6.32E-05 | | | ORF1ab | synonymous_F6302F; nonsynonymous_F6302L |
| 7834 | C | T | 4.05E-03 | | | | | ORF1ab | synonymous_N2523N |
| 22444 | C | T | 3.99E-03 | | | | | S | synonymous_D294D |
| 17410 | C | T | 3.92E-03 | A | 4.43E-04 | | | ORF1ab | nonsynonymous_R5716C; nonsynonymous_R5716S |
| 27384 | T | C | 3.86E-03 | | | | | ORF6 | synonymous_D61D |
| 25433 | C | T | 3.86E-03 | | | | | ORF3a | nonsynonymous_T14I |

| | | | | | | | | | |
|---|---|---|---|---|---|---|---|---|---|
| 19509 | G | A | 3.80E-03 | | | | | ORF1ab | synonymous_E6415E |
| 28 | C | T | 3.80E-03 | A | 1.46E-04 | | | five_prime_UTR | |
| 2244 | T | C | 3.79E-03 | | | | | ORF1ab | nonsynonymous_V660A |
| 19684 | G | T | 3.73E-03 | | | | | ORF1ab | nonsynonymous_V6474L |
| 20755 | A | C | 3.60E-03 | | | | | ORF1ab | nonsynonymous_S6831R |
| 9223 | C | T | 3.54E-03 | | | | | ORF1ab | synonymous_H2986H |
| 10582 | C | T | 3.54E-03 | | | | | ORF1ab | synonymous_D3439D |
| 1515 | A | G | 3.48E-03 | | | | | ORF1ab | nonsynonymous_H417R |
| 884 | C | T | 3.48E-03 | | | | | ORF1ab | nonsynonymous_R207C |
| 29852 | A | T | 3.46E-03 | G | 1.73E-04 | | | three_prime_UTR | |
| 8653 | G | T | 3.42E-03 | | | | | ORF1ab | nonsynonymous_M2796I |
| 26951 | G | A | 3.41E-03 | T | 6.96E-04 | C | 1.90E-04 | M | synonymous_V143V; synonymous_V143V; synonymous_V143V |
| 16887 | C | T | 3.41E-03 | | | | | ORF1ab | synonymous_Y5541Y |
| 29884 | A | G | 3.35E-03 | T | 2.93E-03 | C | 1.26E-03 | three_prime_UTR | |
| 8788 | T | C | 3.35E-03 | | | | | ORF1ab | synonymous_R2841R |
| 29798 | G | A | 3.30E-03 | | | | | three_prime_UTR | |
| 29451 | C | T | 3.29E-03 | | | | | N | nonsynonymous_T393I |
| 13006 | T | C | 3.29E-03 | | | | | ORF1ab | synonymous_A4247A |
| 3373 | C | A | 3.22E-03 | T | 6.32E-05 | | | ORF1ab | nonsynonymous_D1036E; synonymous_D1036D |
| 10851 | C | T | 3.22E-03 | | | | | ORF1ab | nonsynonymous_A3529V |
| 7765 | C | T | 3.22E-03 | | | | | ORF1ab | synonymous_S2500S |
| 8950 | C | T | 3.22E-03 | | | | | ORF1ab | synonymous_I2895I |
| 14 | A | T | 3.13E-03 | G | 1.56E-03 | | | five_prime_UTR | |
| 29895 | A | T | 3.12E-03 | T | 3.12E-03 | C | 2.08E-03 | three_prime_UTR | |
| 28887 | C | T | 3.10E-03 | | | | | N | nonsynonymous_T205I |
| 15 | C | T | 3.07E-03 | A | 1.15E-03 | G | 3.84E-04 | five_prime_UTR | |
| 20578 | G | T | 3.04E-03 | | | | | ORF1ab | nonsynonymous_V6772L |

| | | | | | | | | | |
|---|---|---|---|---|---|---|---|---|---|
| 12710 | C | T | 3.03E-03 | | | | | ORF1ab | synonymous_L4149L |
| 26750 | C | T | 2.91E-03 | | | | | M | synonymous_I76I |
| 29860 | A | C | 2.86E-03 | T | 2.45E-03 | G | 4.09E-04 | three_prime_UTR | |
| 8389 | C | T | 2.85E-03 | | | | | ORF1ab | synonymous_N2708N |
| 7011 | C | T | 2.78E-03 | | | | | ORF1ab | nonsynonymous_A2249V |
| 12880 | C | T | 2.78E-03 | | | | | ORF1ab | synonymous_I4205I |
| 16260 | C | T | 2.72E-03 | | | | | ORF1ab | synonymous_C5332C |
| 2113 | C | T | 2.72E-03 | | | | | ORF1ab | synonymous_I616I |
| 1076 | C | T | 2.72E-03 | | | | | ORF1ab | nonsynonymous_P271S |
| 12832 | G | A | 2.72E-03 | T | 6.32E-05 | | | ORF1ab | synonymous_Q4189Q; nonsynonymous_Q4189H |
| 29795 | G | A | 2.69E-03 | | | | | three_prime_UTR | |
| 29810 | G | A | 2.66E-03 | C | 2.73E-04 | T | 6.82E-05 | three_prime_UTR | |
| 20270 | C | T | 2.66E-03 | | | | | ORF1ab | nonsynonymous_A6669V |
| 21707 | C | T | 2.66E-03 | | | | | S | nonsynonymous_H49Y |
| 18788 | C | T | 2.66E-03 | | | | | ORF1ab | nonsynonymous_T6175I |
| 1401 | G | A | 2.66E-03 | | | | | ORF1ab | nonsynonymous_G379E |
| 4255 | G | T | 2.66E-03 | A | 1.26E-04 | | | ORF1ab | synonymous_P1330P; synonymous_P1330P |
| 6693 | A | G | 2.66E-03 | | | | | ORF1ab | nonsynonymous_K2143R |
| 29800 | G | A | 2.63E-03 | | | | | three_prime_UTR | |
| 20134 | G | T | 2.59E-03 | A | 6.32E-05 | | | ORF1ab | nonsynonymous_V6624L; nonsynonymous_V6624I |
| 2940 | C | T | 2.59E-03 | | | | | ORF1ab | nonsynonymous_P892L |
| 10188 | C | T | 2.59E-03 | | | | | ORF1ab | nonsynonymous_T3308I |
| 21137 | A | G | 2.53E-03 | | | | | ORF1ab | nonsynonymous_K6958R |
| 24197 | G | T | 2.53E-03 | A | 1.26E-04 | | | S | nonsynonymous_A879S; nonsynonymous_A879T |
| 24781 | G | T | 2.53E-03 | | | | | S | nonsynonymous_K1073N |
| 5142 | C | T | 2.53E-03 | | | | | ORF1ab | nonsynonymous_T1626I |

| | | | | | | | | | |
|---|---|---|---|---|---|---|---|---|---|
| 9 | T | A | 2.53E-03 | G | 2.21E-03 | C | 1.58E-03 | five_prime_UTR | |
| 24982 | T | C | 2.49E-03 | | | | | S | synonymous_P1140P |
| 22468 | G | T | 2.47E-03 | | | | | S | synonymous_T302T |
| 29148 | T | C | 2.47E-03 | | | | | N | nonsynonymous_I292T |
| 17690 | C | T | 2.47E-03 | | | | | ORF1ab | nonsynonymous_S5809L |
| 23707 | C | T | 2.47E-03 | | | | | S | synonymous_P715P |
| 4540 | C | T | 2.47E-03 | | | | | ORF1ab | synonymous_Y1425Y |
| 29811 | T | A | 2.46E-03 | G | 1.36E-04 | C | 6.82E-05 | three_prime_UTR | |
| 29809 | T | A | 2.39E-03 | C | 6.82E-05 | | | three_prime_UTR | |
| 11417 | G | T | 2.34E-03 | C | 1.26E-04 | | | ORF1ab | nonsynonymous_V3718F; nonsynonymous_V3718L |
| 28821 | C | A | 2.28E-03 | | | | | N | nonsynonymous_S183Y |
| 3871 | G | T | 2.28E-03 | | | | | ORF1ab | nonsynonymous_K1202N |
| 14877 | C | T | 2.28E-03 | | | | | ORF1ab | synonymous_Y4871Y |
| 21 | C | A | 2.28E-03 | T | 1.90E-03 | G | 1.52E-03 | five_prime_UTR | |
| 29849 | A | T | 2.27E-03 | C | 1.51E-03 | | | three_prime_UTR | |
| 16 | C | G | 2.27E-03 | A | 1.51E-03 | T | 3.78E-04 | five_prime_UTR | |
| 23 | G | A | 2.24E-03 | C | 1.12E-03 | | | five_prime_UTR | |
| 15380 | G | T | 2.21E-03 | | | | | ORF1ab | nonsynonymous_S5039I |
| 7438 | T | C | 2.21E-03 | | | | | ORF1ab | synonymous_Y2391Y |
| 7560 | G | A | 2.21E-03 | | | | | ORF1ab | nonsynonymous_R2432K |
| 25 | T | G | 2.18E-03 | A | 1.46E-03 | C | 1.82E-04 | five_prime_UTR | |
| 11074 | C | T | 2.16E-03 | | | | | ORF1ab | synonymous_F3603F |
| 19999 | G | T | 2.15E-03 | | | | | ORF1ab | nonsynonymous_V6579F |
| 8371 | G | T | 2.15E-03 | | | | | ORF1ab | nonsynonymous_Q2702H |
| 29808 | G | A | 2.11E-03 | | | | | three_prime_UTR | |
| 25669 | C | T | 2.09E-03 | | | | | ORF3a | nonsynonymous_H93Y |
| 21452 | G | T | 2.09E-03 | | | | | ORF1ab | nonsynonymous_G7063V |
| 27299 | T | C | 2.09E-03 | | | | | ORF6 | nonsynonymous_I33T |
| 10948 | A | G | 2.09E-03 | | | | | ORF1ab | synonymous_R3561R |

| | | | | | | | | | |
|---|---|---|---|---|---|---|---|---|---|
| 26233 | G | T | 2.09E-03 | | | | | intergenic | |
| 6027 | C | T | 2.09E-03 | | | | | ORF1ab | nonsynonymous_P1921L |
| 20148 | C | T | 2.02E-03 | | | | | ORF1ab | synonymous_F6628F |
| 17010 | C | T | 2.02E-03 | A | 9.49E-04 | | | ORF1ab | synonymous_I5582I; synonymous_I5582I |
| 6990 | C | T | 2.02E-03 | | | | | ORF1ab | nonsynonymous_S2242F |
| 15354 | T | C | 2.02E-03 | A | 6.32E-05 | | | ORF1ab | synonymous_L5030L; synonymous_L5030L |
| 1820 | G | A | 2.02E-03 | | | | | ORF1ab | nonsynonymous_G519S |
| 29807 | T | A | 1.97E-03 | C | 2.72E-04 | | | three_prime_UTR | |
| 29188 | A | T | 1.96E-03 | | | | | N | synonymous_A305A |
| 29353 | C | T | 1.96E-03 | | | | | N | synonymous_Y360Y |
| 10265 | G | A | 1.96E-03 | | | | | ORF1ab | nonsynonymous_G3334S |
| 9598 | C | T | 1.96E-03 | | | | | ORF1ab | synonymous_Y3111Y |
| 19488 | C | T | 1.90E-03 | | | | | ORF1ab | synonymous_V6408V |
| 29144 | C | T | 1.90E-03 | | | | | N | synonymous_L291L |
| 16381 | G | A | 1.90E-03 | | | | | ORF1ab | nonsynonymous_V5373I |
| 4390 | G | T | 1.90E-03 | | | | | ORF1ab | nonsynonymous_L1375F |
| 17 | T | C | 1.86E-03 | A | 1.11E-03 | G | 7.42E-04 | five_prime_UTR | |
| 21614 | C | T | 1.83E-03 | | | | | S | nonsynonymous_L18F |
| 15540 | C | T | 1.83E-03 | | | | | ORF1ab | synonymous_V5092V |
| 23947 | A | G | 1.83E-03 | | | | | S | synonymous_K795K |
| 14073 | T | C | 1.83E-03 | | | | | ORF1ab | synonymous_D4603D |
| 17126 | T | C | 1.83E-03 | | | | | ORF1ab | nonsynonymous_L5621P |
| 2942 | C | T | 1.83E-03 | | | | | ORF1ab | synonymous_L893L |
| 1594 | C | T | 1.83E-03 | | | | | ORF1ab | synonymous_S443S |
| 1912 | C | T | 1.83E-03 | | | | | ORF1ab | synonymous_S549S |
| 10507 | C | T | 1.83E-03 | | | | | ORF1ab | synonymous_N3414N |
| 27669 | A | G | 1.77E-03 | T | 6.33E-05 | | | ORF7a | synonymous_E92E; nonsynonymous_E92D |

| | | | | | | | | | |
|---|---|---|---|---|---|---|---|---|---|
| 2836 | C | T | 1.77E-03 | | | | | ORF1ab | synonymous_C857C |
| 3564 | G | T | 1.77E-03 | | | | | ORF1ab | nonsynonymous_G1100V |
| 25692 | C | T | 1.77E-03 | A | 6.32E-05 | | | ORF3a | synonymous_G100G; synonymous_G100G |
| 21648 | C | T | 1.77E-03 | | | | | S | nonsynonymous_T29I |
| 26681 | C | T | 1.77E-03 | | | | | M | synonymous_F53F |
| 12781 | C | T | 1.77E-03 | | | | | ORF1ab | synonymous_Y4172Y |
| 7420 | C | T | 1.77E-03 | | | | | ORF1ab | synonymous_I2385I |
| 7479 | A | G | 1.77E-03 | | | | | ORF1ab | nonsynonymous_N2405S |
| 29 | A | C | 1.74E-03 | T | 2.90E-04 | G | 1.45E-04 | five_prime_UTR | |
| 26 | A | T | 1.72E-03 | G | 5.16E-04 | C | 3.44E-04 | five_prime_UTR | |
| 28086 | G | T | 1.71E-03 | | | | | ORF8 | nonsynonymous_A65S |
| 13402 | T | G | 1.71E-03 | | | | | ORF1ab | stopgain_Y4379X |
| 25552 | G | T | 1.71E-03 | | | | | ORF3a | nonsynonymous_A54S |
| 13627 | G | T | 1.71E-03 | | | | | ORF1ab | nonsynonymous_D4455Y |
| 24047 | G | A | 1.71E-03 | | | | | S | nonsynonymous_A829T |
| 24054 | C | T | 1.71E-03 | | | | | S | nonsynonymous_A831V |
| 28338 | A | G | 1.71E-03 | | | | | N | nonsynonymous_D22G |
| 6696 | C | T | 1.71E-03 | | | | | ORF1ab | nonsynonymous_P2144L |
| 29839 | A | T | 1.70E-03 | G | 1.57E-03 | C | 7.27E-04 | three_prime_UTR | |
| 29683 | A | T | 1.65E-03 | | | | | three_prime_UTR | |
| 29825 | G | A | 1.65E-03 | T | 1.37E-04 | | | three_prime_UTR | |
| 28842 | G | T | 1.64E-03 | | | | | N | nonsynonymous_S190I |
| 27281 | G | T | 1.64E-03 | | | | | ORF6 | nonsynonymous_W27L |
| 28519 | C | T | 1.64E-03 | | | | | N | synonymous_D82D |
| 6573 | C | T | 1.64E-03 | | | | | ORF1ab | nonsynonymous_S2103F |
| 14937 | C | T | 1.64E-03 | A | 6.32E-05 | | | ORF1ab | synonymous_D4891D; nonsynonymous_D4891E |
| 12915 | C | T | 1.64E-03 | | | | | ORF1ab | nonsynonymous_T4217I |

| | | | | | | | | | |
|---|---|---|---|---|---|---|---|---|---|
| 2164 | G | A | 1.64E-03 | T | 1.26E-04 | | | ORF1ab | synonymous_E633E; nonsynonymous_E633D |
| 13568 | C | T | 1.58E-03 | | | | | ORF1ab | nonsynonymous_A4435V |
| 23010 | T | C | 1.58E-03 | | | | | S | nonsynonymous_V483A |
| 9438 | C | T | 1.58E-03 | | | | | ORF1ab | nonsynonymous_T3058I |
| 5062 | G | T | 1.58E-03 | A | 1.90E-04 | | | ORF1ab | nonsynonymous_L1599F; synonymous_L1599L |
| 25958 | A | G | 1.58E-03 | | | | | ORF3a | nonsynonymous_Y189C |
| 15243 | G | T | 1.58E-03 | | | | | ORF1ab | nonsynonymous_M4993I |
| 15760 | G | A | 1.58E-03 | | | | | ORF1ab | nonsynonymous_G5166S |
| 17373 | C | T | 1.58E-03 | | | | | ORF1ab | synonymous_A5703A |
| 17877 | T | C | 1.58E-03 | | | | | ORF1ab | synonymous_T5871T |
| 25500 | G | T | 1.58E-03 | A | 1.26E-04 | | | ORF3a | synonymous_P36P; synonymous_P36P |
| 11801 | G | A | 1.58E-03 | | | | | ORF1ab | nonsynonymous_G3846S |
| 1427 | C | T | 1.58E-03 | | | | | ORF1ab | nonsynonymous_H388Y |
| 4402 | T | C | 1.58E-03 | | | | | ORF1ab | synonymous_L1379L |
| 895 | A | G | 1.58E-03 | | | | | ORF1ab | synonymous_K210K |
| 27 | A | C | 1.54E-03 | G | 3.43E-04 | T | 1.71E-04 | five_prime_UTR | |
| 24933 | G | T | 1.52E-03 | | | | | S | nonsynonymous_G1124V |
| 26447 | C | T | 1.52E-03 | G | 1.90E-04 | | | E | nonsynonymous_S68F; nonsynonymous_S68C |
| 9479 | G | T | 1.52E-03 | | | | | ORF1ab | nonsynonymous_G3072C |
| 23185 | C | T | 1.52E-03 | | | | | S | synonymous_F541F |
| 25947 | G | T | 1.52E-03 | C | 1.90E-04 | | | ORF3a | nonsynonymous_Q185H; nonsynonymous_Q185H |
| 11001 | C | T | 1.52E-03 | | | | | ORF1ab | nonsynonymous_T3579I |
| 15438 | G | T | 1.52E-03 | | | | | ORF1ab | nonsynonymous_M5058I |
| 8078 | C | T | 1.52E-03 | G | 6.32E-05 | | | ORF1ab | nonsynonymous_P2605S; nonsynonymous_P2605A |

| | | | | | | | | | |
|---|---|---|---|---|---|---|---|---|---|
| 10712 | C | T | 1.52E-03 | | | | | ORF1ab | nonsynonymous_L3483F |
| 29775 | T | A | 1.50E-03 | C | 1.67E-04 | | | three_prime_UTR | |
| 20316 | C | T | 1.45E-03 | | | | | ORF1ab | synonymous_F6684F |
| 25728 | C | T | 1.45E-03 | | | | | ORF3a | synonymous_V112V |
| 25916 | C | T | 1.45E-03 | | | | | ORF3a | nonsynonymous_T175I |
| 28899 | G | C | 1.45E-03 | T | 3.16E-04 | A | 1.26E-04 | N | nonsynonymous_R209T; nonsynonymous_R209I; nonsynonymous_R209K |
| 25904 | C | T | 1.45E-03 | | | | | ORF3a | nonsynonymous_S171L |
| 17470 | C | T | 1.45E-03 | | | | | ORF1ab | synonymous_L5736L |
| 16912 | G | T | 1.45E-03 | | | | | ORF1ab | nonsynonymous_V5550L |
| 18713 | C | T | 1.45E-03 | A | 6.32E-05 | | | ORF1ab | nonsynonymous_A6150V; nonsynonymous_A6150D |
| 25572 | C | T | 1.45E-03 | | | | | ORF3a | synonymous_S60S |
| 23575 | C | T | 1.45E-03 | | | | | S | synonymous_C671C |
| 28344 | C | A | 1.45E-03 | T | 1.26E-04 | | | N | nonsynonymous_T24N; nonsynonymous_T24I |
| 11109 | C | T | 1.45E-03 | | | | | ORF1ab | nonsynonymous_A3615V |
| 7423 | C | T | 1.45E-03 | | | | | ORF1ab | synonymous_F2386F |
| 11195 | C | T | 1.45E-03 | | | | | ORF1ab | nonsynonymous_L3644F |
| 3311 | C | A | 1.45E-03 | | | | | ORF1ab | nonsynonymous_L1016I |
| 29778 | G | A | 1.45E-03 | | | | | three_prime_UTR | |
| 29826 | T | A | 1.44E-03 | C | 2.06E-04 | G | 6.86E-05 | three_prime_UTR | |
| 29820 | T | A | 1.44E-03 | | | | | three_prime_UTR | |
| 29822 | T | A | 1.44E-03 | | | | | three_prime_UTR | |
| 29791 | T | A | 1.41E-03 | C | 2.69E-04 | | | three_prime_UTR | |
| 27869 | T | C | 1.39E-03 | | | | | ORF7b | synonymous_N38N |
| 28378 | G | T | 1.39E-03 | A | 6.32E-05 | C | 6.32E-05 | N | synonymous_A35A; synonymous_A35A; synonymous_A35A |

| | | | | | | | | | |
|---|---|---|---|---|---|---|---|---|---|
| 24077 | G | T | 1.39E-03 | | | | | S | nonsynonymous_D839Y |
| 17671 | G | T | 1.39E-03 | | | | | ORF1ab | nonsynonymous_V5803F |
| 25523 | G | T | 1.39E-03 | | | | | ORF3a | nonsynonymous_G44V |
| 3096 | C | T | 1.39E-03 | | | | | ORF1ab | nonsynonymous_S944L |
| 4320 | C | T | 1.39E-03 | | | | | ORF1ab | nonsynonymous_A1352V |
| 4838 | A | G | 1.39E-03 | | | | | ORF1ab | nonsynonymous_I1525V |
| 29823 | T | A | 1.37E-03 | C | 1.37E-04 | | | three_prime_UTR | |
| 29821 | T | A | 1.37E-03 | | | | | three_prime_UTR | |
| 29794 | G | A | 1.37E-03 | C | 8.04E-05 | | | three_prime_UTR | |
| 27542 | C | T | 1.33E-03 | | | | | ORF7a | nonsynonymous_A50V |
| 28812 | G | T | 1.33E-03 | | | | | N | nonsynonymous_S180I |
| 29200 | C | T | 1.33E-03 | | | | | N | synonymous_P309P |
| 26461 | C | T | 1.33E-03 | G | 6.32E-05 | | | E | nonsynonymous_L73F; nonsynonymous_L73V |
| 15720 | C | T | 1.33E-03 | | | | | ORF1ab | synonymous_D5152D |
| 10771 | T | C | 1.33E-03 | | | | | ORF1ab | synonymous_Y3502Y |
| 14184 | C | T | 1.33E-03 | | | | | ORF1ab | synonymous_T4640T |
| 14747 | A | G | 1.33E-03 | | | | | ORF1ab | nonsynonymous_E4828G |
| 15277 | C | T | 1.33E-03 | | | | | ORF1ab | nonsynonymous_H5005Y |
| 25528 | C | T | 1.33E-03 | | | | | ORF3a | nonsynonymous_L46F |
| 1684 | C | T | 1.33E-03 | | | | | ORF1ab | synonymous_I473I |
| 8917 | C | T | 1.33E-03 | | | | | ORF1ab | synonymous_F2884F |
| 1437 | C | T | 1.33E-03 | | | | | ORF1ab | nonsynonymous_S391F |
| 5784 | C | T | 1.33E-03 | | | | | ORF1ab | nonsynonymous_T1840I |
| 22 | A | T | 1.31E-03 | C | 1.13E-03 | G | 5.63E-04 | five_prime_UTR | |
| 29781 | G | A | 1.27E-03 | T | 2.00E-04 | | | three_prime_UTR | |
| 27804 | C | T | 1.27E-03 | | | | | ORF7b | synonymous_L17L |
| 28253 | C | T | 1.27E-03 | A | 5.06E-04 | G | 6.33E-05 | ORF8 | synonymous_F120F; nonsynonymous_F120L; nonsynonymous_F120L |

| | | | | | | | | | |
|---|---|---|---|---|---|---|---|---|---|
| 9286 | C | T | 1.26E-03 | | | | | ORF1ab | synonymous_N3007N |
| 14724 | C | T | 1.26E-03 | | | | | ORF1ab | synonymous_F4820F |
| 15842 | C | A | 1.26E-03 | | | | | ORF1ab | nonsynonymous_T5193N |
| 21724 | G | T | 1.26E-03 | C | 6.32E-05 | | | S | nonsynonymous_L54F; nonsynonymous_L54F |
| 24232 | A | G | 1.26E-03 | | | | | S | synonymous_A890A |
| 15960 | C | T | 1.26E-03 | | | | | ORF1ab | synonymous_A5232A |
| 18398 | T | C | 1.26E-03 | | | | | ORF1ab | nonsynonymous_V6045A |
| 21058 | C | T | 1.26E-03 | A | 6.32E-05 | | | ORF1ab | nonsynonymous_P6932S; nonsynonymous_P6932T |
| 23277 | C | T | 1.26E-03 | | | | | S | nonsynonymous_T572I |
| 2018 | A | G | 1.26E-03 | | | | | ORF1ab | nonsynonymous_M585V |
| 3100 | T | G | 1.26E-03 | | | | | ORF1ab | synonymous_T945T |
| 658 | T | C | 1.26E-03 | | | | | ORF1ab | synonymous_A131A |
| 4113 | C | T | 1.26E-03 | | | | | ORF1ab | nonsynonymous_A1283V |
| 619 | C | T | 1.26E-03 | | | | | ORF1ab | synonymous_Y118Y |
| 29779 | G | A | 1.25E-03 | T | 3.96E-04 | C | 1.32E-04 | three_prime_UTR | |
| 29777 | G | A | 1.25E-03 | T | 1.31E-04 | C | 6.57E-05 | three_prime_UTR | |
| 29774 | C | A | 1.25E-03 | T | 7.87E-04 | | | three_prime_UTR | |
| 29828 | G | A | 1.24E-03 | T | 6.88E-05 | | | three_prime_UTR | |
| 19484 | C | T | 1.23E-03 | | | | | ORF1ab | nonsynonymous_A6407V |
| 29803 | C | A | 1.22E-03 | | | | | three_prime_UTR | |
| 29793 | T | A | 1.21E-03 | C | 2.41E-04 | | | three_prime_UTR | |
| 29543 | G | C | 1.20E-03 | T | 3.16E-04 | | | intergenic | |
| 29449 | G | T | 1.20E-03 | | | | | N | synonymous_V392V |
| 29635 | C | T | 1.20E-03 | | | | | ORF10 | synonymous_Y26Y |
| 28826 | C | T | 1.20E-03 | | | | | N | nonsynonymous_R185C |
| 10870 | G | T | 1.20E-03 | | | | | ORF1ab | synonymous_L3535L |
| 18744 | C | T | 1.20E-03 | | | | | ORF1ab | synonymous_Y6160Y |
| 5467 | C | T | 1.20E-03 | | | | | ORF1ab | synonymous_Y1734Y |

| | | | | | | | | | |
|---|---|---|---|---|---|---|---|---|---|
| 22225 | G | T | 1.20E-03 | A | 1.26E-04 | | | S | synonymous_S221S; synonymous_S221S |
| 14912 | A | G | 1.20E-03 | | | | | ORF1ab | nonsynonymous_N4883S |
| 14925 | C | T | 1.20E-03 | | | | | ORF1ab | synonymous_V4887V |
| 17104 | C | T | 1.20E-03 | | | | | ORF1ab | nonsynonymous_H5614Y |
| 26028 | C | T | 1.20E-03 | | | | | ORF3a | synonymous_Y212Y |
| 28629 | C | T | 1.20E-03 | | | | | N | nonsynonymous_A119V |
| 6723 | C | T | 1.20E-03 | | | | | ORF1ab | nonsynonymous_T2153I |
| 1314 | C | T | 1.20E-03 | | | | | ORF1ab | nonsynonymous_T350I |
| 1457 | C | T | 1.20E-03 | | | | | ORF1ab | nonsynonymous_R398C |
| 1943 | C | T | 1.20E-03 | | | | | ORF1ab | nonsynonymous_R560C |
| 8072 | A | G | 1.20E-03 | | | | | ORF1ab | nonsynonymous_N2603D |
| 8293 | C | T | 1.20E-03 | A | 1.90E-04 | | | ORF1ab | synonymous_T2676T; synonymous_T2676T |
| 10277 | C | T | 1.20E-03 | | | | | ORF1ab | nonsynonymous_L3338F |
| 4523 | G | A | 1.20E-03 | | | | | ORF1ab | nonsynonymous_A1420T |
| 5730 | C | T | 1.20E-03 | | | | | ORF1ab | nonsynonymous_T1822I |
| 9514 | A | G | 1.20E-03 | | | | | ORF1ab | synonymous_L3083L |
| 21255 | G | T | 1.14E-03 | C | 5.69E-04 | A | 6.33E-05 | ORF1ab | synonymous_A6997A; synonymous_A6997A; synonymous_A6997A |
| 13517 | C | T | 1.14E-03 | | | | | ORF1ab | nonsynonymous_T4418I |
| 20275 | G | A | 1.14E-03 | T | 3.16E-04 | | | ORF1ab | nonsynonymous_D6671N; nonsynonymous_D6671Y |
| 9474 | C | T | 1.14E-03 | | | | | ORF1ab | nonsynonymous_A3070V |
| 16616 | C | T | 1.14E-03 | | | | | ORF1ab | nonsynonymous_T5451I |
| 21802 | T | C | 1.14E-03 | | | | | S | synonymous_D80D |
| 3931 | T | C | 1.14E-03 | | | | | ORF1ab | synonymous_V1222V |
| 12790 | A | G | 1.14E-03 | | | | | ORF1ab | synonymous_T4175T |
| 3253 | C | T | 1.14E-03 | | | | | ORF1ab | synonymous_D996D |

| | | | | | | | | | |
|---|---|---|---|---|---|---|---|---|---|
| 8090 | C | T | 1.14E-03 | | | | | ORF1ab | nonsynonymous_L2609F |
| 29773 | G | A | 1.11E-03 | T | 8.53E-04 | | | three_prime_UTR | |
| 29804 | T | A | 1.09E-03 | C | 1.36E-04 | | | three_prime_UTR | |
| 228 | C | T | 1.09E-03 | | | | | five_prime_UTR | |
| 29705 | G | T | 1.08E-03 | A | 6.38E-05 | | | three_prime_UTR | |
| 21624 | G | T | 1.08E-03 | A | 6.32E-05 | A | 6.32E-05 | S | nonsynonymous_R21I; nonsynonymous_R21K; nonsynonymous_R21K |
| 22992 | G | A | 1.08E-03 | T | 6.32E-05 | | | S | nonsynonymous_S477N; nonsynonymous_S477I |
| 29253 | C | T | 1.08E-03 | | | | | N | nonsynonymous_S327L |
| 20005 | G | A | 1.08E-03 | | | | | ORF1ab | nonsynonymous_G6581S |
| 26433 | A | G | 1.07E-03 | | | | | E | synonymous_K63K |
| 21855 | C | T | 1.07E-03 | | | | | S | nonsynonymous_S98F |
| 6363 | C | T | 1.07E-03 | | | | | ORF1ab | nonsynonymous_A2033V |
| 15406 | G | A | 1.07E-03 | | | | | ORF1ab | nonsynonymous_A5048T |
| 21911 | C | T | 1.07E-03 | | | | | S | synonymous_L117L |
| 26873 | C | T | 1.07E-03 | | | | | M | synonymous_N117N |
| 3787 | C | T | 1.07E-03 | | | | | ORF1ab | synonymous_Y1174Y |
| 3045 | C | T | 1.07E-03 | | | | | ORF1ab | nonsynonymous_P927L |
| 10376 | C | T | 1.07E-03 | | | | | ORF1ab | nonsynonymous_P3371S |
| 11224 | C | T | 1.07E-03 | | | | | ORF1ab | synonymous_V3653V |
| 12076 | C | T | 1.07E-03 | | | | | ORF1ab | synonymous_N3937N |
| 29772 | T | A | 1.05E-03 | C | 1.31E-04 | | | three_prime_UTR | |
| 29769 | C | A | 1.05E-03 | T | 5.24E-04 | | | three_prime_UTR | |
| 29701 | G | T | 1.02E-03 | C | 6.38E-05 | | | three_prime_UTR | |
| 29801 | C | A | 1.01E-03 | T | 6.76E-05 | | | three_prime_UTR | |
| 27877 | G | T | 1.01E-03 | | | | | ORF7b | nonsynonymous_C41F |
| 29422 | G | T | 1.01E-03 | A | 2.53E-04 | | | N | synonymous_P383P; synonymous_P383P |

| | | | | | | | | | |
|---|---|---|---|---|---|---|---|---|---|
| 29585 | C | T | 1.01E-03 | A | 6.33E-05 | | | ORF10 | nonsynonymous_P10S; nonsynonymous_P10T |
| 29254 | G | T | 1.01E-03 | A | 6.32E-05 | | | N | synonymous_S327S; synonymous_S327S |
| 26936 | C | T | 1.01E-03 | | | | | M | synonymous_L138L |
| 29303 | C | T | 1.01E-03 | | | | | N | nonsynonymous_P344S |
| 28835 | T | C | 1.01E-03 | | | | | N | nonsynonymous_S188P |
| 10874 | A | G | 1.01E-03 | | | | | ORF1ab | nonsynonymous_N3537D |
| 8767 | T | C | 1.01E-03 | | | | | ORF1ab | synonymous_F2834F |
| 13929 | T | C | 1.01E-03 | | | | | ORF1ab | synonymous_Y4555Y |
| 26340 | C | T | 1.01E-03 | | | | | E | synonymous_A32A |
| 9430 | C | T | 1.01E-03 | A | 1.26E-04 | | | ORF1ab | synonymous_I3055I; synonymous_I3055I |
| 11704 | C | T | 1.01E-03 | | | | | ORF1ab | synonymous_Y3813Y |
| 11798 | T | C | 1.01E-03 | | | | | ORF1ab | synonymous_L3845L |
| 1191 | C | T | 1.01E-03 | | | | | ORF1ab | nonsynonymous_P309L |
| 4755 | C | T | 1.01E-03 | | | | | ORF1ab | nonsynonymous_P1497L |
| 4809 | C | T | 1.01E-03 | | | | | ORF1ab | nonsynonymous_S1515F |
| 6504 | C | T | 1.01E-03 | | | | | ORF1ab | nonsynonymous_A2080V |
| 9962 | C | T | 1.01E-03 | | | | | ORF1ab | nonsynonymous_H3233Y |
| 30 | A | T | 9.86E-04 | C | 5.64E-04 | G | 1.41E-04 | five_prime_UTR | |
| 106 | C | T | 9.67E-04 | | | | | five_prime_UTR | |
| 29751 | G | C | 9.59E-04 | A | 2.56E-04 | A | 2.56E-04 | three_prime_UTR | |
| 19366 | C | T | 9.50E-04 | | | | | ORF1ab | nonsynonymous_P6368S |
| 27525 | A | G | 9.50E-04 | | | | | ORF7a | synonymous_S44S |
| 27679 | C | T | 9.50E-04 | | | | | ORF7a | nonsynonymous_L96F |
| 13438 | T | C | 9.49E-04 | | | | | ORF1ab | synonymous_L4391L |
| 16762 | C | T | 9.49E-04 | | | | | ORF1ab | nonsynonymous_L5500F |
| 28977 | C | T | 9.49E-04 | | | | | N | nonsynonymous_S235F |
| 14580 | G | T | 9.48E-04 | | | | | ORF1ab | nonsynonymous_M4772I |

| | | | | | | | | | |
|---|---|---|---|---|---|---|---|---|---|
| 13947 | A | T | 9.48E-04 | | | | | ORF1ab | synonymous_P4561P |
| 14741 | C | T | 9.48E-04 | | | | | ORF1ab | nonsynonymous_S4826F |
| 17014 | G | T | 9.48E-04 | | | | | ORF1ab | nonsynonymous_D5584Y |
| 23422 | C | T | 9.48E-04 | A | 6.32E-05 | | | S | synonymous_V620V; synonymous_V620V |
| 24038 | G | T | 9.48E-04 | | | | | S | nonsynonymous_V826L |
| 26625 | C | T | 9.48E-04 | | | | | M | synonymous_L35L |
| 26677 | T | C | 9.48E-04 | | | | | M | nonsynonymous_I52T |
| 26720 | G | C | 9.48E-04 | | | | | M | synonymous_V66V |
| 28514 | G | T | 9.48E-04 | | | | | N | nonsynonymous_D81Y |
| 3911 | A | G | 9.48E-04 | | | | | ORF1ab | nonsynonymous_T1216A |
| 6255 | C | T | 9.48E-04 | | | | | ORF1ab | nonsynonymous_A1997V |
| 7877 | A | G | 9.48E-04 | | | | | ORF1ab | nonsynonymous_I2538V |
| 10236 | A | G | 9.48E-04 | | | | | ORF1ab | nonsynonymous_K3324R |
| 2455 | C | T | 9.48E-04 | | | | | ORF1ab | synonymous_L730L |
| 541 | C | T | 9.48E-04 | | | | | ORF1ab | synonymous_L92L |
| 618 | A | G | 9.48E-04 | | | | | ORF1ab | nonsynonymous_Y118C |
| 696 | A | C | 9.48E-04 | | | | | ORF1ab | nonsynonymous_D144A |
| 9924 | C | T | 9.48E-04 | | | | | ORF1ab | nonsynonymous_A3220V |
| 9979 | C | T | 9.48E-04 | | | | | ORF1ab | synonymous_L3238L |
| 29802 | C | A | 9.46E-04 | T | 6.76E-05 | T | 6.76E-05 | three_prime_UTR | |
| 29706 | G | T | 8.93E-04 | | | | | three_prime_UTR | |
| 29567 | A | C | 8.86E-04 | G | 6.33E-05 | | | ORF10 | nonsynonymous_I4L; nonsynonymous_I4V |
| 29374 | G | A | 8.86E-04 | | | | | N | synonymous_E367E |
| 25785 | G | T | 8.85E-04 | C | 6.32E-05 | | | ORF3a | nonsynonymous_W131C; nonsynonymous_W131C |
| 4668 | C | T | 8.85E-04 | | | | | ORF1ab | nonsynonymous_S1468F |
| 21584 | T | G | 8.85E-04 | | | | | S | nonsynonymous_L8V |
| 22984 | G | A | 8.85E-04 | | | | | S | synonymous_Q474Q |

| | | | | | | | | | |
|---|---|---|---|---|---|---|---|---|---|
| 27294 | C | T | 8.85E-04 | | | | | ORF6 | synonymous_Y31Y |
| 18756 | G | T | 8.85E-04 | A | 1.26E-04 | | | ORF1ab | synonymous_P6164P; synonymous_P6164P |
| 24095 | G | T | 8.85E-04 | | | | | S | nonsynonymous_A845S |
| 24863 | C | T | 8.85E-04 | | | | | S | nonsynonymous_H1101Y |
| 27005 | C | T | 8.85E-04 | | | | | M | synonymous_I161I |
| 6616 | A | G | 8.85E-04 | | | | | ORF1ab | synonymous_L2117L |
| 9491 | C | T | 8.85E-04 | | | | | ORF1ab | nonsynonymous_H3076Y |
| 10761 | A | G | 8.85E-04 | | | | | ORF1ab | nonsynonymous_K3499R |
| 10789 | C | T | 8.85E-04 | | | | | ORF1ab | synonymous_D3508D |
| 13216 | T | G | 8.85E-04 | | | | | ORF1ab | nonsynonymous_D4317E |
| 15927 | T | G | 8.85E-04 | | | | | ORF1ab | synonymous_L5221L |
| 16329 | C | T | 8.85E-04 | | | | | ORF1ab | synonymous_Y5355Y |
| 17639 | C | T | 8.85E-04 | | | | | ORF1ab | nonsynonymous_S5792L |
| 21742 | C | T | 8.85E-04 | | | | | S | synonymous_S60S |
| 22550 | C | T | 8.85E-04 | | | | | S | nonsynonymous_P330S |
| 22661 | G | T | 8.85E-04 | | | | | S | nonsynonymous_V367F |
| 23311 | G | T | 8.85E-04 | | | | | S | nonsynonymous_E583D |
| 26730 | G | T | 8.85E-04 | A | 6.95E-04 | | | M | nonsynonymous_V70F; nonsynonymous_V70I |
| 28708 | C | T | 8.85E-04 | | | | | N | synonymous_H145H |
| 9445 | T | C | 8.85E-04 | | | | | ORF1ab | synonymous_L3060L |
| 10369 | C | T | 8.85E-04 | | | | | ORF1ab | synonymous_R3368R |
| 2116 | T | A | 8.85E-04 | | | | | ORF1ab | nonsynonymous_F617L |
| 9534 | C | A | 8.85E-04 | T | 2.53E-04 | | | ORF1ab | nonsynonymous_T3090N; nonsynonymous_T3090I |
| 1170 | C | T | 8.85E-04 | | | | | ORF1ab | nonsynonymous_S302F |
| 11839 | A | G | 8.85E-04 | | | | | ORF1ab | synonymous_V3858V |
| 1218 | C | T | 8.85E-04 | | | | | ORF1ab | nonsynonymous_S318L |
| 3215 | A | G | 8.85E-04 | | | | | ORF1ab | nonsynonymous_S984G |

| | | | | | | | | | |
|---|---|---|---|---|---|---|---|---|---|
| 988 | C | T | 8.85E-04 | | | | | ORF1ab | synonymous_Y241Y |
| 29753 | T | C | 8.31E-04 | A | 1.28E-04 | | | three_prime_UTR | |
| 29686 | C | T | 8.26E-04 | A | 1.27E-04 | A | 1.27E-04 | three_prime_UTR | |
| 27751 | A | T | 8.23E-04 | | | | | ORF7a | nonsynonymous_T120S |
| 21204 | G | T | 8.22E-04 | | | | | ORF1ab | nonsynonymous_K6980N |
| 2676 | C | T | 8.22E-04 | | | | | ORF1ab | nonsynonymous_P804L |
| 2764 | G | T | 8.22E-04 | | | | | ORF1ab | synonymous_V833V |
| 29640 | C | T | 8.22E-04 | | | | | ORF10 | nonsynonymous_A28V |
| 29541 | C | T | 8.22E-04 | | | | | intergenic | |
| 25906 | G | T | 8.22E-04 | | | | | ORF3a | nonsynonymous_G172C |
| 5572 | G | T | 8.22E-04 | A | 2.53E-04 | | | ORF1ab | nonsynonymous_M1769I; nonsynonymous_M1769I |
| 23587 | G | C | 8.22E-04 | T | 2.53E-04 | | | S | nonsynonymous_Q675H; nonsynonymous_Q675H |
| 21123 | G | T | 8.22E-04 | | | | | ORF1ab | synonymous_G6953G |
| 25587 | C | T | 8.22E-04 | | | | | ORF3a | synonymous_L65L |
| 14592 | T | G | 8.22E-04 | | | | | ORF1ab | synonymous_S4776S |
| 18583 | G | T | 8.22E-04 | A | 6.32E-05 | | | ORF1ab | nonsynonymous_V6107L; nonsynonymous_V6107I |
| 17808 | G | T | 8.22E-04 | A | 6.32E-05 | A | 6.32E-05 | ORF1ab | nonsynonymous_K5848N; synonymous_K5848K; synonymous_K5848K |
| 19718 | C | T | 8.22E-04 | | | | | ORF1ab | nonsynonymous_T6485I |
| 23849 | T | C | 8.22E-04 | | | | | S | synonymous_L763L |
| 24099 | C | T | 8.22E-04 | | | | | S | nonsynonymous_A846V |
| 26062 | G | T | 8.22E-04 | | | | | ORF3a | nonsynonymous_G224C |
| 27327 | G | T | 8.22E-04 | A | 6.32E-05 | | | ORF6 | nonsynonymous_K42N; synonymous_K42K |
| 3955 | G | T | 8.22E-04 | | | | | ORF1ab | nonsynonymous_K1230N |
| 6026 | C | T | 8.22E-04 | | | | | ORF1ab | nonsynonymous_P1921S |

| | | | | | | | | | |
|---|---|---|---|---|---|---|---|---|---|
| 6606 | C | T | 8.22E-04 | | | | | ORF1ab | nonsynonymous_S2114F |
| 10450 | C | T | 8.22E-04 | | | | | ORF1ab | synonymous_P3395P |
| 11438 | A | G | 8.22E-04 | | | | | ORF1ab | nonsynonymous_N3725D |
| 3619 | C | T | 8.22E-04 | | | | | ORF1ab | synonymous_V1118V |
| 4084 | C | T | 8.22E-04 | | | | | ORF1ab | synonymous_D1273D |
| 8602 | G | A | 8.22E-04 | | | | | ORF1ab | synonymous_V2779V |
| 48 | C | A | 8.19E-04 | | | | | five_prime_UTR | |
| 29831 | C | A | 7.78E-04 | T | 2.12E-04 | | | three_prime_UTR | |
| 29830 | G | A | 7.78E-04 | | | | | three_prime_UTR | |
| 186 | C | T | 7.66E-04 | A | 6.38E-05 | | | five_prime_UTR | |
| 29736 | G | T | 7.66E-04 | A | 1.28E-04 | A | 1.28E-04 | three_prime_UTR | |
| 27741 | C | T | 7.60E-04 | | | | | ORF7a | synonymous_L116L |
| 29614 | C | T | 7.59E-04 | | | | | ORF10 | synonymous_C19C |
| 25844 | C | T | 7.59E-04 | | | | | ORF3a | nonsynonymous_T151I |
| 25593 | G | C | 7.59E-04 | T | 3.79E-04 | | | ORF3a | nonsynonymous_K67N; nonsynonymous_K67N |
| 26456 | C | T | 7.59E-04 | | | | | E | nonsynonymous_P71L |
| 15738 | C | T | 7.59E-04 | | | | | ORF1ab | synonymous_F5158F |
| 20719 | C | T | 7.59E-04 | | | | | ORF1ab | synonymous_L6819L |
| 25047 | C | T | 7.59E-04 | | | | | S | nonsynonymous_P1162L |
| 28845 | G | T | 7.59E-04 | A | 1.26E-04 | | | N | nonsynonymous_R191L; nonsynonymous_R191H |
| 28849 | C | T | 7.59E-04 | G | 1.26E-04 | | | N | synonymous_N192N; nonsynonymous_N192K |
| 5183 | C | T | 7.59E-04 | | | | | ORF1ab | nonsynonymous_P1640S |
| 16078 | G | A | 7.59E-04 | | | | | ORF1ab | nonsynonymous_V5272I |
| 16376 | C | T | 7.59E-04 | | | | | ORF1ab | nonsynonymous_P5371L |
| 16741 | G | T | 7.59E-04 | | | | | ORF1ab | nonsynonymous_V5493F |
| 18110 | C | T | 7.59E-04 | | | | | ORF1ab | nonsynonymous_P5949L |
| 18568 | C | T | 7.59E-04 | | | | | ORF1ab | nonsynonymous_L6102F |

| | | | | | | | | | |
|---|---|---|---|---|---|---|---|---|---|
| 25585 | C | T | 7.59E-04 | | | | | ORF3a | nonsynonymous_L65F |
| 29013 | C | T | 7.59E-04 | | | | | N | nonsynonymous_T247I |
| 9246 | C | T | 7.59E-04 | | | | | ORF1ab | nonsynonymous_A2994V |
| 14122 | G | T | 7.59E-04 | | | | | ORF1ab | nonsynonymous_G4620C |
| 14940 | A | G | 7.59E-04 | | | | | ORF1ab | synonymous_K4892K |
| 15240 | C | T | 7.59E-04 | | | | | ORF1ab | synonymous_N4992N |
| 16206 | G | T | 7.59E-04 | | | | | ORF1ab | nonsynonymous_E5314D |
| 16457 | C | T | 7.59E-04 | | | | | ORF1ab | nonsynonymous_S5398L |
| 19086 | G | T | 7.59E-04 | | | | | ORF1ab | nonsynonymous_K6274N |
| 20796 | A | G | 7.59E-04 | | | | | ORF1ab | synonymous_K6844K |
| 21711 | C | T | 7.59E-04 | | | | | S | nonsynonymous_S50L |
| 24023 | C | T | 7.59E-04 | A | 6.32E-05 | | | S | synonymous_L821L; nonsynonymous_L821I |
| 26013 | C | T | 7.59E-04 | | | | | ORF3a | synonymous_F207F |
| 28739 | G | T | 7.59E-04 | | | | | N | nonsynonymous_A156S |
| 8785 | G | A | 7.59E-04 | | | | | ORF1ab | synonymous_Q2840Q |
| 11572 | C | T | 7.59E-04 | | | | | ORF1ab | synonymous_F3769F |
| 2523 | C | T | 7.59E-04 | | | | | ORF1ab | nonsynonymous_T753I |
| 5869 | C | T | 7.59E-04 | | | | | ORF1ab | synonymous_Y1868Y |
| 9866 | C | T | 7.59E-04 | | | | | ORF1ab | nonsynonymous_L3201F |
| 1101 | C | T | 7.59E-04 | | | | | ORF1ab | nonsynonymous_S279F |
| 12213 | C | T | 7.59E-04 | | | | | ORF1ab | nonsynonymous_S3983F |
| 6226 | T | C | 7.59E-04 | | | | | ORF1ab | synonymous_H1987H |
| 7164 | C | T | 7.59E-04 | | | | | ORF1ab | nonsynonymous_T2300I |
| 29829 | T | A | 7.57E-04 | C | 6.88E-05 | | | three_prime_UTR | |
| 32 | C | T | 7.47E-04 | A | 1.24E-04 | | | five_prime_UTR | |
| 24 | G | A | 7.39E-04 | A | 7.39E-04 | T | 5.54E-04 | five_prime_UTR | |
| 42 | T | C | 7.34E-04 | G | 1.05E-04 | | | five_prime_UTR | |
| 44 | C | A | 7.25E-04 | T | 1.04E-04 | T | 1.04E-04 | five_prime_UTR | |
| 38 | A | G | 7.09E-04 | C | 1.18E-04 | | | five_prime_UTR | |

| | | | | | | | | | |
|---|---|---|---|---|---|---|---|---|---|
| 153 | T | G | 7.03E-04 | C | 6.39E-05 | | | five_prime_UTR | |
| 19518 | G | T | 6.96E-04 | | | | | ORF1ab | nonsynonymous_L6418F |
| 28087 | C | T | 6.96E-04 | | | | | ORF8 | nonsynonymous_A65V |
| 28000 | C | T | 6.96E-04 | | | | | ORF8 | nonsynonymous_P36L |
| 28099 | C | T | 6.96E-04 | | | | | ORF8 | nonsynonymous_S69L |
| 20419 | G | T | 6.96E-04 | | | | | ORF1ab | nonsynonymous_D6719Y |
| 2626 | G | A | 6.96E-04 | | | | | ORF1ab | synonymous_G787G |
| 2775 | A | G | 6.96E-04 | | | | | ORF1ab | nonsynonymous_K837R |
| 23120 | G | T | 6.96E-04 | | | | | S | nonsynonymous_A520S |
| 29625 | C | T | 6.96E-04 | G | 6.33E-05 | | | ORF10 | nonsynonymous_S23F; nonsynonymous_S23C |
| 8655 | C | T | 6.96E-04 | | | | | ORF1ab | nonsynonymous_S2797F |
| 23086 | C | T | 6.96E-04 | | | | | S | synonymous_Y508Y |
| 25855 | G | T | 6.96E-04 | | | | | ORF3a | nonsynonymous_D155Y |
| 25521 | C | T | 6.96E-04 | | | | | ORF3a | synonymous_F43F |
| 26527 | C | T | 6.96E-04 | | | | | M | nonsynonymous_A2V |
| 28857 | G | A | 6.96E-04 | T | 2.53E-04 | | | N | nonsynonymous_R195K; nonsynonymous_R195I |
| 5230 | G | T | 6.96E-04 | | | | | ORF1ab | nonsynonymous_K1655N |
| 10818 | C | T | 6.96E-04 | | | | | ORF1ab | nonsynonymous_A3518V |
| 22606 | A | T | 6.96E-04 | | | | | S | synonymous_A348A |
| 26152 | G | A | 6.96E-04 | T | 1.26E-04 | | | ORF3a | nonsynonymous_G254R; stopgain_G254X |
| 26211 | G | T | 6.96E-04 | | | | | ORF3a | synonymous_V273V |
| 26549 | C | T | 6.96E-04 | | | | | M | synonymous_T9T |
| 13694 | C | T | 6.95E-04 | | | | | ORF1ab | nonsynonymous_T4477I |
| 14195 | C | A | 6.95E-04 | | | | | ORF1ab | nonsynonymous_T4644N |
| 18488 | T | C | 6.95E-04 | | | | | ORF1ab | nonsynonymous_I6075T |
| 19017 | C | T | 6.95E-04 | | | | | ORF1ab | synonymous_F6251F |
| 19602 | C | T | 6.95E-04 | | | | | ORF1ab | synonymous_N6446N |

| | | | | | | | | | |
|---|---|---|---|---|---|---|---|---|---|
| 20991 | G | T | 6.95E-04 | A | 2.53E-04 | C | 6.32E-05 | ORF1ab | nonsynonymous_L6909F; synonymous_L6909L; nonsynonymous_L6909F |
| 23635 | C | T | 6.95E-04 | G | 6.32E-05 | | | S | synonymous_S691S; synonymous_S691S |
| 25485 | T | C | 6.95E-04 | | | | | ORF3a | synonymous_A31A |
| 26111 | C | T | 6.95E-04 | | | | | ORF3a | nonsynonymous_P240L |
| 26895 | C | T | 6.95E-04 | | | | | M | nonsynonymous_H125Y |
| 6446 | G | A | 6.95E-04 | T | 1.90E-04 | | | ORF1ab | nonsynonymous_V2061I; nonsynonymous_V2061F |
| 683 | C | T | 6.95E-04 | | | | | ORF1ab | synonymous_L140L |
| 9724 | C | T | 6.95E-04 | | | | | ORF1ab | synonymous_F3153F |
| 12400 | C | T | 6.95E-04 | | | | | ORF1ab | synonymous_L4045L |
| 1904 | A | G | 6.95E-04 | | | | | ORF1ab | nonsynonymous_I547V |
| 7869 | C | T | 6.95E-04 | | | | | ORF1ab | nonsynonymous_S2535L |
| 7945 | C | T | 6.95E-04 | | | | | ORF1ab | synonymous_Y2560Y |
| 829 | C | T | 6.95E-04 | | | | | ORF1ab | synonymous_N188N |
| 9561 | C | T | 6.95E-04 | | | | | ORF1ab | nonsynonymous_S3099L |
| 10561 | T | C | 6.95E-04 | | | | | ORF1ab | synonymous_T3432T |
| 11379 | C | T | 6.95E-04 | | | | | ORF1ab | nonsynonymous_A3705V |
| 1321 | A | C | 6.95E-04 | | | | | ORF1ab | nonsynonymous_E352D |
| 4444 | G | T | 6.95E-04 | | | | | ORF1ab | synonymous_V1393V |
| 8612 | G | T | 6.95E-04 | | | | | ORF1ab | nonsynonymous_V2783F |
| 9172 | T | C | 6.95E-04 | | | | | ORF1ab | synonymous_L2969L |
| 66 | C | T | 6.52E-04 | | | | | five_prime_UTR | |
| 150 | T | C | 6.40E-04 | | | | | five_prime_UTR | |
| 28093 | C | T | 6.33E-04 | | | | | ORF8 | nonsynonymous_S67F |
| 25798 | A | G | 6.33E-04 | | | | | ORF3a | nonsynonymous_K136E |
| 27715 | G | T | 6.33E-04 | | | | | ORF7a | nonsynonymous_V108L |

| | | | | | | | | | |
|---|---|---|---|---|---|---|---|---|---|
| 21304 | C | A | 6.33E-04 | T | 6.33E-05 | | | ORF1ab | nonsynonymous_R7014S; nonsynonymous_R7014C |
| 29474 | G | T | 6.33E-04 | C | 6.33E-05 | | | N | nonsynonymous_D401Y; nonsynonymous_D401H |
| 29403 | A | G | 6.33E-04 | | | | | N | nonsynonymous_D377G |
| 29627 | C | T | 6.33E-04 | | | | | ORF10 | nonsynonymous_R24C |
| 20178 | C | T | 6.33E-04 | | | | | ORF1ab | synonymous_V6638V |
| 29077 | C | T | 6.33E-04 | | | | | N | synonymous_Y268Y |
| 25687 | G | T | 6.32E-04 | A | 6.32E-05 | | | ORF3a | nonsynonymous_A99S; nonsynonymous_A99T |
| 20031 | C | A | 6.32E-04 | T | 6.32E-05 | | | ORF1ab | synonymous_A6589A; synonymous_A6589A |
| 24919 | C | T | 6.32E-04 | | | | | S | synonymous_N1119N |
| 26466 | G | T | 6.32E-04 | A | 1.26E-04 | | | E | synonymous_L74L; synonymous_L74L |
| 26467 | G | C | 6.32E-04 | | | | | E | nonsynonymous_V75L |
| 29171 | C | T | 6.32E-04 | | | | | N | nonsynonymous_H300Y |
| 6825 | A | C | 6.32E-04 | | | | | ORF1ab | nonsynonymous_N2187T |
| 20887 | G | A | 6.32E-04 | | | | | ORF1ab | nonsynonymous_G6875R |
| 24794 | G | T | 6.32E-04 | | | | | S | nonsynonymous_A1078S |
| 28690 | G | T | 6.32E-04 | | | | | N | nonsynonymous_L139F |
| 28985 | G | T | 6.32E-04 | A | 6.32E-05 | | | N | nonsynonymous_G238C; nonsynonymous_G238S |
| 635 | C | T | 6.32E-04 | | | | | ORF1ab | nonsynonymous_R124C |
| 16362 | C | T | 6.32E-04 | | | | | ORF1ab | synonymous_V5366V |
| 18029 | C | T | 6.32E-04 | | | | | ORF1ab | nonsynonymous_A5922V |
| 19862 | C | T | 6.32E-04 | | | | | ORF1ab | nonsynonymous_A6533V |
| 27059 | C | T | 6.32E-04 | | | | | M | synonymous_Y179Y |
| 16293 | C | T | 6.32E-04 | | | | | ORF1ab | synonymous_C5343C |
| 16852 | G | T | 6.32E-04 | | | | | ORF1ab | nonsynonymous_G5530C |

| | | | | | | | | | |
|---|---|---|---|---|---|---|---|---|---|
| 17531 | T | C | 6.32E-04 | | | | | ORF1ab | nonsynonymous_I5756T |
| 18348 | C | T | 6.32E-04 | | | | | ORF1ab | synonymous_T6028T |
| 21846 | C | T | 6.32E-04 | | | | | S | nonsynonymous_T95I |
| 21995 | T | C | 6.32E-04 | | | | | S | nonsynonymous_Y145H |
| 22729 | T | C | 6.32E-04 | | | | | S | synonymous_D389D |
| 26529 | G | T | 6.32E-04 | | | | | M | nonsynonymous_D3Y |
| 6081 | A | G | 6.32E-04 | | | | | ORF1ab | nonsynonymous_D1939G |
| 14118 | G | T | 6.32E-04 | A | 1.90E-04 | | | ORF1ab | synonymous_T4618T; synonymous_T4618T |
| 17326 | C | T | 6.32E-04 | | | | | ORF1ab | nonsynonymous_P5688S |
| 17678 | C | T | 6.32E-04 | | | | | ORF1ab | nonsynonymous_T5805M |
| 17766 | C | T | 6.32E-04 | | | | | ORF1ab | synonymous_V5834V |
| 17799 | A | G | 6.32E-04 | | | | | ORF1ab | synonymous_V5845V |
| 18508 | C | T | 6.32E-04 | | | | | ORF1ab | nonsynonymous_L6082F |
| 1887 | C | T | 6.32E-04 | | | | | ORF1ab | nonsynonymous_A541V |
| 18928 | C | T | 6.32E-04 | | | | | ORF1ab | nonsynonymous_P6222S |
| 27821 | A | G | 6.32E-04 | | | | | ORF7b | synonymous_L22L |
| 6402 | C | T | 6.32E-04 | | | | | ORF1ab | nonsynonymous_P2046L |
| 7600 | C | T | 6.32E-04 | | | | | ORF1ab | synonymous_C2445C |
| 3140 | C | T | 6.32E-04 | | | | | ORF1ab | nonsynonymous_P959S |
| 4069 | C | T | 6.32E-04 | | | | | ORF1ab | synonymous_A1268A |
| 10015 | C | T | 6.32E-04 | | | | | ORF1ab | synonymous_Y3250Y |
| 11638 | T | C | 6.32E-04 | | | | | ORF1ab | synonymous_T3791T |
| 1163 | A | T | 6.32E-04 | | | | | ORF1ab | nonsynonymous_I300F |
| 11674 | C | T | 6.32E-04 | | | | | ORF1ab | synonymous_Y3803Y |
| 12550 | G | A | 6.32E-04 | | | | | ORF1ab | synonymous_L4095L |
| 13019 | C | T | 6.32E-04 | | | | | ORF1ab | synonymous_L4252L |
| 1319 | G | A | 6.32E-04 | | | | | ORF1ab | nonsynonymous_E352K |
| 1918 | T | C | 6.32E-04 | | | | | ORF1ab | synonymous_T551T |
| 431 | G | A | 6.32E-04 | | | | | ORF1ab | nonsynonymous_V56I |

| | | | | | | | | | |
|---|---|---|---|---|---|---|---|---|---|
| 4949 | T | C | 6.32E-04 | | | | | ORF1ab | synonymous_L1562L |
| 5817 | C | T | 6.32E-04 | | | | | ORF1ab | nonsynonymous_A1851V |
| 7393 | G | T | 6.32E-04 | A | 6.32E-05 | | | ORF1ab | synonymous_P2376P; synonymous_P2376P |
| 33 | C | T | 6.18E-04 | A | 3.71E-04 | G | 1.24E-04 | five_prime_UTR | |
| 29764 | G | A | 5.76E-04 | T | 1.92E-04 | | | three_prime_UTR | |
| 199 | G | T | 5.74E-04 | A | 6.38E-05 | A | 6.38E-05 | five_prime_UTR | |
| 29733 | C | T | 5.74E-04 | A | 6.38E-05 | | | three_prime_UTR | |
| 29708 | C | T | 5.74E-04 | | | | | three_prime_UTR | |
| 27630 | C | T | 5.70E-04 | | | | | ORF7a | synonymous_A79A |
| 28045 | C | T | 5.70E-04 | A | 6.33E-05 | A | 6.33E-05 | ORF8 | nonsynonymous_A51V; nonsynonymous_A51D; nonsynonymous_A51D |
| 3145 | G | T | 5.69E-04 | | | | | ORF1ab | nonsynonymous_L960F |
| 13584 | C | T | 5.69E-04 | | | | | ORF1ab | synonymous_F4440F |
| 2808 | G | A | 5.69E-04 | | | | | ORF1ab | nonsynonymous_R848K |
| 29358 | C | T | 5.69E-04 | A | 6.33E-05 | | | N | nonsynonymous_T362I; nonsynonymous_T362K |
| 29410 | T | C | 5.69E-04 | | | | | N | synonymous_T379T |
| 25782 | C | T | 5.69E-04 | | | | | ORF3a | synonymous_C130C |
| 21627 | C | T | 5.69E-04 | | | | | S | nonsynonymous_T22I |
| 22995 | C | T | 5.69E-04 | | | | | S | nonsynonymous_T478I |
| 21518 | G | T | 5.69E-04 | | | | | ORF1ab | nonsynonymous_R7085I |
| 29284 | C | T | 5.69E-04 | | | | | N | synonymous_I337I |
| 16658 | C | T | 5.69E-04 | | | | | ORF1ab | nonsynonymous_T5465I |
| 21691 | C | T | 5.69E-04 | | | | | S | synonymous_F43F |
| 22592 | G | T | 5.69E-04 | | | | | S | nonsynonymous_A344S |
| 24378 | C | T | 5.69E-04 | | | | | S | nonsynonymous_S939F |
| 28924 | T | C | 5.69E-04 | | | | | N | synonymous_A217A |
| 15860 | A | G | 5.69E-04 | | | | | ORF1ab | nonsynonymous_K5199R |

| | | | | | | | | | |
|---|---|---|---|---|---|---|---|---|---|
| 16017 | C | T | 5.69E-04 | | | | | ORF1ab | synonymous_F5251F |
| 16750 | C | T | 5.69E-04 | | | | | ORF1ab | nonsynonymous_P5496S |
| 17884 | A | G | 5.69E-04 | | | | | ORF1ab | nonsynonymous_T5874A |
| 19645 | G | T | 5.69E-04 | | | | | ORF1ab | nonsynonymous_V6461F |
| 20740 | C | A | 5.69E-04 | | | | | ORF1ab | nonsynonymous_Q6826K |
| 22151 | A | G | 5.69E-04 | | | | | S | nonsynonymous_I197V |
| 23934 | C | T | 5.69E-04 | | | | | S | nonsynonymous_T791I |
| 25494 | G | T | 5.69E-04 | A | 6.32E-05 | | | ORF3a | synonymous_T34T;<br>synonymous_T34T |
| 10626 | C | T | 5.69E-04 | | | | | ORF1ab | nonsynonymous_A3454V |
| 13243 | T | C | 5.69E-04 | | | | | ORF1ab | synonymous_C4326C |
| 13665 | C | T | 5.69E-04 | A | 6.32E-05 | | | ORF1ab | synonymous_H4467H;<br>nonsynonymous_H4467Q |
| 1469 | C | T | 5.69E-04 | | | | | ORF1ab | nonsynonymous_R402C |
| 15426 | A | G | 5.69E-04 | | | | | ORF1ab | synonymous_V5054V |
| 15654 | C | T | 5.69E-04 | | | | | ORF1ab | synonymous_D5130D |
| 16468 | C | T | 5.69E-04 | | | | | ORF1ab | nonsynonymous_P5402S |
| 16813 | G | A | 5.69E-04 | | | | | ORF1ab | nonsynonymous_V5517I |
| 17676 | C | T | 5.69E-04 | | | | | ORF1ab | synonymous_I5804I |
| 17934 | C | T | 5.69E-04 | | | | | ORF1ab | synonymous_T5890T |
| 18264 | C | T | 5.69E-04 | | | | | ORF1ab | synonymous_T6000T |
| 18570 | C | T | 5.69E-04 | | | | | ORF1ab | synonymous_L6102L |
| 19825 | G | T | 5.69E-04 | | | | | ORF1ab | nonsynonymous_V6521L |
| 23191 | C | T | 5.69E-04 | | | | | S | synonymous_F543F |
| 24739 | T | C | 5.69E-04 | | | | | S | synonymous_G1059G |
| 26256 | C | T | 5.69E-04 | | | | | E | synonymous_F4F |
| 28638 | C | T | 5.69E-04 | | | | | N | nonsynonymous_P122L |
| 28727 | G | T | 5.69E-04 | | | | | N | nonsynonymous_A152S |
| 28737 | C | T | 5.69E-04 | | | | | N | nonsynonymous_A155V |
| 3903 | C | T | 5.69E-04 | | | | | ORF1ab | nonsynonymous_P1213L |

| | | | | | | | | | |
|---|---|---|---|---|---|---|---|---|---|
| 5812 | C | T | 5.69E-04 | | | | | ORF1ab | synonymous_D1849D |
| 7279 | C | T | 5.69E-04 | A | 6.32E-05 | | | ORF1ab | synonymous_F2338F; nonsynonymous_F2338L |
| 1514 | C | T | 5.69E-04 | A | 6.32E-05 | | | ORF1ab | nonsynonymous_H417Y; nonsynonymous_H417N |
| 4185 | G | A | 5.69E-04 | T | 6.32E-05 | | | ORF1ab | nonsynonymous_G1307D; nonsynonymous_G1307V |
| 4186 | T | C | 5.69E-04 | | | | | ORF1ab | synonymous_G1307G |
| 4201 | G | T | 5.69E-04 | | | | | ORF1ab | nonsynonymous_M1312I |
| 583 | C | T | 5.69E-04 | | | | | ORF1ab | synonymous_V106V |
| 6636 | C | T | 5.69E-04 | | | | | ORF1ab | nonsynonymous_T2124I |
| 8025 | C | T | 5.69E-04 | | | | | ORF1ab | nonsynonymous_A2587V |
| 10262 | G | A | 5.69E-04 | | | | | ORF1ab | nonsynonymous_A3333T |
| 11537 | A | G | 5.69E-04 | | | | | ORF1ab | nonsynonymous_I3758V |
| 11962 | C | T | 5.69E-04 | | | | | ORF1ab | synonymous_L3899L |
| 1238 | C | T | 5.69E-04 | | | | | ORF1ab | nonsynonymous_H325Y |
| 1570 | T | C | 5.69E-04 | | | | | ORF1ab | synonymous_H435H |
| 2536 | C | T | 5.69E-04 | | | | | ORF1ab | synonymous_V757V |
| 3268 | T | C | 5.69E-04 | | | | | ORF1ab | synonymous_T1001T |
| 3543 | C | T | 5.69E-04 | | | | | ORF1ab | nonsynonymous_T1093I |
| 4551 | C | T | 5.69E-04 | | | | | ORF1ab | nonsynonymous_T1429I |
| 4912 | A | C | 5.69E-04 | | | | | ORF1ab | nonsynonymous_E1549D |
| 4975 | G | T | 5.69E-04 | | | | | ORF1ab | synonymous_V1570V |
| 5826 | C | T | 5.69E-04 | A | 6.32E-05 | | | ORF1ab | nonsynonymous_T1854I; nonsynonymous_T1854K |
| 8367 | C | T | 5.69E-04 | | | | | ORF1ab | nonsynonymous_A2701V |
| 913 | C | T | 5.69E-04 | | | | | ORF1ab | synonymous_S216S |
| 29832 | T | A | 5.67E-04 | G | 7.09E-05 | C | 7.09E-05 | three_prime_UTR | |
| 39 | A | G | 5.44E-04 | T | 3.27E-04 | C | 1.09E-04 | five_prime_UTR | |
| 40 | C | A | 5.31E-04 | T | 4.25E-04 | | | five_prime_UTR | |

| | | | | | | | | | |
|---|---|---|---|---|---|---|---|---|---|
| 41 | T | C | 5.25E-04 | A | 1.05E-04 | | | five_prime_UTR | |
| 29739 | C | T | 5.11E-04 | A | 6.38E-05 | | | three_prime_UTR | |
| 218 | C | T | 5.11E-04 | | | | | five_prime_UTR | |
| 207 | C | T | 5.10E-04 | | | | | five_prime_UTR | |
| 28001 | G | T | 5.06E-04 | | | | | ORF8 | synonymous_P36P |
| 19983 | C | T | 5.06E-04 | | | | | ORF1ab | synonymous_V6573V |
| 27945 | C | T | 5.06E-04 | | | | | ORF8 | stopgain_Q18X |
| 27874 | C | T | 5.06E-04 | | | | | ORF7b | nonsynonymous_T40I |
| 20402 | C | T | 5.06E-04 | | | | | ORF1ab | nonsynonymous_S6713L |
| 2840 | G | A | 5.06E-04 | | | | | ORF1ab | nonsynonymous_A859T |
| 20262 | A | G | 5.06E-04 | | | | | ORF1ab | synonymous_L6666L |
| 25714 | C | T | 5.06E-04 | | | | | ORF3a | nonsynonymous_L108F |
| 17304 | C | T | 5.06E-04 | | | | | ORF1ab | synonymous_V5680V |
| 23127 | C | T | 5.06E-04 | | | | | S | nonsynonymous_A522V |
| 25655 | T | C | 5.06E-04 | | | | | ORF3a | nonsynonymous_V88A |
| 25831 | C | T | 5.06E-04 | | | | | ORF3a | nonsynonymous_L147F |
| 21146 | T | A | 5.06E-04 | | | | | ORF1ab | nonsynonymous_L6961H |
| 22988 | G | A | 5.06E-04 | | | | | S | nonsynonymous_G476S |
| 27476 | C | T | 5.06E-04 | | | | | ORF7a | nonsynonymous_T28I |
| 25603 | C | T | 5.06E-04 | | | | | ORF3a | synonymous_L71L |
| 25624 | C | T | 5.06E-04 | | | | | ORF3a | nonsynonymous_H78Y |
| 29167 | C | T | 5.06E-04 | | | | | N | synonymous_Y298Y |
| 29203 | C | T | 5.06E-04 | | | | | N | synonymous_S310S |
| 29291 | G | A | 5.06E-04 | T | 1.26E-04 | | | N | nonsynonymous_D340N; nonsynonymous_D340Y |
| 29296 | C | T | 5.06E-04 | A | 6.32E-05 | | | N | synonymous_D341D; nonsynonymous_D341E |
| 17502 | C | T | 5.06E-04 | | | | | ORF1ab | synonymous_F5746F |
| 20553 | A | G | 5.06E-04 | | | | | ORF1ab | synonymous_L6763L |
| 2091 | C | T | 5.06E-04 | | | | | ORF1ab | nonsynonymous_T609I |

| | | | | | | | | | |
|---|---|---|---|---|---|---|---|---|---|
| 22103 | G | C | 5.06E-04 | | | | | S | nonsynonymous_G181R |
| 22277 | C | A | 5.06E-04 | | | | | S | nonsynonymous_Q239K |
| 22879 | C | A | 5.06E-04 | | | | | S | nonsynonymous_N439K |
| 23525 | C | T | 5.06E-04 | | | | | S | nonsynonymous_H655Y |
| 25088 | G | T | 5.06E-04 | | | | | S | nonsynonymous_V1176F |
| 25160 | C | T | 5.06E-04 | | | | | S | nonsynonymous_L1200F |
| 25169 | C | T | 5.06E-04 | A | 6.32E-05 | | | S | nonsynonymous_L1203F; nonsynonymous_L1203I |
| 5084 | A | G | 5.06E-04 | | | | | ORF1ab | nonsynonymous_I1607V |
| 14585 | C | T | 5.06E-04 | | | | | ORF1ab | nonsynonymous_A4774V |
| 15237 | C | T | 5.06E-04 | | | | | ORF1ab | synonymous_H4991H |
| 17474 | C | T | 5.06E-04 | | | | | ORF1ab | nonsynonymous_T5737I |
| 17999 | C | T | 5.06E-04 | | | | | ORF1ab | nonsynonymous_T5912I |
| 18656 | C | T | 5.06E-04 | A | 6.32E-05 | | | ORF1ab | nonsynonymous_T6131I; nonsynonymous_T6131N |
| 18723 | T | C | 5.06E-04 | G | 6.32E-05 | | | ORF1ab | synonymous_H6153H; nonsynonymous_H6153Q |
| 18828 | C | T | 5.06E-04 | | | | | ORF1ab | synonymous_V6188V |
| 19884 | C | T | 5.06E-04 | | | | | ORF1ab | synonymous_Y6540Y |
| 21792 | A | T | 5.06E-04 | | | | | S | nonsynonymous_K77M |
| 22309 | G | A | 5.06E-04 | T | 6.32E-05 | | | S | synonymous_L249L; nonsynonymous_L249F |
| 23638 | C | T | 5.06E-04 | | | | | S | synonymous_I692I |
| 24130 | C | T | 5.06E-04 | | | | | S | synonymous_N856N |
| 24133 | C | T | 5.06E-04 | | | | | S | synonymous_G857G |
| 24621 | C | T | 5.06E-04 | | | | | S | nonsynonymous_A1020V |
| 25505 | A | C | 5.06E-04 | T | 1.26E-04 | | | ORF3a | nonsynonymous_Q38P; nonsynonymous_Q38L |
| 25575 | A | C | 5.06E-04 | | | | | ORF3a | nonsynonymous_K61N |
| 26501 | G | T | 5.06E-04 | | | | | intergenic | |

| | | | | | | | | | |
|---|---|---|---|---|---|---|---|---|---|
| 26864 | A | G | 5.06E-04 | | | | | M | synonymous_P114P |
| 28692 | A | C | 5.06E-04 | | | | | N | nonsynonymous_N140T |
| 9344 | C | T | 5.06E-04 | | | | | ORF1ab | nonsynonymous_L3027F |
| 1368 | T | C | 5.06E-04 | | | | | ORF1ab | nonsynonymous_I368T |
| 14079 | T | C | 5.06E-04 | | | | | ORF1ab | synonymous_N4605N |
| 15351 | T | G | 5.06E-04 | | | | | ORF1ab | synonymous_V5029V |
| 16467 | A | G | 5.06E-04 | | | | | ORF1ab | synonymous_P5401P |
| 17944 | G | T | 5.06E-04 | | | | | ORF1ab | nonsynonymous_V5894L |
| 18457 | C | T | 5.06E-04 | | | | | ORF1ab | nonsynonymous_P6065S |
| 18904 | C | T | 5.06E-04 | | | | | ORF1ab | nonsynonymous_R6214C |
| 19029 | C | T | 5.06E-04 | | | | | ORF1ab | synonymous_H6255H |
| 19872 | G | T | 5.06E-04 | | | | | ORF1ab | synonymous_V6536V |
| 20762 | C | T | 5.06E-04 | | | | | ORF1ab | nonsynonymous_T6833I |
| 23533 | C | T | 5.06E-04 | | | | | S | synonymous_N657N |
| 23679 | C | T | 5.06E-04 | | | | | S | nonsynonymous_A706V |
| 25440 | G | A | 5.06E-04 | T | 1.90E-04 | C | 6.32E-05 | ORF3a | synonymous_K16K; nonsynonymous_K16N; nonsynonymous_K16N |
| 26029 | C | A | 5.06E-04 | | | | | ORF3a | nonsynonymous_Q213K |
| 26045 | A | G | 5.06E-04 | | | | | ORF3a | nonsynonymous_Q218R |
| 26124 | C | T | 5.06E-04 | | | | | ORF3a | synonymous_V244V |
| 26615 | G | T | 5.06E-04 | | | | | M | nonsynonymous_W31C |
| 26894 | C | T | 5.06E-04 | | | | | M | synonymous_L124L |
| 6861 | C | T | 5.06E-04 | | | | | ORF1ab | nonsynonymous_A2199V |
| 7292 | T | C | 5.06E-04 | | | | | ORF1ab | synonymous_L2343L |
| 7815 | C | T | 5.06E-04 | | | | | ORF1ab | nonsynonymous_S2517F |
| 10523 | G | C | 5.06E-04 | A | 6.32E-05 | | | ORF1ab | nonsynonymous_V3420L; nonsynonymous_V3420I |
| 11430 | A | G | 5.06E-04 | | | | | ORF1ab | nonsynonymous_Y3722C |
| 1387 | C | T | 5.06E-04 | | | | | ORF1ab | synonymous_H374H |

| | | | | | | | | | |
|---|---|---|---|---|---|---|---|---|---|
| 1625 | C | T | 5.06E-04 | | | | | ORF1ab | nonsynonymous_L454F |
| 2003 | C | T | 5.06E-04 | | | | | ORF1ab | nonsynonymous_L580F |
| 2883 | G | A | 5.06E-04 | | | | | ORF1ab | nonsynonymous_C873Y |
| 2937 | C | T | 5.06E-04 | | | | | ORF1ab | nonsynonymous_T891I |
| 2987 | T | C | 5.06E-04 | | | | | ORF1ab | nonsynonymous_F908L |
| 3393 | C | T | 5.06E-04 | | | | | ORF1ab | nonsynonymous_A1043V |
| 3768 | C | T | 5.06E-04 | | | | | ORF1ab | nonsynonymous_T1168I |
| 3784 | C | T | 5.06E-04 | | | | | ORF1ab | synonymous_V1173V |
| 4050 | A | C | 5.06E-04 | | | | | ORF1ab | nonsynonymous_N1262T |
| 5986 | C | T | 5.06E-04 | | | | | ORF1ab | synonymous_F1907F |
| 761 | A | G | 5.06E-04 | | | | | ORF1ab | nonsynonymous_S166G |
| 8290 | C | T | 5.06E-04 | | | | | ORF1ab | synonymous_L2675L |
| 1148 | G | T | 5.06E-04 | A | 6.32E-05 | | | ORF1ab | nonsynonymous_G295C; nonsynonymous_G295S |
| 1190 | C | T | 5.06E-04 | | | | | ORF1ab | nonsynonymous_P309S |
| 12079 | G | A | 5.06E-04 | C | 6.32E-05 | | | ORF1ab | synonymous_R3938R; nonsynonymous_R3938S |
| 12514 | G | T | 5.06E-04 | C | 6.32E-05 | | | ORF1ab | synonymous_T4083T; synonymous_T4083T |
| 12534 | C | T | 5.06E-04 | | | | | ORF1ab | nonsynonymous_T4090I |
| 12616 | C | T | 5.06E-04 | | | | | ORF1ab | synonymous_D4117D |
| 13072 | C | T | 5.06E-04 | | | | | ORF1ab | synonymous_F4269F |
| 1587 | A | G | 5.06E-04 | | | | | ORF1ab | nonsynonymous_E441G |
| 2266 | T | C | 5.06E-04 | | | | | ORF1ab | synonymous_C667C |
| 2485 | C | T | 5.06E-04 | | | | | ORF1ab | synonymous_I740I |
| 4140 | A | G | 5.06E-04 | | | | | ORF1ab | nonsynonymous_Q1292R |
| 4236 | A | G | 5.06E-04 | | | | | ORF1ab | nonsynonymous_N1324S |
| 7564 | C | T | 5.06E-04 | | | | | ORF1ab | synonymous_S2433S |
| 8092 | C | T | 5.06E-04 | | | | | ORF1ab | synonymous_L2609L |
| 8102 | G | T | 5.06E-04 | | | | | ORF1ab | nonsynonymous_V2613F |

| | | | | | | | | | |
|---|---|---|---|---|---|---|---|---|---|
| 832 | C | T | 5.06E-04 | | | | | ORF1ab | synonymous_N189N |
| 29824 | A | G | 4.80E-04 | | | | | three_prime_UTR | |
| 84 | C | T | 4.54E-04 | A | 6.48E-05 | A | 6.48E-05 | five_prime_UTR | |
| 160 | G | T | 4.47E-04 | A | 3.20E-04 | C | 6.39E-05 | five_prime_UTR | |
| 210 | G | T | 4.47E-04 | A | 6.38E-05 | C | 6.38E-05 | five_prime_UTR | |
| 29732 | C | T | 4.46E-04 | A | 6.38E-05 | A | 6.38E-05 | three_prime_UTR | |
| 527 | C | T | 4.43E-04 | | | | | ORF1ab | synonymous_L88L |
| 19542 | G | T | 4.43E-04 | | | | | ORF1ab | nonsynonymous_M6426I |
| 27915 | G | T | 4.43E-04 | A | 1.90E-04 | | | ORF8 | stopgain_G8X; nonsynonymous_G8R |
| 22335 | G | T | 4.43E-04 | | | | | S | nonsynonymous_W258L |
| 20457 | C | T | 4.43E-04 | | | | | ORF1ab | synonymous_F6731F |
| 29392 | G | T | 4.43E-04 | | | | | N | nonsynonymous_K373N |
| 29546 | C | T | 4.43E-04 | A | 6.33E-05 | | | intergenic | |
| 21242 | C | T | 4.43E-04 | | | | | ORF1ab | nonsynonymous_T6993I |
| 21306 | C | T | 4.43E-04 | | | | | ORF1ab | synonymous_R7014R |
| 20398 | G | A | 4.43E-04 | | | | | ORF1ab | nonsynonymous_E6712K |
| 29402 | G | T | 4.43E-04 | C | 6.33E-05 | | | N | nonsynonymous_D377Y; nonsynonymous_D377H |
| 20355 | A | G | 4.43E-04 | | | | | ORF1ab | synonymous_L6697L |
| 29598 | A | G | 4.43E-04 | | | | | ORF10 | nonsynonymous_Y14C |
| 25731 | C | T | 4.43E-04 | | | | | ORF3a | synonymous_Y113Y |
| 25791 | C | T | 4.43E-04 | | | | | ORF3a | synonymous_C133C |
| 13458 | C | T | 4.43E-04 | | | | | ORF1ab | nonsynonymous_S4398L |
| 21622 | C | T | 4.43E-04 | A | 1.26E-04 | | | S | synonymous_T20T; synonymous_T20T |
| 25658 | C | T | 4.43E-04 | | | | | ORF3a | nonsynonymous_T89I |
| 25775 | G | T | 4.43E-04 | | | | | ORF3a | nonsynonymous_W128L |
| 29140 | G | T | 4.43E-04 | C | 6.32E-05 | | | N | nonsynonymous_Q289H; nonsynonymous_Q289H |

| | | | | | | | | | |
|---|---|---|---|---|---|---|---|---|---|
| 21494 | G | A | 4.43E-04 | | | | | ORF1ab | nonsynonymous_R7077K |
| 21772 | C | T | 4.43E-04 | | | | | S | synonymous_V70V |
| 25879 | G | T | 4.43E-04 | | | | | ORF3a | nonsynonymous_V163L |
| 15857 | C | T | 4.43E-04 | | | | | ORF1ab | nonsynonymous_T5198I |
| 17964 | G | T | 4.43E-04 | A | 3.16E-04 | | | ORF1ab | nonsynonymous_M5900I; nonsynonymous_M5900I |
| 18115 | C | T | 4.43E-04 | | | | | ORF1ab | nonsynonymous_H5951Y |
| 22323 | C | T | 4.43E-04 | | | | | S | nonsynonymous_S254F |
| 25207 | C | T | 4.43E-04 | | | | | S | synonymous_Y1215Y |
| 25606 | G | T | 4.43E-04 | | | | | ORF3a | nonsynonymous_A72S |
| 25644 | G | T | 4.43E-04 | | | | | ORF3a | synonymous_L84L |
| 27434 | C | T | 4.43E-04 | | | | | ORF7a | nonsynonymous_T14I |
| 29272 | C | T | 4.43E-04 | | | | | N | synonymous_Y333Y |
| 5305 | G | A | 4.43E-04 | | | | | ORF1ab | synonymous_L1680L |
| 5573 | T | C | 4.43E-04 | | | | | ORF1ab | nonsynonymous_Y1770H |
| 16575 | C | T | 4.43E-04 | | | | | ORF1ab | synonymous_D5437D |
| 18647 | C | T | 4.43E-04 | | | | | ORF1ab | nonsynonymous_P6128L |
| 2094 | C | T | 4.43E-04 | | | | | ORF1ab | nonsynonymous_S610L |
| 23758 | C | T | 4.43E-04 | | | | | S | synonymous_T732T |
| 25135 | G | T | 4.43E-04 | | | | | S | nonsynonymous_K1191N |
| 25218 | G | T | 4.43E-04 | | | | | S | nonsynonymous_G1219V |
| 25222 | T | C | 4.43E-04 | G | 6.32E-05 | | | S | synonymous_F1220F; nonsynonymous_F1220L |
| 27147 | G | C | 4.43E-04 | T | 2.53E-04 | | | M | nonsynonymous_D209H; nonsynonymous_D209Y |
| 27430 | G | A | 4.43E-04 | | | | | ORF7a | nonsynonymous_A13T |
| 27493 | C | T | 4.43E-04 | | | | | ORF7a | nonsynonymous_P34S |
| 28975 | G | T | 4.43E-04 | A | 6.32E-05 | | | N | nonsynonymous_M234I; nonsynonymous_M234I |
| 5144 | C | T | 4.43E-04 | | | | | ORF1ab | synonymous_L1627L |

| | | | | | | | | | |
|---|---|---|---|---|---|---|---|---|---|
| 5161 | T | A | 4.43E-04 | | | | | ORF1ab | nonsynonymous_F1632L |
| 5170 | C | T | 4.43E-04 | | | | | ORF1ab | synonymous_Y1635Y |
| 16393 | C | T | 4.43E-04 | | | | | ORF1ab | nonsynonymous_P5377S |
| 17898 | C | T | 4.43E-04 | | | | | ORF1ab | synonymous_H5878H |
| 18377 | C | T | 4.43E-04 | | | | | ORF1ab | nonsynonymous_T6038I |
| 19263 | C | T | 4.43E-04 | | | | | ORF1ab | synonymous_N6333N |
| 19713 | T | C | 4.43E-04 | | | | | ORF1ab | synonymous_V6483V |
| 20709 | G | T | 4.43E-04 | | | | | ORF1ab | nonsynonymous_M6815I |
| 20759 | C | T | 4.43E-04 | A | 6.32E-05 | | | ORF1ab | nonsynonymous_A6832V; nonsynonymous_A6832E |
| 23868 | G | T | 4.43E-04 | | | | | S | nonsynonymous_G769V |
| 24586 | G | T | 4.43E-04 | | | | | S | synonymous_V1008V |
| 25463 | C | T | 4.43E-04 | A | 1.26E-04 | | | ORF3a | nonsynonymous_T24I; nonsynonymous_T24N |
| 25511 | C | T | 4.43E-04 | | | | | ORF3a | nonsynonymous_S40L |
| 26512 | T | C | 4.43E-04 | | | | | intergenic | |
| 28603 | C | T | 4.43E-04 | | | | | N | synonymous_F110F |
| 6449 | C | T | 4.43E-04 | | | | | ORF1ab | nonsynonymous_L2062F |
| 7472 | G | A | 4.43E-04 | | | | | ORF1ab | nonsynonymous_G2403S |
| 9422 | G | T | 4.43E-04 | | | | | ORF1ab | nonsynonymous_V3053L |
| 10770 | A | G | 4.43E-04 | | | | | ORF1ab | nonsynonymous_Y3502C |
| 12525 | C | T | 4.43E-04 | | | | | ORF1ab | nonsynonymous_T4087I |
| 13356 | C | T | 4.43E-04 | | | | | ORF1ab | nonsynonymous_T4364I |
| 14120 | C | T | 4.43E-04 | | | | | ORF1ab | nonsynonymous_P4619L |
| 14636 | C | T | 4.43E-04 | | | | | ORF1ab | nonsynonymous_A4791V |
| 15327 | G | T | 4.43E-04 | A | 1.26E-04 | | | ORF1ab | nonsynonymous_M5021I; nonsynonymous_M5021I |
| 15763 | C | T | 4.43E-04 | | | | | ORF1ab | synonymous_L5167L |
| 16054 | C | T | 4.43E-04 | | | | | ORF1ab | nonsynonymous_H5264Y |

| | | | | | | | | | |
|---|---|---|---|---|---|---|---|---|---|
| 16377 | G | T | 4.43E-04 | C | 6.32E-05 | | | ORF1ab | synonymous_P5371P; synonymous_P5371P |
| 17859 | T | C | 4.43E-04 | | | | | ORF1ab | synonymous_Y5865Y |
| 18616 | A | G | 4.43E-04 | | | | | ORF1ab | nonsynonymous_T6118A |
| 19011 | C | T | 4.43E-04 | | | | | ORF1ab | synonymous_D6249D |
| 19140 | C | T | 4.43E-04 | | | | | ORF1ab | synonymous_F6292F |
| 20692 | C | T | 4.43E-04 | A | 6.32E-05 | | | ORF1ab | nonsynonymous_P6810S; nonsynonymous_P6810T |
| 21795 | G | T | 4.43E-04 | A | 6.32E-05 | | | S | nonsynonymous_R78M; nonsynonymous_R78K |
| 23536 | C | T | 4.43E-04 | | | | | S | synonymous_N658N |
| 26060 | C | T | 4.43E-04 | | | | | ORF3a | nonsynonymous_T223I |
| 26660 | G | T | 4.43E-04 | A | 6.32E-05 | | | M | nonsynonymous_L46F; synonymous_L46L |
| 27213 | C | T | 4.43E-04 | | | | | ORF6 | synonymous_L4L |
| 28541 | G | T | 4.43E-04 | | | | | N | nonsynonymous_A90S |
| 28648 | T | C | 4.43E-04 | | | | | N | synonymous_A125A |
| 362 | G | A | 4.43E-04 | | | | | ORF1ab | nonsynonymous_D33N |
| 374 | G | A | 4.43E-04 | | | | | ORF1ab | nonsynonymous_E37K |
| 6926 | C | T | 4.43E-04 | | | | | ORF1ab | nonsynonymous_P2221S |
| 7728 | C | T | 4.43E-04 | A | 1.26E-04 | | | ORF1ab | nonsynonymous_S2488F; nonsynonymous_S2488Y |
| 835 | C | T | 4.43E-04 | | | | | ORF1ab | synonymous_F190F |
| 10440 | C | T | 4.43E-04 | | | | | ORF1ab | nonsynonymous_A3392V |
| 11575 | C | T | 4.43E-04 | A | 6.32E-05 | | | ORF1ab | synonymous_F3770F; nonsynonymous_F3770L |
| 11653 | C | T | 4.43E-04 | | | | | ORF1ab | synonymous_L3796L |
| 12138 | C | T | 4.43E-04 | | | | | ORF1ab | nonsynonymous_A3958V |
| 12488 | C | T | 4.43E-04 | | | | | ORF1ab | nonsynonymous_P4075S |

| | | | | | | | | | |
|---|---|---|---|---|---|---|---|---|---|
| 12685 | G | T | 4.43E-04 | C | 6.32E-05 | | | ORF1ab | nonsynonymous_Q4140H; nonsynonymous_Q4140H |
| 1473 | C | T | 4.43E-04 | | | | | ORF1ab | nonsynonymous_T403I |
| 2933 | C | T | 4.43E-04 | | | | | ORF1ab | nonsynonymous_L890F |
| 3011 | T | C | 4.43E-04 | | | | | ORF1ab | synonymous_L916L |
| 3034 | T | C | 4.43E-04 | | | | | ORF1ab | synonymous_S923S |
| 3116 | A | G | 4.43E-04 | | | | | ORF1ab | nonsynonymous_T951A |
| 3411 | C | T | 4.43E-04 | | | | | ORF1ab | nonsynonymous_A1049V |
| 4683 | C | T | 4.43E-04 | | | | | ORF1ab | nonsynonymous_A1473V |
| 491 | G | A | 4.43E-04 | | | | | ORF1ab | nonsynonymous_A76T |
| 6070 | C | T | 4.43E-04 | | | | | ORF1ab | synonymous_I1935I |
| 6539 | C | T | 4.43E-04 | | | | | ORF1ab | nonsynonymous_H2092Y |
| 7735 | C | T | 4.43E-04 | | | | | ORF1ab | synonymous_I2490I |
| 8032 | A | G | 4.43E-04 | | | | | ORF1ab | synonymous_K2589K |
| 9195 | C | T | 4.43E-04 | | | | | ORF1ab | nonsynonymous_T2977I |
| 9634 | A | T | 4.43E-04 | | | | | ORF1ab | nonsynonymous_L3123F |
| 10138 | C | T | 4.43E-04 | | | | | ORF1ab | synonymous_N3291N |
| 1102 | C | T | 4.43E-04 | | | | | ORF1ab | synonymous_S279S |
| 11747 | C | T | 4.43E-04 | | | | | ORF1ab | synonymous_L3828L |
| 11941 | C | T | 4.43E-04 | | | | | ORF1ab | synonymous_V3892V |
| 12068 | C | T | 4.43E-04 | G | 6.32E-05 | | | ORF1ab | synonymous_L3935L; nonsynonymous_L3935V |
| 12651 | C | T | 4.43E-04 | | | | | ORF1ab | nonsynonymous_T4129I |
| 13119 | C | T | 4.43E-04 | | | | | ORF1ab | nonsynonymous_A4285V |
| 1377 | C | T | 4.43E-04 | | | | | ORF1ab | nonsynonymous_P371L |
| 3743 | C | T | 4.43E-04 | | | | | ORF1ab | nonsynonymous_H1160Y |
| 3817 | C | T | 4.43E-04 | | | | | ORF1ab | synonymous_D1184D |
| 7798 | G | T | 4.43E-04 | | | | | ORF1ab | nonsynonymous_K2511N |
| 8599 | T | C | 4.43E-04 | | | | | ORF1ab | synonymous_L2778L |
| 8626 | C | T | 4.43E-04 | | | | | ORF1ab | synonymous_F2787F |

| | | | | | | | | | |
|---|---|---|---|---|---|---|---|---|---|
| 9130 | G | T | 4.43E-04 | | | | | ORF1ab | synonymous_V2955V |
| 9190 | G | T | 4.43E-04 | C | 1.26E-04 | A | 6.32E-05 | ORF1ab | synonymous_V2975V; synonymous_V2975V; synonymous_V2975V |
| 9803 | C | T | 4.43E-04 | | | | | ORF1ab | synonymous_L3180L |
| 29835 | C | A | 4.37E-04 | T | 1.46E-04 | G | 7.28E-05 | three_prime_UTR | |
| 29834 | T | C | 4.26E-04 | A | 2.13E-04 | G | 7.10E-05 | three_prime_UTR | |
| 29783 | G | T | 4.03E-04 | A | 2.02E-04 | | | three_prime_UTR | |
| 61 | G | T | 3.96E-04 | A | 6.60E-05 | | | five_prime_UTR | |
| 29776 | A | G | 3.94E-04 | T | 6.57E-05 | | | three_prime_UTR | |
| 29743 | C | T | 3.83E-04 | G | 6.39E-05 | | | three_prime_UTR | |
| 203 | C | T | 3.83E-04 | G | 6.38E-05 | | | five_prime_UTR | |
| 222 | C | T | 3.83E-04 | | | | | five_prime_UTR | |
| 29719 | C | T | 3.82E-04 | A | 6.37E-05 | | | three_prime_UTR | |
| 29668 | C | T | 3.80E-04 | | | | | ORF10 | synonymous_L37L |
| 27600 | C | T | 3.80E-04 | | | | | ORF7a | synonymous_D69D |
| 27613 | G | T | 3.80E-04 | | | | | ORF7a | nonsynonymous_V74F |
| 27513 | C | T | 3.80E-04 | | | | | ORF7a | synonymous_Y40Y |
| 27667 | G | T | 3.80E-04 | | | | | ORF7a | stopgain_E92X |
| 27549 | C | T | 3.80E-04 | | | | | ORF7a | synonymous_N52N |
| 27801 | T | C | 3.80E-04 | | | | | ORF7b | nonsynonymous_F16L |
| 27923 | C | T | 3.80E-04 | | | | | ORF8 | synonymous_I10I |
| 27925 | C | T | 3.80E-04 | | | | | ORF8 | nonsynonymous_T11I |
| 27972 | C | T | 3.80E-04 | | | | | ORF8 | stopgain_Q27X |
| 28202 | G | T | 3.80E-04 | A | 1.90E-04 | | | ORF8 | synonymous_S103S; synonymous_S103S |
| 22480 | C | T | 3.80E-04 | | | | | S | synonymous_F306F |
| 29477 | G | T | 3.80E-04 | | | | | N | nonsynonymous_D402Y |
| 5184 | C | T | 3.80E-04 | | | | | ORF1ab | nonsynonymous_P1640L |
| 29555 | C | T | 3.80E-04 | | | | | intergenic | |

| | | | | | | | | | |
|---|---|---|---|---|---|---|---|---|---|
| 20429 | C | T | 3.80E-04 | | | | | ORF1ab | nonsynonymous_P6722L |
| 21305 | G | A | 3.80E-04 | | | | | ORF1ab | nonsynonymous_R7014H |
| 2704 | C | T | 3.80E-04 | | | | | ORF1ab | synonymous_F813F |
| 22120 | C | T | 3.80E-04 | | | | | S | synonymous_F186F |
| 2632 | G | T | 3.80E-04 | | | | | ORF1ab | nonsynonymous_M789I |
| 2773 | C | T | 3.80E-04 | | | | | ORF1ab | synonymous_Y836Y |
| 29398 | G | T | 3.80E-04 | | | | | N | nonsynonymous_K375N |
| 29473 | G | A | 3.80E-04 | | | | | N | synonymous_L400L |
| 13554 | C | T | 3.80E-04 | | | | | ORF1ab | synonymous_Y4430Y |
| 21549 | C | T | 3.80E-04 | | | | | ORF1ab | synonymous_N7095N |
| 25726 | G | T | 3.80E-04 | | | | | ORF3a | nonsynonymous_V112F |
| 29527 | G | A | 3.80E-04 | T | 1.27E-04 | | | N | synonymous_Q418Q; nonsynonymous_Q418H |
| 29593 | G | A | 3.80E-04 | | | | | ORF10 | synonymous_T12T |
| 1707 | C | T | 3.79E-04 | | | | | ORF1ab | nonsynonymous_S481F |
| 5736 | C | T | 3.79E-04 | | | | | ORF1ab | nonsynonymous_A1824V |
| 20051 | C | T | 3.79E-04 | | | | | ORF1ab | nonsynonymous_T6596I |
| 21642 | C | T | 3.79E-04 | | | | | S | nonsynonymous_A27V |
| 23593 | G | T | 3.79E-04 | A | 6.32E-05 | A | 6.32E-05 | S | nonsynonymous_Q677H; synonymous_Q677Q; synonymous_Q677Q |
| 25886 | C | T | 3.79E-04 | | | | | ORF3a | nonsynonymous_S165F |
| 27509 | C | T | 3.79E-04 | | | | | ORF7a | nonsynonymous_T39I |
| 20580 | G | T | 3.79E-04 | A | 1.90E-04 | | | ORF1ab | synonymous_V6772V; synonymous_V6772V |
| 22918 | G | T | 3.79E-04 | | | | | S | synonymous_L452L |
| 28877 | A | T | 3.79E-04 | | | | | N | nonsynonymous_S202C |
| 29179 | G | T | 3.79E-04 | | | | | N | synonymous_P302P |
| 29212 | G | T | 3.79E-04 | A | 6.32E-05 | | | N | synonymous_A313A; synonymous_A313A |

| | | | | | | | | | |
|---|---|---|---|---|---|---|---|---|---|
| 29218 | C | T | 3.79E-04 | G | 6.32E-05 | | | N | synonymous_F315F; nonsynonymous_F315L |
| 29311 | C | T | 3.79E-04 | | | | | N | synonymous_F346F |
| 335 | C | T | 3.79E-04 | | | | | ORF1ab | nonsynonymous_R24C |
| 337 | C | T | 3.79E-04 | | | | | ORF1ab | synonymous_R24R |
| 3994 | G | T | 3.79E-04 | A | 2.53E-04 | | | ORF1ab | synonymous_L1243L; synonymous_L1243L |
| 17004 | C | T | 3.79E-04 | | | | | ORF1ab | synonymous_L5580L |
| 18086 | C | T | 3.79E-04 | A | 2.53E-04 | | | ORF1ab | nonsynonymous_T5941I; nonsynonymous_T5941N |
| 20844 | C | T | 3.79E-04 | A | 6.32E-05 | | | ORF1ab | synonymous_P6860P; synonymous_P6860P |
| 21976 | T | C | 3.79E-04 | | | | | S | synonymous_D138D |
| 22021 | G | T | 3.79E-04 | | | | | S | nonsynonymous_M153I |
| 22295 | C | T | 3.79E-04 | | | | | S | nonsynonymous_H245Y |
| 23590 | T | C | 3.79E-04 | | | | | S | synonymous_T676T |
| 24370 | C | T | 3.79E-04 | | | | | S | synonymous_D936D |
| 25096 | C | T | 3.79E-04 | G | 6.32E-05 | | | S | synonymous_N1178N; nonsynonymous_N1178K |
| 25217 | G | T | 3.79E-04 | | | | | S | nonsynonymous_G1219C |
| 25249 | G | T | 3.79E-04 | | | | | S | nonsynonymous_M1229I |
| 25613 | C | T | 3.79E-04 | | | | | ORF3a | nonsynonymous_S74F |
| 26497 | T | C | 3.79E-04 | | | | | intergenic | |
| 26882 | C | T | 3.79E-04 | | | | | M | synonymous_L120L |
| 27163 | G | T | 3.79E-04 | | | | | M | nonsynonymous_S214I |
| 27382 | G | C | 3.79E-04 | T | 1.90E-04 | | | ORF6 | nonsynonymous_D61H; nonsynonymous_D61Y |
| 28841 | A | G | 3.79E-04 | | | | | N | nonsynonymous_S190G |
| 28852 | T | C | 3.79E-04 | | | | | N | synonymous_S193S |
| 28869 | C | T | 3.79E-04 | | | | | N | nonsynonymous_P199L |

| | | | | | | | | | |
|---|---|---|---|---|---|---|---|---|---|
| 302 | C | T | 3.79E-04 | | | | | ORF1ab | nonsynonymous_H13Y |
| 5178 | C | T | 3.79E-04 | | | | | ORF1ab | nonsynonymous_T1638I |
| 5284 | C | T | 3.79E-04 | | | | | ORF1ab | synonymous_N1673N |
| 11782 | A | G | 3.79E-04 | | | | | ORF1ab | synonymous_K3839K |
| 12621 | C | T | 3.79E-04 | | | | | ORF1ab | nonsynonymous_S4119L |
| 14178 | C | T | 3.79E-04 | | | | | ORF1ab | synonymous_T4638T |
| 14183 | C | T | 3.79E-04 | | | | | ORF1ab | nonsynonymous_T4640I |
| 16075 | G | T | 3.79E-04 | | | | | ORF1ab | nonsynonymous_D5271Y |
| 16846 | G | T | 3.79E-04 | A | 6.32E-05 | | | ORF1ab | nonsynonymous_D5528Y; nonsynonymous_D5528N |
| 17122 | G | T | 3.79E-04 | A | 2.53E-04 | | | ORF1ab | nonsynonymous_A5620S; nonsynonymous_A5620T |
| 17278 | G | T | 3.79E-04 | | | | | ORF1ab | nonsynonymous_V5672L |
| 17562 | G | T | 3.79E-04 | | | | | ORF1ab | synonymous_R5766R |
| 17565 | T | C | 3.79E-04 | | | | | ORF1ab | synonymous_R5767R |
| 18907 | G | T | 3.79E-04 | | | | | ORF1ab | nonsynonymous_V6215F |
| 21974 | G | C | 3.79E-04 | T | 1.90E-04 | | | S | nonsynonymous_D138H; nonsynonymous_D138Y |
| 22201 | G | A | 3.79E-04 | T | 1.90E-04 | | | S | synonymous_V213V; synonymous_V213V |
| 22203 | G | T | 3.79E-04 | | | | | S | nonsynonymous_R214L |
| 22802 | C | G | 3.79E-04 | | | | | S | nonsynonymous_Q414E |
| 23248 | C | T | 3.79E-04 | | | | | S | synonymous_F562F |
| 23856 | G | A | 3.79E-04 | T | 3.16E-04 | | | S | nonsynonymous_R765H; nonsynonymous_R765L |
| 24325 | A | G | 3.79E-04 | | | | | S | synonymous_K921K |
| 26270 | C | T | 3.79E-04 | | | | | E | nonsynonymous_T9I |
| 26781 | C | T | 3.79E-04 | | | | | M | nonsynonymous_L87F |
| 27383 | A | T | 3.79E-04 | | | | | ORF6 | nonsynonymous_D61V |
| 6145 | C | T | 3.79E-04 | | | | | ORF1ab | synonymous_F1960F |

| | | | | | | | | | |
|---|---|---|---|---|---|---|---|---|---|
| 6463 | G | T | 3.79E-04 | | | | | ORF1ab | synonymous_V2066V |
| 6968 | C | T | 3.79E-04 | A | 1.26E-04 | | | ORF1ab | synonymous_L2235L; nonsynonymous_L2235I |
| 7936 | G | T | 3.79E-04 | A | 6.32E-05 | | | ORF1ab | synonymous_A2557A; synonymous_A2557A |
| 9319 | C | T | 3.79E-04 | A | 6.32E-05 | | | ORF1ab | synonymous_F3018F; nonsynonymous_F3018L |
| 10741 | C | T | 3.79E-04 | | | | | ORF1ab | synonymous_D3492D |
| 10889 | C | T | 3.79E-04 | | | | | ORF1ab | nonsynonymous_R3542C |
| 10908 | C | T | 3.79E-04 | | | | | ORF1ab | nonsynonymous_A3548V |
| 13378 | C | T | 3.79E-04 | | | | | ORF1ab | synonymous_T4371T |
| 14407 | C | T | 3.79E-04 | A | 6.32E-05 | | | ORF1ab | nonsynonymous_P4715S; nonsynonymous_P4715T |
| 14622 | G | T | 3.79E-04 | | | | | ORF1ab | synonymous_T4786T |
| 14898 | C | T | 3.79E-04 | | | | | ORF1ab | synonymous_G4878G |
| 15216 | C | T | 3.79E-04 | | | | | ORF1ab | synonymous_S4984S |
| 15957 | G | T | 3.79E-04 | | | | | ORF1ab | synonymous_G5231G |
| 16428 | C | T | 3.79E-04 | | | | | ORF1ab | synonymous_Y5388Y |
| 16456 | T | G | 3.79E-04 | C | 1.26E-04 | | | ORF1ab | nonsynonymous_S5398A; nonsynonymous_S5398P |
| 16824 | A | G | 3.79E-04 | | | | | ORF1ab | synonymous_G5520G |
| 16859 | C | T | 3.79E-04 | | | | | ORF1ab | nonsynonymous_A5532V |
| 17163 | C | T | 3.79E-04 | | | | | ORF1ab | synonymous_C5633C |
| 1722 | C | T | 3.79E-04 | | | | | ORF1ab | nonsynonymous_A486V |
| 17679 | G | T | 3.79E-04 | A | 6.32E-05 | | | ORF1ab | synonymous_T5805T; synonymous_T5805T |
| 17939 | C | T | 3.79E-04 | | | | | ORF1ab | nonsynonymous_A5892V |
| 18401 | C | T | 3.79E-04 | | | | | ORF1ab | nonsynonymous_P6046L |
| 18501 | C | T | 3.79E-04 | | | | | ORF1ab | synonymous_Y6079Y |

| | | | | | | | | | |
|---|---|---|---|---|---|---|---|---|---|
| 18803 | G | T | 3.79E-04 | A | 6.32E-05 | | | ORF1ab | nonsynonymous_S6180I; nonsynonymous_S6180N |
| 18898 | G | T | 3.79E-04 | | | | | ORF1ab | nonsynonymous_V6212F |
| 19018 | C | T | 3.79E-04 | | | | | ORF1ab | nonsynonymous_P6252S |
| 19073 | A | G | 3.79E-04 | | | | | ORF1ab | nonsynonymous_D6270G |
| 19269 | C | T | 3.79E-04 | | | | | ORF1ab | synonymous_N6335N |
| 2161 | A | G | 3.79E-04 | | | | | ORF1ab | synonymous_E632E |
| 23393 | C | T | 3.79E-04 | | | | | S | nonsynonymous_L611F |
| 23453 | C | T | 3.79E-04 | | | | | S | nonsynonymous_P631S |
| 23755 | G | T | 3.79E-04 | C | 6.32E-05 | | | S | nonsynonymous_M731I; nonsynonymous_M731I |
| 25339 | C | T | 3.79E-04 | | | | | S | synonymous_D1259D |
| 26058 | C | T | 3.79E-04 | | | | | ORF3a | synonymous_D222D |
| 26326 | C | T | 3.79E-04 | | | | | E | synonymous_L28L |
| 26640 | G | T | 3.79E-04 | | | | | M | nonsynonymous_A40S |
| 26907 | C | G | 3.79E-04 | A | 2.53E-04 | T | 1.90E-04 | M | nonsynonymous_L129V; nonsynonymous_L129M; synonymous_L129L |
| 27261 | G | T | 3.79E-04 | A | 3.16E-04 | | | ORF6 | nonsynonymous_R20S; synonymous_R20R |
| 27415 | G | T | 3.79E-04 | A | 6.32E-05 | | | ORF7a | nonsynonymous_A8S; nonsynonymous_A8T |
| 28754 | C | T | 3.79E-04 | | | | | N | nonsynonymous_L161F |
| 3738 | C | T | 3.79E-04 | | | | | ORF1ab | nonsynonymous_P1158L |
| 3859 | A | G | 3.79E-04 | | | | | ORF1ab | synonymous_Q1198Q |
| 4321 | C | T | 3.79E-04 | | | | | ORF1ab | synonymous_A1352A |
| 4582 | C | T | 3.79E-04 | | | | | ORF1ab | synonymous_N1439N |
| 6285 | C | T | 3.79E-04 | | | | | ORF1ab | nonsynonymous_T2007I |
| 6385 | C | T | 3.79E-04 | | | | | ORF1ab | synonymous_A2040A |
| 6819 | G | T | 3.79E-04 | | | | | ORF1ab | nonsynonymous_S2185I |

| | | | | | | | | | |
|---|---|---|---|---|---|---|---|---|---|
| 6975 | G | T | 3.79E-04 | | | | | ORF1ab | nonsynonymous_S2237I |
| 7225 | T | C | 3.79E-04 | | | | | ORF1ab | synonymous_A2320A |
| 9565 | C | T | 3.79E-04 | A | 6.32E-05 | | | ORF1ab | synonymous_F3100F; nonsynonymous_F3100L |
| 9857 | C | T | 3.79E-04 | | | | | ORF1ab | synonymous_L3198L |
| 10377 | C | T | 3.79E-04 | | | | | ORF1ab | nonsynonymous_P3371L |
| 11405 | G | A | 3.79E-04 | | | | | ORF1ab | nonsynonymous_V3714I |
| 1281 | C | T | 3.79E-04 | | | | | ORF1ab | nonsynonymous_A339V |
| 1420 | C | T | 3.79E-04 | | | | | ORF1ab | synonymous_A385A |
| 2447 | G | A | 3.79E-04 | | | | | ORF1ab | nonsynonymous_G728S |
| 2507 | C | T | 3.79E-04 | | | | | ORF1ab | nonsynonymous_P748S |
| 2939 | C | T | 3.79E-04 | | | | | ORF1ab | nonsynonymous_P892S |
| 3419 | G | A | 3.79E-04 | | | | | ORF1ab | nonsynonymous_V1052I |
| 3421 | A | G | 3.79E-04 | | | | | ORF1ab | synonymous_V1052V |
| 3593 | C | T | 3.79E-04 | | | | | ORF1ab | nonsynonymous_L1110F |
| 3737 | C | T | 3.79E-04 | | | | | ORF1ab | nonsynonymous_P1158S |
| 4510 | G | A | 3.79E-04 | A | 3.79E-04 | C | 6.32E-05 | ORF1ab | synonymous_V1415V; synonymous_V1415V; synonymous_V1415V |
| 6558 | C | T | 3.79E-04 | | | | | ORF1ab | nonsynonymous_A2098V |
| 6691 | T | C | 3.79E-04 | | | | | ORF1ab | synonymous_A2142A |
| 875 | C | T | 3.79E-04 | | | | | ORF1ab | nonsynonymous_L204F |
| 10070 | A | C | 3.79E-04 | | | | | ORF1ab | nonsynonymous_M3269L |
| 10232 | C | T | 3.79E-04 | | | | | ORF1ab | nonsynonymous_R3323C |
| 10604 | C | T | 3.79E-04 | | | | | ORF1ab | nonsynonymous_P3447S |
| 10702 | C | T | 3.79E-04 | | | | | ORF1ab | synonymous_D3479D |
| 1079 | A | C | 3.79E-04 | | | | | ORF1ab | nonsynonymous_N272H |
| 11222 | G | T | 3.79E-04 | | | | | ORF1ab | nonsynonymous_V3653F |
| 11410 | G | A | 3.79E-04 | | | | | ORF1ab | synonymous_L3715L |
| 11956 | C | T | 3.79E-04 | | | | | ORF1ab | synonymous_D3897D |

| | | | | | | | | | |
|---|---|---|---|---|---|---|---|---|---|
| 12073 | C | T | 3.79E-04 | | | | | ORF1ab | synonymous_D3936D |
| 12473 | C | T | 3.79E-04 | | | | | ORF1ab | synonymous_L4070L |
| 1288 | C | T | 3.79E-04 | | | | | ORF1ab | synonymous_C341C |
| 1355 | G | T | 3.79E-04 | | | | | ORF1ab | nonsynonymous_A364S |
| 1812 | C | T | 3.79E-04 | | | | | ORF1ab | nonsynonymous_A516V |
| 1862 | C | T | 3.79E-04 | | | | | ORF1ab | nonsynonymous_L533F |
| 2508 | C | T | 3.79E-04 | | | | | ORF1ab | nonsynonymous_P748L |
| 3622 | C | T | 3.79E-04 | | | | | ORF1ab | synonymous_G1119G |
| 3653 | C | T | 3.79E-04 | | | | | ORF1ab | nonsynonymous_L1130F |
| 3692 | G | T | 3.79E-04 | | | | | ORF1ab | nonsynonymous_V1143F |
| 476 | A | G | 3.79E-04 | | | | | ORF1ab | nonsynonymous_I71V |
| 4795 | C | T | 3.79E-04 | | | | | ORF1ab | synonymous_S1510S |
| 677 | G | A | 3.79E-04 | | | | | ORF1ab | nonsynonymous_A138T |
| 799 | C | T | 3.79E-04 | | | | | ORF1ab | synonymous_N178N |
| 9170 | C | T | 3.79E-04 | | | | | ORF1ab | nonsynonymous_L2969F |
| 922 | G | A | 3.79E-04 | T | 6.32E-05 | | | ORF1ab | synonymous_L219L; synonymous_L219L |
| 9977 | C | A | 3.79E-04 | | | | | ORF1ab | nonsynonymous_L3238I |
| 29784 | C | T | 3.36E-04 | | | | | three_prime_UTR | |
| 173 | C | T | 3.19E-04 | | | | | five_prime_UTR | |
| 29737 | G | A | 3.19E-04 | T | 2.55E-04 | C | 6.38E-05 | three_prime_UTR | |
| 219 | G | T | 3.19E-04 | | | | | five_prime_UTR | |
| 29722 | C | T | 3.18E-04 | | | | | three_prime_UTR | |
| 22344 | G | T | 3.17E-04 | A | 1.27E-04 | | | S | nonsynonymous_G261V; nonsynonymous_G261D |
| 24389 | A | C | 3.17E-04 | | | | | S | nonsynonymous_S943R |
| 24390 | G | C | 3.17E-04 | T | 1.90E-04 | | | S | nonsynonymous_S943T; nonsynonymous_S943I |
| 27703 | G | T | 3.17E-04 | | | | | ORF7a | nonsynonymous_V104F |
| 27800 | C | T | 3.17E-04 | | | | | ORF7b | synonymous_A15A |

| | | | | | | | | | |
|---|---|---|---|---|---|---|---|---|---|
| 19527 | T | C | 3.17E-04 | | | | | ORF1ab | synonymous_D6421D |
| 19308 | A | G | 3.17E-04 | | | | | ORF1ab | synonymous_K6348K |
| 27924 | A | G | 3.16E-04 | | | | | ORF8 | nonsynonymous_T11A |
| 28118 | T | C | 3.16E-04 | | | | | ORF8 | synonymous_D75D |
| 27879 | C | T | 3.16E-04 | | | | | ORF7b | nonsynonymous_H42Y |
| 21381 | C | T | 3.16E-04 | | | | | ORF1ab | synonymous_S7039S |
| 13476 | C | T | 3.16E-04 | | | | | ORF1ab | nonsynonymous_A4404V |
| 2659 | G | T | 3.16E-04 | | | | | ORF1ab | nonsynonymous_K798N |
| 29405 | G | A | 3.16E-04 | C | 6.33E-05 | | | N | nonsynonymous_E378K; nonsynonymous_E378Q |
| 29421 | C | T | 3.16E-04 | | | | | N | nonsynonymous_P383L |
| 29440 | G | T | 3.16E-04 | A | 1.90E-04 | | | N | nonsynonymous_Q389H; synonymous_Q389Q |
| 13617 | G | T | 3.16E-04 | | | | | ORF1ab | nonsynonymous_K4451N |
| 22432 | C | T | 3.16E-04 | | | | | S | synonymous_D290D |
| 22530 | C | T | 3.16E-04 | | | | | S | nonsynonymous_T323I |
| 13430 | C | T | 3.16E-04 | A | 6.33E-05 | | | ORF1ab | nonsynonymous_P4389S; nonsynonymous_P4389T |
| 13620 | C | T | 3.16E-04 | | | | | ORF1ab | synonymous_D4452D |
| 19961 | C | T | 3.16E-04 | | | | | ORF1ab | nonsynonymous_T6566M |
| 28254 | A | C | 3.16E-04 | T | 1.27E-04 | | | ORF8 | nonsynonymous_I121L; nonsynonymous_I121F |
| 20320 | C | T | 3.16E-04 | | | | | ORF1ab | nonsynonymous_H6686Y |
| 25690 | G | T | 3.16E-04 | | | | | ORF3a | nonsynonymous_G100C |
| 29607 | T | C | 3.16E-04 | | | | | ORF10 | nonsynonymous_L17P |
| 13512 | A | T | 3.16E-04 | | | | | ORF1ab | synonymous_T4416T |
| 13513 | G | T | 3.16E-04 | | | | | ORF1ab | nonsynonymous_G4417C |
| 16466 | C | T | 3.16E-04 | | | | | ORF1ab | nonsynonymous_P5401L |
| 20055 | A | G | 3.16E-04 | | | | | ORF1ab | synonymous_E6597E |
| 21639 | C | T | 3.16E-04 | | | | | S | nonsynonymous_P26L |

| | | | | | | | | | |
|---|---|---|---|---|---|---|---|---|---|
| 25864 | A | G | 3.16E-04 | | | | | ORF3a | nonsynonymous_I158V |
| 21595 | C | T | 3.16E-04 | | | | | S | synonymous_V11V |
| 21637 | C | T | 3.16E-04 | | | | | S | synonymous_P25P |
| 21774 | C | T | 3.16E-04 | | | | | S | nonsynonymous_S71F |
| 28868 | C | T | 3.16E-04 | A | 6.32E-05 | | | N | nonsynonymous_P199S; nonsynonymous_P199T |
| 5239 | C | T | 3.16E-04 | | | | | ORF1ab | synonymous_Y1658Y |
| 19677 | G | T | 3.16E-04 | | | | | ORF1ab | nonsynonymous_Q6471H |
| 20002 | G | T | 3.16E-04 | | | | | ORF1ab | nonsynonymous_D6580Y |
| 20133 | C | T | 3.16E-04 | | | | | ORF1ab | synonymous_A6623A |
| 24904 | C | T | 3.16E-04 | | | | | S | synonymous_I1114I |
| 25177 | G | T | 3.16E-04 | | | | | S | nonsynonymous_K1205N |
| 26885 | C | T | 3.16E-04 | | | | | M | synonymous_N121N |
| 27503 | C | T | 3.16E-04 | | | | | ORF7a | nonsynonymous_S37F |
| 29281 | C | T | 3.16E-04 | G | 6.32E-05 | | | N | synonymous_A336A; synonymous_A336A |
| 13712 | A | G | 3.16E-04 | | | | | ORF1ab | nonsynonymous_K4483R |
| 16726 | C | T | 3.16E-04 | | | | | ORF1ab | nonsynonymous_H5488Y |
| 17550 | C | T | 3.16E-04 | | | | | ORF1ab | synonymous_L5762L |
| 18255 | G | T | 3.16E-04 | | | | | ORF1ab | nonsynonymous_M5997I |
| 23401 | G | T | 3.16E-04 | | | | | S | nonsynonymous_Q613H |
| 24821 | G | T | 3.16E-04 | | | | | S | nonsynonymous_A1087S |
| 25006 | C | T | 3.16E-04 | | | | | S | synonymous_F1148F |
| 25449 | A | G | 3.16E-04 | | | | | ORF3a | synonymous_E19E |
| 26464 | C | T | 3.16E-04 | G | 6.32E-05 | | | E | synonymous_L74L; nonsynonymous_L74V |
| 27167 | C | T | 3.16E-04 | | | | | M | synonymous_D215D |
| 27428 | T | C | 3.16E-04 | | | | | ORF7a | nonsynonymous_L12P |
| 28759 | T | C | 3.16E-04 | | | | | N | synonymous_P162P |

| | | | | | | | | | |
|---|---|---|---|---|---|---|---|---|---|
| 28827 | G | T | 3.16E-04 | A | 6.32E-05 | | | N | nonsynonymous_R185L; nonsynonymous_R185H |
| 28866 | C | T | 3.16E-04 | | | | | N | nonsynonymous_T198I |
| 28948 | C | T | 3.16E-04 | | | | | N | synonymous_D225D |
| 29006 | A | G | 3.16E-04 | | | | | N | nonsynonymous_T245A |
| 5007 | C | T | 3.16E-04 | | | | | ORF1ab | nonsynonymous_T1581M |
| 5157 | C | T | 3.16E-04 | | | | | ORF1ab | nonsynonymous_A1631V |
| 5310 | C | T | 3.16E-04 | | | | | ORF1ab | nonsynonymous_T1682I |
| 6936 | C | T | 3.16E-04 | | | | | ORF1ab | nonsynonymous_S2224F |
| 7042 | G | T | 3.16E-04 | | | | | ORF1ab | nonsynonymous_M2259I |
| 9891 | C | T | 3.16E-04 | | | | | ORF1ab | nonsynonymous_A3209V |
| 11535 | G | T | 3.16E-04 | | | | | ORF1ab | nonsynonymous_G3757V |
| 14599 | C | T | 3.16E-04 | | | | | ORF1ab | synonymous_L4779L |
| 14653 | G | A | 3.16E-04 | T | 2.53E-04 | | | ORF1ab | nonsynonymous_V4797I; nonsynonymous_V4797F |
| 15017 | C | T | 3.16E-04 | | | | | ORF1ab | nonsynonymous_A4918V |
| 15672 | G | T | 3.16E-04 | | | | | ORF1ab | nonsynonymous_E5136D |
| 16101 | C | T | 3.16E-04 | | | | | ORF1ab | synonymous_Y5279Y |
| 16111 | C | T | 3.16E-04 | | | | | ORF1ab | synonymous_L5283L |
| 16607 | A | G | 3.16E-04 | | | | | ORF1ab | nonsynonymous_N5448S |
| 1691 | A | G | 3.16E-04 | | | | | ORF1ab | nonsynonymous_I476V |
| 17427 | G | T | 3.16E-04 | | | | | ORF1ab | synonymous_V5721V |
| 17518 | C | T | 3.16E-04 | | | | | ORF1ab | nonsynonymous_L5752F |
| 17608 | G | T | 3.16E-04 | | | | | ORF1ab | nonsynonymous_D5782Y |
| 18082 | A | G | 3.16E-04 | | | | | ORF1ab | nonsynonymous_I5940V |
| 18167 | C | T | 3.16E-04 | | | | | ORF1ab | nonsynonymous_P5968L |
| 18171 | C | T | 3.16E-04 | A | 2.53E-04 | | | ORF1ab | synonymous_G5969G; synonymous_G5969G |
| 18652 | C | T | 3.16E-04 | | | | | ORF1ab | nonsynonymous_R6130C |

| | | | | | | | | | |
|---|---|---|---|---|---|---|---|---|---|
| 20679 | G | T | 3.16E-04 | A | 1.90E-04 | | | ORF1ab | synonymous_P6805P; synonymous_P6805P |
| 20703 | C | T | 3.16E-04 | A | 6.32E-05 | | | ORF1ab | synonymous_Y6813Y; stopgain_Y6813X |
| 2110 | C | T | 3.16E-04 | | | | | ORF1ab | synonymous_N615N |
| 22713 | C | T | 3.16E-04 | | | | | S | nonsynonymous_P384L |
| 23271 | C | T | 3.16E-04 | A | 6.32E-05 | | | S | nonsynonymous_A570V; nonsynonymous_A570D |
| 23604 | C | T | 3.16E-04 | | | | | S | nonsynonymous_P681L |
| 23896 | C | T | 3.16E-04 | | | | | S | synonymous_T778T |
| 24381 | C | T | 3.16E-04 | | | | | S | nonsynonymous_S940F |
| 25469 | C | T | 3.16E-04 | | | | | ORF3a | nonsynonymous_S26L |
| 25549 | C | T | 3.16E-04 | | | | | ORF3a | nonsynonymous_L53F |
| 25599 | G | T | 3.16E-04 | | | | | ORF3a | nonsynonymous_W69C |
| 26167 | G | T | 3.16E-04 | | | | | ORF3a | nonsynonymous_V259L |
| 27008 | G | T | 3.16E-04 | C | 6.32E-05 | | | M | nonsynonymous_K162N; nonsynonymous_K162N |
| 28368 | G | A | 3.16E-04 | | | | | N | nonsynonymous_R32H |
| 28373 | G | T | 3.16E-04 | | | | | N | nonsynonymous_G34W |
| 28374 | G | A | 3.16E-04 | | | | | N | nonsynonymous_G34E |
| 6362 | G | A | 3.16E-04 | | | | | ORF1ab | nonsynonymous_A2033T |
| 6380 | C | T | 3.16E-04 | | | | | ORF1ab | nonsynonymous_L2039F |
| 6977 | G | A | 3.16E-04 | | | | | ORF1ab | nonsynonymous_V2238I |
| 8886 | T | C | 3.16E-04 | A | 2.53E-04 | | | ORF1ab | nonsynonymous_L2874S; stopgain_L2874X |
| 11293 | T | C | 3.16E-04 | | | | | ORF1ab | synonymous_G3676G |
| 13252 | C | T | 3.16E-04 | | | | | ORF1ab | synonymous_Y4329Y |
| 14104 | T | C | 3.16E-04 | | | | | ORF1ab | nonsynonymous_F4614L |
| 14267 | C | T | 3.16E-04 | | | | | ORF1ab | nonsynonymous_T4668M |
| 14673 | A | G | 3.16E-04 | | | | | ORF1ab | synonymous_K4803K |

| | | | | | | | | | |
|---|---|---|---|---|---|---|---|---|---|
| 14718 | G | T | 3.16E-04 | | | | | ORF1ab | nonsynonymous_K4818N |
| 1487 | G | A | 3.16E-04 | | | | | ORF1ab | nonsynonymous_G408S |
| 15264 | T | C | 3.16E-04 | | | | | ORF1ab | synonymous_D5000D |
| 15366 | T | C | 3.16E-04 | | | | | ORF1ab | synonymous_H5034H |
| 15439 | G | A | 3.16E-04 | | | | | ORF1ab | nonsynonymous_V5059I |
| 1547 | A | G | 3.16E-04 | | | | | ORF1ab | nonsynonymous_S428G |
| 1593 | C | T | 3.16E-04 | | | | | ORF1ab | nonsynonymous_S443F |
| 15972 | A | G | 3.16E-04 | | | | | ORF1ab | synonymous_V5236V |
| 1616 | C | T | 3.16E-04 | | | | | ORF1ab | nonsynonymous_L451F |
| 16474 | A | G | 3.16E-04 | | | | | ORF1ab | nonsynonymous_S5404G |
| 16993 | T | C | 3.16E-04 | | | | | ORF1ab | nonsynonymous_Y5577H |
| 17125 | C | T | 3.16E-04 | A | 6.32E-05 | | | ORF1ab | nonsynonymous_L5621F; nonsynonymous_L5621I |
| 17193 | G | T | 3.16E-04 | | | | | ORF1ab | nonsynonymous_E5643D |
| 17423 | A | G | 3.16E-04 | | | | | ORF1ab | nonsynonymous_Y5720C |
| 17656 | A | G | 3.16E-04 | | | | | ORF1ab | nonsynonymous_M5798V |
| 17673 | T | C | 3.16E-04 | | | | | ORF1ab | synonymous_V5803V |
| 17721 | G | T | 3.16E-04 | | | | | ORF1ab | synonymous_V5819V |
| 18083 | T | C | 3.16E-04 | | | | | ORF1ab | nonsynonymous_I5940T |
| 18526 | C | T | 3.16E-04 | | | | | ORF1ab | nonsynonymous_R6088C |
| 18771 | T | C | 3.16E-04 | | | | | ORF1ab | synonymous_V6169V |
| 18826 | G | T | 3.16E-04 | | | | | ORF1ab | nonsynonymous_V6188F |
| 18863 | T | C | 3.16E-04 | | | | | ORF1ab | nonsynonymous_I6200T |
| 18984 | G | T | 3.16E-04 | C | 6.32E-05 | | | ORF1ab | nonsynonymous_M6240I; nonsynonymous_M6240I |
| 19102 | C | T | 3.16E-04 | | | | | ORF1ab | nonsynonymous_P6280S |
| 19248 | T | C | 3.16E-04 | | | | | ORF1ab | synonymous_T6328T |
| 19254 | G | T | 3.16E-04 | | | | | ORF1ab | synonymous_V6330V |
| 19652 | A | G | 3.16E-04 | | | | | ORF1ab | nonsynonymous_N6463S |
| 19818 | A | G | 3.16E-04 | | | | | ORF1ab | synonymous_V6518V |

| | | | | | | | | | |
|---|---|---|---|---|---|---|---|---|---|
| 21077 | C | T | 3.16E-04 | | | | | ORF1ab | nonsynonymous_T6938I |
| 21721 | C | T | 3.16E-04 | | | | | S | synonymous_D53D |
| 21789 | C | T | 3.16E-04 | | | | | S | nonsynonymous_T76I |
| 21998 | C | T | 3.16E-04 | A | 6.32E-05 | | | S | nonsynonymous_H146Y; nonsynonymous_H146N |
| 22858 | C | T | 3.16E-04 | A | 6.32E-05 | | | S | synonymous_C432C; stopgain_C432X |
| 23518 | G | A | 3.16E-04 | | | | | S | synonymous_G652G |
| 23709 | C | T | 3.16E-04 | | | | | S | nonsynonymous_T716I |
| 24138 | C | T | 3.16E-04 | G | 6.32E-05 | | | S | nonsynonymous_T859I; nonsynonymous_T859S |
| 25340 | G | A | 3.16E-04 | | | | | S | nonsynonymous_D1260N |
| 25413 | C | T | 3.16E-04 | | | | | ORF3a | synonymous_I7I |
| 25460 | C | T | 3.16E-04 | | | | | ORF3a | nonsynonymous_A23V |
| 25466 | C | T | 3.16E-04 | | | | | ORF3a | nonsynonymous_P25L |
| 26002 | C | A | 3.16E-04 | | | | | ORF3a | nonsynonymous_H204N |
| 26094 | T | C | 3.16E-04 | | | | | ORF3a | synonymous_N234N |
| 26109 | G | T | 3.16E-04 | | | | | ORF3a | nonsynonymous_E239D |
| 26314 | G | A | 3.16E-04 | C | 6.32E-05 | | | E | nonsynonymous_V24M; nonsynonymous_V24L |
| 26684 | C | T | 3.16E-04 | | | | | M | synonymous_L54L |
| 28539 | G | T | 3.16E-04 | | | | | N | nonsynonymous_R89I |
| 28562 | G | A | 3.16E-04 | | | | | N | nonsynonymous_G97S |
| 28655 | G | T | 3.16E-04 | C | 6.32E-05 | | | N | nonsynonymous_D128Y; nonsynonymous_D128H |
| 28703 | G | T | 3.16E-04 | C | 1.26E-04 | | | N | nonsynonymous_D144Y; nonsynonymous_D144H |
| 3261 | C | T | 3.16E-04 | A | 6.32E-05 | | | ORF1ab | nonsynonymous_T999I; nonsynonymous_T999K |
| 344 | C | T | 3.16E-04 | | | | | ORF1ab | nonsynonymous_L27F |

| | | | | | | | | | |
|---|---|---|---|---|---|---|---|---|---|
| 354 | G | A | 3.16E-04 | | | | | ORF1ab | nonsynonymous_G30D |
| 3885 | C | A | 3.16E-04 | T | 1.26E-04 | | | ORF1ab | nonsynonymous_P1207H; nonsynonymous_P1207L |
| 3982 | T | C | 3.16E-04 | | | | | ORF1ab | synonymous_V1239V |
| 398 | C | T | 3.16E-04 | | | | | ORF1ab | nonsynonymous_H45Y |
| 6286 | C | T | 3.16E-04 | | | | | ORF1ab | synonymous_T2007T |
| 6843 | C | T | 3.16E-04 | | | | | ORF1ab | nonsynonymous_S2193F |
| 6971 | T | C | 3.16E-04 | | | | | ORF1ab | synonymous_L2236L |
| 7295 | G | T | 3.16E-04 | | | | | ORF1ab | nonsynonymous_A2344S |
| 9737 | A | G | 3.16E-04 | | | | | ORF1ab | nonsynonymous_S3158G |
| 9967 | C | T | 3.16E-04 | | | | | ORF1ab | synonymous_L3234L |
| 10078 | C | T | 3.16E-04 | | | | | ORF1ab | synonymous_F3271F |
| 12049 | C | T | 3.16E-04 | | | | | ORF1ab | synonymous_N3928N |
| 12148 | A | T | 3.16E-04 | | | | | ORF1ab | nonsynonymous_Q3961H |
| 12191 | G | T | 3.16E-04 | | | | | ORF1ab | nonsynonymous_V3976F |
| 12741 | C | T | 3.16E-04 | A | 6.32E-05 | | | ORF1ab | nonsynonymous_T4159I; nonsynonymous_T4159K |
| 1595 | G | A | 3.16E-04 | T | 6.32E-05 | | | ORF1ab | nonsynonymous_E444K; stopgain_E444X |
| 1884 | C | T | 3.16E-04 | | | | | ORF1ab | nonsynonymous_A540V |
| 1997 | C | T | 3.16E-04 | | | | | ORF1ab | synonymous_L578L |
| 2037 | C | T | 3.16E-04 | | | | | ORF1ab | nonsynonymous_A591V |
| 2062 | C | T | 3.16E-04 | | | | | ORF1ab | synonymous_A599A |
| 2388 | C | T | 3.16E-04 | | | | | ORF1ab | nonsynonymous_T708I |
| 3130 | C | T | 3.16E-04 | | | | | ORF1ab | synonymous_Y955Y |
| 3267 | C | T | 3.16E-04 | | | | | ORF1ab | nonsynonymous_T1001I |
| 3472 | A | G | 3.16E-04 | | | | | ORF1ab | synonymous_G1069G |
| 4158 | C | T | 3.16E-04 | | | | | ORF1ab | nonsynonymous_A1298V |
| 4180 | G | T | 3.16E-04 | A | 1.90E-04 | | | ORF1ab | nonsynonymous_K1305N; synonymous_K1305K |

| | | | | | | | | | |
|---|---|---|---|---|---|---|---|---|---|
| 4720 | G | T | 3.16E-04 | A | 1.90E-04 | | | ORF1ab | synonymous_A1485A; synonymous_A1485A |
| 5884 | C | T | 3.16E-04 | | | | | ORF1ab | synonymous_Y1873Y |
| 6541 | C | T | 3.16E-04 | | | | | ORF1ab | synonymous_H2092H |
| 7764 | C | T | 3.16E-04 | | | | | ORF1ab | nonsynonymous_S2500F |
| 7767 | T | C | 3.16E-04 | | | | | ORF1ab | nonsynonymous_I2501T |
| 8179 | G | T | 3.16E-04 | | | | | ORF1ab | synonymous_R2638R |
| 8344 | C | T | 3.16E-04 | | | | | ORF1ab | synonymous_D2693D |
| 920 | C | T | 3.16E-04 | | | | | ORF1ab | synonymous_L219L |
| 10252 | C | T | 3.16E-04 | | | | | ORF1ab | synonymous_F3329F |
| 10335 | C | T | 3.16E-04 | | | | | ORF1ab | nonsynonymous_A3357V |
| 10465 | G | A | 3.16E-04 | | | | | ORF1ab | synonymous_K3400K |
| 11090 | A | T | 3.16E-04 | | | | | ORF1ab | nonsynonymous_N3609Y |
| 11230 | G | T | 3.16E-04 | | | | | ORF1ab | nonsynonymous_M3655I |
| 11313 | T | C | 3.16E-04 | | | | | ORF1ab | nonsynonymous_V3683A |
| 11355 | C | T | 3.16E-04 | | | | | ORF1ab | nonsynonymous_A3697V |
| 1140 | A | C | 3.16E-04 | | | | | ORF1ab | nonsynonymous_K292T |
| 11457 | T | C | 3.16E-04 | | | | | ORF1ab | nonsynonymous_I3731T |
| 11595 | A | G | 3.16E-04 | | | | | ORF1ab | nonsynonymous_Q3777R |
| 11750 | C | T | 3.16E-04 | A | 6.32E-05 | | | ORF1ab | nonsynonymous_L3829F; nonsynonymous_L3829I |
| 12112 | T | C | 3.16E-04 | | | | | ORF1ab | synonymous_S3949S |
| 12119 | C | T | 3.16E-04 | | | | | ORF1ab | nonsynonymous_P3952S |
| 12756 | C | T | 3.16E-04 | | | | | ORF1ab | nonsynonymous_T4164I |
| 12778 | C | T | 3.16E-04 | A | 1.26E-04 | | | ORF1ab | synonymous_Y4171Y; stopgain_Y4171X |
| 12787 | A | G | 3.16E-04 | | | | | ORF1ab | synonymous_T4174T |
| 12809 | C | T | 3.16E-04 | | | | | ORF1ab | nonsynonymous_L4182F |
| 12823 | C | T | 3.16E-04 | | | | | ORF1ab | synonymous_S4186S |
| 12970 | C | T | 3.16E-04 | | | | | ORF1ab | synonymous_N4235N |

| | | | | | | | | | |
|---|---|---|---|---|---|---|---|---|---|
| 1882 | G | T | 3.16E-04 | | | | | ORF1ab | nonsynonymous_E539D |
| 1929 | C | T | 3.16E-04 | | | | | ORF1ab | nonsynonymous_A555V |
| 2215 | A | G | 3.16E-04 | | | | | ORF1ab | synonymous_K650K |
| 2293 | G | T | 3.16E-04 | | | | | ORF1ab | nonsynonymous_Q676H |
| 2869 | A | G | 3.16E-04 | | | | | ORF1ab | synonymous_V868V |
| 3231 | G | T | 3.16E-04 | | | | | ORF1ab | nonsynonymous_G989V |
| 3259 | G | T | 3.16E-04 | A | 1.26E-04 | | | ORF1ab | nonsynonymous_Q998H; synonymous_Q998Q |
| 3567 | G | A | 3.16E-04 | | | | | ORF1ab | nonsynonymous_G1101D |
| 3589 | C | T | 3.16E-04 | | | | | ORF1ab | synonymous_H1108H |
| 4148 | G | T | 3.16E-04 | | | | | ORF1ab | nonsynonymous_V1295F |
| 4202 | C | T | 3.16E-04 | | | | | ORF1ab | synonymous_L1313L |
| 4285 | G | T | 3.16E-04 | | | | | ORF1ab | nonsynonymous_E1340D |
| 472 | G | T | 3.16E-04 | | | | | ORF1ab | synonymous_V69V |
| 4897 | C | T | 3.16E-04 | | | | | ORF1ab | synonymous_F1544F |
| 6031 | C | T | 3.16E-04 | | | | | ORF1ab | synonymous_N1922N |
| 6501 | C | T | 3.16E-04 | | | | | ORF1ab | nonsynonymous_P2079L |
| 7675 | G | T | 3.16E-04 | | | | | ORF1ab | synonymous_A2470A |
| 809 | T | A | 3.16E-04 | | | | | ORF1ab | nonsynonymous_Y182N |
| 9113 | C | T | 3.16E-04 | | | | | ORF1ab | nonsynonymous_P2950S |
| 9165 | C | T | 3.16E-04 | A | 6.32E-05 | | | ORF1ab | nonsynonymous_T2967I; nonsynonymous_T2967N |
| 967 | T | C | 3.16E-04 | | | | | ORF1ab | synonymous_H234H |
| 970 | G | A | 3.16E-04 | | | | | ORF1ab | synonymous_E235E |
| 29818 | A | G | 2.73E-04 | C | 6.82E-05 | | | three_prime_UTR | |
| 55 | A | C | 2.69E-04 | | | | | five_prime_UTR | |
| 91 | G | C | 2.59E-04 | | | | | five_prime_UTR | |
| 92 | C | T | 2.59E-04 | | | | | five_prime_UTR | |
| 99 | T | C | 2.58E-04 | | | | | five_prime_UTR | |
| 101 | G | A | 2.58E-04 | T | 6.45E-05 | | | five_prime_UTR | |

| | | | | | | | | | |
|---|---|---|---|---|---|---|---|---|---|
| 127 | G | T | 2.57E-04 | A | 6.41E-05 | | | five_prime_UTR | |
| 29750 | C | T | 2.56E-04 | G | 6.39E-05 | | | three_prime_UTR | |
| 29745 | G | T | 2.56E-04 | C | 6.39E-05 | | | three_prime_UTR | |
| 183 | C | T | 2.55E-04 | | | | | five_prime_UTR | |
| 204 | G | A | 2.55E-04 | A | 2.55E-04 | C | 1.28E-04 | five_prime_UTR | |
| 211 | T | C | 2.55E-04 | | | | | five_prime_UTR | |
| 242 | G | T | 2.55E-04 | | | | | five_prime_UTR | |
| 29688 | G | T | 2.55E-04 | C | 6.37E-05 | | | three_prime_UTR | |
| 22346 | G | A | 2.53E-04 | A | 2.53E-04 | | | S | nonsynonymous_A262T;<br>nonsynonymous_A262T |
| 27577 | C | T | 2.53E-04 | | | | | ORF7a | stopgain_Q62X |
| 27604 | G | A | 2.53E-04 | T | 1.27E-04 | | | ORF7a | nonsynonymous_V71I;<br>nonsynonymous_V71L |
| 19543 | A | C | 2.53E-04 | | | | | ORF1ab | nonsynonymous_I6427L |
| 27661 | C | T | 2.53E-04 | | | | | ORF7a | stopgain_Q90X |
| 27788 | G | T | 2.53E-04 | | | | | ORF7b | nonsynonymous_L11F |
| 27559 | C | T | 2.53E-04 | | | | | ORF7a | synonymous_L56L |
| 27561 | G | T | 2.53E-04 | | | | | ORF7a | synonymous_L56L |
| 526 | G | T | 2.53E-04 | | | | | ORF1ab | nonsynonymous_E87D |
| 27684 | C | T | 2.53E-04 | | | | | ORF7a | synonymous_Y97Y |
| 27752 | C | T | 2.53E-04 | | | | | ORF7a | nonsynonymous_T120I |
| 27746 | G | T | 2.53E-04 | | | | | ORF7a | nonsynonymous_R118I |
| 19330 | T | C | 2.53E-04 | | | | | ORF1ab | nonsynonymous_F6356L |
| 27920 | C | T | 2.53E-04 | A | 6.33E-05 | A | 6.33E-05 | ORF8 | synonymous_I9I;<br>synonymous_I9I;<br>synonymous_I9I |
| 27999 | C | T | 2.53E-04 | | | | | ORF8 | nonsynonymous_P36S |
| 28044 | G | T | 2.53E-04 | | | | | ORF8 | nonsynonymous_A51S |
| 28105 | T | C | 2.53E-04 | | | | | ORF8 | nonsynonymous_I71T |
| 28139 | C | A | 2.53E-04 | | | | | ORF8 | synonymous_S82S |

| | | | | | | | | | |
|---|---|---|---|---|---|---|---|---|---|
| 22427 | G | A | 2.53E-04 | T | 6.33E-05 | | | S | nonsynonymous_V289I; nonsynonymous_V289L |
| 13571 | G | T | 2.53E-04 | | | | | ORF1ab | nonsynonymous_G4436V |
| 22450 | C | T | 2.53E-04 | | | | | S | synonymous_L296L |
| 26801 | C | G | 2.53E-04 | T | 1.90E-04 | | | M | synonymous_L93L; synonymous_L93L |
| 29425 | G | A | 2.53E-04 | T | 1.90E-04 | | | N | synonymous_Q384Q; nonsynonymous_Q384H |
| 21151 | G | T | 2.53E-04 | | | | | ORF1ab | nonsynonymous_G6963C |
| 2662 | C | T | 2.53E-04 | | | | | ORF1ab | synonymous_Y799Y |
| 29563 | C | T | 2.53E-04 | | | | | ORF10 | synonymous_G2G |
| 2675 | C | T | 2.53E-04 | A | 6.33E-05 | | | ORF1ab | nonsynonymous_P804S; nonsynonymous_P804T |
| 2706 | C | T | 2.53E-04 | | | | | ORF1ab | nonsynonymous_T814I |
| 29441 | C | G | 2.53E-04 | A | 6.33E-05 | | | N | nonsynonymous_Q390E; nonsynonymous_Q390K |
| 20483 | C | T | 2.53E-04 | | | | | ORF1ab | nonsynonymous_S6740F |
| 21635 | C | T | 2.53E-04 | | | | | S | nonsynonymous_P25S |
| 2723 | C | T | 2.53E-04 | | | | | ORF1ab | nonsynonymous_P820S |
| 29420 | C | T | 2.53E-04 | | | | | N | nonsynonymous_P383S |
| 29366 | C | T | 2.53E-04 | | | | | N | nonsynonymous_P365S |
| 20284 | A | G | 2.53E-04 | | | | | ORF1ab | nonsynonymous_I6674V |
| 21658 | C | T | 2.53E-04 | | | | | S | synonymous_F32F |
| 25797 | C | T | 2.53E-04 | | | | | ORF3a | synonymous_S135S |
| 29513 | G | T | 2.53E-04 | | | | | N | nonsynonymous_A414S |
| 13576 | G | T | 2.53E-04 | | | | | ORF1ab | nonsynonymous_A4438S |
| 19955 | C | T | 2.53E-04 | A | 6.32E-05 | A | 6.32E-05 | ORF1ab | nonsynonymous_T6564I; nonsynonymous_T6564N; nonsynonymous_T6564N |
| 21641 | G | T | 2.53E-04 | | | | | S | nonsynonymous_A27S |

| | | | | | | | | | |
|---|---|---|---|---|---|---|---|---|---|
| 23114 | C | T | 2.53E-04 | | | | | S | synonymous_L518L |
| 25650 | G | T | 2.53E-04 | | | | | ORF3a | nonsynonymous_L86F |
| 25710 | C | T | 2.53E-04 | | | | | ORF3a | synonymous_L106L |
| 25769 | G | T | 2.53E-04 | | | | | ORF3a | nonsynonymous_R126M |
| 25822 | A | G | 2.53E-04 | | | | | ORF3a | nonsynonymous_N144D |
| 26534 | C | T | 2.53E-04 | | | | | M | synonymous_S4S |
| 27208 | C | T | 2.53E-04 | | | | | ORF6 | nonsynonymous_H3Y |
| 274 | C | T | 2.53E-04 | A | 1.26E-04 | | | ORF1ab | synonymous_S3S; nonsynonymous_S3R |
| 28895 | G | T | 2.53E-04 | | | | | N | nonsynonymous_A208S |
| 13575 | T | C | 2.53E-04 | | | | | ORF1ab | synonymous_F4437F |
| 21644 | T | C | 2.53E-04 | A | 6.32E-05 | | | S | nonsynonymous_Y28H; nonsynonymous_Y28N |
| 25783 | T | C | 2.53E-04 | | | | | ORF3a | nonsynonymous_W131R |
| 20063 | T | C | 2.53E-04 | | | | | ORF1ab | nonsynonymous_V6600A |
| 21552 | C | T | 2.53E-04 | | | | | ORF1ab | synonymous_N7096N |
| 21590 | C | T | 2.53E-04 | A | 1.26E-04 | | | S | synonymous_L10L; nonsynonymous_L10I |
| 21643 | A | C | 2.53E-04 | | | | | S | synonymous_A27A |
| 21646 | C | T | 2.53E-04 | | | | | S | synonymous_Y28Y |
| 21767 | C | T | 2.53E-04 | | | | | S | nonsynonymous_H69Y |
| 25793 | G | T | 2.53E-04 | A | 6.32E-05 | | | ORF3a | nonsynonymous_R134L; nonsynonymous_R134H |
| 25889 | C | T | 2.53E-04 | | | | | ORF3a | nonsynonymous_S166L |
| 25912 | G | T | 2.53E-04 | | | | | ORF3a | nonsynonymous_G174C |
| 27441 | G | T | 2.53E-04 | | | | | ORF7a | nonsynonymous_E16D |
| 29239 | G | A | 2.53E-04 | T | 1.26E-04 | C | 6.32E-05 | N | nonsynonymous_M322I; nonsynonymous_M322I; nonsynonymous_M322I |

| | | | | | | | | | |
|---|---|---|---|---|---|---|---|---|---|
| 29260 | G | T | 2.53E-04 | A | 1.26E-04 | | | N | synonymous_T329T; synonymous_T329T |
| 29315 | G | T | 2.53E-04 | C | 1.26E-04 | | | N | nonsynonymous_D348Y; nonsynonymous_D348H |
| 5301 | C | T | 2.53E-04 | | | | | ORF1ab | nonsynonymous_A1679V |
| 16694 | C | T | 2.53E-04 | | | | | ORF1ab | nonsynonymous_T5477I |
| 20016 | C | T | 2.53E-04 | | | | | ORF1ab | synonymous_D6584D |
| 22289 | G | T | 2.53E-04 | | | | | S | nonsynonymous_A243S |
| 25214 | C | T | 2.53E-04 | | | | | S | synonymous_L1218L |
| 25244 | G | T | 2.53E-04 | C | 1.26E-04 | | | S | nonsynonymous_V1228L; nonsynonymous_V1228L |
| 27506 | G | T | 2.53E-04 | | | | | ORF7a | nonsynonymous_G38V |
| 28908 | G | T | 2.53E-04 | | | | | N | nonsynonymous_G212V |
| 29085 | C | T | 2.53E-04 | | | | | N | nonsynonymous_T271I |
| 29109 | C | T | 2.53E-04 | | | | | N | nonsynonymous_P279L |
| 29119 | C | T | 2.53E-04 | | | | | N | synonymous_T282T |
| 29149 | C | T | 2.53E-04 | A | 6.32E-05 | | | N | synonymous_I292I; synonymous_I292I |
| 29227 | G | T | 2.53E-04 | A | 1.90E-04 | | | N | synonymous_S318S; synonymous_S318S |
| 29262 | G | T | 2.53E-04 | | | | | N | nonsynonymous_W330L |
| 5340 | C | T | 2.53E-04 | | | | | ORF1ab | nonsynonymous_P1692L |
| 16975 | G | T | 2.53E-04 | | | | | ORF1ab | nonsynonymous_V5571F |
| 20555 | C | T | 2.53E-04 | | | | | ORF1ab | nonsynonymous_S6764F |
| 22104 | G | T | 2.53E-04 | | | | | S | nonsynonymous_G181V |
| 22224 | C | T | 2.53E-04 | G | 6.32E-05 | | | S | nonsynonymous_S221L; nonsynonymous_S221W |
| 24795 | C | T | 2.53E-04 | | | | | S | nonsynonymous_A1078V |
| 24812 | G | T | 2.53E-04 | | | | | S | nonsynonymous_D1084Y |
| 24926 | G | T | 2.53E-04 | | | | | S | nonsynonymous_V1122L |

| | | | | | | | | | |
|---|---|---|---|---|---|---|---|---|---|
| 24928 | G | T | 2.53E-04 | | | | | S | synonymous_V1122V |
| 25166 | G | C | 2.53E-04 | A | 6.32E-05 | | | S | nonsynonymous_E1202Q; nonsynonymous_E1202K |
| 25273 | G | T | 2.53E-04 | C | 1.26E-04 | | | S | nonsynonymous_M1237I; nonsynonymous_M1237I |
| 25276 | C | T | 2.53E-04 | A | 6.32E-05 | | | S | synonymous_T1238T; synonymous_T1238T |
| 25418 | C | A | 2.53E-04 | T | 6.32E-05 | | | ORF3a | nonsynonymous_T9K; nonsynonymous_T9I |
| 25534 | G | T | 2.53E-04 | | | | | ORF3a | nonsynonymous_V48F |
| 26370 | C | T | 2.53E-04 | | | | | E | synonymous_Y42Y |
| 26416 | G | C | 2.53E-04 | T | 1.90E-04 | | | E | nonsynonymous_V58L; nonsynonymous_V58F |
| 26458 | G | T | 2.53E-04 | C | 6.32E-05 | A | 6.32E-05 | E | nonsynonymous_D72Y; nonsynonymous_D72H; nonsynonymous_D72N |
| 26459 | A | G | 2.53E-04 | | | | | E | nonsynonymous_D72G |
| 27092 | C | T | 2.53E-04 | | | | | M | synonymous_D190D |
| 27128 | C | T | 2.53E-04 | | | | | M | synonymous_G202G |
| 27131 | C | T | 2.53E-04 | | | | | M | synonymous_N203N |
| 27498 | C | T | 2.53E-04 | | | | | ORF7a | synonymous_C35C |
| 27500 | C | T | 2.53E-04 | | | | | ORF7a | nonsynonymous_S36F |
| 28830 | C | T | 2.53E-04 | A | 6.32E-05 | | | N | nonsynonymous_S186F; nonsynonymous_S186Y |
| 28844 | C | T | 2.53E-04 | G | 6.32E-05 | | | N | nonsynonymous_R191C; nonsynonymous_R191G |
| 28961 | C | T | 2.53E-04 | | | | | N | nonsynonymous_L230F |
| 5206 | G | T | 2.53E-04 | | | | | ORF1ab | nonsynonymous_M1647I |
| 5221 | C | T | 2.53E-04 | | | | | ORF1ab | synonymous_H1652H |
| 5309 | A | G | 2.53E-04 | | | | | ORF1ab | nonsynonymous_T1682A |

| | | | | | | | | | |
|---|---|---|---|---|---|---|---|---|---|
| 5348 | C | T | 2.53E-04 | | | | | ORF1ab | synonymous_L1695L |
| 5506 | C | T | 2.53E-04 | | | | | ORF1ab | synonymous_V1747V |
| 5538 | A | T | 2.53E-04 | | | | | ORF1ab | nonsynonymous_Q1758L |
| 5551 | T | C | 2.53E-04 | | | | | ORF1ab | synonymous_L1762L |
| 7047 | C | T | 2.53E-04 | | | | | ORF1ab | nonsynonymous_S2261F |
| 10969 | C | T | 2.53E-04 | | | | | ORF1ab | synonymous_F3568F |
| 14829 | G | T | 2.53E-04 | | | | | ORF1ab | nonsynonymous_M4855I |
| 14931 | C | T | 2.53E-04 | | | | | ORF1ab | synonymous_N4889N |
| 16247 | C | T | 2.53E-04 | | | | | ORF1ab | nonsynonymous_A5328V |
| 17249 | C | T | 2.53E-04 | | | | | ORF1ab | nonsynonymous_A5662V |
| 17375 | C | T | 2.53E-04 | | | | | ORF1ab | nonsynonymous_T5704I |
| 17590 | G | T | 2.53E-04 | | | | | ORF1ab | nonsynonymous_V5776L |
| 17615 | A | G | 2.53E-04 | | | | | ORF1ab | nonsynonymous_K5784R |
| 17746 | C | T | 2.53E-04 | | | | | ORF1ab | nonsynonymous_P5828S |
| 18486 | C | T | 2.53E-04 | | | | | ORF1ab | synonymous_L6074L |
| 18639 | A | G | 2.53E-04 | | | | | ORF1ab | synonymous_K6125K |
| 18651 | G | A | 2.53E-04 | T | 1.90E-04 | | | ORF1ab | synonymous_E6129E; nonsynonymous_E6129D |
| 18657 | C | T | 2.53E-04 | | | | | ORF1ab | synonymous_T6131T |
| 18689 | T | A | 2.53E-04 | | | | | ORF1ab | nonsynonymous_F6142Y |
| 18859 | G | T | 2.53E-04 | | | | | ORF1ab | nonsynonymous_A6199S |
| 19145 | C | T | 2.53E-04 | | | | | ORF1ab | nonsynonymous_S6294F |
| 19735 | G | T | 2.53E-04 | | | | | ORF1ab | nonsynonymous_D6491Y |
| 20611 | C | T | 2.53E-04 | | | | | ORF1ab | nonsynonymous_L6783F |
| 20736 | C | T | 2.53E-04 | | | | | ORF1ab | synonymous_D6824D |
| 20823 | C | T | 2.53E-04 | | | | | ORF1ab | synonymous_N6853N |
| 20936 | C | T | 2.53E-04 | | | | | ORF1ab | nonsynonymous_T6891M |
| 22088 | C | T | 2.53E-04 | A | 6.32E-05 | | | S | nonsynonymous_L176F; nonsynonymous_L176I |
| 22207 | T | C | 2.53E-04 | | | | | S | synonymous_D215D |

| | | | | | | | | | |
|---|---|---|---|---|---|---|---|---|---|
| 22708 | G | T | 2.53E-04 | | | | | S | synonymous_V382V |
| 24096 | C | T | 2.53E-04 | | | | | S | nonsynonymous_A845V |
| 24374 | C | T | 2.53E-04 | | | | | S | nonsynonymous_L938F |
| 24382 | C | T | 2.53E-04 | | | | | S | synonymous_S940S |
| 24642 | C | T | 2.53E-04 | | | | | S | nonsynonymous_T1027I |
| 25414 | T | C | 2.53E-04 | | | | | ORF3a | nonsynonymous_F8L |
| 25513 | C | T | 2.53E-04 | | | | | ORF3a | nonsynonymous_L41F |
| 25517 | C | T | 2.53E-04 | | | | | ORF3a | nonsynonymous_P42L |
| 25948 | A | G | 2.53E-04 | | | | | ORF3a | nonsynonymous_I186V |
| 26526 | G | T | 2.53E-04 | | | | | M | nonsynonymous_A2S |
| 26690 | G | T | 2.53E-04 | | | | | M | synonymous_L56L |
| 26959 | G | A | 2.53E-04 | | | | | M | nonsynonymous_R146H |
| 27200 | A | G | 2.53E-04 | | | | | intergenic | |
| 27225 | G | T | 2.53E-04 | | | | | ORF6 | nonsynonymous_Q8H |
| 27371 | C | T | 2.53E-04 | | | | | ORF6 | nonsynonymous_P57L |
| 27810 | T | C | 2.53E-04 | | | | | ORF7b | nonsynonymous_F19L |
| 28300 | G | T | 2.53E-04 | | | | | N | nonsynonymous_Q9H |
| 28321 | G | A | 2.53E-04 | | | | | N | synonymous_T16T |
| 28371 | G | T | 2.53E-04 | A | 6.32E-05 | | | N | nonsynonymous_S33I; nonsynonymous_S33N |
| 28376 | G | T | 2.53E-04 | A | 6.32E-05 | | | N | nonsynonymous_A35S; nonsynonymous_A35T |
| 28628 | G | T | 2.53E-04 | C | 1.90E-04 | | | N | nonsynonymous_A119S; nonsynonymous_A119P |
| 6884 | G | T | 2.53E-04 | A | 6.32E-05 | | | ORF1ab | nonsynonymous_G2207C; nonsynonymous_G2207S |
| 6996 | T | C | 2.53E-04 | | | | | ORF1ab | nonsynonymous_I2244T |
| 7029 | C | T | 2.53E-04 | | | | | ORF1ab | nonsynonymous_S2255F |
| 8885 | T | A | 2.53E-04 | | | | | ORF1ab | nonsynonymous_L2874I |
| 9733 | C | T | 2.53E-04 | | | | | ORF1ab | synonymous_F3156F |

| | | | | | | | | | |
|---|---|---|---|---|---|---|---|---|---|
| 10754 | G | A | 2.53E-04 | | | | | ORF1ab | nonsynonymous_A3497T |
| 10844 | A | G | 2.53E-04 | | | | | ORF1ab | nonsynonymous_M3527V |
| 11833 | C | T | 2.53E-04 | | | | | ORF1ab | synonymous_A3856A |
| 13138 | C | T | 2.53E-04 | | | | | ORF1ab | synonymous_I4291I |
| 13270 | T | C | 2.53E-04 | | | | | ORF1ab | synonymous_D4335D |
| 13740 | C | T | 2.53E-04 | | | | | ORF1ab | synonymous_D4492D |
| 13815 | A | C | 2.53E-04 | | | | | ORF1ab | synonymous_A4517A |
| 14126 | G | A | 2.53E-04 | | | | | ORF1ab | nonsynonymous_S4621N |
| 14133 | T | C | 2.53E-04 | | | | | ORF1ab | synonymous_V4623V |
| 14220 | C | T | 2.53E-04 | | | | | ORF1ab | synonymous_D4652D |
| 14235 | C | T | 2.53E-04 | | | | | ORF1ab | synonymous_Y4657Y |
| 14262 | C | T | 2.53E-04 | | | | | ORF1ab | synonymous_D4666D |
| 14290 | G | T | 2.53E-04 | | | | | ORF1ab | nonsynonymous_D4676Y |
| 14425 | C | A | 2.53E-04 | | | | | ORF1ab | nonsynonymous_L4721I |
| 14792 | G | T | 2.53E-04 | | | | | ORF1ab | nonsynonymous_S4843I |
| 15249 | A | G | 2.53E-04 | | | | | ORF1ab | synonymous_K4995K |
| 15368 | C | T | 2.53E-04 | | | | | ORF1ab | nonsynonymous_T5035I |
| 15480 | C | T | 2.53E-04 | | | | | ORF1ab | synonymous_T5072T |
| 15647 | A | G | 2.53E-04 | | | | | ORF1ab | nonsynonymous_D5128G |
| 15906 | G | A | 2.53E-04 | T | 1.26E-04 | | | ORF1ab | synonymous_Q5214Q; nonsynonymous_Q5214H |
| 15928 | C | T | 2.53E-04 | | | | | ORF1ab | nonsynonymous_P5222S |
| 15982 | G | T | 2.53E-04 | | | | | ORF1ab | nonsynonymous_V5240L |
| 16045 | C | T | 2.53E-04 | | | | | ORF1ab | nonsynonymous_L5261F |
| 16396 | G | T | 2.53E-04 | | | | | ORF1ab | nonsynonymous_G5378C |
| 16858 | G | T | 2.53E-04 | A | 6.32E-05 | | | ORF1ab | nonsynonymous_A5532S; nonsynonymous_A5532T |
| 16915 | C | T | 2.53E-04 | | | | | ORF1ab | synonymous_L5551L |
| 17118 | C | T | 2.53E-04 | | | | | ORF1ab | synonymous_G5618G |
| 17211 | G | T | 2.53E-04 | | | | | ORF1ab | nonsynonymous_L5649F |

| | | | | | | | | | |
|---|---|---|---|---|---|---|---|---|---|
| 17440 | C | T | 2.53E-04 | | | | | ORF1ab | nonsynonymous_P5726S |
| 17733 | C | T | 2.53E-04 | | | | | ORF1ab | synonymous_F5823F |
| 17825 | C | T | 2.53E-04 | | | | | ORF1ab | nonsynonymous_T5854I |
| 17976 | C | T | 2.53E-04 | | | | | ORF1ab | synonymous_D5904D |
| 18312 | C | T | 2.53E-04 | | | | | ORF1ab | synonymous_V6016V |
| 18512 | C | T | 2.53E-04 | | | | | ORF1ab | nonsynonymous_P6083L |
| 18513 | T | C | 2.53E-04 | | | | | ORF1ab | synonymous_P6083P |
| 18630 | T | C | 2.53E-04 | | | | | ORF1ab | synonymous_Y6122Y |
| 18831 | T | C | 2.53E-04 | | | | | ORF1ab | synonymous_H6189H |
| 18972 | G | T | 2.53E-04 | | | | | ORF1ab | nonsynonymous_K6236N |
| 19032 | C | T | 2.53E-04 | | | | | ORF1ab | synonymous_D6256D |
| 19610 | C | T | 2.53E-04 | | | | | ORF1ab | nonsynonymous_T6449I |
| 19656 | G | T | 2.53E-04 | | | | | ORF1ab | nonsynonymous_K6464N |
| 19719 | A | T | 2.53E-04 | | | | | ORF1ab | synonymous_T6485T |
| 19763 | C | T | 2.53E-04 | A | 6.32E-05 | | | ORF1ab | nonsynonymous_T6500I; nonsynonymous_T6500K |
| 19847 | G | A | 2.53E-04 | | | | | ORF1ab | nonsynonymous_G6528D |
| 20647 | C | T | 2.53E-04 | | | | | ORF1ab | nonsynonymous_P6795S |
| 20766 | A | G | 2.53E-04 | | | | | ORF1ab | synonymous_L6834L |
| 21800 | G | T | 2.53E-04 | A | 6.32E-05 | | | S | nonsynonymous_D80Y; nonsynonymous_D80N |
| 21867 | G | T | 2.53E-04 | | | | | S | nonsynonymous_R102I |
| 21920 | G | A | 2.53E-04 | T | 6.32E-05 | | | S | nonsynonymous_V120I; nonsynonymous_V120F |
| 21952 | C | T | 2.53E-04 | G | 6.32E-05 | | | S | synonymous_V130V; synonymous_V130V |
| 23423 | C | T | 2.53E-04 | | | | | S | nonsynonymous_P621S |
| 23426 | G | T | 2.53E-04 | A | 1.90E-04 | C | 1.26E-04 | S | nonsynonymous_V622F; nonsynonymous_V622I; nonsynonymous_V622L |

| | | | | | | | | | |
|---|---|---|---|---|---|---|---|---|---|
| 23430 | C | T | 2.53E-04 | | | | | S | nonsynonymous_A623V |
| 23443 | T | C | 2.53E-04 | | | | | S | synonymous_D627D |
| 23501 | G | T | 2.53E-04 | | | | | S | nonsynonymous_A647S |
| 23524 | A | C | 2.53E-04 | | | | | S | nonsynonymous_E654D |
| 23572 | A | G | 2.53E-04 | | | | | S | nonsynonymous_I670M |
| 23673 | C | T | 2.53E-04 | | | | | S | nonsynonymous_S704L |
| 23975 | A | G | 2.53E-04 | | | | | S | nonsynonymous_I805V |
| 24026 | C | T | 2.53E-04 | | | | | S | nonsynonymous_L822F |
| 24070 | A | T | 2.53E-04 | | | | | S | nonsynonymous_Q836H |
| 25302 | G | T | 2.53E-04 | | | | | S | nonsynonymous_C1247F |
| 25317 | C | T | 2.53E-04 | | | | | S | nonsynonymous_S1252F |
| 25318 | C | T | 2.53E-04 | | | | | S | synonymous_S1252S |
| 25354 | G | T | 2.53E-04 | A | 6.32E-05 | | | S | synonymous_V1264V; synonymous_V1264V |
| 25452 | C | T | 2.53E-04 | | | | | ORF3a | synonymous_I20I |
| 25461 | T | C | 2.53E-04 | | | | | ORF3a | synonymous_A23A |
| 25473 | T | C | 2.53E-04 | | | | | ORF3a | synonymous_D27D |
| 25483 | G | A | 2.53E-04 | T | 6.32E-05 | | | ORF3a | nonsynonymous_A31T; nonsynonymous_A31S |
| 25537 | G | T | 2.53E-04 | | | | | ORF3a | nonsynonymous_G49C |
| 25980 | A | C | 2.53E-04 | | | | | ORF3a | synonymous_G196G |
| 26130 | T | C | 2.53E-04 | | | | | ORF3a | synonymous_I246I |
| 26176 | C | T | 2.53E-04 | | | | | ORF3a | nonsynonymous_P262S |
| 26204 | C | T | 2.53E-04 | | | | | ORF3a | nonsynonymous_T271I |
| 26262 | G | A | 2.53E-04 | | | | | E | synonymous_S6S |
| 26333 | C | T | 2.53E-04 | | | | | E | nonsynonymous_T30I |
| 26545 | T | C | 2.53E-04 | G | 6.32E-05 | | | M | nonsynonymous_I8T; nonsynonymous_I8S |
| 26551 | T | C | 2.53E-04 | | | | | M | nonsynonymous_V10A |
| 26713 | G | T | 2.53E-04 | | | | | M | nonsynonymous_C64F |

| | | | | | | | | | |
|---|---|---|---|---|---|---|---|---|---|
| 26763 | G | T | 2.53E-04 | | | | | M | nonsynonymous_A81S |
| 26849 | G | T | 2.53E-04 | C | 6.32E-05 | | | M | nonsynonymous_M109I; nonsynonymous_M109I |
| 26892 | C | T | 2.53E-04 | | | | | M | nonsynonymous_L124F |
| 27843 | T | C | 2.53E-04 | | | | | ORF7b | nonsynonymous_F30L |
| 28472 | C | T | 2.53E-04 | A | 6.32E-05 | | | N | nonsynonymous_P67S; nonsynonymous_P67T |
| 28480 | A | T | 2.53E-04 | | | | | N | synonymous_G69G |
| 28606 | C | T | 2.53E-04 | | | | | N | synonymous_Y111Y |
| 28621 | G | A | 2.53E-04 | | | | | N | synonymous_G116G |
| 28674 | C | T | 2.53E-04 | | | | | N | nonsynonymous_A134V |
| 28725 | C | T | 2.53E-04 | A | 6.32E-05 | | | N | nonsynonymous_P151L; nonsynonymous_P151H |
| 29009 | G | A | 2.53E-04 | T | 6.32E-05 | | | N | nonsynonymous_V246I; nonsynonymous_V246F |
| 29032 | G | A | 2.53E-04 | | | | | N | synonymous_E253E |
| 361 | A | G | 2.53E-04 | | | | | ORF1ab | synonymous_G32G |
| 364 | C | T | 2.53E-04 | | | | | ORF1ab | synonymous_D33D |
| 544 | A | G | 2.53E-04 | | | | | ORF1ab | synonymous_E93E |
| 6248 | A | C | 2.53E-04 | G | 6.32E-05 | | | ORF1ab | nonsynonymous_N1995H; nonsynonymous_N1995D |
| 6354 | C | T | 2.53E-04 | | | | | ORF1ab | nonsynonymous_S2030L |
| 6394 | T | C | 2.53E-04 | | | | | ORF1ab | synonymous_D2043D |
| 6443 | G | A | 2.53E-04 | | | | | ORF1ab | nonsynonymous_D2060N |
| 6445 | C | T | 2.53E-04 | | | | | ORF1ab | synonymous_D2060D |
| 7081 | C | T | 2.53E-04 | | | | | ORF1ab | synonymous_N2272N |
| 8137 | G | T | 2.53E-04 | | | | | ORF1ab | synonymous_V2624V |
| 1666 | T | C | 2.53E-04 | | | | | ORF1ab | synonymous_F467F |
| 2041 | T | C | 2.53E-04 | | | | | ORF1ab | synonymous_T592T |
| 3086 | T | C | 2.53E-04 | | | | | ORF1ab | nonsynonymous_F941L |

| | | | | | | | | | |
|---|---|---|---|---|---|---|---|---|---|
| 6703 | T | C | 2.53E-04 | | | | | ORF1ab | synonymous_L2146L |
| 6814 | T | C | 2.53E-04 | | | | | ORF1ab | synonymous_T2183T |
| 10105 | G | T | 2.53E-04 | | | | | ORF1ab | nonsynonymous_M3280I |
| 10456 | C | T | 2.53E-04 | | | | | ORF1ab | synonymous_F3397F |
| 11681 | C | T | 2.53E-04 | | | | | ORF1ab | synonymous_L3806L |
| 11758 | C | T | 2.53E-04 | | | | | ORF1ab | synonymous_P3831P |
| 12364 | T | A | 2.53E-04 | | | | | ORF1ab | synonymous_L4033L |
| 13115 | C | T | 2.53E-04 | | | | | ORF1ab | synonymous_L4284L |
| 1648 | C | T | 2.53E-04 | | | | | ORF1ab | synonymous_I461I |
| 1734 | C | T | 2.53E-04 | | | | | ORF1ab | nonsynonymous_T490I |
| 2142 | C | T | 2.53E-04 | | | | | ORF1ab | nonsynonymous_P626L |
| 2269 | A | T | 2.53E-04 | | | | | ORF1ab | synonymous_A668A |
| 2945 | G | A | 2.53E-04 | T | 1.26E-04 | | | ORF1ab | nonsynonymous_G894S; nonsynonymous_G894C |
| 3040 | C | T | 2.53E-04 | | | | | ORF1ab | synonymous_Y925Y |
| 3099 | C | T | 2.53E-04 | | | | | ORF1ab | nonsynonymous_T945I |
| 3425 | C | T | 2.53E-04 | | | | | ORF1ab | nonsynonymous_P1054S |
| 3717 | C | T | 2.53E-04 | | | | | ORF1ab | nonsynonymous_A1151V |
| 4015 | A | T | 2.53E-04 | G | 6.32E-05 | | | ORF1ab | synonymous_T1250T; synonymous_T1250T |
| 4456 | C | T | 2.53E-04 | | | | | ORF1ab | synonymous_A1397A |
| 4504 | G | T | 2.53E-04 | A | 1.26E-04 | | | ORF1ab | nonsynonymous_E1413D; synonymous_E1413E |
| 4655 | C | T | 2.53E-04 | | | | | ORF1ab | nonsynonymous_R1464W |
| 598 | G | T | 2.53E-04 | | | | | ORF1ab | synonymous_V111V |
| 600 | G | A | 2.53E-04 | | | | | ORF1ab | nonsynonymous_G112D |
| 6627 | C | T | 2.53E-04 | | | | | ORF1ab | nonsynonymous_T2121I |
| 6762 | C | T | 2.53E-04 | | | | | ORF1ab | nonsynonymous_T2166I |
| 6781 | C | T | 2.53E-04 | | | | | ORF1ab | synonymous_F2172F |
| 7272 | G | A | 2.53E-04 | | | | | ORF1ab | nonsynonymous_R2336K |

| | | | | | | | | | |
|---|---|---|---|---|---|---|---|---|---|
| 7321 | C | T | 2.53E-04 | | | | | ORF1ab | synonymous_S2352S |
| 7736 | G | A | 2.53E-04 | | | | | ORF1ab | nonsynonymous_V2491I |
| 7819 | C | T | 2.53E-04 | | | | | ORF1ab | synonymous_L2518L |
| 7843 | C | T | 2.53E-04 | | | | | ORF1ab | synonymous_N2526N |
| 7876 | T | A | 2.53E-04 | C | 6.32E-05 | | | ORF1ab | synonymous_P2537P; synonymous_P2537P |
| 8031 | A | G | 2.53E-04 | | | | | ORF1ab | nonsynonymous_K2589R |
| 8074 | C | T | 2.53E-04 | | | | | ORF1ab | synonymous_N2603N |
| 8139 | C | T | 2.53E-04 | | | | | ORF1ab | nonsynonymous_S2625F |
| 8956 | C | T | 2.53E-04 | | | | | ORF1ab | synonymous_Y2897Y |
| 9204 | A | G | 2.53E-04 | | | | | ORF1ab | nonsynonymous_D2980G |
| 10168 | T | C | 2.53E-04 | | | | | ORF1ab | synonymous_C3301C |
| 10478 | A | C | 2.53E-04 | | | | | ORF1ab | nonsynonymous_N3405H |
| 10480 | T | C | 2.53E-04 | | | | | ORF1ab | synonymous_N3405N |
| 11935 | A | C | 2.53E-04 | | | | | ORF1ab | nonsynonymous_Q3890H |
| 1294 | T | C | 2.53E-04 | | | | | ORF1ab | synonymous_F343F |
| 1623 | T | C | 2.53E-04 | | | | | ORF1ab | nonsynonymous_I453T |
| 9070 | T | C | 2.53E-04 | | | | | ORF1ab | synonymous_D2935D |
| 946 | T | C | 2.53E-04 | A | 6.32E-05 | | | ORF1ab | synonymous_G227G; synonymous_G227G |
| 9931 | T | C | 2.53E-04 | | | | | ORF1ab | synonymous_D3222D |
| 10329 | A | G | 2.53E-04 | | | | | ORF1ab | nonsynonymous_D3355G |
| 10479 | A | T | 2.53E-04 | G | 1.26E-04 | | | ORF1ab | nonsynonymous_N3405I; nonsynonymous_N3405S |
| 10579 | A | G | 2.53E-04 | | | | | ORF1ab | synonymous_T3438T |
| 10617 | G | A | 2.53E-04 | | | | | ORF1ab | nonsynonymous_R3451K |
| 10623 | C | T | 2.53E-04 | | | | | ORF1ab | nonsynonymous_T3453I |
| 11596 | G | T | 2.53E-04 | | | | | ORF1ab | nonsynonymous_Q3777H |
| 11606 | C | T | 2.53E-04 | | | | | ORF1ab | synonymous_L3781L |
| 11752 | C | T | 2.53E-04 | | | | | ORF1ab | synonymous_L3829L |

| | | | | | | | | | |
|---|---|---|---|---|---|---|---|---|---|
| 11824 | C | T | 2.53E-04 | | | | | ORF1ab | synonymous_I3853I |
| 11919 | C | T | 2.53E-04 | | | | | ORF1ab | nonsynonymous_S3885F |
| 1204 | C | T | 2.53E-04 | | | | | ORF1ab | synonymous_N313N |
| 12242 | C | T | 2.53E-04 | | | | | ORF1ab | nonsynonymous_R3993C |
| 12439 | C | T | 2.53E-04 | A | 6.32E-05 | | | ORF1ab | synonymous_P4058P; synonymous_P4058P |
| 12459 | C | T | 2.53E-04 | | | | | ORF1ab | nonsynonymous_T4065I |
| 1267 | C | T | 2.53E-04 | | | | | ORF1ab | synonymous_G334G |
| 12747 | C | T | 2.53E-04 | | | | | ORF1ab | nonsynonymous_T4161I |
| 12754 | C | T | 2.53E-04 | | | | | ORF1ab | synonymous_C4163C |
| 12796 | A | C | 2.53E-04 | T | 6.32E-05 | | | ORF1ab | synonymous_G4177G; synonymous_G4177G |
| 12797 | G | A | 2.53E-04 | | | | | ORF1ab | nonsynonymous_G4178S |
| 12820 | A | G | 2.53E-04 | | | | | ORF1ab | synonymous_L4185L |
| 12854 | C | T | 2.53E-04 | | | | | ORF1ab | nonsynonymous_P4197S |
| 12896 | C | T | 2.53E-04 | | | | | ORF1ab | nonsynonymous_P4211S |
| 12986 | A | G | 2.53E-04 | | | | | ORF1ab | nonsynonymous_M4241V |
| 1342 | C | T | 2.53E-04 | | | | | ORF1ab | synonymous_Y359Y |
| 1380 | C | T | 2.53E-04 | | | | | ORF1ab | nonsynonymous_A372V |
| 1385 | C | T | 2.53E-04 | | | | | ORF1ab | nonsynonymous_H374Y |
| 1392 | C | T | 2.53E-04 | | | | | ORF1ab | nonsynonymous_S376L |
| 1419 | C | T | 2.53E-04 | | | | | ORF1ab | nonsynonymous_A385V |
| 1441 | C | T | 2.53E-04 | | | | | ORF1ab | synonymous_G392G |
| 2306 | C | T | 2.53E-04 | | | | | ORF1ab | nonsynonymous_L681F |
| 2528 | G | A | 2.53E-04 | | | | | ORF1ab | nonsynonymous_E755K |
| 3251 | G | T | 2.53E-04 | A | 6.32E-05 | | | ORF1ab | nonsynonymous_D996Y; nonsynonymous_D996N |
| 3634 | C | T | 2.53E-04 | | | | | ORF1ab | synonymous_N1123N |
| 3792 | C | T | 2.53E-04 | | | | | ORF1ab | nonsynonymous_A1176V |
| 4109 | G | A | 2.53E-04 | | | | | ORF1ab | nonsynonymous_D1282N |

| | | | | | | | | | |
|---|---|---|---|---|---|---|---|---|---|
| 4207 | G | T | 2.53E-04 | A | 1.90E-04 | | | ORF1ab | synonymous_A1314A; synonymous_A1314A |
| 4246 | C | T | 2.53E-04 | | | | | ORF1ab | synonymous_T1327T |
| 4423 | C | T | 2.53E-04 | | | | | ORF1ab | synonymous_R1386R |
| 4633 | C | T | 2.53E-04 | | | | | ORF1ab | synonymous_G1456G |
| 4780 | C | T | 2.53E-04 | | | | | ORF1ab | synonymous_I1505I |
| 4866 | G | T | 2.53E-04 | | | | | ORF1ab | nonsynonymous_S1534I |
| 5622 | C | T | 2.53E-04 | | | | | ORF1ab | nonsynonymous_P1786L |
| 5694 | C | T | 2.53E-04 | A | 6.32E-05 | | | ORF1ab | nonsynonymous_P1810L; nonsynonymous_P1810Q |
| 5806 | C | T | 2.53E-04 | | | | | ORF1ab | synonymous_C1847C |
| 613 | G | A | 2.53E-04 | | | | | ORF1ab | synonymous_V116V |
| 6198 | C | T | 2.53E-04 | A | 6.32E-05 | | | ORF1ab | nonsynonymous_S1978F; nonsynonymous_S1978Y |
| 6228 | A | G | 2.53E-04 | | | | | ORF1ab | nonsynonymous_K1988R |
| 679 | C | T | 2.53E-04 | | | | | ORF1ab | synonymous_A138A |
| 7165 | C | T | 2.53E-04 | A | 6.32E-05 | | | ORF1ab | synonymous_T2300T; synonymous_T2300T |
| 7390 | C | T | 2.53E-04 | | | | | ORF1ab | synonymous_A2375A |
| 739 | A | G | 2.53E-04 | | | | | ORF1ab | synonymous_Q158Q |
| 823 | C | T | 2.53E-04 | | | | | ORF1ab | synonymous_V186V |
| 8326 | C | T | 2.53E-04 | | | | | ORF1ab | synonymous_D2687D |
| 8994 | C | T | 2.53E-04 | | | | | ORF1ab | nonsynonymous_T2910I |
| 900 | C | T | 2.53E-04 | | | | | ORF1ab | nonsynonymous_S212L |
| 9163 | C | T | 2.53E-04 | | | | | ORF1ab | synonymous_N2966N |
| 936 | C | T | 2.53E-04 | | | | | ORF1ab | nonsynonymous_T224I |
| 9520 | C | T | 2.53E-04 | A | 6.32E-05 | | | ORF1ab | synonymous_F3085F; nonsynonymous_F3085L |
| 957 | G | A | 2.53E-04 | | | | | ORF1ab | nonsynonymous_C231Y |
| 9749 | A | G | 2.53E-04 | | | | | ORF1ab | nonsynonymous_K3162E |

| | | | | | | | | | |
|---|---|---|---|---|---|---|---|---|---|
| 9778 | C | T | 2.53E-04 | | | | | ORF1ab | synonymous_S3171S |
| 29766 | G | A | 2.27E-04 | T | 7.58E-05 | | | three_prime_UTR | |
| 45 | G | T | 2.07E-04 | C | 1.03E-04 | | | five_prime_UTR | |
| 29813 | A | C | 2.04E-04 | G | 6.81E-05 | | | three_prime_UTR | |
| 29805 | A | G | 2.04E-04 | | | | | three_prime_UTR | |
| 29786 | G | C | 2.02E-04 | A | 6.72E-05 | | | three_prime_UTR | |
| 54 | T | C | 2.02E-04 | A | 1.01E-04 | A | 1.01E-04 | five_prime_UTR | |
| 29782 | A | C | 2.01E-04 | G | 6.71E-05 | | | three_prime_UTR | |
| 56 | G | T | 1.99E-04 | C | 6.64E-05 | | | five_prime_UTR | |
| 29771 | A | G | 1.97E-04 | C | 6.55E-05 | | | three_prime_UTR | |
| 85 | T | C | 1.95E-04 | | | | | five_prime_UTR | |
| 105 | G | T | 1.93E-04 | A | 6.44E-05 | | | five_prime_UTR | |
| 110 | C | T | 1.93E-04 | | | | | five_prime_UTR | |
| 117 | C | T | 1.93E-04 | | | | | five_prime_UTR | |
| 120 | T | C | 1.93E-04 | | | | | five_prime_UTR | |
| 124 | G | A | 1.93E-04 | C | 6.42E-05 | | | five_prime_UTR | |
| 140 | C | T | 1.92E-04 | | | | | five_prime_UTR | |
| 29754 | C | T | 1.92E-04 | A | 6.39E-05 | A | 6.39E-05 | three_prime_UTR | |
| 151 | C | T | 1.92E-04 | | | | | five_prime_UTR | |
| 157 | C | T | 1.92E-04 | G | 6.39E-05 | | | five_prime_UTR | |
| 29752 | A | G | 1.92E-04 | | | | | three_prime_UTR | |
| 161 | A | T | 1.92E-04 | G | 6.39E-05 | | | five_prime_UTR | |
| 29747 | G | A | 1.92E-04 | A | 1.92E-04 | C | 1.28E-04 | three_prime_UTR | |
| 168 | T | C | 1.92E-04 | A | 6.39E-05 | | | five_prime_UTR | |
| 29741 | C | T | 1.92E-04 | G | 6.39E-05 | | | three_prime_UTR | |
| 174 | G | T | 1.92E-04 | A | 1.28E-04 | | | five_prime_UTR | |
| 214 | C | T | 1.92E-04 | | | | | five_prime_UTR | |
| 216 | G | A | 1.92E-04 | | | | | five_prime_UTR | |
| 217 | C | A | 1.92E-04 | T | 6.38E-05 | | | five_prime_UTR | |
| 177 | T | G | 1.92E-04 | | | | | five_prime_UTR | |

| | | | | | | | | | |
|---|---|---|---|---|---|---|---|---|---|
| 182 | T | C | 1.91E-04 | | | | | five_prime_UTR | |
| 221 | T | A | 1.91E-04 | C | 1.28E-04 | G | 6.38E-05 | five_prime_UTR | |
| 213 | G | C | 1.91E-04 | | | | | five_prime_UTR | |
| 233 | C | A | 1.91E-04 | T | 6.38E-05 | | | five_prime_UTR | |
| 254 | C | T | 1.91E-04 | | | | | five_prime_UTR | |
| 29690 | G | T | 1.91E-04 | | | | | three_prime_UTR | |
| 29680 | T | C | 1.91E-04 | | | | | three_prime_UTR | |
| 29675 | C | T | 1.91E-04 | A | 6.35E-05 | | | three_prime_UTR | |
| 690 | C | T | 1.90E-04 | | | | | ORF1ab | nonsynonymous_S142L |
| 19554 | C | T | 1.90E-04 | | | | | ORF1ab | synonymous_G6430G |
| 27688 | C | T | 1.90E-04 | | | | | ORF7a | nonsynonymous_P99S |
| 27595 | C | T | 1.90E-04 | | | | | ORF7a | nonsynonymous_P68S |
| 27580 | T | C | 1.90E-04 | | | | | ORF7a | nonsynonymous_F63L |
| 29666 | C | T | 1.90E-04 | G | 6.33E-05 | | | ORF10 | nonsynonymous_L37F; nonsynonymous_L37V |
| 29670 | C | T | 1.90E-04 | G | 6.33E-05 | | | ORF10 | nonsynonymous_T38I; nonsynonymous_T38R |
| 27593 | G | T | 1.90E-04 | | | | | ORF7a | nonsynonymous_C67F |
| 27588 | T | A | 1.90E-04 | | | | | ORF7a | nonsynonymous_F65L |
| 27618 | T | C | 1.90E-04 | | | | | ORF7a | synonymous_Y75Y |
| 27710 | C | T | 1.90E-04 | | | | | ORF7a | nonsynonymous_A106V |
| 19374 | C | T | 1.90E-04 | | | | | ORF1ab | synonymous_F6370F |
| 27670 | G | T | 1.90E-04 | | | | | ORF7a | nonsynonymous_V93F |
| 27722 | T | C | 1.90E-04 | | | | | ORF7a | nonsynonymous_I110T |
| 506 | C | T | 1.90E-04 | | | | | ORF1ab | nonsynonymous_H81Y |
| 529 | G | T | 1.90E-04 | A | 6.33E-05 | | | ORF1ab | synonymous_L88L; synonymous_L88L |
| 27769 | C | T | 1.90E-04 | | | | | ORF7b | nonsynonymous_S5L |
| 19402 | C | T | 1.90E-04 | | | | | ORF1ab | nonsynonymous_H6380Y |
| 27546 | T | C | 1.90E-04 | | | | | ORF7a | synonymous_D51D |

| | | | | | | | | | |
|---|---|---|---|---|---|---|---|---|---|
| 27673 | C | T | 1.90E-04 | A | 6.33E-05 | | | ORF7a | stopgain_Q94X; nonsynonymous_Q94K |
| 27765 | C | T | 1.90E-04 | | | | | ORF7b | nonsynonymous_L4F |
| 27806 | G | T | 1.90E-04 | A | 6.33E-05 | | | ORF7b | synonymous_L17L; synonymous_L17L |
| 27807 | C | T | 1.90E-04 | | | | | ORF7b | synonymous_L18L |
| 27970 | C | T | 1.90E-04 | | | | | ORF8 | nonsynonymous_T26I |
| 28209 | G | T | 1.90E-04 | C | 6.33E-05 | | | ORF8 | stopgain_E106X; nonsynonymous_E106Q |
| 19512 | C | T | 1.90E-04 | | | | | ORF1ab | synonymous_Y6416Y |
| 22349 | G | T | 1.90E-04 | | | | | S | nonsynonymous_A263S |
| 27948 | G | T | 1.90E-04 | | | | | ORF8 | stopgain_E19X |
| 27962 | G | T | 1.90E-04 | | | | | ORF8 | nonsynonymous_Q23H |
| 27981 | C | T | 1.90E-04 | | | | | ORF8 | nonsynonymous_P30S |
| 28003 | G | T | 1.90E-04 | | | | | ORF8 | nonsynonymous_C37F |
| 28041 | G | T | 1.90E-04 | | | | | ORF8 | stopgain_G50X |
| 28043 | A | G | 1.90E-04 | | | | | ORF8 | synonymous_G50G |
| 28048 | G | T | 1.90E-04 | C | 6.33E-05 | | | ORF8 | nonsynonymous_R52I; nonsynonymous_R52T |
| 28079 | G | T | 1.90E-04 | | | | | ORF8 | synonymous_V62V |
| 28109 | G | T | 1.90E-04 | | | | | ORF8 | nonsynonymous_Q72H |
| 28110 | T | C | 1.90E-04 | | | | | ORF8 | nonsynonymous_Y73H |
| 28167 | G | A | 1.90E-04 | | | | | ORF8 | nonsynonymous_E92K |
| 22404 | A | T | 1.90E-04 | | | | | S | nonsynonymous_E281V |
| 27881 | C | T | 1.90E-04 | | | | | ORF7b | synonymous_H42H |
| 27883 | C | T | 1.90E-04 | | | | | ORF7b | nonsynonymous_A43V |
| 21372 | G | T | 1.90E-04 | | | | | ORF1ab | nonsynonymous_Q7036H |
| 10981 | G | T | 1.90E-04 | | | | | ORF1ab | synonymous_V3572V |
| 20428 | C | T | 1.90E-04 | | | | | ORF1ab | nonsynonymous_P6722S |
| 2607 | C | T | 1.90E-04 | | | | | ORF1ab | nonsynonymous_T781I |

| | | | | | | | | | |
|---|---|---|---|---|---|---|---|---|---|
| 29568 | T | C | 1.90E-04 | | | | | ORF10 | nonsynonymous_I4T |
| 5011 | A | C | 1.90E-04 | | | | | ORF1ab | nonsynonymous_Q1582H |
| 13572 | T | C | 1.90E-04 | | | | | ORF1ab | synonymous_G4436G |
| 21158 | T | C | 1.90E-04 | | | | | ORF1ab | nonsynonymous_V6965A |
| 21221 | C | T | 1.90E-04 | | | | | ORF1ab | nonsynonymous_A6986V |
| 21302 | C | T | 1.90E-04 | | | | | ORF1ab | nonsynonymous_P7013L |
| 21364 | C | T | 1.90E-04 | | | | | ORF1ab | nonsynonymous_P7034S |
| 21365 | C | T | 1.90E-04 | | | | | ORF1ab | nonsynonymous_P7034L |
| 22326 | C | T | 1.90E-04 | | | | | S | nonsynonymous_S255F |
| 22484 | G | T | 1.90E-04 | | | | | S | nonsynonymous_V308L |
| 2724 | C | T | 1.90E-04 | | | | | ORF1ab | nonsynonymous_P820L |
| 2780 | G | T | 1.90E-04 | | | | | ORF1ab | nonsynonymous_V839L |
| 2809 | G | T | 1.90E-04 | | | | | ORF1ab | nonsynonymous_R848S |
| 29445 | C | T | 1.90E-04 | | | | | N | nonsynonymous_T391I |
| 13408 | T | C | 1.90E-04 | | | | | ORF1ab | synonymous_C4381C |
| 20438 | G | T | 1.90E-04 | | | | | ORF1ab | nonsynonymous_S6725I |
| 20480 | C | T | 1.90E-04 | | | | | ORF1ab | nonsynonymous_S6739L |
| 2644 | C | T | 1.90E-04 | | | | | ORF1ab | synonymous_I793I |
| 2725 | A | G | 1.90E-04 | | | | | ORF1ab | synonymous_P820P |
| 2731 | G | A | 1.90E-04 | | | | | ORF1ab | synonymous_K822K |
| 2790 | C | T | 1.90E-04 | | | | | ORF1ab | nonsynonymous_T842I |
| 2842 | C | A | 1.90E-04 | T | 1.27E-04 | | | ORF1ab | synonymous_A859A; synonymous_A859A |
| 2849 | G | T | 1.90E-04 | | | | | ORF1ab | nonsynonymous_V862F |
| 29364 | C | T | 1.90E-04 | G | 6.33E-05 | | | N | nonsynonymous_P364L; nonsynonymous_P364R |
| 29409 | C | T | 1.90E-04 | | | | | N | nonsynonymous_T379I |
| 29415 | C | T | 1.90E-04 | | | | | N | nonsynonymous_A381V |
| 29459 | C | T | 1.90E-04 | | | | | N | nonsynonymous_P396S |
| 29628 | G | T | 1.90E-04 | | | | | ORF10 | nonsynonymous_R24L |

| | | | | | | | | | |
|---|---|---|---|---|---|---|---|---|---|
| 19679 | G | T | 1.90E-04 | | | | | ORF1ab | nonsynonymous_G6472V |
| 20234 | C | T | 1.90E-04 | | | | | ORF1ab | nonsynonymous_P6657L |
| 20315 | T | A | 1.90E-04 | | | | | ORF1ab | nonsynonymous_F6684Y |
| 29586 | C | T | 1.90E-04 | | | | | ORF10 | nonsynonymous_P10L |
| 13487 | C | T | 1.90E-04 | | | | | ORF1ab | nonsynonymous_A4408V |
| 21638 | C | T | 1.90E-04 | | | | | S | nonsynonymous_P26S |
| 23126 | G | T | 1.90E-04 | | | | | S | nonsynonymous_A522S |
| 25660 | G | T | 1.90E-04 | | | | | ORF3a | nonsynonymous_V90F |
| 25721 | C | T | 1.90E-04 | G | 6.33E-05 | | | ORF3a | nonsynonymous_A110V; nonsynonymous_A110G |
| 25744 | A | G | 1.90E-04 | | | | | ORF3a | nonsynonymous_I118V |
| 25767 | G | T | 1.90E-04 | C | 1.27E-04 | | | ORF3a | nonsynonymous_M125I; nonsynonymous_M125I |
| 13381 | C | T | 1.90E-04 | | | | | ORF1ab | synonymous_V4372V |
| 13499 | C | T | 1.90E-04 | | | | | ORF1ab | nonsynonymous_T4412I |
| 13514 | G | A | 1.90E-04 | | | | | ORF1ab | nonsynonymous_G4417D |
| 25665 | C | T | 1.90E-04 | | | | | ORF3a | synonymous_Y91Y |
| 25672 | C | T | 1.90E-04 | A | 6.32E-05 | | | ORF3a | nonsynonymous_L94F; nonsynonymous_L94I |
| 25784 | G | T | 1.90E-04 | C | 6.32E-05 | | | ORF3a | nonsynonymous_W131L; nonsynonymous_W131S |
| 25810 | C | G | 1.90E-04 | T | 1.26E-04 | | | ORF3a | nonsynonymous_L140V; nonsynonymous_L140F |
| 25842 | T | C | 1.90E-04 | | | | | ORF3a | synonymous_H150H |
| 25919 | C | T | 1.90E-04 | | | | | ORF3a | nonsynonymous_T176I |
| 15869 | A | T | 1.90E-04 | | | | | ORF1ab | nonsynonymous_H5202L |
| 1589 | G | A | 1.90E-04 | | | | | ORF1ab | nonsynonymous_G442S |
| 20047 | A | G | 1.90E-04 | | | | | ORF1ab | nonsynonymous_I6595V |
| 20060 | G | T | 1.90E-04 | | | | | ORF1ab | nonsynonymous_S6599I |
| 21458 | T | C | 1.90E-04 | | | | | ORF1ab | nonsynonymous_I7065T |

| | | | | | | | | | |
|---|---|---|---|---|---|---|---|---|---|
| 21462 | T | C | 1.90E-04 | | | | | ORF1ab | synonymous_N7066N |
| 21588 | C | T | 1.90E-04 | | | | | S | nonsynonymous_P9L |
| 21778 | G | A | 1.90E-04 | | | | | S | synonymous_G72G |
| 23029 | C | T | 1.90E-04 | | | | | S | synonymous_Y489Y |
| 27207 | T | C | 1.90E-04 | | | | | ORF6 | synonymous_F2F |
| 28893 | C | T | 1.90E-04 | | | | | N | nonsynonymous_P207L |
| 29250 | C | T | 1.90E-04 | | | | | N | nonsynonymous_P326L |
| 5211 | C | T | 1.90E-04 | | | | | ORF1ab | nonsynonymous_A1649V |
| 8660 | C | T | 1.90E-04 | | | | | ORF1ab | nonsynonymous_H2799Y |
| 10202 | C | T | 1.90E-04 | | | | | ORF1ab | nonsynonymous_L3313F |
| 14645 | C | T | 1.90E-04 | | | | | ORF1ab | nonsynonymous_T4794I |
| 16117 | G | T | 1.90E-04 | | | | | ORF1ab | nonsynonymous_D5285Y |
| 20024 | G | A | 1.90E-04 | T | 1.26E-04 | | | ORF1ab | nonsynonymous_R6587K; nonsynonymous_R6587I |
| 20132 | C | T | 1.90E-04 | | | | | ORF1ab | nonsynonymous_A6623V |
| 20139 | A | G | 1.90E-04 | | | | | ORF1ab | synonymous_K6625K |
| 22964 | A | G | 1.90E-04 | T | 6.32E-05 | | | S | nonsynonymous_I468V; nonsynonymous_I468F |
| 23047 | T | C | 1.90E-04 | | | | | S | synonymous_Y495Y |
| 23589 | C | T | 1.90E-04 | | | | | S | nonsynonymous_T676I |
| 24844 | C | T | 1.90E-04 | | | | | S | synonymous_V1094V |
| 24959 | G | T | 1.90E-04 | | | | | S | nonsynonymous_V1133F |
| 24964 | C | T | 1.90E-04 | | | | | S | synonymous_N1134N |
| 25002 | C | T | 1.90E-04 | | | | | S | nonsynonymous_S1147L |
| 25269 | G | T | 1.90E-04 | | | | | S | nonsynonymous_C1236F |
| 25272 | T | C | 1.90E-04 | | | | | S | nonsynonymous_M1237T |
| 25290 | G | T | 1.90E-04 | A | 6.32E-05 | | | S | nonsynonymous_C1243F; nonsynonymous_C1243Y |
| 28770 | C | T | 1.90E-04 | | | | | N | nonsynonymous_T166I |

| | | | | | | | | | |
|---|---|---|---|---|---|---|---|---|---|
| 28798 | G | A | 1.90E-04 | C | 6.32E-05 | | | N | synonymous_G175G; synonymous_G175G |
| 28968 | G | C | 1.90E-04 | T | 6.32E-05 | | | N | nonsynonymous_S232T; nonsynonymous_S232I |
| 29162 | G | T | 1.90E-04 | | | | | N | nonsynonymous_D297Y |
| 29229 | G | A | 1.90E-04 | | | | | N | nonsynonymous_R319H |
| 29247 | C | T | 1.90E-04 | | | | | N | nonsynonymous_T325I |
| 29259 | C | T | 1.90E-04 | | | | | N | nonsynonymous_T329M |
| 29266 | G | T | 1.90E-04 | | | | | N | nonsynonymous_L331F |
| 29288 | T | C | 1.90E-04 | | | | | N | synonymous_L339L |
| 4999 | C | T | 1.90E-04 | | | | | ORF1ab | synonymous_N1578N |
| 6638 | C | T | 1.90E-04 | | | | | ORF1ab | nonsynonymous_H2125Y |
| 8683 | C | T | 1.90E-04 | A | 6.32E-05 | | | ORF1ab | synonymous_I2806I; synonymous_I2806I |
| 14322 | C | T | 1.90E-04 | | | | | ORF1ab | synonymous_Y4686Y |
| 21405 | A | T | 1.90E-04 | G | 6.32E-05 | | | ORF1ab | nonsynonymous_K7047N; synonymous_K7047K |
| 22281 | C | T | 1.90E-04 | | | | | S | nonsynonymous_T240I |
| 23586 | A | G | 1.90E-04 | | | | | S | nonsynonymous_Q675R |
| 23997 | C | T | 1.90E-04 | | | | | S | nonsynonymous_P812L |
| 24620 | G | T | 1.90E-04 | | | | | S | nonsynonymous_A1020S |
| 25156 | C | T | 1.90E-04 | | | | | S | synonymous_I1198I |
| 25416 | C | T | 1.90E-04 | | | | | ORF3a | synonymous_F8F |
| 25556 | T | G | 1.90E-04 | | | | | ORF3a | nonsynonymous_V55G |
| 25617 | G | T | 1.90E-04 | A | 6.32E-05 | | | ORF3a | nonsynonymous_K75N; synonymous_K75K |
| 26110 | C | T | 1.90E-04 | | | | | ORF3a | nonsynonymous_P240S |
| 26439 | G | T | 1.90E-04 | | | | | E | synonymous_L65L |
| 26452 | G | T | 1.90E-04 | | | | | E | nonsynonymous_V70F |

| | | | | | | | | | |
|---|---|---|---|---|---|---|---|---|---|
| 27047 | G | A | 1.90E-04 | T | 6.32E-05 | | | M | synonymous_T175T; synonymous_T175T |
| 27276 | C | T | 1.90E-04 | | | | | ORF6 | synonymous_S25S |
| 28250 | T | G | 1.90E-04 | | | | | ORF8 | nonsynonymous_D119E |
| 28251 | T | G | 1.90E-04 | C | 6.32E-05 | | | ORF8 | nonsynonymous_F120V; nonsynonymous_F120L |
| 28697 | C | T | 1.90E-04 | | | | | N | nonsynonymous_P142S |
| 28775 | C | T | 1.90E-04 | | | | | N | nonsynonymous_P168S |
| 28792 | A | T | 1.90E-04 | | | | | N | synonymous_A173A |
| 28808 | G | T | 1.90E-04 | A | 6.32E-05 | | | N | nonsynonymous_G179C; nonsynonymous_G179S |
| 28861 | T | C | 1.90E-04 | | | | | N | synonymous_N196N |
| 28873 | C | T | 1.90E-04 | | | | | N | synonymous_G200G |
| 28915 | C | T | 1.90E-04 | | | | | N | synonymous_G214G |
| 28930 | T | C | 1.90E-04 | | | | | N | synonymous_L219L |
| 28957 | C | T | 1.90E-04 | | | | | N | synonymous_N228N |
| 28960 | G | T | 1.90E-04 | | | | | N | nonsynonymous_Q229H |
| 28969 | C | T | 1.90E-04 | A | 6.32E-05 | | | N | synonymous_S232S; nonsynonymous_S232R |
| 29000 | G | T | 1.90E-04 | | | | | N | nonsynonymous_G243C |
| 373 | G | A | 1.90E-04 | | | | | ORF1ab | synonymous_E36E |
| 5136 | A | G | 1.90E-04 | | | | | ORF1ab | nonsynonymous_D1624G |
| 5182 | T | C | 1.90E-04 | | | | | ORF1ab | synonymous_D1639D |
| 5392 | C | T | 1.90E-04 | | | | | ORF1ab | synonymous_N1709N |
| 5465 | T | C | 1.90E-04 | | | | | ORF1ab | nonsynonymous_Y1734H |
| 5512 | C | T | 1.90E-04 | | | | | ORF1ab | synonymous_N1749N |
| 9280 | A | G | 1.90E-04 | | | | | ORF1ab | synonymous_V3005V |
| 13860 | C | T | 1.90E-04 | | | | | ORF1ab | synonymous_D4532D |
| 13994 | C | T | 1.90E-04 | | | | | ORF1ab | nonsynonymous_A4577V |
| 14571 | C | T | 1.90E-04 | | | | | ORF1ab | synonymous_D4769D |

| | | | | | | | | | |
|---|---|---|---|---|---|---|---|---|---|
| 14646 | T | C | 1.90E-04 | | | | | ORF1ab | synonymous_T4794T |
| 14980 | C | T | 1.90E-04 | | | | | ORF1ab | nonsynonymous_L4906F |
| 15850 | G | T | 1.90E-04 | | | | | ORF1ab | nonsynonymous_D5196Y |
| 15910 | G | T | 1.90E-04 | | | | | ORF1ab | nonsynonymous_D5216Y |
| 16049 | C | T | 1.90E-04 | | | | | ORF1ab | nonsynonymous_T5262I |
| 16070 | A | G | 1.90E-04 | | | | | ORF1ab | nonsynonymous_Y5269C |
| 16073 | C | T | 1.90E-04 | | | | | ORF1ab | nonsynonymous_A5270V |
| 16503 | T | C | 1.90E-04 | | | | | ORF1ab | synonymous_V5413V |
| 16515 | T | C | 1.90E-04 | | | | | ORF1ab | synonymous_Y5417Y |
| 16535 | G | T | 1.90E-04 | | | | | ORF1ab | nonsynonymous_S5424I |
| 16622 | G | A | 1.90E-04 | | | | | ORF1ab | nonsynonymous_R5453K |
| 16638 | A | G | 1.90E-04 | | | | | ORF1ab | synonymous_A5458A |
| 17135 | C | T | 1.90E-04 | | | | | ORF1ab | nonsynonymous_P5624L |
| 17331 | G | T | 1.90E-04 | | | | | ORF1ab | nonsynonymous_E5689D |
| 17523 | G | T | 1.90E-04 | | | | | ORF1ab | nonsynonymous_M5753I |
| 17745 | C | T | 1.90E-04 | | | | | ORF1ab | synonymous_N5827N |
| 18126 | T | C | 1.90E-04 | A | 6.32E-05 | | | ORF1ab | synonymous_V5954V; synonymous_V5954V |
| 18129 | C | T | 1.90E-04 | | | | | ORF1ab | synonymous_D5955D |
| 18166 | C | T | 1.90E-04 | | | | | ORF1ab | nonsynonymous_P5968S |
| 18186 | G | T | 1.90E-04 | | | | | ORF1ab | nonsynonymous_M5974I |
| 18686 | G | T | 1.90E-04 | | | | | ORF1ab | nonsynonymous_C6141F |
| 19151 | C | T | 1.90E-04 | | | | | ORF1ab | nonsynonymous_A6296V |
| 20742 | A | G | 1.90E-04 | | | | | ORF1ab | synonymous_Q6826Q |
| 20931 | G | T | 1.90E-04 | | | | | ORF1ab | synonymous_T6889T |
| 21034 | C | T | 1.90E-04 | | | | | ORF1ab | nonsynonymous_L6924F |
| 21110 | C | T | 1.90E-04 | | | | | ORF1ab | nonsynonymous_T6949I |
| 21697 | C | A | 1.90E-04 | A | 1.90E-04 | | | S | synonymous_S45S; synonymous_S45S |
| 21727 | C | T | 1.90E-04 | | | | | S | synonymous_F55F |

| | | | | | | | | | |
|---|---|---|---|---|---|---|---|---|---|
| 22020 | T | C | 1.90E-04 | | | | | S | nonsynonymous_M153T |
| 22051 | G | T | 1.90E-04 | | | | | S | synonymous_A163A |
| 22093 | G | T | 1.90E-04 | | | | | S | nonsynonymous_M177I |
| 22205 | G | T | 1.90E-04 | T | 1.90E-04 | A | 6.32E-05 | S | nonsynonymous_D215Y; nonsynonymous_D215Y; nonsynonymous_D215N |
| 22303 | T | G | 1.90E-04 | | | | | S | nonsynonymous_S247R |
| 22320 | A | G | 1.90E-04 | | | | | S | nonsynonymous_D253G |
| 22712 | C | T | 1.90E-04 | | | | | S | nonsynonymous_P384S |
| 23557 | C | T | 1.90E-04 | | | | | S | synonymous_P665P |
| 23606 | C | T | 1.90E-04 | | | | | S | nonsynonymous_R682W |
| 23796 | A | G | 1.90E-04 | | | | | S | nonsynonymous_D745G |
| 23909 | G | T | 1.90E-04 | | | | | S | nonsynonymous_A783S |
| 23987 | C | T | 1.90E-04 | | | | | S | nonsynonymous_P809S |
| 24117 | C | T | 1.90E-04 | | | | | S | nonsynonymous_A852V |
| 24121 | A | G | 1.90E-04 | | | | | S | synonymous_Q853Q |
| 24202 | T | C | 1.90E-04 | | | | | S | synonymous_G880G |
| 24236 | G | T | 1.90E-04 | | | | | S | nonsynonymous_A892S |
| 24348 | G | T | 1.90E-04 | C | 1.26E-04 | | | S | nonsynonymous_S929I; nonsynonymous_S929T |
| 24652 | A | T | 1.90E-04 | | | | | S | synonymous_S1030S |
| 24680 | G | T | 1.90E-04 | | | | | S | nonsynonymous_V1040F |
| 25009 | G | A | 1.90E-04 | | | | | S | synonymous_K1149K |
| 25012 | G | T | 1.90E-04 | | | | | S | nonsynonymous_E1150D |
| 25046 | C | T | 1.90E-04 | G | 1.26E-04 | | | S | nonsynonymous_P1162S; nonsynonymous_P1162A |
| 25051 | T | C | 1.90E-04 | | | | | S | synonymous_D1163D |
| 25541 | T | C | 1.90E-04 | | | | | ORF3a | nonsynonymous_V50A |
| 25553 | C | T | 1.90E-04 | | | | | ORF3a | nonsynonymous_A54V |
| 25559 | T | G | 1.90E-04 | | | | | ORF3a | nonsynonymous_F56C |

| | | | | | | | | | |
|---|---|---|---|---|---|---|---|---|---|
| 25584 | C | T | 1.90E-04 | | | | | ORF3a | synonymous_T64T |
| 26022 | C | T | 1.90E-04 | | | | | ORF3a | synonymous_D210D |
| 26753 | C | T | 1.90E-04 | | | | | M | synonymous_T77T |
| 26927 | A | G | 1.90E-04 | | | | | M | synonymous_E135E |
| 26984 | C | T | 1.90E-04 | | | | | M | synonymous_H154H |
| 27021 | G | C | 1.90E-04 | | | | | M | nonsynonymous_E167Q |
| 27204 | G | T | 1.90E-04 | | | | | ORF6 | nonsynonymous_M1I |
| 27812 | C | T | 1.90E-04 | | | | | ORF7b | synonymous_F19F |
| 28310 | C | A | 1.90E-04 | A | 1.90E-04 | | | N | nonsynonymous_P13T; nonsynonymous_P13T |
| 28329 | G | T | 1.90E-04 | A | 1.26E-04 | | | N | nonsynonymous_G19V; nonsynonymous_G19E |
| 28331 | C | T | 1.90E-04 | | | | | N | nonsynonymous_P20S |
| 28380 | G | T | 1.90E-04 | | | | | N | nonsynonymous_R36L |
| 28390 | A | G | 1.90E-04 | C | 6.32E-05 | | | N | synonymous_Q39Q; nonsynonymous_Q39H |
| 28395 | G | A | 1.90E-04 | T | 6.32E-05 | | | N | nonsynonymous_R41Q; nonsynonymous_R41L |
| 28396 | G | A | 1.90E-04 | A | 1.90E-04 | | | N | synonymous_R41R; synonymous_R41R |
| 28399 | C | T | 1.90E-04 | | | | | N | synonymous_P42P |
| 28432 | C | T | 1.90E-04 | | | | | N | synonymous_F53F |
| 28677 | C | T | 1.90E-04 | | | | | N | nonsynonymous_T135I |
| 28744 | C | T | 1.90E-04 | | | | | N | synonymous_I157I |
| 29321 | G | T | 1.90E-04 | | | | | N | nonsynonymous_V350F |
| 334 | T | C | 1.90E-04 | | | | | ORF1ab | synonymous_V23V |
| 6361 | C | T | 1.90E-04 | | | | | ORF1ab | synonymous_D2032D |
| 6525 | C | T | 1.90E-04 | | | | | ORF1ab | nonsynonymous_T2087I |
| 6982 | C | T | 1.90E-04 | | | | | ORF1ab | synonymous_C2239C |
| 7006 | C | T | 1.90E-04 | | | | | ORF1ab | synonymous_T2247T |

| | | | | | | | | | |
|---|---|---|---|---|---|---|---|---|---|
| 7020 | T | G | 1.90E-04 | C | 6.32E-05 | | | ORF1ab | nonsynonymous_V2252G; nonsynonymous_V2252A |
| 8809 | C | T | 1.90E-04 | | | | | ORF1ab | synonymous_D2848D |
| 8882 | A | G | 1.90E-04 | | | | | ORF1ab | nonsynonymous_I2873V |
| 8883 | T | C | 1.90E-04 | | | | | ORF1ab | nonsynonymous_I2873T |
| 10753 | G | T | 1.90E-04 | | | | | ORF1ab | synonymous_V3496V |
| 10755 | C | T | 1.90E-04 | | | | | ORF1ab | nonsynonymous_A3497V |
| 10840 | A | G | 1.90E-04 | | | | | ORF1ab | synonymous_L3525L |
| 10960 | T | C | 1.90E-04 | | | | | ORF1ab | synonymous_G3565G |
| 10989 | C | T | 1.90E-04 | | | | | ORF1ab | nonsynonymous_T3575I |
| 11003 | C | T | 1.90E-04 | G | 6.32E-05 | | | ORF1ab | nonsynonymous_H3580Y; nonsynonymous_H3580D |
| 11474 | A | G | 1.90E-04 | | | | | ORF1ab | nonsynonymous_I3737V |
| 11991 | A | G | 1.90E-04 | | | | | ORF1ab | nonsynonymous_E3909G |
| 13176 | C | T | 1.90E-04 | | | | | ORF1ab | nonsynonymous_T4304I |
| 13265 | A | T | 1.90E-04 | | | | | ORF1ab | nonsynonymous_I4334L |
| 1326 | C | T | 1.90E-04 | | | | | ORF1ab | nonsynonymous_A354V |
| 13345 | T | C | 1.90E-04 | | | | | ORF1ab | synonymous_P4360P |
| 13749 | G | T | 1.90E-04 | | | | | ORF1ab | nonsynonymous_K4495N |
| 13768 | A | G | 1.90E-04 | | | | | ORF1ab | nonsynonymous_M4502V |
| 13774 | C | T | 1.90E-04 | | | | | ORF1ab | nonsynonymous_P4504S |
| 13965 | C | T | 1.90E-04 | | | | | ORF1ab | synonymous_Y4567Y |
| 14006 | C | A | 1.90E-04 | | | | | ORF1ab | nonsynonymous_T4581K |
| 14198 | C | T | 1.90E-04 | G | 6.32E-05 | | | ORF1ab | nonsynonymous_A4645V; nonsynonymous_A4645G |
| 14199 | A | G | 1.90E-04 | | | | | ORF1ab | synonymous_A4645A |
| 14214 | C | T | 1.90E-04 | | | | | ORF1ab | synonymous_D4650D |
| 14241 | G | T | 1.90E-04 | | | | | ORF1ab | nonsynonymous_K4659N |
| 14422 | C | T | 1.90E-04 | | | | | ORF1ab | nonsynonymous_P4720S |
| 14604 | A | G | 1.90E-04 | | | | | ORF1ab | synonymous_L4780L |

| | | | | | | | | | |
|---|---|---|---|---|---|---|---|---|---|
| 14676 | C | T | 1.90E-04 | | | | | ORF1ab | synonymous_P4804P |
| 14688 | C | T | 1.90E-04 | | | | | ORF1ab | synonymous_N4808N |
| 14708 | C | T | 1.90E-04 | | | | | ORF1ab | nonsynonymous_A4815V |
| 14712 | G | A | 1.90E-04 | | | | | ORF1ab | synonymous_V4816V |
| 14737 | A | G | 1.90E-04 | | | | | ORF1ab | nonsynonymous_S4825G |
| 14766 | T | C | 1.90E-04 | | | | | ORF1ab | synonymous_F4834F |
| 14889 | C | T | 1.90E-04 | | | | | ORF1ab | synonymous_Y4875Y |
| 15016 | G | T | 1.90E-04 | | | | | ORF1ab | nonsynonymous_A4918S |
| 15024 | C | T | 1.90E-04 | | | | | ORF1ab | synonymous_F4920F |
| 15222 | C | T | 1.90E-04 | | | | | ORF1ab | synonymous_F4986F |
| 15246 | A | G | 1.90E-04 | | | | | ORF1ab | synonymous_L4994L |
| 15371 | C | T | 1.90E-04 | | | | | ORF1ab | nonsynonymous_T5036M |
| 15405 | A | G | 1.90E-04 | | | | | ORF1ab | synonymous_L5047L |
| 15477 | A | G | 1.90E-04 | | | | | ORF1ab | synonymous_G5071G |
| 15535 | G | T | 1.90E-04 | | | | | ORF1ab | nonsynonymous_A5091S |
| 15780 | G | T | 1.90E-04 | | | | | ORF1ab | nonsynonymous_K5172N |
| 15888 | T | C | 1.90E-04 | | | | | ORF1ab | synonymous_H5208H |
| 15919 | G | T | 1.90E-04 | | | | | ORF1ab | nonsynonymous_V5219L |
| 15978 | T | C | 1.90E-04 | | | | | ORF1ab | synonymous_D5238D |
| 16047 | T | C | 1.90E-04 | | | | | ORF1ab | synonymous_L5261L |
| 16163 | C | T | 1.90E-04 | | | | | ORF1ab | nonsynonymous_T5300I |
| 16178 | C | T | 1.90E-04 | | | | | ORF1ab | nonsynonymous_S5305L |
| 16375 | C | T | 1.90E-04 | | | | | ORF1ab | nonsynonymous_P5371S |
| 16448 | A | G | 1.90E-04 | | | | | ORF1ab | nonsynonymous_Y5395C |
| 16795 | G | T | 1.90E-04 | | | | | ORF1ab | nonsynonymous_V5511L |
| 16806 | C | T | 1.90E-04 | | | | | ORF1ab | synonymous_N5514N |
| 16830 | C | T | 1.90E-04 | | | | | ORF1ab | synonymous_Y5522Y |
| 16864 | G | A | 1.90E-04 | | | | | ORF1ab | nonsynonymous_V5534I |
| 16869 | C | T | 1.90E-04 | | | | | ORF1ab | synonymous_Y5535Y |
| 16935 | G | T | 1.90E-04 | | | | | ORF1ab | nonsynonymous_M5557I |

| | | | | | | | | | |
|---|---|---|---|---|---|---|---|---|---|
| 16950 | T | C | 1.90E-04 | | | | | ORF1ab | synonymous_P5562P |
| 17140 | G | T | 1.90E-04 | | | | | ORF1ab | nonsynonymous_A5626S |
| 17172 | C | A | 1.90E-04 | | | | | ORF1ab | synonymous_A5636A |
| 17259 | G | T | 1.90E-04 | | | | | ORF1ab | nonsynonymous_E5665D |
| 17338 | G | T | 1.90E-04 | | | | | ORF1ab | nonsynonymous_A5692S |
| 17403 | C | T | 1.90E-04 | | | | | ORF1ab | synonymous_A5713A |
| 17430 | C | T | 1.90E-04 | | | | | ORF1ab | synonymous_Y5722Y |
| 17658 | G | T | 1.90E-04 | | | | | ORF1ab | nonsynonymous_M5798I |
| 17712 | A | G | 1.90E-04 | | | | | ORF1ab | synonymous_Q5816Q |
| 17894 | C | T | 1.90E-04 | | | | | ORF1ab | nonsynonymous_A5877V |
| 18079 | G | T | 1.90E-04 | | | | | ORF1ab | nonsynonymous_V5939L |
| 18252 | C | T | 1.90E-04 | | | | | ORF1ab | synonymous_N5996N |
| 18395 | C | T | 1.90E-04 | | | | | ORF1ab | nonsynonymous_A6044V |
| 18603 | T | C | 1.90E-04 | | | | | ORF1ab | synonymous_H6113H |
| 18814 | C | T | 1.90E-04 | | | | | ORF1ab | synonymous_L6184L |
| 18834 | T | C | 1.90E-04 | | | | | ORF1ab | synonymous_G6190G |
| 18894 | C | T | 1.90E-04 | G | 6.32E-05 | | | ORF1ab | synonymous_C6210C; nonsynonymous_C6210W |
| 18927 | T | C | 1.90E-04 | A | 6.32E-05 | | | ORF1ab | synonymous_Y6221Y; stopgain_Y6221X |
| 18985 | G | T | 1.90E-04 | | | | | ORF1ab | nonsynonymous_V6241F |
| 18996 | T | C | 1.90E-04 | | | | | ORF1ab | synonymous_A6244A |
| 19072 | G | T | 1.90E-04 | | | | | ORF1ab | nonsynonymous_D6270Y |
| 19101 | G | T | 1.90E-04 | | | | | ORF1ab | nonsynonymous_Q6279H |
| 19854 | C | T | 1.90E-04 | | | | | ORF1ab | synonymous_D6530D |
| 20628 | C | T | 1.90E-04 | | | | | ORF1ab | synonymous_G6788G |
| 20667 | A | G | 1.90E-04 | | | | | ORF1ab | synonymous_Q6801Q |
| 20756 | G | T | 1.90E-04 | | | | | ORF1ab | nonsynonymous_S6831I |
| 21072 | T | C | 1.90E-04 | | | | | ORF1ab | synonymous_N6936N |
| 21805 | C | T | 1.90E-04 | | | | | S | synonymous_N81N |

| | | | | | | | | | |
|---|---|---|---|---|---|---|---|---|---|
| 21830 | G | T | 1.90E-04 | | | | | S | nonsynonymous_V90F |
| 21859 | C | T | 1.90E-04 | | | | | S | synonymous_N99N |
| 21999 | A | G | 1.90E-04 | | | | | S | nonsynonymous_H146R |
| 22009 | C | T | 1.90E-04 | | | | | S | synonymous_N149N |
| 22687 | C | T | 1.90E-04 | | | | | S | synonymous_S375S |
| 23170 | C | T | 1.90E-04 | | | | | S | synonymous_N536N |
| 23230 | C | T | 1.90E-04 | A | 6.32E-05 | | | S | synonymous_N556N; nonsynonymous_N556K |
| 23282 | G | T | 1.90E-04 | | | | | S | nonsynonymous_D574Y |
| 23287 | T | C | 1.90E-04 | G | 1.26E-04 | | | S | synonymous_A575A; synonymous_A575A |
| 23678 | G | T | 1.90E-04 | | | | | S | nonsynonymous_A706S |
| 23782 | G | T | 1.90E-04 | | | | | S | nonsynonymous_M740I |
| 23870 | A | G | 1.90E-04 | | | | | S | nonsynonymous_I770V |
| 23893 | C | T | 1.90E-04 | | | | | S | synonymous_N777N |
| 24076 | T | C | 1.90E-04 | | | | | S | synonymous_G838G |
| 2526 | A | C | 1.90E-04 | | | | | ORF1ab | nonsynonymous_E754A |
| 25305 | G | C | 1.90E-04 | T | 6.32E-05 | | | S | nonsynonymous_C1248S; nonsynonymous_C1248F |
| 25315 | A | G | 1.90E-04 | | | | | S | synonymous_G1251G |
| 25316 | T | C | 1.90E-04 | | | | | S | nonsynonymous_S1252P |
| 25352 | G | T | 1.90E-04 | | | | | S | nonsynonymous_V1264L |
| 25455 | G | T | 1.90E-04 | | | | | ORF3a | nonsynonymous_K21N |
| 25456 | G | T | 1.90E-04 | | | | | ORF3a | nonsynonymous_D22Y |
| 25487 | C | T | 1.90E-04 | | | | | ORF3a | nonsynonymous_T32I |
| 26023 | T | C | 1.90E-04 | | | | | ORF3a | nonsynonymous_Y211H |
| 26032 | C | T | 1.90E-04 | | | | | ORF3a | synonymous_L214L |
| 26172 | G | A | 1.90E-04 | | | | | ORF3a | nonsynonymous_M260I |
| 26195 | C | T | 1.90E-04 | | | | | ORF3a | nonsynonymous_T268M |

| | | | | | | | | | |
|---|---|---|---|---|---|---|---|---|---|
| 26319 | A | T | 1.90E-04 | G | 6.32E-05 | | | E | synonymous_V25V; synonymous_V25V |
| 26727 | G | T | 1.90E-04 | | | | | M | nonsynonymous_A69S |
| 26746 | G | T | 1.90E-04 | | | | | M | nonsynonymous_W75L |
| 26775 | G | T | 1.90E-04 | | | | | M | nonsynonymous_A85S |
| 26858 | C | T | 1.90E-04 | | | | | M | synonymous_F112F |
| 27238 | G | T | 1.90E-04 | | | | | ORF6 | stopgain_E13X |
| 27253 | A | G | 1.90E-04 | | | | | ORF6 | nonsynonymous_I18V |
| 27263 | C | T | 1.90E-04 | | | | | ORF6 | nonsynonymous_T21I |
| 27331 | C | T | 1.90E-04 | | | | | ORF6 | synonymous_L44L |
| 27348 | T | C | 1.90E-04 | | | | | ORF6 | synonymous_Y49Y |
| 27364 | G | T | 1.90E-04 | C | 1.26E-04 | | | ORF6 | stopgain_E55X; nonsynonymous_E55Q |
| 27416 | C | T | 1.90E-04 | | | | | ORF7a | nonsynonymous_A8V |
| 27451 | T | A | 1.90E-04 | | | | | ORF7a | nonsynonymous_Y20N |
| 27463 | G | T | 1.90E-04 | | | | | ORF7a | nonsynonymous_V24F |
| 28289 | C | A | 1.90E-04 | A | 1.90E-04 | | | N | nonsynonymous_P6T; nonsynonymous_P6T |
| 28512 | C | T | 1.90E-04 | | | | | N | nonsynonymous_P80L |
| 28567 | C | T | 1.90E-04 | | | | | N | synonymous_D98D |
| 28615 | A | G | 1.90E-04 | | | | | N | synonymous_G114G |
| 28723 | T | C | 1.90E-04 | | | | | N | synonymous_N150N |
| 28724 | C | T | 1.90E-04 | | | | | N | nonsynonymous_P151S |
| 29029 | T | C | 1.90E-04 | | | | | N | synonymous_A252A |
| 3046 | A | G | 1.90E-04 | | | | | ORF1ab | synonymous_P927P |
| 3340 | G | T | 1.90E-04 | | | | | ORF1ab | synonymous_V1025V |
| 3874 | C | T | 1.90E-04 | | | | | ORF1ab | synonymous_I1203I |
| 3889 | A | G | 1.90E-04 | | | | | ORF1ab | synonymous_K1208K |
| 3892 | G | T | 1.90E-04 | | | | | ORF1ab | nonsynonymous_E1209D |
| 3992 | C | T | 1.90E-04 | | | | | ORF1ab | synonymous_L1243L |

| | | | | | | | | | |
|---|---|---|---|---|---|---|---|---|---|
| 4543 | C | T | 1.90E-04 | | | | | ORF1ab | synonymous_T1426T |
| 593 | C | T | 1.90E-04 | | | | | ORF1ab | nonsynonymous_H110Y |
| 601 | C | T | 1.90E-04 | | | | | ORF1ab | synonymous_G112G |
| 6120 | C | T | 1.90E-04 | | | | | ORF1ab | nonsynonymous_S1952L |
| 6317 | C | T | 1.90E-04 | A | 6.32E-05 | | | ORF1ab | nonsynonymous_P2018S; nonsynonymous_P2018T |
| 6329 | T | C | 1.90E-04 | | | | | ORF1ab | nonsynonymous_S2022P |
| 6352 | G | T | 1.90E-04 | A | 6.32E-05 | | | ORF1ab | nonsynonymous_K2029N; synonymous_K2029K |
| 6384 | C | T | 1.90E-04 | | | | | ORF1ab | nonsynonymous_A2040V |
| 6400 | A | G | 1.90E-04 | | | | | ORF1ab | synonymous_K2045K |
| 6466 | A | G | 1.90E-04 | | | | | ORF1ab | synonymous_K2067K |
| 6469 | T | C | 1.90E-04 | | | | | ORF1ab | synonymous_T2068T |
| 6472 | C | T | 1.90E-04 | | | | | ORF1ab | synonymous_T2069T |
| 6487 | C | T | 1.90E-04 | | | | | ORF1ab | synonymous_D2074D |
| 6543 | C | T | 1.90E-04 | | | | | ORF1ab | nonsynonymous_T2093I |
| 6582 | C | T | 1.90E-04 | | | | | ORF1ab | nonsynonymous_T2106I |
| 6604 | A | G | 1.90E-04 | | | | | ORF1ab | synonymous_L2113L |
| 6628 | C | T | 1.90E-04 | | | | | ORF1ab | synonymous_T2121T |
| 6633 | C | T | 1.90E-04 | | | | | ORF1ab | nonsynonymous_A2123V |
| 6730 | C | T | 1.90E-04 | | | | | ORF1ab | synonymous_N2155N |
| 6813 | C | T | 1.90E-04 | | | | | ORF1ab | nonsynonymous_T2183I |
| 6894 | G | T | 1.90E-04 | | | | | ORF1ab | nonsynonymous_C2210F |
| 7093 | C | T | 1.90E-04 | | | | | ORF1ab | synonymous_V2276V |
| 7124 | C | T | 1.90E-04 | | | | | ORF1ab | nonsynonymous_P2287S |
| 7267 | C | T | 1.90E-04 | | | | | ORF1ab | synonymous_F2334F |
| 7929 | A | G | 1.90E-04 | | | | | ORF1ab | nonsynonymous_K2555R |
| 8016 | C | T | 1.90E-04 | | | | | ORF1ab | nonsynonymous_A2584V |
| 8676 | G | T | 1.90E-04 | | | | | ORF1ab | nonsynonymous_S2804I |
| 8681 | A | G | 1.90E-04 | | | | | ORF1ab | nonsynonymous_I2806V |

| | | | | | | | | | |
|---|---|---|---|---|---|---|---|---|---|
| 8733 | C | T | 1.90E-04 | | | | | ORF1ab | nonsynonymous_T2823I |
| 9442 | C | T | 1.90E-04 | | | | | ORF1ab | synonymous_C3059C |
| 10116 | C | T | 1.90E-04 | | | | | ORF1ab | nonsynonymous_T3284I |
| 10401 | C | T | 1.90E-04 | | | | | ORF1ab | nonsynonymous_A3379V |
| 10449 | C | T | 1.90E-04 | | | | | ORF1ab | nonsynonymous_P3395L |
| 11032 | T | G | 1.90E-04 | C | 6.32E-05 | | | ORF1ab | synonymous_T3589T; synonymous_T3589T |
| 11036 | C | T | 1.90E-04 | | | | | ORF1ab | nonsynonymous_L3591F |
| 11200 | C | T | 1.90E-04 | | | | | ORF1ab | synonymous_A3645A |
| 12115 | C | T | 1.90E-04 | | | | | ORF1ab | synonymous_S3950S |
| 12188 | G | T | 1.90E-04 | | | | | ORF1ab | nonsynonymous_V3975F |
| 12199 | A | G | 1.90E-04 | | | | | ORF1ab | synonymous_K3978K |
| 12513 | C | T | 1.90E-04 | | | | | ORF1ab | nonsynonymous_T4083M |
| 12923 | C | A | 1.90E-04 | A | 1.90E-04 | | | ORF1ab | nonsynonymous_P4220T; nonsynonymous_P4220T |
| 1346 | C | T | 1.90E-04 | | | | | ORF1ab | nonsynonymous_P361S |
| 1415 | C | T | 1.90E-04 | | | | | ORF1ab | nonsynonymous_L384F |
| 1646 | A | G | 1.90E-04 | | | | | ORF1ab | nonsynonymous_I461V |
| 1663 | C | T | 1.90E-04 | | | | | ORF1ab | synonymous_D466D |
| 1685 | G | A | 1.90E-04 | | | | | ORF1ab | nonsynonymous_A474T |
| 1714 | C | T | 1.90E-04 | | | | | ORF1ab | synonymous_S483S |
| 1911 | C | T | 1.90E-04 | | | | | ORF1ab | nonsynonymous_S549F |
| 2035 | G | T | 1.90E-04 | | | | | ORF1ab | nonsynonymous_L590F |
| 2061 | C | T | 1.90E-04 | | | | | ORF1ab | nonsynonymous_A599V |
| 2070 | C | T | 1.90E-04 | | | | | ORF1ab | nonsynonymous_T602I |
| 2189 | C | T | 1.90E-04 | | | | | ORF1ab | nonsynonymous_L642F |
| 2196 | A | C | 1.90E-04 | | | | | ORF1ab | nonsynonymous_D644A |
| 2399 | C | T | 1.90E-04 | | | | | ORF1ab | nonsynonymous_H712Y |
| 2918 | C | T | 1.90E-04 | | | | | ORF1ab | nonsynonymous_P885S |
| 2980 | C | T | 1.90E-04 | | | | | ORF1ab | synonymous_Y905Y |

| | | | | | | | | | |
|---|---|---|---|---|---|---|---|---|---|
| 3052 | G | A | 1.90E-04 | | | | | ORF1ab | synonymous_E929E |
| 3119 | G | A | 1.90E-04 | | | | | ORF1ab | nonsynonymous_E952K |
| 3312 | T | C | 1.90E-04 | | | | | ORF1ab | nonsynonymous_L1016P |
| 3429 | C | T | 1.90E-04 | | | | | ORF1ab | nonsynonymous_T1055I |
| 3656 | C | T | 1.90E-04 | | | | | ORF1ab | nonsynonymous_L1131F |
| 3923 | C | A | 1.90E-04 | T | 1.26E-04 | | | ORF1ab | nonsynonymous_P1220T;<br>nonsynonymous_P1220S |
| 392 | C | T | 1.90E-04 | | | | | ORF1ab | nonsynonymous_R43C |
| 4206 | C | T | 1.90E-04 | | | | | ORF1ab | nonsynonymous_A1314V |
| 4230 | C | T | 1.90E-04 | A | 6.32E-05 | | | ORF1ab | nonsynonymous_T1322I;<br>nonsynonymous_T1322K |
| 4586 | C | T | 1.90E-04 | | | | | ORF1ab | synonymous_L1441L |
| 487 | G | A | 1.90E-04 | | | | | ORF1ab | synonymous_S74S |
| 5629 | G | T | 1.90E-04 | | | | | ORF1ab | synonymous_T1788T |
| 5654 | C | T | 1.90E-04 | | | | | ORF1ab | synonymous_L1797L |
| 5950 | G | T | 1.90E-04 | | | | | ORF1ab | nonsynonymous_K1895N |
| 6123 | G | A | 1.90E-04 | | | | | ORF1ab | nonsynonymous_R1953K |
| 6163 | T | A | 1.90E-04 | | | | | ORF1ab | nonsynonymous_D1966E |
| 6507 | A | G | 1.90E-04 | C | 1.26E-04 | | | ORF1ab | nonsynonymous_N2081S;<br>nonsynonymous_N2081T |
| 6548 | C | T | 1.90E-04 | | | | | ORF1ab | synonymous_L2095L |
| 6578 | C | T | 1.90E-04 | | | | | ORF1ab | nonsynonymous_L2105F |
| 6646 | A | G | 1.90E-04 | | | | | ORF1ab | synonymous_L2127L |
| 6695 | C | A | 1.90E-04 | T | 6.32E-05 | | | ORF1ab | nonsynonymous_P2144T;<br>nonsynonymous_P2144S |
| 6701 | C | T | 1.90E-04 | | | | | ORF1ab | nonsynonymous_L2146F |
| 6720 | C | T | 1.90E-04 | | | | | ORF1ab | nonsynonymous_T2152I |
| 7224 | C | T | 1.90E-04 | | | | | ORF1ab | nonsynonymous_A2320V |
| 7318 | C | T | 1.90E-04 | A | 6.32E-05 | | | ORF1ab | synonymous_F2351F;<br>nonsynonymous_F2351L |

| | | | | | | | | | |
|---|---|---|---|---|---|---|---|---|---|
| 7334 | C | T | 1.90E-04 | | | | | ORF1ab | nonsynonymous_H2357Y |
| 7756 | G | T | 1.90E-04 | | | | | ORF1ab | nonsynonymous_K2497N |
| 7824 | A | G | 1.90E-04 | | | | | ORF1ab | nonsynonymous_H2520R |
| 7851 | C | T | 1.90E-04 | | | | | ORF1ab | nonsynonymous_A2529V |
| 8170 | A | G | 1.90E-04 | | | | | ORF1ab | synonymous_S2635S |
| 9166 | C | T | 1.90E-04 | | | | | ORF1ab | synonymous_T2967T |
| 9661 | C | T | 1.90E-04 | | | | | ORF1ab | synonymous_F3132F |
| 9727 | T | C | 1.90E-04 | | | | | ORF1ab | synonymous_Y3154Y |
| 10030 | C | T | 1.90E-04 | | | | | ORF1ab | synonymous_T3255T |
| 10339 | T | C | 1.90E-04 | | | | | ORF1ab | synonymous_N3358N |
| 10525 | C | T | 1.90E-04 | | | | | ORF1ab | synonymous_V3420V |
| 10631 | G | A | 1.90E-04 | | | | | ORF1ab | nonsynonymous_A3456T |
| 1077 | C | T | 1.90E-04 | A | 6.32E-05 | | | ORF1ab | nonsynonymous_P271L;<br>nonsynonymous_P271Q |
| 11102 | C | T | 1.90E-04 | | | | | ORF1ab | nonsynonymous_P3613S |
| 11117 | A | G | 1.90E-04 | | | | | ORF1ab | nonsynonymous_I3618V |
| 11182 | T | C | 1.90E-04 | | | | | ORF1ab | synonymous_F3639F |
| 11191 | T | G | 1.90E-04 | A | 6.32E-05 | | | ORF1ab | synonymous_P3642P;<br>synonymous_P3642P |
| 11298 | A | G | 1.90E-04 | | | | | ORF1ab | nonsynonymous_K3678R |
| 11344 | C | T | 1.90E-04 | | | | | ORF1ab | synonymous_I3693I |
| 11455 | C | T | 1.90E-04 | | | | | ORF1ab | synonymous_A3730A |
| 1149 | G | T | 1.90E-04 | A | 6.32E-05 | | | ORF1ab | nonsynonymous_G295V;<br>nonsynonymous_G295D |
| 1150 | C | T | 1.90E-04 | | | | | ORF1ab | synonymous_G295G |
| 11514 | C | T | 1.90E-04 | | | | | ORF1ab | nonsynonymous_T3750I |
| 11516 | G | A | 1.90E-04 | | | | | ORF1ab | nonsynonymous_V3751I |
| 11521 | G | T | 1.90E-04 | | | | | ORF1ab | nonsynonymous_M3752I |
| 11522 | T | G | 1.90E-04 | | | | | ORF1ab | nonsynonymous_F3753V |
| 11590 | A | G | 1.90E-04 | | | | | ORF1ab | synonymous_T3775T |

| | | | | | | | | | |
|---|---|---|---|---|---|---|---|---|---|
| 1165 | T | C | 1.90E-04 | | | | | ORF1ab | synonymous_I300I |
| 11660 | T | A | 1.90E-04 | | | | | ORF1ab | nonsynonymous_L3799I |
| 11767 | C | T | 1.90E-04 | | | | | ORF1ab | synonymous_S3834S |
| 1177 | T | C | 1.90E-04 | | | | | ORF1ab | synonymous_Y304Y |
| 1181 | G | T | 1.90E-04 | | | | | ORF1ab | nonsynonymous_V306F |
| 11831 | G | A | 1.90E-04 | T | 6.32E-05 | | | ORF1ab | nonsynonymous_A3856T; nonsynonymous_A3856S |
| 11902 | C | T | 1.90E-04 | | | | | ORF1ab | synonymous_L3879L |
| 12070 | G | T | 1.90E-04 | | | | | ORF1ab | synonymous_L3935L |
| 12084 | C | T | 1.90E-04 | | | | | ORF1ab | nonsynonymous_T3940I |
| 1211 | T | G | 1.90E-04 | | | | | ORF1ab | nonsynonymous_C316G |
| 1213 | C | T | 1.90E-04 | | | | | ORF1ab | synonymous_C316C |
| 12286 | A | G | 1.90E-04 | | | | | ORF1ab | synonymous_Q4007Q |
| 12357 | C | T | 1.90E-04 | | | | | ORF1ab | nonsynonymous_T4031I |
| 12418 | T | C | 1.90E-04 | | | | | ORF1ab | synonymous_N4051N |
| 1263 | C | T | 1.90E-04 | | | | | ORF1ab | nonsynonymous_T333M |
| 12722 | T | C | 1.90E-04 | | | | | ORF1ab | nonsynonymous_S4153P |
| 12729 | C | T | 1.90E-04 | | | | | ORF1ab | nonsynonymous_A4155V |
| 12779 | T | C | 1.90E-04 | | | | | ORF1ab | nonsynonymous_Y4172H |
| 12789 | C | T | 1.90E-04 | | | | | ORF1ab | nonsynonymous_T4175I |
| 12885 | C | T | 1.90E-04 | | | | | ORF1ab | nonsynonymous_T4207I |
| 12924 | C | T | 1.90E-04 | | | | | ORF1ab | nonsynonymous_P4220L |
| 12933 | C | T | 1.90E-04 | | | | | ORF1ab | nonsynonymous_P4223L |
| 12988 | G | T | 1.90E-04 | | | | | ORF1ab | nonsynonymous_M4241I |
| 13064 | T | C | 1.90E-04 | | | | | ORF1ab | synonymous_L4267L |
| 1412 | A | G | 1.90E-04 | | | | | ORF1ab | nonsynonymous_S383G |
| 1430 | A | G | 1.90E-04 | | | | | ORF1ab | nonsynonymous_N389D |
| 1592 | T | C | 1.90E-04 | | | | | ORF1ab | nonsynonymous_S443P |
| 1909 | C | T | 1.90E-04 | | | | | ORF1ab | synonymous_F548F |
| 1914 | G | A | 1.90E-04 | | | | | ORF1ab | nonsynonymous_R550H |

| | | | | | | | | | |
|---|---|---|---|---|---|---|---|---|---|
| 2219 | A | G | 1.90E-04 | | | | | ORF1ab | nonsynonymous_I652V |
| 2469 | C | T | 1.90E-04 | | | | | ORF1ab | nonsynonymous_A735V |
| 3171 | T | C | 1.90E-04 | A | 6.32E-05 | | | ORF1ab | nonsynonymous_L969P; nonsynonymous_L969H |
| 3178 | T | G | 1.90E-04 | | | | | ORF1ab | synonymous_P971P |
| 3234 | A | G | 1.90E-04 | | | | | ORF1ab | nonsynonymous_Q990R |
| 3518 | G | T | 1.90E-04 | | | | | ORF1ab | nonsynonymous_V1085F |
| 3587 | C | T | 1.90E-04 | | | | | ORF1ab | nonsynonymous_H1108Y |
| 3628 | T | C | 1.90E-04 | | | | | ORF1ab | synonymous_N1121N |
| 3689 | G | A | 1.90E-04 | | | | | ORF1ab | nonsynonymous_E1142K |
| 3710 | T | C | 1.90E-04 | | | | | ORF1ab | synonymous_L1149L |
| 3741 | T | C | 1.90E-04 | G | 6.32E-05 | | | ORF1ab | nonsynonymous_I1159T; nonsynonymous_I1159R |
| 3793 | T | C | 1.90E-04 | | | | | ORF1ab | synonymous_A1176A |
| 4276 | C | T | 1.90E-04 | | | | | ORF1ab | synonymous_Y1337Y |
| 4399 | G | T | 1.90E-04 | | | | | ORF1ab | nonsynonymous_M1378I |
| 4544 | A | T | 1.90E-04 | | | | | ORF1ab | nonsynonymous_S1427C |
| 4548 | A | G | 1.90E-04 | | | | | ORF1ab | nonsynonymous_K1428R |
| 4827 | C | T | 1.90E-04 | | | | | ORF1ab | nonsynonymous_T1521I |
| 4890 | C | T | 1.90E-04 | | | | | ORF1ab | nonsynonymous_T1542I |
| 4904 | G | T | 1.90E-04 | | | | | ORF1ab | nonsynonymous_D1547Y |
| 4946 | T | G | 1.90E-04 | C | 6.32E-05 | | | ORF1ab | nonsynonymous_S1561A; nonsynonymous_S1561P |
| 4965 | C | T | 1.90E-04 | | | | | ORF1ab | nonsynonymous_T1567I |
| 5717 | C | T | 1.90E-04 | | | | | ORF1ab | nonsynonymous_H1818Y |
| 5724 | C | T | 1.90E-04 | | | | | ORF1ab | nonsynonymous_T1820I |
| 5845 | A | T | 1.90E-04 | | | | | ORF1ab | nonsynonymous_K1860N |
| 5855 | A | G | 1.90E-04 | | | | | ORF1ab | nonsynonymous_T1864A |
| 5860 | T | C | 1.90E-04 | | | | | ORF1ab | synonymous_D1865D |
| 5907 | C | T | 1.90E-04 | | | | | ORF1ab | nonsynonymous_T1881I |

| | | | | | | | | | |
|---|---|---|---|---|---|---|---|---|---|
| 5917 | G | A | 1.90E-04 | | | | | ORF1ab | synonymous_L1884L |
| 6133 | A | G | 1.90E-04 | | | | | ORF1ab | synonymous_K1956K |
| 652 | A | T | 1.90E-04 | | | | | ORF1ab | nonsynonymous_K129N |
| 700 | A | G | 1.90E-04 | | | | | ORF1ab | synonymous_L145L |
| 703 | C | T | 1.90E-04 | | | | | ORF1ab | synonymous_G146G |
| 714 | G | A | 1.90E-04 | | | | | ORF1ab | nonsynonymous_G150D |
| 7216 | T | C | 1.90E-04 | | | | | ORF1ab | synonymous_D2317D |
| 721 | T | C | 1.90E-04 | | | | | ORF1ab | synonymous_D152D |
| 7504 | C | T | 1.90E-04 | | | | | ORF1ab | synonymous_Y2413Y |
| 7528 | C | T | 1.90E-04 | | | | | ORF1ab | synonymous_V2421V |
| 7621 | T | C | 1.90E-04 | | | | | ORF1ab | synonymous_C2452C |
| 7644 | C | T | 1.90E-04 | | | | | ORF1ab | nonsynonymous_A2460V |
| 8299 | C | T | 1.90E-04 | | | | | ORF1ab | synonymous_N2678N |
| 8386 | C | T | 1.90E-04 | | | | | ORF1ab | synonymous_H2707H |
| 8391 | T | C | 1.90E-04 | | | | | ORF1ab | nonsynonymous_I2709T |
| 8394 | C | T | 1.90E-04 | | | | | ORF1ab | nonsynonymous_A2710V |
| 843 | C | T | 1.90E-04 | | | | | ORF1ab | nonsynonymous_P193L |
| 854 | C | T | 1.90E-04 | | | | | ORF1ab | nonsynonymous_P197S |
| 885 | G | T | 1.90E-04 | | | | | ORF1ab | nonsynonymous_R207L |
| 8964 | C | T | 1.90E-04 | | | | | ORF1ab | nonsynonymous_S2900L |
| 9073 | C | T | 1.90E-04 | | | | | ORF1ab | synonymous_T2936T |
| 9111 | G | A | 1.90E-04 | | | | | ORF1ab | nonsynonymous_R2949H |
| 915 | A | G | 1.90E-04 | | | | | ORF1ab | nonsynonymous_E217G |
| 9176 | G | A | 1.90E-04 | C | 6.32E-05 | | | ORF1ab | nonsynonymous_G2971S; nonsynonymous_G2971R |
| 9243 | A | G | 1.90E-04 | | | | | ORF1ab | nonsynonymous_E2993G |
| 931 | T | C | 1.90E-04 | | | | | ORF1ab | synonymous_I222I |
| 9526 | G | T | 1.90E-04 | | | | | ORF1ab | nonsynonymous_M3087I |
| 9623 | G | T | 1.90E-04 | | | | | ORF1ab | nonsynonymous_V3120F |
| 9693 | C | T | 1.90E-04 | | | | | ORF1ab | nonsynonymous_A3143V |

| | | | | | | | | | |
|---|---|---|---|---|---|---|---|---|---|
| 9802 | G | T | 1.90E-04 | | | | | ORF1ab | synonymous_A3179A |
| 9848 | A | G | 1.90E-04 | | | | | ORF1ab | nonsynonymous_S3195G |
| 9996 | C | T | 1.90E-04 | | | | | ORF1ab | nonsynonymous_S3244L |
| 29765 | T | A | 1.52E-04 | A | 1.52E-04 | G | 7.58E-05 | three_prime_UTR | |
| 29827 | A | G | 1.37E-04 | C | 6.87E-05 | | | three_prime_UTR | |
| 29815 | A | G | 1.36E-04 | T | 6.81E-05 | | | three_prime_UTR | |
| 29789 | T | A | 1.35E-04 | G | 6.73E-05 | | | three_prime_UTR | |
| 29797 | A | C | 1.35E-04 | T | 6.73E-05 | T | 6.73E-05 | three_prime_UTR | |
| 29796 | A | T | 1.35E-04 | | | | | three_prime_UTR | |
| 59 | C | T | 1.33E-04 | | | | | five_prime_UTR | |
| 64 | C | T | 1.31E-04 | | | | | five_prime_UTR | |
| 76 | T | A | 1.30E-04 | | | | | five_prime_UTR | |
| 78 | T | A | 1.30E-04 | G | 6.50E-05 | | | five_prime_UTR | |
| 83 | T | C | 1.30E-04 | | | | | five_prime_UTR | |
| 98 | C | T | 1.29E-04 | | | | | five_prime_UTR | |
| 100 | C | A | 1.29E-04 | | | | | five_prime_UTR | |
| 102 | G | A | 1.29E-04 | | | | | five_prime_UTR | |
| 116 | G | A | 1.29E-04 | | | | | five_prime_UTR | |
| 126 | A | T | 1.28E-04 | | | | | five_prime_UTR | |
| 125 | C | T | 1.28E-04 | | | | | five_prime_UTR | |
| 136 | A | G | 1.28E-04 | | | | | five_prime_UTR | |
| 141 | T | C | 1.28E-04 | A | 6.41E-05 | | | five_prime_UTR | |
| 148 | T | C | 1.28E-04 | | | | | five_prime_UTR | |
| 29759 | G | T | 1.28E-04 | A | 6.41E-05 | | | three_prime_UTR | |
| 144 | T | C | 1.28E-04 | | | | | five_prime_UTR | |
| 29762 | C | T | 1.28E-04 | A | 6.41E-05 | | | three_prime_UTR | |
| 29755 | G | A | 1.28E-04 | | | | | three_prime_UTR | |
| 29744 | G | A | 1.28E-04 | T | 6.39E-05 | C | 6.39E-05 | three_prime_UTR | |
| 167 | G | T | 1.28E-04 | T | 1.28E-04 | A | 6.39E-05 | five_prime_UTR | |
| 29738 | C | T | 1.28E-04 | | | | | three_prime_UTR | |

| | | | | | | | | | |
|---|---|---|---|---|---|---|---|---|---|
| 193 | C | T | 1.28E-04 | | | | | five_prime_UTR | |
| 198 | G | A | 1.28E-04 | T | 6.38E-05 | | | five_prime_UTR | |
| 225 | C | T | 1.28E-04 | | | | | five_prime_UTR | |
| 229 | A | C | 1.28E-04 | | | | | five_prime_UTR | |
| 208 | G | T | 1.28E-04 | | | | | five_prime_UTR | |
| 232 | T | C | 1.28E-04 | G | 6.38E-05 | | | five_prime_UTR | |
| 29715 | G | A | 1.28E-04 | | | | | three_prime_UTR | |
| 29717 | G | T | 1.28E-04 | | | | | three_prime_UTR | |
| 234 | T | C | 1.28E-04 | | | | | five_prime_UTR | |
| 29730 | C | T | 1.28E-04 | G | 6.38E-05 | | | three_prime_UTR | |
| 29731 | A | T | 1.28E-04 | | | | | three_prime_UTR | |
| 29724 | C | T | 1.27E-04 | | | | | three_prime_UTR | |
| 29695 | A | G | 1.27E-04 | | | | | three_prime_UTR | |
| 29692 | G | T | 1.27E-04 | | | | | three_prime_UTR | |
| 263 | A | T | 1.27E-04 | G | 6.36E-05 | | | five_prime_UTR | |
| 264 | A | G | 1.27E-04 | | | | | five_prime_UTR | |
| 29679 | C | T | 1.27E-04 | | | | | three_prime_UTR | |
| 520 | G | T | 1.27E-04 | | | | | ORF1ab | nonsynonymous_M85I |
| 11075 | T | C | 1.27E-04 | | | | | ORF1ab | nonsynonymous_F3604L |
| 8022 | T | G | 1.27E-04 | | | | | ORF1ab | nonsynonymous_V2586G |
| 512 | C | T | 1.27E-04 | | | | | ORF1ab | nonsynonymous_H83Y |
| 27532 | C | T | 1.27E-04 | | | | | ORF7a | nonsynonymous_H47Y |
| 27654 | C | T | 1.27E-04 | | | | | ORF7a | synonymous_F87F |
| 19525 | G | T | 1.27E-04 | A | 6.33E-05 | | | ORF1ab | nonsynonymous_D6421Y; nonsynonymous_D6421N |
| 27651 | G | C | 1.27E-04 | T | 6.33E-05 | | | ORF7a | synonymous_L86L; synonymous_L86L |
| 19454 | C | T | 1.27E-04 | | | | | ORF1ab | nonsynonymous_T6397M |
| 27625 | C | T | 1.27E-04 | | | | | ORF7a | nonsynonymous_R78C |
| 27698 | T | G | 1.27E-04 | | | | | ORF7a | nonsynonymous_L102R |

| | | | | | | | | | |
|---|---|---|---|---|---|---|---|---|---|
| 27706 | G | T | 1.27E-04 | | | | | ORF7a | nonsynonymous_A105S |
| 27709 | G | T | 1.27E-04 | | | | | ORF7a | nonsynonymous_A106S |
| 27727 | C | T | 1.27E-04 | A | 6.33E-05 | | | ORF7a | nonsynonymous_L112F; nonsynonymous_L112I |
| 27740 | T | C | 1.27E-04 | | | | | ORF7a | nonsynonymous_L116P |
| 27798 | G | T | 1.27E-04 | A | 6.33E-05 | | | ORF7b | nonsynonymous_A15S; nonsynonymous_A15T |
| 27799 | C | T | 1.27E-04 | | | | | ORF7b | nonsynonymous_A15V |
| 532 | A | G | 1.27E-04 | | | | | ORF1ab | synonymous_V89V |
| 21762 | C | T | 1.27E-04 | | | | | S | nonsynonymous_A67V |
| 19328 | C | T | 1.27E-04 | | | | | ORF1ab | nonsynonymous_A6355V |
| 19417 | G | T | 1.27E-04 | | | | | ORF1ab | nonsynonymous_V6385L |
| 19429 | G | T | 1.27E-04 | | | | | ORF1ab | nonsynonymous_D6389Y |
| 19521 | T | C | 1.27E-04 | | | | | ORF1ab | synonymous_Y6419Y |
| 27545 | A | G | 1.27E-04 | | | | | ORF7a | nonsynonymous_D51G |
| 28178 | G | T | 1.27E-04 | | | | | ORF8 | nonsynonymous_L95F |
| 19325 | C | T | 1.27E-04 | | | | | ORF1ab | nonsynonymous_P6354L |
| 19391 | C | T | 1.27E-04 | | | | | ORF1ab | nonsynonymous_P6376L |
| 19476 | A | G | 1.27E-04 | | | | | ORF1ab | synonymous_L6404L |
| 27754 | G | T | 1.27E-04 | | | | | ORF7a | stopgain_E121X |
| 28054 | C | T | 1.27E-04 | | | | | ORF8 | nonsynonymous_S54L |
| 28153 | C | T | 1.27E-04 | | | | | ORF8 | nonsynonymous_T87I |
| 19392 | A | G | 1.27E-04 | | | | | ORF1ab | synonymous_P6376P |
| 27911 | C | T | 1.27E-04 | | | | | ORF8 | synonymous_F6F |
| 27952 | G | T | 1.27E-04 | | | | | ORF8 | nonsynonymous_C20F |
| 27967 | G | T | 1.27E-04 | | | | | ORF8 | nonsynonymous_C25F |
| 27987 | G | T | 1.27E-04 | A | 6.33E-05 | | | ORF8 | nonsynonymous_V32L; nonsynonymous_V32I |
| 28005 | C | T | 1.27E-04 | | | | | ORF8 | nonsynonymous_P38S |

| | | | | | | | | | |
|---|---|---|---|---|---|---|---|---|---|
| 28027 | G | T | 1.27E-04 | T | 1.27E-04 | | | ORF8 | nonsynonymous_W45L; nonsynonymous_W45L |
| 28060 | C | T | 1.27E-04 | | | | | ORF8 | nonsynonymous_P56L |
| 28081 | A | G | 1.27E-04 | C | 6.33E-05 | | | ORF8 | nonsynonymous_D63G; nonsynonymous_D63A |
| 28121 | C | T | 1.27E-04 | | | | | ORF8 | synonymous_I76I |
| 28155 | A | G | 1.27E-04 | | | | | ORF8 | nonsynonymous_I88V |
| 28175 | A | G | 1.27E-04 | | | | | ORF8 | synonymous_K94K |
| 28179 | G | A | 1.27E-04 | T | 6.33E-05 | | | ORF8 | nonsynonymous_G96S; nonsynonymous_G96C |
| 28195 | G | T | 1.27E-04 | | | | | ORF8 | nonsynonymous_R101L |
| 28221 | G | T | 1.27E-04 | | | | | ORF8 | stopgain_E110X |
| 28229 | T | G | 1.27E-04 | | | | | ORF8 | nonsynonymous_H112Q |
| 27870 | G | T | 1.27E-04 | A | 6.33E-05 | | | ORF7b | stopgain_E39X; nonsynonymous_E39K |
| 27882 | G | A | 1.27E-04 | | | | | ORF7b | nonsynonymous_A43T |
| 27849 | C | T | 1.27E-04 | | | | | ORF7b | nonsynonymous_L32F |
| 27893 | C | T | 1.27E-04 | | | | | intergenic | |
| 27895 | T | C | 1.27E-04 | | | | | ORF8 | nonsynonymous_M1T |
| 6869 | A | T | 1.27E-04 | | | | | ORF1ab | nonsynonymous_T2202S |
| 22363 | G | T | 1.27E-04 | | | | | S | synonymous_V267V |
| 22487 | G | C | 1.27E-04 | | | | | S | nonsynonymous_E309Q |
| 2832 | A | G | 1.27E-04 | | | | | ORF1ab | nonsynonymous_K856R |
| 22374 | A | G | 1.27E-04 | | | | | S | nonsynonymous_Q271R |
| 22388 | C | T | 1.27E-04 | | | | | S | synonymous_L276L |
| 2590 | A | G | 1.27E-04 | | | | | ORF1ab | synonymous_E775E |
| 29551 | A | G | 1.27E-04 | | | | | intergenic | |
| 29557 | G | T | 1.27E-04 | | | | | intergenic | |
| 21352 | A | G | 1.27E-04 | | | | | ORF1ab | nonsynonymous_R7030G |
| 21386 | C | T | 1.27E-04 | | | | | ORF1ab | nonsynonymous_S7041F |

| | | | | | | | | | |
|---|---|---|---|---|---|---|---|---|---|
| 29399 | G | T | 1.27E-04 | | | | | N | nonsynonymous_A376S |
| 29573 | G | A | 1.27E-04 | | | | | ORF10 | nonsynonymous_V6I |
| 5194 | G | T | 1.27E-04 | | | | | ORF1ab | synonymous_L1643L |
| 21156 | C | T | 1.27E-04 | | | | | ORF1ab | synonymous_S6964S |
| 21225 | G | T | 1.27E-04 | | | | | ORF1ab | nonsynonymous_W6987C |
| 21282 | A | G | 1.27E-04 | | | | | ORF1ab | synonymous_G7006G |
| 21294 | T | A | 1.27E-04 | | | | | ORF1ab | synonymous_L7010L |
| 21295 | G | A | 1.27E-04 | | | | | ORF1ab | nonsynonymous_G7011S |
| 21296 | G | A | 1.27E-04 | | | | | ORF1ab | nonsynonymous_G7011D |
| 21551 | A | T | 1.27E-04 | | | | | ORF1ab | nonsynonymous_N7096I |
| 2198 | G | A | 1.27E-04 | | | | | ORF1ab | nonsynonymous_G645S |
| 22458 | C | T | 1.27E-04 | | | | | S | nonsynonymous_T299I |
| 2652 | C | T | 1.27E-04 | | | | | ORF1ab | nonsynonymous_T796I |
| 2695 | C | T | 1.27E-04 | | | | | ORF1ab | synonymous_N810N |
| 2698 | T | C | 1.27E-04 | | | | | ORF1ab | synonymous_N811N |
| 2710 | C | G | 1.27E-04 | | | | | ORF1ab | synonymous_L815L |
| 2782 | G | T | 1.27E-04 | | | | | ORF1ab | synonymous_V839V |
| 29360 | T | C | 1.27E-04 | | | | | N | nonsynonymous_F363L |
| 29414 | G | A | 1.27E-04 | T | 6.33E-05 | | | N | nonsynonymous_A381T; nonsynonymous_A381S |
| 29446 | T | C | 1.27E-04 | | | | | N | synonymous_T391T |
| 29463 | C | T | 1.27E-04 | | | | | N | nonsynonymous_A397V |
| 29509 | C | T | 1.27E-04 | | | | | N | synonymous_S412S |
| 29381 | A | C | 1.27E-04 | | | | | N | nonsynonymous_K370Q |
| 29395 | G | C | 1.27E-04 | | | | | N | nonsynonymous_K374N |
| 20233 | C | T | 1.27E-04 | | | | | ORF1ab | nonsynonymous_P6657S |
| 20274 | G | T | 1.27E-04 | | | | | ORF1ab | nonsynonymous_M6670I |
| 20383 | G | T | 1.27E-04 | | | | | ORF1ab | nonsynonymous_A6707S |
| 20401 | T | G | 1.27E-04 | | | | | ORF1ab | nonsynonymous_S6713A |
| 20434 | G | T | 1.27E-04 | | | | | ORF1ab | nonsynonymous_D6724Y |

| | | | | | | | | | |
|---|---|---|---|---|---|---|---|---|---|
| 20477 | G | T | 1.27E-04 | | | | | ORF1ab | nonsynonymous_G6738V |
| 22436 | G | T | 1.27E-04 | | | | | S | nonsynonymous_A292S |
| 23116 | A | T | 1.27E-04 | | | | | S | synonymous_L518L |
| 2625 | G | A | 1.27E-04 | | | | | ORF1ab | nonsynonymous_G787E |
| 2638 | C | T | 1.27E-04 | | | | | ORF1ab | synonymous_L791L |
| 2641 | A | G | 1.27E-04 | | | | | ORF1ab | synonymous_E792E |
| 29362 | C | T | 1.27E-04 | | | | | N | synonymous_F363F |
| 29370 | C | T | 1.27E-04 | | | | | N | nonsynonymous_T366I |
| 29389 | A | G | 1.27E-04 | | | | | N | synonymous_K372K |
| 29413 | A | G | 1.27E-04 | | | | | N | synonymous_Q380Q |
| 29416 | C | T | 1.27E-04 | | | | | N | synonymous_A381A |
| 29462 | G | T | 1.27E-04 | C | 6.33E-05 | | | N | nonsynonymous_A397S; nonsynonymous_A397P |
| 29467 | A | T | 1.27E-04 | | | | | N | synonymous_A398A |
| 29482 | C | T | 1.27E-04 | | | | | N | synonymous_F403F |
| 29518 | C | T | 1.27E-04 | A | 6.33E-05 | | | N | synonymous_D415D; nonsynonymous_D415E |
| 29587 | G | A | 1.27E-04 | T | 6.33E-05 | T | 6.33E-05 | ORF10 | synonymous_P10P; synonymous_P10P; synonymous_P10P |
| 29642 | C | T | 1.27E-04 | A | 6.33E-05 | | | ORF10 | stopgain_Q29X; nonsynonymous_Q29K |
| 13384 | C | T | 1.27E-04 | | | | | ORF1ab | synonymous_C4373C |
| 20232 | A | G | 1.27E-04 | | | | | ORF1ab | synonymous_K6656K |
| 20235 | C | A | 1.27E-04 | A | 1.27E-04 | | | ORF1ab | synonymous_P6657P; synonymous_P6657P |
| 20238 | G | T | 1.27E-04 | | | | | ORF1ab | nonsynonymous_R6658S |
| 20377 | G | A | 1.27E-04 | | | | | ORF1ab | nonsynonymous_G6705R |
| 25720 | G | T | 1.27E-04 | | | | | ORF3a | nonsynonymous_A110S |
| 29511 | G | T | 1.27E-04 | | | | | N | nonsynonymous_S413I |

| | | | | | | | | | |
|---|---|---|---|---|---|---|---|---|---|
| 29623 | T | C | 1.27E-04 | | | | | ORF10 | synonymous_N22N |
| 29645 | G | T | 1.27E-04 | | | | | ORF10 | nonsynonymous_V30L |
| 29648 | G | T | 1.27E-04 | | | | | ORF10 | nonsynonymous_D31Y |
| 29657 | A | T | 1.27E-04 | | | | | ORF10 | nonsynonymous_N34Y |
| 6941 | C | T | 1.27E-04 | | | | | ORF1ab | synonymous_L2226L |
| 13548 | C | T | 1.27E-04 | | | | | ORF1ab | synonymous_D4428D |
| 19962 | G | T | 1.27E-04 | A | 6.33E-05 | C | 6.33E-05 | ORF1ab | synonymous_T6566T; synonymous_T6566T; synonymous_T6566T |
| 20276 | A | G | 1.27E-04 | | | | | ORF1ab | nonsynonymous_D6671G |
| 20281 | T | C | 1.27E-04 | | | | | ORF1ab | nonsynonymous_F6673L |
| 20283 | C | T | 1.27E-04 | | | | | ORF1ab | synonymous_F6673F |
| 20323 | A | G | 1.27E-04 | | | | | ORF1ab | nonsynonymous_I6687V |
| 20326 | G | A | 1.27E-04 | | | | | ORF1ab | nonsynonymous_V6688I |
| 20334 | A | G | 1.27E-04 | | | | | ORF1ab | synonymous_G6690G |
| 20371 | C | T | 1.27E-04 | A | 6.33E-05 | | | ORF1ab | synonymous_L6703L; nonsynonymous_L6703M |
| 22899 | G | T | 1.27E-04 | | | | | S | nonsynonymous_G446V |
| 25677 | G | T | 1.27E-04 | | | | | ORF3a | nonsynonymous_L95F |
| 25703 | C | T | 1.27E-04 | A | 6.33E-05 | | | ORF3a | nonsynonymous_P104L; nonsynonymous_P104H |
| 25724 | T | C | 1.27E-04 | | | | | ORF3a | nonsynonymous_L111S |
| 26533 | C | T | 1.27E-04 | | | | | M | nonsynonymous_S4F |
| 275 | C | A | 1.27E-04 | | | | | ORF1ab | nonsynonymous_L4I |
| 283 | T | C | 1.27E-04 | | | | | ORF1ab | synonymous_P6P |
| 29528 | G | T | 1.27E-04 | | | | | N | nonsynonymous_A419S |
| 29592 | C | T | 1.27E-04 | | | | | ORF10 | nonsynonymous_T12M |
| 5257 | A | G | 1.27E-04 | | | | | ORF1ab | synonymous_L1664L |
| 13426 | C | T | 1.26E-04 | | | | | ORF1ab | synonymous_R4387R |
| 13432 | C | T | 1.26E-04 | | | | | ORF1ab | synonymous_P4389P |

| | | | | | | | | | |
|---|---|---|---|---|---|---|---|---|---|
| 13459 | G | T | 1.26E-04 | | | | | ORF1ab | synonymous_S4398S |
| 13500 | A | G | 1.26E-04 | | | | | ORF1ab | synonymous_T4412T |
| 13503 | G | T | 1.26E-04 | A | 6.32E-05 | | | ORF1ab | synonymous_P4413P; synonymous_P4413P |
| 13515 | C | T | 1.26E-04 | | | | | ORF1ab | synonymous_G4417G |
| 13532 | T | C | 1.26E-04 | | | | | ORF1ab | nonsynonymous_V4423A |
| 15868 | C | T | 1.26E-04 | | | | | ORF1ab | nonsynonymous_H5202Y |
| 20170 | G | T | 1.26E-04 | | | | | ORF1ab | nonsynonymous_G6636C |
| 21676 | C | T | 1.26E-04 | | | | | S | synonymous_Y38Y |
| 23123 | C | T | 1.26E-04 | | | | | S | nonsynonymous_P521S |
| 23124 | C | G | 1.26E-04 | | | | | S | nonsynonymous_P521R |
| 25066 | C | T | 1.26E-04 | | | | | S | synonymous_D1168D |
| 25691 | G | T | 1.26E-04 | | | | | ORF3a | nonsynonymous_G100V |
| 25708 | C | T | 1.26E-04 | | | | | ORF3a | nonsynonymous_L106F |
| 25771 | C | A | 1.26E-04 | T | 6.32E-05 | | | ORF3a | nonsynonymous_L127I; nonsynonymous_L127F |
| 25776 | G | T | 1.26E-04 | | | | | ORF3a | nonsynonymous_W128C |
| 25827 | T | C | 1.26E-04 | | | | | ORF3a | synonymous_Y145Y |
| 25854 | C | T | 1.26E-04 | | | | | ORF3a | synonymous_Y154Y |
| 25859 | A | G | 1.26E-04 | | | | | ORF3a | nonsynonymous_Y156C |
| 280 | C | T | 1.26E-04 | | | | | ORF1ab | synonymous_V5V |
| 28902 | T | C | 1.26E-04 | | | | | N | nonsynonymous_M210T |
| 29244 | T | C | 1.26E-04 | | | | | N | nonsynonymous_V324A |
| 5554 | G | T | 1.26E-04 | A | 6.32E-05 | | | ORF1ab | nonsynonymous_K1763N; synonymous_K1763K |
| 8790 | G | T | 1.26E-04 | | | | | ORF1ab | nonsynonymous_G2842V |
| 19960 | A | G | 1.26E-04 | | | | | ORF1ab | nonsynonymous_T6566A |
| 20125 | G | A | 1.26E-04 | | | | | ORF1ab | nonsynonymous_G6621R |
| 21467 | T | C | 1.26E-04 | | | | | ORF1ab | nonsynonymous_M7068T |

| | | | | | | | | | |
|---|---|---|---|---|---|---|---|---|---|
| 21520 | G | T | 1.26E-04 | A | 6.32E-05 | | | ORF1ab | nonsynonymous_V7086F; nonsynonymous_V7086I |
| 21524 | T | G | 1.26E-04 | | | | | ORF1ab | nonsynonymous_V7087G |
| 21600 | G | T | 1.26E-04 | A | 6.32E-05 | | | S | nonsynonymous_S13I; nonsynonymous_S13N |
| 21604 | G | T | 1.26E-04 | | | | | S | nonsynonymous_Q14H |
| 21621 | C | T | 1.26E-04 | | | | | S | nonsynonymous_T20I |
| 21625 | A | G | 1.26E-04 | | | | | S | synonymous_R21R |
| 21628 | T | G | 1.26E-04 | | | | | S | synonymous_T22T |
| 21987 | G | A | 1.26E-04 | T | 6.32E-05 | C | 6.32E-05 | S | nonsynonymous_G142D; nonsynonymous_G142V; nonsynonymous_G142A |
| 21989 | G | T | 1.26E-04 | | | | | S | nonsynonymous_V143F |
| 22141 | G | A | 1.26E-04 | | | | | S | synonymous_V193V |
| 23009 | G | T | 1.26E-04 | A | 6.32E-05 | | | S | nonsynonymous_V483F; nonsynonymous_V483I |
| 23032 | T | C | 1.26E-04 | | | | | S | synonymous_F490F |
| 25641 | G | T | 1.26E-04 | C | 6.32E-05 | | | ORF3a | nonsynonymous_L83F; nonsynonymous_L83F |
| 271 | G | T | 1.26E-04 | | | | | ORF1ab | nonsynonymous_E2D |
| 285 | G | A | 1.26E-04 | T | 6.32E-05 | | | ORF1ab | nonsynonymous_G7D; nonsynonymous_G7V |
| 29067 | C | T | 1.26E-04 | | | | | N | nonsynonymous_T265I |
| 5338 | T | C | 1.26E-04 | | | | | ORF1ab | synonymous_N1691N |
| 6944 | A | G | 1.26E-04 | | | | | ORF1ab | nonsynonymous_I2227V |
| 11020 | C | T | 1.26E-04 | | | | | ORF1ab | synonymous_L3585L |
| 11272 | G | T | 1.26E-04 | | | | | ORF1ab | nonsynonymous_M3669I |
| 19572 | C | T | 1.26E-04 | | | | | ORF1ab | synonymous_Y6436Y |
| 20006 | G | T | 1.26E-04 | A | 6.32E-05 | | | ORF1ab | nonsynonymous_G6581V; nonsynonymous_G6581D |

| | | | | | | | | | |
|---|---|---|---|---|---|---|---|---|---|
| 20014 | G | T | 1.26E-04 | | | | | ORF1ab | nonsynonymous_D6584Y |
| 20064 | T | C | 1.26E-04 | | | | | ORF1ab | synonymous_V6600V |
| 20091 | C | T | 1.26E-04 | | | | | ORF1ab | synonymous_P6609P |
| 20097 | A | G | 1.26E-04 | | | | | ORF1ab | synonymous_Q6611Q |
| 20128 | G | A | 1.26E-04 | | | | | ORF1ab | nonsynonymous_E6622K |
| 20151 | T | C | 1.26E-04 | | | | | ORF1ab | synonymous_N6629N |
| 21781 | C | T | 1.26E-04 | | | | | S | synonymous_T73T |
| 21784 | T | A | 1.26E-04 | | | | | S | nonsynonymous_N74K |
| 22227 | C | T | 1.26E-04 | | | | | S | nonsynonymous_A222V |
| 22894 | G | A | 1.26E-04 | | | | | S | synonymous_K444K |
| 22922 | A | C | 1.26E-04 | | | | | S | synonymous_R454R |
| 22986 | C | T | 1.26E-04 | | | | | S | nonsynonymous_A475V |
| 25000 | C | T | 1.26E-04 | A | 6.32E-05 | | | S | synonymous_D1146D;<br>nonsynonymous_D1146E |
| 25055 | G | T | 1.26E-04 | | | | | S | nonsynonymous_D1165Y |
| 25056 | A | G | 1.26E-04 | | | | | S | nonsynonymous_D1165G |
| 25060 | A | G | 1.26E-04 | | | | | S | synonymous_L1166L |
| 25266 | G | T | 1.26E-04 | | | | | S | nonsynonymous_C1235F |
| 25275 | C | T | 1.26E-04 | | | | | S | nonsynonymous_T1238I |
| 25297 | G | A | 1.26E-04 | | | | | S | synonymous_K1245K |
| 25555 | G | T | 1.26E-04 | | | | | ORF3a | nonsynonymous_V55F |
| 25614 | C | T | 1.26E-04 | | | | | ORF3a | synonymous_S74S |
| 266 | A | T | 1.26E-04 | | | | | ORF1ab | nonsynonymous_M1L |
| 27103 | C | T | 1.26E-04 | | | | | M | nonsynonymous_A194V |
| 27110 | C | T | 1.26E-04 | | | | | M | synonymous_Y196Y |
| 27433 | A | G | 1.26E-04 | | | | | ORF7a | nonsynonymous_T14A |
| 27478 | G | T | 1.26E-04 | A | 6.32E-05 | | | ORF7a | nonsynonymous_V29L;<br>nonsynonymous_V29I |
| 27488 | A | T | 1.26E-04 | | | | | ORF7a | nonsynonymous_K32I |
| 27494 | C | T | 1.26E-04 | | | | | ORF7a | nonsynonymous_P34L |

| | | | | | | | | | |
|---|---|---|---|---|---|---|---|---|---|
| 28786 | C | T | 1.26E-04 | | | | | N | synonymous_F171F |
| 28884 | G | T | 1.26E-04 | | | | | N | nonsynonymous_G204V |
| 28923 | C | T | 1.26E-04 | | | | | N | nonsynonymous_A217V |
| 289 | C | T | 1.26E-04 | | | | | ORF1ab | synonymous_F8F |
| 29044 | G | A | 1.26E-04 | | | | | N | synonymous_K257K |
| 29050 | G | A | 1.26E-04 | | | | | N | synonymous_R259R |
| 29104 | T | C | 1.26E-04 | | | | | N | synonymous_R277R |
| 29177 | C | T | 1.26E-04 | | | | | N | nonsynonymous_P302S |
| 29194 | T | C | 1.26E-04 | | | | | N | synonymous_F307F |
| 29195 | G | T | 1.26E-04 | | | | | N | nonsynonymous_A308S |
| 29199 | C | T | 1.26E-04 | | | | | N | nonsynonymous_P309L |
| 29238 | T | C | 1.26E-04 | | | | | N | nonsynonymous_M322T |
| 29249 | C | T | 1.26E-04 | | | | | N | nonsynonymous_P326S |
| 29256 | G | T | 1.26E-04 | | | | | N | nonsynonymous_G328V |
| 29264 | T | C | 1.26E-04 | | | | | N | synonymous_L331L |
| 29269 | C | T | 1.26E-04 | | | | | N | synonymous_T332T |
| 29274 | C | T | 1.26E-04 | | | | | N | nonsynonymous_T334I |
| 29302 | T | C | 1.26E-04 | | | | | N | synonymous_D343D |
| 5425 | G | T | 1.26E-04 | | | | | ORF1ab | nonsynonymous_K1720N |
| 5497 | C | T | 1.26E-04 | | | | | ORF1ab | synonymous_C1744C |
| 5521 | T | C | 1.26E-04 | | | | | ORF1ab | synonymous_C1752C |
| 5561 | G | A | 1.26E-04 | | | | | ORF1ab | nonsynonymous_E1766K |
| 5575 | C | T | 1.26E-04 | | | | | ORF1ab | synonymous_Y1770Y |
| 842 | C | T | 1.26E-04 | | | | | ORF1ab | nonsynonymous_P193S |
| 1093 | C | T | 1.26E-04 | | | | | ORF1ab | synonymous_P276P |
| 13624 | G | T | 1.26E-04 | | | | | ORF1ab | nonsynonymous_D4454Y |
| 14277 | G | T | 1.26E-04 | | | | | ORF1ab | nonsynonymous_R4671S |
| 16064 | A | G | 1.26E-04 | | | | | ORF1ab | nonsynonymous_Q5267R |
| 16716 | C | T | 1.26E-04 | | | | | ORF1ab | synonymous_D5484D |

| | | | | | | | | | |
|---|---|---|---|---|---|---|---|---|---|
| 16733 | C | T | 1.26E-04 | A | 6.32E-05 | | | ORF1ab | nonsynonymous_S5490L; stopgain_S5490X |
| 16949 | C | T | 1.26E-04 | | | | | ORF1ab | nonsynonymous_P5562L |
| 17079 | G | T | 1.26E-04 | | | | | ORF1ab | nonsynonymous_Q5605H |
| 18176 | C | T | 1.26E-04 | | | | | ORF1ab | nonsynonymous_P5971L |
| 18246 | C | T | 1.26E-04 | | | | | ORF1ab | synonymous_Y5994Y |
| 20569 | G | T | 1.26E-04 | | | | | ORF1ab | nonsynonymous_V6769F |
| 20582 | C | T | 1.26E-04 | | | | | ORF1ab | nonsynonymous_T6773I |
| 20594 | C | T | 1.26E-04 | | | | | ORF1ab | nonsynonymous_T6777I |
| 20860 | A | G | 1.26E-04 | | | | | ORF1ab | nonsynonymous_I6866V |
| 20893 | G | T | 1.26E-04 | | | | | ORF1ab | nonsynonymous_A6877S |
| 21391 | T | G | 1.26E-04 | | | | | ORF1ab | nonsynonymous_F7043V |
| 21417 | A | T | 1.26E-04 | | | | | ORF1ab | nonsynonymous_K7051N |
| 22017 | G | T | 1.26E-04 | | | | | S | nonsynonymous_W152L |
| 22032 | T | C | 1.26E-04 | | | | | S | nonsynonymous_F157S |
| 22124 | A | G | 1.26E-04 | | | | | S | nonsynonymous_N188D |
| 22132 | G | A | 1.26E-04 | | | | | S | synonymous_R190R |
| 22246 | T | C | 1.26E-04 | | | | | S | synonymous_D228D |
| 22264 | C | T | 1.26E-04 | | | | | S | synonymous_N234N |
| 22290 | C | T | 1.26E-04 | | | | | S | nonsynonymous_A243V |
| 22624 | C | T | 1.26E-04 | A | 6.32E-05 | | | S | synonymous_N354N; nonsynonymous_N354K |
| 23592 | A | G | 1.26E-04 | | | | | S | nonsynonymous_Q677R |
| 23683 | C | T | 1.26E-04 | | | | | S | synonymous_Y707Y |
| 24102 | G | A | 1.26E-04 | A | 1.26E-04 | | | S | nonsynonymous_R847K; nonsynonymous_R847K |
| 24193 | G | T | 1.26E-04 | | | | | S | synonymous_L877L |
| 24797 | C | T | 1.26E-04 | A | 6.32E-05 | | | S | nonsynonymous_P1079S; nonsynonymous_P1079T |
| 24811 | T | A | 1.26E-04 | | | | | S | nonsynonymous_H1083Q |

| | | | | | | | | | |
|---|---|---|---|---|---|---|---|---|---|
| 24867 | G | T | 1.26E-04 | C | 6.32E-05 | | | S | nonsynonymous_W1102L; nonsynonymous_W1102S |
| 24872 | G | T | 1.26E-04 | A | 6.32E-05 | | | S | nonsynonymous_V1104L; nonsynonymous_V1104I |
| 24887 | T | C | 1.26E-04 | | | | | S | nonsynonymous_F1109L |
| 24910 | T | C | 1.26E-04 | | | | | S | synonymous_T1116T |
| 24923 | T | C | 1.26E-04 | | | | | S | nonsynonymous_F1121L |
| 24952 | A | G | 1.26E-04 | | | | | S | nonsynonymous_I1130M |
| 24990 | C | T | 1.26E-04 | | | | | S | nonsynonymous_P1143L |
| 25049 | G | T | 1.26E-04 | | | | | S | nonsynonymous_D1163Y |
| 25106 | G | C | 1.26E-04 | | | | | S | nonsynonymous_E1182Q |
| 25162 | C | A | 1.26E-04 | G | 6.32E-05 | | | S | synonymous_L1200L; synonymous_L1200L |
| 25163 | C | A | 1.26E-04 | T | 6.32E-05 | | | S | nonsynonymous_Q1201K; stopgain_Q1201X |
| 25187 | T | C | 1.26E-04 | | | | | S | nonsynonymous_Y1209H |
| 25209 | T | C | 1.26E-04 | | | | | S | nonsynonymous_I1216T |
| 25250 | G | T | 1.26E-04 | | | | | S | nonsynonymous_V1230L |
| 25252 | G | A | 1.26E-04 | A | 1.26E-04 | C | 6.32E-05 | S | synonymous_V1230V; synonymous_V1230V; synonymous_V1230V |
| 25396 | G | T | 1.26E-04 | | | | | ORF3a | nonsynonymous_D2Y |
| 25459 | G | T | 1.26E-04 | | | | | ORF3a | nonsynonymous_A23S |
| 25568 | C | A | 1.26E-04 | | | | | ORF3a | nonsynonymous_A59D |
| 25609 | C | T | 1.26E-04 | | | | | ORF3a | nonsynonymous_L73F |
| 25611 | C | T | 1.26E-04 | | | | | ORF3a | synonymous_L73L |
| 25612 | T | G | 1.26E-04 | | | | | ORF3a | nonsynonymous_S74A |
| 25615 | A | G | 1.26E-04 | | | | | ORF3a | nonsynonymous_K75E |
| 26227 | G | A | 1.26E-04 | | | | | intergenic | |
| 26389 | G | T | 1.26E-04 | | | | | E | nonsynonymous_V49L |

| | | | | | | | | | |
|---|---|---|---|---|---|---|---|---|---|
| 26408 | C | T | 1.26E-04 | | | | | E | nonsynonymous_S55F |
| 26450 | G | T | 1.26E-04 | | | | | E | nonsynonymous_R69I |
| 26499 | C | G | 1.26E-04 | | | | | intergenic | |
| 26973 | A | G | 1.26E-04 | | | | | M | nonsynonymous_I151V |
| 27020 | A | G | 1.26E-04 | | | | | M | synonymous_K166K |
| 27038 | A | T | 1.26E-04 | G | 6.32E-05 | | | M | synonymous_T172T; synonymous_T172T |
| 27145 | C | T | 1.26E-04 | | | | | M | nonsynonymous_T208I |
| 27296 | T | C | 1.26E-04 | | | | | ORF6 | nonsynonymous_I32T |
| 27426 | A | G | 1.26E-04 | | | | | ORF7a | synonymous_T11T |
| 28233 | G | A | 1.26E-04 | A | 1.26E-04 | | | ORF8 | nonsynonymous_V114I; nonsynonymous_V114I |
| 28245 | T | G | 1.26E-04 | | | | | ORF8 | nonsynonymous_L118V |
| 28377 | C | T | 1.26E-04 | | | | | N | nonsynonymous_A35V |
| 28758 | C | T | 1.26E-04 | | | | | N | nonsynonymous_P162L |
| 28765 | A | G | 1.26E-04 | | | | | N | synonymous_G164G |
| 28776 | C | A | 1.26E-04 | T | 6.32E-05 | | | N | nonsynonymous_P168Q; nonsynonymous_P168L |
| 28789 | C | T | 1.26E-04 | | | | | N | synonymous_Y172Y |
| 28791 | C | T | 1.26E-04 | | | | | N | nonsynonymous_A173V |
| 28807 | C | T | 1.26E-04 | | | | | N | synonymous_G178G |
| 28855 | A | G | 1.26E-04 | | | | | N | synonymous_S194S |
| 28860 | A | T | 1.26E-04 | | | | | N | nonsynonymous_N196I |
| 28867 | T | C | 1.26E-04 | | | | | N | synonymous_T198T |
| 28871 | G | A | 1.26E-04 | | | | | N | nonsynonymous_G200S |
| 28932 | C | T | 1.26E-04 | | | | | N | nonsynonymous_A220V |
| 28954 | G | A | 1.26E-04 | | | | | N | synonymous_L227L |
| 28955 | A | T | 1.26E-04 | G | 6.32E-05 | | | N | nonsynonymous_N228Y; nonsynonymous_N228D |
| 28978 | T | A | 1.26E-04 | | | | | N | synonymous_S235S |

| | | | | | | | | | |
|---|---|---|---|---|---|---|---|---|---|
| 300 | C | T | 1.26E-04 | | | | | ORF1ab | nonsynonymous_T12I |
| 301 | A | G | 1.26E-04 | | | | | ORF1ab | synonymous_T12T |
| 5008 | G | A | 1.26E-04 | T | 6.32E-05 | | | ORF1ab | synonymous_T1581T; synonymous_T1581T |
| 5036 | G | A | 1.26E-04 | | | | | ORF1ab | nonsynonymous_G1591R |
| 5040 | A | G | 1.26E-04 | | | | | ORF1ab | nonsynonymous_Q1592R |
| 5094 | A | G | 1.26E-04 | | | | | ORF1ab | nonsynonymous_H1610R |
| 5098 | T | C | 1.26E-04 | | | | | ORF1ab | synonymous_N1611N |
| 5140 | C | T | 1.26E-04 | | | | | ORF1ab | synonymous_D1625D |
| 5155 | G | A | 1.26E-04 | | | | | ORF1ab | synonymous_E1630E |
| 5223 | C | T | 1.26E-04 | | | | | ORF1ab | nonsynonymous_T1653I |
| 5253 | G | A | 1.26E-04 | | | | | ORF1ab | nonsynonymous_G1663D |
| 5411 | G | T | 1.26E-04 | | | | | ORF1ab | nonsynonymous_A1716S |
| 5413 | C | T | 1.26E-04 | | | | | ORF1ab | synonymous_A1716A |
| 5457 | C | T | 1.26E-04 | A | 6.32E-05 | | | ORF1ab | nonsynonymous_T1731I; nonsynonymous_T1731K |
| 5518 | G | T | 1.26E-04 | | | | | ORF1ab | synonymous_V1751V |
| 5539 | G | A | 1.26E-04 | | | | | ORF1ab | synonymous_Q1758Q |
| 5544 | C | T | 1.26E-04 | | | | | ORF1ab | nonsynonymous_T1760I |
| 6364 | G | T | 1.26E-04 | A | 6.32E-05 | | | ORF1ab | synonymous_A2033A; synonymous_A2033A |
| 7038 | G | A | 1.26E-04 | | | | | ORF1ab | nonsynonymous_G2258D |
| 7043 | C | T | 1.26E-04 | | | | | ORF1ab | nonsynonymous_P2260S |
| 7312 | G | C | 1.26E-04 | A | 6.32E-05 | | | ORF1ab | nonsynonymous_L2349F; synonymous_L2349L |
| 10834 | C | T | 1.26E-04 | | | | | ORF1ab | synonymous_A3523A |
| 10836 | T | C | 1.26E-04 | | | | | ORF1ab | nonsynonymous_V3524A |
| 13656 | T | C | 1.26E-04 | | | | | ORF1ab | synonymous_V4464V |
| 13667 | C | T | 1.26E-04 | | | | | ORF1ab | nonsynonymous_T4468I |
| 13921 | G | T | 1.26E-04 | | | | | ORF1ab | nonsynonymous_D4553Y |

| | | | | | | | | | |
|---|---|---|---|---|---|---|---|---|---|
| 14230 | C | T | 1.26E-04 | | | | | ORF1ab | nonsynonymous_P4656S |
| 14371 | G | T | 1.26E-04 | | | | | ORF1ab | nonsynonymous_A4703S |
| 14511 | G | T | 1.26E-04 | | | | | ORF1ab | nonsynonymous_Q4749H |
| 14583 | C | T | 1.26E-04 | | | | | ORF1ab | synonymous_H4773H |
| 14593 | G | A | 1.26E-04 | | | | | ORF1ab | nonsynonymous_G4777S |
| 14657 | C | T | 1.26E-04 | | | | | ORF1ab | nonsynonymous_A4798V |
| 14831 | G | T | 1.26E-04 | | | | | ORF1ab | nonsynonymous_C4856F |
| 14874 | G | T | 1.26E-04 | | | | | ORF1ab | nonsynonymous_K4870N |
| 15444 | G | T | 1.26E-04 | | | | | ORF1ab | nonsynonymous_M5060I |
| 15579 | C | T | 1.26E-04 | | | | | ORF1ab | synonymous_N5105N |
| 15834 | A | G | 1.26E-04 | | | | | ORF1ab | synonymous_K5190K |
| 15852 | C | T | 1.26E-04 | | | | | ORF1ab | synonymous_D5196D |
| 15895 | C | T | 1.26E-04 | | | | | ORF1ab | synonymous_L5211L |
| 16095 | A | G | 1.26E-04 | | | | | ORF1ab | synonymous_L5277L |
| 16197 | G | T | 1.26E-04 | | | | | ORF1ab | nonsynonymous_E5311D |
| 16489 | G | T | 1.26E-04 | | | | | ORF1ab | nonsynonymous_A5409S |
| 16534 | A | G | 1.26E-04 | | | | | ORF1ab | nonsynonymous_S5424G |
| 16568 | C | T | 1.26E-04 | | | | | ORF1ab | nonsynonymous_T5435I |
| 16579 | A | G | 1.26E-04 | | | | | ORF1ab | nonsynonymous_T5439A |
| 16580 | C | T | 1.26E-04 | | | | | ORF1ab | nonsynonymous_T5439I |
| 16617 | T | C | 1.26E-04 | | | | | ORF1ab | synonymous_T5451T |
| 16628 | A | G | 1.26E-04 | | | | | ORF1ab | nonsynonymous_K5455R |
| 16647 | G | T | 1.26E-04 | | | | | ORF1ab | synonymous_T5461T |
| 16671 | T | C | 1.26E-04 | | | | | ORF1ab | synonymous_F5469F |
| 16857 | T | C | 1.26E-04 | | | | | ORF1ab | synonymous_D5531D |
| 17285 | C | T | 1.26E-04 | | | | | ORF1ab | nonsynonymous_S5674L |
| 17528 | C | T | 1.26E-04 | | | | | ORF1ab | nonsynonymous_T5755I |
| 17573 | C | T | 1.26E-04 | | | | | ORF1ab | nonsynonymous_A5770V |
| 17686 | G | T | 1.26E-04 | | | | | ORF1ab | nonsynonymous_V5808F |
| 18114 | A | G | 1.26E-04 | | | | | ORF1ab | synonymous_T5950T |

| | | | | | | | | | |
|---|---|---|---|---|---|---|---|---|---|
| 18149 | G | T | 1.26E-04 | | | | | ORF1ab | nonsynonymous_G5962V |
| 18153 | A | G | 1.26E-04 | | | | | ORF1ab | synonymous_L5963L |
| 18213 | G | T | 1.26E-04 | | | | | ORF1ab | nonsynonymous_M5983I |
| 18351 | T | C | 1.26E-04 | | | | | ORF1ab | synonymous_N6029N |
| 18452 | C | T | 1.26E-04 | | | | | ORF1ab | nonsynonymous_A6063V |
| 18600 | A | T | 1.26E-04 | | | | | ORF1ab | synonymous_A6112A |
| 18674 | G | A | 1.26E-04 | | | | | ORF1ab | nonsynonymous_R6137K |
| 18677 | G | T | 1.26E-04 | | | | | ORF1ab | nonsynonymous_R6138L |
| 18681 | C | T | 1.26E-04 | | | | | ORF1ab | synonymous_A6139A |
| 18688 | T | C | 1.26E-04 | | | | | ORF1ab | nonsynonymous_F6142L |
| 18710 | A | G | 1.26E-04 | | | | | ORF1ab | nonsynonymous_Y6149C |
| 18747 | C | T | 1.26E-04 | | | | | ORF1ab | synonymous_V6161V |
| 18802 | A | G | 1.26E-04 | | | | | ORF1ab | nonsynonymous_S6180G |
| 18938 | G | A | 1.26E-04 | | | | | ORF1ab | nonsynonymous_G6225D |
| 19118 | C | T | 1.26E-04 | | | | | ORF1ab | nonsynonymous_A6285V |
| 19665 | T | A | 1.26E-04 | | | | | ORF1ab | nonsynonymous_F6467L |
| 19667 | A | G | 1.26E-04 | | | | | ORF1ab | nonsynonymous_D6468G |
| 20615 | G | T | 1.26E-04 | | | | | ORF1ab | nonsynonymous_W6784L |
| 20670 | G | A | 1.26E-04 | | | | | ORF1ab | synonymous_A6802A |
| 20787 | T | C | 1.26E-04 | | | | | ORF1ab | synonymous_N6841N |
| 20801 | C | T | 1.26E-04 | | | | | ORF1ab | nonsynonymous_T6846I |
| 20925 | G | T | 1.26E-04 | | | | | ORF1ab | nonsynonymous_L6887F |
| 21051 | G | A | 1.26E-04 | | | | | ORF1ab | nonsynonymous_M6929I |
| 21694 | A | G | 1.26E-04 | | | | | S | synonymous_R44R |
| 21698 | T | G | 1.26E-04 | | | | | S | nonsynonymous_S46A |
| 21793 | G | T | 1.26E-04 | C | 6.32E-05 | | | S | nonsynonymous_K77N; nonsynonymous_K77N |
| 21811 | C | T | 1.26E-04 | | | | | S | synonymous_V83V |
| 22000 | C | T | 1.26E-04 | | | | | S | synonymous_H146H |

| | | | | | | | | | |
|---|---|---|---|---|---|---|---|---|---|
| 22014 | G | T | 1.26E-04 | A | 6.32E-05 | | | S | nonsynonymous_S151I; nonsynonymous_S151N |
| 22033 | C | A | 1.26E-04 | | | | | S | nonsynonymous_F157L |
| 22037 | G | A | 1.26E-04 | | | | | S | nonsynonymous_V159I |
| 22042 | T | C | 1.26E-04 | | | | | S | synonymous_Y160Y |
| 22054 | T | C | 1.26E-04 | | | | | S | synonymous_N164N |
| 22096 | C | T | 1.26E-04 | | | | | S | synonymous_D178D |
| 22101 | A | T | 1.26E-04 | | | | | S | nonsynonymous_E180V |
| 22155 | A | G | 1.26E-04 | | | | | S | nonsynonymous_D198G |
| 22206 | A | G | 1.26E-04 | | | | | S | nonsynonymous_D215G |
| 22208 | C | T | 1.26E-04 | | | | | S | nonsynonymous_L216F |
| 22213 | T | C | 1.26E-04 | | | | | S | synonymous_P217P |
| 22319 | G | T | 1.26E-04 | | | | | S | nonsynonymous_D253Y |
| 22574 | T | C | 1.26E-04 | | | | | S | nonsynonymous_F338L |
| 22622 | A | G | 1.26E-04 | | | | | S | nonsynonymous_N354D |
| 23202 | C | T | 1.26E-04 | | | | | S | nonsynonymous_T547I |
| 23520 | C | T | 1.26E-04 | | | | | S | nonsynonymous_A653V |
| 23569 | T | C | 1.26E-04 | | | | | S | synonymous_G669G |
| 23599 | T | C | 1.26E-04 | A | 6.32E-05 | | | S | synonymous_N679N; nonsynonymous_N679K |
| 23605 | T | G | 1.26E-04 | | | | | S | synonymous_P681P |
| 23610 | G | A | 1.26E-04 | C | 6.32E-05 | | | S | nonsynonymous_R683Q; nonsynonymous_R683P |
| 23611 | G | A | 1.26E-04 | | | | | S | synonymous_R683R |
| 23613 | C | T | 1.26E-04 | | | | | S | nonsynonymous_A684V |
| 23625 | C | T | 1.26E-04 | | | | | S | nonsynonymous_A688V |
| 23986 | T | C | 1.26E-04 | | | | | S | synonymous_D808D |
| 24000 | G | T | 1.26E-04 | | | | | S | nonsynonymous_S813I |
| 24173 | G | T | 1.26E-04 | | | | | S | nonsynonymous_A871S |
| 24199 | G | T | 1.26E-04 | | | | | S | synonymous_A879A |

| | | | | | | | | | |
|---|---|---|---|---|---|---|---|---|---|
| 24213 | C | T | 1.26E-04 | | | | | S | nonsynonymous_S884F |
| 24237 | C | T | 1.26E-04 | | | | | S | nonsynonymous_A892V |
| 24240 | C | T | 1.26E-04 | | | | | S | nonsynonymous_A893V |
| 24257 | G | T | 1.26E-04 | | | | | S | nonsynonymous_A899S |
| 24289 | T | C | 1.26E-04 | | | | | S | synonymous_I909I |
| 24337 | C | T | 1.26E-04 | | | | | S | synonymous_N925N |
| 24410 | G | C | 1.26E-04 | | | | | S | nonsynonymous_D950H |
| 24484 | A | T | 1.26E-04 | | | | | S | synonymous_S974S |
| 24488 | G | T | 1.26E-04 | | | | | S | nonsynonymous_V976F |
| 24628 | T | C | 1.26E-04 | | | | | S | synonymous_A1022A |
| 24667 | A | G | 1.26E-04 | | | | | S | synonymous_G1035G |
| 24710 | A | G | 1.26E-04 | | | | | S | nonsynonymous_M1050V |
| 25050 | A | G | 1.26E-04 | | | | | S | nonsynonymous_D1163G |
| 25477 | G | T | 1.26E-04 | | | | | ORF3a | nonsynonymous_V29F |
| 25526 | G | T | 1.26E-04 | | | | | ORF3a | nonsynonymous_W45L |
| 25571 | C | T | 1.26E-04 | | | | | ORF3a | nonsynonymous_S60F |
| 25598 | G | T | 1.26E-04 | | | | | ORF3a | nonsynonymous_W69L |
| 25946 | A | G | 1.26E-04 | | | | | ORF3a | nonsynonymous_Q185R |
| 25976 | C | A | 1.26E-04 | | | | | ORF3a | nonsynonymous_S195Y |
| 26507 | A | T | 1.26E-04 | | | | | intergenic | |
| 26754 | G | T | 1.26E-04 | | | | | M | nonsynonymous_G78C |
| 26828 | G | T | 1.26E-04 | | | | | M | synonymous_L102L |
| 26945 | A | T | 1.26E-04 | | | | | M | synonymous_G141G |
| 26972 | T | C | 1.26E-04 | | | | | M | synonymous_R150R |
| 26995 | G | A | 1.26E-04 | | | | | M | nonsynonymous_R158H |
| 27011 | C | T | 1.26E-04 | | | | | M | synonymous_D163D |
| 27012 | C | T | 1.26E-04 | A | 6.32E-05 | | | M | synonymous_L164L; nonsynonymous_L164M |
| 27044 | A | C | 1.26E-04 | | | | | M | synonymous_R174R |
| 27201 | G | T | 1.26E-04 | | | | | intergenic | |

| | | | | | | | | | |
|---|---|---|---|---|---|---|---|---|---|
| 27247 | C | T | 1.26E-04 | | | | | ORF6 | synonymous_L16L |
| 27297 | C | T | 1.26E-04 | | | | | ORF6 | synonymous_I32I |
| 27335 | C | T | 1.26E-04 | | | | | ORF6 | nonsynonymous_T45I |
| 27358 | G | T | 1.26E-04 | | | | | ORF6 | nonsynonymous_D53Y |
| 27390 | G | C | 1.26E-04 | T | 6.32E-05 | | | intergenic | |
| 27393 | C | T | 1.26E-04 | | | | | intergenic | |
| 27398 | A | G | 1.26E-04 | | | | | ORF7a | nonsynonymous_K2R |
| 27406 | C | T | 1.26E-04 | | | | | ORF7a | nonsynonymous_L5F |
| 27813 | C | T | 1.26E-04 | | | | | ORF7b | nonsynonymous_L20F |
| 28308 | C | T | 1.26E-04 | | | | | N | nonsynonymous_A12V |
| 28315 | C | T | 1.26E-04 | | | | | N | synonymous_R14R |
| 28362 | G | T | 1.26E-04 | | | | | N | nonsynonymous_G30V |
| 28393 | T | C | 1.26E-04 | | | | | N | synonymous_R40R |
| 28435 | C | T | 1.26E-04 | | | | | N | synonymous_T54T |
| 28646 | G | A | 1.26E-04 | | | | | N | nonsynonymous_A125T |
| 28651 | C | T | 1.26E-04 | | | | | N | synonymous_N126N |
| 29347 | C | T | 1.26E-04 | | | | | N | synonymous_D358D |
| 319 | G | A | 1.26E-04 | | | | | ORF1ab | synonymous_L18L |
| 327 | T | C | 1.26E-04 | | | | | ORF1ab | nonsynonymous_L21S |
| 3873 | T | C | 1.26E-04 | | | | | ORF1ab | nonsynonymous_I1203T |
| 4011 | T | C | 1.26E-04 | | | | | ORF1ab | nonsynonymous_L1249P |
| 6309 | G | A | 1.26E-04 | T | 6.32E-05 | | | ORF1ab | nonsynonymous_S2015N; nonsynonymous_S2015I |
| 6896 | C | T | 1.26E-04 | | | | | ORF1ab | synonymous_L2211L |
| 7065 | G | A | 1.26E-04 | | | | | ORF1ab | nonsynonymous_R2267K |
| 7084 | T | C | 1.26E-04 | | | | | ORF1ab | synonymous_S2273S |
| 7302 | T | C | 1.26E-04 | | | | | ORF1ab | nonsynonymous_I2346T |
| 8818 | C | T | 1.26E-04 | | | | | ORF1ab | synonymous_C2851C |
| 8855 | T | C | 1.26E-04 | | | | | ORF1ab | nonsynonymous_F2864L |
| 9333 | C | T | 1.26E-04 | | | | | ORF1ab | nonsynonymous_A3023V |

| | | | | | | | | | |
|---|---|---|---|---|---|---|---|---|---|
| 9360 | C | T | 1.26E-04 | | | | | ORF1ab | nonsynonymous_T3032I |
| 9419 | A | G | 1.26E-04 | | | | | ORF1ab | nonsynonymous_I3052V |
| 9429 | T | C | 1.26E-04 | | | | | ORF1ab | nonsynonymous_I3055T |
| 9483 | A | G | 1.26E-04 | | | | | ORF1ab | nonsynonymous_E3073G |
| 10156 | C | T | 1.26E-04 | | | | | ORF1ab | synonymous_D3297D |
| 10615 | C | T | 1.26E-04 | | | | | ORF1ab | synonymous_D3450D |
| 10776 | C | T | 1.26E-04 | | | | | ORF1ab | nonsynonymous_P3504L |
| 10828 | A | C | 1.26E-04 | | | | | ORF1ab | synonymous_G3521G |
| 10830 | T | C | 1.26E-04 | | | | | ORF1ab | nonsynonymous_I3522T |
| 10912 | A | G | 1.26E-04 | | | | | ORF1ab | synonymous_L3549L |
| 10958 | G | T | 1.26E-04 | | | | | ORF1ab | nonsynonymous_G3565C |
| 10965 | C | T | 1.26E-04 | | | | | ORF1ab | nonsynonymous_T3567I |
| 10979 | G | A | 1.26E-04 | | | | | ORF1ab | nonsynonymous_V3572M |
| 11008 | C | T | 1.26E-04 | | | | | ORF1ab | synonymous_H3581H |
| 12141 | C | A | 1.26E-04 | | | | | ORF1ab | nonsynonymous_T3959N |
| 13175 | A | G | 1.26E-04 | | | | | ORF1ab | nonsynonymous_T4304A |
| 13201 | G | T | 1.26E-04 | | | | | ORF1ab | synonymous_P4312P |
| 13210 | T | C | 1.26E-04 | G | 6.32E-05 | | | ORF1ab | synonymous_N4315N; nonsynonymous_N4315K |
| 13225 | C | T | 1.26E-04 | G | 6.32E-05 | | | ORF1ab | synonymous_S4320S; synonymous_S4320S |
| 13273 | T | C | 1.26E-04 | | | | | ORF1ab | synonymous_H4336H |
| 13308 | A | G | 1.26E-04 | | | | | ORF1ab | nonsynonymous_K4348R |
| 13335 | C | T | 1.26E-04 | | | | | ORF1ab | nonsynonymous_A4357V |
| 13338 | A | G | 1.26E-04 | | | | | ORF1ab | nonsynonymous_N4358S |
| 13348 | G | T | 1.26E-04 | | | | | ORF1ab | synonymous_V4361V |
| 13366 | C | T | 1.26E-04 | | | | | ORF1ab | synonymous_N4367N |
| 13690 | G | A | 1.26E-04 | | | | | ORF1ab | nonsynonymous_E4476K |
| 13723 | G | A | 1.26E-04 | T | 6.32E-05 | | | ORF1ab | nonsynonymous_A4487T; nonsynonymous_A4487S |

| | | | | | | | | | |
|---|---|---|---|---|---|---|---|---|---|
| 13812 | G | A | 1.26E-04 | | | | | ORF1ab | nonsynonymous_M4516I |
| 13835 | G | T | 1.26E-04 | | | | | ORF1ab | nonsynonymous_R4524M |
| 13845 | T | C | 1.26E-04 | | | | | ORF1ab | synonymous_D4527D |
| 13902 | T | C | 1.26E-04 | | | | | ORF1ab | synonymous_D4546D |
| 13957 | C | T | 1.26E-04 | G | 6.32E-05 | | | ORF1ab | nonsynonymous_R4565C; nonsynonymous_R4565G |
| 13960 | G | A | 1.26E-04 | | | | | ORF1ab | nonsynonymous_V4566I |
| 13998 | G | A | 1.26E-04 | | | | | ORF1ab | synonymous_L4578L |
| 14012 | A | T | 1.26E-04 | | | | | ORF1ab | nonsynonymous_Q4583L |
| 14177 | C | T | 1.26E-04 | | | | | ORF1ab | nonsynonymous_T4638I |
| 14185 | A | T | 1.26E-04 | | | | | ORF1ab | nonsynonymous_R4641W |
| 14186 | G | A | 1.26E-04 | T | 6.32E-05 | | | ORF1ab | nonsynonymous_R4641K; nonsynonymous_R4641M |
| 14259 | T | C | 1.26E-04 | | | | | ORF1ab | synonymous_Y4665Y |
| 14270 | A | C | 1.26E-04 | | | | | ORF1ab | nonsynonymous_E4669A |
| 14274 | G | T | 1.26E-04 | T | 1.26E-04 | | | ORF1ab | nonsynonymous_E4670D; nonsynonymous_E4670D |
| 14313 | T | C | 1.26E-04 | | | | | ORF1ab | synonymous_D4683D |
| 14316 | G | T | 1.26E-04 | | | | | ORF1ab | nonsynonymous_Q4684H |
| 14350 | G | T | 1.26E-04 | | | | | ORF1ab | nonsynonymous_D4696Y |
| 14391 | C | T | 1.26E-04 | | | | | ORF1ab | synonymous_F4709F |
| 14396 | C | T | 1.26E-04 | | | | | ORF1ab | nonsynonymous_T4711I |
| 14554 | C | T | 1.26E-04 | | | | | ORF1ab | nonsynonymous_L4764F |
| 14697 | C | T | 1.26E-04 | | | | | ORF1ab | synonymous_F4811F |
| 14768 | C | T | 1.26E-04 | | | | | ORF1ab | nonsynonymous_A4835V |
| 14772 | G | T | 1.26E-04 | | | | | ORF1ab | nonsynonymous_Q4836H |
| 14773 | G | A | 1.26E-04 | T | 6.32E-05 | | | ORF1ab | nonsynonymous_D4837N; nonsynonymous_D4837Y |
| 14784 | T | C | 1.26E-04 | | | | | ORF1ab | synonymous_A4840A |
| 14817 | T | C | 1.26E-04 | | | | | ORF1ab | synonymous_N4851N |

| | | | | | | | | | |
|---|---|---|---|---|---|---|---|---|---|
| 14836 | A | G | 1.26E-04 | | | | | ORF1ab | nonsynonymous_I4858V |
| 14988 | T | C | 1.26E-04 | | | | | ORF1ab | synonymous_Y4908Y |
| 15026 | C | T | 1.26E-04 | | | | | ORF1ab | nonsynonymous_A4921V |
| 15048 | C | A | 1.26E-04 | T | 6.32E-05 | | | ORF1ab | synonymous_I4928I; synonymous_I4928I |
| 15166 | T | C | 1.26E-04 | | | | | ORF1ab | synonymous_L4968L |
| 15194 | C | T | 1.26E-04 | | | | | ORF1ab | nonsynonymous_A4977V |
| 15199 | G | A | 1.26E-04 | | | | | ORF1ab | nonsynonymous_V4979I |
| 15202 | G | T | 1.26E-04 | | | | | ORF1ab | nonsynonymous_V4980L |
| 15336 | T | C | 1.26E-04 | | | | | ORF1ab | synonymous_I5024I |
| 15372 | G | T | 1.26E-04 | | | | | ORF1ab | synonymous_T5036T |
| 15418 | G | T | 1.26E-04 | | | | | ORF1ab | nonsynonymous_A5052S |
| 15450 | C | T | 1.26E-04 | | | | | ORF1ab | synonymous_G5062G |
| 15451 | G | A | 1.26E-04 | | | | | ORF1ab | nonsynonymous_G5063S |
| 15504 | T | G | 1.26E-04 | | | | | ORF1ab | synonymous_A5080A |
| 15555 | T | C | 1.26E-04 | | | | | ORF1ab | synonymous_N5097N |
| 15593 | A | G | 1.26E-04 | | | | | ORF1ab | nonsynonymous_K5110R |
| 15594 | G | A | 1.26E-04 | T | 6.32E-05 | | | ORF1ab | synonymous_K5110K; nonsynonymous_K5110N |
| 15613 | C | T | 1.26E-04 | | | | | ORF1ab | nonsynonymous_H5117Y |
| 15657 | A | G | 1.26E-04 | | | | | ORF1ab | synonymous_T5131T |
| 15752 | C | T | 1.26E-04 | | | | | ORF1ab | nonsynonymous_A5163V |
| 15779 | A | G | 1.26E-04 | G | 1.26E-04 | | | ORF1ab | nonsynonymous_K5172R; nonsynonymous_K5172R |
| 15804 | T | C | 1.26E-04 | | | | | ORF1ab | synonymous_Y5180Y |
| 15952 | C | T | 1.26E-04 | | | | | ORF1ab | synonymous_L5230L |
| 16173 | C | T | 1.26E-04 | | | | | ORF1ab | synonymous_N5303N |
| 16188 | G | T | 1.26E-04 | | | | | ORF1ab | nonsynonymous_W5308C |
| 16203 | T | C | 1.26E-04 | | | | | ORF1ab | synonymous_Y5313Y |
| 16221 | G | A | 1.26E-04 | | | | | ORF1ab | synonymous_P5319P |

| | | | | | | | | | |
|---|---|---|---|---|---|---|---|---|---|
| 16222 | C | T | 1.26E-04 | | | | | ORF1ab | nonsynonymous_H5320Y |
| 16323 | C | T | 1.26E-04 | | | | | ORF1ab | synonymous_C5353C |
| 16338 | C | T | 1.26E-04 | | | | | ORF1ab | synonymous_V5358V |
| 16355 | A | G | 1.26E-04 | | | | | ORF1ab | nonsynonymous_K5364R |
| 16365 | G | T | 1.26E-04 | | | | | ORF1ab | nonsynonymous_L5367F |
| 16387 | A | T | 1.26E-04 | | | | | ORF1ab | nonsynonymous_N5375Y |
| 16863 | T | C | 1.26E-04 | | | | | ORF1ab | synonymous_V5533V |
| 16906 | T | C | 1.26E-04 | | | | | ORF1ab | nonsynonymous_Y5548H |
| 16942 | A | G | 1.26E-04 | | | | | ORF1ab | nonsynonymous_S5560G |
| 16968 | G | A | 1.26E-04 | T | 6.32E-05 | | | ORF1ab | synonymous_E5568E; nonsynonymous_E5568D |
| 16984 | A | G | 1.26E-04 | | | | | ORF1ab | nonsynonymous_T5574A |
| 16996 | C | T | 1.26E-04 | | | | | ORF1ab | nonsynonymous_P5578S |
| 17002 | C | T | 1.26E-04 | | | | | ORF1ab | nonsynonymous_L5580F |
| 17012 | C | T | 1.26E-04 | | | | | ORF1ab | nonsynonymous_S5583L |
| 17069 | C | A | 1.26E-04 | | | | | ORF1ab | nonsynonymous_S5602Y |
| 17110 | G | T | 1.26E-04 | | | | | ORF1ab | nonsynonymous_A5616S |
| 17149 | G | A | 1.26E-04 | | | | | ORF1ab | nonsynonymous_V5629M |
| 17158 | G | A | 1.26E-04 | | | | | ORF1ab | nonsynonymous_A5632T |
| 17270 | A | G | 1.26E-04 | | | | | ORF1ab | nonsynonymous_K5669R |
| 17302 | G | T | 1.26E-04 | | | | | ORF1ab | nonsynonymous_V5680F |
| 17339 | C | T | 1.26E-04 | | | | | ORF1ab | nonsynonymous_A5692V |
| 17372 | C | T | 1.26E-04 | | | | | ORF1ab | nonsynonymous_A5703V |
| 17397 | C | T | 1.26E-04 | | | | | ORF1ab | synonymous_V5711V |
| 17407 | T | C | 1.26E-04 | | | | | ORF1ab | synonymous_L5715L |
| 17443 | G | T | 1.26E-04 | | | | | ORF1ab | nonsynonymous_A5727S |
| 17632 | G | A | 1.26E-04 | T | 6.32E-05 | | | ORF1ab | nonsynonymous_D5790N; nonsynonymous_D5790Y |
| 17634 | C | A | 1.26E-04 | T | 6.32E-05 | | | ORF1ab | nonsynonymous_D5790E; synonymous_D5790D |

| | | | | | | | | | |
|---|---|---|---|---|---|---|---|---|---|
| 17659 | T | G | 1.26E-04 | | | | | ORF1ab | nonsynonymous_F5799V |
| 17688 | T | C | 1.26E-04 | | | | | ORF1ab | synonymous_V5808V |
| 17835 | T | C | 1.26E-04 | | | | | ORF1ab | synonymous_V5857V |
| 17850 | C | T | 1.26E-04 | | | | | ORF1ab | synonymous_G5862G |
| 17949 | C | T | 1.26E-04 | | | | | ORF1ab | synonymous_G5895G |
| 18020 | G | A | 1.26E-04 | T | 6.32E-05 | | | ORF1ab | nonsynonymous_R5919K; nonsynonymous_R5919M |
| 18248 | C | T | 1.26E-04 | | | | | ORF1ab | nonsynonymous_P5995L |
| 18267 | C | T | 1.26E-04 | | | | | ORF1ab | synonymous_R6001R |
| 18281 | G | T | 1.26E-04 | | | | | ORF1ab | nonsynonymous_R6006I |
| 18318 | G | A | 1.26E-04 | T | 6.32E-05 | | | ORF1ab | synonymous_G6018G; synonymous_G6018G |
| 18325 | G | T | 1.26E-04 | | | | | ORF1ab | nonsynonymous_A6021S |
| 18379 | G | A | 1.26E-04 | | | | | ORF1ab | nonsynonymous_G6039S |
| 18394 | G | T | 1.26E-04 | | | | | ORF1ab | nonsynonymous_A6044S |
| 18411 | T | C | 1.26E-04 | | | | | ORF1ab | synonymous_Y6049Y |
| 18412 | G | T | 1.26E-04 | A | 6.32E-05 | | | ORF1ab | nonsynonymous_V6050F; nonsynonymous_V6050I |
| 18433 | G | A | 1.26E-04 | | | | | ORF1ab | nonsynonymous_D6057N |
| 18495 | T | C | 1.26E-04 | A | 6.32E-05 | | | ORF1ab | synonymous_L6077L; synonymous_L6077L |
| 18511 | C | T | 1.26E-04 | G | 6.32E-05 | | | ORF1ab | nonsynonymous_P6083S; nonsynonymous_P6083A |
| 18555 | C | T | 1.26E-04 | | | | | ORF1ab | synonymous_D6097D |
| 18557 | C | T | 1.26E-04 | | | | | ORF1ab | nonsynonymous_T6098I |
| 18754 | C | G | 1.26E-04 | | | | | ORF1ab | nonsynonymous_P6164A |
| 18755 | C | T | 1.26E-04 | | | | | ORF1ab | nonsynonymous_P6164L |
| 18765 | T | C | 1.26E-04 | | | | | ORF1ab | synonymous_I6167I |
| 18858 | T | C | 1.26E-04 | | | | | ORF1ab | synonymous_D6198D |
| 18888 | C | T | 1.26E-04 | | | | | ORF1ab | synonymous_H6208H |

| | | | | | | | | | |
|---|---|---|---|---|---|---|---|---|---|
| 1895 | G | T | 1.26E-04 | | | | | ORF1ab | nonsynonymous_V544L |
| 18979 | C | T | 1.26E-04 | | | | | ORF1ab | nonsynonymous_H6239Y |
| 18981 | C | T | 1.26E-04 | G | 6.32E-05 | | | ORF1ab | synonymous_H6239H;<br>nonsynonymous_H6239Q |
| 19006 | G | T | 1.26E-04 | | | | | ORF1ab | nonsynonymous_A6248S |
| 19098 | A | G | 1.26E-04 | | | | | ORF1ab | synonymous_A6278A |
| 19126 | A | G | 1.26E-04 | | | | | ORF1ab | nonsynonymous_I6288V |
| 19166 | A | G | 1.26E-04 | | | | | ORF1ab | nonsynonymous_K6301R |
| 19182 | A | G | 1.26E-04 | | | | | ORF1ab | synonymous_V6306V |
| 19186 | C | T | 1.26E-04 | | | | | ORF1ab | synonymous_L6308L |
| 19586 | C | A | 1.26E-04 | | | | | ORF1ab | nonsynonymous_T6441N |
| 19627 | G | A | 1.26E-04 | | | | | ORF1ab | nonsynonymous_E6455K |
| 19632 | T | C | 1.26E-04 | A | 6.32E-05 | | | ORF1ab | synonymous_N6456N;<br>nonsynonymous_N6456K |
| 19635 | G | A | 1.26E-04 | | | | | ORF1ab | synonymous_V6457V |
| 19637 | C | T | 1.26E-04 | | | | | ORF1ab | nonsynonymous_A6458V |
| 1968 | C | T | 1.26E-04 | | | | | ORF1ab | nonsynonymous_T568I |
| 19721 | A | C | 1.26E-04 | | | | | ORF1ab | nonsynonymous_K6486T |
| 1973 | C | T | 1.26E-04 | | | | | ORF1ab | synonymous_L570L |
| 19791 | T | C | 1.26E-04 | | | | | ORF1ab | synonymous_L6509L |
| 19800 | G | T | 1.26E-04 | A | 6.32E-05 | | | ORF1ab | nonsynonymous_K6512N;<br>synonymous_K6512K |
| 19813 | C | T | 1.26E-04 | | | | | ORF1ab | nonsynonymous_P6517S |
| 19881 | C | T | 1.26E-04 | | | | | ORF1ab | synonymous_D6539D |
| 19891 | G | T | 1.26E-04 | | | | | ORF1ab | nonsynonymous_D6543Y |
| 19895 | C | T | 1.26E-04 | | | | | ORF1ab | nonsynonymous_A6544V |
| 2023 | C | T | 1.26E-04 | | | | | ORF1ab | synonymous_F586F |
| 20635 | G | A | 1.26E-04 | | | | | ORF1ab | nonsynonymous_E6791K |
| 20645 | A | G | 1.26E-04 | | | | | ORF1ab | nonsynonymous_Y6794C |
| 20646 | C | T | 1.26E-04 | | | | | ORF1ab | synonymous_Y6794Y |

| | | | | | | | | | |
|---|---|---|---|---|---|---|---|---|---|
| 20698 | C | T | 1.26E-04 | | | | | ORF1ab | nonsynonymous_L6812F |
| 20725 | G | T | 1.26E-04 | | | | | ORF1ab | stopgain_E6821X |
| 20790 | C | T | 1.26E-04 | A | 6.32E-05 | | | ORF1ab | synonymous_V6842V;<br>synonymous_V6842V |
| 20980 | G | A | 1.26E-04 | T | 6.32E-05 | T | 6.32E-05 | ORF1ab | nonsynonymous_D6906N;<br>nonsynonymous_D6906Y;<br>nonsynonymous_D6906Y |
| 20995 | G | T | 1.26E-04 | | | | | ORF1ab | nonsynonymous_G6911C |
| 21014 | A | G | 1.26E-04 | | | | | ORF1ab | nonsynonymous_H6917R |
| 21108 | C | T | 1.26E-04 | | | | | ORF1ab | synonymous_F6948F |
| 21114 | C | T | 1.26E-04 | | | | | ORF1ab | synonymous_Y6950Y |
| 21145 | C | T | 1.26E-04 | | | | | ORF1ab | nonsynonymous_L6961F |
| 21736 | C | T | 1.26E-04 | | | | | S | synonymous_F58F |
| 21745 | T | C | 1.26E-04 | | | | | S | synonymous_N61N |
| 21801 | A | C | 1.26E-04 | | | | | S | nonsynonymous_D80A |
| 21812 | C | T | 1.26E-04 | | | | | S | synonymous_L84L |
| 21825 | A | C | 1.26E-04 | | | | | S | nonsynonymous_D88A |
| 21843 | C | T | 1.26E-04 | | | | | S | nonsynonymous_S94F |
| 21850 | G | T | 1.26E-04 | | | | | S | nonsynonymous_E96D |
| 21890 | T | C | 1.26E-04 | | | | | S | synonymous_L110L |
| 21893 | G | A | 1.26E-04 | | | | | S | nonsynonymous_D111N |
| 21917 | A | G | 1.26E-04 | | | | | S | nonsynonymous_I119V |
| 22003 | A | G | 1.26E-04 | | | | | S | synonymous_K147K |
| 22638 | G | A | 1.26E-04 | | | | | S | nonsynonymous_S359N |
| 22663 | C | A | 1.26E-04 | | | | | S | synonymous_V367V |
| 22735 | C | T | 1.26E-04 | | | | | S | synonymous_C391C |
| 22865 | G | T | 1.26E-04 | | | | | S | nonsynonymous_A435S |
| 22875 | C | T | 1.26E-04 | | | | | S | nonsynonymous_S438F |
| 23155 | T | C | 1.26E-04 | | | | | S | synonymous_T531T |
| 23206 | C | T | 1.26E-04 | | | | | S | synonymous_G548G |

| | | | | | | | | | |
|---|---|---|---|---|---|---|---|---|---|
| 23220 | C | T | 1.26E-04 | A | 6.32E-05 | | | S | nonsynonymous_T553I; nonsynonymous_T553N |
| 23284 | T | C | 1.26E-04 | | | | | S | synonymous_D574D |
| 23285 | G | T | 1.26E-04 | | | | | S | nonsynonymous_A575S |
| 23315 | C | T | 1.26E-04 | | | | | S | nonsynonymous_L585F |
| 23359 | A | G | 1.26E-04 | | | | | S | synonymous_T599T |
| 23383 | G | A | 1.26E-04 | | | | | S | synonymous_Q607Q |
| 23402 | G | A | 1.26E-04 | | | | | S | nonsynonymous_D614N |
| 23415 | C | T | 1.26E-04 | | | | | S | nonsynonymous_T618I |
| 23439 | C | T | 1.26E-04 | | | | | S | nonsynonymous_A626V |
| 23481 | C | T | 1.26E-04 | | | | | S | nonsynonymous_S640F |
| 23670 | A | G | 1.26E-04 | | | | | S | nonsynonymous_N703S |
| 23677 | T | C | 1.26E-04 | | | | | S | synonymous_V705V |
| 23701 | C | T | 1.26E-04 | | | | | S | synonymous_A713A |
| 23702 | A | T | 1.26E-04 | | | | | S | nonsynonymous_I714L |
| 23779 | A | G | 1.26E-04 | | | | | S | synonymous_T739T |
| 23873 | G | T | 1.26E-04 | | | | | S | nonsynonymous_A771S |
| 23874 | C | T | 1.26E-04 | | | | | S | nonsynonymous_A771V |
| 23876 | G | A | 1.26E-04 | | | | | S | nonsynonymous_V772I |
| 24063 | T | C | 1.26E-04 | | | | | S | nonsynonymous_I834T |
| 24434 | G | T | 1.26E-04 | | | | | S | nonsynonymous_A958S |
| 24436 | T | C | 1.26E-04 | | | | | S | synonymous_A958A |
| 24442 | C | T | 1.26E-04 | A | 6.32E-05 | | | S | synonymous_N960N; nonsynonymous_N960K |
| 24751 | G | T | 1.26E-04 | | | | | S | nonsynonymous_L1063F |
| 24755 | G | T | 1.26E-04 | | | | | S | nonsynonymous_V1065L |
| 24757 | G | T | 1.26E-04 | | | | | S | synonymous_V1065V |
| 25311 | G | T | 1.26E-04 | | | | | S | nonsynonymous_C1250F |
| 25314 | G | T | 1.26E-04 | | | | | S | nonsynonymous_G1251V |
| 25323 | G | T | 1.26E-04 | | | | | S | nonsynonymous_C1254F |

| | | | | | | | | | |
|---|---|---|---|---|---|---|---|---|---|
| 25395 | G | T | 1.26E-04 | | | | | ORF3a | nonsynonymous_M1I |
| 25405 | A | G | 1.26E-04 | | | | | ORF3a | nonsynonymous_M5V |
| 25437 | G | T | 1.26E-04 | | | | | ORF3a | nonsynonymous_L15F |
| 25471 | G | T | 1.26E-04 | T | 1.26E-04 | | | ORF3a | nonsynonymous_D27Y; nonsynonymous_D27Y |
| 25480 | C | T | 1.26E-04 | | | | | ORF3a | nonsynonymous_R30C |
| 25499 | C | T | 1.26E-04 | | | | | ORF3a | nonsynonymous_P36L |
| 25539 | C | T | 1.26E-04 | | | | | ORF3a | synonymous_G49G |
| 25936 | C | T | 1.26E-04 | | | | | ORF3a | nonsynonymous_H182Y |
| 25981 | G | A | 1.26E-04 | | | | | ORF3a | nonsynonymous_V197I |
| 26037 | C | T | 1.26E-04 | | | | | ORF3a | synonymous_Y215Y |
| 26042 | C | T | 1.26E-04 | | | | | ORF3a | nonsynonymous_T217I |
| 26078 | C | T | 1.26E-04 | | | | | ORF3a | nonsynonymous_T229I |
| 26143 | G | T | 1.26E-04 | | | | | ORF3a | nonsynonymous_G251C |
| 26151 | C | T | 1.26E-04 | | | | | ORF3a | synonymous_S253S |
| 26183 | A | G | 1.26E-04 | | | | | ORF3a | nonsynonymous_Y264C |
| 26198 | C | T | 1.26E-04 | | | | | ORF3a | nonsynonymous_T269M |
| 26209 | G | C | 1.26E-04 | T | 6.32E-05 | | | ORF3a | nonsynonymous_V273L; nonsynonymous_V273L |
| 26226 | A | G | 1.26E-04 | | | | | intergenic | |
| 26313 | C | T | 1.26E-04 | | | | | E | synonymous_F23F |
| 26320 | T | C | 1.26E-04 | | | | | E | nonsynonymous_F26L |
| 26566 | A | G | 1.26E-04 | | | | | M | nonsynonymous_K15R |
| 26589 | G | C | 1.26E-04 | T | 6.32E-05 | | | M | nonsynonymous_V23L; nonsynonymous_V23L |
| 26606 | C | T | 1.26E-04 | | | | | M | synonymous_F28F |
| 26622 | C | T | 1.26E-04 | | | | | M | nonsynonymous_L34F |
| 26642 | C | T | 1.26E-04 | A | 6.32E-05 | | | M | synonymous_A40A; synonymous_A40A |
| 26645 | C | T | 1.26E-04 | | | | | M | synonymous_N41N |

| | | | | | | | | | |
|---|---|---|---|---|---|---|---|---|---|
| 26758 | G | T | 1.26E-04 | | | | | M | nonsynonymous_G79V |
| 26768 | C | T | 1.26E-04 | | | | | M | synonymous_I82I |
| 26779 | G | T | 1.26E-04 | | | | | M | nonsynonymous_C86F |
| 26834 | G | T | 1.26E-04 | | | | | M | synonymous_A104A |
| 26840 | G | T | 1.26E-04 | | | | | M | synonymous_T106T |
| 26842 | G | A | 1.26E-04 | T | 6.32E-05 | | | M | nonsynonymous_R107H; nonsynonymous_R107L |
| 27214 | G | T | 1.26E-04 | | | | | ORF6 | nonsynonymous_V5F |
| 27217 | G | T | 1.26E-04 | | | | | ORF6 | nonsynonymous_D6Y |
| 27219 | C | T | 1.26E-04 | | | | | ORF6 | synonymous_D6D |
| 27222 | T | C | 1.26E-04 | | | | | ORF6 | synonymous_F7F |
| 27233 | T | C | 1.26E-04 | | | | | ORF6 | nonsynonymous_I11T |
| 27237 | A | G | 1.26E-04 | T | 6.32E-05 | | | ORF6 | synonymous_A12A; synonymous_A12A |
| 27338 | A | G | 1.26E-04 | | | | | ORF6 | nonsynonymous_E46G |
| 27407 | T | C | 1.26E-04 | | | | | ORF7a | nonsynonymous_L5P |
| 27420 | G | T | 1.26E-04 | | | | | ORF7a | synonymous_L9L |
| 28290 | C | T | 1.26E-04 | | | | | N | nonsynonymous_P6L |
| 28292 | C | A | 1.26E-04 | | | | | N | nonsynonymous_Q7K |
| 28297 | T | C | 1.26E-04 | | | | | N | synonymous_N8N |
| 28305 | A | C | 1.26E-04 | G | 6.32E-05 | | | N | nonsynonymous_N11T; nonsynonymous_N11S |
| 28320 | C | T | 1.26E-04 | | | | | N | nonsynonymous_T16M |
| 28340 | T | A | 1.26E-04 | C | 6.32E-05 | | | N | nonsynonymous_S23T; nonsynonymous_S23P |
| 28457 | G | A | 1.26E-04 | | | | | N | nonsynonymous_E62K |
| 28460 | G | A | 1.26E-04 | | | | | N | nonsynonymous_D63N |
| 28482 | A | G | 1.26E-04 | | | | | N | nonsynonymous_Q70R |
| 28487 | G | C | 1.26E-04 | A | 6.32E-05 | | | N | nonsynonymous_V72L; nonsynonymous_V72I |

| | | | | | | | | | |
|---|---|---|---|---|---|---|---|---|---|
| 28528 | C | T | 1.26E-04 | | | | | N | synonymous_G85G |
| 28546 | C | A | 1.26E-04 | | | | | N | synonymous_T91T |
| 28582 | T | C | 1.26E-04 | | | | | N | synonymous_D103D |
| 28609 | C | T | 1.26E-04 | | | | | N | synonymous_Y112Y |
| 28685 | G | T | 1.26E-04 | | | | | N | nonsynonymous_A138S |
| 28706 | C | T | 1.26E-04 | | | | | N | nonsynonymous_H145Y |
| 28709 | A | T | 1.26E-04 | | | | | N | nonsynonymous_I146F |
| 28728 | C | T | 1.26E-04 | | | | | N | nonsynonymous_A152V |
| 28748 | C | T | 1.26E-04 | | | | | N | synonymous_L159L |
| 29020 | A | G | 1.26E-04 | | | | | N | synonymous_K249K |
| 3338 | G | T | 1.26E-04 | | | | | ORF1ab | nonsynonymous_V1025L |
| 3531 | A | G | 1.26E-04 | | | | | ORF1ab | nonsynonymous_D1089G |
| 370 | G | T | 1.26E-04 | | | | | ORF1ab | synonymous_V35V |
| 3842 | A | G | 1.26E-04 | | | | | ORF1ab | nonsynonymous_M1193V |
| 3866 | C | G | 1.26E-04 | A | 6.32E-05 | | | ORF1ab | nonsynonymous_Q1201E; nonsynonymous_Q1201K |
| 3917 | A | G | 1.26E-04 | | | | | ORF1ab | nonsynonymous_S1218G |
| 4010 | C | T | 1.26E-04 | | | | | ORF1ab | nonsynonymous_L1249F |
| 4261 | G | A | 1.26E-04 | | | | | ORF1ab | synonymous_Q1332Q |
| 5264 | A | G | 1.26E-04 | | | | | ORF1ab | nonsynonymous_I1667V |
| 6109 | G | A | 1.26E-04 | | | | | ORF1ab | synonymous_K1948K |
| 6266 | G | A | 1.26E-04 | | | | | ORF1ab | nonsynonymous_A2001T |
| 6324 | A | G | 1.26E-04 | | | | | ORF1ab | nonsynonymous_E2020G |
| 6336 | C | T | 1.26E-04 | | | | | ORF1ab | nonsynonymous_S2024L |
| 6341 | G | A | 1.26E-04 | | | | | ORF1ab | nonsynonymous_D2026N |
| 6347 | C | T | 1.26E-04 | | | | | ORF1ab | synonymous_L2028L |
| 6358 | G | A | 1.26E-04 | | | | | ORF1ab | synonymous_E2031E |
| 6371 | A | G | 1.26E-04 | | | | | ORF1ab | nonsynonymous_M2036V |
| 6401 | C | T | 1.26E-04 | G | 6.32E-05 | | | ORF1ab | nonsynonymous_P2046S; nonsynonymous_P2046A |

| | | | | | | | | | |
|---|---|---|---|---|---|---|---|---|---|
| 6419 | G | A | 1.26E-04 | | | | | ORF1ab | nonsynonymous_V2052M |
| 6669 | G | T | 1.26E-04 | | | | | ORF1ab | nonsynonymous_W2135L |
| 6734 | G | A | 1.26E-04 | | | | | ORF1ab | nonsynonymous_V2157I |
| 6791 | T | A | 1.26E-04 | | | | | ORF1ab | nonsynonymous_L2176M |
| 6883 | C | T | 1.26E-04 | | | | | ORF1ab | synonymous_V2206V |
| 6989 | T | C | 1.26E-04 | | | | | ORF1ab | nonsynonymous_S2242P |
| 7083 | C | T | 1.26E-04 | | | | | ORF1ab | nonsynonymous_S2273F |
| 8131 | G | T | 1.26E-04 | | | | | ORF1ab | nonsynonymous_K2622N |
| 8667 | A | G | 1.26E-04 | | | | | ORF1ab | nonsynonymous_D2801G |
| 8692 | C | T | 1.26E-04 | | | | | ORF1ab | synonymous_Y2809Y |
| 8697 | C | T | 1.26E-04 | | | | | ORF1ab | nonsynonymous_A2811V |
| 8707 | T | C | 1.26E-04 | | | | | ORF1ab | synonymous_G2814G |
| 8722 | C | T | 1.26E-04 | | | | | ORF1ab | synonymous_D2819D |
| 8727 | C | T | 1.26E-04 | | | | | ORF1ab | nonsynonymous_A2821V |
| 8777 | T | C | 1.26E-04 | | | | | ORF1ab | nonsynonymous_F2838L |
| 9141 | G | A | 1.26E-04 | | | | | ORF1ab | nonsynonymous_G2959D |
| 9307 | A | G | 1.26E-04 | | | | | ORF1ab | synonymous_L3014L |
| 9443 | C | T | 1.26E-04 | | | | | ORF1ab | nonsynonymous_L3060F |
| 9642 | T | C | 1.26E-04 | | | | | ORF1ab | nonsynonymous_I3126T |
| 9856 | G | T | 1.26E-04 | | | | | ORF1ab | synonymous_V3197V |
| 10096 | G | A | 1.26E-04 | | | | | ORF1ab | synonymous_E3277E |
| 10150 | T | C | 1.26E-04 | | | | | ORF1ab | synonymous_L3295L |
| 10360 | G | A | 1.26E-04 | | | | | ORF1ab | synonymous_K3365K |
| 10368 | G | A | 1.26E-04 | | | | | ORF1ab | nonsynonymous_R3368H |
| 10537 | C | T | 1.26E-04 | | | | | ORF1ab | synonymous_Y3424Y |
| 11152 | C | T | 1.26E-04 | | | | | ORF1ab | synonymous_V3629V |
| 11289 | C | T | 1.26E-04 | | | | | ORF1ab | nonsynonymous_S3675F |
| 11635 | T | C | 1.26E-04 | | | | | ORF1ab | synonymous_C3790C |
| 11842 | G | T | 1.26E-04 | A | 6.32E-05 | | | ORF1ab | nonsynonymous_Q3859H; synonymous_Q3859Q |

| | | | | | | | | | |
|---|---|---|---|---|---|---|---|---|---|
| 11866 | C | T | 1.26E-04 | | | | | ORF1ab | synonymous_C3867C |
| 12015 | T | G | 1.26E-04 | | | | | ORF1ab | nonsynonymous_V3917G |
| 12037 | T | C | 1.26E-04 | | | | | ORF1ab | synonymous_A3924A |
| 12067 | G | T | 1.26E-04 | | | | | ORF1ab | nonsynonymous_M3934I |
| 12102 | C | T | 1.26E-04 | | | | | ORF1ab | nonsynonymous_S3946L |
| 12123 | C | T | 1.26E-04 | | | | | ORF1ab | nonsynonymous_S3953L |
| 12131 | G | A | 1.26E-04 | | | | | ORF1ab | nonsynonymous_A3956T |
| 1230 | A | G | 1.26E-04 | | | | | ORF1ab | nonsynonymous_K322R |
| 12374 | C | T | 1.26E-04 | | | | | ORF1ab | nonsynonymous_L4037F |
| 12828 | T | C | 1.26E-04 | | | | | ORF1ab | nonsynonymous_L4188S |
| 12838 | G | T | 1.26E-04 | | | | | ORF1ab | nonsynonymous_L4191F |
| 1348 | C | T | 1.26E-04 | | | | | ORF1ab | synonymous_P361P |
| 1439 | G | T | 1.26E-04 | | | | | ORF1ab | nonsynonymous_G392C |
| 1471 | C | T | 1.26E-04 | | | | | ORF1ab | synonymous_R402R |
| 1477 | T | A | 1.26E-04 | | | | | ORF1ab | synonymous_I404I |
| 1519 | C | T | 1.26E-04 | | | | | ORF1ab | synonymous_N418N |
| 1524 | G | A | 1.26E-04 | | | | | ORF1ab | nonsynonymous_C420Y |
| 1545 | C | T | 1.26E-04 | | | | | ORF1ab | nonsynonymous_A427V |
| 1620 | A | G | 1.26E-04 | | | | | ORF1ab | nonsynonymous_E452G |
| 1627 | C | T | 1.26E-04 | | | | | ORF1ab | synonymous_L454L |
| 1670 | C | T | 1.26E-04 | | | | | ORF1ab | nonsynonymous_L469F |
| 1710 | C | T | 1.26E-04 | | | | | ORF1ab | nonsynonymous_A482V |
| 1729 | G | A | 1.26E-04 | T | 6.32E-05 | | | ORF1ab | synonymous_V488V; synonymous_V488V |
| 1743 | G | A | 1.26E-04 | | | | | ORF1ab | nonsynonymous_G493D |
| 1747 | G | A | 1.26E-04 | | | | | ORF1ab | synonymous_L494L |
| 1758 | C | T | 1.26E-04 | | | | | ORF1ab | nonsynonymous_A498V |
| 1779 | C | T | 1.26E-04 | | | | | ORF1ab | nonsynonymous_S505F |
| 1815 | A | G | 1.26E-04 | | | | | ORF1ab | nonsynonymous_K517R |
| 1919 | C | T | 1.26E-04 | | | | | ORF1ab | nonsynonymous_L552F |

| | | | | | | | | | |
|---|---|---|---|---|---|---|---|---|---|
| 1944 | G | A | 1.26E-04 | | | | | ORF1ab | nonsynonymous_R560H |
| 1978 | T | A | 1.26E-04 | | | | | ORF1ab | nonsynonymous_D571E |
| 1980 | G | A | 1.26E-04 | | | | | ORF1ab | nonsynonymous_G572E |
| 1999 | G | T | 1.26E-04 | | | | | ORF1ab | synonymous_L578L |
| 2086 | G | T | 1.26E-04 | | | | | ORF1ab | nonsynonymous_Q607H |
| 2095 | G | A | 1.26E-04 | | | | | ORF1ab | synonymous_S610S |
| 2100 | G | T | 1.26E-04 | | | | | ORF1ab | nonsynonymous_W612L |
| 2109 | A | G | 1.26E-04 | | | | | ORF1ab | nonsynonymous_N615S |
| 2156 | C | T | 1.26E-04 | | | | | ORF1ab | nonsynonymous_L631F |
| 2380 | A | G | 1.26E-04 | | | | | ORF1ab | synonymous_L705L |
| 2393 | G | A | 1.26E-04 | T | 6.32E-05 | | | ORF1ab | nonsynonymous_V710I; nonsynonymous_V710F |
| 2422 | G | A | 1.26E-04 | | | | | ORF1ab | synonymous_K719K |
| 2527 | G | A | 1.26E-04 | A | 1.26E-04 | | | ORF1ab | synonymous_E754E; synonymous_E754E |
| 2897 | G | A | 1.26E-04 | | | | | ORF1ab | nonsynonymous_A878T |
| 2915 | C | A | 1.26E-04 | | | | | ORF1ab | nonsynonymous_Q884K |
| 2969 | A | G | 1.26E-04 | | | | | ORF1ab | nonsynonymous_M902V |
| 2997 | C | T | 1.26E-04 | | | | | ORF1ab | nonsynonymous_S911F |
| 3076 | A | T | 1.26E-04 | | | | | ORF1ab | nonsynonymous_E937D |
| 3094 | A | G | 1.26E-04 | | | | | ORF1ab | synonymous_P943P |
| 3122 | G | T | 1.26E-04 | | | | | ORF1ab | nonsynonymous_D953Y |
| 3157 | C | T | 1.26E-04 | | | | | ORF1ab | synonymous_A964A |
| 3365 | C | T | 1.26E-04 | | | | | ORF1ab | nonsynonymous_L1034F |
| 3433 | G | T | 1.26E-04 | | | | | ORF1ab | synonymous_V1056V |
| 3447 | C | T | 1.26E-04 | | | | | ORF1ab | nonsynonymous_A1061V |
| 3466 | T | C | 1.26E-04 | | | | | ORF1ab | synonymous_H1067H |
| 3523 | A | G | 1.26E-04 | | | | | ORF1ab | synonymous_E1086E |
| 3728 | G | T | 1.26E-04 | | | | | ORF1ab | nonsynonymous_G1155C |
| 3966 | C | T | 1.26E-04 | | | | | ORF1ab | nonsynonymous_A1234V |

| | | | | | | | | | |
|---|---|---|---|---|---|---|---|---|---|
| 3969 | G | A | 1.26E-04 | | | | | ORF1ab | nonsynonymous_C1235Y |
| 3987 | C | T | 1.26E-04 | | | | | ORF1ab | nonsynonymous_T1241I |
| 4059 | C | A | 1.26E-04 | | | | | ORF1ab | nonsynonymous_P1265Q |
| 4066 | T | C | 1.26E-04 | | | | | ORF1ab | synonymous_S1267S |
| 4105 | G | T | 1.26E-04 | | | | | ORF1ab | nonsynonymous_K1280N |
| 4166 | A | G | 1.26E-04 | | | | | ORF1ab | nonsynonymous_I1301V |
| 4656 | G | A | 1.26E-04 | C | 6.32E-05 | | | ORF1ab | nonsynonymous_R1464Q; nonsynonymous_R1464P |
| 4698 | C | T | 1.26E-04 | | | | | ORF1ab | nonsynonymous_S1478F |
| 4728 | G | T | 1.26E-04 | | | | | ORF1ab | nonsynonymous_G1488V |
| 4832 | C | T | 1.26E-04 | | | | | ORF1ab | synonymous_L1523L |
| 4886 | C | T | 1.26E-04 | | | | | ORF1ab | nonsynonymous_P1541S |
| 4893 | C | T | 1.26E-04 | | | | | ORF1ab | nonsynonymous_T1543I |
| 4993 | C | T | 1.26E-04 | | | | | ORF1ab | synonymous_N1576N |
| 5608 | A | G | 1.26E-04 | | | | | ORF1ab | synonymous_K1781K |
| 575 | C | T | 1.26E-04 | | | | | ORF1ab | nonsynonymous_L104F |
| 578 | G | A | 1.26E-04 | | | | | ORF1ab | nonsynonymous_G105S |
| 5849 | C | T | 1.26E-04 | | | | | ORF1ab | nonsynonymous_P1862S |
| 586 | T | C | 1.26E-04 | | | | | ORF1ab | synonymous_L107L |
| 606 | T | C | 1.26E-04 | | | | | ORF1ab | nonsynonymous_I114T |
| 6195 | C | T | 1.26E-04 | | | | | ORF1ab | nonsynonymous_P1977L |
| 643 | C | T | 1.26E-04 | | | | | ORF1ab | synonymous_N126N |
| 6538 | C | T | 1.26E-04 | | | | | ORF1ab | synonymous_G2091G |
| 6561 | A | G | 1.26E-04 | | | | | ORF1ab | nonsynonymous_Y2099C |
| 6769 | T | C | 1.26E-04 | | | | | ORF1ab | synonymous_Y2168Y |
| 680 | G | A | 1.26E-04 | | | | | ORF1ab | nonsynonymous_D139N |
| 6865 | G | T | 1.26E-04 | | | | | ORF1ab | nonsynonymous_K2200N |
| 7143 | G | A | 1.26E-04 | | | | | ORF1ab | nonsynonymous_S2293N |
| 7392 | C | T | 1.26E-04 | | | | | ORF1ab | nonsynonymous_P2376L |
| 7770 | A | G | 1.26E-04 | | | | | ORF1ab | nonsynonymous_H2502R |

| | | | | | | | | | |
|---|---|---|---|---|---|---|---|---|---|
| 7831 | T | C | 1.26E-04 | | | | | ORF1ab | synonymous_V2522V |
| 7844 | C | A | 1.26E-04 | | | | | ORF1ab | nonsynonymous_L2527M |
| 7926 | C | T | 1.26E-04 | | | | | ORF1ab | nonsynonymous_A2554V |
| 7990 | A | C | 1.26E-04 | | | | | ORF1ab | synonymous_A2575A |
| 8004 | T | C | 1.26E-04 | | | | | ORF1ab | nonsynonymous_V2580A |
| 8040 | A | G | 1.26E-04 | | | | | ORF1ab | nonsynonymous_D2592G |
| 8140 | C | T | 1.26E-04 | | | | | ORF1ab | synonymous_S2625S |
| 8203 | A | G | 1.26E-04 | | | | | ORF1ab | synonymous_V2646V |
| 8356 | T | G | 1.26E-04 | | | | | ORF1ab | synonymous_R2697R |
| 8357 | C | T | 1.26E-04 | | | | | ORF1ab | nonsynonymous_H2698Y |
| 8431 | T | C | 1.26E-04 | | | | | ORF1ab | synonymous_S2722S |
| 934 | C | T | 1.26E-04 | | | | | ORF1ab | synonymous_D223D |
| 9532 | C | T | 1.26E-04 | | | | | ORF1ab | synonymous_F3089F |
| 960 | G | A | 1.26E-04 | | | | | ORF1ab | nonsynonymous_R232H |
| 9650 | A | G | 1.26E-04 | | | | | ORF1ab | nonsynonymous_M3129V |
| 9685 | A | G | 1.26E-04 | | | | | ORF1ab | nonsynonymous_I3140M |
| 9951 | C | T | 1.26E-04 | | | | | ORF1ab | nonsynonymous_A3229V |
| 10036 | C | T | 1.26E-04 | | | | | ORF1ab | synonymous_I3257I |
| 10157 | G | T | 1.26E-04 | | | | | ORF1ab | nonsynonymous_V3298L |
| 10159 | A | G | 1.26E-04 | | | | | ORF1ab | synonymous_V3298V |
| 1015 | A | G | 1.26E-04 | | | | | ORF1ab | synonymous_E250E |
| 10198 | C | A | 1.26E-04 | | | | | ORF1ab | nonsynonymous_D3311E |
| 10208 | C | T | 1.26E-04 | | | | | ORF1ab | nonsynonymous_P3315S |
| 10279 | C | T | 1.26E-04 | | | | | ORF1ab | synonymous_L3338L |
| 10291 | A | G | 1.26E-04 | | | | | ORF1ab | synonymous_G3342G |
| 1030 | T | C | 1.26E-04 | | | | | ORF1ab | synonymous_F255F |
| 10341 | C | T | 1.26E-04 | | | | | ORF1ab | nonsynonymous_P3359L |
| 10384 | G | T | 1.26E-04 | | | | | ORF1ab | nonsynonymous_Q3373H |
| 10508 | A | G | 1.26E-04 | | | | | ORF1ab | nonsynonymous_I3415V |
| 10519 | C | T | 1.26E-04 | | | | | ORF1ab | synonymous_D3418D |

| | | | | | | | | | |
|---|---|---|---|---|---|---|---|---|---|
| 10521 | G | A | 1.26E-04 | T | 6.32E-05 | | | ORF1ab | nonsynonymous_C3419Y; nonsynonymous_C3419F |
| 10573 | T | C | 1.26E-04 | | | | | ORF1ab | synonymous_A3436A |
| 10576 | C | T | 1.26E-04 | | | | | ORF1ab | synonymous_G3437G |
| 10605 | C | T | 1.26E-04 | | | | | ORF1ab | nonsynonymous_P3447L |
| 10632 | C | T | 1.26E-04 | | | | | ORF1ab | nonsynonymous_A3456V |
| 10641 | C | T | 1.26E-04 | | | | | ORF1ab | nonsynonymous_T3459M |
| 10728 | C | T | 1.26E-04 | | | | | ORF1ab | nonsynonymous_T3488I |
| 1098 | A | G | 1.26E-04 | | | | | ORF1ab | nonsynonymous_N278S |
| 11121 | T | C | 1.26E-04 | | | | | ORF1ab | nonsynonymous_I3619T |
| 11161 | G | A | 1.26E-04 | | | | | ORF1ab | synonymous_K3632K |
| 11173 | C | T | 1.26E-04 | | | | | ORF1ab | synonymous_L3636L |
| 11221 | G | T | 1.26E-04 | | | | | ORF1ab | nonsynonymous_M3652I |
| 11233 | T | A | 1.26E-04 | G | 6.32E-05 | | | ORF1ab | synonymous_P3656P; synonymous_P3656P |
| 1123 | A | G | 1.26E-04 | | | | | ORF1ab | synonymous_P286P |
| 1125 | G | A | 1.26E-04 | | | | | ORF1ab | nonsynonymous_R287K |
| 11335 | G | T | 1.26E-04 | | | | | ORF1ab | synonymous_V3690V |
| 11365 | G | T | 1.26E-04 | | | | | ORF1ab | synonymous_V3700V |
| 11376 | G | T | 1.26E-04 | | | | | ORF1ab | nonsynonymous_G3704V |
| 1145 | G | A | 1.26E-04 | | | | | ORF1ab | nonsynonymous_D294N |
| 11464 | G | A | 1.26E-04 | T | 6.32E-05 | | | ORF1ab | nonsynonymous_M3733I; nonsynonymous_M3733I |
| 11471 | C | T | 1.26E-04 | | | | | ORF1ab | nonsynonymous_L3736F |
| 11519 | A | G | 1.26E-04 | | | | | ORF1ab | nonsynonymous_M3752V |
| 11534 | G | A | 1.26E-04 | | | | | ORF1ab | nonsynonymous_G3757S |
| 11557 | G | A | 1.26E-04 | A | 1.26E-04 | | | ORF1ab | synonymous_E3764E; synonymous_E3764E |
| 11563 | C | T | 1.26E-04 | | | | | ORF1ab | synonymous_C3766C |
| 11620 | C | T | 1.26E-04 | | | | | ORF1ab | synonymous_F3785F |

| | | | | | | | | | |
|---|---|---|---|---|---|---|---|---|---|
| 11634 | G | T | 1.26E-04 | | | | | ORF1ab | nonsynonymous_C3790F |
| 11651 | C | T | 1.26E-04 | | | | | ORF1ab | nonsynonymous_L3796F |
| 11663 | C | T | 1.26E-04 | | | | | ORF1ab | nonsynonymous_L3800F |
| 11781 | A | G | 1.26E-04 | | | | | ORF1ab | nonsynonymous_K3839R |
| 11858 | G | A | 1.26E-04 | | | | | ORF1ab | nonsynonymous_V3865I |
| 11878 | C | T | 1.26E-04 | A | 6.32E-05 | | | ORF1ab | synonymous_V3871V; synonymous_V3871V |
| 11937 | G | A | 1.26E-04 | | | | | ORF1ab | nonsynonymous_C3891Y |
| 11959 | T | C | 1.26E-04 | | | | | ORF1ab | synonymous_I3898I |
| 11974 | T | C | 1.26E-04 | | | | | ORF1ab | synonymous_D3903D |
| 11979 | C | T | 1.26E-04 | | | | | ORF1ab | nonsynonymous_T3905I |
| 12020 | C | T | 1.26E-04 | | | | | ORF1ab | nonsynonymous_L3919F |
| 12025 | C | T | 1.26E-04 | | | | | ORF1ab | synonymous_S3920S |
| 12027 | T | C | 1.26E-04 | | | | | ORF1ab | nonsynonymous_M3921T |
| 12052 | G | T | 1.26E-04 | A | 6.32E-05 | | | ORF1ab | nonsynonymous_K3929N; synonymous_K3929K |
| 12053 | C | T | 1.26E-04 | | | | | ORF1ab | nonsynonymous_L3930F |
| 12071 | G | A | 1.26E-04 | | | | | ORF1ab | nonsynonymous_D3936N |
| 12085 | C | T | 1.26E-04 | | | | | ORF1ab | synonymous_T3940T |
| 12091 | A | G | 1.26E-04 | | | | | ORF1ab | synonymous_Q3942Q |
| 12100 | C | T | 1.26E-04 | | | | | ORF1ab | synonymous_A3945A |
| 12111 | G | A | 1.26E-04 | | | | | ORF1ab | nonsynonymous_S3949N |
| 12121 | A | G | 1.26E-04 | | | | | ORF1ab | synonymous_P3952P |
| 12274 | G | A | 1.26E-04 | T | 6.32E-05 | | | ORF1ab | synonymous_K4003K; nonsynonymous_K4003N |
| 12283 | T | C | 1.26E-04 | | | | | ORF1ab | synonymous_D4006D |
| 12322 | G | A | 1.26E-04 | | | | | ORF1ab | synonymous_E4019E |
| 12393 | A | G | 1.26E-04 | | | | | ORF1ab | nonsynonymous_D4043G |
| 12395 | G | A | 1.26E-04 | | | | | ORF1ab | nonsynonymous_A4044T |
| 12415 | C | T | 1.26E-04 | | | | | ORF1ab | synonymous_N4050N |

| | | | | | | | | | |
|---|---|---|---|---|---|---|---|---|---|
| 12484 | C | T | 1.26E-04 | | | | | ORF1ab | synonymous_V4073V |
| 12528 | C | T | 1.26E-04 | | | | | ORF1ab | nonsynonymous_T4088I |
| 12529 | A | G | 1.26E-04 | | | | | ORF1ab | synonymous_T4088T |
| 12546 | C | T | 1.26E-04 | | | | | ORF1ab | nonsynonymous_A4094V |
| 12556 | A | G | 1.26E-04 | | | | | ORF1ab | synonymous_E4097E |
| 12565 | G | T | 1.26E-04 | C | 6.32E-05 | | | ORF1ab | nonsynonymous_Q4100H; nonsynonymous_Q4100H |
| 1260 | A | T | 1.26E-04 | | | | | ORF1ab | nonsynonymous_Q332L |
| 12664 | C | T | 1.26E-04 | | | | | ORF1ab | synonymous_A4133A |
| 12672 | C | T | 1.26E-04 | | | | | ORF1ab | nonsynonymous_A4136V |
| 12739 | T | C | 1.26E-04 | | | | | ORF1ab | synonymous_T4158T |
| 12757 | T | C | 1.26E-04 | | | | | ORF1ab | synonymous_T4164T |
| 12759 | A | C | 1.26E-04 | G | 6.32E-05 | | | ORF1ab | nonsynonymous_D4165A; nonsynonymous_D4165G |
| 12769 | G | T | 1.26E-04 | | | | | ORF1ab | synonymous_A4168A |
| 12793 | G | T | 1.26E-04 | | | | | ORF1ab | nonsynonymous_K4176N |
| 13051 | C | T | 1.26E-04 | | | | | ORF1ab | synonymous_A4262A |
| 13059 | C | T | 1.26E-04 | | | | | ORF1ab | nonsynonymous_T4265I |
| 1306 | G | T | 1.26E-04 | | | | | ORF1ab | nonsynonymous_E347D |
| 1308 | A | G | 1.26E-04 | | | | | ORF1ab | nonsynonymous_N348S |
| 13118 | G | T | 1.26E-04 | | | | | ORF1ab | nonsynonymous_A4285S |
| 1331 | A | G | 1.26E-04 | | | | | ORF1ab | nonsynonymous_T356A |
| 1350 | A | T | 1.26E-04 | | | | | ORF1ab | nonsynonymous_Q362L |
| 1404 | C | T | 1.26E-04 | | | | | ORF1ab | nonsynonymous_P380L |
| 1455 | T | C | 1.26E-04 | | | | | ORF1ab | nonsynonymous_L397P |
| 1580 | G | A | 1.26E-04 | | | | | ORF1ab | nonsynonymous_V439I |
| 1598 | G | A | 1.26E-04 | C | 6.32E-05 | | | ORF1ab | nonsynonymous_G445S; nonsynonymous_G445R |
| 1599 | G | T | 1.26E-04 | | | | | ORF1ab | nonsynonymous_G445V |
| 1630 | A | C | 1.26E-04 | | | | | ORF1ab | nonsynonymous_Q455H |

| | | | | | | | | | |
|---|---|---|---|---|---|---|---|---|---|
| 1813 | T | A | 1.26E-04 | | | | | ORF1ab | synonymous_A516A |
| 1817 | A | G | 1.26E-04 | | | | | ORF1ab | nonsynonymous_K518E |
| 1846 | A | T | 1.26E-04 | C | 6.32E-05 | | | ORF1ab | nonsynonymous_K527N; nonsynonymous_K527N |
| 1853 | C | T | 1.26E-04 | | | | | ORF1ab | synonymous_L530L |
| 1869 | C | T | 1.26E-04 | | | | | ORF1ab | nonsynonymous_A535V |
| 2232 | C | T | 1.26E-04 | | | | | ORF1ab | nonsynonymous_A656V |
| 2238 | A | G | 1.26E-04 | | | | | ORF1ab | nonsynonymous_E658G |
| 2249 | G | A | 1.26E-04 | | | | | ORF1ab | nonsynonymous_G662R |
| 2268 | C | T | 1.26E-04 | | | | | ORF1ab | nonsynonymous_A668V |
| 2334 | C | T | 1.26E-04 | | | | | ORF1ab | nonsynonymous_A690V |
| 2445 | C | T | 1.26E-04 | | | | | ORF1ab | nonsynonymous_T727I |
| 2446 | T | C | 1.26E-04 | | | | | ORF1ab | synonymous_T727T |
| 2453 | C | T | 1.26E-04 | A | 6.32E-05 | | | ORF1ab | nonsynonymous_L730F; nonsynonymous_L730I |
| 2459 | C | T | 1.26E-04 | | | | | ORF1ab | nonsynonymous_P732S |
| 2476 | A | G | 1.26E-04 | | | | | ORF1ab | synonymous_K737K |
| 2499 | A | C | 1.26E-04 | G | 6.32E-05 | | | ORF1ab | nonsynonymous_E745A; nonsynonymous_E745G |
| 2501 | A | G | 1.26E-04 | | | | | ORF1ab | nonsynonymous_T746A |
| 2509 | C | T | 1.26E-04 | | | | | ORF1ab | synonymous_P748P |
| 2550 | A | G | 1.26E-04 | | | | | ORF1ab | nonsynonymous_D762G |
| 2551 | T | C | 1.26E-04 | | | | | ORF1ab | synonymous_D762D |
| 2565 | A | G | 1.26E-04 | | | | | ORF1ab | nonsynonymous_E767G |
| 2862 | C | T | 1.26E-04 | A | 6.32E-05 | | | ORF1ab | nonsynonymous_T866I; nonsynonymous_T866K |
| 2875 | G | A | 1.26E-04 | | | | | ORF1ab | synonymous_E870E |
| 2880 | C | T | 1.26E-04 | | | | | ORF1ab | nonsynonymous_A872V |
| 3176 | C | T | 1.26E-04 | | | | | ORF1ab | nonsynonymous_P971S |

| | | | | | | | | | |
|---|---|---|---|---|---|---|---|---|---|
| 3242 | G | A | 1.26E-04 | A | 1.26E-04 | | | ORF1ab | nonsynonymous_G993S; nonsynonymous_G993S |
| 3293 | C | T | 1.26E-04 | | | | | ORF1ab | nonsynonymous_P1010S |
| 3492 | A | C | 1.26E-04 | | | | | ORF1ab | nonsynonymous_N1076T |
| 3517 | A | G | 1.26E-04 | | | | | ORF1ab | synonymous_Q1084Q |
| 3558 | A | G | 1.26E-04 | | | | | ORF1ab | nonsynonymous_K1098R |
| 3559 | A | G | 1.26E-04 | | | | | ORF1ab | synonymous_K1098K |
| 3582 | G | A | 1.26E-04 | T | 6.32E-05 | | | ORF1ab | nonsynonymous_S1106N; nonsynonymous_S1106I |
| 3602 | C | T | 1.26E-04 | | | | | ORF1ab | nonsynonymous_H1113Y |
| 3625 | A | C | 1.26E-04 | | | | | ORF1ab | synonymous_P1120P |
| 3635 | A | G | 1.26E-04 | | | | | ORF1ab | nonsynonymous_K1124E |
| 3688 | C | T | 1.26E-04 | | | | | ORF1ab | synonymous_H1141H |
| 3736 | C | T | 1.26E-04 | | | | | ORF1ab | synonymous_D1157D |
| 3766 | T | C | 1.26E-04 | | | | | ORF1ab | synonymous_D1167D |
| 3811 | C | T | 1.26E-04 | | | | | ORF1ab | synonymous_L1182L |
| 4095 | C | T | 1.26E-04 | | | | | ORF1ab | nonsynonymous_T1277I |
| 4127 | G | A | 1.26E-04 | | | | | ORF1ab | nonsynonymous_G1288S |
| 4162 | G | T | 1.26E-04 | | | | | ORF1ab | synonymous_V1299V |
| 4235 | A | G | 1.26E-04 | T | 6.32E-05 | | | ORF1ab | nonsynonymous_N1324D; nonsynonymous_N1324Y |
| 4269 | A | G | 1.26E-04 | | | | | ORF1ab | nonsynonymous_N1335S |
| 4300 | G | T | 1.26E-04 | | | | | ORF1ab | synonymous_V1345V |
| 4386 | A | G | 1.26E-04 | | | | | ORF1ab | nonsynonymous_N1374S |
| 443 | G | A | 1.26E-04 | | | | | ORF1ab | nonsynonymous_V60I |
| 4494 | A | G | 1.26E-04 | | | | | ORF1ab | nonsynonymous_K1410R |
| 4579 | T | A | 1.26E-04 | | | | | ORF1ab | synonymous_L1438L |
| 4591 | T | C | 1.26E-04 | | | | | ORF1ab | synonymous_N1442N |
| 4609 | G | T | 1.26E-04 | | | | | ORF1ab | nonsynonymous_M1448I |
| 4631 | G | A | 1.26E-04 | | | | | ORF1ab | nonsynonymous_G1456S |

| | | | | | | | | | |
|---|---|---|---|---|---|---|---|---|---|
| 4763 | C | T | 1.26E-04 | | | | | ORF1ab | nonsynonymous_H1500Y |
| 4878 | C | T | 1.26E-04 | | | | | ORF1ab | nonsynonymous_T1538I |
| 4884 | A | G | 1.26E-04 | | | | | ORF1ab | nonsynonymous_N1540S |
| 4925 | G | T | 1.26E-04 | | | | | ORF1ab | nonsynonymous_D1554Y |
| 4940 | C | A | 1.26E-04 | A | 1.26E-04 | | | ORF1ab | nonsynonymous_L1559I; nonsynonymous_L1559I |
| 4990 | C | T | 1.26E-04 | | | | | ORF1ab | synonymous_D1575D |
| 534 | C | T | 1.26E-04 | | | | | ORF1ab | nonsynonymous_A90V |
| 5665 | G | A | 1.26E-04 | | | | | ORF1ab | synonymous_Q1800Q |
| 5672 | C | T | 1.26E-04 | | | | | ORF1ab | nonsynonymous_P1803S |
| 5700 | C | A | 1.26E-04 | | | | | ORF1ab | nonsynonymous_A1812D |
| 5777 | C | T | 1.26E-04 | | | | | ORF1ab | nonsynonymous_H1838Y |
| 5838 | A | C | 1.26E-04 | G | 6.32E-05 | | | ORF1ab | nonsynonymous_E1858A; nonsynonymous_E1858G |
| 5886 | C | T | 1.26E-04 | | | | | ORF1ab | nonsynonymous_T1874I |
| 5974 | C | T | 1.26E-04 | | | | | ORF1ab | synonymous_D1903D |
| 5977 | T | C | 1.26E-04 | | | | | ORF1ab | synonymous_N1904N |
| 6025 | T | C | 1.26E-04 | | | | | ORF1ab | synonymous_Y1920Y |
| 6035 | A | G | 1.26E-04 | | | | | ORF1ab | nonsynonymous_S1924G |
| 611 | G | A | 1.26E-04 | | | | | ORF1ab | nonsynonymous_V116M |
| 614 | G | A | 1.26E-04 | T | 6.32E-05 | | | ORF1ab | nonsynonymous_A117T; nonsynonymous_A117S |
| 6178 | T | C | 1.26E-04 | | | | | ORF1ab | synonymous_D1971D |
| 621 | G | A | 1.26E-04 | | | | | ORF1ab | nonsynonymous_R119H |
| 6220 | G | A | 1.26E-04 | | | | | ORF1ab | synonymous_L1985L |
| 673 | C | T | 1.26E-04 | | | | | ORF1ab | synonymous_Y136Y |
| 697 | C | T | 1.26E-04 | | | | | ORF1ab | synonymous_D144D |
| 701 | G | T | 1.26E-04 | | | | | ORF1ab | nonsynonymous_G146C |
| 715 | C | T | 1.26E-04 | | | | | ORF1ab | synonymous_G150G |
| 7162 | C | T | 1.26E-04 | | | | | ORF1ab | synonymous_D2299D |

| | | | | | | | | | |
|---|---|---|---|---|---|---|---|---|---|
| 7173 | C | T | 1.26E-04 | | | | | ORF1ab | nonsynonymous_S2303F |
| 7214 | G | A | 1.26E-04 | | | | | ORF1ab | nonsynonymous_D2317N |
| 7217 | T | C | 1.26E-04 | | | | | ORF1ab | synonymous_L2318L |
| 7231 | C | T | 1.26E-04 | | | | | ORF1ab | synonymous_G2322G |
| 7452 | G | T | 1.26E-04 | | | | | ORF1ab | nonsynonymous_S2396I |
| 7672 | T | C | 1.26E-04 | | | | | ORF1ab | synonymous_V2469V |
| 8118 | C | T | 1.26E-04 | | | | | ORF1ab | nonsynonymous_A2618V |
| 815 | C | T | 1.26E-04 | | | | | ORF1ab | nonsynonymous_R184C |
| 8208 | C | T | 1.26E-04 | | | | | ORF1ab | nonsynonymous_T2648I |
| 8240 | C | T | 1.26E-04 | | | | | ORF1ab | nonsynonymous_H2659Y |
| 8247 | C | T | 1.26E-04 | | | | | ORF1ab | nonsynonymous_S2661F |
| 8266 | C | T | 1.26E-04 | | | | | ORF1ab | synonymous_G2667G |
| 8274 | G | A | 1.26E-04 | | | | | ORF1ab | nonsynonymous_C2670Y |
| 8311 | C | T | 1.26E-04 | | | | | ORF1ab | synonymous_N2682N |
| 8316 | C | T | 1.26E-04 | | | | | ORF1ab | nonsynonymous_T2684I |
| 8318 | C | A | 1.26E-04 | | | | | ORF1ab | nonsynonymous_P2685T |
| 8320 | C | T | 1.26E-04 | | | | | ORF1ab | synonymous_P2685P |
| 845 | G | T | 1.26E-04 | | | | | ORF1ab | nonsynonymous_D194Y |
| 8481 | C | T | 1.26E-04 | | | | | ORF1ab | nonsynonymous_P2739L |
| 850 | C | T | 1.26E-04 | | | | | ORF1ab | synonymous_G195G |
| 8600 | G | T | 1.26E-04 | | | | | ORF1ab | nonsynonymous_V2779L |
| 881 | G | A | 1.26E-04 | | | | | ORF1ab | nonsynonymous_A206T |
| 8987 | T | A | 1.26E-04 | C | 6.32E-05 | | | ORF1ab | nonsynonymous_F2908I; nonsynonymous_F2908L |
| 9067 | T | C | 1.26E-04 | | | | | ORF1ab | synonymous_Y2934Y |
| 9118 | C | T | 1.26E-04 | | | | | ORF1ab | synonymous_D2951D |
| 9124 | T | C | 1.26E-04 | | | | | ORF1ab | synonymous_R2953R |
| 9145 | T | A | 1.26E-04 | | | | | ORF1ab | synonymous_S2960S |
| 9180 | C | T | 1.26E-04 | | | | | ORF1ab | nonsynonymous_S2972F |

| | | | | | | | | | |
|---|---|---|---|---|---|---|---|---|---|
| 9242 | G | T | 1.26E-04 | C | 6.32E-05 | | | ORF1ab | stopgain_E2993X; nonsynonymous_E2993Q |
| 9505 | T | C | 1.26E-04 | | | | | ORF1ab | synonymous_F3080F |
| 9510 | C | T | 1.26E-04 | | | | | ORF1ab | nonsynonymous_T3082I |
| 9518 | T | C | 1.26E-04 | | | | | ORF1ab | nonsynonymous_F3085L |
| 9697 | T | C | 1.26E-04 | | | | | ORF1ab | synonymous_Y3144Y |
| 9743 | T | C | 1.26E-04 | | | | | ORF1ab | nonsynonymous_Y3160H |
| 9811 | C | T | 1.26E-04 | | | | | ORF1ab | synonymous_T3182T |
| 9836 | C | T | 1.26E-04 | | | | | ORF1ab | synonymous_L3191L |
| 9943 | C | T | 1.26E-04 | | | | | ORF1ab | synonymous_Y3226Y |
| 9946 | A | G | 1.26E-04 | | | | | ORF1ab | synonymous_R3227R |
| 994 | A | G | 1.26E-04 | | | | | ORF1ab | synonymous_E243E |
| 9968 | G | A | 1.26E-04 | | | | | ORF1ab | nonsynonymous_A3235T |
| 9995 | T | G | 1.26E-04 | | | | | ORF1ab | nonsynonymous_S3244A |
| 43 | T | C | 1.05E-04 | | | | | five_prime_UTR | |
| 46 | A | G | 1.03E-04 | | | | | five_prime_UTR | |
| 47 | T | C | 1.03E-04 | | | | | five_prime_UTR | |
| 49 | T | A | 1.02E-04 | | | | | five_prime_UTR | |
| 51 | T | C | 1.01E-04 | | | | | five_prime_UTR | |
| 52 | T | G | 1.01E-04 | G | 1.01E-04 | | | five_prime_UTR | |
| 53 | G | T | 1.01E-04 | | | | | five_prime_UTR | |
| 29792 | A | G | 7.86E-05 | | | | | three_prime_UTR | |
| 29833 | A | C | 7.09E-05 | G | 7.09E-05 | | | three_prime_UTR | |
| 29819 | A | T | 6.82E-05 | | | | | three_prime_UTR | |
| 29814 | A | C | 6.81E-05 | | | | | three_prime_UTR | |
| 29812 | A | C | 6.81E-05 | | | | | three_prime_UTR | |
| 29788 | C | A | 6.73E-05 | A | 6.73E-05 | | | three_prime_UTR | |
| 29787 | C | A | 6.73E-05 | | | | | three_prime_UTR | |
| 29785 | T | G | 6.72E-05 | | | | | three_prime_UTR | |
| 29790 | A | C | 6.72E-05 | | | | | three_prime_UTR | |

| | | | | | | | | | |
|---|---|---|---|---|---|---|---|---|---|
| 29780 | A | G | 6.68E-05 | G | 6.68E-05 | | | three_prime_UTR | |
| 57 | A | T | 6.63E-05 | T | 6.63E-05 | | | five_prime_UTR | |
| 58 | T | A | 6.63E-05 | | | | | five_prime_UTR | |
| 60 | T | C | 6.62E-05 | | | | | five_prime_UTR | |
| 62 | T | A | 6.59E-05 | | | | | five_prime_UTR | |
| 63 | T | A | 6.57E-05 | | | | | five_prime_UTR | |
| 65 | T | C | 6.53E-05 | | | | | five_prime_UTR | |
| 68 | A | T | 6.52E-05 | | | | | five_prime_UTR | |
| 71 | C | A | 6.52E-05 | | | | | five_prime_UTR | |
| 69 | A | G | 6.52E-05 | | | | | five_prime_UTR | |
| 70 | A | T | 6.52E-05 | | | | | five_prime_UTR | |
| 74 | A | C | 6.52E-05 | | | | | five_prime_UTR | |
| 77 | T | C | 6.50E-05 | A | 6.50E-05 | | | five_prime_UTR | |
| 75 | C | A | 6.50E-05 | | | | | five_prime_UTR | |
| 79 | A | G | 6.49E-05 | C | 6.49E-05 | | | five_prime_UTR | |
| 80 | A | T | 6.49E-05 | T | 6.49E-05 | | | five_prime_UTR | |
| 82 | A | T | 6.48E-05 | | | | | five_prime_UTR | |
| 86 | G | A | 6.48E-05 | | | | | five_prime_UTR | |
| 90 | G | T | 6.47E-05 | | | | | five_prime_UTR | |
| 96 | C | A | 6.47E-05 | | | | | five_prime_UTR | |
| 97 | A | G | 6.47E-05 | | | | | five_prime_UTR | |
| 1606 | T | C | 6.46E-05 | | | | | ORF1ab | synonymous_N447N |
| 1607 | G | A | 6.46E-05 | | | | | ORF1ab | nonsynonymous_D448N |
| 103 | C | T | 6.45E-05 | | | | | five_prime_UTR | |
| 104 | T | A | 6.45E-05 | | | | | five_prime_UTR | |
| 111 | T | A | 6.43E-05 | C | 6.43E-05 | | | five_prime_UTR | |
| 112 | T | G | 6.43E-05 | | | | | five_prime_UTR | |
| 115 | T | C | 6.43E-05 | | | | | five_prime_UTR | |
| 118 | A | G | 6.43E-05 | | | | | five_prime_UTR | |
| 119 | C | G | 6.43E-05 | | | | | five_prime_UTR | |

| | | | | | | | | |
|---|---|---|---|---|---|---|---|---|
| 121 | C | T | 6.42E-05 | | | | | five_prime_UTR |
| 123 | C | G | 6.42E-05 | | | | | five_prime_UTR |
| 122 | A | T | 6.42E-05 | | | | | five_prime_UTR |
| 128 | T | C | 6.42E-05 | | | | | five_prime_UTR |
| 142 | A | T | 6.41E-05 | T | 6.41E-05 | | | five_prime_UTR |
| 29760 | T | A | 6.41E-05 | | | | | three_prime_UTR |
| 147 | C | A | 6.41E-05 | A | 6.41E-05 | | | five_prime_UTR |
| 29768 | A | C | 6.41E-05 | | | | | three_prime_UTR |
| 29758 | T | C | 6.40E-05 | A | 6.40E-05 | | | three_prime_UTR |
| 149 | G | A | 6.40E-05 | A | 6.40E-05 | | | five_prime_UTR |
| 29756 | A | T | 6.40E-05 | | | | | three_prime_UTR |
| 29757 | G | A | 6.40E-05 | | | | | three_prime_UTR |
| 152 | G | C | 6.39E-05 | | | | | five_prime_UTR |
| 154 | T | A | 6.39E-05 | | | | | five_prime_UTR |
| 158 | A | C | 6.39E-05 | | | | | five_prime_UTR |
| 159 | G | A | 6.39E-05 | | | | | five_prime_UTR |
| 162 | C | A | 6.39E-05 | | | | | five_prime_UTR |
| 156 | A | G | 6.39E-05 | | | | | five_prime_UTR |
| 163 | A | G | 6.39E-05 | C | 6.39E-05 | | | five_prime_UTR |
| 165 | G | A | 6.39E-05 | A | 6.39E-05 | | | five_prime_UTR |
| 29746 | A | G | 6.39E-05 | | | | | three_prime_UTR |
| 29748 | T | C | 6.39E-05 | | | | | three_prime_UTR |
| 164 | C | G | 6.39E-05 | | | | | five_prime_UTR |
| 169 | A | C | 6.39E-05 | | | | | five_prime_UTR |
| 171 | C | A | 6.39E-05 | | | | | five_prime_UTR |
| 172 | T | C | 6.39E-05 | | | | | five_prime_UTR |
| 191 | T | C | 6.39E-05 | | | | | five_prime_UTR |
| 29740 | A | T | 6.39E-05 | | | | | three_prime_UTR |
| 190 | C | G | 6.38E-05 | | | | | five_prime_UTR |
| 180 | C | T | 6.38E-05 | | | | | five_prime_UTR |

| | | | | | | | | |
|---|---|---|---|---|---|---|---|---|
| 224 | T | A | 6.38E-05 | | | | | five_prime_UTR |
| 227 | G | A | 6.38E-05 | | | | | five_prime_UTR |
| 176 | C | T | 6.38E-05 | | | | | five_prime_UTR |
| 202 | T | G | 6.38E-05 | G | 6.38E-05 | | | five_prime_UTR |
| 223 | A | G | 6.38E-05 | | | | | five_prime_UTR |
| 197 | C | T | 6.38E-05 | | | | | five_prime_UTR |
| 201 | T | A | 6.38E-05 | | | | | five_prime_UTR |
| 200 | T | C | 6.38E-05 | | | | | five_prime_UTR |
| 205 | T | G | 6.38E-05 | G | 6.38E-05 | | | five_prime_UTR |
| 246 | G | T | 6.38E-05 | | | | | five_prime_UTR |
| 206 | C | G | 6.38E-05 | | | | | five_prime_UTR |
| 252 | G | T | 6.38E-05 | | | | | five_prime_UTR |
| 29735 | A | C | 6.38E-05 | | | | | three_prime_UTR |
| 212 | T | A | 6.38E-05 | A | 6.38E-05 | | | five_prime_UTR |
| 244 | C | T | 6.38E-05 | | | | | five_prime_UTR |
| 247 | G | T | 6.38E-05 | | | | | five_prime_UTR |
| 248 | G | T | 6.38E-05 | | | | | five_prime_UTR |
| 29716 | A | T | 6.38E-05 | T | 6.38E-05 | | | three_prime_UTR |
| 237 | G | T | 6.38E-05 | | | | | five_prime_UTR |
| 29707 | A | T | 6.38E-05 | | | | | three_prime_UTR |
| 235 | A | G | 6.38E-05 | | | | | five_prime_UTR |
| 29714 | A | G | 6.38E-05 | | | | | three_prime_UTR |
| 29702 | G | A | 6.38E-05 | | | | | three_prime_UTR |
| 29729 | T | G | 6.38E-05 | | | | | three_prime_UTR |
| 29703 | G | C | 6.37E-05 | | | | | three_prime_UTR |
| 256 | G | A | 6.37E-05 | | | | | five_prime_UTR |
| 29728 | T | G | 6.37E-05 | G | 6.37E-05 | | | three_prime_UTR |
| 29725 | A | G | 6.37E-05 | | | | | three_prime_UTR |
| 29698 | T | A | 6.37E-05 | | | | | three_prime_UTR |
| 29697 | A | T | 6.37E-05 | T | 6.37E-05 | | | three_prime_UTR |

| | | | | | | | | | |
|---|---|---|---|---|---|---|---|---|---|
| 29720 | A | G | 6.37E-05 | | | | | three_prime_UTR | |
| 29696 | C | T | 6.37E-05 | | | | | three_prime_UTR | |
| 29693 | T | C | 6.36E-05 | | | | | three_prime_UTR | |
| 260 | G | C | 6.36E-05 | | | | | five_prime_UTR | |
| 261 | G | T | 6.36E-05 | T | 6.36E-05 | | | five_prime_UTR | |
| 29685 | T | A | 6.36E-05 | A | 6.36E-05 | | | three_prime_UTR | |
| 29678 | T | C | 6.36E-05 | | | | | three_prime_UTR | |
| 29677 | A | G | 6.35E-05 | | | | | three_prime_UTR | |
| 29676 | A | G | 6.35E-05 | | | | | three_prime_UTR | |
| 518 | A | G | 6.35E-05 | | | | | ORF1ab | nonsynonymous_M85V |
| 25202 | T | C | 6.34E-05 | | | | | S | nonsynonymous_W1214R |
| 694 | T | A | 6.34E-05 | | | | | ORF1ab | nonsynonymous_F143L |
| 22516 | T | C | 6.34E-05 | | | | | S | synonymous_F318F |
| 22340 | G | T | 6.34E-05 | | | | | S | nonsynonymous_A260S |
| 22515 | T | C | 6.34E-05 | | | | | S | nonsynonymous_F318S |
| 510 | G | A | 6.34E-05 | | | | | ORF1ab | nonsynonymous_G82D |
| 19344 | T | C | 6.34E-05 | | | | | ORF1ab | synonymous_A6360A |
| 521 | G | A | 6.34E-05 | | | | | ORF1ab | nonsynonymous_V86I |
| 19329 | T | C | 6.34E-05 | | | | | ORF1ab | synonymous_A6355A |
| 22447 | T | C | 6.33E-05 | | | | | S | synonymous_P295P |
| 29669 | A | T | 6.33E-05 | | | | | ORF10 | nonsynonymous_T38S |
| 19984 | T | C | 6.33E-05 | | | | | ORF1ab | nonsynonymous_F6574L |
| 27578 | A | T | 6.33E-05 | | | | | ORF7a | nonsynonymous_Q62L |
| 29667 | T | C | 6.33E-05 | | | | | ORF10 | nonsynonymous_L37P |
| 27516 | G | T | 6.33E-05 | | | | | ORF7a | nonsynonymous_E41D |
| 27603 | C | A | 6.33E-05 | A | 6.33E-05 | | | ORF7a | synonymous_G70G; synonymous_G70G |
| 27612 | C | T | 6.33E-05 | T | 6.33E-05 | | | ORF7a | synonymous_H73H; synonymous_H73H |
| 27617 | A | G | 6.33E-05 | | | | | ORF7a | nonsynonymous_Y75C |

| | | | | | | | | | |
|---|---|---|---|---|---|---|---|---|---|
| 29673 | A | C | 6.33E-05 | | | | | ORF10 | stoploss_X39S |
| 29674 | G | T | 6.33E-05 | | | | | ORF10 | stoploss_X39Y |
| 27572 | G | T | 6.33E-05 | | | | | ORF7a | nonsynonymous_S60I |
| 27573 | C | T | 6.33E-05 | | | | | ORF7a | synonymous_S60S |
| 27615 | C | G | 6.33E-05 | | | | | ORF7a | synonymous_V74V |
| 27621 | G | A | 6.33E-05 | A | 6.33E-05 | | | ORF7a | synonymous_Q76Q; synonymous_Q76Q |
| 27791 | C | T | 6.33E-05 | | | | | ORF7b | synonymous_C12C |
| 29672 | T | C | 6.33E-05 | | | | | ORF10 | stoploss_X39Q |
| 523 | T | G | 6.33E-05 | | | | | ORF1ab | synonymous_V86V |
| 27652 | T | C | 6.33E-05 | | | | | ORF7a | nonsynonymous_F87L |
| 19355 | T | G | 6.33E-05 | | | | | ORF1ab | stopgain_L6364X |
| 27514 | G | T | 6.33E-05 | | | | | ORF7a | stopgain_E41X |
| 27622 | T | G | 6.33E-05 | | | | | ORF7a | nonsynonymous_L77V |
| 27641 | C | T | 6.33E-05 | | | | | ORF7a | nonsynonymous_S83L |
| 27657 | C | T | 6.33E-05 | | | | | ORF7a | synonymous_I88I |
| 19398 | G | T | 6.33E-05 | | | | | ORF1ab | nonsynonymous_E6378D |
| 19407 | A | G | 6.33E-05 | | | | | ORF1ab | synonymous_G6381G |
| 26709 | G | A | 6.33E-05 | A | 6.33E-05 | | | M | nonsynonymous_A63T; nonsynonymous_A63T |
| 27522 | T | C | 6.33E-05 | | | | | ORF7a | synonymous_N43N |
| 27564 | T | G | 6.33E-05 | | | | | ORF7a | synonymous_T57T |
| 27623 | T | A | 6.33E-05 | | | | | ORF7a | stopgain_L77X |
| 27624 | A | G | 6.33E-05 | | | | | ORF7a | synonymous_L77L |
| 27629 | C | T | 6.33E-05 | | | | | ORF7a | nonsynonymous_A79V |
| 27633 | A | G | 6.33E-05 | | | | | ORF7a | synonymous_R80R |
| 27645 | T | C | 6.33E-05 | | | | | ORF7a | synonymous_P84P |
| 27646 | A | C | 6.33E-05 | | | | | ORF7a | nonsynonymous_K85Q |
| 27659 | G | T | 6.33E-05 | | | | | ORF7a | nonsynonymous_R89I |
| 27680 | T | G | 6.33E-05 | | | | | ORF7a | nonsynonymous_L96R |

| | | | | | | | | | |
|---|---|---|---|---|---|---|---|---|---|
| 27707 | C | T | 6.33E-05 | | | | | ORF7a | nonsynonymous_A105V |
| 27708 | G | T | 6.33E-05 | | | | | ORF7a | synonymous_A105A |
| 27784 | A | G | 6.33E-05 | | | | | ORF7b | nonsynonymous_Y10C |
| 503 | C | T | 6.33E-05 | | | | | ORF1ab | nonsynonymous_P80S |
| 19421 | C | T | 6.33E-05 | | | | | ORF1ab | nonsynonymous_S6386L |
| 27515 | A | G | 6.33E-05 | | | | | ORF7a | nonsynonymous_E41G |
| 27555 | T | C | 6.33E-05 | | | | | ORF7a | synonymous_F54F |
| 27556 | G | T | 6.33E-05 | | | | | ORF7a | nonsynonymous_A55S |
| 27560 | T | C | 6.33E-05 | | | | | ORF7a | nonsynonymous_L56P |
| 27562 | A | T | 6.33E-05 | T | 6.33E-05 | | | ORF7a | nonsynonymous_T57S; nonsynonymous_T57S |
| 27563 | C | T | 6.33E-05 | | | | | ORF7a | nonsynonymous_T57I |
| 27565 | T | A | 6.33E-05 | | | | | ORF7a | nonsynonymous_C58S |
| 27626 | G | A | 6.33E-05 | | | | | ORF7a | nonsynonymous_R78H |
| 27631 | A | T | 6.33E-05 | | | | | ORF7a | stopgain_R80X |
| 27632 | G | A | 6.33E-05 | | | | | ORF7a | nonsynonymous_R80K |
| 27671 | T | G | 6.33E-05 | | | | | ORF7a | nonsynonymous_V93G |
| 27686 | C | T | 6.33E-05 | | | | | ORF7a | nonsynonymous_S98F |
| 27689 | C | T | 6.33E-05 | | | | | ORF7a | nonsynonymous_P99L |
| 27735 | C | T | 6.33E-05 | | | | | ORF7a | synonymous_F114F |
| 27736 | A | G | 6.33E-05 | | | | | ORF7a | nonsynonymous_T115A |
| 27759 | A | G | 6.33E-05 | | | | | ORF7a | stoploss_X122W |
| 27761 | T | C | 6.33E-05 | | | | | ORF7b | synonymous_I2I |
| 27785 | T | C | 6.33E-05 | | | | | ORF7b | synonymous_Y10Y |
| 28215 | T | G | 6.33E-05 | | | | | ORF8 | nonsynonymous_F108V |
| 29387 | A | T | 6.33E-05 | | | | | N | stopgain_K372X |
| 19449 | T | C | 6.33E-05 | | | | | ORF1ab | synonymous_S6395S |
| 22384 | T | C | 6.33E-05 | | | | | S | synonymous_T274T |
| 27554 | T | C | 6.33E-05 | | | | | ORF7a | nonsynonymous_F54S |
| 19536 | C | T | 6.33E-05 | | | | | ORF1ab | synonymous_N6424N |

| | | | | | | | | | |
|---|---|---|---|---|---|---|---|---|---|
| 22350 | C | T | 6.33E-05 | | | | | S | nonsynonymous_A263V |
| 27519 | C | T | 6.33E-05 | | | | | ORF7a | synonymous_G42G |
| 27524 | C | T | 6.33E-05 | | | | | ORF7a | nonsynonymous_S44L |
| 27527 | C | T | 6.33E-05 | | | | | ORF7a | nonsynonymous_P45L |
| 27535 | C | T | 6.33E-05 | | | | | ORF7a | nonsynonymous_P48S |
| 27536 | C | A | 6.33E-05 | | | | | ORF7a | nonsynonymous_P48H |
| 27538 | C | T | 6.33E-05 | | | | | ORF7a | synonymous_L49L |
| 27553 | T | G | 6.33E-05 | G | 6.33E-05 | | | ORF7a | nonsynonymous_F54V; nonsynonymous_F54V |
| 27762 | G | A | 6.33E-05 | A | 6.33E-05 | | | ORF7b | nonsynonymous_E3K; nonsynonymous_E3K |
| 19300 | G | T | 6.33E-05 | | | | | ORF1ab | nonsynonymous_V6346L |
| 19327 | G | A | 6.33E-05 | A | 6.33E-05 | | | ORF1ab | nonsynonymous_A6355T; nonsynonymous_A6355T |
| 19343 | C | T | 6.33E-05 | | | | | ORF1ab | nonsynonymous_A6360V |
| 19403 | A | G | 6.33E-05 | | | | | ORF1ab | nonsynonymous_H6380R |
| 19420 | T | C | 6.33E-05 | | | | | ORF1ab | nonsynonymous_S6386P |
| 19479 | T | C | 6.33E-05 | | | | | ORF1ab | synonymous_G6405G |
| 19519 | T | A | 6.33E-05 | | | | | ORF1ab | nonsynonymous_Y6419N |
| 19547 | C | T | 6.33E-05 | | | | | ORF1ab | nonsynonymous_S6428L |
| 22393 | A | G | 6.33E-05 | | | | | S | synonymous_L277L |
| 27548 | A | G | 6.33E-05 | | | | | ORF7a | nonsynonymous_N52S |
| 27676 | G | C | 6.33E-05 | | | | | ORF7a | nonsynonymous_E95Q |
| 27678 | A | G | 6.33E-05 | | | | | ORF7a | synonymous_E95E |
| 27750 | G | C | 6.33E-05 | | | | | ORF7a | nonsynonymous_K119N |
| 27758 | G | T | 6.33E-05 | | | | | ORF7a | stoploss_X122L |
| 27763 | A | G | 6.33E-05 | | | | | ORF7b | nonsynonymous_E3G |
| 27775 | T | C | 6.33E-05 | | | | | ORF7b | nonsynonymous_I7T |
| 27776 | T | G | 6.33E-05 | | | | | ORF7b | nonsynonymous_I7M |

| | | | | | | | | | |
|---|---|---|---|---|---|---|---|---|---|
| 27777 | G | A | 6.33E-05 | A | 6.33E-05 | | | ORF7b | nonsynonymous_D8N; nonsynonymous_D8N |
| 27803 | T | C | 6.33E-05 | | | | | ORF7b | synonymous_F16F |
| 28057 | C | T | 6.33E-05 | | | | | ORF8 | nonsynonymous_A55V |
| 28090 | G | C | 6.33E-05 | | | | | ORF8 | nonsynonymous_G66A |
| 28094 | T | C | 6.33E-05 | | | | | ORF8 | synonymous_S67S |
| 28212 | G | A | 6.33E-05 | | | | | ORF8 | nonsynonymous_D107N |
| 19352 | A | C | 6.33E-05 | | | | | ORF1ab | nonsynonymous_N6363T |
| 19367 | C | T | 6.33E-05 | | | | | ORF1ab | nonsynonymous_P6368L |
| 19414 | G | A | 6.33E-05 | A | 6.33E-05 | | | ORF1ab | nonsynonymous_V6384I; nonsynonymous_V6384I |
| 19559 | G | T | 6.33E-05 | | | | | ORF1ab | nonsynonymous_S6432I |
| 22351 | A | T | 6.33E-05 | | | | | S | synonymous_A263A |
| 27748 | A | C | 6.33E-05 | | | | | ORF7a | nonsynonymous_K119Q |
| 27757 | T | A | 6.33E-05 | | | | | ORF7a | stoploss_X122R |
| 27919 | T | C | 6.33E-05 | | | | | ORF8 | nonsynonymous_I9T |
| 28008 | A | G | 6.33E-05 | | | | | ORF8 | nonsynonymous_I39V |
| 28028 | G | T | 6.33E-05 | | | | | ORF8 | nonsynonymous_W45C |
| 28052 | A | G | 6.33E-05 | | | | | ORF8 | synonymous_K53K |
| 28085 | G | T | 6.33E-05 | T | 6.33E-05 | | | ORF8 | nonsynonymous_E64D; nonsynonymous_E64D |
| 28089 | G | A | 6.33E-05 | A | 6.33E-05 | | | ORF8 | nonsynonymous_G66S; nonsynonymous_G66S |
| 28214 | C | A | 6.33E-05 | | | | | ORF8 | nonsynonymous_D107E |
| 19331 | T | G | 6.33E-05 | | | | | ORF1ab | nonsynonymous_F6356C |
| 19455 | G | A | 6.33E-05 | | | | | ORF1ab | synonymous_T6397T |
| 19517 | T | C | 6.33E-05 | | | | | ORF1ab | nonsynonymous_L6418S |
| 21757 | C | T | 6.33E-05 | | | | | S | synonymous_F65F |
| 21761 | G | T | 6.33E-05 | | | | | S | nonsynonymous_A67S |
| 22380 | G | A | 6.33E-05 | | | | | S | nonsynonymous_R273K |

| | | | | | | | | | |
|---|---|---|---|---|---|---|---|---|---|
| 27916 | G | A | 6.33E-05 | A | 6.33E-05 | | | ORF8 | nonsynonymous_G8E; nonsynonymous_G8E |
| 27922 | T | G | 6.33E-05 | G | 6.33E-05 | | | ORF8 | nonsynonymous_I10S; nonsynonymous_I10S |
| 27926 | A | G | 6.33E-05 | | | | | ORF8 | synonymous_T11T |
| 27937 | C | T | 6.33E-05 | | | | | ORF8 | nonsynonymous_A15V |
| 27938 | A | G | 6.33E-05 | | | | | ORF8 | synonymous_A15A |
| 27942 | C | T | 6.33E-05 | | | | | ORF8 | nonsynonymous_H17Y |
| 27944 | C | T | 6.33E-05 | | | | | ORF8 | synonymous_H17H |
| 27977 | T | G | 6.33E-05 | | | | | ORF8 | nonsynonymous_H28Q |
| 27983 | A | T | 6.33E-05 | | | | | ORF8 | synonymous_P30P |
| 27985 | A | G | 6.33E-05 | | | | | ORF8 | nonsynonymous_Y31C |
| 27986 | T | A | 6.33E-05 | A | 6.33E-05 | | | ORF8 | stopgain_Y31X; stopgain_Y31X |
| 27989 | A | G | 6.33E-05 | | | | | ORF8 | synonymous_V32V |
| 27990 | G | A | 6.33E-05 | | | | | ORF8 | nonsynonymous_V33I |
| 27996 | G | T | 6.33E-05 | | | | | ORF8 | nonsynonymous_D35Y |
| 27998 | C | T | 6.33E-05 | | | | | ORF8 | synonymous_D35D |
| 28026 | T | A | 6.33E-05 | | | | | ORF8 | nonsynonymous_W45R |
| 28055 | A | G | 6.33E-05 | | | | | ORF8 | synonymous_S54S |
| 28059 | C | T | 6.33E-05 | | | | | ORF8 | nonsynonymous_P56S |
| 28061 | T | C | 6.33E-05 | | | | | ORF8 | synonymous_P56P |
| 28067 | T | A | 6.33E-05 | | | | | ORF8 | synonymous_I58I |
| 28083 | G | T | 6.33E-05 | | | | | ORF8 | stopgain_E64X |
| 28088 | T | C | 6.33E-05 | | | | | ORF8 | synonymous_A65A |
| 28102 | C | T | 6.33E-05 | | | | | ORF8 | nonsynonymous_P70L |
| 28104 | A | T | 6.33E-05 | | | | | ORF8 | nonsynonymous_I71F |
| 28115 | C | T | 6.33E-05 | | | | | ORF8 | synonymous_I74I |
| 28116 | G | A | 6.33E-05 | | | | | ORF8 | nonsynonymous_D75N |
| 28119 | A | G | 6.33E-05 | | | | | ORF8 | nonsynonymous_I76V |

| | | | | | | | | | |
|---|---|---|---|---|---|---|---|---|---|
| 28122 | G | A | 6.33E-05 | A | 6.33E-05 | | | ORF8 | nonsynonymous_G77S; nonsynonymous_G77S |
| 28135 | T | C | 6.33E-05 | | | | | ORF8 | nonsynonymous_V81A |
| 28145 | A | C | 6.33E-05 | | | | | ORF8 | nonsynonymous_L84F |
| 28151 | T | C | 6.33E-05 | | | | | ORF8 | synonymous_F86F |
| 28157 | T | C | 6.33E-05 | | | | | ORF8 | synonymous_I88I |
| 28160 | T | C | 6.33E-05 | | | | | ORF8 | synonymous_N89N |
| 28164 | C | A | 6.33E-05 | | | | | ORF8 | nonsynonymous_Q91K |
| 28171 | C | T | 6.33E-05 | | | | | ORF8 | nonsynonymous_P93L |
| 28183 | G | A | 6.33E-05 | | | | | ORF8 | nonsynonymous_S97N |
| 28191 | G | T | 6.33E-05 | | | | | ORF8 | nonsynonymous_V100L |
| 28194 | C | T | 6.33E-05 | | | | | ORF8 | nonsynonymous_R101C |
| 28199 | T | C | 6.33E-05 | | | | | ORF8 | synonymous_C102C |
| 28225 | A | G | 6.33E-05 | | | | | ORF8 | nonsynonymous_Y111C |
| 19480 | G | A | 6.33E-05 | | | | | ORF1ab | nonsynonymous_G6406S |
| 19492 | A | G | 6.33E-05 | | | | | ORF1ab | nonsynonymous_R6410G |
| 19560 | C | T | 6.33E-05 | | | | | ORF1ab | synonymous_S6432S |
| 22449 | T | A | 6.33E-05 | | | | | S | nonsynonymous_L296H |
| 29427 | G | T | 6.33E-05 | | | | | N | nonsynonymous_R385I |
| 19285 | G | A | 6.33E-05 | | | | | ORF1ab | nonsynonymous_G6341S |
| 22381 | G | T | 6.33E-05 | | | | | S | nonsynonymous_R273S |
| 22474 | A | G | 6.33E-05 | | | | | S | synonymous_K304K |
| 27852 | G | T | 6.33E-05 | | | | | ORF7b | stopgain_E33X |
| 27860 | A | G | 6.33E-05 | | | | | ORF7b | synonymous_Q35Q |
| 27861 | G | T | 6.33E-05 | | | | | ORF7b | nonsynonymous_D36Y |
| 27863 | T | C | 6.33E-05 | | | | | ORF7b | synonymous_D36D |
| 27864 | C | T | 6.33E-05 | | | | | ORF7b | nonsynonymous_H37Y |
| 27875 | T | C | 6.33E-05 | | | | | ORF7b | synonymous_T40T |
| 27880 | A | C | 6.33E-05 | | | | | ORF7b | nonsynonymous_H42P |
| 27890 | G | T | 6.33E-05 | | | | | intergenic | |

| | | | | | | | | | |
|---|---|---|---|---|---|---|---|---|---|
| 27894 | A | G | 6.33E-05 | | | | | ORF8 | nonsynonymous_M1V |
| 27897 | A | G | 6.33E-05 | | | | | ORF8 | nonsynonymous_K2E |
| 27900 | T | C | 6.33E-05 | | | | | ORF8 | nonsynonymous_F3L |
| 27902 | T | C | 6.33E-05 | | | | | ORF8 | synonymous_F3F |
| 27906 | G | T | 6.33E-05 | | | | | ORF8 | nonsynonymous_V5F |
| 18109 | C | T | 6.33E-05 | | | | | ORF1ab | nonsynonymous_P5949S |
| 22361 | G | T | 6.33E-05 | | | | | S | nonsynonymous_V267L |
| 29560 | G | T | 6.33E-05 | | | | | ORF10 | nonsynonymous_M1I |
| 18506 | G | A | 6.33E-05 | | | | | ORF1ab | nonsynonymous_G6081E |
| 21245 | A | G | 6.33E-05 | | | | | ORF1ab | nonsynonymous_N6994S |
| 21263 | C | T | 6.33E-05 | | | | | ORF1ab | nonsynonymous_S7000F |
| 22391 | T | C | 6.33E-05 | | | | | S | synonymous_L277L |
| 22414 | C | T | 6.33E-05 | | | | | S | synonymous_T284T |
| 22419 | C | T | 6.33E-05 | | | | | S | nonsynonymous_T286I |
| 29382 | A | G | 6.33E-05 | | | | | N | nonsynonymous_K370R |
| 29552 | A | G | 6.33E-05 | | | | | intergenic | |
| 3144 | T | C | 6.33E-05 | | | | | ORF1ab | nonsynonymous_L960S |
| 22455 | A | G | 6.33E-05 | | | | | S | nonsynonymous_E298G |
| 2586 | T | A | 6.33E-05 | | | | | ORF1ab | nonsynonymous_V774D |
| 2588 | G | A | 6.33E-05 | | | | | ORF1ab | nonsynonymous_E775K |
| 2591 | G | A | 6.33E-05 | | | | | ORF1ab | nonsynonymous_A776T |
| 2592 | C | A | 6.33E-05 | A | 6.33E-05 | | | ORF1ab | nonsynonymous_A776D; nonsynonymous_A776D |
| 2737 | T | A | 6.33E-05 | | | | | ORF1ab | synonymous_T824T |
| 29428 | A | G | 6.33E-05 | | | | | N | synonymous_R385R |
| 29545 | C | A | 6.33E-05 | A | 6.33E-05 | | | intergenic | |
| 20472 | A | G | 6.33E-05 | | | | | ORF1ab | synonymous_Q6736Q |
| 20473 | A | C | 6.33E-05 | | | | | ORF1ab | nonsynonymous_T6737P |
| 21334 | G | T | 6.33E-05 | | | | | ORF1ab | nonsynonymous_A7024S |
| 21550 | A | C | 6.33E-05 | | | | | ORF1ab | nonsynonymous_N7096H |

| | | | | | | | | | |
|---|---|---|---|---|---|---|---|---|---|
| 22355 | T | A | 6.33E-05 | | | | | S | nonsynonymous_Y265N |
| 22441 | T | C | 6.33E-05 | | | | | S | synonymous_L293L |
| 22466 | A | G | 6.33E-05 | | | | | S | nonsynonymous_T302A |
| 22501 | T | C | 6.33E-05 | | | | | S | synonymous_Y313Y |
| 22511 | A | G | 6.33E-05 | | | | | S | nonsynonymous_N317D |
| 2668 | C | T | 6.33E-05 | | | | | ORF1ab | synonymous_A801A |
| 2707 | A | G | 6.33E-05 | | | | | ORF1ab | synonymous_T814T |
| 29562 | G | A | 6.33E-05 | | | | | ORF10 | nonsynonymous_G2D |
| 29570 | A | G | 6.33E-05 | | | | | ORF10 | nonsynonymous_N5D |
| 29575 | T | A | 6.33E-05 | | | | | ORF10 | synonymous_V6V |
| 5192 | C | T | 6.33E-05 | | | | | ORF1ab | synonymous_L1643L |
| 20436 | C | T | 6.33E-05 | | | | | ORF1ab | synonymous_D6724D |
| 21147 | T | C | 6.33E-05 | | | | | ORF1ab | synonymous_L6961L |
| 21159 | G | A | 6.33E-05 | | | | | ORF1ab | synonymous_V6965V |
| 21188 | A | G | 6.33E-05 | | | | | ORF1ab | nonsynonymous_N6975S |
| 21190 | G | A | 6.33E-05 | | | | | ORF1ab | nonsynonymous_A6976T |
| 21194 | A | G | 6.33E-05 | | | | | ORF1ab | nonsynonymous_D6977G |
| 21205 | C | A | 6.33E-05 | | | | | ORF1ab | nonsynonymous_L6981I |
| 21206 | T | G | 6.33E-05 | | | | | ORF1ab | nonsynonymous_L6981R |
| 21220 | G | A | 6.33E-05 | A | 6.33E-05 | | | ORF1ab | nonsynonymous_A6986T; nonsynonymous_A6986T |
| 21254 | C | T | 6.33E-05 | | | | | ORF1ab | nonsynonymous_A6997V |
| 21267 | A | G | 6.33E-05 | | | | | ORF1ab | synonymous_E7001E |
| 21268 | G | A | 6.33E-05 | A | 6.33E-05 | | | ORF1ab | nonsynonymous_A7002T; nonsynonymous_A7002T |
| 21315 | A | G | 6.33E-05 | | | | | ORF1ab | nonsynonymous_I7017M |
| 21316 | G | A | 6.33E-05 | | | | | ORF1ab | nonsynonymous_D7018N |
| 21342 | C | T | 6.33E-05 | | | | | ORF1ab | synonymous_Y7026Y |
| 21356 | A | G | 6.33E-05 | | | | | ORF1ab | nonsynonymous_N7031S |
| 21357 | T | G | 6.33E-05 | | | | | ORF1ab | nonsynonymous_N7031K |

| | | | | | | | | | |
|---|---|---|---|---|---|---|---|---|---|
| 21359 | C | T | 6.33E-05 | | | | | ORF1ab | nonsynonymous_T7032I |
| 21368 | T | C | 6.33E-05 | | | | | ORF1ab | nonsynonymous_I7035T |
| 21380 | C | T | 6.33E-05 | | | | | ORF1ab | nonsynonymous_S7039F |
| 22356 | A | G | 6.33E-05 | | | | | S | nonsynonymous_Y265C |
| 22424 | G | T | 6.33E-05 | | | | | S | nonsynonymous_A288S |
| 22482 | C | T | 6.33E-05 | | | | | S | nonsynonymous_T307I |
| 22498 | C | T | 6.33E-05 | | | | | S | synonymous_I312I |
| 2570 | C | T | 6.33E-05 | | | | | ORF1ab | nonsynonymous_P769S |
| 2572 | T | C | 6.33E-05 | | | | | ORF1ab | synonymous_P769P |
| 2574 | C | T | 6.33E-05 | | | | | ORF1ab | nonsynonymous_T770I |
| 25891 | A | G | 6.33E-05 | | | | | ORF3a | nonsynonymous_I167V |
| 2609 | C | T | 6.33E-05 | | | | | ORF1ab | nonsynonymous_P782S |
| 2627 | C | T | 6.33E-05 | | | | | ORF1ab | nonsynonymous_L788F |
| 2648 | G | C | 6.33E-05 | | | | | ORF1ab | nonsynonymous_D795H |
| 2651 | A | G | 6.33E-05 | | | | | ORF1ab | nonsynonymous_T796A |
| 2653 | A | G | 6.33E-05 | | | | | ORF1ab | synonymous_T796T |
| 2672 | G | T | 6.33E-05 | | | | | ORF1ab | nonsynonymous_A803S |
| 2674 | A | G | 6.33E-05 | | | | | ORF1ab | synonymous_A803A |
| 2680 | T | C | 6.33E-05 | | | | | ORF1ab | synonymous_N805N |
| 2685 | T | C | 6.33E-05 | | | | | ORF1ab | nonsynonymous_M807T |
| 2686 | G | A | 6.33E-05 | | | | | ORF1ab | nonsynonymous_M807I |
| 2697 | A | T | 6.33E-05 | | | | | ORF1ab | nonsynonymous_N811I |
| 2701 | C | T | 6.33E-05 | | | | | ORF1ab | synonymous_T812T |
| 2715 | G | T | 6.33E-05 | | | | | ORF1ab | nonsynonymous_G817V |
| 2716 | C | T | 6.33E-05 | | | | | ORF1ab | synonymous_G817G |
| 2717 | G | A | 6.33E-05 | | | | | ORF1ab | nonsynonymous_G818S |
| 2778 | G | A | 6.33E-05 | | | | | ORF1ab | nonsynonymous_S838N |
| 2786 | A | G | 6.33E-05 | | | | | ORF1ab | nonsynonymous_I841V |
| 2800 | T | A | 6.33E-05 | | | | | ORF1ab | synonymous_L845L |

| | | | | | | | | | |
|---|---|---|---|---|---|---|---|---|---|
| 2801 | G | A | 6.33E-05 | A | 6.33E-05 | | | ORF1ab | nonsynonymous_D846N; nonsynonymous_D846N |
| 29372 | G | A | 6.33E-05 | | | | | N | nonsynonymous_E367K |
| 29377 | T | A | 6.33E-05 | | | | | N | synonymous_P368P |
| 29391 | A | T | 6.33E-05 | T | 6.33E-05 | | | N | nonsynonymous_K373M; nonsynonymous_K373M |
| 29433 | A | G | 6.33E-05 | | | | | N | nonsynonymous_K387R |
| 29444 | A | T | 6.33E-05 | | | | | N | nonsynonymous_T391S |
| 29447 | G | T | 6.33E-05 | T | 6.33E-05 | | | N | nonsynonymous_V392L; nonsynonymous_V392L |
| 29470 | T | A | 6.33E-05 | A | 6.33E-05 | | | N | nonsynonymous_D399E; nonsynonymous_D399E |
| 29475 | A | T | 6.33E-05 | | | | | N | nonsynonymous_D401V |
| 29487 | A | T | 6.33E-05 | | | | | N | nonsynonymous_K405I |
| 29489 | C | T | 6.33E-05 | | | | | N | stopgain_Q406X |
| 29498 | C | G | 6.33E-05 | | | | | N | nonsynonymous_Q409E |
| 29502 | C | G | 6.33E-05 | | | | | N | nonsynonymous_S410C |
| 29508 | G | C | 6.33E-05 | | | | | N | nonsynonymous_S412T |
| 29588 | T | A | 6.33E-05 | | | | | ORF10 | nonsynonymous_F11I |
| 29632 | C | T | 6.33E-05 | | | | | ORF10 | synonymous_N25N |
| 29654 | G | C | 6.33E-05 | | | | | ORF10 | nonsynonymous_V33L |
| 20464 | G | C | 6.33E-05 | T | 6.33E-05 | | | ORF1ab | nonsynonymous_D6734H; nonsynonymous_D6734Y |
| 2745 | A | C | 6.33E-05 | T | 6.33E-05 | | | ORF1ab | nonsynonymous_D827A; nonsynonymous_D827V |
| 2817 | A | C | 6.33E-05 | G | 6.33E-05 | | | ORF1ab | nonsynonymous_K851T; nonsynonymous_K851R |
| 13608 | C | T | 6.33E-05 | | | | | ORF1ab | synonymous_F4448F |
| 13618 | G | T | 6.33E-05 | | | | | ORF1ab | nonsynonymous_D4452Y |
| 20249 | A | T | 6.33E-05 | | | | | ORF1ab | nonsynonymous_E6662V |

| | | | | | | | | | |
|---|---|---|---|---|---|---|---|---|---|
| 20367 | T | C | 6.33E-05 | | | | | ORF1ab | synonymous_H6701H |
| 20397 | G | A | 6.33E-05 | | | | | ORF1ab | synonymous_K6711K |
| 20404 | C | T | 6.33E-05 | | | | | ORF1ab | nonsynonymous_P6714S |
| 20405 | C | A | 6.33E-05 | A | 6.33E-05 | | | ORF1ab | nonsynonymous_P6714H; nonsynonymous_P6714H |
| 20430 | T | C | 6.33E-05 | | | | | ORF1ab | synonymous_P6722P |
| 20433 | G | T | 6.33E-05 | | | | | ORF1ab | nonsynonymous_M6723I |
| 20437 | A | G | 6.33E-05 | | | | | ORF1ab | nonsynonymous_S6725G |
| 20481 | A | G | 6.33E-05 | | | | | ORF1ab | synonymous_S6739S |
| 20489 | G | T | 6.33E-05 | | | | | ORF1ab | nonsynonymous_C6742F |
| 22369 | T | C | 6.33E-05 | | | | | S | synonymous_Y269Y |
| 2612 | G | C | 6.33E-05 | | | | | ORF1ab | nonsynonymous_V783L |
| 2635 | G | A | 6.33E-05 | | | | | ORF1ab | synonymous_L790L |
| 2647 | A | T | 6.33E-05 | | | | | ORF1ab | nonsynonymous_K794N |
| 2708 | C | T | 6.33E-05 | | | | | ORF1ab | nonsynonymous_L815F |
| 2749 | C | T | 6.33E-05 | | | | | ORF1ab | synonymous_D828D |
| 2761 | A | T | 6.33E-05 | | | | | ORF1ab | nonsynonymous_E832D |
| 2803 | T | A | 6.33E-05 | | | | | ORF1ab | nonsynonymous_D846E |
| 2805 | A | G | 6.33E-05 | | | | | ORF1ab | nonsynonymous_E847G |
| 2812 | T | C | 6.33E-05 | | | | | ORF1ab | synonymous_I849I |
| 2845 | T | C | 6.33E-05 | | | | | ORF1ab | synonymous_Y860Y |
| 29380 | A | G | 6.33E-05 | | | | | N | synonymous_K369K |
| 29383 | G | A | 6.33E-05 | A | 6.33E-05 | | | N | synonymous_K370K; synonymous_K370K |
| 29384 | G | T | 6.33E-05 | | | | | N | nonsynonymous_D371Y |
| 29390 | A | C | 6.33E-05 | | | | | N | nonsynonymous_K373Q |
| 29406 | A | C | 6.33E-05 | | | | | N | nonsynonymous_E378A |
| 29417 | T | C | 6.33E-05 | | | | | N | synonymous_L382L |
| 29418 | T | C | 6.33E-05 | | | | | N | nonsynonymous_L382S |
| 29419 | A | G | 6.33E-05 | | | | | N | synonymous_L382L |

| | | | | | | | | | |
|---|---|---|---|---|---|---|---|---|---|
| 29423 | C | A | 6.33E-05 | | | | | N | nonsynonymous_Q384K |
| 29430 | A | G | 6.33E-05 | | | | | N | nonsynonymous_Q386R |
| 29431 | G | T | 6.33E-05 | | | | | N | nonsynonymous_Q386H |
| 29432 | A | T | 6.33E-05 | | | | | N | stopgain_K387X |
| 29435 | A | T | 6.33E-05 | | | | | N | stopgain_K388X |
| 29438 | C | G | 6.33E-05 | | | | | N | nonsynonymous_Q389E |
| 29439 | A | T | 6.33E-05 | | | | | N | nonsynonymous_Q389L |
| 29452 | T | A | 6.33E-05 | | | | | N | synonymous_T393T |
| 29454 | T | G | 6.33E-05 | | | | | N | nonsynonymous_L394R |
| 29456 | C | A | 6.33E-05 | | | | | N | nonsynonymous_L395I |
| 29457 | T | A | 6.33E-05 | | | | | N | nonsynonymous_L395H |
| 29458 | T | C | 6.33E-05 | | | | | N | synonymous_L395L |
| 29460 | C | A | 6.33E-05 | | | | | N | nonsynonymous_P396H |
| 29464 | T | A | 6.33E-05 | | | | | N | synonymous_A397A |
| 29465 | G | T | 6.33E-05 | | | | | N | nonsynonymous_A398S |
| 29466 | C | T | 6.33E-05 | T | 6.33E-05 | | | N | nonsynonymous_A398V; nonsynonymous_A398V |
| 29468 | G | T | 6.33E-05 | | | | | N | nonsynonymous_D399Y |
| 29471 | T | C | 6.33E-05 | | | | | N | synonymous_L400L |
| 29483 | T | A | 6.33E-05 | | | | | N | nonsynonymous_S404T |
| 29486 | A | T | 6.33E-05 | | | | | N | stopgain_K405X |
| 29496 | A | G | 6.33E-05 | | | | | N | nonsynonymous_Q408R |
| 29499 | A | G | 6.33E-05 | | | | | N | nonsynonymous_Q409R |
| 29542 | A | T | 6.33E-05 | | | | | intergenic | |
| 29580 | C | T | 6.33E-05 | | | | | ORF10 | nonsynonymous_A8V |
| 29620 | G | T | 6.33E-05 | | | | | ORF10 | nonsynonymous_M21I |
| 29630 | A | T | 6.33E-05 | | | | | ORF10 | nonsynonymous_N25Y |
| 29647 | A | G | 6.33E-05 | | | | | ORF10 | synonymous_V30V |
| 29650 | T | C | 6.33E-05 | | | | | ORF10 | synonymous_D31D |
| 29659 | C | T | 6.33E-05 | | | | | ORF10 | synonymous_N34N |

| | | | | | | | | | |
|---|---|---|---|---|---|---|---|---|---|
| 13555 | A | T | 6.33E-05 | | | | | ORF1ab | nonsynonymous_N4431Y |
| 13599 | T | C | 6.33E-05 | | | | | ORF1ab | synonymous_C4445C |
| 13616 | A | G | 6.33E-05 | | | | | ORF1ab | nonsynonymous_K4451R |
| 20239 | A | T | 6.33E-05 | | | | | ORF1ab | nonsynonymous_S6659C |
| 20250 | A | T | 6.33E-05 | | | | | ORF1ab | nonsynonymous_E6662D |
| 20255 | A | G | 6.33E-05 | | | | | ORF1ab | nonsynonymous_D6664G |
| 20271 | T | C | 6.33E-05 | | | | | ORF1ab | synonymous_A6669A |
| 20295 | T | C | 6.33E-05 | | | | | ORF1ab | synonymous_Y6677Y |
| 20302 | G | A | 6.33E-05 | | | | | ORF1ab | nonsynonymous_E6680K |
| 20305 | G | T | 6.33E-05 | | | | | ORF1ab | nonsynonymous_G6681C |
| 20306 | G | C | 6.33E-05 | | | | | ORF1ab | nonsynonymous_G6681A |
| 20308 | T | C | 6.33E-05 | | | | | ORF1ab | nonsynonymous_Y6682H |
| 20310 | T | A | 6.33E-05 | | | | | ORF1ab | stopgain_Y6682X |
| 20312 | C | A | 6.33E-05 | | | | | ORF1ab | nonsynonymous_A6683D |
| 20313 | C | T | 6.33E-05 | | | | | ORF1ab | synonymous_A6683A |
| 20356 | G | T | 6.33E-05 | | | | | ORF1ab | nonsynonymous_G6698C |
| 20378 | G | T | 6.33E-05 | | | | | ORF1ab | nonsynonymous_G6705V |
| 21562 | A | G | 6.33E-05 | | | | | intergenic | |
| 22328 | T | C | 6.33E-05 | | | | | S | nonsynonymous_S256P |
| 22429 | A | G | 6.33E-05 | | | | | S | synonymous_V289V |
| 22524 | A | T | 6.33E-05 | | | | | S | nonsynonymous_Q321L |
| 22525 | A | G | 6.33E-05 | | | | | S | synonymous_Q321Q |
| 23122 | A | T | 6.33E-05 | | | | | S | synonymous_A520A |
| 29361 | T | C | 6.33E-05 | | | | | N | nonsynonymous_F363S |
| 29363 | C | T | 6.33E-05 | | | | | N | nonsynonymous_P364S |
| 29522 | A | T | 6.33E-05 | | | | | N | nonsynonymous_T417S |
| 29525 | C | T | 6.33E-05 | | | | | N | stopgain_Q418X |
| 29579 | G | A | 6.33E-05 | | | | | ORF10 | nonsynonymous_A8T |
| 29582 | T | C | 6.33E-05 | | | | | ORF10 | nonsynonymous_F9L |
| 29583 | T | A | 6.33E-05 | | | | | ORF10 | nonsynonymous_F9Y |

| | | | | | | | | | |
|---|---|---|---|---|---|---|---|---|---|
| 29584 | T | A | 6.33E-05 | | | | | ORF10 | nonsynonymous_F9L |
| 29589 | T | C | 6.33E-05 | | | | | ORF10 | nonsynonymous_F11S |
| 29596 | A | G | 6.33E-05 | | | | | ORF10 | nonsynonymous_I13M |
| 29599 | T | C | 6.33E-05 | | | | | ORF10 | synonymous_Y14Y |
| 29616 | G | A | 6.33E-05 | | | | | ORF10 | nonsynonymous_R20K |
| 29618 | A | T | 6.33E-05 | | | | | ORF10 | nonsynonymous_M21L |
| 29629 | T | C | 6.33E-05 | | | | | ORF10 | synonymous_R24R |
| 29633 | T | C | 6.33E-05 | | | | | ORF10 | nonsynonymous_Y26H |
| 29637 | T | G | 6.33E-05 | | | | | ORF10 | nonsynonymous_I27R |
| 29649 | A | G | 6.33E-05 | | | | | ORF10 | nonsynonymous_D31G |
| 29651 | G | A | 6.33E-05 | | | | | ORF10 | nonsynonymous_V32I |
| 29655 | T | C | 6.33E-05 | | | | | ORF10 | nonsynonymous_V33A |
| 29658 | A | G | 6.33E-05 | | | | | ORF10 | nonsynonymous_N34S |
| 29661 | T | A | 6.33E-05 | | | | | ORF10 | nonsynonymous_F35Y |
| 670 | T | C | 6.33E-05 | | | | | ORF1ab | synonymous_S135S |
| 8026 | A | T | 6.33E-05 | | | | | ORF1ab | synonymous_A2587A |
| 8651 | A | G | 6.33E-05 | | | | | ORF1ab | nonsynonymous_M2796V |
| 9039 | C | T | 6.33E-05 | | | | | ORF1ab | nonsynonymous_A2925V |
| 13413 | G | C | 6.33E-05 | | | | | ORF1ab | nonsynonymous_C4383S |
| 13492 | C | T | 6.33E-05 | | | | | ORF1ab | nonsynonymous_R4410C |
| 19574 | A | T | 6.33E-05 | | | | | ORF1ab | nonsynonymous_K6437I |
| 20251 | A | G | 6.33E-05 | | | | | ORF1ab | nonsynonymous_I6663V |
| 20256 | T | C | 6.33E-05 | | | | | ORF1ab | synonymous_D6664D |
| 20259 | C | A | 6.33E-05 | A | 6.33E-05 | | | ORF1ab | nonsynonymous_F6665L; nonsynonymous_F6665L |
| 20272 | A | G | 6.33E-05 | G | 6.33E-05 | | | ORF1ab | nonsynonymous_M6670V; nonsynonymous_M6670V |
| 20280 | A | T | 6.33E-05 | T | 6.33E-05 | | | ORF1ab | nonsynonymous_E6672D; nonsynonymous_E6672D |
| 20286 | T | C | 6.33E-05 | | | | | ORF1ab | synonymous_I6674I |

| | | | | | | | | | |
|---|---|---|---|---|---|---|---|---|---|
| 20288 | A | G | 6.33E-05 | | | | | ORF1ab | nonsynonymous_E6675G |
| 20321 | A | C | 6.33E-05 | | | | | ORF1ab | nonsynonymous_H6686P |
| 20325 | C | A | 6.33E-05 | A | 6.33E-05 | | | ORF1ab | synonymous_I6687I; synonymous_I6687I |
| 20327 | T | C | 6.33E-05 | | | | | ORF1ab | nonsynonymous_V6688A |
| 20330 | A | C | 6.33E-05 | | | | | ORF1ab | nonsynonymous_Y6689S |
| 20332 | G | A | 6.33E-05 | | | | | ORF1ab | nonsynonymous_G6690R |
| 20333 | G | A | 6.33E-05 | | | | | ORF1ab | nonsynonymous_G6690E |
| 20335 | G | A | 6.33E-05 | | | | | ORF1ab | nonsynonymous_D6691N |
| 20341 | A | T | 6.33E-05 | | | | | ORF1ab | nonsynonymous_S6693C |
| 20342 | G | A | 6.33E-05 | | | | | ORF1ab | nonsynonymous_S6693N |
| 20346 | T | A | 6.33E-05 | | | | | ORF1ab | nonsynonymous_H6694Q |
| 20350 | C | G | 6.33E-05 | | | | | ORF1ab | nonsynonymous_Q6696E |
| 20368 | C | T | 6.33E-05 | | | | | ORF1ab | synonymous_L6702L |
| 20369 | T | C | 6.33E-05 | | | | | ORF1ab | nonsynonymous_L6702P |
| 20374 | A | G | 6.33E-05 | | | | | ORF1ab | nonsynonymous_I6704V |
| 20375 | T | C | 6.33E-05 | | | | | ORF1ab | nonsynonymous_I6704T |
| 20379 | A | G | 6.33E-05 | | | | | ORF1ab | synonymous_G6705G |
| 20387 | A | G | 6.33E-05 | | | | | ORF1ab | nonsynonymous_K6708R |
| 20389 | C | A | 6.33E-05 | A | 6.33E-05 | | | ORF1ab | nonsynonymous_R6709S; nonsynonymous_R6709S |
| 20390 | G | A | 6.33E-05 | | | | | ORF1ab | nonsynonymous_R6709H |
| 20891 | T | G | 6.33E-05 | | | | | ORF1ab | nonsynonymous_V6876G |
| 21489 | A | C | 6.33E-05 | | | | | ORF1ab | nonsynonymous_K7075N |
| 21508 | G | A | 6.33E-05 | | | | | ORF1ab | nonsynonymous_E7082K |
| 21585 | T | G | 6.33E-05 | | | | | S | nonsynonymous_L8W |
| 22144 | T | C | 6.33E-05 | | | | | S | synonymous_F194F |
| 23119 | T | A | 6.33E-05 | | | | | S | nonsynonymous_H519Q |
| 25649 | T | G | 6.33E-05 | | | | | ORF3a | nonsynonymous_L86W |
| 25743 | T | C | 6.33E-05 | | | | | ORF3a | synonymous_S117S |

| | | | | | | | | | |
|---|---|---|---|---|---|---|---|---|---|
| 25770 | G | A | 6.33E-05 | A | 6.33E-05 | | | ORF3a | synonymous_R126R; synonymous_R126R |
| 25900 | A | G | 6.33E-05 | | | | | ORF3a | nonsynonymous_T170A |
| 29022 | C | T | 6.33E-05 | | | | | N | nonsynonymous_S250F |
| 29514 | C | T | 6.33E-05 | | | | | N | nonsynonymous_A414V |
| 29515 | T | A | 6.33E-05 | C | 6.33E-05 | | | N | synonymous_A414A; synonymous_A414A |
| 29516 | G | C | 6.33E-05 | | | | | N | nonsynonymous_D415H |
| 29519 | T | A | 6.33E-05 | | | | | N | nonsynonymous_S416T |
| 29520 | C | T | 6.33E-05 | T | 6.33E-05 | | | N | nonsynonymous_S416L; nonsynonymous_S416L |
| 29523 | C | T | 6.33E-05 | | | | | N | nonsynonymous_T417I |
| 29526 | A | G | 6.33E-05 | | | | | N | nonsynonymous_Q418R |
| 29529 | C | A | 6.33E-05 | | | | | N | nonsynonymous_A419D |
| 29530 | C | A | 6.33E-05 | A | 6.33E-05 | | | N | synonymous_A419A; synonymous_A419A |
| 29535 | C | T | 6.33E-05 | | | | | intergenic | |
| 29597 | T | A | 6.33E-05 | | | | | ORF10 | nonsynonymous_Y14N |
| 29604 | T | A | 6.33E-05 | | | | | ORF10 | nonsynonymous_L16Q |
| 29609 | T | C | 6.33E-05 | | | | | ORF10 | synonymous_L18L |
| 8658 | A | G | 6.33E-05 | | | | | ORF1ab | nonsynonymous_K2798R |
| 11071 | G | C | 6.32E-05 | | | | | ORF1ab | nonsynonymous_L3602F |
| 13379 | G | A | 6.32E-05 | | | | | ORF1ab | nonsynonymous_V4372I |
| 13410 | G | A | 6.32E-05 | A | 6.32E-05 | | | ORF1ab | nonsynonymous_S4382N; nonsynonymous_S4382N |
| 13415 | G | C | 6.32E-05 | | | | | ORF1ab | nonsynonymous_D4384H |
| 13419 | A | T | 6.32E-05 | | | | | ORF1ab | nonsynonymous_Q4385L |
| 13434 | T | C | 6.32E-05 | | | | | ORF1ab | nonsynonymous_M4390T |
| 13443 | C | T | 6.32E-05 | | | | | ORF1ab | nonsynonymous_S4393L |
| 13445 | G | T | 6.32E-05 | | | | | ORF1ab | nonsynonymous_A4394S |

| | | | | | | | | | |
|---|---|---|---|---|---|---|---|---|---|
| 13501 | C | T | 6.32E-05 | | | | | ORF1ab | nonsynonymous_P4413S |
| 13502 | C | T | 6.32E-05 | | | | | ORF1ab | nonsynonymous_P4413L |
| 13506 | C | T | 6.32E-05 | | | | | ORF1ab | synonymous_C4414C |
| 13510 | A | G | 6.32E-05 | | | | | ORF1ab | nonsynonymous_T4416A |
| 13511 | C | T | 6.32E-05 | | | | | ORF1ab | nonsynonymous_T4416I |
| 13520 | G | A | 6.32E-05 | | | | | ORF1ab | nonsynonymous_S4419N |
| 13521 | T | C | 6.32E-05 | | | | | ORF1ab | synonymous_S4419S |
| 13567 | G | A | 6.32E-05 | | | | | ORF1ab | nonsynonymous_A4435T |
| 13585 | C | A | 6.32E-05 | | | | | ORF1ab | nonsynonymous_L4441I |
| 13592 | C | T | 6.32E-05 | | | | | ORF1ab | nonsynonymous_T4443I |
| 14487 | A | G | 6.32E-05 | G | 6.32E-05 | | | ORF1ab | synonymous_R4741R; synonymous_R4741R |
| 14501 | T | C | 6.32E-05 | | | | | ORF1ab | nonsynonymous_V4746A |
| 18690 | T | A | 6.32E-05 | | | | | ORF1ab | nonsynonymous_F6142L |
| 19973 | C | T | 6.32E-05 | | | | | ORF1ab | nonsynonymous_P6570L |
| 20011 | G | T | 6.32E-05 | | | | | ORF1ab | nonsynonymous_V6583L |
| 20056 | G | A | 6.32E-05 | | | | | ORF1ab | nonsynonymous_G6598S |
| 20180 | A | T | 6.32E-05 | | | | | ORF1ab | nonsynonymous_Q6639L |
| 21606 | G | T | 6.32E-05 | | | | | S | nonsynonymous_C15F |
| 21632 | T | A | 6.32E-05 | A | 6.32E-05 | | | S | nonsynonymous_L24I; nonsynonymous_L24I |
| 22114 | T | C | 6.32E-05 | | | | | S | synonymous_G184G |
| 22962 | A | T | 6.32E-05 | | | | | S | nonsynonymous_D467V |
| 25652 | T | A | 6.32E-05 | A | 6.32E-05 | | | ORF3a | nonsynonymous_F87Y; nonsynonymous_F87Y |
| 25654 | G | T | 6.32E-05 | | | | | ORF3a | nonsynonymous_V88L |
| 25656 | A | G | 6.32E-05 | | | | | ORF3a | synonymous_V88V |
| 25667 | C | T | 6.32E-05 | | | | | ORF3a | nonsynonymous_S92L |
| 25670 | A | C | 6.32E-05 | | | | | ORF3a | nonsynonymous_H93P |
| 25671 | C | T | 6.32E-05 | | | | | ORF3a | synonymous_H93H |

| | | | | | | | | | |
|---|---|---|---|---|---|---|---|---|---|
| 25680 | C | T | 6.32E-05 | | | | | ORF3a | synonymous_L96L |
| 25681 | G | C | 6.32E-05 | | | | | ORF3a | nonsynonymous_V97L |
| 25685 | C | T | 6.32E-05 | | | | | ORF3a | nonsynonymous_A98V |
| 25693 | C | T | 6.32E-05 | | | | | ORF3a | nonsynonymous_L101F |
| 25711 | T | C | 6.32E-05 | | | | | ORF3a | nonsynonymous_Y107H |
| 25718 | A | G | 6.32E-05 | | | | | ORF3a | nonsynonymous_Y109C |
| 25725 | A | G | 6.32E-05 | | | | | ORF3a | synonymous_L111L |
| 25729 | T | C | 6.32E-05 | | | | | ORF3a | nonsynonymous_Y113H |
| 25732 | T | G | 6.32E-05 | | | | | ORF3a | nonsynonymous_F114V |
| 25733 | T | G | 6.32E-05 | | | | | ORF3a | nonsynonymous_F114C |
| 25740 | G | T | 6.32E-05 | | | | | ORF3a | nonsynonymous_Q116H |
| 25757 | G | A | 6.32E-05 | | | | | ORF3a | nonsynonymous_R122K |
| 25766 | T | C | 6.32E-05 | | | | | ORF3a | nonsynonymous_M125T |
| 25777 | C | T | 6.32E-05 | | | | | ORF3a | nonsynonymous_L129F |
| 25801 | A | C | 6.32E-05 | | | | | ORF3a | nonsynonymous_N137H |
| 25803 | C | T | 6.32E-05 | | | | | ORF3a | synonymous_N137N |
| 25806 | A | T | 6.32E-05 | | | | | ORF3a | synonymous_P138P |
| 25815 | T | C | 6.32E-05 | | | | | ORF3a | synonymous_Y141Y |
| 25819 | G | T | 6.32E-05 | | | | | ORF3a | nonsynonymous_A143S |
| 25821 | C | T | 6.32E-05 | | | | | ORF3a | synonymous_A143A |
| 25824 | C | T | 6.32E-05 | | | | | ORF3a | synonymous_N144N |
| 25835 | G | T | 6.32E-05 | | | | | ORF3a | nonsynonymous_C148F |
| 25838 | G | T | 6.32E-05 | | | | | ORF3a | nonsynonymous_W149L |
| 25847 | A | G | 6.32E-05 | | | | | ORF3a | nonsynonymous_N152S |
| 25848 | T | C | 6.32E-05 | | | | | ORF3a | synonymous_N152N |
| 25850 | G | A | 6.32E-05 | | | | | ORF3a | nonsynonymous_C153Y |
| 25853 | A | G | 6.32E-05 | | | | | ORF3a | nonsynonymous_Y154C |
| 25858 | T | A | 6.32E-05 | | | | | ORF3a | nonsynonymous_Y156N |
| 25907 | G | T | 6.32E-05 | | | | | ORF3a | nonsynonymous_G172V |
| 27151 | A | T | 6.32E-05 | | | | | M | nonsynonymous_H210L |

| | | | | | | | | | |
|---|---|---|---|---|---|---|---|---|---|
| 284 | G | T | 6.32E-05 | | | | | ORF1ab | nonsynonymous_G7C |
| 28903 | G | T | 6.32E-05 | T | 6.32E-05 | | | N | nonsynonymous_M210I; nonsynonymous_M210I |
| 28916 | G | A | 6.32E-05 | | | | | N | nonsynonymous_G215S |
| 292 | C | T | 6.32E-05 | | | | | ORF1ab | synonymous_N9N |
| 5339 | C | T | 6.32E-05 | | | | | ORF1ab | nonsynonymous_P1692S |
| 539 | C | T | 6.32E-05 | | | | | ORF1ab | nonsynonymous_L92F |
| 13687 | G | T | 6.32E-05 | | | | | ORF1ab | stopgain_E4475X |
| 19938 | C | T | 6.32E-05 | | | | | ORF1ab | synonymous_D6558D |
| 19939 | A | C | 6.32E-05 | | | | | ORF1ab | nonsynonymous_I6559L |
| 19945 | A | G | 6.32E-05 | | | | | ORF1ab | nonsynonymous_K6561E |
| 19947 | G | T | 6.32E-05 | | | | | ORF1ab | nonsynonymous_K6561N |
| 19952 | C | T | 6.32E-05 | | | | | ORF1ab | nonsynonymous_P6563L |
| 19958 | A | T | 6.32E-05 | | | | | ORF1ab | nonsynonymous_E6565V |
| 19959 | A | C | 6.32E-05 | | | | | ORF1ab | nonsynonymous_E6565D |
| 19969 | G | A | 6.32E-05 | | | | | ORF1ab | nonsynonymous_A6569T |
| 19970 | C | T | 6.32E-05 | | | | | ORF1ab | nonsynonymous_A6569V |
| 19972 | C | T | 6.32E-05 | | | | | ORF1ab | nonsynonymous_P6570S |
| 19974 | A | G | 6.32E-05 | | | | | ORF1ab | synonymous_P6570P |
| 20059 | A | T | 6.32E-05 | | | | | ORF1ab | nonsynonymous_S6599C |
| 20167 | G | T | 6.32E-05 | | | | | ORF1ab | nonsynonymous_D6635Y |
| 21428 | C | T | 6.32E-05 | | | | | ORF1ab | nonsynonymous_T7055I |
| 21431 | C | T | 6.32E-05 | | | | | ORF1ab | nonsynonymous_A7056V |
| 21440 | C | T | 6.32E-05 | | | | | ORF1ab | nonsynonymous_S7059F |
| 21446 | A | G | 6.32E-05 | | | | | ORF1ab | nonsynonymous_K7061R |
| 21459 | C | T | 6.32E-05 | | | | | ORF1ab | synonymous_I7065I |
| 21481 | C | T | 6.32E-05 | | | | | ORF1ab | nonsynonymous_L7073F |
| 21485 | G | T | 6.32E-05 | | | | | ORF1ab | nonsynonymous_S7074I |
| 21511 | A | G | 6.32E-05 | | | | | ORF1ab | nonsynonymous_N7083D |
| 21516 | C | T | 6.32E-05 | | | | | ORF1ab | synonymous_N7084N |

| | | | | | | | | | |
|---|---|---|---|---|---|---|---|---|---|
| 21536 | A | T | 6.32E-05 | | | | | ORF1ab | nonsynonymous_D7091V |
| 21537 | T | A | 6.32E-05 | A | 6.32E-05 | | | ORF1ab | nonsynonymous_D7091E; nonsynonymous_D7091E |
| 21538 | G | T | 6.32E-05 | | | | | ORF1ab | nonsynonymous_V7092F |
| 21539 | T | A | 6.32E-05 | | | | | ORF1ab | nonsynonymous_V7092D |
| 21556 | A | G | 6.32E-05 | | | | | intergenic | |
| 21568 | T | A | 6.32E-05 | | | | | S | nonsynonymous_F2L |
| 21578 | G | A | 6.32E-05 | | | | | S | nonsynonymous_V6I |
| 21587 | C | T | 6.32E-05 | | | | | S | nonsynonymous_P9S |
| 21616 | T | C | 6.32E-05 | | | | | S | synonymous_L18L |
| 21629 | C | A | 6.32E-05 | A | 6.32E-05 | | | S | nonsynonymous_Q23K; nonsynonymous_Q23K |
| 21636 | C | T | 6.32E-05 | | | | | S | nonsynonymous_P25L |
| 21651 | A | G | 6.32E-05 | | | | | S | nonsynonymous_N30S |
| 21656 | T | A | 6.32E-05 | | | | | S | nonsynonymous_F32I |
| 21770 | G | T | 6.32E-05 | | | | | S | nonsynonymous_V70F |
| 21775 | T | C | 6.32E-05 | | | | | S | synonymous_S71S |
| 21776 | G | A | 6.32E-05 | | | | | S | nonsynonymous_G72R |
| 21986 | G | A | 6.32E-05 | | | | | S | nonsynonymous_G142S |
| 22115 | A | G | 6.32E-05 | | | | | S | nonsynonymous_N185D |
| 22139 | G | T | 6.32E-05 | | | | | S | nonsynonymous_V193L |
| 23030 | T | C | 6.32E-05 | | | | | S | nonsynonymous_F490L |
| 23042 | T | C | 6.32E-05 | | | | | S | nonsynonymous_S494P |
| 23069 | G | T | 6.32E-05 | | | | | S | nonsynonymous_V503F |
| 23083 | A | G | 6.32E-05 | | | | | S | synonymous_P507P |
| 23088 | G | A | 6.32E-05 | | | | | S | nonsynonymous_R509K |
| 24056 | G | A | 6.32E-05 | A | 6.32E-05 | | | S | nonsynonymous_G832S; nonsynonymous_G832S |
| 25061 | G | A | 6.32E-05 | | | | | S | nonsynonymous_G1167S |
| 25105 | A | G | 6.32E-05 | | | | | S | synonymous_K1181K |

| | | | | | | | | | |
|---|---|---|---|---|---|---|---|---|---|
| 25198 | G | A | 6.32E-05 | | | | | S | stopgain_W1212X |
| 25245 | T | G | 6.32E-05 | | | | | S | nonsynonymous_V1228G |
| 25621 | G | T | 6.32E-05 | | | | | ORF3a | nonsynonymous_V77F |
| 25885 | T | G | 6.32E-05 | | | | | ORF3a | nonsynonymous_S165A |
| 25899 | T | C | 6.32E-05 | | | | | ORF3a | synonymous_I169I |
| 25905 | A | T | 6.32E-05 | | | | | ORF3a | synonymous_S171S |
| 25909 | G | A | 6.32E-05 | A | 6.32E-05 | | | ORF3a | nonsynonymous_D173N; nonsynonymous_D173N |
| 270 | A | G | 6.32E-05 | | | | | ORF1ab | nonsynonymous_E2G |
| 28676 | A | C | 6.32E-05 | | | | | N | nonsynonymous_T135P |
| 28894 | T | C | 6.32E-05 | | | | | N | synonymous_P207P |
| 28904 | G | T | 6.32E-05 | | | | | N | nonsynonymous_A211S |
| 29052 | A | C | 6.32E-05 | | | | | N | nonsynonymous_Q260P |
| 29070 | A | G | 6.32E-05 | | | | | N | nonsynonymous_K266R |
| 29072 | G | C | 6.32E-05 | | | | | N | nonsynonymous_A267P |
| 29075 | T | C | 6.32E-05 | | | | | N | nonsynonymous_Y268H |
| 29080 | T | G | 6.32E-05 | | | | | N | nonsynonymous_N269K |
| 29081 | G | T | 6.32E-05 | | | | | N | nonsynonymous_V270L |
| 29115 | A | T | 6.32E-05 | T | 6.32E-05 | | | N | nonsynonymous_Q281L; nonsynonymous_Q281L |
| 29116 | A | G | 6.32E-05 | | | | | N | synonymous_Q281Q |
| 29117 | A | T | 6.32E-05 | | | | | N | nonsynonymous_T282S |
| 29134 | G | T | 6.32E-05 | | | | | N | synonymous_G287G |
| 29138 | C | T | 6.32E-05 | | | | | N | stopgain_Q289X |
| 29197 | C | T | 6.32E-05 | | | | | N | synonymous_A308A |
| 29219 | G | A | 6.32E-05 | | | | | N | nonsynonymous_G316R |
| 29290 | G | A | 6.32E-05 | A | 6.32E-05 | | | N | synonymous_L339L; synonymous_L339L |
| 29314 | A | T | 6.32E-05 | | | | | N | nonsynonymous_K347N |
| 5321 | A | G | 6.32E-05 | | | | | ORF1ab | nonsynonymous_I1686V |

| | | | | | | | | | |
|---|---|---|---|---|---|---|---|---|---|
| 5332 | G | T | 6.32E-05 | | | | | ORF1ab | nonsynonymous_K1689N |
| 8905 | C | T | 6.32E-05 | | | | | ORF1ab | synonymous_D2880D |
| 11271 | T | C | 6.32E-05 | | | | | ORF1ab | nonsynonymous_M3669T |
| 20033 | G | C | 6.32E-05 | | | | | ORF1ab | nonsynonymous_R6590P |
| 20157 | T | C | 6.32E-05 | | | | | ORF1ab | synonymous_Y6631Y |
| 20529 | T | C | 6.32E-05 | | | | | ORF1ab | synonymous_V6755V |
| 21994 | T | C | 6.32E-05 | | | | | S | synonymous_Y144Y |
| 22939 | T | C | 6.32E-05 | | | | | S | synonymous_S459S |
| 22965 | T | C | 6.32E-05 | | | | | S | nonsynonymous_I468T |
| 22973 | G | C | 6.32E-05 | | | | | S | nonsynonymous_E471Q |
| 22981 | T | C | 6.32E-05 | | | | | S | synonymous_Y473Y |
| 23031 | T | C | 6.32E-05 | | | | | S | nonsynonymous_F490S |
| 23036 | T | C | 6.32E-05 | | | | | S | synonymous_L492L |
| 23084 | T | C | 6.32E-05 | | | | | S | nonsynonymous_Y508H |
| 25102 | A | C | 6.32E-05 | | | | | S | nonsynonymous_Q1180H |
| 25228 | T | C | 6.32E-05 | | | | | S | synonymous_A1222A |
| 25645 | T | C | 6.32E-05 | | | | | ORF3a | synonymous_L85L |
| 26409 | T | C | 6.32E-05 | | | | | E | synonymous_S55S |
| 27210 | T | C | 6.32E-05 | | | | | ORF6 | synonymous_H3H |
| 27267 | T | C | 6.32E-05 | | | | | ORF6 | synonymous_F22F |
| 27482 | T | C | 6.32E-05 | | | | | ORF7a | nonsynonymous_L30P |
| 28889 | T | C | 6.32E-05 | | | | | N | nonsynonymous_S206P |
| 29047 | T | C | 6.32E-05 | | | | | N | synonymous_P258P |
| 29059 | T | C | 6.32E-05 | | | | | N | synonymous_R262R |
| 29082 | T | C | 6.32E-05 | | | | | N | nonsynonymous_V270A |
| 29097 | G | C | 6.32E-05 | | | | | N | nonsynonymous_G275A |
| 29111 | G | C | 6.32E-05 | | | | | N | nonsynonymous_E280Q |
| 29113 | A | C | 6.32E-05 | | | | | N | nonsynonymous_E280D |
| 29122 | A | C | 6.32E-05 | | | | | N | nonsynonymous_Q283H |
| 29128 | T | C | 6.32E-05 | | | | | N | synonymous_N285N |

| | | | | | | | | | |
|---|---|---|---|---|---|---|---|---|---|
| 29130 | T | C | 6.32E-05 | | | | | N | nonsynonymous_F286S |
| 29132 | G | C | 6.32E-05 | | | | | N | nonsynonymous_G287R |
| 29146 | A | C | 6.32E-05 | | | | | N | synonymous_L291L |
| 29151 | G | C | 6.32E-05 | | | | | N | nonsynonymous_R293T |
| 29161 | T | C | 6.32E-05 | | | | | N | synonymous_T296T |
| 29182 | A | C | 6.32E-05 | | | | | N | nonsynonymous_Q303H |
| 29193 | T | C | 6.32E-05 | | | | | N | nonsynonymous_F307S |
| 29225 | T | C | 6.32E-05 | | | | | N | nonsynonymous_S318P |
| 29261 | T | C | 6.32E-05 | | | | | N | nonsynonymous_W330R |
| 29287 | A | C | 6.32E-05 | | | | | N | nonsynonymous_K338N |
| 5104 | T | C | 6.32E-05 | | | | | ORF1ab | synonymous_H1613H |
| 5158 | T | C | 6.32E-05 | | | | | ORF1ab | synonymous_A1631A |
| 5464 | T | C | 6.32E-05 | | | | | ORF1ab | synonymous_S1733S |
| 15866 | C | G | 6.32E-05 | | | | | ORF1ab | nonsynonymous_P5201R |
| 17822 | C | T | 6.32E-05 | | | | | ORF1ab | nonsynonymous_P5853L |
| 18693 | C | G | 6.32E-05 | | | | | ORF1ab | synonymous_S6143S |
| 19571 | A | T | 6.32E-05 | | | | | ORF1ab | nonsynonymous_Y6436F |
| 20077 | C | A | 6.32E-05 | | | | | ORF1ab | nonsynonymous_P6605T |
| 20081 | C | T | 6.32E-05 | | | | | ORF1ab | nonsynonymous_S6606F |
| 20137 | A | G | 6.32E-05 | | | | | ORF1ab | nonsynonymous_K6625E |
| 20142 | A | G | 6.32E-05 | | | | | ORF1ab | synonymous_T6626T |
| 20173 | G | T | 6.32E-05 | | | | | ORF1ab | nonsynonymous_V6637F |
| 20196 | T | A | 6.32E-05 | | | | | ORF1ab | synonymous_T6644T |
| 21786 | G | A | 6.32E-05 | A | 6.32E-05 | | | S | nonsynonymous_G75D; nonsynonymous_G75D |
| 21980 | T | G | 6.32E-05 | | | | | S | nonsynonymous_F140V |
| 22016 | T | A | 6.32E-05 | | | | | S | nonsynonymous_W152R |
| 22218 | G | A | 6.32E-05 | | | | | S | nonsynonymous_G219D |
| 22286 | C | T | 6.32E-05 | | | | | S | nonsynonymous_L242F |
| 22882 | T | G | 6.32E-05 | | | | | S | nonsynonymous_N440K |

| | | | | | | | | | |
|---|---|---|---|---|---|---|---|---|---|
| 22893 | A | G | 6.32E-05 | | | | | S | nonsynonymous_K444R |
| 22917 | T | G | 6.32E-05 | | | | | S | nonsynonymous_L452R |
| 22936 | G | T | 6.32E-05 | | | | | S | nonsynonymous_K458N |
| 22945 | C | T | 6.32E-05 | | | | | S | synonymous_L461L |
| 22976 | A | G | 6.32E-05 | | | | | S | nonsynonymous_I472V |
| 22987 | C | A | 6.32E-05 | A | 6.32E-05 | A | 6.32E-05 | S | synonymous_A475A; synonymous_A475A; synonymous_A475A |
| 22993 | C | A | 6.32E-05 | A | 6.32E-05 | | | S | nonsynonymous_S477R; nonsynonymous_S477R |
| 23015 | G | A | 6.32E-05 | | | | | S | nonsynonymous_G485S |
| 23034 | C | G | 6.32E-05 | | | | | S | nonsynonymous_P491R |
| 23059 | C | T | 6.32E-05 | | | | | S | synonymous_P499P |
| 23090 | G | T | 6.32E-05 | | | | | S | nonsynonymous_V510L |
| 23130 | C | A | 6.32E-05 | | | | | S | nonsynonymous_T523N |
| 23595 | C | T | 6.32E-05 | | | | | S | nonsynonymous_T678I |
| 23597 | A | T | 6.32E-05 | | | | | S | nonsynonymous_N679Y |
| 24914 | G | T | 6.32E-05 | | | | | S | nonsynonymous_D1118Y |
| 24998 | G | A | 6.32E-05 | | | | | S | nonsynonymous_D1146N |
| 25064 | G | A | 6.32E-05 | C | 6.32E-05 | | | S | nonsynonymous_D1168N; nonsynonymous_D1168H |
| 25186 | G | T | 6.32E-05 | C | 6.32E-05 | | | S | nonsynonymous_Q1208H; nonsynonymous_Q1208H |
| 25197 | G | T | 6.32E-05 | | | | | S | nonsynonymous_W1212L |
| 25230 | G | T | 6.32E-05 | | | | | S | nonsynonymous_G1223V |
| 25231 | C | T | 6.32E-05 | T | 6.32E-05 | | | S | synonymous_G1223G; synonymous_G1223G |
| 25232 | T | A | 6.32E-05 | | | | | S | nonsynonymous_L1224M |
| 25239 | C | T | 6.32E-05 | | | | | S | nonsynonymous_A1226V |
| 25267 | C | T | 6.32E-05 | | | | | S | synonymous_C1235C |

| | | | | | | | | | |
|---|---|---|---|---|---|---|---|---|---|
| 25618 | G | A | 6.32E-05 | | | | | ORF3a | nonsynonymous_G76S |
| 25625 | A | G | 6.32E-05 | | | | | ORF3a | nonsynonymous_H78R |
| 25626 | C | T | 6.32E-05 | | | | | ORF3a | synonymous_H78H |
| 25630 | G | A | 6.32E-05 | A | 6.32E-05 | C | 6.32E-05 | ORF3a | nonsynonymous_V80I; nonsynonymous_V80I; nonsynonymous_V80L |
| 25631 | T | A | 6.32E-05 | | | | | ORF3a | nonsynonymous_V80D |
| 25636 | A | G | 6.32E-05 | | | | | ORF3a | nonsynonymous_N82D |
| 25642 | C | T | 6.32E-05 | | | | | ORF3a | synonymous_L84L |
| 26396 | T | A | 6.32E-05 | | | | | E | nonsynonymous_L51H |
| 26428 | G | T | 6.32E-05 | | | | | E | nonsynonymous_V62F |
| 27474 | A | G | 6.32E-05 | | | | | ORF7a | synonymous_T27T |
| 27490 | G | T | 6.32E-05 | | | | | ORF7a | stopgain_E33X |
| 27505 | G | A | 6.32E-05 | | | | | ORF7a | nonsynonymous_G38R |
| 28232 | C | T | 6.32E-05 | | | | | ORF8 | synonymous_D113D |
| 28814 | C | G | 6.32E-05 | | | | | N | nonsynonymous_Q181E |
| 28874 | A | G | 6.32E-05 | | | | | N | nonsynonymous_S201G |
| 28885 | A | G | 6.32E-05 | | | | | N | synonymous_G204G |
| 28888 | T | A | 6.32E-05 | | | | | N | synonymous_T205T |
| 28907 | G | T | 6.32E-05 | | | | | N | nonsynonymous_G212C |
| 28990 | A | G | 6.32E-05 | | | | | N | synonymous_Q239Q |
| 29046 | C | T | 6.32E-05 | | | | | N | nonsynonymous_P258L |
| 29048 | C | A | 6.32E-05 | | | | | N | synonymous_R259R |
| 29053 | A | T | 6.32E-05 | | | | | N | nonsynonymous_Q260H |
| 29058 | G | A | 6.32E-05 | A | 6.32E-05 | | | N | nonsynonymous_R262H; nonsynonymous_R262H |
| 29060 | A | G | 6.32E-05 | | | | | N | nonsynonymous_T263A |
| 29084 | A | G | 6.32E-05 | | | | | N | nonsynonymous_T271A |
| 29087 | C | A | 6.32E-05 | | | | | N | nonsynonymous_Q272K |

| | | | | | | | | | |
|---|---|---|---|---|---|---|---|---|---|
| 29088 | A | T | 6.32E-05 | T | 6.32E-05 | | | N | nonsynonymous_Q272L; nonsynonymous_Q272L |
| 29089 | A | G | 6.32E-05 | G | 6.32E-05 | | | N | synonymous_Q272Q; synonymous_Q272Q |
| 29090 | G | T | 6.32E-05 | | | | | N | nonsynonymous_A273S |
| 29091 | C | T | 6.32E-05 | | | | | N | nonsynonymous_A273V |
| 29098 | C | T | 6.32E-05 | | | | | N | synonymous_G275G |
| 290 | A | T | 6.32E-05 | | | | | ORF1ab | nonsynonymous_N9Y |
| 29100 | G | T | 6.32E-05 | | | | | N | nonsynonymous_R276I |
| 29102 | C | T | 6.32E-05 | | | | | N | nonsynonymous_R277C |
| 29103 | G | A | 6.32E-05 | | | | | N | nonsynonymous_R277H |
| 29105 | G | A | 6.32E-05 | | | | | N | nonsynonymous_G278S |
| 29108 | C | T | 6.32E-05 | | | | | N | nonsynonymous_P279S |
| 29112 | A | T | 6.32E-05 | | | | | N | nonsynonymous_E280V |
| 29114 | C | T | 6.32E-05 | | | | | N | stopgain_Q281X |
| 29118 | C | T | 6.32E-05 | T | 6.32E-05 | | | N | nonsynonymous_T282I; nonsynonymous_T282I |
| 29120 | C | A | 6.32E-05 | | | | | N | nonsynonymous_Q283K |
| 29121 | A | G | 6.32E-05 | | | | | N | nonsynonymous_Q283R |
| 29123 | G | A | 6.32E-05 | | | | | N | nonsynonymous_G284R |
| 29124 | G | T | 6.32E-05 | | | | | N | nonsynonymous_G284V |
| 29125 | A | G | 6.32E-05 | | | | | N | synonymous_G284G |
| 29135 | G | A | 6.32E-05 | | | | | N | nonsynonymous_D288N |
| 29150 | A | G | 6.32E-05 | | | | | N | nonsynonymous_R293G |
| 29154 | A | T | 6.32E-05 | | | | | N | nonsynonymous_Q294L |
| 29157 | G | T | 6.32E-05 | | | | | N | nonsynonymous_G295V |
| 29159 | A | G | 6.32E-05 | | | | | N | nonsynonymous_T296A |
| 29160 | C | A | 6.32E-05 | A | 6.32E-05 | | | N | nonsynonymous_T296N; nonsynonymous_T296N |
| 29165 | T | A | 6.32E-05 | | | | | N | nonsynonymous_Y298N |

| | | | | | | | | | |
|---|---|---|---|---|---|---|---|---|---|
| 29166 | A | G | 6.32E-05 | | | | | N | nonsynonymous_Y298C |
| 29168 | A | T | 6.32E-05 | | | | | N | stopgain_K299X |
| 29172 | A | T | 6.32E-05 | | | | | N | nonsynonymous_H300L |
| 29175 | G | A | 6.32E-05 | | | | | N | stopgain_W301X |
| 29176 | G | A | 6.32E-05 | | | | | N | stopgain_W301X |
| 29185 | T | A | 6.32E-05 | | | | | N | synonymous_I304I |
| 29196 | C | T | 6.32E-05 | | | | | N | nonsynonymous_A308V |
| 29198 | C | A | 6.32E-05 | | | | | N | nonsynonymous_P309T |
| 29202 | G | T | 6.32E-05 | | | | | N | nonsynonymous_S310I |
| 29206 | T | G | 6.32E-05 | G | 6.32E-05 | | | N | synonymous_A311A; synonymous_A311A |
| 29207 | T | G | 6.32E-05 | | | | | N | nonsynonymous_S312A |
| 29208 | C | T | 6.32E-05 | | | | | N | nonsynonymous_S312L |
| 29210 | G | A | 6.32E-05 | | | | | N | nonsynonymous_A313T |
| 29220 | G | A | 6.32E-05 | | | | | N | nonsynonymous_G316E |
| 29222 | A | G | 6.32E-05 | | | | | N | nonsynonymous_M317V |
| 29223 | T | A | 6.32E-05 | | | | | N | nonsynonymous_M317K |
| 29230 | C | T | 6.32E-05 | | | | | N | synonymous_R319R |
| 29231 | A | G | 6.32E-05 | | | | | N | nonsynonymous_I320V |
| 29234 | G | T | 6.32E-05 | C | 6.32E-05 | | | N | nonsynonymous_G321C; nonsynonymous_G321R |
| 29235 | G | T | 6.32E-05 | | | | | N | nonsynonymous_G321V |
| 29242 | A | G | 6.32E-05 | | | | | N | synonymous_E323E |
| 29248 | A | G | 6.32E-05 | | | | | N | synonymous_T325T |
| 29258 | A | G | 6.32E-05 | | | | | N | nonsynonymous_T329A |
| 29263 | G | A | 6.32E-05 | | | | | N | stopgain_W330X |
| 29268 | C | A | 6.32E-05 | A | 6.32E-05 | | | N | nonsynonymous_T332N; nonsynonymous_T332N |
| 29270 | T | A | 6.32E-05 | | | | | N | nonsynonymous_Y333N |
| 29273 | A | G | 6.32E-05 | | | | | N | nonsynonymous_T334A |

| | | | | | | | | | |
|---|---|---|---|---|---|---|---|---|---|
| 29275 | A | T | 6.32E-05 | T | 6.32E-05 | | | N | synonymous_T334T; synonymous_T334T |
| 29276 | G | A | 6.32E-05 | | | | | N | nonsynonymous_G335S |
| 29278 | T | G | 6.32E-05 | | | | | N | synonymous_G335G |
| 29280 | C | A | 6.32E-05 | | | | | N | nonsynonymous_A336D |
| 29282 | A | T | 6.32E-05 | T | 6.32E-05 | | | N | nonsynonymous_I337F; nonsynonymous_I337F |
| 29283 | T | G | 6.32E-05 | | | | | N | nonsynonymous_I337S |
| 29285 | A | T | 6.32E-05 | | | | | N | stopgain_K338X |
| 29286 | A | G | 6.32E-05 | | | | | N | nonsynonymous_K338R |
| 29292 | A | G | 6.32E-05 | | | | | N | nonsynonymous_D340G |
| 29295 | A | G | 6.32E-05 | | | | | N | nonsynonymous_D341G |
| 29298 | A | G | 6.32E-05 | | | | | N | nonsynonymous_K342R |
| 29301 | A | T | 6.32E-05 | T | 6.32E-05 | | | N | nonsynonymous_D343V; nonsynonymous_D343V |
| 29304 | C | T | 6.32E-05 | | | | | N | nonsynonymous_P344L |
| 29307 | A | T | 6.32E-05 | | | | | N | nonsynonymous_N345I |
| 29308 | T | G | 6.32E-05 | G | 6.32E-05 | | | N | nonsynonymous_N345K; nonsynonymous_N345K |
| 29310 | T | G | 6.32E-05 | | | | | N | nonsynonymous_F346C |
| 29313 | A | G | 6.32E-05 | | | | | N | nonsynonymous_K347R |
| 4175 | A | G | 6.32E-05 | | | | | ORF1ab | nonsynonymous_K1304E |
| 5079 | C | T | 6.32E-05 | | | | | ORF1ab | nonsynonymous_T1605I |
| 5175 | C | T | 6.32E-05 | | | | | ORF1ab | nonsynonymous_T1637I |
| 5337 | A | G | 6.32E-05 | | | | | ORF1ab | nonsynonymous_N1691S |
| 5353 | A | G | 6.32E-05 | | | | | ORF1ab | synonymous_Q1696Q |
| 5399 | G | T | 6.32E-05 | | | | | ORF1ab | nonsynonymous_A1712S |
| 5421 | A | G | 6.32E-05 | | | | | ORF1ab | nonsynonymous_N1719S |
| 5462 | A | G | 6.32E-05 | | | | | ORF1ab | nonsynonymous_S1733G |
| 5526 | C | T | 6.32E-05 | | | | | ORF1ab | nonsynonymous_T1754I |

| | | | | | | | | | |
|---|---|---|---|---|---|---|---|---|---|
| 5555 | G | A | 6.32E-05 | | | | | ORF1ab | nonsynonymous_G1764S |
| 5580 | G | T | 6.32E-05 | | | | | ORF1ab | nonsynonymous_G1772V |
| 7017 | G | A | 6.32E-05 | A | 6.32E-05 | | | ORF1ab | nonsynonymous_G2251D; nonsynonymous_G2251D |
| 9370 | T | G | 6.32E-05 | | | | | ORF1ab | nonsynonymous_I3035M |
| 24364 | T | C | 6.32E-05 | A | 6.32E-05 | A | 6.32E-05 | S | synonymous_I934I; synonymous_I934I; synonymous_I934I |
| 25068 | T | C | 6.32E-05 | G | 6.32E-05 | | | S | nonsynonymous_I1169T; nonsynonymous_I1169S |
| 25191 | T | C | 6.32E-05 | A | 6.32E-05 | | | S | nonsynonymous_I1210T; nonsynonymous_I1210K |
| 10129 | T | C | 6.32E-05 | | | | | ORF1ab | synonymous_T3288T |
| 11270 | A | G | 6.32E-05 | | | | | ORF1ab | nonsynonymous_M3669V |
| 13914 | T | C | 6.32E-05 | | | | | ORF1ab | synonymous_N4550N |
| 14335 | G | T | 6.32E-05 | | | | | ORF1ab | nonsynonymous_V4691F |
| 14707 | G | T | 6.32E-05 | | | | | ORF1ab | nonsynonymous_A4815S |
| 14913 | C | T | 6.32E-05 | | | | | ORF1ab | synonymous_N4883N |
| 15393 | T | G | 6.32E-05 | | | | | ORF1ab | synonymous_R5043R |
| 15864 | A | G | 6.32E-05 | | | | | ORF1ab | synonymous_G5200G |
| 16068 | G | T | 6.32E-05 | | | | | ORF1ab | nonsynonymous_E5268D |
| 16305 | A | G | 6.32E-05 | | | | | ORF1ab | synonymous_P5347P |
| 16528 | G | T | 6.32E-05 | | | | | ORF1ab | nonsynonymous_V5422F |
| 16553 | T | A | 6.32E-05 | | | | | ORF1ab | nonsynonymous_F5430Y |
| 16655 | C | T | 6.32E-05 | | | | | ORF1ab | nonsynonymous_A5464V |
| 16699 | C | T | 6.32E-05 | | | | | ORF1ab | nonsynonymous_R5479C |
| 17536 | C | T | 6.32E-05 | | | | | ORF1ab | nonsynonymous_P5758S |
| 17568 | T | C | 6.32E-05 | | | | | ORF1ab | synonymous_C5768C |
| 17572 | G | T | 6.32E-05 | | | | | ORF1ab | nonsynonymous_A5770S |
| 18040 | G | T | 6.32E-05 | | | | | ORF1ab | nonsynonymous_A5926S |

| | | | | | | | | | |
|---|---|---|---|---|---|---|---|---|---|
| 18105 | G | T | 6.32E-05 | | | | | ORF1ab | nonsynonymous_Q5947H |
| 18214 | G | T | 6.32E-05 | | | | | ORF1ab | nonsynonymous_G5984C |
| 18680 | C | A | 6.32E-05 | A | 6.32E-05 | | | ORF1ab | nonsynonymous_A6139D; nonsynonymous_A6139D |
| 18700 | T | G | 6.32E-05 | | | | | ORF1ab | nonsynonymous_S6146A |
| 18912 | C | T | 6.32E-05 | | | | | ORF1ab | synonymous_D6216D |
| 19152 | C | T | 6.32E-05 | | | | | ORF1ab | synonymous_A6296A |
| 19846 | G | A | 6.32E-05 | | | | | ORF1ab | nonsynonymous_G6528S |
| 20527 | G | T | 6.32E-05 | | | | | ORF1ab | nonsynonymous_V6755F |
| 20541 | A | G | 6.32E-05 | | | | | ORF1ab | synonymous_K6759K |
| 20544 | C | T | 6.32E-05 | | | | | ORF1ab | synonymous_S6760S |
| 20548 | G | T | 6.32E-05 | | | | | ORF1ab | nonsynonymous_D6762Y |
| 20568 | G | A | 6.32E-05 | A | 6.32E-05 | | | ORF1ab | synonymous_K6768K; synonymous_K6768K |
| 20598 | A | G | 6.32E-05 | | | | | ORF1ab | synonymous_E6778E |
| 20668 | G | A | 6.32E-05 | | | | | ORF1ab | nonsynonymous_A6802T |
| 20852 | T | A | 6.32E-05 | | | | | ORF1ab | nonsynonymous_M6863K |
| 20898 | A | G | 6.32E-05 | | | | | ORF1ab | synonymous_P6878P |
| 20930 | C | T | 6.32E-05 | | | | | ORF1ab | nonsynonymous_T6889M |
| 21005 | C | T | 6.32E-05 | | | | | ORF1ab | nonsynonymous_A6914V |
| 2129 | G | A | 6.32E-05 | | | | | ORF1ab | nonsynonymous_E622K |
| 21404 | A | G | 6.32E-05 | | | | | ORF1ab | nonsynonymous_K7047R |
| 21410 | C | T | 6.32E-05 | | | | | ORF1ab | nonsynonymous_P7049L |
| 21411 | C | T | 6.32E-05 | | | | | ORF1ab | synonymous_P7049P |
| 2182 | A | G | 6.32E-05 | | | | | ORF1ab | synonymous_V639V |
| 21997 | C | A | 6.32E-05 | A | 6.32E-05 | | | S | stopgain_Y145X; stopgain_Y145X |
| 22102 | A | C | 6.32E-05 | | | | | S | nonsynonymous_E180D |
| 22168 | A | C | 6.32E-05 | | | | | S | nonsynonymous_K202N |
| 22186 | G | A | 6.32E-05 | | | | | S | synonymous_T208T |

| | | | | | | | | | |
|---|---|---|---|---|---|---|---|---|---|
| 22273 | G | A | 6.32E-05 | | | | | S | synonymous_R237R |
| 22285 | A | C | 6.32E-05 | | | | | S | nonsynonymous_L241F |
| 22291 | T | C | 6.32E-05 | | | | | S | synonymous_A243A |
| 22299 | G | T | 6.32E-05 | | | | | S | nonsynonymous_R246I |
| 22313 | C | T | 6.32E-05 | | | | | S | nonsynonymous_P251S |
| 23585 | C | A | 6.32E-05 | | | | | S | nonsynonymous_Q675K |
| 23588 | A | T | 6.32E-05 | | | | | S | nonsynonymous_T676S |
| 23612 | G | A | 6.32E-05 | | | | | S | nonsynonymous_A684T |
| 24334 | C | T | 6.32E-05 | | | | | S | synonymous_A924A |
| 24380 | T | A | 6.32E-05 | | | | | S | nonsynonymous_S940T |
| 24475 | T | A | 6.32E-05 | | | | | S | synonymous_G971G |
| 24520 | A | G | 6.32E-05 | | | | | S | synonymous_K986K |
| 24784 | C | T | 6.32E-05 | | | | | S | synonymous_N1074N |
| 24788 | A | T | 6.32E-05 | | | | | S | nonsynonymous_T1076S |
| 24796 | T | C | 6.32E-05 | | | | | S | synonymous_A1078A |
| 24805 | T | C | 6.32E-05 | | | | | S | synonymous_I1081I |
| 24817 | A | C | 6.32E-05 | | | | | S | synonymous_G1085G |
| 24818 | A | T | 6.32E-05 | | | | | S | stopgain_K1086X |
| 24819 | A | G | 6.32E-05 | | | | | S | nonsynonymous_K1086R |
| 24832 | T | C | 6.32E-05 | | | | | S | synonymous_P1090P |
| 24834 | G | T | 6.32E-05 | | | | | S | nonsynonymous_R1091L |
| 24851 | T | A | 6.32E-05 | | | | | S | nonsynonymous_S1097T |
| 24865 | C | T | 6.32E-05 | | | | | S | synonymous_H1101H |
| 24877 | A | G | 6.32E-05 | | | | | S | synonymous_T1105T |
| 24896 | C | T | 6.32E-05 | | | | | S | nonsynonymous_P1112S |
| 24902 | A | G | 6.32E-05 | | | | | S | nonsynonymous_I1114V |
| 24907 | T | C | 6.32E-05 | | | | | S | synonymous_I1115I |
| 24912 | C | T | 6.32E-05 | | | | | S | nonsynonymous_T1117I |
| 24947 | G | C | 6.32E-05 | | | | | S | nonsynonymous_V1129L |
| 24956 | A | G | 6.32E-05 | | | | | S | nonsynonymous_I1132V |

| | | | | | | | | | |
|---|---|---|---|---|---|---|---|---|---|
| 24977 | G | T | 6.32E-05 | | | | | S | nonsynonymous_D1139Y |
| 24979 | T | C | 6.32E-05 | | | | | S | synonymous_D1139D |
| 24985 | G | T | 6.32E-05 | | | | | S | nonsynonymous_L1141F |
| 24995 | T | C | 6.32E-05 | | | | | S | synonymous_L1145L |
| 25065 | A | G | 6.32E-05 | | | | | S | nonsynonymous_D1168G |
| 25067 | A | T | 6.32E-05 | | | | | S | nonsynonymous_I1169F |
| 25070 | T | C | 6.32E-05 | | | | | S | nonsynonymous_S1170P |
| 25072 | T | C | 6.32E-05 | | | | | S | synonymous_S1170S |
| 25073 | G | T | 6.32E-05 | | | | | S | nonsynonymous_G1171C |
| 25090 | T | C | 6.32E-05 | | | | | S | synonymous_V1176V |
| 25093 | A | C | 6.32E-05 | | | | | S | synonymous_V1177V |
| 25095 | A | T | 6.32E-05 | | | | | S | nonsynonymous_N1178I |
| 25099 | T | G | 6.32E-05 | | | | | S | nonsynonymous_I1179M |
| 25100 | C | G | 6.32E-05 | | | | | S | nonsynonymous_Q1180E |
| 25104 | A | G | 6.32E-05 | | | | | S | nonsynonymous_K1181R |
| 25108 | A | C | 6.32E-05 | | | | | S | nonsynonymous_E1182D |
| 25110 | T | C | 6.32E-05 | | | | | S | nonsynonymous_I1183T |
| 25112 | G | T | 6.32E-05 | | | | | S | nonsynonymous_D1184Y |
| 25113 | A | T | 6.32E-05 | | | | | S | nonsynonymous_D1184V |
| 25115 | C | A | 6.32E-05 | | | | | S | nonsynonymous_R1185S |
| 25116 | G | A | 6.32E-05 | A | 6.32E-05 | | | S | nonsynonymous_R1185H; nonsynonymous_R1185H |
| 25122 | A | G | 6.32E-05 | | | | | S | nonsynonymous_N1187S |
| 25123 | T | A | 6.32E-05 | | | | | S | nonsynonymous_N1187K |
| 25125 | A | C | 6.32E-05 | | | | | S | nonsynonymous_E1188A |
| 25126 | G | A | 6.32E-05 | | | | | S | synonymous_E1188E |
| 25128 | T | G | 6.32E-05 | | | | | S | nonsynonymous_V1189G |
| 25131 | C | T | 6.32E-05 | T | 6.32E-05 | | | S | nonsynonymous_A1190V; nonsynonymous_A1190V |
| 25132 | C | T | 6.32E-05 | | | | | S | synonymous_A1190A |

| | | | | | | | | | |
|---|---|---|---|---|---|---|---|---|---|
| 25133 | A | T | 6.32E-05 | | | | | S | stopgain_K1191X |
| 25134 | A | T | 6.32E-05 | | | | | S | nonsynonymous_K1191M |
| 25136 | A | G | 6.32E-05 | | | | | S | nonsynonymous_N1192D |
| 25137 | A | T | 6.32E-05 | | | | | S | nonsynonymous_N1192I |
| 25140 | T | C | 6.32E-05 | | | | | S | nonsynonymous_L1193S |
| 25143 | A | G | 6.32E-05 | | | | | S | nonsynonymous_N1194S |
| 25146 | A | G | 6.32E-05 | | | | | S | nonsynonymous_E1195G |
| 25150 | T | A | 6.32E-05 | | | | | S | synonymous_S1196S |
| 25152 | T | A | 6.32E-05 | | | | | S | nonsynonymous_L1197H |
| 25153 | C | T | 6.32E-05 | | | | | S | synonymous_L1197L |
| 25155 | T | C | 6.32E-05 | | | | | S | nonsynonymous_I1198T |
| 25158 | A | G | 6.32E-05 | | | | | S | nonsynonymous_D1199G |
| 25168 | A | C | 6.32E-05 | | | | | S | nonsynonymous_E1202D |
| 25170 | T | A | 6.32E-05 | | | | | S | nonsynonymous_L1203H |
| 25171 | T | A | 6.32E-05 | | | | | S | synonymous_L1203L |
| 25176 | A | T | 6.32E-05 | | | | | S | nonsynonymous_K1205M |
| 25179 | A | T | 6.32E-05 | | | | | S | nonsynonymous_Y1206F |
| 25181 | G | A | 6.32E-05 | | | | | S | nonsynonymous_E1207K |
| 25182 | A | T | 6.32E-05 | | | | | S | nonsynonymous_E1207V |
| 25183 | G | T | 6.32E-05 | | | | | S | nonsynonymous_E1207D |
| 25184 | C | A | 6.32E-05 | | | | | S | nonsynonymous_Q1208K |
| 25189 | T | C | 6.32E-05 | | | | | S | synonymous_Y1209Y |
| 25193 | A | T | 6.32E-05 | | | | | S | stopgain_K1211X |
| 25194 | A | C | 6.32E-05 | | | | | S | nonsynonymous_K1211T |
| 25200 | C | T | 6.32E-05 | | | | | S | nonsynonymous_P1213L |
| 25203 | G | A | 6.32E-05 | | | | | S | stopgain_W1214X |
| 25205 | T | C | 6.32E-05 | | | | | S | nonsynonymous_Y1215H |
| 25220 | T | C | 6.32E-05 | | | | | S | nonsynonymous_F1220L |
| 25223 | A | G | 6.32E-05 | | | | | S | nonsynonymous_I1221V |
| 25226 | G | A | 6.32E-05 | | | | | S | nonsynonymous_A1222T |

| | | | | | | | | | |
|---|---|---|---|---|---|---|---|---|---|
| 25240 | C | A | 6.32E-05 | | | | | S | synonymous_A1226A |
| 25243 | A | G | 6.32E-05 | | | | | S | nonsynonymous_I1227M |
| 25254 | C | A | 6.32E-05 | | | | | S | nonsynonymous_T1231K |
| 25255 | A | C | 6.32E-05 | | | | | S | synonymous_T1231T |
| 25257 | T | A | 6.32E-05 | | | | | S | nonsynonymous_I1232N |
| 25258 | T | C | 6.32E-05 | | | | | S | synonymous_I1232I |
| 25259 | A | G | 6.32E-05 | | | | | S | nonsynonymous_M1233V |
| 25260 | T | C | 6.32E-05 | | | | | S | nonsynonymous_M1233T |
| 25261 | G | A | 6.32E-05 | | | | | S | nonsynonymous_M1233I |
| 25262 | C | G | 6.32E-05 | | | | | S | nonsynonymous_L1234V |
| 25282 | C | T | 6.32E-05 | | | | | S | synonymous_C1240C |
| 25283 | T | G | 6.32E-05 | | | | | S | nonsynonymous_C1241G |
| 25284 | G | A | 6.32E-05 | | | | | S | nonsynonymous_C1241Y |
| 25287 | G | A | 6.32E-05 | | | | | S | nonsynonymous_S1242N |
| 25288 | T | A | 6.32E-05 | | | | | S | nonsynonymous_S1242R |
| 25292 | C | T | 6.32E-05 | | | | | S | nonsynonymous_L1244F |
| 25294 | C | T | 6.32E-05 | | | | | S | synonymous_L1244L |
| 25298 | G | A | 6.32E-05 | | | | | S | nonsynonymous_G1246S |
| 25299 | G | A | 6.32E-05 | | | | | S | nonsynonymous_G1246D |
| 25300 | C | T | 6.32E-05 | | | | | S | synonymous_G1246G |
| 25301 | T | C | 6.32E-05 | | | | | S | nonsynonymous_C1247R |
| 25424 | G | T | 6.32E-05 | | | | | ORF3a | nonsynonymous_G11V |
| 25507 | G | A | 6.32E-05 | | | | | ORF3a | nonsynonymous_A39T |
| 25519 | T | A | 6.32E-05 | | | | | ORF3a | nonsynonymous_F43I |
| 25522 | G | A | 6.32E-05 | | | | | ORF3a | nonsynonymous_G44R |
| 26359 | T | A | 6.32E-05 | | | | | E | nonsynonymous_L39M |
| 26365 | G | T | 6.32E-05 | | | | | E | nonsynonymous_A41S |
| 26392 | A | G | 6.32E-05 | | | | | E | nonsynonymous_S50G |
| 26398 | G | A | 6.32E-05 | | | | | E | nonsynonymous_V52I |
| 26404 | C | T | 6.32E-05 | | | | | E | nonsynonymous_P54S |

| | | | | | | | | | |
|---|---|---|---|---|---|---|---|---|---|
| 26426 | G | A | 6.32E-05 | | | | | E | nonsynonymous_R61H |
| 26449 | A | C | 6.32E-05 | | | | | E | synonymous_R69R |
| 26453 | T | C | 6.32E-05 | | | | | E | nonsynonymous_V70A |
| 26455 | C | G | 6.32E-05 | | | | | E | nonsynonymous_P71A |
| 26457 | T | A | 6.32E-05 | | | | | E | synonymous_P71P |
| 26462 | T | A | 6.32E-05 | | | | | E | nonsynonymous_L73H |
| 26478 | C | T | 6.32E-05 | | | | | intergenic | |
| 26483 | T | C | 6.32E-05 | | | | | intergenic | |
| 26708 | A | G | 6.32E-05 | | | | | M | synonymous_L62L |
| 26789 | C | T | 6.32E-05 | | | | | M | synonymous_G89G |
| 26815 | C | A | 6.32E-05 | | | | | M | nonsynonymous_A98D |
| 26918 | G | T | 6.32E-05 | | | | | M | synonymous_P132P |
| 26985 | C | T | 6.32E-05 | | | | | M | nonsynonymous_H155Y |
| 26987 | T | C | 6.32E-05 | | | | | M | synonymous_H155H |
| 269 | G | A | 6.32E-05 | | | | | ORF1ab | nonsynonymous_E2K |
| 27039 | T | A | 6.32E-05 | | | | | M | nonsynonymous_S173T |
| 27040 | C | A | 6.32E-05 | | | | | M | stopgain_S173X |
| 27056 | T | C | 6.32E-05 | | | | | M | synonymous_Y178Y |
| 27063 | T | C | 6.32E-05 | | | | | M | synonymous_L181L |
| 27074 | G | A | 6.32E-05 | | | | | M | synonymous_S184S |
| 27090 | G | A | 6.32E-05 | | | | | M | nonsynonymous_D190N |
| 27095 | A | T | 6.32E-05 | | | | | M | synonymous_S191S |
| 27098 | T | G | 6.32E-05 | | | | | M | synonymous_G192G |
| 27120 | A | T | 6.32E-05 | | | | | M | nonsynonymous_R200W |
| 27135 | A | T | 6.32E-05 | | | | | M | stopgain_K205X |
| 27136 | A | T | 6.32E-05 | | | | | M | nonsynonymous_K205I |
| 27137 | A | T | 6.32E-05 | | | | | M | nonsynonymous_K205N |
| 27139 | T | C | 6.32E-05 | | | | | M | nonsynonymous_L206S |
| 27143 | C | T | 6.32E-05 | | | | | M | synonymous_N207N |
| 27150 | C | T | 6.32E-05 | | | | | M | nonsynonymous_H210Y |

| | | | | | | | | | |
|---|---|---|---|---|---|---|---|---|---|
| 27191 | A | G | 6.32E-05 | | | | | M | synonymous_X223X |
| 27272 | T | C | 6.32E-05 | | | | | ORF6 | nonsynonymous_V24A |
| 27273 | T | C | 6.32E-05 | | | | | ORF6 | synonymous_V24V |
| 27280 | T | C | 6.32E-05 | | | | | ORF6 | nonsynonymous_W27R |
| 27282 | G | A | 6.32E-05 | | | | | ORF6 | stopgain_W27X |
| 27285 | T | C | 6.32E-05 | | | | | ORF6 | synonymous_N28N |
| 27292 | T | C | 6.32E-05 | | | | | ORF6 | nonsynonymous_Y31H |
| 27427 | C | T | 6.32E-05 | | | | | ORF7a | nonsynonymous_L12F |
| 27495 | T | A | 6.32E-05 | A | 6.32E-05 | | | ORF7a | synonymous_P34P; synonymous_P34P |
| 27497 | G | T | 6.32E-05 | | | | | ORF7a | nonsynonymous_C35F |
| 27499 | T | C | 6.32E-05 | | | | | ORF7a | nonsynonymous_S36P |
| 27502 | T | A | 6.32E-05 | | | | | ORF7a | nonsynonymous_S37T |
| 27504 | T | A | 6.32E-05 | | | | | ORF7a | synonymous_S37S |
| 27847 | C | T | 6.32E-05 | | | | | ORF7b | nonsynonymous_S31L |
| 28236 | C | T | 6.32E-05 | | | | | ORF8 | nonsynonymous_R115C |
| 28237 | G | T | 6.32E-05 | | | | | ORF8 | nonsynonymous_R115L |
| 28238 | T | A | 6.32E-05 | | | | | ORF8 | synonymous_R115R |
| 28239 | G | T | 6.32E-05 | | | | | ORF8 | nonsynonymous_V116F |
| 28240 | T | C | 6.32E-05 | | | | | ORF8 | nonsynonymous_V116A |
| 28242 | G | C | 6.32E-05 | | | | | ORF8 | nonsynonymous_V117L |
| 28325 | G | A | 6.32E-05 | A | 6.32E-05 | | | N | nonsynonymous_G18S; nonsynonymous_G18S |
| 28367 | C | T | 6.32E-05 | | | | | N | nonsynonymous_R32C |
| 28392 | G | A | 6.32E-05 | A | 6.32E-05 | | | N | nonsynonymous_R40H; nonsynonymous_R40H |
| 28409 | C | T | 6.32E-05 | | | | | N | nonsynonymous_P46S |
| 28760 | C | A | 6.32E-05 | | | | | N | nonsynonymous_Q163K |
| 28761 | A | G | 6.32E-05 | | | | | N | nonsynonymous_Q163R |
| 28762 | A | G | 6.32E-05 | | | | | N | synonymous_Q163Q |

| | | | | | | | | | |
|---|---|---|---|---|---|---|---|---|---|
| 28781 | G | T | 6.32E-05 | | | | | N | nonsynonymous_G170C |
| 28811 | A | T | 6.32E-05 | T | 6.32E-05 | | | N | nonsynonymous_S180C; nonsynonymous_S180C |
| 28817 | G | A | 6.32E-05 | A | 6.32E-05 | | | N | nonsynonymous_A182T; nonsynonymous_A182T |
| 28818 | C | T | 6.32E-05 | | | | | N | nonsynonymous_A182V |
| 28819 | C | T | 6.32E-05 | | | | | N | synonymous_A182A |
| 28820 | T | A | 6.32E-05 | | | | | N | nonsynonymous_S183T |
| 28823 | T | C | 6.32E-05 | | | | | N | nonsynonymous_S184P |
| 28833 | C | T | 6.32E-05 | | | | | N | nonsynonymous_S187L |
| 28834 | A | G | 6.32E-05 | | | | | N | synonymous_S187S |
| 28846 | C | T | 6.32E-05 | | | | | N | synonymous_R191R |
| 28848 | A | G | 6.32E-05 | | | | | N | nonsynonymous_N192S |
| 28850 | A | T | 6.32E-05 | | | | | N | nonsynonymous_S193C |
| 28858 | A | G | 6.32E-05 | | | | | N | synonymous_R195R |
| 28876 | C | T | 6.32E-05 | | | | | N | synonymous_S201S |
| 28910 | A | T | 6.32E-05 | | | | | N | nonsynonymous_N213Y |
| 28912 | T | C | 6.32E-05 | | | | | N | synonymous_N213N |
| 28913 | G | A | 6.32E-05 | A | 6.32E-05 | | | N | nonsynonymous_G214S; nonsynonymous_G214S |
| 28919 | G | A | 6.32E-05 | | | | | N | nonsynonymous_D216N |
| 28925 | G | T | 6.32E-05 | | | | | N | nonsynonymous_A218S |
| 28926 | C | T | 6.32E-05 | | | | | N | nonsynonymous_A218V |
| 28931 | G | A | 6.32E-05 | | | | | N | nonsynonymous_A220T |
| 28937 | C | T | 6.32E-05 | | | | | N | synonymous_L222L |
| 28940 | C | T | 6.32E-05 | | | | | N | synonymous_L223L |
| 28963 | T | A | 6.32E-05 | | | | | N | synonymous_L230L |
| 28976 | T | C | 6.32E-05 | | | | | N | nonsynonymous_S235P |
| 29001 | G | A | 6.32E-05 | | | | | N | nonsynonymous_G243D |
| 5000 | C | T | 6.32E-05 | | | | | ORF1ab | nonsynonymous_L1579F |

| | | | | | | | | | |
|---|---|---|---|---|---|---|---|---|---|
| 5002 | C | T | 6.32E-05 | | | | | ORF1ab | synonymous_L1579L |
| 5005 | C | T | 6.32E-05 | | | | | ORF1ab | synonymous_H1580H |
| 5012 | G | A | 6.32E-05 | | | | | ORF1ab | nonsynonymous_V1583I |
| 5017 | G | A | 6.32E-05 | A | 6.32E-05 | | | ORF1ab | synonymous_V1584V; synonymous_V1584V |
| 5018 | G | A | 6.32E-05 | | | | | ORF1ab | nonsynonymous_D1585N |
| 5039 | C | A | 6.32E-05 | | | | | ORF1ab | nonsynonymous_Q1592K |
| 5052 | C | T | 6.32E-05 | | | | | ORF1ab | nonsynonymous_P1596L |
| 5063 | G | A | 6.32E-05 | | | | | ORF1ab | nonsynonymous_D1600N |
| 5116 | A | C | 6.32E-05 | | | | | ORF1ab | synonymous_T1617T |
| 5124 | T | G | 6.32E-05 | | | | | ORF1ab | nonsynonymous_V1620G |
| 5148 | G | T | 6.32E-05 | | | | | ORF1ab | nonsynonymous_R1628L |
| 5150 | G | A | 6.32E-05 | | | | | ORF1ab | nonsynonymous_V1629I |
| 5151 | T | C | 6.32E-05 | | | | | ORF1ab | nonsynonymous_V1629A |
| 5165 | T | C | 6.32E-05 | | | | | ORF1ab | nonsynonymous_Y1634H |
| 5173 | C | T | 6.32E-05 | | | | | ORF1ab | synonymous_H1636H |
| 5176 | A | G | 6.32E-05 | | | | | ORF1ab | synonymous_T1637T |
| 5210 | G | T | 6.32E-05 | | | | | ORF1ab | nonsynonymous_A1649S |
| 5220 | A | C | 6.32E-05 | | | | | ORF1ab | nonsynonymous_H1652P |
| 5224 | T | A | 6.32E-05 | | | | | ORF1ab | synonymous_T1653T |
| 5240 | C | G | 6.32E-05 | | | | | ORF1ab | nonsynonymous_P1659A |
| 5357 | G | T | 6.32E-05 | | | | | ORF1ab | nonsynonymous_A1698S |
| 5365 | C | T | 6.32E-05 | T | 6.32E-05 | | | ORF1ab | synonymous_Y1700Y; synonymous_Y1700Y |
| 5384 | G | T | 6.32E-05 | | | | | ORF1ab | nonsynonymous_A1707S |
| 5385 | C | T | 6.32E-05 | | | | | ORF1ab | nonsynonymous_A1707V |
| 5405 | A | G | 6.32E-05 | | | | | ORF1ab | nonsynonymous_I1714V |
| 5406 | T | C | 6.32E-05 | | | | | ORF1ab | nonsynonymous_I1714T |
| 5407 | C | T | 6.32E-05 | | | | | ORF1ab | synonymous_I1714I |
| 5409 | T | A | 6.32E-05 | | | | | ORF1ab | stopgain_L1715X |

| | | | | | | | | | |
|---|---|---|---|---|---|---|---|---|---|
| 5429 | G | T | 6.32E-05 | | | | | ORF1ab | nonsynonymous_V1722L |
| 5432 | G | T | 6.32E-05 | | | | | ORF1ab | nonsynonymous_G1723C |
| 5446 | T | C | 6.32E-05 | | | | | ORF1ab | synonymous_D1727D |
| 5449 | T | C | 6.32E-05 | | | | | ORF1ab | synonymous_V1728V |
| 5477 | C | T | 6.32E-05 | | | | | ORF1ab | nonsynonymous_H1738Y |
| 5485 | T | G | 6.32E-05 | | | | | ORF1ab | nonsynonymous_N1740K |
| 5513 | G | A | 6.32E-05 | | | | | ORF1ab | nonsynonymous_V1750M |
| 5536 | A | G | 6.32E-05 | | | | | ORF1ab | synonymous_Q1757Q |
| 5542 | G | A | 6.32E-05 | | | | | ORF1ab | synonymous_Q1759Q |
| 5547 | C | T | 6.32E-05 | | | | | ORF1ab | nonsynonymous_T1761I |
| 5548 | C | T | 6.32E-05 | | | | | ORF1ab | synonymous_T1761T |
| 5567 | G | A | 6.32E-05 | | | | | ORF1ab | nonsynonymous_V1768I |
| 6553 | G | T | 6.32E-05 | | | | | ORF1ab | nonsynonymous_M2096I |
| 6963 | T | A | 6.32E-05 | | | | | ORF1ab | nonsynonymous_F2233Y |
| 7016 | G | A | 6.32E-05 | | | | | ORF1ab | nonsynonymous_G2251S |
| 7027 | G | T | 6.32E-05 | | | | | ORF1ab | nonsynonymous_M2254I |
| 7082 | T | C | 6.32E-05 | | | | | ORF1ab | nonsynonymous_S2273P |
| 7310 | T | A | 6.32E-05 | | | | | ORF1ab | nonsynonymous_L2349M |
| 8770 | C | T | 6.32E-05 | | | | | ORF1ab | synonymous_D2835D |
| 8772 | C | T | 6.32E-05 | | | | | ORF1ab | nonsynonymous_T2836I |
| 8873 | C | T | 6.32E-05 | | | | | ORF1ab | nonsynonymous_P2870S |
| 8895 | C | T | 6.32E-05 | | | | | ORF1ab | nonsynonymous_T2877I |
| 9281 | C | A | 6.32E-05 | | | | | ORF1ab | nonsynonymous_L3006I |
| 9283 | T | G | 6.32E-05 | | | | | ORF1ab | synonymous_L3006L |
| 9369 | T | A | 6.32E-05 | | | | | ORF1ab | nonsynonymous_I3035N |
| 9388 | G | T | 6.32E-05 | | | | | ORF1ab | nonsynonymous_L3041F |
| 9471 | G | C | 6.32E-05 | | | | | ORF1ab | nonsynonymous_R3069T |
| 9530 | T | G | 6.32E-05 | | | | | ORF1ab | nonsynonymous_F3089V |
| 10207 | C | T | 6.32E-05 | | | | | ORF1ab | synonymous_N3314N |
| 10846 | G | T | 6.32E-05 | | | | | ORF1ab | nonsynonymous_M3527I |

| | | | | | | | | | |
|---|---|---|---|---|---|---|---|---|---|
| 10963 | T | A | 6.32E-05 | | | | | ORF1ab | synonymous_V3566V |
| 13193 | G | T | 6.32E-05 | | | | | ORF1ab | nonsynonymous_V4310F |
| 13240 | G | A | 6.32E-05 | | | | | ORF1ab | synonymous_S4325S |
| 13640 | A | G | 6.32E-05 | | | | | ORF1ab | nonsynonymous_D4459G |
| 13684 | C | T | 6.32E-05 | | | | | ORF1ab | nonsynonymous_H4474Y |
| 13724 | C | T | 6.32E-05 | | | | | ORF1ab | nonsynonymous_A4487V |
| 13748 | A | G | 6.32E-05 | | | | | ORF1ab | nonsynonymous_K4495R |
| 13920 | G | T | 6.32E-05 | | | | | ORF1ab | nonsynonymous_K4552N |
| 14022 | T | A | 6.32E-05 | | | | | ORF1ab | nonsynonymous_D4586E |
| 14224 | A | G | 6.32E-05 | | | | | ORF1ab | nonsynonymous_T4654A |
| 14318 | C | T | 6.32E-05 | | | | | ORF1ab | nonsynonymous_T4685I |
| 14323 | C | T | 6.32E-05 | | | | | ORF1ab | nonsynonymous_H4687Y |
| 14330 | A | G | 6.32E-05 | | | | | ORF1ab | nonsynonymous_N4689S |
| 14336 | T | C | 6.32E-05 | | | | | ORF1ab | nonsynonymous_V4691A |
| 1433 | G | A | 6.32E-05 | | | | | ORF1ab | nonsynonymous_E390K |
| 14411 | C | T | 6.32E-05 | | | | | ORF1ab | nonsynonymous_T4716I |
| 14485 | A | G | 6.32E-05 | | | | | ORF1ab | nonsynonymous_R4741G |
| 14514 | T | C | 6.32E-05 | | | | | ORF1ab | synonymous_D4750D |
| 14529 | C | T | 6.32E-05 | | | | | ORF1ab | synonymous_S4755S |
| 14565 | T | A | 6.32E-05 | | | | | ORF1ab | synonymous_A4767A |
| 14576 | C | T | 6.32E-05 | | | | | ORF1ab | nonsynonymous_A4771V |
| 14584 | G | A | 6.32E-05 | | | | | ORF1ab | nonsynonymous_A4774T |
| 14644 | A | G | 6.32E-05 | | | | | ORF1ab | nonsynonymous_T4794A |
| 14709 | T | C | 6.32E-05 | | | | | ORF1ab | synonymous_A4815A |
| 15154 | C | T | 6.32E-05 | | | | | ORF1ab | nonsynonymous_H4964Y |
| 15189 | A | G | 6.32E-05 | | | | | ORF1ab | synonymous_R4975R |
| 15190 | G | T | 6.32E-05 | | | | | ORF1ab | stopgain_G4976X |
| 15191 | G | T | 6.32E-05 | | | | | ORF1ab | nonsynonymous_G4976V |
| 15714 | C | T | 6.32E-05 | | | | | ORF1ab | synonymous_L5150L |
| 15769 | G | A | 6.32E-05 | | | | | ORF1ab | nonsynonymous_A5169T |

| | | | | | | | | | |
|---|---|---|---|---|---|---|---|---|---|
| 15810 | C | T | 6.32E-05 | | | | | ORF1ab | synonymous_N5182N |
| 15847 | A | T | 6.32E-05 | T | 6.32E-05 | | | ORF1ab | nonsynonymous_T5195S; nonsynonymous_T5195S |
| 15848 | C | T | 6.32E-05 | | | | | ORF1ab | nonsynonymous_T5195I |
| 15849 | T | C | 6.32E-05 | | | | | ORF1ab | synonymous_T5195T |
| 15861 | A | G | 6.32E-05 | | | | | ORF1ab | synonymous_K5199K |
| 16085 | A | G | 6.32E-05 | | | | | ORF1ab | nonsynonymous_H5274R |
| 16114 | C | T | 6.32E-05 | | | | | ORF1ab | nonsynonymous_H5284Y |
| 16141 | G | T | 6.32E-05 | | | | | ORF1ab | nonsynonymous_D5293Y |
| 16299 | T | C | 6.32E-05 | | | | | ORF1ab | synonymous_R5345R |
| 16311 | A | G | 6.32E-05 | | | | | ORF1ab | synonymous_L5349L |
| 16325 | G | C | 6.32E-05 | | | | | ORF1ab | nonsynonymous_C5354S |
| 16521 | T | C | 6.32E-05 | | | | | ORF1ab | synonymous_N5419N |
| 16523 | C | A | 6.32E-05 | | | | | ORF1ab | nonsynonymous_T5420K |
| 16536 | C | T | 6.32E-05 | | | | | ORF1ab | synonymous_S5424S |
| 16545 | T | C | 6.32E-05 | | | | | ORF1ab | synonymous_V5427V |
| 16554 | T | C | 6.32E-05 | | | | | ORF1ab | synonymous_F5430F |
| 16592 | A | G | 6.32E-05 | | | | | ORF1ab | nonsynonymous_D5443G |
| 16596 | C | T | 6.32E-05 | | | | | ORF1ab | synonymous_Y5444Y |
| 16626 | C | T | 6.32E-05 | | | | | ORF1ab | synonymous_L5454L |
| 16650 | C | A | 6.32E-05 | | | | | ORF1ab | synonymous_L5462L |
| 16682 | A | G | 6.32E-05 | | | | | ORF1ab | nonsynonymous_Y5473C |
| 16687 | A | G | 6.32E-05 | | | | | ORF1ab | nonsynonymous_I5475V |
| 16700 | G | T | 6.32E-05 | | | | | ORF1ab | nonsynonymous_R5479L |
| 16701 | T | C | 6.32E-05 | | | | | ORF1ab | synonymous_R5479R |
| 16718 | G | T | 6.32E-05 | | | | | ORF1ab | nonsynonymous_R5485I |
| 16744 | G | A | 6.32E-05 | | | | | ORF1ab | nonsynonymous_G5494S |
| 16748 | A | G | 6.32E-05 | | | | | ORF1ab | nonsynonymous_K5495R |
| 16946 | C | T | 6.32E-05 | | | | | ORF1ab | nonsynonymous_A5561V |
| 17019 | G | T | 6.32E-05 | | | | | ORF1ab | nonsynonymous_E5585D |

| | | | | | | | | | |
|---|---|---|---|---|---|---|---|---|---|
| 17137 | T | A | 6.32E-05 | | | | | ORF1ab | nonsynonymous_S5625T |
| 17202 | A | G | 6.32E-05 | | | | | ORF1ab | synonymous_L5646L |
| 1730 | G | A | 6.32E-05 | | | | | ORF1ab | nonsynonymous_E489K |
| 17475 | T | C | 6.32E-05 | | | | | ORF1ab | synonymous_T5737T |
| 17506 | T | A | 6.32E-05 | | | | | ORF1ab | nonsynonymous_S5748T |
| 17533 | G | A | 6.32E-05 | | | | | ORF1ab | nonsynonymous_G5757S |
| 17541 | C | T | 6.32E-05 | | | | | ORF1ab | synonymous_D5759D |
| 17542 | A | G | 6.32E-05 | G | 6.32E-05 | | | ORF1ab | nonsynonymous_M5760V; nonsynonymous_M5760V |
| 17555 | C | T | 6.32E-05 | | | | | ORF1ab | nonsynonymous_T5764I |
| 17567 | G | A | 6.32E-05 | | | | | ORF1ab | nonsynonymous_C5768Y |
| 17569 | C | T | 6.32E-05 | | | | | ORF1ab | nonsynonymous_P5769S |
| 17574 | T | A | 6.32E-05 | A | 6.32E-05 | | | ORF1ab | synonymous_A5770A; synonymous_A5770A |
| 17599 | T | A | 6.32E-05 | | | | | ORF1ab | nonsynonymous_L5779M |
| 17602 | G | A | 6.32E-05 | | | | | ORF1ab | nonsynonymous_V5780I |
| 17622 | A | G | 6.32E-05 | | | | | ORF1ab | synonymous_K5786K |
| 17626 | C | T | 6.32E-05 | | | | | ORF1ab | nonsynonymous_H5788Y |
| 17940 | A | T | 6.32E-05 | | | | | ORF1ab | synonymous_A5892A |
| 17977 | C | T | 6.32E-05 | | | | | ORF1ab | nonsynonymous_L5905F |
| 18099 | T | A | 6.32E-05 | A | 6.32E-05 | | | ORF1ab | synonymous_P5945P; synonymous_P5945P |
| 18103 | C | T | 6.32E-05 | | | | | ORF1ab | stopgain_Q5947X |
| 18121 | A | G | 6.32E-05 | | | | | ORF1ab | nonsynonymous_S5953G |
| 18131 | C | T | 6.32E-05 | | | | | ORF1ab | nonsynonymous_T5956I |
| 18134 | A | G | 6.32E-05 | | | | | ORF1ab | nonsynonymous_K5957R |
| 18138 | C | G | 6.32E-05 | | | | | ORF1ab | nonsynonymous_F5958L |
| 18147 | A | G | 6.32E-05 | | | | | ORF1ab | synonymous_E5961E |
| 18158 | T | C | 6.32E-05 | | | | | ORF1ab | nonsynonymous_V5965A |
| 18162 | C | T | 6.32E-05 | | | | | ORF1ab | synonymous_D5966D |

| | | | | | | | | | |
|---|---|---|---|---|---|---|---|---|---|
| 18163 | A | G | 6.32E-05 | | | | | ORF1ab | nonsynonymous_I5967V |
| 18170 | G | A | 6.32E-05 | | | | | ORF1ab | nonsynonymous_G5969D |
| 18175 | C | T | 6.32E-05 | | | | | ORF1ab | nonsynonymous_P5971S |
| 18180 | G | A | 6.32E-05 | | | | | ORF1ab | synonymous_K5972K |
| 18189 | C | T | 6.32E-05 | | | | | ORF1ab | synonymous_T5975T |
| 18196 | A | C | 6.32E-05 | | | | | ORF1ab | synonymous_R5978R |
| 18210 | G | T | 6.32E-05 | | | | | ORF1ab | nonsynonymous_M5982I |
| 18212 | T | A | 6.32E-05 | | | | | ORF1ab | nonsynonymous_M5983K |
| 18225 | G | T | 6.32E-05 | | | | | ORF1ab | nonsynonymous_M5987I |
| 18230 | A | T | 6.32E-05 | | | | | ORF1ab | nonsynonymous_Y5989F |
| 18233 | A | T | 6.32E-05 | | | | | ORF1ab | nonsynonymous_Q5990L |
| 18490 | C | G | 6.32E-05 | | | | | ORF1ab | nonsynonymous_P6076A |
| 18534 | G | C | 6.32E-05 | | | | | ORF1ab | nonsynonymous_K6090N |
| 18535 | A | T | 6.32E-05 | | | | | ORF1ab | nonsynonymous_I6091F |
| 18612 | G | C | 6.32E-05 | | | | | ORF1ab | nonsynonymous_E6116D |
| 18640 | A | T | 6.32E-05 | | | | | ORF1ab | nonsynonymous_I6126L |
| 18646 | C | T | 6.32E-05 | | | | | ORF1ab | nonsynonymous_P6128S |
| 18653 | G | T | 6.32E-05 | T | 6.32E-05 | | | ORF1ab | nonsynonymous_R6130L;<br>nonsynonymous_R6130L |
| 18655 | A | C | 6.32E-05 | | | | | ORF1ab | nonsynonymous_T6131P |
| 18662 | G | T | 6.32E-05 | | | | | ORF1ab | nonsynonymous_C6133F |
| 18663 | T | A | 6.32E-05 | | | | | ORF1ab | stopgain_C6133X |
| 18664 | C | A | 6.32E-05 | | | | | ORF1ab | nonsynonymous_L6134I |
| 18665 | T | C | 6.32E-05 | | | | | ORF1ab | nonsynonymous_L6134P |
| 18668 | G | A | 6.32E-05 | | | | | ORF1ab | nonsynonymous_C6135Y |
| 18670 | G | T | 6.32E-05 | | | | | ORF1ab | nonsynonymous_D6136Y |
| 18679 | G | A | 6.32E-05 | | | | | ORF1ab | nonsynonymous_A6139T |
| 18682 | A | G | 6.32E-05 | G | 6.32E-05 | | | ORF1ab | nonsynonymous_T6140A;<br>nonsynonymous_T6140A |
| 18683 | C | A | 6.32E-05 | | | | | ORF1ab | nonsynonymous_T6140K |

| | | | | | | | | | |
|---|---|---|---|---|---|---|---|---|---|
| 18685 | T | A | 6.32E-05 | | | | | ORF1ab | nonsynonymous_C6141S |
| 18692 | C | A | 6.32E-05 | A | 6.32E-05 | | | ORF1ab | nonsynonymous_S6143Y; nonsynonymous_S6143Y |
| 18695 | C | A | 6.32E-05 | | | | | ORF1ab | nonsynonymous_T6144N |
| 18698 | C | A | 6.32E-05 | A | 6.32E-05 | | | ORF1ab | nonsynonymous_A6145D; nonsynonymous_A6145D |
| 18702 | A | T | 6.32E-05 | T | 6.32E-05 | | | ORF1ab | synonymous_S6146S; synonymous_S6146S |
| 18703 | G | A | 6.32E-05 | | | | | ORF1ab | nonsynonymous_D6147N |
| 18705 | C | T | 6.32E-05 | | | | | ORF1ab | synonymous_D6147D |
| 18707 | C | G | 6.32E-05 | | | | | ORF1ab | nonsynonymous_T6148S |
| 18708 | T | G | 6.32E-05 | | | | | ORF1ab | synonymous_T6148T |
| 18709 | T | C | 6.32E-05 | | | | | ORF1ab | nonsynonymous_Y6149H |
| 18712 | G | A | 6.32E-05 | | | | | ORF1ab | nonsynonymous_A6150T |
| 18714 | C | G | 6.32E-05 | | | | | ORF1ab | synonymous_A6150A |
| 18718 | T | C | 6.32E-05 | | | | | ORF1ab | nonsynonymous_W6152R |
| 18719 | G | T | 6.32E-05 | | | | | ORF1ab | nonsynonymous_W6152L |
| 18721 | C | A | 6.32E-05 | | | | | ORF1ab | nonsynonymous_H6153N |
| 18726 | T | G | 6.32E-05 | | | | | ORF1ab | nonsynonymous_H6154Q |
| 18728 | C | G | 6.32E-05 | | | | | ORF1ab | nonsynonymous_S6155C |
| 18729 | T | G | 6.32E-05 | | | | | ORF1ab | synonymous_S6155S |
| 18764 | T | C | 6.32E-05 | | | | | ORF1ab | nonsynonymous_I6167T |
| 18823 | C | T | 6.32E-05 | | | | | ORF1ab | stopgain_Q6187X |
| 18844 | G | A | 6.32E-05 | A | 6.32E-05 | | | ORF1ab | nonsynonymous_V6194I; nonsynonymous_V6194I |
| 18909 | T | A | 6.32E-05 | | | | | ORF1ab | synonymous_V6215V |
| 18910 | G | A | 6.32E-05 | | | | | ORF1ab | nonsynonymous_D6216N |
| 18915 | G | A | 6.32E-05 | | | | | ORF1ab | stopgain_W6217X |
| 18937 | G | A | 6.32E-05 | | | | | ORF1ab | nonsynonymous_G6225S |
| 19149 | T | C | 6.32E-05 | | | | | ORF1ab | synonymous_Y6295Y |

| | | | | | | | | | |
|---|---|---|---|---|---|---|---|---|---|
| 19164 | C | T | 6.32E-05 | | | | | ORF1ab | synonymous_D6300D |
| 19175 | A | C | 6.32E-05 | | | | | ORF1ab | nonsynonymous_D6304A |
| 19203 | T | C | 6.32E-05 | | | | | ORF1ab | synonymous_N6313N |
| 19273 | C | T | 6.32E-05 | | | | | ORF1ab | nonsynonymous_P6337S |
| 19579 | T | A | 6.32E-05 | | | | | ORF1ab | nonsynonymous_F6439I |
| 19580 | T | A | 6.32E-05 | | | | | ORF1ab | nonsynonymous_F6439Y |
| 19582 | G | C | 6.32E-05 | | | | | ORF1ab | nonsynonymous_D6440H |
| 19590 | T | C | 6.32E-05 | | | | | ORF1ab | synonymous_Y6442Y |
| 19626 | A | G | 6.32E-05 | | | | | ORF1ab | synonymous_L6454L |
| 19648 | G | T | 6.32E-05 | | | | | ORF1ab | nonsynonymous_V6462L |
| 19658 | G | A | 6.32E-05 | | | | | ORF1ab | nonsynonymous_G6465E |
| 19660 | C | A | 6.32E-05 | | | | | ORF1ab | nonsynonymous_H6466N |
| 19662 | C | T | 6.32E-05 | | | | | ORF1ab | synonymous_H6466H |
| 19668 | T | C | 6.32E-05 | | | | | ORF1ab | synonymous_D6468D |
| 19686 | A | G | 6.32E-05 | | | | | ORF1ab | synonymous_V6474V |
| 19711 | G | A | 6.32E-05 | | | | | ORF1ab | nonsynonymous_V6483I |
| 19814 | C | T | 6.32E-05 | | | | | ORF1ab | nonsynonymous_P6517L |
| 20610 | G | T | 6.32E-05 | | | | | ORF1ab | nonsynonymous_M6782I |
| 20649 | A | C | 6.32E-05 | | | | | ORF1ab | synonymous_P6795P |
| 20676 | A | G | 6.32E-05 | | | | | ORF1ab | synonymous_Q6804Q |
| 20799 | T | C | 6.32E-05 | | | | | ORF1ab | synonymous_Y6845Y |
| 20806 | C | T | 6.32E-05 | | | | | ORF1ab | synonymous_L6848L |
| 20827 | T | C | 6.32E-05 | | | | | ORF1ab | synonymous_L6855L |
| 20923 | T | C | 6.32E-05 | | | | | ORF1ab | synonymous_L6887L |
| 20941 | C | T | 6.32E-05 | | | | | ORF1ab | nonsynonymous_L6893F |
| 20989 | T | A | 6.32E-05 | | | | | ORF1ab | nonsynonymous_L6909M |
| 21686 | G | C | 6.32E-05 | | | | | S | nonsynonymous_V42L |
| 21699 | C | T | 6.32E-05 | | | | | S | nonsynonymous_S46L |
| 21700 | A | T | 6.32E-05 | | | | | S | synonymous_S46S |
| 21746 | G | T | 6.32E-05 | | | | | S | nonsynonymous_V62F |

| | | | | | | | | | |
|---|---|---|---|---|---|---|---|---|---|
| 21753 | G | T | 6.32E-05 | | | | | S | nonsynonymous_W64L |
| 21941 | G | T | 6.32E-05 | | | | | S | nonsynonymous_V127F |
| 22019 | A | G | 6.32E-05 | | | | | S | nonsynonymous_M153V |
| 22030 | G | T | 6.32E-05 | | | | | S | nonsynonymous_E156D |
| 22047 | G | T | 6.32E-05 | | | | | S | nonsynonymous_S162I |
| 22075 | C | T | 6.32E-05 | | | | | S | synonymous_V171V |
| 22097 | C | T | 6.32E-05 | | | | | S | nonsynonymous_L179F |
| 22108 | A | C | 6.32E-05 | | | | | S | nonsynonymous_K182N |
| 22111 | G | T | 6.32E-05 | | | | | S | nonsynonymous_Q183H |
| 22131 | G | A | 6.32E-05 | | | | | S | nonsynonymous_R190K |
| 22171 | A | G | 6.32E-05 | | | | | S | nonsynonymous_I203M |
| 22179 | A | G | 6.32E-05 | | | | | S | nonsynonymous_K206R |
| 22180 | G | A | 6.32E-05 | | | | | S | synonymous_K206K |
| 22199 | G | T | 6.32E-05 | | | | | S | nonsynonymous_V213L |
| 22214 | C | G | 6.32E-05 | | | | | S | nonsynonymous_Q218E |
| 22215 | A | T | 6.32E-05 | | | | | S | nonsynonymous_Q218L |
| 22234 | A | G | 6.32E-05 | | | | | S | synonymous_E224E |
| 22244 | G | T | 6.32E-05 | T | 6.32E-05 | | | S | nonsynonymous_D228Y; nonsynonymous_D228Y |
| 22254 | T | C | 6.32E-05 | | | | | S | nonsynonymous_I231T |
| 22255 | A | C | 6.32E-05 | | | | | S | synonymous_I231I |
| 22256 | G | T | 6.32E-05 | | | | | S | nonsynonymous_G232C |
| 22258 | T | A | 6.32E-05 | A | 6.32E-05 | | | S | synonymous_G232G; synonymous_G232G |
| 22260 | T | G | 6.32E-05 | | | | | S | nonsynonymous_I233S |
| 22261 | T | A | 6.32E-05 | | | | | S | synonymous_I233I |
| 22278 | A | G | 6.32E-05 | | | | | S | nonsynonymous_Q239R |
| 22280 | A | C | 6.32E-05 | | | | | S | nonsynonymous_T240P |
| 22300 | A | C | 6.32E-05 | | | | | S | nonsynonymous_R246S |
| 22302 | G | A | 6.32E-05 | | | | | S | nonsynonymous_S247N |

| | | | | | | | | | |
|---|---|---|---|---|---|---|---|---|---|
| 22305 | A | C | 6.32E-05 | | | | | S | nonsynonymous_Y248S |
| 22312 | T | C | 6.32E-05 | | | | | S | synonymous_T250T |
| 22314 | C | T | 6.32E-05 | | | | | S | nonsynonymous_P251L |
| 22318 | T | C | 6.32E-05 | | | | | S | synonymous_G252G |
| 22324 | T | C | 6.32E-05 | | | | | S | synonymous_S254S |
| 22570 | C | T | 6.32E-05 | | | | | S | synonymous_C336C |
| 22571 | C | T | 6.32E-05 | | | | | S | nonsynonymous_P337S |
| 22583 | G | A | 6.32E-05 | | | | | S | nonsynonymous_V341I |
| 22591 | C | T | 6.32E-05 | | | | | S | synonymous_N343N |
| 22594 | C | T | 6.32E-05 | | | | | S | synonymous_A344A |
| 22652 | G | T | 6.32E-05 | | | | | S | nonsynonymous_D364Y |
| 22672 | T | C | 6.32E-05 | | | | | S | synonymous_N370N |
| 22707 | T | A | 6.32E-05 | | | | | S | nonsynonymous_V382E |
| 22715 | A | G | 6.32E-05 | | | | | S | nonsynonymous_T385A |
| 22725 | A | C | 6.32E-05 | | | | | S | nonsynonymous_N388T |
| 22726 | T | C | 6.32E-05 | | | | | S | synonymous_N388N |
| 22747 | C | T | 6.32E-05 | | | | | S | synonymous_V395V |
| 22784 | A | T | 6.32E-05 | | | | | S | stopgain_R408X |
| 22803 | A | G | 6.32E-05 | G | 6.32E-05 | | | S | nonsynonymous_Q414R; nonsynonymous_Q414R |
| 23280 | C | T | 6.32E-05 | | | | | S | nonsynonymous_T573I |
| 23429 | G | T | 6.32E-05 | | | | | S | nonsynonymous_A623S |
| 23551 | C | T | 6.32E-05 | | | | | S | synonymous_D663D |
| 23600 | T | C | 6.32E-05 | | | | | S | nonsynonymous_S680P |
| 23601 | C | T | 6.32E-05 | | | | | S | nonsynonymous_S680F |
| 23607 | G | A | 6.32E-05 | | | | | S | nonsynonymous_R682Q |
| 23608 | G | T | 6.32E-05 | | | | | S | synonymous_R682R |
| 23614 | A | G | 6.32E-05 | | | | | S | synonymous_A684A |
| 23618 | A | G | 6.32E-05 | | | | | S | nonsynonymous_S686G |
| 23621 | G | A | 6.32E-05 | | | | | S | nonsynonymous_V687I |

| | | | | | | | | | |
|---|---|---|---|---|---|---|---|---|---|
| 23624 | G | T | 6.32E-05 | | | | | S | nonsynonymous_A688S |
| 23672 | T | C | 6.32E-05 | | | | | S | nonsynonymous_S704P |
| 23816 | C | T | 6.32E-05 | | | | | S | nonsynonymous_L752F |
| 23905 | T | C | 6.32E-05 | | | | | S | synonymous_V781V |
| 23922 | A | G | 6.32E-05 | | | | | S | nonsynonymous_Q787R |
| 23926 | T | C | 6.32E-05 | | | | | S | synonymous_I788I |
| 23927 | T | G | 6.32E-05 | | | | | S | nonsynonymous_Y789D |
| 23936 | C | A | 6.32E-05 | | | | | S | nonsynonymous_P792T |
| 23937 | C | T | 6.32E-05 | | | | | S | nonsynonymous_P792L |
| 23942 | A | T | 6.32E-05 | | | | | S | nonsynonymous_I794F |
| 23984 | G | A | 6.32E-05 | | | | | S | nonsynonymous_D808N |
| 23985 | A | G | 6.32E-05 | | | | | S | nonsynonymous_D808G |
| 23996 | C | T | 6.32E-05 | | | | | S | nonsynonymous_P812S |
| 24040 | G | A | 6.32E-05 | A | 6.32E-05 | | | S | synonymous_V826V; synonymous_V826V |
| 24106 | C | T | 6.32E-05 | | | | | S | synonymous_D848D |
| 24110 | A | T | 6.32E-05 | | | | | S | nonsynonymous_I850F |
| 24123 | A | G | 6.32E-05 | | | | | S | nonsynonymous_K854R |
| 24194 | T | C | 6.32E-05 | | | | | S | synonymous_L878L |
| 24198 | C | T | 6.32E-05 | | | | | S | nonsynonymous_A879V |
| 24205 | A | T | 6.32E-05 | | | | | S | synonymous_T881T |
| 24208 | C | T | 6.32E-05 | | | | | S | synonymous_I882I |
| 24214 | T | C | 6.32E-05 | | | | | S | synonymous_S884S |
| 24221 | A | G | 6.32E-05 | | | | | S | nonsynonymous_T887A |
| 24227 | G | A | 6.32E-05 | | | | | S | nonsynonymous_G889S |
| 24238 | T | G | 6.32E-05 | | | | | S | synonymous_A892A |
| 24241 | A | T | 6.32E-05 | | | | | S | synonymous_A893A |
| 24253 | A | G | 6.32E-05 | | | | | S | synonymous_P897P |
| 24286 | T | G | 6.32E-05 | | | | | S | synonymous_G908G |
| 24310 | C | T | 6.32E-05 | | | | | S | synonymous_L916L |

| | | | | | | | | | |
|---|---|---|---|---|---|---|---|---|---|
| 24311 | T | C | 6.32E-05 | | | | | S | nonsynonymous_Y917H |
| 24319 | C | T | 6.32E-05 | | | | | S | synonymous_N919N |
| 24332 | G | T | 6.32E-05 | | | | | S | nonsynonymous_A924S |
| 24340 | A | T | 6.32E-05 | | | | | S | nonsynonymous_Q926H |
| 24351 | C | T | 6.32E-05 | | | | | S | nonsynonymous_A930V |
| 24357 | G | A | 6.32E-05 | | | | | S | nonsynonymous_G932D |
| 24358 | C | A | 6.32E-05 | | | | | S | synonymous_G932G |
| 24367 | A | G | 6.32E-05 | G | 6.32E-05 | | | S | synonymous_Q935Q; synonymous_Q935Q |
| 24372 | C | T | 6.32E-05 | | | | | S | nonsynonymous_S937L |
| 24373 | A | G | 6.32E-05 | | | | | S | synonymous_S937S |
| 24376 | T | A | 6.32E-05 | A | 6.32E-05 | | | S | synonymous_L938L; synonymous_L938L |
| 24388 | A | T | 6.32E-05 | | | | | S | synonymous_A942A |
| 24444 | C | T | 6.32E-05 | | | | | S | nonsynonymous_T961M |
| 24450 | T | A | 6.32E-05 | | | | | S | nonsynonymous_V963D |
| 24454 | A | G | 6.32E-05 | | | | | S | synonymous_K964K |
| 24463 | C | A | 6.32E-05 | | | | | S | nonsynonymous_S967R |
| 24466 | C | T | 6.32E-05 | | | | | S | synonymous_S968S |
| 24471 | T | C | 6.32E-05 | | | | | S | nonsynonymous_F970S |
| 24502 | C | T | 6.32E-05 | | | | | S | synonymous_I980I |
| 24503 | C | T | 6.32E-05 | | | | | S | nonsynonymous_L981F |
| 24529 | T | A | 6.32E-05 | | | | | S | synonymous_A989A |
| 24565 | T | G | 6.32E-05 | | | | | S | synonymous_L1001L |
| 24624 | C | T | 6.32E-05 | | | | | S | nonsynonymous_S1021F |
| 24626 | G | T | 6.32E-05 | | | | | S | nonsynonymous_A1022S |
| 24632 | C | T | 6.32E-05 | | | | | S | nonsynonymous_L1024F |
| 24685 | T | C | 6.32E-05 | | | | | S | synonymous_D1041D |
| 24712 | G | A | 6.32E-05 | | | | | S | nonsynonymous_M1050I |
| 24715 | C | T | 6.32E-05 | | | | | S | synonymous_S1051S |

| | | | | | | | | | |
|---|---|---|---|---|---|---|---|---|---|
| 24748 | C | T | 6.32E-05 | | | | | S | synonymous_F1062F |
| 25010 | G | A | 6.32E-05 | | | | | S | nonsynonymous_E1150K |
| 25031 | A | T | 6.32E-05 | | | | | S | stopgain_K1157X |
| 25037 | C | T | 6.32E-05 | | | | | S | nonsynonymous_H1159Y |
| 25048 | A | T | 6.32E-05 | | | | | S | synonymous_P1162P |
| 25378 | C | T | 6.32E-05 | | | | | S | synonymous_Y1272Y |
| 25493 | C | T | 6.32E-05 | | | | | ORF3a | nonsynonymous_T34M |
| 25496 | T | C | 6.32E-05 | | | | | ORF3a | nonsynonymous_I35T |
| 25509 | C | T | 6.32E-05 | | | | | ORF3a | synonymous_A39A |
| 25514 | T | C | 6.32E-05 | | | | | ORF3a | nonsynonymous_L41P |
| 25516 | C | T | 6.32E-05 | | | | | ORF3a | nonsynonymous_P42S |
| 25540 | G | A | 6.32E-05 | | | | | ORF3a | nonsynonymous_V50I |
| 25546 | C | A | 6.32E-05 | A | 6.32E-05 | | | ORF3a | nonsynonymous_L52I; nonsynonymous_L52I |
| 25554 | T | G | 6.32E-05 | | | | | ORF3a | synonymous_A54A |
| 25566 | C | T | 6.32E-05 | | | | | ORF3a | synonymous_S58S |
| 25567 | G | A | 6.32E-05 | A | 6.32E-05 | | | ORF3a | nonsynonymous_A59T; nonsynonymous_A59T |
| 25577 | T | C | 6.32E-05 | | | | | ORF3a | nonsynonymous_I62T |
| 25578 | C | T | 6.32E-05 | | | | | ORF3a | synonymous_I62I |
| 25580 | T | C | 6.32E-05 | | | | | ORF3a | nonsynonymous_I63T |
| 25589 | A | C | 6.32E-05 | | | | | ORF3a | nonsynonymous_K66T |
| 25594 | A | C | 6.32E-05 | | | | | ORF3a | synonymous_R68R |
| 25956 | T | C | 6.32E-05 | | | | | ORF3a | synonymous_G188G |
| 25957 | T | C | 6.32E-05 | | | | | ORF3a | nonsynonymous_Y189H |
| 26019 | A | G | 6.32E-05 | | | | | ORF3a | synonymous_S209S |
| 26199 | G | T | 6.32E-05 | | | | | ORF3a | synonymous_T269T |
| 26232 | T | C | 6.32E-05 | | | | | intergenic | |
| 26263 | G | A | 6.32E-05 | | | | | E | nonsynonymous_E7K |
| 26509 | C | T | 6.32E-05 | | | | | intergenic | |

| | | | | | | | | | |
|---|---|---|---|---|---|---|---|---|---|
| 26513 | A | G | 6.32E-05 | | | | | intergenic | |
| 26536 | A | G | 6.32E-05 | | | | | M | nonsynonymous_N5S |
| 26542 | C | T | 6.32E-05 | | | | | M | nonsynonymous_T7I |
| 26596 | G | T | 6.32E-05 | | | | | M | nonsynonymous_G25V |
| 26634 | G | T | 6.32E-05 | | | | | M | nonsynonymous_A38S |
| 26780 | T | C | 6.32E-05 | | | | | M | synonymous_C86C |
| 26818 | C | T | 6.32E-05 | | | | | M | nonsynonymous_S99F |
| 26861 | T | C | 6.32E-05 | | | | | M | synonymous_N113N |
| 26921 | T | C | 6.32E-05 | | | | | M | synonymous_L133L |
| 26922 | C | T | 6.32E-05 | | | | | M | synonymous_L134L |
| 26929 | G | A | 6.32E-05 | | | | | M | nonsynonymous_S136N |
| 26934 | C | A | 6.32E-05 | | | | | M | nonsynonymous_L138I |
| 26942 | C | T | 6.32E-05 | | | | | M | synonymous_I140I |
| 26947 | C | T | 6.32E-05 | | | | | M | nonsynonymous_A142V |
| 26966 | T | A | 6.32E-05 | | | | | M | nonsynonymous_H148Q |
| 26967 | C | T | 6.32E-05 | | | | | M | nonsynonymous_L149F |
| 26981 | A | G | 6.32E-05 | | | | | M | synonymous_G153G |
| 26994 | C | T | 6.32E-05 | | | | | M | nonsynonymous_R158C |
| 26996 | C | T | 6.32E-05 | | | | | M | synonymous_R158R |
| 27007 | A | G | 6.32E-05 | | | | | M | nonsynonymous_K162R |
| 27014 | G | T | 6.32E-05 | | | | | M | synonymous_L164L |
| 27023 | A | G | 6.32E-05 | | | | | M | synonymous_E167E |
| 27032 | T | C | 6.32E-05 | | | | | M | synonymous_V170V |
| 27176 | T | G | 6.32E-05 | | | | | M | synonymous_A218A |
| 27198 | A | G | 6.32E-05 | | | | | intergenic | |
| 27372 | A | C | 6.32E-05 | | | | | ORF6 | synonymous_P57P |
| 27373 | A | T | 6.32E-05 | | | | | ORF6 | nonsynonymous_M58L |
| 27374 | T | C | 6.32E-05 | | | | | ORF6 | nonsynonymous_M58T |
| 27376 | G | T | 6.32E-05 | | | | | ORF6 | stopgain_E59X |
| 27378 | G | T | 6.32E-05 | | | | | ORF6 | nonsynonymous_E59D |

| | | | | | | | | | |
|---|---|---|---|---|---|---|---|---|---|
| 27380 | T | C | 6.32E-05 | | | | | ORF6 | nonsynonymous_I60T |
| 27389 | C | T | 6.32E-05 | | | | | intergenic | |
| 27392 | A | G | 6.32E-05 | | | | | intergenic | |
| 27397 | A | G | 6.32E-05 | | | | | ORF7a | nonsynonymous_K2E |
| 27401 | T | C | 6.32E-05 | | | | | ORF7a | nonsynonymous_I3T |
| 27402 | T | C | 6.32E-05 | | | | | ORF7a | synonymous_I3I |
| 27444 | T | C | 6.32E-05 | | | | | ORF7a | synonymous_L17L |
| 27447 | T | C | 6.32E-05 | | | | | ORF7a | synonymous_Y18Y |
| 27809 | A | T | 6.32E-05 | T | 6.32E-05 | | | ORF7b | synonymous_L18L; synonymous_L18L |
| 28307 | G | A | 6.32E-05 | A | 6.32E-05 | | | N | nonsynonymous_A12T; nonsynonymous_A12T |
| 28361 | G | A | 6.32E-05 | A | 6.32E-05 | | | N | nonsynonymous_G30R; nonsynonymous_G30R |
| 28369 | C | T | 6.32E-05 | | | | | N | synonymous_R32R |
| 28381 | A | T | 6.32E-05 | | | | | N | synonymous_R36R |
| 28416 | A | T | 6.32E-05 | | | | | N | nonsynonymous_N48I |
| 28423 | G | A | 6.32E-05 | | | | | N | synonymous_A50A |
| 28451 | G | C | 6.32E-05 | | | | | N | nonsynonymous_G60R |
| 28455 | A | G | 6.32E-05 | | | | | N | nonsynonymous_K61R |
| 28631 | G | A | 6.32E-05 | | | | | N | nonsynonymous_G120R |
| 28693 | T | G | 6.32E-05 | G | 6.32E-05 | | | N | nonsynonymous_N140K; nonsynonymous_N140K |
| 28720 | C | T | 6.32E-05 | | | | | N | synonymous_R149R |
| 29334 | A | G | 6.32E-05 | | | | | N | nonsynonymous_N354S |
| 29348 | G | T | 6.32E-05 | | | | | N | nonsynonymous_A359S |
| 332 | G | T | 6.32E-05 | | | | | ORF1ab | nonsynonymous_V23F |
| 3828 | C | T | 6.32E-05 | | | | | ORF1ab | nonsynonymous_S1188L |
| 385 | A | T | 6.32E-05 | T | 6.32E-05 | | | ORF1ab | synonymous_S40S; synonymous_S40S |

| | | | | | | | | | |
|---|---|---|---|---|---|---|---|---|---|
| 388 | G | A | 6.32E-05 | A | 6.32E-05 | | | ORF1ab | synonymous_E41E; synonymous_E41E |
| 6274 | T | C | 6.32E-05 | | | | | ORF1ab | synonymous_Y2003Y |
| 6459 | A | G | 6.32E-05 | | | | | ORF1ab | nonsynonymous_N2065S |
| 6474 | A | G | 6.32E-05 | | | | | ORF1ab | nonsynonymous_E2070G |
| 6664 | C | A | 6.32E-05 | A | 6.32E-05 | | | ORF1ab | synonymous_V2133V; synonymous_V2133V |
| 6872 | G | A | 6.32E-05 | | | | | ORF1ab | nonsynonymous_V2203I |
| 6880 | T | C | 6.32E-05 | | | | | ORF1ab | synonymous_S2205S |
| 6954 | T | C | 6.32E-05 | | | | | ORF1ab | nonsynonymous_I2230T |
| 6956 | A | C | 6.32E-05 | | | | | ORF1ab | nonsynonymous_I2231L |
| 6964 | T | C | 6.32E-05 | | | | | ORF1ab | synonymous_F2233F |
| 6997 | C | T | 6.32E-05 | | | | | ORF1ab | synonymous_I2244I |
| 7000 | C | T | 6.32E-05 | | | | | ORF1ab | synonymous_Y2245Y |
| 7007 | G | A | 6.32E-05 | | | | | ORF1ab | nonsynonymous_A2248T |
| 7022 | T | C | 6.32E-05 | | | | | ORF1ab | synonymous_L2253L |
| 7025 | A | G | 6.32E-05 | | | | | ORF1ab | nonsynonymous_M2254V |
| 7034 | T | A | 6.32E-05 | | | | | ORF1ab | nonsynonymous_L2257I |
| 7037 | G | T | 6.32E-05 | | | | | ORF1ab | nonsynonymous_G2258C |
| 7040 | A | G | 6.32E-05 | | | | | ORF1ab | nonsynonymous_M2259V |
| 7060 | T | G | 6.32E-05 | | | | | ORF1ab | synonymous_G2265G |
| 7063 | C | T | 6.32E-05 | | | | | ORF1ab | synonymous_Y2266Y |
| 7289 | G | T | 6.32E-05 | | | | | ORF1ab | stopgain_G2342X |
| 7313 | T | G | 6.32E-05 | | | | | ORF1ab | nonsynonymous_F2350V |
| 8647 | T | C | 6.32E-05 | | | | | ORF1ab | synonymous_H2794H |
| 8649 | T | C | 6.32E-05 | | | | | ORF1ab | nonsynonymous_V2795A |
| 8671 | T | G | 6.32E-05 | | | | | ORF1ab | nonsynonymous_F2802L |
| 8717 | C | T | 6.32E-05 | | | | | ORF1ab | nonsynonymous_R2818C |
| 8763 | A | G | 6.32E-05 | | | | | ORF1ab | nonsynonymous_D2833G |
| 8768 | G | A | 6.32E-05 | | | | | ORF1ab | nonsynonymous_D2835N |

| | | | | | | | | | |
|---|---|---|---|---|---|---|---|---|---|
| 8812 | A | T | 6.32E-05 | | | | | ORF1ab | nonsynonymous_K2849N |
| 8831 | G | T | 6.32E-05 | | | | | ORF1ab | nonsynonymous_A2856S |
| 8832 | C | T | 6.32E-05 | | | | | ORF1ab | nonsynonymous_A2856V |
| 8834 | G | A | 6.32E-05 | | | | | ORF1ab | nonsynonymous_V2857I |
| 8836 | C | T | 6.32E-05 | | | | | ORF1ab | synonymous_V2857V |
| 8841 | C | T | 6.32E-05 | | | | | ORF1ab | nonsynonymous_T2859I |
| 8844 | G | T | 6.32E-05 | | | | | ORF1ab | nonsynonymous_R2860I |
| 8852 | G | A | 6.32E-05 | | | | | ORF1ab | nonsynonymous_G2863S |
| 8861 | G | A | 6.32E-05 | A | 6.32E-05 | | | ORF1ab | nonsynonymous_V2866M; nonsynonymous_V2866M |
| 8862 | T | A | 6.32E-05 | | | | | ORF1ab | nonsynonymous_V2866E |
| 8863 | G | T | 6.32E-05 | | | | | ORF1ab | synonymous_V2866V |
| 8867 | G | A | 6.32E-05 | | | | | ORF1ab | nonsynonymous_G2868S |
| 8874 | C | T | 6.32E-05 | | | | | ORF1ab | nonsynonymous_P2870L |
| 8876 | G | A | 6.32E-05 | | | | | ORF1ab | nonsynonymous_G2871S |
| 8880 | C | T | 6.32E-05 | | | | | ORF1ab | nonsynonymous_T2872M |
| 8891 | A | T | 6.32E-05 | T | 6.32E-05 | | | ORF1ab | nonsynonymous_T2876S; nonsynonymous_T2876S |
| 8911 | G | T | 6.32E-05 | | | | | ORF1ab | nonsynonymous_L2882F |
| 9249 | G | A | 6.32E-05 | | | | | ORF1ab | nonsynonymous_G2995D |
| 9274 | A | G | 6.32E-05 | | | | | ORF1ab | synonymous_R3003R |
| 9314 | G | T | 6.32E-05 | | | | | ORF1ab | nonsynonymous_V3017F |
| 9317 | T | G | 6.32E-05 | | | | | ORF1ab | nonsynonymous_F3018V |
| 9318 | T | A | 6.32E-05 | | | | | ORF1ab | nonsynonymous_F3018Y |
| 9320 | T | A | 6.32E-05 | | | | | ORF1ab | nonsynonymous_C3019S |
| 9348 | C | T | 6.32E-05 | T | 6.32E-05 | | | ORF1ab | nonsynonymous_T3028I; nonsynonymous_T3028I |
| 9349 | T | C | 6.32E-05 | | | | | ORF1ab | synonymous_T3028T |
| 9357 | T | A | 6.32E-05 | | | | | ORF1ab | nonsynonymous_F3031Y |
| 9372 | A | G | 6.32E-05 | | | | | ORF1ab | nonsynonymous_Q3036R |

| | | | | | | | | | |
|---|---|---|---|---|---|---|---|---|---|
| 9383 | G | C | 6.32E-05 | | | | | ORF1ab | nonsynonymous_A3040P |
| 9389 | G | T | 6.32E-05 | | | | | ORF1ab | nonsynonymous_D3042Y |
| 9391 | C | T | 6.32E-05 | | | | | ORF1ab | synonymous_D3042D |
| 9398 | G | A | 6.32E-05 | | | | | ORF1ab | nonsynonymous_A3045T |
| 9405 | T | G | 6.32E-05 | | | | | ORF1ab | nonsynonymous_I3047R |
| 9408 | T | C | 6.32E-05 | | | | | ORF1ab | nonsynonymous_V3048A |
| 9409 | A | G | 6.32E-05 | | | | | ORF1ab | synonymous_V3048V |
| 9412 | T | C | 6.32E-05 | | | | | ORF1ab | synonymous_A3049A |
| 9414 | G | C | 6.32E-05 | | | | | ORF1ab | nonsynonymous_G3050A |
| 9448 | C | T | 6.32E-05 | | | | | ORF1ab | synonymous_A3061A |
| 9463 | G | T | 6.32E-05 | | | | | ORF1ab | nonsynonymous_R3066S |
| 10039 | C | T | 6.32E-05 | | | | | ORF1ab | synonymous_T3258T |
| 10748 | C | T | 6.32E-05 | | | | | ORF1ab | nonsynonymous_L3495F |
| 10785 | A | G | 6.32E-05 | | | | | ORF1ab | nonsynonymous_Q3507R |
| 10805 | G | A | 6.32E-05 | | | | | ORF1ab | nonsynonymous_G3514R |
| 10825 | T | A | 6.32E-05 | | | | | ORF1ab | synonymous_T3520T |
| 10831 | T | C | 6.32E-05 | | | | | ORF1ab | synonymous_I3522I |
| 1087 | A | G | 6.32E-05 | | | | | ORF1ab | synonymous_V274V |
| 10881 | T | C | 6.32E-05 | | | | | ORF1ab | nonsynonymous_M3539T |
| 10888 | A | G | 6.32E-05 | | | | | ORF1ab | synonymous_G3541G |
| 10933 | T | C | 6.32E-05 | | | | | ORF1ab | synonymous_P3556P |
| 10952 | T | A | 6.32E-05 | | | | | ORF1ab | nonsynonymous_C3563S |
| 10954 | C | T | 6.32E-05 | | | | | ORF1ab | synonymous_C3563C |
| 10956 | C | T | 6.32E-05 | | | | | ORF1ab | nonsynonymous_S3564L |
| 10983 | A | G | 6.32E-05 | | | | | ORF1ab | nonsynonymous_K3573R |
| 10985 | A | G | 6.32E-05 | | | | | ORF1ab | nonsynonymous_R3574G |
| 10987 | A | G | 6.32E-05 | | | | | ORF1ab | synonymous_R3574R |
| 10990 | A | G | 6.32E-05 | | | | | ORF1ab | synonymous_T3575T |
| 10994 | A | G | 6.32E-05 | | | | | ORF1ab | nonsynonymous_K3577E |
| 10995 | A | G | 6.32E-05 | | | | | ORF1ab | nonsynonymous_K3577R |

| | | | | | | | | | |
|---|---|---|---|---|---|---|---|---|---|
| 11005 | C | T | 6.32E-05 | | | | | ORF1ab | synonymous_H3580H |
| 11268 | A | G | 6.32E-05 | | | | | ORF1ab | nonsynonymous_D3668G |
| 11566 | T | A | 6.32E-05 | | | | | ORF1ab | synonymous_P3767P |
| 12149 | G | A | 6.32E-05 | | | | | ORF1ab | nonsynonymous_E3962K |
| 12167 | G | A | 6.32E-05 | | | | | ORF1ab | nonsynonymous_V3968I |
| 12355 | G | C | 6.32E-05 | | | | | ORF1ab | nonsynonymous_Q4030H |
| 12382 | G | T | 6.32E-05 | | | | | ORF1ab | nonsynonymous_K4039N |
| 13147 | T | C | 6.32E-05 | | | | | ORF1ab | synonymous_C4294C |
| 13157 | T | C | 6.32E-05 | | | | | ORF1ab | synonymous_L4298L |
| 13159 | G | A | 6.32E-05 | | | | | ORF1ab | synonymous_L4298L |
| 13170 | C | T | 6.32E-05 | | | | | ORF1ab | nonsynonymous_T4302I |
| 13171 | T | C | 6.32E-05 | | | | | ORF1ab | synonymous_T4302T |
| 13183 | G | A | 6.32E-05 | | | | | ORF1ab | synonymous_Q4306Q |
| 13198 | A | G | 6.32E-05 | | | | | ORF1ab | synonymous_T4311T |
| 13205 | G | T | 6.32E-05 | | | | | ORF1ab | nonsynonymous_A4314S |
| 13212 | T | A | 6.32E-05 | A | 6.32E-05 | | | ORF1ab | nonsynonymous_M4316K; nonsynonymous_M4316K |
| 13226 | T | C | 6.32E-05 | | | | | ORF1ab | nonsynonymous_F4321L |
| 13255 | C | T | 6.32E-05 | | | | | ORF1ab | synonymous_C4330C |
| 13264 | C | T | 6.32E-05 | | | | | ORF1ab | synonymous_H4333H |
| 13269 | A | G | 6.32E-05 | | | | | ORF1ab | nonsynonymous_D4335G |
| 13275 | C | T | 6.32E-05 | | | | | ORF1ab | nonsynonymous_P4337L |
| 13286 | G | T | 6.32E-05 | | | | | ORF1ab | stopgain_G4341X |
| 13297 | C | A | 6.32E-05 | A | 6.32E-05 | | | ORF1ab | nonsynonymous_D4344E; nonsynonymous_D4344E |
| 13309 | G | T | 6.32E-05 | | | | | ORF1ab | nonsynonymous_K4348N |
| 13329 | C | T | 6.32E-05 | | | | | ORF1ab | nonsynonymous_T4355I |
| 13358 | C | T | 6.32E-05 | | | | | ORF1ab | nonsynonymous_L4365F |
| 13376 | A | G | 6.32E-05 | | | | | ORF1ab | nonsynonymous_T4371A |
| 13377 | C | A | 6.32E-05 | | | | | ORF1ab | nonsynonymous_T4371N |

| | | | | | | | | | |
|---|---|---|---|---|---|---|---|---|---|
| 13668 | T | C | 6.32E-05 | | | | | ORF1ab | synonymous_T4468T |
| 13672 | T | G | 6.32E-05 | G | 6.32E-05 | | | ORF1ab | nonsynonymous_S4470A; nonsynonymous_S4470A |
| 13679 | A | G | 6.32E-05 | | | | | ORF1ab | nonsynonymous_Y4472C |
| 1367 | A | C | 6.32E-05 | | | | | ORF1ab | nonsynonymous_I368L |
| 13683 | A | G | 6.32E-05 | | | | | ORF1ab | synonymous_Q4473Q |
| 13703 | A | G | 6.32E-05 | | | | | ORF1ab | nonsynonymous_N4480S |
| 13711 | A | G | 6.32E-05 | | | | | ORF1ab | nonsynonymous_K4483E |
| 13716 | T | C | 6.32E-05 | | | | | ORF1ab | synonymous_D4484D |
| 13721 | C | T | 6.32E-05 | | | | | ORF1ab | nonsynonymous_P4486L |
| 13728 | T | C | 6.32E-05 | | | | | ORF1ab | synonymous_V4488V |
| 13743 | C | T | 6.32E-05 | | | | | ORF1ab | synonymous_F4493F |
| 13755 | A | G | 6.32E-05 | | | | | ORF1ab | synonymous_R4497R |
| 13758 | A | G | 6.32E-05 | | | | | ORF1ab | nonsynonymous_I4498M |
| 13771 | G | T | 6.32E-05 | | | | | ORF1ab | nonsynonymous_V4503L |
| 13782 | A | G | 6.32E-05 | | | | | ORF1ab | nonsynonymous_I4506M |
| 13821 | C | T | 6.32E-05 | | | | | ORF1ab | synonymous_L4519L |
| 13852 | A | G | 6.32E-05 | | | | | ORF1ab | nonsynonymous_N4530D |
| 13857 | T | C | 6.32E-05 | | | | | ORF1ab | synonymous_C4531C |
| 13859 | A | G | 6.32E-05 | G | 6.32E-05 | | | ORF1ab | nonsynonymous_D4532G; nonsynonymous_D4532G |
| 13863 | A | C | 6.32E-05 | | | | | ORF1ab | synonymous_T4533T |
| 13868 | A | G | 6.32E-05 | | | | | ORF1ab | nonsynonymous_K4535R |
| 13887 | C | T | 6.32E-05 | | | | | ORF1ab | synonymous_Y4541Y |
| 13900 | G | T | 6.32E-05 | | | | | ORF1ab | nonsynonymous_D4546Y |
| 13901 | A | G | 6.32E-05 | | | | | ORF1ab | nonsynonymous_D4546G |
| 13922 | A | T | 6.32E-05 | | | | | ORF1ab | nonsynonymous_D4553V |
| 13928 | A | G | 6.32E-05 | | | | | ORF1ab | nonsynonymous_Y4555C |
| 13932 | T | C | 6.32E-05 | | | | | ORF1ab | synonymous_D4556D |
| 13950 | T | C | 6.32E-05 | | | | | ORF1ab | synonymous_D4562D |

| | | | | | | | | | |
|---|---|---|---|---|---|---|---|---|---|
| 13951 | A | T | 6.32E-05 | | | | | ORF1ab | nonsynonymous_I4563L |
| 13959 | C | T | 6.32E-05 | | | | | ORF1ab | synonymous_R4565R |
| 13975 | G | A | 6.32E-05 | A | 6.32E-05 | | | ORF1ab | nonsynonymous_G4571S;<br>nonsynonymous_G4571S |
| 13982 | G | T | 6.32E-05 | | | | | ORF1ab | nonsynonymous_R4573L |
| 13990 | C | A | 6.32E-05 | | | | | ORF1ab | nonsynonymous_Q4576K |
| 13992 | A | T | 6.32E-05 | | | | | ORF1ab | nonsynonymous_Q4576H |
| 13993 | G | A | 6.32E-05 | A | 6.32E-05 | | | ORF1ab | nonsynonymous_A4577T;<br>nonsynonymous_A4577T |
| 13995 | T | G | 6.32E-05 | | | | | ORF1ab | synonymous_A4577A |
| 13997 | T | A | 6.32E-05 | | | | | ORF1ab | stopgain_L4578X |
| 13999 | T | C | 6.32E-05 | | | | | ORF1ab | synonymous_L4579L |
| 14001 | A | G | 6.32E-05 | | | | | ORF1ab | synonymous_L4579L |
| 14004 | A | C | 6.32E-05 | | | | | ORF1ab | nonsynonymous_K4580N |
| 14005 | A | G | 6.32E-05 | | | | | ORF1ab | nonsynonymous_T4581A |
| 14008 | G | A | 6.32E-05 | | | | | ORF1ab | nonsynonymous_V4582I |
| 14016 | C | T | 6.32E-05 | | | | | ORF1ab | synonymous_F4584F |
| 14028 | G | T | 6.32E-05 | | | | | ORF1ab | nonsynonymous_M4588I |
| 14034 | T | C | 6.32E-05 | | | | | ORF1ab | synonymous_N4590N |
| 14041 | A | C | 6.32E-05 | | | | | ORF1ab | nonsynonymous_I4593L |
| 14055 | G | T | 6.32E-05 | | | | | ORF1ab | synonymous_L4597L |
| 14097 | C | T | 6.32E-05 | | | | | ORF1ab | synonymous_F4611F |
| 14103 | T | C | 6.32E-05 | | | | | ORF1ab | synonymous_D4613D |
| 14109 | A | G | 6.32E-05 | | | | | ORF1ab | nonsynonymous_I4615M |
| 14117 | C | T | 6.32E-05 | | | | | ORF1ab | nonsynonymous_T4618M |
| 14121 | A | G | 6.32E-05 | | | | | ORF1ab | synonymous_P4619P |
| 14125 | A | G | 6.32E-05 | | | | | ORF1ab | nonsynonymous_S4621G |
| 14141 | T | C | 6.32E-05 | | | | | ORF1ab | nonsynonymous_V4626A |
| 14145 | T | C | 6.32E-05 | | | | | ORF1ab | synonymous_D4627D |
| 14148 | T | C | 6.32E-05 | | | | | ORF1ab | synonymous_S4628S |

| | | | | | | | | | |
|---|---|---|---|---|---|---|---|---|---|
| 14151 | T | C | 6.32E-05 | | | | | ORF1ab | synonymous_Y4629Y |
| 14172 | A | G | 6.32E-05 | | | | | ORF1ab | nonsynonymous_I4636M |
| 14187 | G | A | 6.32E-05 | | | | | ORF1ab | synonymous_R4641R |
| 14189 | C | T | 6.32E-05 | | | | | ORF1ab | nonsynonymous_A4642V |
| 14191 | T | C | 6.32E-05 | | | | | ORF1ab | synonymous_L4643L |
| 14197 | G | A | 6.32E-05 | A | 6.32E-05 | | | ORF1ab | nonsynonymous_A4645T; nonsynonymous_A4645T |
| 14201 | A | T | 6.32E-05 | | | | | ORF1ab | nonsynonymous_E4646V |
| 14209 | G | A | 6.32E-05 | A | 6.32E-05 | | | ORF1ab | nonsynonymous_V4649I; nonsynonymous_V4649I |
| 14212 | G | A | 6.32E-05 | | | | | ORF1ab | nonsynonymous_D4650N |
| 14229 | G | T | 6.32E-05 | | | | | ORF1ab | nonsynonymous_K4655N |
| 14268 | G | A | 6.32E-05 | | | | | ORF1ab | synonymous_T4668T |
| 14280 | A | G | 6.32E-05 | | | | | ORF1ab | synonymous_L4672L |
| 14286 | C | T | 6.32E-05 | | | | | ORF1ab | synonymous_L4674L |
| 14294 | G | A | 6.32E-05 | | | | | ORF1ab | nonsynonymous_R4677H |
| 14295 | T | C | 6.32E-05 | | | | | ORF1ab | synonymous_R4677R |
| 14310 | G | A | 6.32E-05 | | | | | ORF1ab | stopgain_W4682X |
| 14352 | C | T | 6.32E-05 | | | | | ORF1ab | synonymous_D4696D |
| 14362 | C | T | 6.32E-05 | | | | | ORF1ab | synonymous_L4700L |
| 14383 | G | T | 6.32E-05 | | | | | ORF1ab | nonsynonymous_V4707F |
| 14394 | T | A | 6.32E-05 | A | 6.32E-05 | | | ORF1ab | synonymous_S4710S; synonymous_S4710S |
| 14409 | T | C | 6.32E-05 | | | | | ORF1ab | synonymous_P4715P |
| 14430 | G | A | 6.32E-05 | | | | | ORF1ab | synonymous_V4722V |
| 14436 | A | G | 6.32E-05 | | | | | ORF1ab | synonymous_K4724K |
| 14443 | G | T | 6.32E-05 | | | | | ORF1ab | nonsynonymous_V4727F |
| 14451 | T | G | 6.32E-05 | | | | | ORF1ab | synonymous_G4729G |
| 14538 | T | C | 6.32E-05 | | | | | ORF1ab | synonymous_L4758L |
| 14539 | A | C | 6.32E-05 | | | | | ORF1ab | nonsynonymous_S4759R |

| | | | | | | | | | |
|---|---|---|---|---|---|---|---|---|---|
| 14540 | G | C | 6.32E-05 | | | | | ORF1ab | nonsynonymous_S4759T |
| 14549 | A | G | 6.32E-05 | | | | | ORF1ab | nonsynonymous_E4762G |
| 14597 | A | G | 6.32E-05 | | | | | ORF1ab | nonsynonymous_N4778S |
| 14616 | C | T | 6.32E-05 | | | | | ORF1ab | synonymous_R4784R |
| 14618 | C | A | 6.32E-05 | | | | | ORF1ab | nonsynonymous_T4785N |
| 14619 | T | C | 6.32E-05 | | | | | ORF1ab | synonymous_T4785T |
| 14621 | C | T | 6.32E-05 | | | | | ORF1ab | nonsynonymous_T4786M |
| 14638 | G | T | 6.32E-05 | | | | | ORF1ab | nonsynonymous_A4792S |
| 14649 | C | T | 6.32E-05 | | | | | ORF1ab | synonymous_N4795N |
| 14656 | G | T | 6.32E-05 | | | | | ORF1ab | nonsynonymous_A4798S |
| 14666 | C | T | 6.32E-05 | | | | | ORF1ab | nonsynonymous_T4801I |
| 14675 | C | G | 6.32E-05 | | | | | ORF1ab | nonsynonymous_P4804R |
| 14679 | T | C | 6.32E-05 | | | | | ORF1ab | synonymous_G4805G |
| 14682 | T | G | 6.32E-05 | | | | | ORF1ab | nonsynonymous_N4806K |
| 14691 | A | G | 6.32E-05 | | | | | ORF1ab | synonymous_K4809K |
| 14692 | G | A | 6.32E-05 | | | | | ORF1ab | nonsynonymous_D4810N |
| 14706 | T | C | 6.32E-05 | | | | | ORF1ab | synonymous_F4814F |
| 14710 | G | A | 6.32E-05 | | | | | ORF1ab | nonsynonymous_V4816M |
| 14740 | T | C | 6.32E-05 | | | | | ORF1ab | nonsynonymous_S4826P |
| 14757 | C | T | 6.32E-05 | | | | | ORF1ab | synonymous_H4831H |
| 14760 | C | T | 6.32E-05 | | | | | ORF1ab | synonymous_F4832F |
| 14763 | C | A | 6.32E-05 | | | | | ORF1ab | nonsynonymous_F4833L |
| 14785 | G | A | 6.32E-05 | | | | | ORF1ab | nonsynonymous_A4841T |
| 14788 | A | G | 6.32E-05 | | | | | ORF1ab | nonsynonymous_I4842V |
| 14790 | C | T | 6.32E-05 | | | | | ORF1ab | synonymous_I4842I |
| 14808 | T | A | 6.32E-05 | | | | | ORF1ab | stopgain_Y4848X |
| 14821 | C | T | 6.32E-05 | | | | | ORF1ab | nonsynonymous_P4853S |
| 14823 | A | T | 6.32E-05 | | | | | ORF1ab | synonymous_P4853P |
| 14832 | T | C | 6.32E-05 | | | | | ORF1ab | synonymous_C4856C |
| 14863 | G | A | 6.32E-05 | | | | | ORF1ab | nonsynonymous_V4867I |

| | | | | | | | | | |
|---|---|---|---|---|---|---|---|---|---|
| 14866 | G | T | 6.32E-05 | | | | | ORF1ab | nonsynonymous_V4868F |
| 14882 | A | G | 6.32E-05 | | | | | ORF1ab | nonsynonymous_D4873G |
| 14907 | T | G | 6.32E-05 | | | | | ORF1ab | nonsynonymous_N4881K |
| 14932 | C | T | 6.32E-05 | | | | | ORF1ab | synonymous_L4890L |
| 14996 | T | C | 6.32E-05 | | | | | ORF1ab | nonsynonymous_M4911T |
| 15033 | A | C | 6.32E-05 | | | | | ORF1ab | synonymous_T4923T |
| 15049 | C | T | 6.32E-05 | | | | | ORF1ab | nonsynonymous_P4929S |
| 15051 | T | C | 6.32E-05 | | | | | ORF1ab | synonymous_P4929P |
| 15064 | A | C | 6.32E-05 | | | | | ORF1ab | nonsynonymous_M4934L |
| 15079 | G | A | 6.32E-05 | | | | | ORF1ab | nonsynonymous_A4939T |
| 15089 | C | T | 6.32E-05 | | | | | ORF1ab | nonsynonymous_A4942V |
| 15100 | G | A | 6.32E-05 | | | | | ORF1ab | nonsynonymous_A4946T |
| 15101 | C | A | 6.32E-05 | | | | | ORF1ab | nonsynonymous_A4946D |
| 15102 | T | C | 6.32E-05 | | | | | ORF1ab | synonymous_A4946A |
| 15105 | C | A | 6.32E-05 | A | 6.32E-05 | | | ORF1ab | synonymous_R4947R; synonymous_R4947R |
| 15119 | T | A | 6.32E-05 | | | | | ORF1ab | nonsynonymous_V4952D |
| 15120 | C | A | 6.32E-05 | | | | | ORF1ab | synonymous_V4952V |
| 15126 | C | T | 6.32E-05 | | | | | ORF1ab | synonymous_I4954I |
| 15129 | T | C | 6.32E-05 | | | | | ORF1ab | synonymous_C4955C |
| 15138 | G | T | 6.32E-05 | | | | | ORF1ab | nonsynonymous_M4958I |
| 15141 | C | T | 6.32E-05 | | | | | ORF1ab | synonymous_T4959T |
| 15147 | A | G | 6.32E-05 | | | | | ORF1ab | synonymous_R4961R |
| 15168 | G | A | 6.32E-05 | | | | | ORF1ab | synonymous_L4968L |
| 15177 | A | G | 6.32E-05 | | | | | ORF1ab | nonsynonymous_I4971M |
| 15181 | G | A | 6.32E-05 | A | 6.32E-05 | | | ORF1ab | nonsynonymous_A4973T; nonsynonymous_A4973T |
| 15182 | C | G | 6.32E-05 | | | | | ORF1ab | nonsynonymous_A4973G |
| 15204 | A | G | 6.32E-05 | | | | | ORF1ab | synonymous_V4980V |
| 15205 | A | G | 6.32E-05 | | | | | ORF1ab | nonsynonymous_I4981V |

| | | | | | | | | | |
|---|---|---|---|---|---|---|---|---|---|
| 15207 | T | A | 6.32E-05 | | | | | ORF1ab | synonymous_I4981I |
| 15213 | A | G | 6.32E-05 | | | | | ORF1ab | synonymous_T4983T |
| 15225 | T | C | 6.32E-05 | | | | | ORF1ab | synonymous_Y4987Y |
| 15226 | G | A | 6.32E-05 | | | | | ORF1ab | nonsynonymous_G4988S |
| 15257 | A | G | 6.32E-05 | | | | | ORF1ab | nonsynonymous_Y4998C |
| 15273 | C | T | 6.32E-05 | | | | | ORF1ab | synonymous_N5003N |
| 15303 | A | G | 6.32E-05 | | | | | ORF1ab | synonymous_K5013K |
| 15328 | C | T | 6.32E-05 | | | | | ORF1ab | nonsynonymous_L5022F |
| 15346 | C | A | 6.32E-05 | A | 6.32E-05 | | | ORF1ab | nonsynonymous_L5028I; nonsynonymous_L5028I |
| 15348 | T | A | 6.32E-05 | | | | | ORF1ab | synonymous_L5028L |
| 15349 | G | C | 6.32E-05 | | | | | ORF1ab | nonsynonymous_V5029L |
| 15352 | C | T | 6.32E-05 | | | | | ORF1ab | nonsynonymous_L5030F |
| 15359 | G | A | 6.32E-05 | | | | | ORF1ab | nonsynonymous_R5032H |
| 15384 | G | T | 6.32E-05 | | | | | ORF1ab | nonsynonymous_L5040F |
| 15399 | T | C | 6.32E-05 | | | | | ORF1ab | synonymous_Y5045Y |
| 15407 | C | T | 6.32E-05 | | | | | ORF1ab | nonsynonymous_A5048V |
| 15414 | G | A | 6.32E-05 | | | | | ORF1ab | synonymous_E5050E |
| 15429 | G | A | 6.32E-05 | | | | | ORF1ab | synonymous_L5055L |
| 15448 | G | A | 6.32E-05 | | | | | ORF1ab | nonsynonymous_G5062S |
| 15460 | T | C | 6.32E-05 | | | | | ORF1ab | nonsynonymous_Y5066H |
| 15462 | T | C | 6.32E-05 | | | | | ORF1ab | synonymous_Y5066Y |
| 15463 | G | T | 6.32E-05 | | | | | ORF1ab | nonsynonymous_V5067F |
| 15491 | A | G | 6.32E-05 | | | | | ORF1ab | nonsynonymous_D5076G |
| 15492 | T | A | 6.32E-05 | | | | | ORF1ab | nonsynonymous_D5076E |
| 15525 | C | T | 6.32E-05 | | | | | ORF1ab | synonymous_N5087N |
| 15539 | T | C | 6.32E-05 | | | | | ORF1ab | nonsynonymous_V5092A |
| 15543 | G | T | 6.32E-05 | | | | | ORF1ab | synonymous_T5093T |
| 15553 | A | G | 6.32E-05 | | | | | ORF1ab | nonsynonymous_N5097D |
| 15571 | G | T | 6.32E-05 | | | | | ORF1ab | nonsynonymous_D5103Y |

| | | | | | | | | | |
|---|---|---|---|---|---|---|---|---|---|
| 15574 | G | A | 6.32E-05 | | | | | ORF1ab | nonsynonymous_G5104S |
| 15575 | G | T | 6.32E-05 | | | | | ORF1ab | nonsynonymous_G5104V |
| 15596 | A | G | 6.32E-05 | | | | | ORF1ab | nonsynonymous_Y5111C |
| 15597 | T | C | 6.32E-05 | | | | | ORF1ab | synonymous_Y5111Y |
| 15603 | C | T | 6.32E-05 | | | | | ORF1ab | synonymous_R5113R |
| 15607 | T | C | 6.32E-05 | | | | | ORF1ab | synonymous_L5115L |
| 15615 | C | T | 6.32E-05 | | | | | ORF1ab | synonymous_H5117H |
| 15624 | T | C | 6.32E-05 | | | | | ORF1ab | synonymous_Y5120Y |
| 15627 | G | T | 6.32E-05 | | | | | ORF1ab | nonsynonymous_E5121D |
| 15636 | T | A | 6.32E-05 | | | | | ORF1ab | stopgain_Y5124X |
| 15638 | G | A | 6.32E-05 | | | | | ORF1ab | nonsynonymous_R5125K |
| 15640 | A | T | 6.32E-05 | | | | | ORF1ab | nonsynonymous_N5126Y |
| 15641 | A | C | 6.32E-05 | | | | | ORF1ab | nonsynonymous_N5126T |
| 15648 | T | C | 6.32E-05 | | | | | ORF1ab | synonymous_D5128D |
| 15651 | T | C | 6.32E-05 | | | | | ORF1ab | synonymous_V5129V |
| 15664 | G | A | 6.32E-05 | | | | | ORF1ab | nonsynonymous_V5134M |
| 15665 | T | C | 6.32E-05 | | | | | ORF1ab | nonsynonymous_V5134A |
| 15684 | T | C | 6.32E-05 | | | | | ORF1ab | synonymous_Y5140Y |
| 15688 | C | T | 6.32E-05 | | | | | ORF1ab | nonsynonymous_R5142C |
| 15696 | T | G | 6.32E-05 | | | | | ORF1ab | nonsynonymous_H5144Q |
| 15708 | G | T | 6.32E-05 | T | 6.32E-05 | | | ORF1ab | nonsynonymous_M5148I; nonsynonymous_M5148I |
| 15709 | A | G | 6.32E-05 | | | | | ORF1ab | nonsynonymous_I5149V |
| 15743 | G | A | 6.32E-05 | | | | | ORF1ab | nonsynonymous_S5160N |
| 15755 | C | T | 6.32E-05 | | | | | ORF1ab | nonsynonymous_S5164F |
| 15792 | A | G | 6.32E-05 | | | | | ORF1ab | synonymous_S5176S |
| 15821 | T | G | 6.32E-05 | | | | | ORF1ab | nonsynonymous_M5186R |
| 15898 | G | T | 6.32E-05 | | | | | ORF1ab | nonsynonymous_V5212F |
| 15907 | G | A | 6.32E-05 | | | | | ORF1ab | nonsynonymous_G5215S |
| 15908 | G | T | 6.32E-05 | | | | | ORF1ab | nonsynonymous_G5215V |

| | | | | | | | | | |
|---|---|---|---|---|---|---|---|---|---|
| 15909 | T | A | 6.32E-05 | | | | | ORF1ab | synonymous_G5215G |
| 15914 | A | G | 6.32E-05 | | | | | ORF1ab | nonsynonymous_D5217G |
| 15915 | T | C | 6.32E-05 | | | | | ORF1ab | synonymous_D5217D |
| 15924 | C | T | 6.32E-05 | | | | | ORF1ab | synonymous_Y5220Y |
| 15958 | G | A | 6.32E-05 | | | | | ORF1ab | nonsynonymous_A5232T |
| 15963 | C | T | 6.32E-05 | | | | | ORF1ab | synonymous_G5233G |
| 15989 | C | T | 6.32E-05 | | | | | ORF1ab | nonsynonymous_T5242I |
| 16008 | T | C | 6.32E-05 | | | | | ORF1ab | synonymous_I5248I |
| 1604 | A | G | 6.32E-05 | | | | | ORF1ab | nonsynonymous_N447D |
| 16053 | A | G | 6.32E-05 | | | | | ORF1ab | synonymous_K5263K |
| 16060 | A | G | 6.32E-05 | | | | | ORF1ab | nonsynonymous_N5266D |
| 16061 | A | G | 6.32E-05 | | | | | ORF1ab | nonsynonymous_N5266S |
| 1609 | C | T | 6.32E-05 | | | | | ORF1ab | synonymous_D448D |
| 16137 | G | T | 6.32E-05 | | | | | ORF1ab | nonsynonymous_M5291I |
| 16146 | G | T | 6.32E-05 | | | | | ORF1ab | nonsynonymous_M5294I |
| 16166 | A | G | 6.32E-05 | | | | | ORF1ab | nonsynonymous_N5301S |
| 16179 | A | G | 6.32E-05 | | | | | ORF1ab | synonymous_S5305S |
| 16192 | C | T | 6.32E-05 | | | | | ORF1ab | nonsynonymous_P5310S |
| 16230 | C | T | 6.32E-05 | | | | | ORF1ab | synonymous_V5322V |
| 16242 | T | C | 6.32E-05 | | | | | ORF1ab | synonymous_V5326V |
| 16245 | G | A | 6.32E-05 | | | | | ORF1ab | synonymous_G5327G |
| 16272 | T | G | 6.32E-05 | G | 6.32E-05 | | | ORF1ab | synonymous_T5336T; synonymous_T5336T |
| 16332 | C | T | 6.32E-05 | | | | | ORF1ab | synonymous_D5356D |
| 16342 | T | C | 6.32E-05 | | | | | ORF1ab | nonsynonymous_S5360P |
| 16349 | C | T | 6.32E-05 | | | | | ORF1ab | nonsynonymous_S5362L |
| 16386 | C | T | 6.32E-05 | | | | | ORF1ab | synonymous_C5374C |
| 16394 | C | T | 6.32E-05 | | | | | ORF1ab | nonsynonymous_P5377L |
| 16409 | C | T | 6.32E-05 | | | | | ORF1ab | nonsynonymous_T5382I |
| 16410 | A | G | 6.32E-05 | | | | | ORF1ab | synonymous_T5382T |

| | | | | | | | | | |
|---|---|---|---|---|---|---|---|---|---|
| 16417 | A | G | 6.32E-05 | | | | | ORF1ab | nonsynonymous_T5385A |
| 16419 | T | C | 6.32E-05 | | | | | ORF1ab | synonymous_T5385T |
| 16422 | A | C | 6.32E-05 | | | | | ORF1ab | nonsynonymous_Q5386H |
| 16425 | T | A | 6.32E-05 | | | | | ORF1ab | synonymous_L5387L |
| 16438 | A | G | 6.32E-05 | | | | | ORF1ab | nonsynonymous_M5392V |
| 16455 | A | G | 6.32E-05 | | | | | ORF1ab | synonymous_K5397K |
| 16469 | C | T | 6.32E-05 | | | | | ORF1ab | nonsynonymous_P5402L |
| 16470 | C | T | 6.32E-05 | | | | | ORF1ab | synonymous_P5402P |
| 16812 | A | G | 6.32E-05 | | | | | ORF1ab | synonymous_K5516K |
| 1681 | G | A | 6.32E-05 | | | | | ORF1ab | synonymous_E472E |
| 16838 | A | C | 6.32E-05 | | | | | ORF1ab | nonsynonymous_E5525A |
| 16839 | A | G | 6.32E-05 | | | | | ORF1ab | synonymous_E5525E |
| 16841 | A | G | 6.32E-05 | | | | | ORF1ab | nonsynonymous_K5526R |
| 16848 | C | T | 6.32E-05 | | | | | ORF1ab | synonymous_D5528D |
| 16853 | G | T | 6.32E-05 | | | | | ORF1ab | nonsynonymous_G5530V |
| 16856 | A | G | 6.32E-05 | | | | | ORF1ab | nonsynonymous_D5531G |
| 16877 | C | T | 6.32E-05 | | | | | ORF1ab | nonsynonymous_T5538I |
| 16881 | A | G | 6.32E-05 | | | | | ORF1ab | synonymous_T5539T |
| 16890 | A | G | 6.32E-05 | | | | | ORF1ab | synonymous_K5542K |
| 16897 | G | T | 6.32E-05 | | | | | ORF1ab | nonsynonymous_V5545F |
| 16909 | T | A | 6.32E-05 | | | | | ORF1ab | nonsynonymous_F5549I |
| 16939 | T | C | 6.32E-05 | | | | | ORF1ab | synonymous_L5559L |
| 16941 | A | G | 6.32E-05 | | | | | ORF1ab | synonymous_L5559L |
| 16945 | G | T | 6.32E-05 | | | | | ORF1ab | nonsynonymous_A5561S |
| 16947 | A | G | 6.32E-05 | | | | | ORF1ab | synonymous_A5561A |
| 16952 | C | T | 6.32E-05 | | | | | ORF1ab | nonsynonymous_T5563I |
| 16957 | G | A | 6.32E-05 | A | 6.32E-05 | | | ORF1ab | nonsynonymous_V5565M; nonsynonymous_V5565M |
| 16960 | C | T | 6.32E-05 | | | | | ORF1ab | nonsynonymous_P5566S |
| 16979 | G | T | 6.32E-05 | | | | | ORF1ab | nonsynonymous_R5572I |

| | | | | | | | | | |
|---|---|---|---|---|---|---|---|---|---|
| 17000 | C | T | 6.32E-05 | | | | | ORF1ab | nonsynonymous_T5579I |
| 17049 | G | A | 6.32E-05 | A | 6.32E-05 | | | ORF1ab | synonymous_K5595K; synonymous_K5595K |
| 17058 | G | A | 6.32E-05 | | | | | ORF1ab | nonsynonymous_M5598I |
| 17059 | C | A | 6.32E-05 | | | | | ORF1ab | nonsynonymous_Q5599K |
| 17075 | T | A | 6.32E-05 | | | | | ORF1ab | nonsynonymous_L5604H |
| 17091 | T | C | 6.32E-05 | | | | | ORF1ab | synonymous_G5609G |
| 17097 | T | C | 6.32E-05 | | | | | ORF1ab | synonymous_G5611G |
| 17105 | A | G | 6.32E-05 | | | | | ORF1ab | nonsynonymous_H5614R |
| 17116 | G | T | 6.32E-05 | | | | | ORF1ab | nonsynonymous_G5618C |
| 17141 | C | A | 6.32E-05 | | | | | ORF1ab | nonsynonymous_A5626D |
| 17142 | T | C | 6.32E-05 | | | | | ORF1ab | synonymous_A5626A |
| 17151 | G | A | 6.32E-05 | | | | | ORF1ab | synonymous_V5629V |
| 17154 | T | C | 6.32E-05 | | | | | ORF1ab | synonymous_Y5630Y |
| 17165 | C | T | 6.32E-05 | | | | | ORF1ab | nonsynonymous_S5634F |
| 17185 | C | T | 6.32E-05 | | | | | ORF1ab | synonymous_L5641L |
| 17197 | G | A | 6.32E-05 | | | | | ORF1ab | nonsynonymous_A5645T |
| 17199 | A | G | 6.32E-05 | | | | | ORF1ab | synonymous_A5645A |
| 1719 | G | A | 6.32E-05 | A | 6.32E-05 | | | ORF1ab | nonsynonymous_S485N; nonsynonymous_S485N |
| 17200 | T | A | 6.32E-05 | | | | | ORF1ab | nonsynonymous_L5646I |
| 17215 | A | T | 6.32E-05 | T | 6.32E-05 | | | ORF1ab | nonsynonymous_I5651L; nonsynonymous_I5651L |
| 17222 | A | G | 6.32E-05 | | | | | ORF1ab | nonsynonymous_K5653R |
| 17229 | T | C | 6.32E-05 | | | | | ORF1ab | synonymous_S5655S |
| 17236 | A | G | 6.32E-05 | | | | | ORF1ab | nonsynonymous_I5658V |
| 1723 | T | C | 6.32E-05 | | | | | ORF1ab | synonymous_A486A |
| 17242 | G | A | 6.32E-05 | | | | | ORF1ab | nonsynonymous_A5660T |
| 17252 | G | T | 6.32E-05 | | | | | ORF1ab | nonsynonymous_R5663L |
| 17271 | A | C | 6.32E-05 | | | | | ORF1ab | nonsynonymous_K5669N |

| | | | | | | | | | |
|---|---|---|---|---|---|---|---|---|---|
| 17274 | C | T | 6.32E-05 | | | | | ORF1ab | synonymous_F5670F |
| 17280 | G | T | 6.32E-05 | | | | | ORF1ab | synonymous_V5672V |
| 17287 | A | G | 6.32E-05 | | | | | ORF1ab | nonsynonymous_T5675A |
| 17288 | C | T | 6.32E-05 | | | | | ORF1ab | nonsynonymous_T5675I |
| 17290 | T | C | 6.32E-05 | | | | | ORF1ab | synonymous_L5676L |
| 17314 | G | A | 6.32E-05 | | | | | ORF1ab | nonsynonymous_V5684I |
| 17334 | G | T | 6.32E-05 | | | | | ORF1ab | synonymous_T5690T |
| 1736 | G | A | 6.32E-05 | | | | | ORF1ab | nonsynonymous_V491M |
| 17376 | A | G | 6.32E-05 | | | | | ORF1ab | synonymous_T5704T |
| 17380 | T | C | 6.32E-05 | | | | | ORF1ab | nonsynonymous_Y5706H |
| 17415 | T | C | 6.32E-05 | | | | | ORF1ab | synonymous_A5717A |
| 17457 | A | G | 6.32E-05 | | | | | ORF1ab | synonymous_A5731A |
| 17460 | A | G | 6.32E-05 | | | | | ORF1ab | synonymous_P5732P |
| 17642 | C | T | 6.32E-05 | | | | | ORF1ab | nonsynonymous_A5793V |
| 17661 | T | C | 6.32E-05 | | | | | ORF1ab | synonymous_F5799F |
| 17664 | T | C | 6.32E-05 | | | | | ORF1ab | synonymous_Y5800Y |
| 17694 | T | C | 6.32E-05 | | | | | ORF1ab | synonymous_S5810S |
| 17697 | A | T | 6.32E-05 | | | | | ORF1ab | synonymous_A5811A |
| 17723 | T | C | 6.32E-05 | | | | | ORF1ab | nonsynonymous_V5820A |
| 17750 | C | T | 6.32E-05 | | | | | ORF1ab | nonsynonymous_A5829V |
| 17754 | G | T | 6.32E-05 | | | | | ORF1ab | nonsynonymous_W5830C |
| 17762 | C | T | 6.32E-05 | | | | | ORF1ab | nonsynonymous_A5833V |
| 17790 | G | T | 6.32E-05 | | | | | ORF1ab | nonsynonymous_Q5842H |
| 17795 | C | T | 6.32E-05 | | | | | ORF1ab | nonsynonymous_A5844V |
| 17802 | C | T | 6.32E-05 | | | | | ORF1ab | synonymous_A5846A |
| 17805 | A | T | 6.32E-05 | | | | | ORF1ab | synonymous_S5847S |
| 17814 | G | T | 6.32E-05 | | | | | ORF1ab | nonsynonymous_L5850F |
| 17817 | A | G | 6.32E-05 | | | | | ORF1ab | synonymous_G5851G |
| 17844 | A | G | 6.32E-05 | | | | | ORF1ab | synonymous_S5860S |
| 17892 | A | G | 6.32E-05 | | | | | ORF1ab | synonymous_T5876T |

| | | | | | | | | | |
|---|---|---|---|---|---|---|---|---|---|
| 17893 | G | T | 6.32E-05 | | | | | ORF1ab | nonsynonymous_A5877S |
| 17928 | T | C | 6.32E-05 | | | | | ORF1ab | synonymous_A5888A |
| 17938 | G | A | 6.32E-05 | | | | | ORF1ab | nonsynonymous_A5892T |
| 17950 | A | G | 6.32E-05 | | | | | ORF1ab | nonsynonymous_I5896V |
| 17959 | A | G | 6.32E-05 | | | | | ORF1ab | nonsynonymous_I5899V |
| 17961 | A | G | 6.32E-05 | | | | | ORF1ab | nonsynonymous_I5899M |
| 17985 | C | T | 6.32E-05 | | | | | ORF1ab | synonymous_D5907D |
| 17993 | A | G | 6.32E-05 | | | | | ORF1ab | nonsynonymous_Q5910R |
| 17994 | A | C | 6.32E-05 | | | | | ORF1ab | nonsynonymous_Q5910H |
| 18015 | A | G | 6.32E-05 | | | | | ORF1ab | synonymous_P5917P |
| 18017 | G | T | 6.32E-05 | | | | | ORF1ab | nonsynonymous_R5918L |
| 18019 | A | G | 6.32E-05 | | | | | ORF1ab | nonsynonymous_R5919G |
| 18027 | G | T | 6.32E-05 | | | | | ORF1ab | synonymous_V5921V |
| 18028 | G | T | 6.32E-05 | | | | | ORF1ab | nonsynonymous_A5922S |
| 18078 | G | A | 6.32E-05 | | | | | ORF1ab | synonymous_K5938K |
| 18090 | G | A | 6.32E-05 | A | 6.32E-05 | | | ORF1ab | synonymous_G5942G; synonymous_G5942G |
| 18093 | A | G | 6.32E-05 | | | | | ORF1ab | synonymous_L5943L |
| 18238 | A | T | 6.32E-05 | | | | | ORF1ab | nonsynonymous_N5992Y |
| 18245 | A | T | 6.32E-05 | | | | | ORF1ab | nonsynonymous_Y5994F |
| 18250 | A | T | 6.32E-05 | | | | | ORF1ab | nonsynonymous_N5996Y |
| 18253 | A | T | 6.32E-05 | | | | | ORF1ab | nonsynonymous_M5997L |
| 18259 | A | T | 6.32E-05 | | | | | ORF1ab | nonsynonymous_I5999F |
| 18260 | T | C | 6.32E-05 | | | | | ORF1ab | nonsynonymous_I5999T |
| 18261 | C | T | 6.32E-05 | | | | | ORF1ab | synonymous_I5999I |
| 18266 | G | T | 6.32E-05 | | | | | ORF1ab | nonsynonymous_R6001L |
| 18269 | A | T | 6.32E-05 | | | | | ORF1ab | nonsynonymous_E6002V |
| 18277 | A | T | 6.32E-05 | | | | | ORF1ab | nonsynonymous_I6005L |
| 18278 | T | C | 6.32E-05 | | | | | ORF1ab | nonsynonymous_I6005T |
| 18279 | A | T | 6.32E-05 | | | | | ORF1ab | synonymous_I6005I |

| | | | | | | | | | |
|---|---|---|---|---|---|---|---|---|---|
| 18282 | A | T | 6.32E-05 | | | | | ORF1ab | nonsynonymous_R6006S |
| 18284 | A | T | 6.32E-05 | | | | | ORF1ab | nonsynonymous_H6007L |
| 18286 | G | T | 6.32E-05 | | | | | ORF1ab | nonsynonymous_V6008L |
| 18288 | A | T | 6.32E-05 | | | | | ORF1ab | synonymous_V6008V |
| 18294 | A | T | 6.32E-05 | | | | | ORF1ab | synonymous_A6010A |
| 18297 | G | T | 6.32E-05 | | | | | ORF1ab | nonsynonymous_W6011C |
| 18298 | A | T | 6.32E-05 | | | | | ORF1ab | nonsynonymous_I6012F |
| 18300 | T | C | 6.32E-05 | | | | | ORF1ab | synonymous_I6012I |
| 18303 | C | T | 6.32E-05 | | | | | ORF1ab | synonymous_G6013G |
| 18310 | G | A | 6.32E-05 | | | | | ORF1ab | nonsynonymous_V6016I |
| 18322 | C | T | 6.32E-05 | | | | | ORF1ab | nonsynonymous_H6020Y |
| 18326 | C | T | 6.32E-05 | | | | | ORF1ab | nonsynonymous_A6021V |
| 18329 | C | T | 6.32E-05 | | | | | ORF1ab | nonsynonymous_T6022I |
| 18333 | A | T | 6.32E-05 | | | | | ORF1ab | nonsynonymous_R6023S |
| 18340 | G | A | 6.32E-05 | | | | | ORF1ab | nonsynonymous_V6026I |
| 18345 | T | C | 6.32E-05 | | | | | ORF1ab | synonymous_G6027G |
| 18364 | C | T | 6.32E-05 | | | | | ORF1ab | synonymous_L6034L |
| 18371 | T | A | 6.32E-05 | | | | | ORF1ab | nonsynonymous_F6036Y |
| 18380 | G | T | 6.32E-05 | | | | | ORF1ab | nonsynonymous_G6039V |
| 18381 | T | C | 6.32E-05 | | | | | ORF1ab | synonymous_G6039G |
| 18384 | T | C | 6.32E-05 | | | | | ORF1ab | synonymous_V6040V |
| 18391 | G | A | 6.32E-05 | | | | | ORF1ab | nonsynonymous_V6043I |
| 18402 | T | C | 6.32E-05 | | | | | ORF1ab | synonymous_P6046P |
| 18410 | A | G | 6.32E-05 | | | | | ORF1ab | nonsynonymous_Y6049C |
| 18417 | T | C | 6.32E-05 | | | | | ORF1ab | synonymous_D6051D |
| 18431 | C | T | 6.32E-05 | | | | | ORF1ab | nonsynonymous_T6056I |
| 18436 | T | C | 6.32E-05 | | | | | ORF1ab | nonsynonymous_F6058L |
| 18441 | C | T | 6.32E-05 | | | | | ORF1ab | synonymous_S6059S |
| 18462 | G | T | 6.32E-05 | | | | | ORF1ab | synonymous_P6066P |
| 18471 | T | C | 6.32E-05 | | | | | ORF1ab | synonymous_D6069D |

| | | | | | | | | | |
|---|---|---|---|---|---|---|---|---|---|
| 18481 | C | T | 6.32E-05 | | | | | ORF1ab | nonsynonymous_H6073Y |
| 18483 | C | T | 6.32E-05 | | | | | ORF1ab | synonymous_H6073H |
| 18494 | T | C | 6.32E-05 | | | | | ORF1ab | nonsynonymous_L6077P |
| 18499 | T | A | 6.32E-05 | | | | | ORF1ab | nonsynonymous_Y6079N |
| 18500 | A | G | 6.32E-05 | | | | | ORF1ab | nonsynonymous_Y6079C |
| 18502 | A | G | 6.32E-05 | | | | | ORF1ab | nonsynonymous_K6080E |
| 18509 | T | C | 6.32E-05 | | | | | ORF1ab | nonsynonymous_L6082P |
| 18514 | T | A | 6.32E-05 | | | | | ORF1ab | nonsynonymous_W6084R |
| 18515 | G | A | 6.32E-05 | | | | | ORF1ab | stopgain_W6084X |
| 18516 | G | T | 6.32E-05 | | | | | ORF1ab | nonsynonymous_W6084C |
| 18519 | T | C | 6.32E-05 | | | | | ORF1ab | synonymous_N6085N |
| 18521 | T | C | 6.32E-05 | | | | | ORF1ab | nonsynonymous_V6086A |
| 18529 | A | T | 6.32E-05 | | | | | ORF1ab | nonsynonymous_I6089L |
| 18530 | T | G | 6.32E-05 | | | | | ORF1ab | nonsynonymous_I6089R |
| 18533 | A | C | 6.32E-05 | | | | | ORF1ab | nonsynonymous_K6090T |
| 18538 | G | C | 6.32E-05 | | | | | ORF1ab | nonsynonymous_V6092L |
| 18546 | G | T | 6.32E-05 | T | 6.32E-05 | | | ORF1ab | nonsynonymous_M6094I; nonsynonymous_M6094I |
| 18549 | A | C | 6.32E-05 | | | | | ORF1ab | nonsynonymous_L6095F |
| 18553 | G | A | 6.32E-05 | | | | | ORF1ab | nonsynonymous_D6097N |
| 18558 | A | G | 6.32E-05 | | | | | ORF1ab | synonymous_T6098T |
| 18559 | C | T | 6.32E-05 | | | | | ORF1ab | nonsynonymous_L6099F |
| 18564 | A | C | 6.32E-05 | | | | | ORF1ab | nonsynonymous_K6100N |
| 18567 | T | A | 6.32E-05 | | | | | ORF1ab | nonsynonymous_N6101K |
| 18571 | T | G | 6.32E-05 | | | | | ORF1ab | nonsynonymous_S6103A |
| 18572 | C | T | 6.32E-05 | | | | | ORF1ab | nonsynonymous_S6103F |
| 18573 | T | G | 6.32E-05 | G | 6.32E-05 | | | ORF1ab | synonymous_S6103S; synonymous_S6103S |
| 18576 | C | T | 6.32E-05 | | | | | ORF1ab | synonymous_D6104D |
| 18582 | C | T | 6.32E-05 | | | | | ORF1ab | synonymous_V6106V |

| | | | | | | | | | |
|---|---|---|---|---|---|---|---|---|---|
| 18585 | A | G | 6.32E-05 | | | | | ORF1ab | synonymous_V6107V |
| 18586 | T | A | 6.32E-05 | | | | | ORF1ab | nonsynonymous_F6108I |
| 18589 | G | T | 6.32E-05 | | | | | ORF1ab | nonsynonymous_V6109F |
| 18590 | T | C | 6.32E-05 | | | | | ORF1ab | nonsynonymous_V6109A |
| 18591 | C | A | 6.32E-05 | | | | | ORF1ab | synonymous_V6109V |
| 18592 | T | C | 6.32E-05 | | | | | ORF1ab | synonymous_L6110L |
| 18593 | T | C | 6.32E-05 | | | | | ORF1ab | nonsynonymous_L6110S |
| 18596 | G | A | 6.32E-05 | | | | | ORF1ab | stopgain_W6111X |
| 18598 | G | C | 6.32E-05 | | | | | ORF1ab | nonsynonymous_A6112P |
| 18599 | C | T | 6.32E-05 | | | | | ORF1ab | nonsynonymous_A6112V |
| 18607 | T | A | 6.32E-05 | | | | | ORF1ab | nonsynonymous_F6115I |
| 18615 | G | A | 6.32E-05 | | | | | ORF1ab | synonymous_L6117L |
| 18624 | G | A | 6.32E-05 | | | | | ORF1ab | nonsynonymous_M6120I |
| 18625 | A | C | 6.32E-05 | | | | | ORF1ab | nonsynonymous_K6121Q |
| 18627 | G | A | 6.32E-05 | | | | | ORF1ab | synonymous_K6121K |
| 18629 | A | G | 6.32E-05 | | | | | ORF1ab | nonsynonymous_Y6122C |
| 18632 | T | G | 6.32E-05 | | | | | ORF1ab | nonsynonymous_F6123C |
| 18731 | T | A | 6.32E-05 | A | 6.32E-05 | | | ORF1ab | nonsynonymous_I6156N; nonsynonymous_I6156N |
| 18732 | T | A | 6.32E-05 | | | | | ORF1ab | synonymous_I6156I |
| 18733 | G | A | 6.32E-05 | | | | | ORF1ab | nonsynonymous_G6157R |
| 18735 | A | C | 6.32E-05 | | | | | ORF1ab | synonymous_G6157G |
| 18738 | T | G | 6.32E-05 | | | | | ORF1ab | nonsynonymous_F6158L |
| 18739 | G | T | 6.32E-05 | | | | | ORF1ab | nonsynonymous_D6159Y |
| 18740 | A | G | 6.32E-05 | | | | | ORF1ab | nonsynonymous_D6159G |
| 18741 | T | G | 6.32E-05 | | | | | ORF1ab | nonsynonymous_D6159E |
| 18742 | T | C | 6.32E-05 | | | | | ORF1ab | nonsynonymous_Y6160H |
| 18743 | A | T | 6.32E-05 | | | | | ORF1ab | nonsynonymous_Y6160F |
| 18745 | G | T | 6.32E-05 | | | | | ORF1ab | nonsynonymous_V6161F |
| 18750 | T | C | 6.32E-05 | | | | | ORF1ab | synonymous_Y6162Y |

| | | | | | | | | | |
|---|---|---|---|---|---|---|---|---|---|
| 18751 | A | C | 6.32E-05 | | | | | ORF1ab | nonsynonymous_N6163H |
| 18757 | T | C | 6.32E-05 | | | | | ORF1ab | nonsynonymous_F6165L |
| 18759 | T | C | 6.32E-05 | | | | | ORF1ab | synonymous_F6165F |
| 18761 | T | G | 6.32E-05 | | | | | ORF1ab | nonsynonymous_M6166R |
| 18762 | G | T | 6.32E-05 | | | | | ORF1ab | nonsynonymous_M6166I |
| 18763 | A | T | 6.32E-05 | | | | | ORF1ab | nonsynonymous_I6167F |
| 18766 | G | T | 6.32E-05 | | | | | ORF1ab | nonsynonymous_D6168Y |
| 18768 | T | C | 6.32E-05 | | | | | ORF1ab | synonymous_D6168D |
| 18769 | G | T | 6.32E-05 | | | | | ORF1ab | nonsynonymous_V6169F |
| 18774 | A | C | 6.32E-05 | | | | | ORF1ab | nonsynonymous_Q6170H |
| 18775 | C | A | 6.32E-05 | | | | | ORF1ab | nonsynonymous_Q6171K |
| 18780 | G | A | 6.32E-05 | A | 6.32E-05 | | | ORF1ab | stopgain_W6172X; stopgain_W6172X |
| 18781 | G | T | 6.32E-05 | | | | | ORF1ab | nonsynonymous_G6173C |
| 18782 | G | A | 6.32E-05 | A | 6.32E-05 | | | ORF1ab | nonsynonymous_G6173D; nonsynonymous_G6173D |
| 18783 | T | C | 6.32E-05 | | | | | ORF1ab | synonymous_G6173G |
| 18786 | T | C | 6.32E-05 | | | | | ORF1ab | synonymous_F6174F |
| 18791 | G | A | 6.32E-05 | | | | | ORF1ab | nonsynonymous_G6176D |
| 18793 | A | G | 6.32E-05 | | | | | ORF1ab | nonsynonymous_N6177D |
| 18794 | A | T | 6.32E-05 | T | 6.32E-05 | | | ORF1ab | nonsynonymous_N6177I; nonsynonymous_N6177I |
| 18795 | C | T | 6.32E-05 | | | | | ORF1ab | synonymous_N6177N |
| 18796 | C | T | 6.32E-05 | | | | | ORF1ab | synonymous_L6178L |
| 18797 | T | C | 6.32E-05 | | | | | ORF1ab | nonsynonymous_L6178P |
| 18799 | C | T | 6.32E-05 | | | | | ORF1ab | stopgain_Q6179X |
| 18801 | A | C | 6.32E-05 | | | | | ORF1ab | nonsynonymous_Q6179H |
| 18807 | C | A | 6.32E-05 | | | | | ORF1ab | nonsynonymous_N6181K |
| 18808 | C | T | 6.32E-05 | | | | | ORF1ab | nonsynonymous_H6182Y |
| 18810 | T | C | 6.32E-05 | | | | | ORF1ab | synonymous_H6182H |

| | | | | | | | | | |
|---|---|---|---|---|---|---|---|---|---|
| 18812 | A | G | 6.32E-05 | | | | | ORF1ab | nonsynonymous_D6183G |
| 18818 | A | G | 6.32E-05 | | | | | ORF1ab | nonsynonymous_Y6185C |
| 18821 | G | T | 6.32E-05 | T | 6.32E-05 | | | ORF1ab | nonsynonymous_C6186F; nonsynonymous_C6186F |
| 18825 | A | G | 6.32E-05 | | | | | ORF1ab | synonymous_Q6187Q |
| 18830 | A | G | 6.32E-05 | | | | | ORF1ab | nonsynonymous_H6189R |
| 18833 | G | T | 6.32E-05 | | | | | ORF1ab | nonsynonymous_G6190V |
| 18836 | A | C | 6.32E-05 | | | | | ORF1ab | nonsynonymous_N6191T |
| 18837 | T | C | 6.32E-05 | | | | | ORF1ab | synonymous_N6191N |
| 18838 | G | T | 6.32E-05 | | | | | ORF1ab | nonsynonymous_A6192S |
| 18839 | C | A | 6.32E-05 | | | | | ORF1ab | nonsynonymous_A6192E |
| 18846 | A | G | 6.32E-05 | | | | | ORF1ab | synonymous_V6194V |
| 18847 | G | T | 6.32E-05 | | | | | ORF1ab | nonsynonymous_A6195S |
| 18848 | C | A | 6.32E-05 | | | | | ORF1ab | nonsynonymous_A6195D |
| 18849 | T | C | 6.32E-05 | | | | | ORF1ab | synonymous_A6195A |
| 18851 | G | A | 6.32E-05 | | | | | ORF1ab | nonsynonymous_S6196N |
| 18853 | T | C | 6.32E-05 | | | | | ORF1ab | nonsynonymous_C6197R |
| 18854 | G | T | 6.32E-05 | | | | | ORF1ab | nonsynonymous_C6197F |
| 18856 | G | T | 6.32E-05 | | | | | ORF1ab | nonsynonymous_D6198Y |
| 18862 | A | C | 6.32E-05 | | | | | ORF1ab | nonsynonymous_I6200L |
| 18867 | G | T | 6.32E-05 | | | | | ORF1ab | nonsynonymous_M6201I |
| 18868 | A | C | 6.32E-05 | | | | | ORF1ab | nonsynonymous_T6202P |
| 18870 | T | A | 6.32E-05 | | | | | ORF1ab | synonymous_T6202T |
| 18874 | T | A | 6.32E-05 | | | | | ORF1ab | nonsynonymous_C6204S |
| 18875 | G | A | 6.32E-05 | | | | | ORF1ab | nonsynonymous_C6204Y |
| 18876 | T | G | 6.32E-05 | | | | | ORF1ab | nonsynonymous_C6204W |
| 18878 | T | C | 6.32E-05 | | | | | ORF1ab | nonsynonymous_L6205P |
| 18879 | A | C | 6.32E-05 | | | | | ORF1ab | synonymous_L6205L |
| 18880 | G | T | 6.32E-05 | | | | | ORF1ab | nonsynonymous_A6206S |
| 18881 | C | T | 6.32E-05 | | | | | ORF1ab | nonsynonymous_A6206V |

| | | | | | | | | | |
|---|---|---|---|---|---|---|---|---|---|
| 18885 | C | A | 6.32E-05 | A | 6.32E-05 | | | ORF1ab | synonymous_V6207V; synonymous_V6207V |
| 18887 | A | G | 6.32E-05 | | | | | ORF1ab | nonsynonymous_H6208R |
| 18889 | G | A | 6.32E-05 | | | | | ORF1ab | nonsynonymous_E6209K |
| 18890 | A | G | 6.32E-05 | | | | | ORF1ab | nonsynonymous_E6209G |
| 18891 | G | A | 6.32E-05 | | | | | ORF1ab | synonymous_E6209E |
| 18896 | T | A | 6.32E-05 | | | | | ORF1ab | nonsynonymous_F6211Y |
| 18902 | A | G | 6.32E-05 | | | | | ORF1ab | nonsynonymous_K6213R |
| 18919 | A | C | 6.32E-05 | | | | | ORF1ab | nonsynonymous_I6219L |
| 18921 | T | G | 6.32E-05 | | | | | ORF1ab | nonsynonymous_I6219M |
| 18925 | T | A | 6.32E-05 | | | | | ORF1ab | nonsynonymous_Y6221N |
| 18929 | C | T | 6.32E-05 | | | | | ORF1ab | nonsynonymous_P6222L |
| 18931 | A | G | 6.32E-05 | | | | | ORF1ab | nonsynonymous_I6223V |
| 18940 | G | T | 6.32E-05 | | | | | ORF1ab | nonsynonymous_D6226Y |
| 18943 | G | T | 6.32E-05 | T | 6.32E-05 | | | ORF1ab | stopgain_E6227X; stopgain_E6227X |
| 18945 | A | G | 6.32E-05 | | | | | ORF1ab | synonymous_E6227E |
| 18946 | C | G | 6.32E-05 | | | | | ORF1ab | nonsynonymous_L6228V |
| 18951 | G | T | 6.32E-05 | | | | | ORF1ab | nonsynonymous_K6229N |
| 18954 | T | C | 6.32E-05 | | | | | ORF1ab | synonymous_I6230I |
| 18956 | A | G | 6.32E-05 | | | | | ORF1ab | nonsynonymous_N6231S |
| 18957 | T | C | 6.32E-05 | | | | | ORF1ab | synonymous_N6231N |
| 18958 | G | A | 6.32E-05 | | | | | ORF1ab | nonsynonymous_A6232T |
| 18959 | C | T | 6.32E-05 | | | | | ORF1ab | nonsynonymous_A6232V |
| 18960 | G | A | 6.32E-05 | | | | | ORF1ab | synonymous_A6232A |
| 18961 | G | A | 6.32E-05 | | | | | ORF1ab | nonsynonymous_A6233T |
| 18963 | T | C | 6.32E-05 | | | | | ORF1ab | synonymous_A6233A |
| 18969 | A | G | 6.32E-05 | | | | | ORF1ab | synonymous_R6235R |
| 18970 | A | T | 6.32E-05 | | | | | ORF1ab | stopgain_K6236X |
| 18971 | A | G | 6.32E-05 | | | | | ORF1ab | nonsynonymous_K6236R |

| | | | | | | | | | |
|---|---|---|---|---|---|---|---|---|---|
| 18973 | G | T | 6.32E-05 | | | | | ORF1ab | nonsynonymous_V6237F |
| 18975 | T | A | 6.32E-05 | | | | | ORF1ab | synonymous_V6237V |
| 19003 | T | A | 6.32E-05 | | | | | ORF1ab | nonsynonymous_L6247I |
| 19009 | G | T | 6.32E-05 | | | | | ORF1ab | nonsynonymous_D6249Y |
| 19015 | T | A | 6.32E-05 | | | | | ORF1ab | nonsynonymous_F6251I |
| 19019 | C | T | 6.32E-05 | | | | | ORF1ab | nonsynonymous_P6252L |
| 19027 | C | T | 6.32E-05 | | | | | ORF1ab | nonsynonymous_H6255Y |
| 19033 | A | G | 6.32E-05 | | | | | ORF1ab | nonsynonymous_I6257V |
| 19049 | C | T | 6.32E-05 | | | | | ORF1ab | nonsynonymous_A6262V |
| 19065 | T | C | 6.32E-05 | | | | | ORF1ab | synonymous_P6267P |
| 19071 | T | A | 6.32E-05 | | | | | ORF1ab | synonymous_A6269A |
| 19117 | G | T | 6.32E-05 | | | | | ORF1ab | nonsynonymous_A6285S |
| 19132 | G | A | 6.32E-05 | | | | | ORF1ab | nonsynonymous_E6290K |
| 19135 | T | C | 6.32E-05 | | | | | ORF1ab | synonymous_L6291L |
| 19137 | A | T | 6.32E-05 | | | | | ORF1ab | nonsynonymous_L6291F |
| 19172 | C | T | 6.32E-05 | | | | | ORF1ab | nonsynonymous_T6303I |
| 19184 | G | T | 6.32E-05 | | | | | ORF1ab | nonsynonymous_C6307F |
| 19185 | C | T | 6.32E-05 | | | | | ORF1ab | synonymous_C6307C |
| 19190 | T | A | 6.32E-05 | | | | | ORF1ab | nonsynonymous_F6309Y |
| 19199 | G | A | 6.32E-05 | | | | | ORF1ab | nonsynonymous_C6312Y |
| 19209 | T | C | 6.32E-05 | | | | | ORF1ab | synonymous_D6315D |
| 19215 | T | C | 6.32E-05 | | | | | ORF1ab | synonymous_Y6317Y |
| 19220 | C | T | 6.32E-05 | | | | | ORF1ab | nonsynonymous_A6319V |
| 19255 | C | T | 6.32E-05 | | | | | ORF1ab | synonymous_L6331L |
| 19264 | C | T | 6.32E-05 | | | | | ORF1ab | nonsynonymous_L6334F |
| 1927 | T | A | 6.32E-05 | | | | | ORF1ab | synonymous_T554T |
| 19599 | G | T | 6.32E-05 | | | | | ORF1ab | nonsynonymous_W6445C |
| 19646 | T | C | 6.32E-05 | | | | | ORF1ab | nonsynonymous_V6461A |
| 19687 | C | T | 6.32E-05 | | | | | ORF1ab | nonsynonymous_P6475S |
| 19700 | T | G | 6.32E-05 | | | | | ORF1ab | nonsynonymous_I6479S |

| | | | | | | | | | |
|---|---|---|---|---|---|---|---|---|---|
| 19705 | A | G | 6.32E-05 | | | | | ORF1ab | nonsynonymous_N6481D |
| 19720 | A | G | 6.32E-05 | | | | | ORF1ab | nonsynonymous_K6486E |
| 19723 | G | T | 6.32E-05 | | | | | ORF1ab | nonsynonymous_V6487F |
| 19731 | T | C | 6.32E-05 | | | | | ORF1ab | synonymous_G6489G |
| 19783 | T | C | 6.32E-05 | | | | | ORF1ab | nonsynonymous_F6507L |
| 19788 | G | T | 6.32E-05 | | | | | ORF1ab | nonsynonymous_E6508D |
| 19836 | C | T | 6.32E-05 | | | | | ORF1ab | synonymous_L6524L |
| 19861 | G | A | 6.32E-05 | | | | | ORF1ab | nonsynonymous_A6533T |
| 19865 | A | G | 6.32E-05 | | | | | ORF1ab | nonsynonymous_N6534S |
| 19883 | A | G | 6.32E-05 | | | | | ORF1ab | nonsynonymous_Y6540C |
| 19900 | G | A | 6.32E-05 | | | | | ORF1ab | nonsynonymous_A6546T |
| 19902 | A | G | 6.32E-05 | | | | | ORF1ab | synonymous_A6546A |
| 19903 | C | T | 6.32E-05 | T | 6.32E-05 | | | ORF1ab | nonsynonymous_H6547Y; nonsynonymous_H6547Y |
| 19905 | T | C | 6.32E-05 | | | | | ORF1ab | synonymous_H6547H |
| 19909 | T | G | 6.32E-05 | | | | | ORF1ab | nonsynonymous_S6549A |
| 19910 | C | T | 6.32E-05 | | | | | ORF1ab | nonsynonymous_S6549F |
| 20205 | T | A | 6.32E-05 | | | | | ORF1ab | synonymous_T6647T |
| 20229 | T | C | 6.32E-05 | | | | | ORF1ab | synonymous_F6655F |
| 2028 | C | T | 6.32E-05 | | | | | ORF1ab | nonsynonymous_S588F |
| 2036 | G | A | 6.32E-05 | | | | | ORF1ab | nonsynonymous_A591T |
| 20603 | C | T | 6.32E-05 | | | | | ORF1ab | nonsynonymous_S6780L |
| 20621 | A | G | 6.32E-05 | | | | | ORF1ab | nonsynonymous_K6786R |
| 20622 | A | G | 6.32E-05 | | | | | ORF1ab | synonymous_K6786K |
| 20623 | G | A | 6.32E-05 | | | | | ORF1ab | nonsynonymous_D6787N |
| 20626 | G | C | 6.32E-05 | | | | | ORF1ab | nonsynonymous_G6788R |
| 20627 | G | A | 6.32E-05 | | | | | ORF1ab | nonsynonymous_G6788D |
| 20629 | C | T | 6.32E-05 | | | | | ORF1ab | nonsynonymous_H6789Y |
| 20630 | A | T | 6.32E-05 | | | | | ORF1ab | nonsynonymous_H6789L |
| 20632 | G | T | 6.32E-05 | | | | | ORF1ab | nonsynonymous_V6790L |

| | | | | | | | | | |
|---|---|---|---|---|---|---|---|---|---|
| 20634 | A | G | 6.32E-05 | | | | | ORF1ab | synonymous_V6790V |
| 20637 | A | G | 6.32E-05 | | | | | ORF1ab | synonymous_E6791E |
| 20639 | C | G | 6.32E-05 | | | | | ORF1ab | nonsynonymous_T6792R |
| 20650 | A | T | 6.32E-05 | T | 6.32E-05 | | | ORF1ab | stopgain_K6796X; stopgain_K6796X |
| 20652 | A | T | 6.32E-05 | | | | | ORF1ab | nonsynonymous_K6796N |
| 20653 | T | G | 6.32E-05 | | | | | ORF1ab | nonsynonymous_L6797V |
| 20654 | T | G | 6.32E-05 | | | | | ORF1ab | stopgain_L6797X |
| 20658 | A | G | 6.32E-05 | | | | | ORF1ab | synonymous_Q6798Q |
| 20660 | C | A | 6.32E-05 | | | | | ORF1ab | nonsynonymous_S6799Y |
| 20661 | T | C | 6.32E-05 | | | | | ORF1ab | synonymous_S6799S |
| 20663 | G | A | 6.32E-05 | | | | | ORF1ab | nonsynonymous_S6800N |
| 20665 | C | T | 6.32E-05 | | | | | ORF1ab | stopgain_Q6801X |
| 20691 | G | A | 6.32E-05 | | | | | ORF1ab | nonsynonymous_M6809I |
| 20696 | A | T | 6.32E-05 | | | | | ORF1ab | nonsynonymous_N6811I |
| 20699 | T | C | 6.32E-05 | | | | | ORF1ab | nonsynonymous_L6812P |
| 20700 | T | A | 6.32E-05 | | | | | ORF1ab | synonymous_L6812L |
| 20704 | A | C | 6.32E-05 | | | | | ORF1ab | nonsynonymous_K6814Q |
| 20706 | A | G | 6.32E-05 | | | | | ORF1ab | synonymous_K6814K |
| 20708 | T | C | 6.32E-05 | | | | | ORF1ab | nonsynonymous_M6815T |
| 20711 | A | G | 6.32E-05 | | | | | ORF1ab | nonsynonymous_Q6816R |
| 20714 | G | A | 6.32E-05 | | | | | ORF1ab | nonsynonymous_R6817K |
| 20715 | A | C | 6.32E-05 | | | | | ORF1ab | nonsynonymous_R6817S |
| 20717 | T | G | 6.32E-05 | | | | | ORF1ab | nonsynonymous_M6818R |
| 20721 | A | C | 6.32E-05 | | | | | ORF1ab | synonymous_L6819L |
| 20722 | T | A | 6.32E-05 | | | | | ORF1ab | nonsynonymous_L6820I |
| 20724 | A | C | 6.32E-05 | | | | | ORF1ab | nonsynonymous_L6820F |
| 20728 | A | G | 6.32E-05 | | | | | ORF1ab | nonsynonymous_K6822E |
| 20729 | A | T | 6.32E-05 | | | | | ORF1ab | nonsynonymous_K6822M |
| 20747 | A | G | 6.32E-05 | | | | | ORF1ab | nonsynonymous_Y6828C |

| | | | | | | | | | |
|---|---|---|---|---|---|---|---|---|---|
| 20749 | G | T | 6.32E-05 | | | | | ORF1ab | nonsynonymous_G6829C |
| 20750 | G | C | 6.32E-05 | | | | | ORF1ab | nonsynonymous_G6829A |
| 20752 | G | A | 6.32E-05 | A | 6.32E-05 | | | ORF1ab | nonsynonymous_D6830N; nonsynonymous_D6830N |
| 20753 | A | T | 6.32E-05 | | | | | ORF1ab | nonsynonymous_D6830V |
| 20758 | G | A | 6.32E-05 | | | | | ORF1ab | nonsynonymous_A6832T |
| 20760 | A | T | 6.32E-05 | | | | | ORF1ab | synonymous_A6832A |
| 20761 | A | T | 6.32E-05 | | | | | ORF1ab | nonsynonymous_T6833S |
| 20767 | C | T | 6.32E-05 | | | | | ORF1ab | nonsynonymous_P6835S |
| 20768 | C | T | 6.32E-05 | | | | | ORF1ab | nonsynonymous_P6835L |
| 20770 | A | G | 6.32E-05 | | | | | ORF1ab | nonsynonymous_K6836E |
| 20772 | A | T | 6.32E-05 | | | | | ORF1ab | nonsynonymous_K6836N |
| 20773 | G | A | 6.32E-05 | A | 6.32E-05 | | | ORF1ab | nonsynonymous_G6837S; nonsynonymous_G6837S |
| 20778 | A | T | 6.32E-05 | | | | | ORF1ab | synonymous_I6838I |
| 20779 | A | T | 6.32E-05 | | | | | ORF1ab | nonsynonymous_M6839L |
| 20782 | A | T | 6.32E-05 | | | | | ORF1ab | nonsynonymous_M6840L |
| 20788 | G | A | 6.32E-05 | | | | | ORF1ab | nonsynonymous_V6842I |
| 20791 | G | A | 6.32E-05 | | | | | ORF1ab | nonsynonymous_A6843T |
| 20792 | C | T | 6.32E-05 | | | | | ORF1ab | nonsynonymous_A6843V |
| 20793 | A | T | 6.32E-05 | | | | | ORF1ab | synonymous_A6843A |
| 20833 | T | C | 6.32E-05 | | | | | ORF1ab | synonymous_L6857L |
| 20946 | C | T | 6.32E-05 | | | | | ORF1ab | synonymous_V6894V |
| 20962 | G | A | 6.32E-05 | | | | | ORF1ab | nonsynonymous_D6900N |
| 20987 | C | T | 6.32E-05 | | | | | ORF1ab | nonsynonymous_T6908I |
| 21004 | G | T | 6.32E-05 | | | | | ORF1ab | nonsynonymous_A6914S |
| 21024 | T | C | 6.32E-05 | | | | | ORF1ab | synonymous_N6920N |
| 21037 | A | G | 6.32E-05 | | | | | ORF1ab | nonsynonymous_I6925V |
| 21040 | A | G | 6.32E-05 | | | | | ORF1ab | nonsynonymous_I6926V |
| 21043 | A | G | 6.32E-05 | | | | | ORF1ab | nonsynonymous_S6927G |

| | | | | | | | | | |
|---|---|---|---|---|---|---|---|---|---|
| 21111 | T | G | 6.32E-05 | | | | | ORF1ab | synonymous_T6949T |
| 21139 | C | T | 6.32E-05 | | | | | ORF1ab | synonymous_L6959L |
| 21141 | A | G | 6.32E-05 | | | | | ORF1ab | synonymous_L6959L |
| 21142 | G | T | 6.32E-05 | | | | | ORF1ab | nonsynonymous_A6960S |
| 21144 | T | C | 6.32E-05 | | | | | ORF1ab | synonymous_A6960A |
| 2143 | C | A | 6.32E-05 | | | | | ORF1ab | synonymous_P626P |
| 2160 | A | G | 6.32E-05 | | | | | ORF1ab | nonsynonymous_E632G |
| 21703 | T | A | 6.32E-05 | | | | | S | synonymous_V47V |
| 21709 | T | C | 6.32E-05 | | | | | S | synonymous_H49H |
| 21809 | G | T | 6.32E-05 | | | | | S | nonsynonymous_V83F |
| 21817 | A | T | 6.32E-05 | | | | | S | synonymous_P85P |
| 21836 | T | C | 6.32E-05 | | | | | S | nonsynonymous_F92L |
| 21839 | G | T | 6.32E-05 | | | | | S | nonsynonymous_A93S |
| 21851 | A | G | 6.32E-05 | | | | | S | nonsynonymous_K97E |
| 21853 | G | A | 6.32E-05 | | | | | S | synonymous_K97K |
| 21868 | A | T | 6.32E-05 | | | | | S | nonsynonymous_R102S |
| 21872 | T | A | 6.32E-05 | | | | | S | nonsynonymous_W104R |
| 21883 | T | C | 6.32E-05 | | | | | S | synonymous_G107G |
| 21891 | T | A | 6.32E-05 | | | | | S | stopgain_L110X |
| 21896 | T | C | 6.32E-05 | | | | | S | nonsynonymous_S112P |
| 21898 | G | A | 6.32E-05 | | | | | S | synonymous_S112S |
| 21910 | C | T | 6.32E-05 | | | | | S | synonymous_S116S |
| 21914 | C | T | 6.32E-05 | | | | | S | nonsynonymous_L118F |
| 21931 | T | C | 6.32E-05 | | | | | S | synonymous_A123A |
| 21935 | A | C | 6.32E-05 | | | | | S | nonsynonymous_N125H |
| 21944 | A | T | 6.32E-05 | | | | | S | nonsynonymous_I128F |
| 21968 | T | C | 6.32E-05 | | | | | S | nonsynonymous_C136R |
| 21973 | T | C | 6.32E-05 | | | | | S | synonymous_N137N |
| 22005 | A | G | 6.32E-05 | | | | | S | nonsynonymous_N148S |
| 22006 | C | A | 6.32E-05 | | | | | S | nonsynonymous_N148K |

| | | | | | | | | | |
|---|---|---|---|---|---|---|---|---|---|
| 22007 | A | C | 6.32E-05 | | | | | S | nonsynonymous_N149H |
| 22551 | C | T | 6.32E-05 | | | | | S | nonsynonymous_P330L |
| 22604 | G | A | 6.32E-05 | | | | | S | nonsynonymous_A348T |
| 22642 | C | T | 6.32E-05 | | | | | S | synonymous_N360N |
| 22695 | A | G | 6.32E-05 | | | | | S | nonsynonymous_K378R |
| 22698 | G | T | 6.32E-05 | | | | | S | nonsynonymous_C379F |
| 22739 | A | C | 6.32E-05 | | | | | S | nonsynonymous_T393P |
| 22745 | G | A | 6.32E-05 | | | | | S | nonsynonymous_V395I |
| 22757 | T | C | 6.32E-05 | | | | | S | nonsynonymous_S399P |
| 22776 | A | T | 6.32E-05 | | | | | S | nonsynonymous_D405V |
| 22780 | A | G | 6.32E-05 | | | | | S | synonymous_E406E |
| 22785 | G | T | 6.32E-05 | | | | | S | nonsynonymous_R408I |
| 22787 | C | G | 6.32E-05 | | | | | S | nonsynonymous_Q409E |
| 22792 | C | A | 6.32E-05 | A | 6.32E-05 | | | S | synonymous_I410I; synonymous_I410I |
| 22795 | T | C | 6.32E-05 | | | | | S | synonymous_A411A |
| 22798 | A | G | 6.32E-05 | | | | | S | synonymous_P412P |
| 22801 | G | T | 6.32E-05 | | | | | S | synonymous_G413G |
| 22811 | A | T | 6.32E-05 | | | | | S | stopgain_K417X |
| 22813 | G | T | 6.32E-05 | | | | | S | nonsynonymous_K417N |
| 22855 | C | A | 6.32E-05 | | | | | S | synonymous_G431G |
| 22861 | T | A | 6.32E-05 | A | 6.32E-05 | | | S | synonymous_V433V; synonymous_V433V |
| 22863 | T | A | 6.32E-05 | | | | | S | nonsynonymous_I434K |
| 2288 | G | A | 6.32E-05 | | | | | ORF1ab | nonsynonymous_V675I |
| 23147 | A | G | 6.32E-05 | | | | | S | nonsynonymous_K529E |
| 23149 | G | C | 6.32E-05 | | | | | S | nonsynonymous_K529N |
| 23173 | A | G | 6.32E-05 | | | | | S | synonymous_K537K |
| 23182 | T | C | 6.32E-05 | | | | | S | synonymous_N540N |
| 23224 | G | T | 6.32E-05 | | | | | S | nonsynonymous_E554D |

| | | | | | | | | | |
|---|---|---|---|---|---|---|---|---|---|
| 23229 | A | G | 6.32E-05 | | | | | S | nonsynonymous_N556S |
| 23235 | A | G | 6.32E-05 | | | | | S | nonsynonymous_K558R |
| 23236 | G | T | 6.32E-05 | | | | | S | nonsynonymous_K558N |
| 23242 | G | T | 6.32E-05 | | | | | S | synonymous_L560L |
| 23244 | C | T | 6.32E-05 | | | | | S | nonsynonymous_P561L |
| 23254 | A | G | 6.32E-05 | | | | | S | synonymous_Q564Q |
| 23255 | T | C | 6.32E-05 | | | | | S | nonsynonymous_F565L |
| 23269 | T | A | 6.32E-05 | | | | | S | synonymous_I569I |
| 23270 | G | A | 6.32E-05 | | | | | S | nonsynonymous_A570T |
| 23272 | T | C | 6.32E-05 | | | | | S | synonymous_A570A |
| 23281 | T | C | 6.32E-05 | | | | | S | synonymous_T573T |
| 23320 | C | T | 6.32E-05 | | | | | S | synonymous_D586D |
| 23335 | T | A | 6.32E-05 | | | | | S | synonymous_S591S |
| 23347 | C | T | 6.32E-05 | | | | | S | synonymous_V595V |
| 23348 | A | C | 6.32E-05 | | | | | S | nonsynonymous_S596R |
| 23349 | G | C | 6.32E-05 | | | | | S | nonsynonymous_S596T |
| 23356 | A | T | 6.32E-05 | | | | | S | synonymous_I598I |
| 23367 | C | T | 6.32E-05 | | | | | S | nonsynonymous_T602I |
| 23371 | T | A | 6.32E-05 | | | | | S | nonsynonymous_N603K |
| 23380 | C | T | 6.32E-05 | | | | | S | synonymous_N606N |
| 23392 | T | C | 6.32E-05 | | | | | S | synonymous_V610V |
| 23405 | G | T | 6.32E-05 | T | 6.32E-05 | | | S | nonsynonymous_V615F; nonsynonymous_V615F |
| 23413 | C | T | 6.32E-05 | | | | | S | synonymous_C617C |
| 23416 | A | T | 6.32E-05 | | | | | S | synonymous_T618T |
| 23419 | A | G | 6.32E-05 | | | | | S | synonymous_E619E |
| 23427 | T | C | 6.32E-05 | | | | | S | nonsynonymous_V622A |
| 23437 | T | A | 6.32E-05 | | | | | S | nonsynonymous_H625Q |
| 23440 | A | G | 6.32E-05 | | | | | S | synonymous_A626A |
| 23447 | C | G | 6.32E-05 | | | | | S | nonsynonymous_L629V |

| | | | | | | | | | |
|---|---|---|---|---|---|---|---|---|---|
| 23450 | A | T | 6.32E-05 | | | | | S | nonsynonymous_T630S |
| 23469 | A | T | 6.32E-05 | | | | | S | nonsynonymous_Y636F |
| 23480 | T | G | 6.32E-05 | | | | | S | nonsynonymous_S640A |
| 23482 | T | C | 6.32E-05 | | | | | S | synonymous_S640S |
| 23487 | T | G | 6.32E-05 | | | | | S | nonsynonymous_V642G |
| 23489 | T | C | 6.32E-05 | | | | | S | nonsynonymous_F643L |
| 23491 | T | G | 6.32E-05 | | | | | S | nonsynonymous_F643L |
| 23503 | A | T | 6.32E-05 | | | | | S | synonymous_A647A |
| 23506 | C | T | 6.32E-05 | | | | | S | synonymous_G648G |
| 23522 | G | A | 6.32E-05 | A | 6.32E-05 | | | S | nonsynonymous_E654K; nonsynonymous_E654K |
| 23538 | C | T | 6.32E-05 | | | | | S | nonsynonymous_S659L |
| 23558 | A | G | 6.32E-05 | | | | | S | nonsynonymous_I666V |
| 23561 | G | A | 6.32E-05 | | | | | S | nonsynonymous_G667S |
| 23628 | G | T | 6.32E-05 | | | | | S | nonsynonymous_S689I |
| 23634 | C | T | 6.32E-05 | | | | | S | nonsynonymous_S691F |
| 23691 | A | C | 6.32E-05 | | | | | S | nonsynonymous_N710T |
| 23712 | A | C | 6.32E-05 | | | | | S | nonsynonymous_N717T |
| 23717 | A | G | 6.32E-05 | | | | | S | nonsynonymous_T719A |
| 23720 | A | G | 6.32E-05 | | | | | S | nonsynonymous_I720V |
| 23730 | C | T | 6.32E-05 | | | | | S | nonsynonymous_T723I |
| 23733 | C | T | 6.32E-05 | | | | | S | nonsynonymous_T724I |
| 23734 | A | T | 6.32E-05 | | | | | S | synonymous_T724T |
| 23741 | C | T | 6.32E-05 | | | | | S | synonymous_L727L |
| 23749 | G | T | 6.32E-05 | | | | | S | synonymous_V729V |
| 23785 | C | T | 6.32E-05 | | | | | S | synonymous_Y741Y |
| 23801 | A | G | 6.32E-05 | | | | | S | nonsynonymous_T747A |
| 23813 | A | G | 6.32E-05 | | | | | S | nonsynonymous_N751D |
| 23844 | C | T | 6.32E-05 | | | | | S | nonsynonymous_T761I |
| 23851 | A | G | 6.32E-05 | | | | | S | synonymous_L763L |

| | | | | | | | | | |
|---|---|---|---|---|---|---|---|---|---|
| 23854 | C | T | 6.32E-05 | | | | | S | synonymous_N764N |
| 23865 | C | T | 6.32E-05 | | | | | S | nonsynonymous_T768I |
| 23867 | G | A | 6.32E-05 | | | | | S | nonsynonymous_G769R |
| 23884 | A | G | 6.32E-05 | | | | | S | synonymous_Q774Q |
| 23895 | C | A | 6.32E-05 | A | 6.32E-05 | | | S | nonsynonymous_T778N; nonsynonymous_T778N |
| 23900 | G | C | 6.32E-05 | | | | | S | nonsynonymous_E780Q |
| 23901 | A | T | 6.32E-05 | | | | | S | nonsynonymous_E780V |
| 23911 | A | G | 6.32E-05 | | | | | S | synonymous_A783A |
| 23952 | T | G | 6.32E-05 | | | | | S | nonsynonymous_F797C |
| 23972 | C | A | 6.32E-05 | | | | | S | nonsynonymous_Q804K |
| 23989 | A | G | 6.32E-05 | | | | | S | synonymous_P809P |
| 23992 | A | T | 6.32E-05 | | | | | S | synonymous_S810S |
| 24010 | A | C | 6.32E-05 | | | | | S | synonymous_S816S |
| 24011 | T | C | 6.32E-05 | | | | | S | nonsynonymous_F817L |
| 24014 | A | G | 6.32E-05 | | | | | S | nonsynonymous_I818V |
| 24042 | C | T | 6.32E-05 | | | | | S | nonsynonymous_T827I |
| 2404 | A | T | 6.32E-05 | | | | | ORF1ab | synonymous_S713S |
| 24064 | C | T | 6.32E-05 | | | | | S | synonymous_I834I |
| 2406 | A | G | 6.32E-05 | | | | | ORF1ab | nonsynonymous_K714R |
| 24071 | T | C | 6.32E-05 | | | | | S | nonsynonymous_Y837H |
| 24075 | G | A | 6.32E-05 | A | 6.32E-05 | | | S | nonsynonymous_G838D; nonsynonymous_G838D |
| 24083 | C | A | 6.32E-05 | | | | | S | nonsynonymous_L841I |
| 24086 | G | A | 6.32E-05 | | | | | S | nonsynonymous_G842S |
| 24140 | G | T | 6.32E-05 | T | 6.32E-05 | | | S | nonsynonymous_V860F; nonsynonymous_V860F |
| 24141 | T | A | 6.32E-05 | | | | | S | nonsynonymous_V860D |
| 24142 | T | A | 6.32E-05 | | | | | S | synonymous_V860V |
| 24143 | T | A | 6.32E-05 | | | | | S | nonsynonymous_L861M |

| | | | | | | | | | |
|---|---|---|---|---|---|---|---|---|---|
| 24144 | T | A | 6.32E-05 | | | | | S | stopgain_L861X |
| 24151 | T | C | 6.32E-05 | | | | | S | synonymous_P863P |
| 24152 | T | C | 6.32E-05 | | | | | S | synonymous_L864L |
| 24156 | T | C | 6.32E-05 | | | | | S | nonsynonymous_L865P |
| 24171 | T | G | 6.32E-05 | | | | | S | nonsynonymous_I870S |
| 24174 | C | T | 6.32E-05 | | | | | S | nonsynonymous_A871V |
| 24405 | T | G | 6.32E-05 | | | | | S | nonsynonymous_L948R |
| 24422 | C | A | 6.32E-05 | | | | | S | nonsynonymous_Q954K |
| 24439 | A | G | 6.32E-05 | | | | | S | synonymous_L959L |
| 24735 | A | T | 6.32E-05 | | | | | S | nonsynonymous_H1058L |
| 24749 | T | C | 6.32E-05 | | | | | S | synonymous_L1063L |
| 24759 | C | A | 6.32E-05 | | | | | S | nonsynonymous_T1066N |
| 24762 | A | T | 6.32E-05 | | | | | S | nonsynonymous_Y1067F |
| 25036 | T | C | 6.32E-05 | | | | | S | synonymous_N1158N |
| 25038 | A | T | 6.32E-05 | | | | | S | nonsynonymous_H1159L |
| 25039 | T | C | 6.32E-05 | | | | | S | synonymous_H1159H |
| 25320 | G | T | 6.32E-05 | | | | | S | nonsynonymous_C1253F |
| 25347 | A | G | 6.32E-05 | | | | | S | nonsynonymous_E1262G |
| 25356 | T | C | 6.32E-05 | | | | | S | nonsynonymous_L1265P |
| 25366 | C | T | 6.32E-05 | | | | | S | synonymous_V1268V |
| 25367 | A | C | 6.32E-05 | | | | | S | nonsynonymous_K1269Q |
| 25368 | A | G | 6.32E-05 | | | | | S | nonsynonymous_K1269R |
| 25373 | C | G | 6.32E-05 | | | | | S | nonsynonymous_H1271D |
| 25377 | A | T | 6.32E-05 | | | | | S | nonsynonymous_Y1272F |
| 25379 | A | T | 6.32E-05 | | | | | S | nonsynonymous_T1273S |
| 25380 | C | T | 6.32E-05 | | | | | S | nonsynonymous_T1273I |
| 25382 | T | C | 6.32E-05 | | | | | S | stoploss_X1274Q |
| 25386 | C | T | 6.32E-05 | | | | | intergenic | |
| 25394 | T | C | 6.32E-05 | | | | | ORF3a | nonsynonymous_M1T |
| 25407 | G | T | 6.32E-05 | | | | | ORF3a | nonsynonymous_M5I |

| | | | | | | | | | |
|---|---|---|---|---|---|---|---|---|---|
| 25410 | A | C | 6.32E-05 | | | | | ORF3a | nonsynonymous_R6S |
| 25430 | T | C | 6.32E-05 | | | | | ORF3a | nonsynonymous_V13A |
| 25436 | T | A | 6.32E-05 | | | | | ORF3a | stopgain_L15X |
| 25438 | A | C | 6.32E-05 | | | | | ORF3a | nonsynonymous_K16Q |
| 25439 | A | T | 6.32E-05 | | | | | ORF3a | nonsynonymous_K16M |
| 25443 | A | C | 6.32E-05 | | | | | ORF3a | nonsynonymous_Q17H |
| 25445 | G | T | 6.32E-05 | | | | | ORF3a | nonsynonymous_G18V |
| 25448 | A | C | 6.32E-05 | | | | | ORF3a | nonsynonymous_E19A |
| 25454 | A | G | 6.32E-05 | | | | | ORF3a | nonsynonymous_K21R |
| 25464 | T | C | 6.32E-05 | | | | | ORF3a | synonymous_T24T |
| 25468 | T | C | 6.32E-05 | | | | | ORF3a | nonsynonymous_S26P |
| 25476 | T | C | 6.32E-05 | | | | | ORF3a | synonymous_F28F |
| 25482 | C | T | 6.32E-05 | | | | | ORF3a | synonymous_R30R |
| 25489 | G | T | 6.32E-05 | | | | | ORF3a | nonsynonymous_A33S |
| 25536 | T | C | 6.32E-05 | | | | | ORF3a | synonymous_V48V |
| 25538 | G | A | 6.32E-05 | | | | | ORF3a | nonsynonymous_G49D |
| 25937 | A | G | 6.32E-05 | | | | | ORF3a | nonsynonymous_H182R |
| 25938 | T | C | 6.32E-05 | | | | | ORF3a | synonymous_H182H |
| 25962 | T | G | 6.32E-05 | | | | | ORF3a | synonymous_T190T |
| 25964 | A | G | 6.32E-05 | | | | | ORF3a | nonsynonymous_E191G |
| 25965 | A | C | 6.32E-05 | | | | | ORF3a | nonsynonymous_E191D |
| 25971 | G | T | 6.32E-05 | | | | | ORF3a | nonsynonymous_W193C |
| 25974 | A | G | 6.32E-05 | | | | | ORF3a | synonymous_E194E |
| 25983 | A | G | 6.32E-05 | | | | | ORF3a | synonymous_V197V |
| 25989 | C | T | 6.32E-05 | | | | | ORF3a | synonymous_D199D |
| 25993 | G | A | 6.32E-05 | | | | | ORF3a | nonsynonymous_V201I |
| 26006 | G | C | 6.32E-05 | | | | | ORF3a | nonsynonymous_S205T |
| 26009 | A | G | 6.32E-05 | | | | | ORF3a | nonsynonymous_Y206C |
| 26020 | G | T | 6.32E-05 | | | | | ORF3a | nonsynonymous_D210Y |
| 26031 | G | T | 6.32E-05 | | | | | ORF3a | nonsynonymous_Q213H |

| | | | | | | | | | |
|---|---|---|---|---|---|---|---|---|---|
| 26038 | T | C | 6.32E-05 | | | | | ORF3a | nonsynonymous_S216P |
| 26048 | T | G | 6.32E-05 | | | | | ORF3a | nonsynonymous_L219W |
| 26055 | A | G | 6.32E-05 | | | | | ORF3a | synonymous_T221T |
| 26056 | G | A | 6.32E-05 | | | | | ORF3a | nonsynonymous_D222N |
| 26063 | G | T | 6.32E-05 | | | | | ORF3a | nonsynonymous_G224V |
| 26065 | G | T | 6.32E-05 | | | | | ORF3a | nonsynonymous_V225F |
| 26084 | T | C | 6.32E-05 | | | | | ORF3a | nonsynonymous_F231S |
| 26101 | G | T | 6.32E-05 | | | | | ORF3a | nonsynonymous_V237F |
| 26106 | T | G | 6.32E-05 | | | | | ORF3a | nonsynonymous_D238E |
| 26108 | A | T | 6.32E-05 | | | | | ORF3a | nonsynonymous_E239V |
| 26114 | A | T | 6.32E-05 | | | | | ORF3a | nonsynonymous_E241V |
| 26116 | G | A | 6.32E-05 | A | 6.32E-05 | | | ORF3a | nonsynonymous_E242K; nonsynonymous_E242K |
| 26131 | C | T | 6.32E-05 | | | | | ORF3a | nonsynonymous_H247Y |
| 26141 | A | T | 6.32E-05 | | | | | ORF3a | nonsynonymous_D250V |
| 26150 | C | A | 6.32E-05 | A | 6.32E-05 | | | ORF3a | nonsynonymous_S253Y; nonsynonymous_S253Y |
| 26171 | T | A | 6.32E-05 | | | | | ORF3a | nonsynonymous_M260K |
| 26177 | C | T | 6.32E-05 | | | | | ORF3a | nonsynonymous_P262L |
| 26181 | T | G | 6.32E-05 | | | | | ORF3a | nonsynonymous_I263M |
| 26191 | C | T | 6.32E-05 | | | | | ORF3a | nonsynonymous_P267S |
| 26192 | C | T | 6.32E-05 | | | | | ORF3a | nonsynonymous_P267L |
| 26196 | G | A | 6.32E-05 | A | 6.32E-05 | | | ORF3a | synonymous_T268T; synonymous_T268T |
| 26208 | C | T | 6.32E-05 | | | | | ORF3a | synonymous_S272S |
| 26225 | A | G | 6.32E-05 | | | | | intergenic | |
| 26230 | G | T | 6.32E-05 | | | | | intergenic | |
| 26234 | A | G | 6.32E-05 | | | | | intergenic | |
| 26261 | C | T | 6.32E-05 | | | | | E | nonsynonymous_S6L |
| 26265 | A | T | 6.32E-05 | | | | | E | nonsynonymous_E7D |

| | | | | | | | | | |
|---|---|---|---|---|---|---|---|---|---|
| 26266 | G | A | 6.32E-05 | | | | | E | nonsynonymous_E8K |
| 26272 | G | T | 6.32E-05 | | | | | E | nonsynonymous_G10C |
| 26274 | T | C | 6.32E-05 | | | | | E | synonymous_G10G |
| 26294 | T | C | 6.32E-05 | | | | | E | nonsynonymous_V17A |
| 26301 | T | G | 6.32E-05 | G | 6.32E-05 | | | E | synonymous_L19L; synonymous_L19L |
| 26307 | T | A | 6.32E-05 | | | | | E | synonymous_L21L |
| 26308 | G | T | 6.32E-05 | | | | | E | nonsynonymous_A22S |
| 26351 | C | T | 6.32E-05 | | | | | E | nonsynonymous_A36V |
| 26353 | C | T | 6.32E-05 | | | | | E | nonsynonymous_L37F |
| 26354 | T | A | 6.32E-05 | | | | | E | nonsynonymous_L37H |
| 26358 | A | C | 6.32E-05 | | | | | E | synonymous_R38R |
| 26557 | A | G | 6.32E-05 | | | | | M | nonsynonymous_E12G |
| 26570 | C | T | 6.32E-05 | | | | | M | synonymous_L16L |
| 26573 | T | C | 6.32E-05 | | | | | M | synonymous_L17L |
| 26593 | T | C | 6.32E-05 | | | | | M | nonsynonymous_I24T |
| 26598 | T | C | 6.32E-05 | | | | | M | nonsynonymous_F26L |
| 26601 | C | T | 6.32E-05 | | | | | M | synonymous_L27L |
| 26607 | C | T | 6.32E-05 | | | | | M | nonsynonymous_L29F |
| 26609 | T | C | 6.32E-05 | | | | | M | synonymous_L29L |
| 26612 | A | T | 6.32E-05 | T | 6.32E-05 | | | M | synonymous_T30T; synonymous_T30T |
| 26620 | G | T | 6.32E-05 | | | | | M | nonsynonymous_C33F |
| 26631 | T | A | 6.32E-05 | | | | | M | nonsynonymous_F37I |
| 26636 | C | T | 6.32E-05 | | | | | M | synonymous_A38A |
| 26641 | C | T | 6.32E-05 | | | | | M | nonsynonymous_A40V |
| 26650 | A | G | 6.32E-05 | | | | | M | nonsynonymous_N43S |
| 26651 | T | A | 6.32E-05 | | | | | M | nonsynonymous_N43K |
| 26679 | T | C | 6.32E-05 | | | | | M | nonsynonymous_F53L |
| 26693 | A | G | 6.32E-05 | | | | | M | synonymous_L57L |

| | | | | | | | | | |
|---|---|---|---|---|---|---|---|---|---|
| 26718 | G | T | 6.32E-05 | | | | | M | nonsynonymous_V66L |
| 26721 | C | T | 6.32E-05 | | | | | M | nonsynonymous_L67F |
| 26723 | T | A | 6.32E-05 | | | | | M | synonymous_L67L |
| 26726 | T | C | 6.32E-05 | | | | | M | synonymous_A68A |
| 26728 | C | T | 6.32E-05 | | | | | M | nonsynonymous_A69V |
| 26732 | T | C | 6.32E-05 | | | | | M | synonymous_V70V |
| 26755 | G | C | 6.32E-05 | | | | | M | nonsynonymous_G78A |
| 26756 | T | A | 6.32E-05 | | | | | M | synonymous_G78G |
| 26759 | A | T | 6.32E-05 | | | | | M | synonymous_G79G |
| 26761 | T | C | 6.32E-05 | | | | | M | nonsynonymous_I80T |
| 26774 | G | T | 6.32E-05 | | | | | M | nonsynonymous_M84I |
| 26822 | C | T | 6.32E-05 | | | | | M | synonymous_F100F |
| 26823 | A | C | 6.32E-05 | | | | | M | synonymous_R101R |
| 26833 | C | T | 6.32E-05 | | | | | M | nonsynonymous_A104V |
| 26839 | C | T | 6.32E-05 | | | | | M | nonsynonymous_T106M |
| 26847 | A | G | 6.32E-05 | | | | | M | nonsynonymous_M109V |
| 26871 | A | C | 6.32E-05 | | | | | M | nonsynonymous_N117H |
| 26902 | C | A | 6.32E-05 | | | | | M | nonsynonymous_T127N |
| 27215 | T | C | 6.32E-05 | | | | | ORF6 | nonsynonymous_V5A |
| 27223 | C | T | 6.32E-05 | | | | | ORF6 | stopgain_Q8X |
| 27226 | G | T | 6.32E-05 | | | | | ORF6 | nonsynonymous_V9F |
| 27228 | T | A | 6.32E-05 | | | | | ORF6 | synonymous_V9V |
| 27230 | C | T | 6.32E-05 | | | | | ORF6 | nonsynonymous_T10I |
| 27232 | A | G | 6.32E-05 | | | | | ORF6 | nonsynonymous_I11V |
| 27235 | G | A | 6.32E-05 | | | | | ORF6 | nonsynonymous_A12T |
| 27246 | A | C | 6.32E-05 | | | | | ORF6 | nonsynonymous_L15F |
| 27248 | T | C | 6.32E-05 | | | | | ORF6 | nonsynonymous_L16P |
| 27255 | T | G | 6.32E-05 | | | | | ORF6 | nonsynonymous_I18M |
| 27259 | A | G | 6.32E-05 | | | | | ORF6 | nonsynonymous_R20G |
| 27302 | A | G | 6.32E-05 | | | | | ORF6 | nonsynonymous_N34S |

| | | | | | | | | | |
|---|---|---|---|---|---|---|---|---|---|
| 27317 | A | G | 6.32E-05 | | | | | ORF6 | nonsynonymous_N39S |
| 27322 | T | G | 6.32E-05 | | | | | ORF6 | nonsynonymous_S41A |
| 27340 | A | C | 6.32E-05 | | | | | ORF6 | nonsynonymous_N47H |
| 27342 | T | G | 6.32E-05 | | | | | ORF6 | nonsynonymous_N47K |
| 27344 | A | G | 6.32E-05 | | | | | ORF6 | nonsynonymous_K48R |
| 27350 | C | T | 6.32E-05 | | | | | ORF6 | nonsynonymous_S50F |
| 27354 | A | G | 6.32E-05 | | | | | ORF6 | synonymous_Q51Q |
| 27356 | T | C | 6.32E-05 | | | | | ORF6 | nonsynonymous_L52S |
| 27363 | A | G | 6.32E-05 | G | 6.32E-05 | | | ORF6 | synonymous_E54E; synonymous_E54E |
| 27365 | A | T | 6.32E-05 | | | | | ORF6 | nonsynonymous_E55V |
| 27366 | G | T | 6.32E-05 | T | 6.32E-05 | | | ORF6 | nonsynonymous_E55D; nonsynonymous_E55D |
| 27367 | C | G | 6.32E-05 | | | | | ORF6 | nonsynonymous_Q56E |
| 27369 | A | T | 6.32E-05 | | | | | ORF6 | nonsynonymous_Q56H |
| 27419 | T | C | 6.32E-05 | | | | | ORF7a | nonsynonymous_L9P |
| 27421 | A | G | 6.32E-05 | G | 6.32E-05 | | | ORF7a | nonsynonymous_I10V; nonsynonymous_I10V |
| 27459 | G | T | 6.32E-05 | | | | | ORF7a | nonsynonymous_E22D |
| 27461 | G | T | 6.32E-05 | | | | | ORF7a | nonsynonymous_C23F |
| 27829 | T | C | 6.32E-05 | | | | | ORF7b | nonsynonymous_L25P |
| 27832 | T | C | 6.32E-05 | | | | | ORF7b | nonsynonymous_I26T |
| 27838 | T | A | 6.32E-05 | | | | | ORF7b | nonsynonymous_F28Y |
| 27842 | G | T | 6.32E-05 | | | | | ORF7b | nonsynonymous_W29C |
| 27844 | T | G | 6.32E-05 | | | | | ORF7b | nonsynonymous_F30C |
| 28256 | C | T | 6.32E-05 | | | | | ORF8 | synonymous_I121I |
| 28280 | G | A | 6.32E-05 | A | 6.32E-05 | A | 6.32E-05 | N | nonsynonymous_D3N; nonsynonymous_D3N; nonsynonymous_D3N |
| 28281 | A | T | 6.32E-05 | | | | | N | nonsynonymous_D3V |

| | | | | | | | | | |
|---|---|---|---|---|---|---|---|---|---|
| 28282 | T | A | 6.32E-05 | | | | | N | nonsynonymous_D3E |
| 28283 | A | G | 6.32E-05 | | | | | N | nonsynonymous_N4D |
| 28287 | G | T | 6.32E-05 | | | | | N | nonsynonymous_G5V |
| 28288 | A | G | 6.32E-05 | | | | | N | synonymous_G5G |
| 28291 | C | T | 6.32E-05 | | | | | N | synonymous_P6P |
| 28313 | C | T | 6.32E-05 | | | | | N | nonsynonymous_R14C |
| 28314 | G | A | 6.32E-05 | | | | | N | nonsynonymous_R14H |
| 28330 | A | G | 6.32E-05 | | | | | N | synonymous_G19G |
| 28332 | C | T | 6.32E-05 | | | | | N | nonsynonymous_P20L |
| 28337 | G | T | 6.32E-05 | | | | | N | nonsynonymous_D22Y |
| 28346 | G | T | 6.32E-05 | | | | | N | nonsynonymous_G25C |
| 28347 | G | T | 6.32E-05 | | | | | N | nonsynonymous_G25V |
| 28352 | A | T | 6.32E-05 | | | | | N | nonsynonymous_N27Y |
| 28354 | C | T | 6.32E-05 | | | | | N | synonymous_N27N |
| 28355 | C | G | 6.32E-05 | | | | | N | nonsynonymous_Q28E |
| 28356 | A | G | 6.32E-05 | | | | | N | nonsynonymous_Q28R |
| 28357 | G | C | 6.32E-05 | | | | | N | nonsynonymous_Q28H |
| 28359 | A | G | 6.32E-05 | | | | | N | nonsynonymous_N29S |
| 28458 | A | T | 6.32E-05 | | | | | N | nonsynonymous_E62V |
| 28483 | A | G | 6.32E-05 | | | | | N | synonymous_Q70Q |
| 28492 | A | G | 6.32E-05 | | | | | N | synonymous_P73P |
| 28507 | C | T | 6.32E-05 | | | | | N | synonymous_S78S |
| 28511 | C | A | 6.32E-05 | A | 6.32E-05 | | | N | nonsynonymous_P80T; nonsynonymous_P80T |
| 28516 | T | C | 6.32E-05 | | | | | N | synonymous_D81D |
| 28531 | C | T | 6.32E-05 | | | | | N | synonymous_Y86Y |
| 28543 | T | C | 6.32E-05 | | | | | N | synonymous_A90A |
| 28557 | G | T | 6.32E-05 | | | | | N | nonsynonymous_R95L |
| 28572 | A | G | 6.32E-05 | | | | | N | nonsynonymous_K100R |
| 28576 | G | A | 6.32E-05 | | | | | N | nonsynonymous_M101I |

| | | | | | | | | | |
|---|---|---|---|---|---|---|---|---|---|
| 28610 | C | A | 6.32E-05 | | | | | N | nonsynonymous_L113I |
| 28622 | C | T | 6.32E-05 | | | | | N | nonsynonymous_P117S |
| 28627 | A | G | 6.32E-05 | | | | | N | synonymous_E118E |
| 28632 | G | C | 6.32E-05 | | | | | N | nonsynonymous_G120A |
| 28642 | T | C | 6.32E-05 | | | | | N | synonymous_Y123Y |
| 28663 | C | T | 6.32E-05 | | | | | N | synonymous_I130I |
| 28678 | T | A | 6.32E-05 | | | | | N | synonymous_T135T |
| 28695 | C | T | 6.32E-05 | | | | | N | nonsynonymous_T141I |
| 28696 | A | T | 6.32E-05 | | | | | N | synonymous_T141T |
| 28705 | T | G | 6.32E-05 | | | | | N | nonsynonymous_D144E |
| 28712 | G | A | 6.32E-05 | | | | | N | nonsynonymous_G147S |
| 28714 | C | T | 6.32E-05 | | | | | N | synonymous_G147G |
| 28716 | C | T | 6.32E-05 | | | | | N | nonsynonymous_T148I |
| 28732 | C | T | 6.32E-05 | | | | | N | synonymous_N153N |
| 28733 | A | T | 6.32E-05 | | | | | N | nonsynonymous_N154Y |
| 28741 | A | T | 6.32E-05 | | | | | N | synonymous_A156A |
| 28743 | T | C | 6.32E-05 | | | | | N | nonsynonymous_I157T |
| 28747 | G | T | 6.32E-05 | | | | | N | synonymous_V158V |
| 28752 | A | C | 6.32E-05 | | | | | N | nonsynonymous_Q160P |
| 29008 | T | C | 6.32E-05 | | | | | N | synonymous_T245T |
| 29010 | T | C | 6.32E-05 | | | | | N | nonsynonymous_V246A |
| 29012 | A | G | 6.32E-05 | | | | | N | nonsynonymous_T247A |
| 29017 | G | T | 6.32E-05 | | | | | N | nonsynonymous_K248N |
| 29019 | A | G | 6.32E-05 | | | | | N | nonsynonymous_K249R |
| 29025 | C | T | 6.32E-05 | | | | | N | nonsynonymous_A251V |
| 29027 | G | T | 6.32E-05 | | | | | N | nonsynonymous_A252S |
| 29031 | A | G | 6.32E-05 | | | | | N | nonsynonymous_E253G |
| 29033 | G | T | 6.32E-05 | | | | | N | nonsynonymous_A254S |
| 29037 | C | T | 6.32E-05 | | | | | N | nonsynonymous_S255F |
| 29323 | C | A | 6.32E-05 | | | | | N | synonymous_V350V |

| | | | | | | | | | |
|---|---|---|---|---|---|---|---|---|---|
| 29324 | A | G | 6.32E-05 | | | | | N | nonsynonymous_I351V |
| 29337 | A | G | 6.32E-05 | | | | | N | nonsynonymous_K355R |
| 29338 | G | T | 6.32E-05 | | | | | N | nonsynonymous_K355N |
| 29339 | C | G | 6.32E-05 | | | | | N | nonsynonymous_H356D |
| 29340 | A | C | 6.32E-05 | | | | | N | nonsynonymous_H356P |
| 29344 | T | C | 6.32E-05 | | | | | N | synonymous_I357I |
| 29345 | G | C | 6.32E-05 | | | | | N | nonsynonymous_D358H |
| 2973 | C | T | 6.32E-05 | | | | | ORF1ab | nonsynonymous_A903V |
| 3003 | A | G | 6.32E-05 | | | | | ORF1ab | nonsynonymous_E913G |
| 3014 | G | A | 6.32E-05 | | | | | ORF1ab | nonsynonymous_A917T |
| 3243 | G | T | 6.32E-05 | | | | | ORF1ab | nonsynonymous_G993V |
| 3328 | G | T | 6.32E-05 | | | | | ORF1ab | nonsynonymous_Q1021H |
| 3331 | T | C | 6.32E-05 | | | | | ORF1ab | synonymous_T1022T |
| 3426 | C | T | 6.32E-05 | | | | | ORF1ab | nonsynonymous_P1054L |
| 346 | C | T | 6.32E-05 | | | | | ORF1ab | synonymous_L27L |
| 3476 | G | A | 6.32E-05 | | | | | ORF1ab | nonsynonymous_V1071I |
| 3551 | C | A | 6.32E-05 | | | | | ORF1ab | nonsynonymous_P1096T |
| 355 | C | T | 6.32E-05 | | | | | ORF1ab | synonymous_G30G |
| 3638 | G | T | 6.32E-05 | | | | | ORF1ab | nonsynonymous_G1125C |
| 368 | G | A | 6.32E-05 | A | 6.32E-05 | | | ORF1ab | nonsynonymous_V35M; nonsynonymous_V35M |
| 3778 | A | G | 6.32E-05 | | | | | ORF1ab | synonymous_T1171T |
| 3830 | A | G | 6.32E-05 | | | | | ORF1ab | nonsynonymous_S1189G |
| 3831 | G | A | 6.32E-05 | | | | | ORF1ab | nonsynonymous_S1189N |
| 3832 | C | T | 6.32E-05 | | | | | ORF1ab | synonymous_S1189S |
| 3845 | A | G | 6.32E-05 | | | | | ORF1ab | nonsynonymous_K1194E |
| 3854 | A | G | 6.32E-05 | | | | | ORF1ab | nonsynonymous_K1197E |
| 3860 | G | A | 6.32E-05 | | | | | ORF1ab | nonsynonymous_V1199I |
| 3862 | T | A | 6.32E-05 | | | | | ORF1ab | synonymous_V1199V |
| 3863 | G | A | 6.32E-05 | | | | | ORF1ab | nonsynonymous_E1200K |

| | | | | | | | | | |
|---|---|---|---|---|---|---|---|---|---|
| 3872 | A | T | 6.32E-05 | | | | | ORF1ab | nonsynonymous_I1203F |
| 3877 | T | A | 6.32E-05 | | | | | ORF1ab | synonymous_A1204A |
| 3894 | A | G | 6.32E-05 | | | | | ORF1ab | nonsynonymous_E1210G |
| 3896 | G | T | 6.32E-05 | | | | | ORF1ab | nonsynonymous_V1211F |
| 3906 | T | C | 6.32E-05 | | | | | ORF1ab | nonsynonymous_F1214S |
| 3924 | C | T | 6.32E-05 | | | | | ORF1ab | nonsynonymous_P1220L |
| 4006 | G | A | 6.32E-05 | A | 6.32E-05 | | | ORF1ab | synonymous_K1247K; synonymous_K1247K |
| 4008 | T | C | 6.32E-05 | | | | | ORF1ab | nonsynonymous_F1248S |
| 4012 | C | T | 6.32E-05 | | | | | ORF1ab | synonymous_L1249L |
| 4017 | A | T | 6.32E-05 | | | | | ORF1ab | nonsynonymous_E1251V |
| 4060 | A | G | 6.32E-05 | | | | | ORF1ab | synonymous_P1265P |
| 4307 | A | C | 6.32E-05 | | | | | ORF1ab | nonsynonymous_K1348Q |
| 4316 | A | G | 6.32E-05 | | | | | ORF1ab | nonsynonymous_S1351G |
| 4455 | C | T | 6.32E-05 | | | | | ORF1ab | nonsynonymous_A1397V |
| 488 | G | A | 6.32E-05 | | | | | ORF1ab | nonsynonymous_D75N |
| 4900 | C | T | 6.32E-05 | | | | | ORF1ab | synonymous_H1545H |
| 5273 | G | A | 6.32E-05 | | | | | ORF1ab | nonsynonymous_A1670T |
| 5278 | T | C | 6.32E-05 | | | | | ORF1ab | synonymous_D1671D |
| 5281 | C | T | 6.32E-05 | | | | | ORF1ab | synonymous_N1672N |
| 558 | G | T | 6.32E-05 | | | | | ORF1ab | nonsynonymous_G98V |
| 580 | T | A | 6.32E-05 | | | | | ORF1ab | synonymous_G105G |
| 6151 | C | T | 6.32E-05 | | | | | ORF1ab | synonymous_D1962D |
| 6250 | C | T | 6.32E-05 | | | | | ORF1ab | synonymous_N1995N |
| 6258 | C | T | 6.32E-05 | | | | | ORF1ab | nonsynonymous_T1998I |
| 6264 | A | G | 6.32E-05 | | | | | ORF1ab | nonsynonymous_K2000R |
| 6267 | C | T | 6.32E-05 | | | | | ORF1ab | nonsynonymous_A2001V |
| 6273 | A | G | 6.32E-05 | | | | | ORF1ab | nonsynonymous_Y2003C |
| 6276 | A | G | 6.32E-05 | | | | | ORF1ab | nonsynonymous_K2004R |
| 6277 | A | G | 6.32E-05 | | | | | ORF1ab | synonymous_K2004K |

| | | | | | | | | | |
|---|---|---|---|---|---|---|---|---|---|
| 6294 | T | C | 6.32E-05 | | | | | ORF1ab | nonsynonymous_I2010T |
| 6295 | A | G | 6.32E-05 | | | | | ORF1ab | nonsynonymous_I2010M |
| 6308 | A | G | 6.32E-05 | | | | | ORF1ab | nonsynonymous_S2015G |
| 6318 | C | T | 6.32E-05 | | | | | ORF1ab | nonsynonymous_P2018L |
| 6330 | C | T | 6.32E-05 | | | | | ORF1ab | nonsynonymous_S2022L |
| 6331 | A | G | 6.32E-05 | | | | | ORF1ab | synonymous_S2022S |
| 6337 | G | A | 6.32E-05 | | | | | ORF1ab | synonymous_S2024S |
| 6345 | T | C | 6.32E-05 | | | | | ORF1ab | nonsynonymous_V2027A |
| 6349 | G | A | 6.32E-05 | | | | | ORF1ab | synonymous_L2028L |
| 6350 | A | G | 6.32E-05 | | | | | ORF1ab | nonsynonymous_K2029E |
| 6366 | A | G | 6.32E-05 | | | | | ORF1ab | nonsynonymous_Q2034R |
| 6368 | G | A | 6.32E-05 | | | | | ORF1ab | nonsynonymous_G2035R |
| 6373 | G | T | 6.32E-05 | | | | | ORF1ab | nonsynonymous_M2036I |
| 6395 | C | T | 6.32E-05 | | | | | ORF1ab | synonymous_L2044L |
| 6408 | C | T | 6.32E-05 | | | | | ORF1ab | nonsynonymous_S2048F |
| 6424 | A | C | 6.32E-05 | | | | | ORF1ab | nonsynonymous_E2053D |
| 6428 | C | T | 6.32E-05 | | | | | ORF1ab | nonsynonymous_P2055S |
| 6430 | T | C | 6.32E-05 | | | | | ORF1ab | synonymous_P2055P |
| 6441 | A | G | 6.32E-05 | | | | | ORF1ab | nonsynonymous_K2059R |
| 6442 | A | T | 6.32E-05 | T | 6.32E-05 | | | ORF1ab | nonsynonymous_K2059N; nonsynonymous_K2059N |
| 6444 | A | G | 6.32E-05 | | | | | ORF1ab | nonsynonymous_D2060G |
| 6454 | G | T | 6.32E-05 | | | | | ORF1ab | nonsynonymous_E2063D |
| 6461 | G | T | 6.32E-05 | | | | | ORF1ab | nonsynonymous_V2066L |
| 6476 | G | T | 6.32E-05 | | | | | ORF1ab | nonsynonymous_V2071F |
| 6486 | A | G | 6.32E-05 | | | | | ORF1ab | nonsynonymous_D2074G |
| 6614 | T | C | 6.32E-05 | | | | | ORF1ab | synonymous_L2117L |
| 6629 | C | T | 6.32E-05 | | | | | ORF1ab | nonsynonymous_L2122F |
| 6653 | G | A | 6.32E-05 | | | | | ORF1ab | nonsynonymous_V2130I |
| 6666 | C | T | 6.32E-05 | | | | | ORF1ab | nonsynonymous_P2134L |

| | | | | | | | | | |
|---|---|---|---|---|---|---|---|---|---|
| 6786 | C | G | 6.32E-05 | | | | | ORF1ab | nonsynonymous_T2174S |
| 6834 | T | G | 6.32E-05 | | | | | ORF1ab | nonsynonymous_I2190S |
| 6838 | A | G | 6.32E-05 | | | | | ORF1ab | synonymous_K2191K |
| 6842 | T | A | 6.32E-05 | | | | | ORF1ab | nonsynonymous_S2193T |
| 6844 | T | C | 6.32E-05 | | | | | ORF1ab | synonymous_S2193S |
| 6846 | T | C | 6.32E-05 | | | | | ORF1ab | nonsynonymous_M2194T |
| 6853 | T | C | 6.32E-05 | | | | | ORF1ab | synonymous_T2196T |
| 6870 | C | T | 6.32E-05 | | | | | ORF1ab | nonsynonymous_T2202I |
| 6871 | T | C | 6.32E-05 | | | | | ORF1ab | synonymous_T2202T |
| 6905 | T | G | 6.32E-05 | | | | | ORF1ab | nonsynonymous_S2214A |
| 6925 | A | T | 6.32E-05 | | | | | ORF1ab | synonymous_S2220S |
| 6927 | C | T | 6.32E-05 | | | | | ORF1ab | nonsynonymous_P2221L |
| 6986 | G | T | 6.32E-05 | | | | | ORF1ab | nonsynonymous_G2241C |
| 7064 | A | G | 6.32E-05 | | | | | ORF1ab | nonsynonymous_R2267G |
| 7079 | A | G | 6.32E-05 | | | | | ORF1ab | nonsynonymous_N2272D |
| 7095 | C | T | 6.32E-05 | | | | | ORF1ab | nonsynonymous_T2277I |
| 7098 | T | C | 6.32E-05 | | | | | ORF1ab | nonsynonymous_I2278T |
| 7100 | G | T | 6.32E-05 | | | | | ORF1ab | nonsynonymous_A2279S |
| 7101 | C | T | 6.32E-05 | | | | | ORF1ab | nonsynonymous_A2279V |
| 7105 | C | T | 6.32E-05 | | | | | ORF1ab | synonymous_T2280T |
| 7106 | T | C | 6.32E-05 | | | | | ORF1ab | nonsynonymous_Y2281H |
| 7113 | C | T | 6.32E-05 | | | | | ORF1ab | nonsynonymous_T2283I |
| 7119 | C | T | 6.32E-05 | | | | | ORF1ab | nonsynonymous_S2285F |
| 7128 | G | A | 6.32E-05 | | | | | ORF1ab | nonsynonymous_C2288Y |
| 7144 | T | A | 6.32E-05 | | | | | ORF1ab | nonsynonymous_S2293R |
| 7270 | T | C | 6.32E-05 | | | | | ORF1ab | synonymous_T2335T |
| 7273 | G | A | 6.32E-05 | | | | | ORF1ab | synonymous_R2336R |
| 7283 | G | A | 6.32E-05 | | | | | ORF1ab | nonsynonymous_V2340I |
| 7286 | C | A | 6.32E-05 | | | | | ORF1ab | nonsynonymous_L2341I |
| 7290 | G | A | 6.32E-05 | | | | | ORF1ab | nonsynonymous_G2342E |

| | | | | | | | | | |
|---|---|---|---|---|---|---|---|---|---|
| 7291 | A | G | 6.32E-05 | | | | | ORF1ab | synonymous_G2342G |
| 7296 | C | A | 6.32E-05 | | | | | ORF1ab | nonsynonymous_A2344D |
| 7299 | C | A | 6.32E-05 | A | 6.32E-05 | | | ORF1ab | nonsynonymous_A2345E;<br>nonsynonymous_A2345E |
| 7316 | T | G | 6.32E-05 | | | | | ORF1ab | nonsynonymous_F2351V |
| 7345 | T | A | 6.32E-05 | | | | | ORF1ab | nonsynonymous_S2360R |
| 7360 | G | A | 6.32E-05 | A | 6.32E-05 | | | ORF1ab | nonsynonymous_M2365I;<br>nonsynonymous_M2365I |
| 7705 | A | T | 6.32E-05 | | | | | ORF1ab | synonymous_P2480P |
| 7710 | A | G | 6.32E-05 | | | | | ORF1ab | nonsynonymous_N2482S |
| 7749 | C | T | 6.32E-05 | | | | | ORF1ab | nonsynonymous_T2495I |
| 7814 | T | C | 6.32E-05 | | | | | ORF1ab | nonsynonymous_S2517P |
| 7846 | G | A | 6.32E-05 | | | | | ORF1ab | synonymous_L2527L |
| 7930 | A | G | 6.32E-05 | | | | | ORF1ab | synonymous_K2555K |
| 7938 | C | G | 6.32E-05 | | | | | ORF1ab | nonsynonymous_S2558C |
| 7970 | A | G | 6.32E-05 | | | | | ORF1ab | nonsynonymous_I2569V |
| 7976 | T | C | 6.32E-05 | | | | | ORF1ab | synonymous_L2571L |
| 7998 | C | T | 6.32E-05 | | | | | ORF1ab | nonsynonymous_S2578F |
| 8175 | C | T | 6.32E-05 | | | | | ORF1ab | nonsynonymous_A2637V |
| 8637 | C | T | 6.32E-05 | | | | | ORF1ab | nonsynonymous_T2791I |
| 8645 | C | T | 6.32E-05 | | | | | ORF1ab | nonsynonymous_H2794Y |
| 8682 | T | C | 6.32E-05 | | | | | ORF1ab | nonsynonymous_I2806T |
| 8704 | T | C | 6.32E-05 | | | | | ORF1ab | synonymous_D2813D |
| 8713 | C | A | 6.32E-05 | | | | | ORF1ab | synonymous_V2816V |
| 8728 | A | G | 6.32E-05 | | | | | ORF1ab | synonymous_A2821A |
| 8729 | T | C | 6.32E-05 | | | | | ORF1ab | nonsynonymous_S2822P |
| 8730 | C | T | 6.32E-05 | | | | | ORF1ab | nonsynonymous_S2822F |
| 8739 | C | T | 6.32E-05 | | | | | ORF1ab | nonsynonymous_T2825I |
| 8752 | C | T | 6.32E-05 | | | | | ORF1ab | synonymous_N2829N |
| 8800 | T | C | 6.32E-05 | | | | | ORF1ab | synonymous_Y2845Y |

| | | | | | | | | | |
|---|---|---|---|---|---|---|---|---|---|
| 9248 | G | A | 6.32E-05 | | | | | ORF1ab | nonsynonymous_G2995S |
| 9250 | T | C | 6.32E-05 | | | | | ORF1ab | synonymous_G2995G |
| 9259 | A | G | 6.32E-05 | | | | | ORF1ab | synonymous_V2998V |
| 9264 | C | T | 6.32E-05 | | | | | ORF1ab | nonsynonymous_T3000I |
| 9267 | G | A | 6.32E-05 | A | 6.32E-05 | | | ORF1ab | nonsynonymous_S3001N; nonsynonymous_S3001N |
| 9269 | G | A | 6.32E-05 | | | | | ORF1ab | nonsynonymous_G3002S |
| 9304 | T | C | 6.32E-05 | | | | | ORF1ab | synonymous_S3013S |
| 9436 | A | T | 6.32E-05 | | | | | ORF1ab | synonymous_V3057V |
| 9437 | A | T | 6.32E-05 | | | | | ORF1ab | nonsynonymous_T3058S |
| 9454 | T | C | 6.32E-05 | | | | | ORF1ab | synonymous_Y3063Y |
| 9473 | G | A | 6.32E-05 | | | | | ORF1ab | nonsynonymous_A3070T |
| 9485 | T | C | 6.32E-05 | | | | | ORF1ab | nonsynonymous_Y3074H |
| 9496 | A | G | 6.32E-05 | | | | | ORF1ab | synonymous_V3077V |
| 9640 | T | C | 6.32E-05 | | | | | ORF1ab | synonymous_H3125H |
| 9665 | C | T | 6.32E-05 | | | | | ORF1ab | nonsynonymous_P3134S |
| 9683 | A | G | 6.32E-05 | | | | | ORF1ab | nonsynonymous_I3140V |
| 9863 | C | T | 6.32E-05 | | | | | ORF1ab | nonsynonymous_P3200S |
| 2153 | T | C | 6.32E-05 | A | 6.32E-05 | | | ORF1ab | nonsynonymous_W630R; nonsynonymous_W630R |
| 675 | G | C | 6.32E-05 | A | 6.32E-05 | A | 6.32E-05 | ORF1ab | nonsynonymous_G137A; nonsynonymous_G137D; nonsynonymous_G137D |
| 10054 | G | A | 6.32E-05 | | | | | ORF1ab | synonymous_Q3263Q |
| 10074 | C | T | 6.32E-05 | | | | | ORF1ab | nonsynonymous_A3270V |
| 10076 | T | C | 6.32E-05 | | | | | ORF1ab | nonsynonymous_F3271L |
| 10081 | A | T | 6.32E-05 | | | | | ORF1ab | synonymous_P3272P |
| 10135 | T | C | 6.32E-05 | | | | | ORF1ab | synonymous_L3290L |
| 10177 | T | C | 6.32E-05 | | | | | ORF1ab | synonymous_H3304H |
| 1018 | G | A | 6.32E-05 | | | | | ORF1ab | synonymous_L251L |

| | | | | | | | | | |
|---|---|---|---|---|---|---|---|---|---|
| 10203 | T | A | 6.32E-05 | | | | | ORF1ab | nonsynonymous_L3313H |
| 10332 | C | T | 6.32E-05 | | | | | ORF1ab | nonsynonymous_T3356I |
| 11042 | G | T | 6.32E-05 | | | | | ORF1ab | nonsynonymous_V3593F |
| 11093 | G | A | 6.32E-05 | | | | | ORF1ab | nonsynonymous_A3610T |
| 11095 | C | T | 6.32E-05 | | | | | ORF1ab | synonymous_A3610A |
| 1112 | A | G | 6.32E-05 | | | | | ORF1ab | nonsynonymous_T283A |
| 1126 | G | T | 6.32E-05 | | | | | ORF1ab | nonsynonymous_R287S |
| 11290 | T | C | 6.32E-05 | | | | | ORF1ab | synonymous_S3675S |
| 11291 | G | A | 6.32E-05 | | | | | ORF1ab | nonsynonymous_G3676S |
| 11339 | C | T | 6.32E-05 | | | | | ORF1ab | synonymous_L3692L |
| 11396 | C | T | 6.32E-05 | T | 6.32E-05 | | | ORF1ab | nonsynonymous_L3711F; nonsynonymous_L3711F |
| 11416 | C | A | 6.32E-05 | A | 6.32E-05 | | | ORF1ab | synonymous_L3717L; synonymous_L3717L |
| 11515 | T | A | 6.32E-05 | | | | | ORF1ab | synonymous_T3750T |
| 11730 | A | G | 6.32E-05 | | | | | ORF1ab | nonsynonymous_Y3822C |
| 11755 | A | G | 6.32E-05 | | | | | ORF1ab | synonymous_P3830P |
| 11807 | G | A | 6.32E-05 | | | | | ORF1ab | nonsynonymous_G3848S |
| 11996 | A | G | 6.32E-05 | | | | | ORF1ab | nonsynonymous_M3911V |
| 12038 | G | A | 6.32E-05 | | | | | ORF1ab | nonsynonymous_V3925I |
| 12041 | G | C | 6.32E-05 | | | | | ORF1ab | nonsynonymous_D3926H |
| 12043 | C | T | 6.32E-05 | | | | | ORF1ab | synonymous_D3926D |
| 12130 | A | G | 6.32E-05 | | | | | ORF1ab | synonymous_A3955A |
| 12142 | T | C | 6.32E-05 | | | | | ORF1ab | synonymous_T3959T |
| 12143 | G | T | 6.32E-05 | | | | | ORF1ab | nonsynonymous_A3960S |
| 12151 | A | G | 6.32E-05 | | | | | ORF1ab | synonymous_E3962E |
| 12153 | C | T | 6.32E-05 | | | | | ORF1ab | nonsynonymous_A3963V |
| 12160 | G | C | 6.32E-05 | | | | | ORF1ab | nonsynonymous_E3965D |
| 12171 | C | T | 6.32E-05 | | | | | ORF1ab | nonsynonymous_A3969V |
| 12180 | A | G | 6.32E-05 | | | | | ORF1ab | nonsynonymous_D3972G |

| | | | | | | | | | |
|---|---|---|---|---|---|---|---|---|---|
| 12184 | T | C | 6.32E-05 | | | | | ORF1ab | synonymous_S3973S |
| 12202 | G | C | 6.32E-05 | | | | | ORF1ab | nonsynonymous_K3979N |
| 12331 | G | C | 6.32E-05 | | | | | ORF1ab | nonsynonymous_R4022S |
| 1251 | C | T | 6.32E-05 | | | | | ORF1ab | nonsynonymous_T329I |
| 12520 | T | C | 6.32E-05 | | | | | ORF1ab | synonymous_D4085D |
| 1252 | T | G | 6.32E-05 | | | | | ORF1ab | synonymous_T329T |
| 1258 | G | T | 6.32E-05 | | | | | ORF1ab | nonsynonymous_W331C |
| 12613 | G | A | 6.32E-05 | | | | | ORF1ab | nonsynonymous_M4116I |
| 12638 | C | A | 6.32E-05 | | | | | ORF1ab | nonsynonymous_P4125T |
| 12669 | C | T | 6.32E-05 | | | | | ORF1ab | nonsynonymous_S4135F |
| 12815 | C | T | 6.32E-05 | | | | | ORF1ab | synonymous_L4184L |
| 12827 | T | G | 6.32E-05 | | | | | ORF1ab | nonsynonymous_L4188V |
| 12974 | C | T | 6.32E-05 | | | | | ORF1ab | synonymous_L4237L |
| 13011 | C | T | 6.32E-05 | | | | | ORF1ab | nonsynonymous_T4249I |
| 13095 | C | T | 6.32E-05 | | | | | ORF1ab | nonsynonymous_A4277V |
| 13124 | G | A | 6.32E-05 | | | | | ORF1ab | nonsynonymous_G4287R |
| 13126 | G | A | 6.32E-05 | | | | | ORF1ab | synonymous_G4287G |
| 13133 | C | T | 6.32E-05 | | | | | ORF1ab | nonsynonymous_P4290S |
| 1362 | T | G | 6.32E-05 | | | | | ORF1ab | nonsynonymous_V366G |
| 1470 | G | A | 6.32E-05 | A | 6.32E-05 | | | ORF1ab | nonsynonymous_R402H; nonsynonymous_R402H |
| 1475 | A | C | 6.32E-05 | | | | | ORF1ab | nonsynonymous_I404L |
| 1483 | T | A | 6.32E-05 | | | | | ORF1ab | nonsynonymous_F406L |
| 1494 | T | C | 6.32E-05 | | | | | ORF1ab | nonsynonymous_V410A |
| 1495 | G | T | 6.32E-05 | | | | | ORF1ab | synonymous_V410V |
| 1508 | G | A | 6.32E-05 | | | | | ORF1ab | nonsynonymous_G415S |
| 1516 | T | C | 6.32E-05 | | | | | ORF1ab | synonymous_H417H |
| 1537 | T | C | 6.32E-05 | | | | | ORF1ab | synonymous_V424V |
| 1548 | G | A | 6.32E-05 | | | | | ORF1ab | nonsynonymous_S428N |
| 1551 | C | T | 6.32E-05 | | | | | ORF1ab | nonsynonymous_A429V |

| | | | | | | | | | |
|---|---|---|---|---|---|---|---|---|---|
| 1555 | C | A | 6.32E-05 | | | | | ORF1ab | nonsynonymous_N430K |
| 1573 | A | G | 6.32E-05 | | | | | ORF1ab | synonymous_T436T |
| 1576 | T | C | 6.32E-05 | | | | | ORF1ab | synonymous_G437G |
| 1637 | A | T | 6.32E-05 | | | | | ORF1ab | stopgain_K458X |
| 1642 | C | T | 6.32E-05 | | | | | ORF1ab | synonymous_V459V |
| 1643 | A | G | 6.32E-05 | | | | | ORF1ab | nonsynonymous_N460D |
| 1671 | T | C | 6.32E-05 | | | | | ORF1ab | nonsynonymous_L469P |
| 1679 | G | A | 6.32E-05 | | | | | ORF1ab | nonsynonymous_E472K |
| 1711 | T | A | 6.32E-05 | | | | | ORF1ab | synonymous_A482A |
| 1713 | C | A | 6.32E-05 | | | | | ORF1ab | nonsynonymous_S483Y |
| 1717 | A | G | 6.32E-05 | | | | | ORF1ab | synonymous_T484T |
| 1721 | G | T | 6.32E-05 | | | | | ORF1ab | nonsynonymous_A486S |
| 1738 | G | T | 6.32E-05 | | | | | ORF1ab | synonymous_V491V |
| 1740 | A | C | 6.32E-05 | | | | | ORF1ab | nonsynonymous_K492T |
| 1750 | T | C | 6.32E-05 | | | | | ORF1ab | synonymous_D495D |
| 1762 | C | T | 6.32E-05 | | | | | ORF1ab | synonymous_F499F |
| 1780 | C | T | 6.32E-05 | | | | | ORF1ab | synonymous_S505S |
| 1782 | G | A | 6.32E-05 | | | | | ORF1ab | nonsynonymous_C506Y |
| 1824 | C | T | 6.32E-05 | | | | | ORF1ab | nonsynonymous_A520V |
| 1954 | G | T | 6.32E-05 | | | | | ORF1ab | nonsynonymous_Q563H |
| 1957 | G | T | 6.32E-05 | | | | | ORF1ab | nonsynonymous_K564N |
| 1959 | C | T | 6.32E-05 | | | | | ORF1ab | nonsynonymous_A565V |
| 1960 | C | T | 6.32E-05 | | | | | ORF1ab | synonymous_A565A |
| 1964 | A | C | 6.32E-05 | | | | | ORF1ab | nonsynonymous_I567L |
| 1967 | A | G | 6.32E-05 | | | | | ORF1ab | nonsynonymous_T568A |
| 1972 | A | G | 6.32E-05 | | | | | ORF1ab | nonsynonymous_I569M |
| 1983 | T | C | 6.32E-05 | | | | | ORF1ab | nonsynonymous_I573T |
| 1984 | T | C | 6.32E-05 | | | | | ORF1ab | synonymous_I573I |
| 2004 | T | G | 6.32E-05 | | | | | ORF1ab | nonsynonymous_L580R |
| 2022 | T | G | 6.32E-05 | | | | | ORF1ab | nonsynonymous_F586C |

| | | | | | | | | | |
|---|---|---|---|---|---|---|---|---|---|
| 2040 | C | T | 6.32E-05 | | | | | ORF1ab | nonsynonymous_T592I |
| 2044 | C | T | 6.32E-05 | | | | | ORF1ab | synonymous_N593N |
| 2048 | C | T | 6.32E-05 | | | | | ORF1ab | synonymous_L595L |
| 2051 | G | A | 6.32E-05 | | | | | ORF1ab | nonsynonymous_V596I |
| 2058 | T | C | 6.32E-05 | | | | | ORF1ab | nonsynonymous_M598T |
| 2075 | G | A | 6.32E-05 | | | | | ORF1ab | nonsynonymous_G604S |
| 2102 | C | T | 6.32E-05 | | | | | ORF1ab | synonymous_L613L |
| 2106 | C | T | 6.32E-05 | | | | | ORF1ab | nonsynonymous_T614I |
| 2111 | A | T | 6.32E-05 | T | 6.32E-05 | | | ORF1ab | nonsynonymous_I616F; nonsynonymous_I616F |
| 2117 | G | T | 6.32E-05 | | | | | ORF1ab | nonsynonymous_G618C |
| 2123 | G | A | 6.32E-05 | | | | | ORF1ab | nonsynonymous_V620I |
| 2138 | A | G | 6.32E-05 | | | | | ORF1ab | nonsynonymous_K625E |
| 2141 | C | A | 6.32E-05 | | | | | ORF1ab | nonsynonymous_P626T |
| 2144 | G | T | 6.32E-05 | | | | | ORF1ab | nonsynonymous_V627F |
| 2147 | C | T | 6.32E-05 | | | | | ORF1ab | nonsynonymous_L628F |
| 2158 | T | C | 6.32E-05 | | | | | ORF1ab | synonymous_L631L |
| 2165 | A | G | 6.32E-05 | | | | | ORF1ab | nonsynonymous_K634E |
| 2169 | T | C | 6.32E-05 | | | | | ORF1ab | nonsynonymous_F635S |
| 2173 | G | A | 6.32E-05 | | | | | ORF1ab | synonymous_K636K |
| 2190 | T | A | 6.32E-05 | | | | | ORF1ab | nonsynonymous_L642H |
| 2197 | C | T | 6.32E-05 | | | | | ORF1ab | synonymous_D644D |
| 2245 | C | T | 6.32E-05 | | | | | ORF1ab | synonymous_V660V |
| 2345 | A | T | 6.32E-05 | | | | | ORF1ab | nonsynonymous_I694F |
| 2357 | G | A | 6.32E-05 | | | | | ORF1ab | nonsynonymous_A698T |
| 2358 | C | T | 6.32E-05 | | | | | ORF1ab | nonsynonymous_A698V |
| 2361 | A | G | 6.32E-05 | | | | | ORF1ab | nonsynonymous_K699R |
| 2369 | G | A | 6.32E-05 | | | | | ORF1ab | nonsynonymous_A702T |
| 2371 | C | T | 6.32E-05 | | | | | ORF1ab | synonymous_A702A |
| 2383 | T | G | 6.32E-05 | | | | | ORF1ab | synonymous_G706G |

| | | | | | | | | | |
|---|---|---|---|---|---|---|---|---|---|
| 2397 | C | T | 6.32E-05 | | | | | ORF1ab | nonsynonymous_T711M |
| 2401 | C | A | 6.32E-05 | | | | | ORF1ab | nonsynonymous_H712Q |
| 2437 | A | C | 6.32E-05 | | | | | ORF1ab | nonsynonymous_R724S |
| 2514 | A | G | 6.32E-05 | | | | | ORF1ab | nonsynonymous_E750G |
| 2516 | G | A | 6.32E-05 | | | | | ORF1ab | nonsynonymous_V751M |
| 2517 | T | A | 6.32E-05 | | | | | ORF1ab | nonsynonymous_V751E |
| 2518 | G | A | 6.32E-05 | A | 6.32E-05 | | | ORF1ab | synonymous_V751V; synonymous_V751V |
| 2885 | G | T | 6.32E-05 | | | | | ORF1ab | nonsynonymous_V874F |
| 2893 | A | G | 6.32E-05 | | | | | ORF1ab | synonymous_A876A |
| 2900 | G | A | 6.32E-05 | | | | | ORF1ab | nonsynonymous_V879I |
| 2922 | T | C | 6.32E-05 | | | | | ORF1ab | nonsynonymous_V886A |
| 2931 | T | C | 6.32E-05 | | | | | ORF1ab | nonsynonymous_L889S |
| 2932 | A | G | 6.32E-05 | | | | | ORF1ab | synonymous_L889L |
| 2944 | G | A | 6.32E-05 | | | | | ORF1ab | synonymous_L893L |
| 2952 | A | G | 6.32E-05 | | | | | ORF1ab | nonsynonymous_D896G |
| 2956 | A | T | 6.32E-05 | | | | | ORF1ab | nonsynonymous_L897F |
| 2957 | G | T | 6.32E-05 | | | | | ORF1ab | nonsynonymous_D898Y |
| 2971 | G | T | 6.32E-05 | | | | | ORF1ab | nonsynonymous_M902I |
| 2981 | T | C | 6.32E-05 | | | | | ORF1ab | nonsynonymous_Y906H |
| 2982 | A | G | 6.32E-05 | | | | | ORF1ab | nonsynonymous_Y906C |
| 2986 | A | G | 6.32E-05 | | | | | ORF1ab | synonymous_L907L |
| 2991 | A | G | 6.32E-05 | | | | | ORF1ab | nonsynonymous_D909G |
| 2992 | T | C | 6.32E-05 | | | | | ORF1ab | synonymous_D909D |
| 3013 | G | A | 6.32E-05 | A | 6.32E-05 | | | ORF1ab | synonymous_L916L; synonymous_L916L |
| 3022 | T | A | 6.32E-05 | | | | | ORF1ab | nonsynonymous_H919Q |
| 3025 | G | T | 6.32E-05 | | | | | ORF1ab | nonsynonymous_M920I |
| 3027 | A | G | 6.32E-05 | | | | | ORF1ab | nonsynonymous_Y921C |
| 3047 | G | T | 6.32E-05 | | | | | ORF1ab | nonsynonymous_D928Y |

| | | | | | | | | | |
|---|---|---|---|---|---|---|---|---|---|
| 3058 | A | G | 6.32E-05 | | | | | ORF1ab | synonymous_E931E |
| 3065 | G | T | 6.32E-05 | | | | | ORF1ab | nonsynonymous_G934C |
| 3068 | G | A | 6.32E-05 | | | | | ORF1ab | nonsynonymous_D935N |
| 3069 | A | G | 6.32E-05 | | | | | ORF1ab | nonsynonymous_D935G |
| 3081 | A | C | 6.32E-05 | | | | | ORF1ab | nonsynonymous_E939A |
| 3093 | C | T | 6.32E-05 | | | | | ORF1ab | nonsynonymous_P943L |
| 3109 | G | T | 6.32E-05 | | | | | ORF1ab | nonsynonymous_E948D |
| 3113 | G | C | 6.32E-05 | | | | | ORF1ab | nonsynonymous_G950R |
| 3117 | C | T | 6.32E-05 | | | | | ORF1ab | nonsynonymous_T951I |
| 3118 | T | C | 6.32E-05 | | | | | ORF1ab | synonymous_T951T |
| 3132 | A | G | 6.32E-05 | | | | | ORF1ab | nonsynonymous_Q956R |
| 3146 | G | C | 6.32E-05 | | | | | ORF1ab | nonsynonymous_E961Q |
| 3156 | C | T | 6.32E-05 | | | | | ORF1ab | nonsynonymous_A964V |
| 3160 | T | C | 6.32E-05 | | | | | ORF1ab | synonymous_T965T |
| 3163 | T | A | 6.32E-05 | | | | | ORF1ab | synonymous_S966S |
| 3180 | A | T | 6.32E-05 | | | | | ORF1ab | nonsynonymous_E972V |
| 3181 | A | T | 6.32E-05 | | | | | ORF1ab | nonsynonymous_E972D |
| 3246 | G | T | 6.32E-05 | | | | | ORF1ab | nonsynonymous_S994I |
| 3250 | G | T | 6.32E-05 | | | | | ORF1ab | nonsynonymous_E995D |
| 3317 | C | T | 6.32E-05 | | | | | ORF1ab | nonsynonymous_P1018S |
| 3320 | G | T | 6.32E-05 | | | | | ORF1ab | nonsynonymous_V1019F |
| 3323 | G | T | 6.32E-05 | | | | | ORF1ab | nonsynonymous_V1020F |
| 3330 | C | T | 6.32E-05 | | | | | ORF1ab | nonsynonymous_T1022I |
| 3350 | A | G | 6.32E-05 | | | | | ORF1ab | nonsynonymous_S1029G |
| 3369 | C | T | 6.32E-05 | | | | | ORF1ab | nonsynonymous_T1035I |
| 3371 | G | A | 6.32E-05 | | | | | ORF1ab | nonsynonymous_D1036N |
| 3375 | A | C | 6.32E-05 | | | | | ORF1ab | nonsynonymous_N1037T |
| 3376 | T | C | 6.32E-05 | | | | | ORF1ab | synonymous_N1037N |
| 3381 | A | G | 6.32E-05 | | | | | ORF1ab | nonsynonymous_Y1039C |
| 3388 | A | G | 6.32E-05 | | | | | ORF1ab | synonymous_K1041K |

| | | | | | | | | | |
|---|---|---|---|---|---|---|---|---|---|
| 3398 | A | G | 6.32E-05 | | | | | ORF1ab | nonsynonymous_I1045V |
| 3406 | A | T | 6.32E-05 | T | 6.32E-05 | | | ORF1ab | nonsynonymous_E1047D; nonsynonymous_E1047D |
| 3410 | G | A | 6.32E-05 | | | | | ORF1ab | nonsynonymous_A1049T |
| 3416 | A | C | 6.32E-05 | | | | | ORF1ab | nonsynonymous_K1051Q |
| 3428 | A | C | 6.32E-05 | | | | | ORF1ab | nonsynonymous_T1055P |
| 3445 | A | G | 6.32E-05 | | | | | ORF1ab | synonymous_A1060A |
| 3450 | A | G | 6.32E-05 | | | | | ORF1ab | nonsynonymous_N1062S |
| 3455 | T | C | 6.32E-05 | | | | | ORF1ab | nonsynonymous_Y1064H |
| 3463 | A | G | 6.32E-05 | | | | | ORF1ab | synonymous_K1066K |
| 3467 | G | T | 6.32E-05 | | | | | ORF1ab | stopgain_G1068X |
| 3478 | T | C | 6.32E-05 | | | | | ORF1ab | synonymous_V1071V |
| 3502 | T | A | 6.32E-05 | | | | | ORF1ab | synonymous_T1079T |
| 3514 | G | T | 6.32E-05 | | | | | ORF1ab | nonsynonymous_M1083I |
| 3522 | A | G | 6.32E-05 | | | | | ORF1ab | nonsynonymous_E1086G |
| 3578 | T | C | 6.32E-05 | | | | | ORF1ab | synonymous_L1105L |
| 3695 | C | T | 6.32E-05 | | | | | ORF1ab | synonymous_L1144L |
| 3698 | C | A | 6.32E-05 | A | 6.32E-05 | | | ORF1ab | nonsynonymous_L1145I; nonsynonymous_L1145I |
| 3725 | T | G | 6.32E-05 | | | | | ORF1ab | nonsynonymous_F1154V |
| 3727 | T | C | 6.32E-05 | | | | | ORF1ab | synonymous_F1154F |
| 3759 | G | T | 6.32E-05 | | | | | ORF1ab | nonsynonymous_C1165F |
| 3821 | C | A | 6.32E-05 | | | | | ORF1ab | nonsynonymous_L1186I |
| 3920 | A | T | 6.32E-05 | | | | | ORF1ab | stopgain_K1219X |
| 3921 | A | T | 6.32E-05 | T | 6.32E-05 | | | ORF1ab | nonsynonymous_K1219I; nonsynonymous_K1219I |
| 3947 | G | T | 6.32E-05 | | | | | ORF1ab | nonsynonymous_D1228Y |
| 3961 | C | T | 6.32E-05 | | | | | ORF1ab | synonymous_I1232I |
| 3971 | G | T | 6.32E-05 | | | | | ORF1ab | nonsynonymous_V1236F |
| 3977 | G | A | 6.32E-05 | | | | | ORF1ab | nonsynonymous_E1238K |

| | | | | | | | | | |
|---|---|---|---|---|---|---|---|---|---|
| 3990 | C | T | 6.32E-05 | | | | | ORF1ab | nonsynonymous_T1242I |
| 3995 | G | A | 6.32E-05 | | | | | ORF1ab | nonsynonymous_E1244K |
| 400 | T | C | 6.32E-05 | | | | | ORF1ab | synonymous_H45H |
| 4014 | C | T | 6.32E-05 | | | | | ORF1ab | nonsynonymous_T1250I |
| 4016 | G | C | 6.32E-05 | | | | | ORF1ab | nonsynonymous_E1251Q |
| 401 | C | T | 6.32E-05 | | | | | ORF1ab | nonsynonymous_L46F |
| 4034 | A | G | 6.32E-05 | | | | | ORF1ab | nonsynonymous_I1257V |
| 4047 | G | A | 6.32E-05 | | | | | ORF1ab | nonsynonymous_G1261D |
| 4048 | C | T | 6.32E-05 | | | | | ORF1ab | synonymous_G1261G |
| 4068 | C | T | 6.32E-05 | | | | | ORF1ab | nonsynonymous_A1268V |
| 4080 | G | A | 6.32E-05 | | | | | ORF1ab | nonsynonymous_S1272N |
| 4165 | T | A | 6.32E-05 | | | | | ORF1ab | synonymous_V1300V |
| 4170 | C | G | 6.32E-05 | | | | | ORF1ab | nonsynonymous_P1302R |
| 4171 | T | A | 6.32E-05 | | | | | ORF1ab | synonymous_P1302P |
| 4173 | C | T | 6.32E-05 | | | | | ORF1ab | nonsynonymous_T1303I |
| 4181 | G | A | 6.32E-05 | | | | | ORF1ab | nonsynonymous_A1306T |
| 4184 | G | A | 6.32E-05 | | | | | ORF1ab | nonsynonymous_G1307S |
| 4252 | C | T | 6.32E-05 | | | | | ORF1ab | synonymous_Y1329Y |
| 4258 | T | C | 6.32E-05 | | | | | ORF1ab | synonymous_G1331G |
| 4280 | G | A | 6.32E-05 | | | | | ORF1ab | nonsynonymous_V1339I |
| 4288 | G | T | 6.32E-05 | | | | | ORF1ab | nonsynonymous_E1341D |
| 4507 | T | C | 6.32E-05 | | | | | ORF1ab | synonymous_G1414G |
| 4517 | T | A | 6.32E-05 | | | | | ORF1ab | nonsynonymous_Y1418N |
| 4672 | C | T | 6.32E-05 | | | | | ORF1ab | synonymous_L1469L |
| 4678 | G | A | 6.32E-05 | | | | | ORF1ab | synonymous_V1471V |
| 4684 | T | C | 6.32E-05 | | | | | ORF1ab | synonymous_A1473A |
| 4687 | A | T | 6.32E-05 | | | | | ORF1ab | synonymous_T1474T |
| 4704 | C | T | 6.32E-05 | | | | | ORF1ab | nonsynonymous_P1480L |
| 4706 | G | C | 6.32E-05 | | | | | ORF1ab | nonsynonymous_D1481H |
| 4729 | T | A | 6.32E-05 | | | | | ORF1ab | synonymous_G1488G |

| | | | | | | | | | |
|---|---|---|---|---|---|---|---|---|---|
| 4731 | A | G | 6.32E-05 | | | | | ORF1ab | nonsynonymous_Y1489C |
| 4739 | T | C | 6.32E-05 | | | | | ORF1ab | nonsynonymous_S1492P |
| 4752 | C | T | 6.32E-05 | | | | | ORF1ab | nonsynonymous_T1496I |
| 4754 | C | T | 6.32E-05 | | | | | ORF1ab | nonsynonymous_P1497S |
| 4815 | C | T | 6.32E-05 | | | | | ORF1ab | nonsynonymous_S1517F |
| 486 | C | T | 6.32E-05 | | | | | ORF1ab | nonsynonymous_S74L |
| 4891 | C | T | 6.32E-05 | | | | | ORF1ab | synonymous_T1542T |
| 489 | A | G | 6.32E-05 | | | | | ORF1ab | nonsynonymous_D75G |
| 4931 | C | A | 6.32E-05 | | | | | ORF1ab | nonsynonymous_L1556I |
| 5593 | T | C | 6.32E-05 | | | | | ORF1ab | synonymous_Y1776Y |
| 5609 | G | T | 6.32E-05 | | | | | ORF1ab | nonsynonymous_G1782C |
| 560 | C | T | 6.32E-05 | | | | | ORF1ab | nonsynonymous_R99C |
| 5628 | C | T | 6.32E-05 | | | | | ORF1ab | nonsynonymous_T1788M |
| 563 | A | T | 6.32E-05 | | | | | ORF1ab | nonsynonymous_S100C |
| 5657 | G | A | 6.32E-05 | A | 6.32E-05 | | | ORF1ab | nonsynonymous_V1798I; nonsynonymous_V1798I |
| 565 | T | C | 6.32E-05 | | | | | ORF1ab | synonymous_S100S |
| 577 | T | A | 6.32E-05 | | | | | ORF1ab | synonymous_L104L |
| 5829 | A | G | 6.32E-05 | | | | | ORF1ab | nonsynonymous_K1855R |
| 5846 | G | A | 6.32E-05 | | | | | ORF1ab | nonsynonymous_G1861S |
| 5880 | G | A | 6.32E-05 | | | | | ORF1ab | nonsynonymous_S1872N |
| 589 | C | A | 6.32E-05 | | | | | ORF1ab | synonymous_V108V |
| 5941 | T | G | 6.32E-05 | G | 6.32E-05 | | | ORF1ab | nonsynonymous_I1892M; nonsynonymous_I1892M |
| 5944 | C | T | 6.32E-05 | | | | | ORF1ab | synonymous_D1893D |
| 595 | T | A | 6.32E-05 | | | | | ORF1ab | nonsynonymous_H110Q |
| 5996 | C | T | 6.32E-05 | | | | | ORF1ab | nonsynonymous_P1911S |
| 6016 | C | A | 6.32E-05 | A | 6.32E-05 | | | ORF1ab | nonsynonymous_N1917K; nonsynonymous_N1917K |
| 6032 | G | A | 6.32E-05 | | | | | ORF1ab | nonsynonymous_A1923T |

| | | | | | | | | | |
|---|---|---|---|---|---|---|---|---|---|
| 6033 | C | T | 6.32E-05 | | | | | ORF1ab | nonsynonymous_A1923V |
| 6052 | G | A | 6.32E-05 | | | | | ORF1ab | synonymous_K1929K |
| 608 | C | T | 6.32E-05 | | | | | ORF1ab | nonsynonymous_P115S |
| 609 | C | T | 6.32E-05 | | | | | ORF1ab | nonsynonymous_P115L |
| 6190 | C | T | 6.32E-05 | | | | | ORF1ab | synonymous_Y1975Y |
| 6196 | C | T | 6.32E-05 | | | | | ORF1ab | synonymous_P1977P |
| 6202 | T | A | 6.32E-05 | | | | | ORF1ab | nonsynonymous_F1979L |
| 622 | C | T | 6.32E-05 | | | | | ORF1ab | synonymous_R119R |
| 627 | T | A | 6.32E-05 | | | | | ORF1ab | nonsynonymous_V121D |
| 6513 | G | T | 6.32E-05 | | | | | ORF1ab | nonsynonymous_S2083I |
| 6530 | G | A | 6.32E-05 | | | | | ORF1ab | nonsynonymous_E2089K |
| 6532 | G | T | 6.32E-05 | | | | | ORF1ab | nonsynonymous_E2089D |
| 6536 | G | T | 6.32E-05 | | | | | ORF1ab | nonsynonymous_G2091C |
| 6542 | A | G | 6.32E-05 | | | | | ORF1ab | nonsynonymous_T2093A |
| 6544 | A | G | 6.32E-05 | | | | | ORF1ab | synonymous_T2093T |
| 6555 | C | T | 6.32E-05 | | | | | ORF1ab | nonsynonymous_A2097V |
| 6568 | C | T | 6.32E-05 | | | | | ORF1ab | synonymous_D2101D |
| 6569 | A | G | 6.32E-05 | | | | | ORF1ab | nonsynonymous_N2102D |
| 6584 | A | G | 6.32E-05 | | | | | ORF1ab | nonsynonymous_I2107V |
| 6589 | G | T | 6.32E-05 | | | | | ORF1ab | nonsynonymous_K2108N |
| 6593 | C | T | 6.32E-05 | | | | | ORF1ab | nonsynonymous_P2110S |
| 6618 | G | A | 6.32E-05 | | | | | ORF1ab | nonsynonymous_G2118D |
| 6630 | T | C | 6.32E-05 | | | | | ORF1ab | nonsynonymous_L2122P |
| 6639 | A | G | 6.32E-05 | | | | | ORF1ab | nonsynonymous_H2125R |
| 6648 | C | T | 6.32E-05 | | | | | ORF1ab | nonsynonymous_A2128V |
| 6662 | G | A | 6.32E-05 | | | | | ORF1ab | nonsynonymous_V2133I |
| 6673 | T | C | 6.32E-05 | | | | | ORF1ab | synonymous_D2136D |
| 6689 | G | A | 6.32E-05 | | | | | ORF1ab | nonsynonymous_A2142T |
| 6690 | C | T | 6.32E-05 | | | | | ORF1ab | nonsynonymous_A2142V |
| 6706 | C | T | 6.32E-05 | | | | | ORF1ab | synonymous_N2147N |

| | | | | | | | | | |
|---|---|---|---|---|---|---|---|---|---|
| 6707 | A | G | 6.32E-05 | | | | | ORF1ab | nonsynonymous_K2148E |
| 6708 | A | T | 6.32E-05 | | | | | ORF1ab | nonsynonymous_K2148I |
| 6710 | G | T | 6.32E-05 | | | | | ORF1ab | nonsynonymous_V2149F |
| 6715 | T | C | 6.32E-05 | | | | | ORF1ab | synonymous_V2150V |
| 6719 | A | G | 6.32E-05 | | | | | ORF1ab | nonsynonymous_T2152A |
| 6729 | A | G | 6.32E-05 | G | 6.32E-05 | | | ORF1ab | nonsynonymous_N2155S; nonsynonymous_N2155S |
| 6731 | A | T | 6.32E-05 | | | | | ORF1ab | nonsynonymous_I2156L |
| 6748 | A | G | 6.32E-05 | | | | | ORF1ab | synonymous_L2161L |
| 6752 | C | T | 6.32E-05 | | | | | ORF1ab | nonsynonymous_R2163C |
| 6760 | T | C | 6.32E-05 | | | | | ORF1ab | synonymous_C2165C |
| 6779 | T | A | 6.32E-05 | | | | | ORF1ab | nonsynonymous_F2172I |
| 6782 | T | C | 6.32E-05 | | | | | ORF1ab | nonsynonymous_F2173L |
| 6821 | A | G | 6.32E-05 | | | | | ORF1ab | nonsynonymous_T2186A |
| 6852 | C | T | 6.32E-05 | | | | | ORF1ab | nonsynonymous_T2196I |
| 7203 | C | T | 6.32E-05 | | | | | ORF1ab | nonsynonymous_S2313F |
| 7249 | T | G | 6.32E-05 | | | | | ORF1ab | nonsynonymous_F2328L |
| 7332 | T | C | 6.32E-05 | | | | | ORF1ab | nonsynonymous_V2356A |
| 7337 | T | G | 6.32E-05 | | | | | ORF1ab | nonsynonymous_F2358V |
| 7338 | T | A | 6.32E-05 | | | | | ORF1ab | nonsynonymous_F2358Y |
| 7342 | T | A | 6.32E-05 | | | | | ORF1ab | synonymous_I2359I |
| 7348 | T | A | 6.32E-05 | | | | | ORF1ab | nonsynonymous_N2361K |
| 7351 | T | G | 6.32E-05 | | | | | ORF1ab | synonymous_S2362S |
| 7355 | C | A | 6.32E-05 | | | | | ORF1ab | nonsynonymous_L2364I |
| 7357 | T | A | 6.32E-05 | | | | | ORF1ab | synonymous_L2364L |
| 7431 | C | T | 6.32E-05 | | | | | ORF1ab | nonsynonymous_S2389L |
| 7444 | A | G | 6.32E-05 | | | | | ORF1ab | synonymous_V2393V |
| 7573 | C | T | 6.32E-05 | | | | | ORF1ab | synonymous_V2436V |
| 7704 | C | T | 6.32E-05 | | | | | ORF1ab | nonsynonymous_P2480L |
| 7716 | C | T | 6.32E-05 | | | | | ORF1ab | nonsynonymous_T2484I |

| | | | | | | | | | |
|---|---|---|---|---|---|---|---|---|---|
| 7732 | C | T | 6.32E-05 | | | | | ORF1ab | synonymous_Y2489Y |
| 7734 | T | G | 6.32E-05 | | | | | ORF1ab | nonsynonymous_I2490S |
| 7743 | G | T | 6.32E-05 | | | | | ORF1ab | nonsynonymous_S2493I |
| 7747 | T | C | 6.32E-05 | | | | | ORF1ab | synonymous_V2494V |
| 7755 | A | G | 6.32E-05 | | | | | ORF1ab | nonsynonymous_K2497R |
| 7757 | A | G | 6.32E-05 | | | | | ORF1ab | nonsynonymous_N2498D |
| 7766 | A | G | 6.32E-05 | | | | | ORF1ab | nonsynonymous_I2501V |
| 7771 | T | C | 6.32E-05 | | | | | ORF1ab | synonymous_H2502H |
| 7772 | C | T | 6.32E-05 | | | | | ORF1ab | nonsynonymous_L2503F |
| 7816 | T | A | 6.32E-05 | | | | | ORF1ab | synonymous_S2517S |
| 7818 | T | C | 6.32E-05 | | | | | ORF1ab | nonsynonymous_L2518P |
| 7829 | G | T | 6.32E-05 | | | | | ORF1ab | nonsynonymous_V2522F |
| 7837 | A | C | 6.32E-05 | | | | | ORF1ab | nonsynonymous_L2524F |
| 7857 | A | G | 6.32E-05 | | | | | ORF1ab | nonsynonymous_N2531S |
| 7866 | G | T | 6.32E-05 | | | | | ORF1ab | nonsynonymous_G2534V |
| 7873 | G | T | 6.32E-05 | | | | | ORF1ab | nonsynonymous_L2536F |
| 7878 | T | G | 6.32E-05 | | | | | ORF1ab | nonsynonymous_I2538S |
| 7881 | A | G | 6.32E-05 | | | | | ORF1ab | nonsynonymous_N2539S |
| 7901 | A | G | 6.32E-05 | | | | | ORF1ab | nonsynonymous_K2546E |
| 7919 | T | C | 6.32E-05 | | | | | ORF1ab | nonsynonymous_S2552P |
| 8001 | A | C | 6.32E-05 | | | | | ORF1ab | nonsynonymous_D2579A |
| 8013 | G | A | 6.32E-05 | | | | | ORF1ab | nonsynonymous_S2583N |
| 8017 | G | T | 6.32E-05 | | | | | ORF1ab | synonymous_A2584A |
| 8043 | C | T | 6.32E-05 | | | | | ORF1ab | nonsynonymous_A2593V |
| 8053 | T | C | 6.32E-05 | | | | | ORF1ab | synonymous_N2596N |
| 8068 | T | C | 6.32E-05 | | | | | ORF1ab | synonymous_T2601T |
| 8083 | G | T | 6.32E-05 | | | | | ORF1ab | nonsynonymous_M2606I |
| 808 | A | T | 6.32E-05 | | | | | ORF1ab | synonymous_A181A |
| 8125 | T | C | 6.32E-05 | | | | | ORF1ab | synonymous_L2620L |
| 8127 | C | A | 6.32E-05 | | | | | ORF1ab | nonsynonymous_A2621E |

| | | | | | | | | | |
|---|---|---|---|---|---|---|---|---|---|
| 8134 | T | C | 6.32E-05 | | | | | ORF1ab | synonymous_N2623N |
| 8145 | A | G | 6.32E-05 | | | | | ORF1ab | nonsynonymous_D2627G |
| 8156 | T | G | 6.32E-05 | | | | | ORF1ab | nonsynonymous_S2631A |
| 8160 | C | T | 6.32E-05 | | | | | ORF1ab | nonsynonymous_T2632I |
| 8173 | A | T | 6.32E-05 | | | | | ORF1ab | synonymous_A2636A |
| 8174 | G | A | 6.32E-05 | | | | | ORF1ab | nonsynonymous_A2637T |
| 8182 | A | C | 6.32E-05 | | | | | ORF1ab | nonsynonymous_Q2639H |
| 8223 | A | C | 6.32E-05 | | | | | ORF1ab | nonsynonymous_E2653A |
| 8347 | T | A | 6.32E-05 | | | | | ORF1ab | stopgain_C2694X |
| 8352 | C | T | 6.32E-05 | | | | | ORF1ab | nonsynonymous_A2696V |
| 8353 | G | A | 6.32E-05 | A | 6.32E-05 | | | ORF1ab | synonymous_A2696A; synonymous_A2696A |
| 840 | G | A | 6.32E-05 | | | | | ORF1ab | nonsynonymous_G192D |
| 8422 | G | A | 6.32E-05 | | | | | ORF1ab | nonsynonymous_M2719I |
| 8476 | C | T | 6.32E-05 | | | | | ORF1ab | synonymous_N2737N |
| 8571 | G | T | 6.32E-05 | | | | | ORF1ab | nonsynonymous_W2769L |
| 8605 | C | T | 6.32E-05 | | | | | ORF1ab | synonymous_F2780F |
| 8914 | T | G | 6.32E-05 | | | | | ORF1ab | nonsynonymous_H2883Q |
| 8916 | T | G | 6.32E-05 | | | | | ORF1ab | nonsynonymous_F2884C |
| 8924 | A | G | 6.32E-05 | | | | | ORF1ab | nonsynonymous_R2887G |
| 8926 | A | T | 6.32E-05 | | | | | ORF1ab | nonsynonymous_R2887S |
| 9136 | G | T | 6.32E-05 | | | | | ORF1ab | nonsynonymous_M2957I |
| 9207 | C | T | 6.32E-05 | | | | | ORF1ab | nonsynonymous_S2981F |
| 9208 | T | G | 6.32E-05 | G | 6.32E-05 | | | ORF1ab | synonymous_S2981S; synonymous_S2981S |
| 9519 | T | G | 6.32E-05 | G | 6.32E-05 | | | ORF1ab | nonsynonymous_F3085C; nonsynonymous_F3085C |
| 9541 | C | T | 6.32E-05 | | | | | ORF1ab | synonymous_L3092L |
| 9559 | C | T | 6.32E-05 | | | | | ORF1ab | synonymous_Y3098Y |
| 9652 | G | T | 6.32E-05 | | | | | ORF1ab | nonsynonymous_M3129I |

| | | | | | | | | | |
|---|---|---|---|---|---|---|---|---|---|
| 9653 | G | T | 6.32E-05 | | | | | ORF1ab | nonsynonymous_V3130F |
| 9667 | T | C | 6.32E-05 | | | | | ORF1ab | synonymous_P3134P |
| 9808 | C | T | 6.32E-05 | | | | | ORF1ab | synonymous_C3181C |
| 9817 | G | A | 6.32E-05 | | | | | ORF1ab | synonymous_L3184L |
| 9844 | G | T | 6.32E-05 | | | | | ORF1ab | nonsynonymous_L3193F |
| 9973 | G | A | 6.32E-05 | | | | | ORF1ab | synonymous_K3236K |
| 9053 | G | C | 6.32E-05 | A | 6.32E-05 | A | 6.32E-05 | ORF1ab | nonsynonymous_V2930L; nonsynonymous_V2930I; nonsynonymous_V2930I |
| 10002 | C | T | 6.32E-05 | | | | | ORF1ab | nonsynonymous_S3246F |
| 10006 | T | C | 6.32E-05 | | | | | ORF1ab | synonymous_D3247D |
| 10024 | A | G | 6.32E-05 | | | | | ORF1ab | synonymous_P3253P |
| 10029 | C | T | 6.32E-05 | | | | | ORF1ab | nonsynonymous_T3255I |
| 10038 | C | T | 6.32E-05 | | | | | ORF1ab | nonsynonymous_T3258I |
| 10073 | G | A | 6.32E-05 | | | | | ORF1ab | nonsynonymous_A3270T |
| 10098 | G | A | 6.32E-05 | | | | | ORF1ab | nonsynonymous_G3278D |
| 10152 | A | C | 6.32E-05 | | | | | ORF1ab | nonsynonymous_D3296A |
| 10176 | A | C | 6.32E-05 | | | | | ORF1ab | nonsynonymous_H3304P |
| 10181 | A | C | 6.32E-05 | | | | | ORF1ab | nonsynonymous_I3306L |
| 10186 | C | T | 6.32E-05 | | | | | ORF1ab | synonymous_C3307C |
| 10201 | G | T | 6.32E-05 | | | | | ORF1ab | nonsynonymous_M3312I |
| 10255 | G | T | 6.32E-05 | | | | | ORF1ab | nonsynonymous_L3330F |
| 10256 | G | T | 6.32E-05 | | | | | ORF1ab | nonsynonymous_V3331L |
| 10261 | G | T | 6.32E-05 | | | | | ORF1ab | nonsynonymous_Q3332H |
| 10350 | C | T | 6.32E-05 | | | | | ORF1ab | nonsynonymous_P3362L |
| 10354 | G | T | 6.32E-05 | | | | | ORF1ab | nonsynonymous_K3363N |
| 10396 | G | T | 6.32E-05 | | | | | ORF1ab | synonymous_V3377V |
| 10416 | C | T | 6.32E-05 | | | | | ORF1ab | nonsynonymous_S3384L |
| 10421 | T | C | 6.32E-05 | | | | | ORF1ab | nonsynonymous_S3386P |
| 10448 | C | T | 6.32E-05 | | | | | ORF1ab | nonsynonymous_P3395S |

| | | | | | | | | | |
|---|---|---|---|---|---|---|---|---|---|
| 1044 | C | T | 6.32E-05 | | | | | ORF1ab | nonsynonymous_A260V |
| 10458 | C | T | 6.32E-05 | | | | | ORF1ab | nonsynonymous_T3398I |
| 10512 | A | G | 6.32E-05 | | | | | ORF1ab | nonsynonymous_D3416G |
| 10516 | T | C | 6.32E-05 | | | | | ORF1ab | synonymous_Y3417Y |
| 10532 | T | A | 6.32E-05 | | | | | ORF1ab | nonsynonymous_C3423S |
| 10533 | G | A | 6.32E-05 | | | | | ORF1ab | nonsynonymous_C3423Y |
| 10556 | C | T | 6.32E-05 | | | | | ORF1ab | nonsynonymous_P3431S |
| 10560 | C | T | 6.32E-05 | | | | | ORF1ab | nonsynonymous_T3432I |
| 10562 | G | A | 6.32E-05 | | | | | ORF1ab | nonsynonymous_G3433R |
| 10572 | C | T | 6.32E-05 | | | | | ORF1ab | nonsynonymous_A3436V |
| 10586 | G | C | 6.32E-05 | | | | | ORF1ab | nonsynonymous_E3441Q |
| 10600 | T | C | 6.32E-05 | | | | | ORF1ab | synonymous_Y3445Y |
| 10610 | G | T | 6.32E-05 | | | | | ORF1ab | nonsynonymous_V3449F |
| 10630 | A | G | 6.32E-05 | | | | | ORF1ab | synonymous_Q3455Q |
| 10635 | C | T | 6.32E-05 | | | | | ORF1ab | nonsynonymous_A3457V |
| 10636 | T | C | 6.32E-05 | | | | | ORF1ab | synonymous_A3457A |
| 10642 | G | T | 6.32E-05 | | | | | ORF1ab | synonymous_T3459T |
| 10645 | C | T | 6.32E-05 | | | | | ORF1ab | synonymous_D3460D |
| 10650 | C | T | 6.32E-05 | | | | | ORF1ab | nonsynonymous_T3462I |
| 10667 | T | C | 6.32E-05 | | | | | ORF1ab | synonymous_L3468L |
| 10670 | G | C | 6.32E-05 | | | | | ORF1ab | nonsynonymous_A3469P |
| 1068 | G | A | 6.32E-05 | | | | | ORF1ab | nonsynonymous_G268E |
| 10692 | T | C | 6.32E-05 | | | | | ORF1ab | nonsynonymous_I3476T |
| 10697 | G | A | 6.32E-05 | | | | | ORF1ab | nonsynonymous_G3478R |
| 1069 | G | A | 6.32E-05 | | | | | ORF1ab | synonymous_G268G |
| 10711 | T | C | 6.32E-05 | | | | | ORF1ab | synonymous_F3482F |
| 10720 | A | G | 6.32E-05 | | | | | ORF1ab | synonymous_R3485R |
| 10722 | T | C | 6.32E-05 | | | | | ORF1ab | nonsynonymous_F3486S |
| 10725 | C | T | 6.32E-05 | | | | | ORF1ab | nonsynonymous_T3487I |
| 1072 | A | C | 6.32E-05 | | | | | ORF1ab | nonsynonymous_E269D |

| | | | | | | | | | |
|---|---|---|---|---|---|---|---|---|---|
| 1085 | G | T | 6.32E-05 | | | | | ORF1ab | nonsynonymous_V274L |
| 1096 | A | G | 6.32E-05 | | | | | ORF1ab | synonymous_L277L |
| 1099 | T | C | 6.32E-05 | | | | | ORF1ab | synonymous_N278N |
| 1100 | T | G | 6.32E-05 | | | | | ORF1ab | nonsynonymous_S279A |
| 1106 | A | T | 6.32E-05 | | | | | ORF1ab | nonsynonymous_I281F |
| 11094 | C | T | 6.32E-05 | | | | | ORF1ab | nonsynonymous_A3610V |
| 11101 | A | G | 6.32E-05 | | | | | ORF1ab | synonymous_L3612L |
| 11107 | T | C | 6.32E-05 | | | | | ORF1ab | synonymous_F3614F |
| 11108 | G | A | 6.32E-05 | | | | | ORF1ab | nonsynonymous_A3615T |
| 11113 | G | T | 6.32E-05 | | | | | ORF1ab | nonsynonymous_M3616I |
| 11119 | T | C | 6.32E-05 | | | | | ORF1ab | synonymous_I3618I |
| 11124 | C | T | 6.32E-05 | | | | | ORF1ab | nonsynonymous_A3620V |
| 11138 | G | T | 6.32E-05 | | | | | ORF1ab | nonsynonymous_A3625S |
| 11139 | C | T | 6.32E-05 | | | | | ORF1ab | nonsynonymous_A3625V |
| 1113 | C | T | 6.32E-05 | | | | | ORF1ab | nonsynonymous_T283I |
| 11149 | T | C | 6.32E-05 | | | | | ORF1ab | synonymous_F3628F |
| 11154 | A | C | 6.32E-05 | | | | | ORF1ab | nonsynonymous_K3630T |
| 1115 | A | G | 6.32E-05 | | | | | ORF1ab | nonsynonymous_I284V |
| 1117 | T | C | 6.32E-05 | | | | | ORF1ab | synonymous_I284I |
| 11194 | T | C | 6.32E-05 | | | | | ORF1ab | synonymous_S3643S |
| 1119 | A | C | 6.32E-05 | | | | | ORF1ab | nonsynonymous_Q285P |
| 11202 | C | T | 6.32E-05 | | | | | ORF1ab | nonsynonymous_T3646I |
| 11207 | G | C | 6.32E-05 | | | | | ORF1ab | nonsynonymous_A3648P |
| 11212 | T | C | 6.32E-05 | | | | | ORF1ab | synonymous_Y3649Y |
| 11217 | A | G | 6.32E-05 | | | | | ORF1ab | nonsynonymous_N3651S |
| 1121 | C | T | 6.32E-05 | | | | | ORF1ab | nonsynonymous_P286S |
| 11232 | C | T | 6.32E-05 | | | | | ORF1ab | nonsynonymous_P3656L |
| 11234 | G | A | 6.32E-05 | | | | | ORF1ab | nonsynonymous_A3657T |
| 11235 | C | T | 6.32E-05 | | | | | ORF1ab | nonsynonymous_A3657V |
| 11244 | T | C | 6.32E-05 | | | | | ORF1ab | nonsynonymous_V3660A |

| | | | | | | | | | |
|---|---|---|---|---|---|---|---|---|---|
| 11300 | C | A | 6.32E-05 | A | 6.32E-05 | | | ORF1ab | nonsynonymous_L3679I; nonsynonymous_L3679I |
| 11304 | A | G | 6.32E-05 | | | | | ORF1ab | nonsynonymous_K3680R |
| 11308 | C | T | 6.32E-05 | | | | | ORF1ab | synonymous_D3681D |
| 11310 | G | T | 6.32E-05 | | | | | ORF1ab | nonsynonymous_C3682F |
| 11314 | T | C | 6.32E-05 | | | | | ORF1ab | synonymous_V3683V |
| 11322 | C | T | 6.32E-05 | | | | | ORF1ab | nonsynonymous_A3686V |
| 11345 | C | T | 6.32E-05 | | | | | ORF1ab | nonsynonymous_L3694F |
| 1134 | A | T | 6.32E-05 | | | | | ORF1ab | nonsynonymous_K290M |
| 11356 | A | T | 6.32E-05 | | | | | ORF1ab | synonymous_A3697A |
| 11369 | G | A | 6.32E-05 | | | | | ORF1ab | nonsynonymous_D3702N |
| 11375 | G | A | 6.32E-05 | | | | | ORF1ab | nonsynonymous_G3704S |
| 11399 | A | G | 6.32E-05 | | | | | ORF1ab | nonsynonymous_M3712V |
| 11412 | C | T | 6.32E-05 | | | | | ORF1ab | nonsynonymous_T3716I |
| 11414 | C | T | 6.32E-05 | | | | | ORF1ab | nonsynonymous_L3717F |
| 11419 | T | C | 6.32E-05 | | | | | ORF1ab | synonymous_V3718V |
| 11425 | A | G | 6.32E-05 | | | | | ORF1ab | synonymous_K3720K |
| 11439 | A | G | 6.32E-05 | | | | | ORF1ab | nonsynonymous_N3725S |
| 11442 | C | T | 6.32E-05 | | | | | ORF1ab | nonsynonymous_A3726V |
| 11451 | A | G | 6.32E-05 | | | | | ORF1ab | nonsynonymous_Q3729R |
| 11453 | G | T | 6.32E-05 | | | | | ORF1ab | nonsynonymous_A3730S |
| 11456 | A | T | 6.32E-05 | | | | | ORF1ab | nonsynonymous_I3731F |
| 11460 | C | A | 6.32E-05 | | | | | ORF1ab | nonsynonymous_S3732Y |
| 11491 | T | A | 6.32E-05 | | | | | ORF1ab | synonymous_S3742S |
| 11499 | C | A | 6.32E-05 | | | | | ORF1ab | stopgain_S3745X |
| 11501 | G | C | 6.32E-05 | | | | | ORF1ab | nonsynonymous_G3746R |
| 11527 | G | A | 6.32E-05 | A | 6.32E-05 | | | ORF1ab | synonymous_L3754L; synonymous_L3754L |
| 11528 | G | A | 6.32E-05 | A | 6.32E-05 | | | ORF1ab | nonsynonymous_A3755T; nonsynonymous_A3755T |

| | | | | | | | | | |
|---|---|---|---|---|---|---|---|---|---|
| 11538 | T | C | 6.32E-05 | | | | | ORF1ab | nonsynonymous_I3758T |
| 11540 | G | T | 6.32E-05 | | | | | ORF1ab | nonsynonymous_V3759F |
| 11548 | G | A | 6.32E-05 | | | | | ORF1ab | nonsynonymous_M3761I |
| 11552 | G | T | 6.32E-05 | | | | | ORF1ab | nonsynonymous_V3763F |
| 1155 | T | C | 6.32E-05 | | | | | ORF1ab | nonsynonymous_M297T |
| 11561 | T | C | 6.32E-05 | | | | | ORF1ab | nonsynonymous_C3766R |
| 1156 | G | A | 6.32E-05 | | | | | ORF1ab | nonsynonymous_M297I |
| 11591 | C | T | 6.32E-05 | | | | | ORF1ab | nonsynonymous_L3776F |
| 11601 | T | C | 6.32E-05 | | | | | ORF1ab | nonsynonymous_I3779T |
| 11636 | A | T | 6.32E-05 | | | | | ORF1ab | nonsynonymous_T3791S |
| 11645 | T | A | 6.32E-05 | | | | | ORF1ab | nonsynonymous_F3794I |
| 11665 | C | T | 6.32E-05 | | | | | ORF1ab | synonymous_L3800L |
| 11671 | C | T | 6.32E-05 | | | | | ORF1ab | synonymous_R3802R |
| 11698 | T | C | 6.32E-05 | | | | | ORF1ab | synonymous_Y3811Y |
| 11700 | A | G | 6.32E-05 | | | | | ORF1ab | nonsynonymous_D3812G |
| 11707 | A | G | 6.32E-05 | | | | | ORF1ab | synonymous_L3814L |
| 11709 | T | A | 6.32E-05 | | | | | ORF1ab | nonsynonymous_V3815D |
| 11714 | A | G | 6.32E-05 | | | | | ORF1ab | nonsynonymous_T3817A |
| 11719 | G | A | 6.32E-05 | | | | | ORF1ab | synonymous_Q3818Q |
| 11736 | A | G | 6.32E-05 | | | | | ORF1ab | nonsynonymous_N3824S |
| 11737 | T | C | 6.32E-05 | | | | | ORF1ab | synonymous_N3824N |
| 11742 | A | T | 6.32E-05 | T | 6.32E-05 | | | ORF1ab | nonsynonymous_Q3826L; nonsynonymous_Q3826L |
| 11748 | T | A | 6.32E-05 | | | | | ORF1ab | nonsynonymous_L3828Q |
| 11753 | C | T | 6.32E-05 | | | | | ORF1ab | nonsynonymous_P3830S |
| 11761 | G | T | 6.32E-05 | | | | | ORF1ab | nonsynonymous_K3832N |
| 11764 | T | A | 6.32E-05 | | | | | ORF1ab | nonsynonymous_N3833K |
| 11768 | A | T | 6.32E-05 | | | | | ORF1ab | nonsynonymous_I3835L |
| 11771 | G | A | 6.32E-05 | | | | | ORF1ab | nonsynonymous_D3836N |
| 11788 | C | T | 6.32E-05 | | | | | ORF1ab | synonymous_N3841N |

| | | | | | | | | | |
|---|---|---|---|---|---|---|---|---|---|
| 11789 | A | G | 6.32E-05 | | | | | ORF1ab | nonsynonymous_I3842V |
| 1178 | C | T | 6.32E-05 | | | | | ORF1ab | nonsynonymous_P305S |
| 11795 | T | A | 6.32E-05 | | | | | ORF1ab | nonsynonymous_L3844M |
| 11820 | G | A | 6.32E-05 | | | | | ORF1ab | nonsynonymous_C3852Y |
| 11822 | A | G | 6.32E-05 | | | | | ORF1ab | nonsynonymous_I3853V |
| 11827 | A | T | 6.32E-05 | | | | | ORF1ab | nonsynonymous_K3854N |
| 11830 | A | G | 6.32E-05 | | | | | ORF1ab | synonymous_V3855V |
| 11832 | C | T | 6.32E-05 | | | | | ORF1ab | nonsynonymous_A3856V |
| 11857 | T | C | 6.32E-05 | | | | | ORF1ab | synonymous_D3864D |
| 1185 | C | T | 6.32E-05 | | | | | ORF1ab | nonsynonymous_A307V |
| 11868 | C | T | 6.32E-05 | | | | | ORF1ab | nonsynonymous_T3868I |
| 1186 | G | C | 6.32E-05 | | | | | ORF1ab | synonymous_A307A |
| 11871 | C | T | 6.32E-05 | | | | | ORF1ab | nonsynonymous_S3869L |
| 11875 | A | G | 6.32E-05 | | | | | ORF1ab | synonymous_V3870V |
| 11879 | T | C | 6.32E-05 | | | | | ORF1ab | synonymous_L3872L |
| 11882 | C | T | 6.32E-05 | | | | | ORF1ab | nonsynonymous_L3873F |
| 11884 | C | T | 6.32E-05 | | | | | ORF1ab | synonymous_L3873L |
| 11892 | T | A | 6.32E-05 | | | | | ORF1ab | stopgain_L3876X |
| 11900 | C | T | 6.32E-05 | | | | | ORF1ab | nonsynonymous_L3879F |
| 11918 | T | G | 6.32E-05 | | | | | ORF1ab | nonsynonymous_S3885A |
| 11926 | G | A | 6.32E-05 | | | | | ORF1ab | synonymous_L3887L |
| 1193 | A | G | 6.32E-05 | | | | | ORF1ab | nonsynonymous_N310D |
| 11957 | A | G | 6.32E-05 | | | | | ORF1ab | nonsynonymous_I3898V |
| 11960 | C | A | 6.32E-05 | A | 6.32E-05 | | | ORF1ab | nonsynonymous_L3899I; nonsynonymous_L3899I |
| 11966 | G | T | 6.32E-05 | | | | | ORF1ab | nonsynonymous_A3901S |
| 11987 | T | A | 6.32E-05 | | | | | ORF1ab | nonsynonymous_F3908I |
| 12005 | C | T | 6.32E-05 | | | | | ORF1ab | synonymous_L3914L |
| 12008 | C | T | 6.32E-05 | | | | | ORF1ab | nonsynonymous_L3915F |

| | | | | | | | | | |
|---|---|---|---|---|---|---|---|---|---|
| 12014 | G | A | 6.32E-05 | A | 6.32E-05 | | | ORF1ab | nonsynonymous_V3917I; nonsynonymous_V3917I |
| 12016 | T | C | 6.32E-05 | | | | | ORF1ab | synonymous_V3917V |
| 12045 | T | C | 6.32E-05 | | | | | ORF1ab | nonsynonymous_I3927T |
| 12066 | T | C | 6.32E-05 | | | | | ORF1ab | nonsynonymous_M3934T |
| 12081 | C | T | 6.32E-05 | | | | | ORF1ab | nonsynonymous_A3939V |
| 12082 | A | G | 6.32E-05 | | | | | ORF1ab | synonymous_A3939A |
| 12094 | T | C | 6.32E-05 | | | | | ORF1ab | synonymous_A3943A |
| 12097 | A | G | 6.32E-05 | | | | | ORF1ab | nonsynonymous_I3944M |
| 1210 | G | A | 6.32E-05 | | | | | ORF1ab | nonsynonymous_M315I |
| 12110 | A | G | 6.32E-05 | | | | | ORF1ab | nonsynonymous_S3949G |
| 12116 | C | T | 6.32E-05 | | | | | ORF1ab | nonsynonymous_L3951F |
| 12124 | A | T | 6.32E-05 | | | | | ORF1ab | synonymous_S3953S |
| 12208 | G | T | 6.32E-05 | | | | | ORF1ab | nonsynonymous_K3981N |
| 12225 | C | T | 6.32E-05 | | | | | ORF1ab | nonsynonymous_A3987V |
| 12229 | A | G | 6.32E-05 | | | | | ORF1ab | synonymous_K3988K |
| 12244 | T | C | 6.32E-05 | | | | | ORF1ab | synonymous_R3993R |
| 12248 | G | T | 6.32E-05 | | | | | ORF1ab | nonsynonymous_A3995S |
| 12249 | C | T | 6.32E-05 | | | | | ORF1ab | nonsynonymous_A3995V |
| 12253 | C | T | 6.32E-05 | | | | | ORF1ab | synonymous_A3996A |
| 1225 | C | A | 6.32E-05 | A | 6.32E-05 | | | ORF1ab | synonymous_L320L; synonymous_L320L |
| 12265 | G | C | 6.32E-05 | | | | | ORF1ab | nonsynonymous_K4000N |
| 12268 | G | A | 6.32E-05 | | | | | ORF1ab | synonymous_L4001L |
| 12271 | A | G | 6.32E-05 | | | | | ORF1ab | synonymous_E4002E |
| 12310 | G | A | 6.32E-05 | A | 6.32E-05 | | | ORF1ab | synonymous_Q4015Q; synonymous_Q4015Q |
| 12334 | A | T | 6.32E-05 | | | | | ORF1ab | synonymous_A4023A |
| 12342 | C | T | 6.32E-05 | | | | | ORF1ab | nonsynonymous_T4026I |
| 12347 | G | A | 6.32E-05 | | | | | ORF1ab | nonsynonymous_A4028T |

| | | | | | | | | | |
|---|---|---|---|---|---|---|---|---|---|
| 12358 | A | C | 6.32E-05 | | | | | ORF1ab | synonymous_T4031T |
| 12359 | A | T | 6.32E-05 | | | | | ORF1ab | nonsynonymous_M4032L |
| 12360 | T | C | 6.32E-05 | | | | | ORF1ab | nonsynonymous_M4032T |
| 12361 | G | T | 6.32E-05 | | | | | ORF1ab | nonsynonymous_M4032I |
| 12362 | C | T | 6.32E-05 | | | | | ORF1ab | nonsynonymous_L4033F |
| 12367 | C | A | 6.32E-05 | | | | | ORF1ab | nonsynonymous_F4034L |
| 12369 | C | T | 6.32E-05 | | | | | ORF1ab | nonsynonymous_T4035I |
| 12378 | G | A | 6.32E-05 | | | | | ORF1ab | nonsynonymous_R4038K |
| 1237 | T | C | 6.32E-05 | | | | | ORF1ab | synonymous_D324D |
| 12386 | G | T | 6.32E-05 | | | | | ORF1ab | nonsynonymous_D4041Y |
| 12390 | A | G | 6.32E-05 | | | | | ORF1ab | nonsynonymous_N4042S |
| 12396 | C | T | 6.32E-05 | | | | | ORF1ab | nonsynonymous_A4044V |
| 12403 | C | T | 6.32E-05 | | | | | ORF1ab | synonymous_N4046N |
| 12404 | A | T | 6.32E-05 | | | | | ORF1ab | nonsynonymous_N4047Y |
| 12410 | A | G | 6.32E-05 | | | | | ORF1ab | nonsynonymous_I4049V |
| 12427 | T | C | 6.32E-05 | | | | | ORF1ab | synonymous_D4054D |
| 12434 | G | A | 6.32E-05 | | | | | ORF1ab | nonsynonymous_V4057I |
| 12452 | C | T | 6.32E-05 | | | | | ORF1ab | nonsynonymous_P4063S |
| 1245 | G | A | 6.32E-05 | | | | | ORF1ab | nonsynonymous_G327D |
| 12462 | C | T | 6.32E-05 | | | | | ORF1ab | nonsynonymous_T4066I |
| 12464 | G | T | 6.32E-05 | | | | | ORF1ab | nonsynonymous_A4067S |
| 12467 | G | T | 6.32E-05 | | | | | ORF1ab | nonsynonymous_A4068S |
| 12469 | C | T | 6.32E-05 | | | | | ORF1ab | synonymous_A4068A |
| 12475 | A | T | 6.32E-05 | | | | | ORF1ab | synonymous_L4070L |
| 12479 | G | A | 6.32E-05 | | | | | ORF1ab | nonsynonymous_V4072I |
| 12483 | T | A | 6.32E-05 | | | | | ORF1ab | nonsynonymous_V4073D |
| 12485 | A | T | 6.32E-05 | | | | | ORF1ab | nonsynonymous_I4074L |
| 12489 | C | G | 6.32E-05 | | | | | ORF1ab | nonsynonymous_P4075R |
| 12491 | G | T | 6.32E-05 | | | | | ORF1ab | nonsynonymous_D4076Y |
| 12505 | T | C | 6.32E-05 | | | | | ORF1ab | synonymous_Y4080Y |

| | | | | | | | | | |
|---|---|---|---|---|---|---|---|---|---|
| 12507 | A | G | 6.32E-05 | | | | | ORF1ab | nonsynonymous_K4081R |
| 12511 | T | C | 6.32E-05 | | | | | ORF1ab | synonymous_N4082N |
| 12521 | G | A | 6.32E-05 | | | | | ORF1ab | nonsynonymous_G4086S |
| 12522 | G | A | 6.32E-05 | | | | | ORF1ab | nonsynonymous_G4086D |
| 12530 | T | C | 6.32E-05 | | | | | ORF1ab | nonsynonymous_F4089L |
| 12533 | A | G | 6.32E-05 | | | | | ORF1ab | nonsynonymous_T4090A |
| 12538 | T | A | 6.32E-05 | | | | | ORF1ab | stopgain_Y4091X |
| 12545 | G | A | 6.32E-05 | | | | | ORF1ab | nonsynonymous_A4094T |
| 12547 | A | T | 6.32E-05 | | | | | ORF1ab | synonymous_A4094A |
| 12551 | T | A | 6.32E-05 | | | | | ORF1ab | nonsynonymous_W4096R |
| 12555 | A | C | 6.32E-05 | | | | | ORF1ab | nonsynonymous_E4097A |
| 12566 | G | T | 6.32E-05 | | | | | ORF1ab | nonsynonymous_V4101F |
| 12571 | A | G | 6.32E-05 | | | | | ORF1ab | synonymous_V4102V |
| 12572 | G | T | 6.32E-05 | | | | | ORF1ab | nonsynonymous_D4103Y |
| 12578 | G | T | 6.32E-05 | | | | | ORF1ab | nonsynonymous_D4105Y |
| 12582 | G | T | 6.32E-05 | | | | | ORF1ab | nonsynonymous_S4106I |
| 12597 | T | A | 6.32E-05 | | | | | ORF1ab | nonsynonymous_L4111H |
| 12600 | G | A | 6.32E-05 | | | | | ORF1ab | nonsynonymous_S4112N |
| 12605 | A | G | 6.32E-05 | | | | | ORF1ab | nonsynonymous_I4114V |
| 12629 | T | C | 6.32E-05 | | | | | ORF1ab | synonymous_L4122L |
| 1262 | A | G | 6.32E-05 | | | | | ORF1ab | nonsynonymous_T333A |
| 12631 | A | G | 6.32E-05 | | | | | ORF1ab | synonymous_L4122L |
| 12647 | G | A | 6.32E-05 | A | 6.32E-05 | | | ORF1ab | nonsynonymous_V4128I; nonsynonymous_V4128I |
| 1264 | G | A | 6.32E-05 | A | 6.32E-05 | | | ORF1ab | synonymous_T333T; synonymous_T333T |
| 12654 | C | A | 6.32E-05 | | | | | ORF1ab | nonsynonymous_A4130D |
| 1265 | G | A | 6.32E-05 | | | | | ORF1ab | nonsynonymous_G334S |
| 12660 | G | C | 6.32E-05 | | | | | ORF1ab | nonsynonymous_R4132T |
| 12661 | G | T | 6.32E-05 | | | | | ORF1ab | nonsynonymous_R4132S |

| | | | | | | | | | |
|---|---|---|---|---|---|---|---|---|---|
| 12676 | C | T | 6.32E-05 | | | | | ORF1ab | synonymous_V4137V |
| 12686 | A | T | 6.32E-05 | | | | | ORF1ab | nonsynonymous_N4141Y |
| 1268 | G | A | 6.32E-05 | | | | | ORF1ab | nonsynonymous_D335N |
| 12691 | T | C | 6.32E-05 | | | | | ORF1ab | synonymous_N4142N |
| 1269 | A | G | 6.32E-05 | | | | | ORF1ab | nonsynonymous_D335G |
| 12704 | G | T | 6.32E-05 | | | | | ORF1ab | nonsynonymous_V4147F |
| 1270 | T | C | 6.32E-05 | | | | | ORF1ab | synonymous_D335D |
| 12730 | T | G | 6.32E-05 | | | | | ORF1ab | synonymous_A4155A |
| 12744 | A | T | 6.32E-05 | | | | | ORF1ab | nonsynonymous_Q4160L |
| 12745 | A | G | 6.32E-05 | | | | | ORF1ab | synonymous_Q4160Q |
| 1274 | G | T | 6.32E-05 | | | | | ORF1ab | nonsynonymous_V337F |
| 12773 | G | T | 6.32E-05 | | | | | ORF1ab | nonsynonymous_A4170S |
| 12776 | T | G | 6.32E-05 | | | | | ORF1ab | nonsynonymous_Y4171D |
| 1278 | A | G | 6.32E-05 | | | | | ORF1ab | nonsynonymous_K338R |
| 12792 | A | T | 6.32E-05 | | | | | ORF1ab | nonsynonymous_K4176M |
| 12794 | G | T | 6.32E-05 | | | | | ORF1ab | stopgain_G4177X |
| 12798 | G | A | 6.32E-05 | | | | | ORF1ab | nonsynonymous_G4178D |
| 12806 | G | A | 6.32E-05 | | | | | ORF1ab | nonsynonymous_V4181I |
| 12839 | A | C | 6.32E-05 | | | | | ORF1ab | nonsynonymous_K4192Q |
| 12845 | G | T | 6.32E-05 | | | | | ORF1ab | nonsynonymous_A4194S |
| 12849 | G | A | 6.32E-05 | | | | | ORF1ab | nonsynonymous_R4195K |
| 12866 | G | A | 6.32E-05 | | | | | ORF1ab | nonsynonymous_G4201R |
| 12870 | C | T | 6.32E-05 | | | | | ORF1ab | nonsynonymous_T4202I |
| 12876 | C | T | 6.32E-05 | | | | | ORF1ab | nonsynonymous_T4204I |
| 12897 | C | A | 6.32E-05 | A | 6.32E-05 | | | ORF1ab | nonsynonymous_P4211Q; nonsynonymous_P4211Q |
| 12907 | G | T | 6.32E-05 | | | | | ORF1ab | nonsynonymous_R4214S |
| 12910 | T | A | 6.32E-05 | | | | | ORF1ab | nonsynonymous_F4215L |
| 12911 | G | T | 6.32E-05 | | | | | ORF1ab | nonsynonymous_V4216F |
| 12918 | A | G | 6.32E-05 | | | | | ORF1ab | nonsynonymous_D4218G |

| | | | | | | | | | |
|---|---|---|---|---|---|---|---|---|---|
| 12936 | A | C | 6.32E-05 | | | | | ORF1ab | nonsynonymous_K4224T |
| 12939 | T | G | 6.32E-05 | | | | | ORF1ab | nonsynonymous_V4225G |
| 12940 | G | T | 6.32E-05 | | | | | ORF1ab | synonymous_V4225V |
| 12950 | T | A | 6.32E-05 | | | | | ORF1ab | nonsynonymous_Y4229N |
| 12964 | A | G | 6.32E-05 | | | | | ORF1ab | synonymous_G4233G |
| 12976 | A | T | 6.32E-05 | | | | | ORF1ab | synonymous_L4237L |
| 12979 | T | C | 6.32E-05 | | | | | ORF1ab | synonymous_N4238N |
| 12982 | A | G | 6.32E-05 | | | | | ORF1ab | synonymous_R4239R |
| 12992 | C | T | 6.32E-05 | | | | | ORF1ab | nonsynonymous_L4243F |
| 12995 | G | C | 6.32E-05 | | | | | ORF1ab | nonsynonymous_G4244R |
| 12998 | A | G | 6.32E-05 | | | | | ORF1ab | nonsynonymous_S4245G |
| 1299 | G | T | 6.32E-05 | | | | | ORF1ab | nonsynonymous_G345V |
| 13002 | T | C | 6.32E-05 | | | | | ORF1ab | nonsynonymous_L4246S |
| 13003 | A | T | 6.32E-05 | | | | | ORF1ab | nonsynonymous_L4246F |
| 13004 | G | T | 6.32E-05 | | | | | ORF1ab | nonsynonymous_A4247S |
| 13005 | C | T | 6.32E-05 | | | | | ORF1ab | nonsynonymous_A4247V |
| 13026 | C | T | 6.32E-05 | | | | | ORF1ab | nonsynonymous_A4254V |
| 13046 | C | T | 6.32E-05 | | | | | ORF1ab | nonsynonymous_P4261S |
| 13047 | C | T | 6.32E-05 | | | | | ORF1ab | nonsynonymous_P4261L |
| 13077 | C | T | 6.32E-05 | | | | | ORF1ab | nonsynonymous_A4271V |
| 13078 | T | C | 6.32E-05 | | | | | ORF1ab | synonymous_A4271A |
| 13082 | G | T | 6.32E-05 | | | | | ORF1ab | nonsynonymous_A4273S |
| 13090 | T | C | 6.32E-05 | | | | | ORF1ab | synonymous_D4275D |
| 13105 | C | T | 6.32E-05 | | | | | ORF1ab | synonymous_Y4280Y |
| 13112 | T | A | 6.32E-05 | | | | | ORF1ab | nonsynonymous_Y4283N |
| 1329 | C | T | 6.32E-05 | | | | | ORF1ab | nonsynonymous_T355I |
| 1333 | T | A | 6.32E-05 | | | | | ORF1ab | synonymous_T356T |
| 1334 | T | A | 6.32E-05 | | | | | ORF1ab | nonsynonymous_C357S |
| 1347 | C | T | 6.32E-05 | | | | | ORF1ab | nonsynonymous_P361L |
| 1359 | T | C | 6.32E-05 | | | | | ORF1ab | nonsynonymous_V365A |

| | | | | | | | | | |
|---|---|---|---|---|---|---|---|---|---|
| 1372 | T | C | 6.32E-05 | | | | | ORF1ab | synonymous_Y369Y |
| 1376 | C | T | 6.32E-05 | | | | | ORF1ab | nonsynonymous_P371S |
| 1381 | A | G | 6.32E-05 | | | | | ORF1ab | synonymous_A372A |
| 1399 | A | T | 6.32E-05 | T | 6.32E-05 | | | ORF1ab | synonymous_V378V;<br>synonymous_V378V |
| 1405 | T | C | 6.32E-05 | | | | | ORF1ab | synonymous_P380P |
| 1417 | T | G | 6.32E-05 | | | | | ORF1ab | synonymous_L384L |
| 1458 | G | T | 6.32E-05 | | | | | ORF1ab | nonsynonymous_R398L |
| 1461 | A | G | 6.32E-05 | | | | | ORF1ab | nonsynonymous_K399R |
| 1466 | G | T | 6.32E-05 | | | | | ORF1ab | nonsynonymous_G401C |
| 1467 | G | A | 6.32E-05 | | | | | ORF1ab | nonsynonymous_G401D |
| 1560 | G | T | 6.32E-05 | | | | | ORF1ab | nonsynonymous_G432V |
| 1566 | A | C | 6.32E-05 | | | | | ORF1ab | nonsynonymous_N434T |
| 1568 | C | T | 6.32E-05 | | | | | ORF1ab | nonsynonymous_H435Y |
| 1582 | T | C | 6.32E-05 | | | | | ORF1ab | synonymous_V439V |
| 1583 | G | A | 6.32E-05 | | | | | ORF1ab | nonsynonymous_G440R |
| 1601 | C | A | 6.32E-05 | | | | | ORF1ab | nonsynonymous_L446I |
| 1622 | A | G | 6.32E-05 | | | | | ORF1ab | nonsynonymous_I453V |
| 1632 | A | T | 6.32E-05 | | | | | ORF1ab | nonsynonymous_K456I |
| 1635 | A | C | 6.32E-05 | | | | | ORF1ab | nonsynonymous_E457A |
| 1805 | G | A | 6.32E-05 | | | | | ORF1ab | nonsynonymous_G514R |
| 1825 | C | T | 6.32E-05 | | | | | ORF1ab | synonymous_A520A |
| 1835 | G | A | 6.32E-05 | | | | | ORF1ab | nonsynonymous_G524S |
| 1842 | A | G | 6.32E-05 | | | | | ORF1ab | nonsynonymous_Q526R |
| 1848 | C | T | 6.32E-05 | | | | | ORF1ab | nonsynonymous_S528L |
| 1852 | A | G | 6.32E-05 | | | | | ORF1ab | nonsynonymous_I529M |
| 1858 | T | G | 6.32E-05 | | | | | ORF1ab | nonsynonymous_S531R |
| 1859 | C | T | 6.32E-05 | | | | | ORF1ab | nonsynonymous_P532S |
| 1860 | C | T | 6.32E-05 | | | | | ORF1ab | nonsynonymous_P532L |

| | | | | | | | | | |
|---|---|---|---|---|---|---|---|---|---|
| 1874 | G | A | 6.32E-05 | A | 6.32E-05 | | | ORF1ab | nonsynonymous_A537T; nonsynonymous_A537T |
| 1878 | C | T | 6.32E-05 | | | | | ORF1ab | nonsynonymous_S538L |
| 1889 | C | T | 6.32E-05 | | | | | ORF1ab | nonsynonymous_R542C |
| 1891 | T | G | 6.32E-05 | | | | | ORF1ab | synonymous_R542R |
| 1896 | T | C | 6.32E-05 | | | | | ORF1ab | nonsynonymous_V544A |
| 1900 | A | G | 6.32E-05 | | | | | ORF1ab | synonymous_R545R |
| 1906 | T | C | 6.32E-05 | | | | | ORF1ab | synonymous_I547I |
| 1913 | C | T | 6.32E-05 | | | | | ORF1ab | nonsynonymous_R550C |
| 1915 | C | T | 6.32E-05 | | | | | ORF1ab | synonymous_R550R |
| 1936 | T | C | 6.32E-05 | | | | | ORF1ab | synonymous_N557N |
| 1938 | C | T | 6.32E-05 | | | | | ORF1ab | nonsynonymous_S558F |
| 2227 | C | T | 6.32E-05 | | | | | ORF1ab | synonymous_T654T |
| 2228 | T | G | 6.32E-05 | | | | | ORF1ab | nonsynonymous_C655G |
| 2229 | G | A | 6.32E-05 | | | | | ORF1ab | nonsynonymous_C655Y |
| 2231 | G | T | 6.32E-05 | | | | | ORF1ab | nonsynonymous_A656S |
| 2239 | A | C | 6.32E-05 | | | | | ORF1ab | nonsynonymous_E658D |
| 2246 | G | A | 6.32E-05 | A | 6.32E-05 | | | ORF1ab | nonsynonymous_G661S; nonsynonymous_G661S |
| 2250 | G | A | 6.32E-05 | | | | | ORF1ab | nonsynonymous_G662E |
| 2254 | A | G | 6.32E-05 | | | | | ORF1ab | synonymous_Q663Q |
| 2274 | A | C | 6.32E-05 | | | | | ORF1ab | nonsynonymous_E670A |
| 2277 | T | C | 6.32E-05 | | | | | ORF1ab | nonsynonymous_I671T |
| 2281 | G | A | 6.32E-05 | A | 6.32E-05 | | | ORF1ab | synonymous_K672K; synonymous_K672K |
| 2286 | G | A | 6.32E-05 | | | | | ORF1ab | nonsynonymous_S674N |
| 2290 | T | G | 6.32E-05 | G | 6.32E-05 | | | ORF1ab | synonymous_V675V; synonymous_V675V |
| 2323 | G | T | 6.32E-05 | | | | | ORF1ab | nonsynonymous_L686F |
| 2434 | C | T | 6.32E-05 | | | | | ORF1ab | synonymous_S723S |

| | | | | | | | | | |
|---|---|---|---|---|---|---|---|---|---|
| 2443 | A | G | 6.32E-05 | | | | | ORF1ab | synonymous_E726E |
| 2444 | A | G | 6.32E-05 | | | | | ORF1ab | nonsynonymous_T727A |
| 2448 | G | A | 6.32E-05 | A | 6.32E-05 | | | ORF1ab | nonsynonymous_G728D;<br>nonsynonymous_G728D |
| 2456 | A | T | 6.32E-05 | | | | | ORF1ab | nonsynonymous_M731L |
| 2458 | G | T | 6.32E-05 | | | | | ORF1ab | nonsynonymous_M731I |
| 2461 | T | A | 6.32E-05 | | | | | ORF1ab | synonymous_P732P |
| 2468 | G | A | 6.32E-05 | | | | | ORF1ab | nonsynonymous_A735T |
| 2471 | C | T | 6.32E-05 | | | | | ORF1ab | nonsynonymous_P736S |
| 2475 | A | T | 6.32E-05 | T | 6.32E-05 | | | ORF1ab | nonsynonymous_K737I;<br>nonsynonymous_K737I |
| 2488 | C | T | 6.32E-05 | | | | | ORF1ab | synonymous_F741F |
| 2494 | G | A | 6.32E-05 | | | | | ORF1ab | synonymous_E743E |
| 2497 | A | G | 6.32E-05 | | | | | ORF1ab | synonymous_G744G |
| 2500 | A | G | 6.32E-05 | | | | | ORF1ab | synonymous_E745E |
| 2502 | C | A | 6.32E-05 | | | | | ORF1ab | nonsynonymous_T746K |
| 2522 | A | G | 6.32E-05 | | | | | ORF1ab | nonsynonymous_T753A |
| 2533 | T | A | 6.32E-05 | | | | | ORF1ab | synonymous_V756V |
| 2534 | G | A | 6.32E-05 | | | | | ORF1ab | nonsynonymous_V757I |
| 2542 | A | G | 6.32E-05 | | | | | ORF1ab | synonymous_K759K |
| 2543 | A | G | 6.32E-05 | | | | | ORF1ab | nonsynonymous_T760A |
| 2549 | G | C | 6.32E-05 | | | | | ORF1ab | nonsynonymous_D762H |
| 2556 | A | T | 6.32E-05 | | | | | ORF1ab | nonsynonymous_Q764L |
| 2559 | C | T | 6.32E-05 | | | | | ORF1ab | nonsynonymous_P765L |
| 2561 | T | C | 6.32E-05 | | | | | ORF1ab | synonymous_L766L |
| 2851 | T | C | 6.32E-05 | | | | | ORF1ab | synonymous_V862V |
| 2863 | A | G | 6.32E-05 | | | | | ORF1ab | synonymous_T866T |
| 2871 | A | G | 6.32E-05 | | | | | ORF1ab | nonsynonymous_N869S |
| 2878 | C | T | 6.32E-05 | | | | | ORF1ab | synonymous_F871F |
| 2879 | G | T | 6.32E-05 | | | | | ORF1ab | nonsynonymous_A872S |

| | | | | | | | | | |
|---|---|---|---|---|---|---|---|---|---|
| 2881 | C | A | 6.32E-05 | | | | | ORF1ab | synonymous_A872A |
| 3168 | C | T | 6.32E-05 | | | | | ORF1ab | nonsynonymous_A968V |
| 3170 | C | T | 6.32E-05 | | | | | ORF1ab | nonsynonymous_L969F |
| 3174 | A | T | 6.32E-05 | | | | | ORF1ab | nonsynonymous_Q970L |
| 3175 | A | T | 6.32E-05 | T | 6.32E-05 | | | ORF1ab | nonsynonymous_Q970H; nonsynonymous_Q970H |
| 3179 | G | A | 6.32E-05 | | | | | ORF1ab | nonsynonymous_E972K |
| 3224 | A | G | 6.32E-05 | | | | | ORF1ab | nonsynonymous_T987A |
| 3235 | A | G | 6.32E-05 | G | 6.32E-05 | | | ORF1ab | synonymous_Q990Q; synonymous_Q990Q |
| 3244 | C | T | 6.32E-05 | | | | | ORF1ab | synonymous_G993G |
| 3245 | A | T | 6.32E-05 | | | | | ORF1ab | nonsynonymous_S994C |
| 3248 | G | A | 6.32E-05 | | | | | ORF1ab | nonsynonymous_E995K |
| 3260 | A | G | 6.32E-05 | | | | | ORF1ab | nonsynonymous_T999A |
| 3263 | A | G | 6.32E-05 | | | | | ORF1ab | nonsynonymous_T1000A |
| 3264 | C | T | 6.32E-05 | | | | | ORF1ab | nonsynonymous_T1000I |
| 3283 | T | A | 6.32E-05 | | | | | ORF1ab | synonymous_V1006V |
| 3287 | G | T | 6.32E-05 | | | | | ORF1ab | nonsynonymous_V1008F |
| 3299 | T | C | 6.32E-05 | | | | | ORF1ab | synonymous_L1012L |
| 3304 | G | A | 6.32E-05 | | | | | ORF1ab | synonymous_E1013E |
| 3307 | G | A | 6.32E-05 | | | | | ORF1ab | nonsynonymous_M1014I |
| 3486 | C | T | 6.32E-05 | | | | | ORF1ab | nonsynonymous_A1074V |
| 3511 | C | T | 6.32E-05 | | | | | ORF1ab | synonymous_A1082A |
| 3539 | G | T | 6.32E-05 | | | | | ORF1ab | nonsynonymous_A1092S |
| 3540 | C | T | 6.32E-05 | | | | | ORF1ab | nonsynonymous_A1092V |
| 3541 | T | C | 6.32E-05 | | | | | ORF1ab | synonymous_A1092A |
| 3546 | A | T | 6.32E-05 | | | | | ORF1ab | nonsynonymous_N1094I |
| 3548 | G | A | 6.32E-05 | | | | | ORF1ab | nonsynonymous_G1095R |
| 3552 | C | T | 6.32E-05 | | | | | ORF1ab | nonsynonymous_P1096L |
| 3566 | G | A | 6.32E-05 | | | | | ORF1ab | nonsynonymous_G1101S |

| | | | | | | | | | |
|---|---|---|---|---|---|---|---|---|---|
| 3569 | A | G | 6.32E-05 | | | | | ORF1ab | nonsynonymous_S1102G |
| 3580 | A | G | 6.32E-05 | | | | | ORF1ab | synonymous_L1105L |
| 3604 | C | T | 6.32E-05 | | | | | ORF1ab | synonymous_H1113H |
| 3607 | T | C | 6.32E-05 | | | | | ORF1ab | synonymous_C1114C |
| 3608 | C | A | 6.32E-05 | | | | | ORF1ab | nonsynonymous_L1115I |
| 3610 | T | C | 6.32E-05 | | | | | ORF1ab | synonymous_L1115L |
| 3660 | A | C | 6.32E-05 | | | | | ORF1ab | nonsynonymous_K1132T |
| 3666 | C | T | 6.32E-05 | | | | | ORF1ab | nonsynonymous_A1134V |
| 3685 | G | T | 6.32E-05 | | | | | ORF1ab | nonsynonymous_Q1140H |
| 3686 | C | A | 6.32E-05 | A | 6.32E-05 | | | ORF1ab | nonsynonymous_H1141N; nonsynonymous_H1141N |
| 3719 | G | A | 6.32E-05 | | | | | ORF1ab | nonsynonymous_G1152S |
| 3729 | G | A | 6.32E-05 | | | | | ORF1ab | nonsynonymous_G1155D |
| 3730 | T | C | 6.32E-05 | | | | | ORF1ab | synonymous_G1155G |
| 3732 | C | T | 6.32E-05 | | | | | ORF1ab | nonsynonymous_A1156V |
| 3745 | T | C | 6.32E-05 | | | | | ORF1ab | synonymous_H1160H |
| 3763 | A | G | 6.32E-05 | | | | | ORF1ab | synonymous_V1166V |
| 3765 | A | G | 6.32E-05 | | | | | ORF1ab | nonsynonymous_D1167G |
| 3790 | A | G | 6.32E-05 | | | | | ORF1ab | synonymous_L1175L |
| 3801 | A | C | 6.32E-05 | | | | | ORF1ab | nonsynonymous_D1179A |
| 4090 | C | T | 6.32E-05 | | | | | ORF1ab | synonymous_D1275D |
| 4091 | A | G | 6.32E-05 | | | | | ORF1ab | nonsynonymous_I1276V |
| 409 | T | A | 6.32E-05 | | | | | ORF1ab | nonsynonymous_D48E |
| 410 | G | T | 6.32E-05 | | | | | ORF1ab | nonsynonymous_G49C |
| 4111 | T | C | 6.32E-05 | | | | | ORF1ab | synonymous_D1282D |
| 4114 | T | G | 6.32E-05 | | | | | ORF1ab | synonymous_A1283A |
| 4117 | A | G | 6.32E-05 | | | | | ORF1ab | synonymous_P1284P |
| 4123 | A | G | 6.32E-05 | | | | | ORF1ab | nonsynonymous_I1286M |
| 4138 | T | C | 6.32E-05 | | | | | ORF1ab | synonymous_V1291V |
| 4149 | T | C | 6.32E-05 | | | | | ORF1ab | nonsynonymous_V1295A |

| | | | | | | | | | |
|---|---|---|---|---|---|---|---|---|---|
| 4198 | A | T | 6.32E-05 | T | 6.32E-05 | | | ORF1ab | nonsynonymous_E1311D; nonsynonymous_E1311D |
| 4200 | T | G | 6.32E-05 | | | | | ORF1ab | nonsynonymous_M1312R |
| 4208 | A | C | 6.32E-05 | | | | | ORF1ab | nonsynonymous_K1315Q |
| 421 | C | T | 6.32E-05 | | | | | ORF1ab | synonymous_G52G |
| 4220 | A | G | 6.32E-05 | | | | | ORF1ab | nonsynonymous_K1319E |
| 4226 | C | T | 6.32E-05 | | | | | ORF1ab | nonsynonymous_P1321S |
| 4234 | C | T | 6.32E-05 | | | | | ORF1ab | synonymous_D1323D |
| 4266 | T | C | 6.32E-05 | | | | | ORF1ab | nonsynonymous_L1334S |
| 4273 | T | G | 6.32E-05 | | | | | ORF1ab | synonymous_G1336G |
| 4274 | T | A | 6.32E-05 | | | | | ORF1ab | nonsynonymous_Y1337N |
| 4282 | A | G | 6.32E-05 | | | | | ORF1ab | synonymous_V1339V |
| 428 | G | A | 6.32E-05 | | | | | ORF1ab | nonsynonymous_E55K |
| 4305 | A | G | 6.32E-05 | | | | | ORF1ab | nonsynonymous_K1347R |
| 4331 | C | T | 6.32E-05 | | | | | ORF1ab | synonymous_L1356L |
| 4336 | A | G | 6.32E-05 | | | | | ORF1ab | synonymous_P1357P |
| 4345 | C | T | 6.32E-05 | | | | | ORF1ab | synonymous_I1360I |
| 4346 | T | A | 6.32E-05 | | | | | ORF1ab | nonsynonymous_S1361T |
| 4351 | T | C | 6.32E-05 | | | | | ORF1ab | synonymous_N1362N |
| 4360 | A | C | 6.32E-05 | | | | | ORF1ab | nonsynonymous_Q1365H |
| 4370 | G | A | 6.32E-05 | | | | | ORF1ab | nonsynonymous_G1369R |
| 4392 | G | A | 6.32E-05 | | | | | ORF1ab | nonsynonymous_R1376Q |
| 4393 | A | T | 6.32E-05 | | | | | ORF1ab | synonymous_R1376R |
| 4396 | A | G | 6.32E-05 | | | | | ORF1ab | synonymous_E1377E |
| 439 | A | G | 6.32E-05 | | | | | ORF1ab | synonymous_K58K |
| 4400 | C | T | 6.32E-05 | | | | | ORF1ab | nonsynonymous_L1379F |
| 4414 | A | G | 6.32E-05 | | | | | ORF1ab | synonymous_E1383E |
| 4426 | A | G | 6.32E-05 | | | | | ORF1ab | synonymous_K1387K |
| 4427 | T | C | 6.32E-05 | | | | | ORF1ab | synonymous_L1388L |

| | | | | | | | | | |
|---|---|---|---|---|---|---|---|---|---|
| 4436 | G | A | 6.32E-05 | A | 6.32E-05 | | | ORF1ab | nonsynonymous_V1391I; nonsynonymous_V1391I |
| 4448 | A | G | 6.32E-05 | | | | | ORF1ab | nonsynonymous_T1395A |
| 4457 | A | G | 6.32E-05 | | | | | ORF1ab | nonsynonymous_I1398V |
| 4463 | T | C | 6.32E-05 | | | | | ORF1ab | nonsynonymous_S1400P |
| 4467 | C | T | 6.32E-05 | | | | | ORF1ab | nonsynonymous_T1401I |
| 4473 | A | G | 6.32E-05 | | | | | ORF1ab | nonsynonymous_Q1403R |
| 4475 | C | T | 6.32E-05 | | | | | ORF1ab | nonsynonymous_R1404C |
| 447 | T | G | 6.32E-05 | | | | | ORF1ab | nonsynonymous_L61W |
| 4495 | A | G | 6.32E-05 | | | | | ORF1ab | synonymous_K1410K |
| 449 | C | T | 6.32E-05 | | | | | ORF1ab | nonsynonymous_P62S |
| 4508 | G | A | 6.32E-05 | | | | | ORF1ab | nonsynonymous_V1415M |
| 450 | C | T | 6.32E-05 | | | | | ORF1ab | nonsynonymous_P62L |
| 4528 | A | G | 6.32E-05 | | | | | ORF1ab | synonymous_R1421R |
| 4545 | G | T | 6.32E-05 | | | | | ORF1ab | nonsynonymous_S1427I |
| 4546 | T | A | 6.32E-05 | | | | | ORF1ab | nonsynonymous_S1427R |
| 4547 | A | G | 6.32E-05 | | | | | ORF1ab | nonsynonymous_K1428E |
| 4550 | A | G | 6.32E-05 | | | | | ORF1ab | nonsynonymous_T1429A |
| 4555 | T | A | 6.32E-05 | | | | | ORF1ab | synonymous_T1430T |
| 4560 | C | T | 6.32E-05 | | | | | ORF1ab | nonsynonymous_A1432V |
| 4567 | T | A | 6.32E-05 | | | | | ORF1ab | synonymous_L1434L |
| 4573 | C | T | 6.32E-05 | | | | | ORF1ab | synonymous_N1436N |
| 4575 | C | T | 6.32E-05 | | | | | ORF1ab | nonsynonymous_T1437I |
| 4577 | C | T | 6.32E-05 | | | | | ORF1ab | nonsynonymous_L1438F |
| 457 | T | C | 6.32E-05 | | | | | ORF1ab | synonymous_L64L |
| 4580 | A | C | 6.32E-05 | | | | | ORF1ab | nonsynonymous_N1439H |
| 4595 | A | G | 6.32E-05 | G | 6.32E-05 | | | ORF1ab | nonsynonymous_T1444A; nonsynonymous_T1444A |
| 4596 | C | T | 6.32E-05 | | | | | ORF1ab | nonsynonymous_T1444I |
| 4597 | T | C | 6.32E-05 | | | | | ORF1ab | synonymous_T1444T |

| | | | | | | | | | |
|---|---|---|---|---|---|---|---|---|---|
| 4608 | T | A | 6.32E-05 | | | | | ORF1ab | nonsynonymous_M1448K |
| 460 | A | G | 6.32E-05 | | | | | ORF1ab | synonymous_E65E |
| 4624 | A | G | 6.32E-05 | | | | | ORF1ab | synonymous_V1453V |
| 4630 | T | C | 6.32E-05 | | | | | ORF1ab | synonymous_H1455H |
| 4632 | G | A | 6.32E-05 | | | | | ORF1ab | nonsynonymous_G1456D |
| 4642 | G | A | 6.32E-05 | | | | | ORF1ab | synonymous_L1459L |
| 4644 | A | G | 6.32E-05 | | | | | ORF1ab | nonsynonymous_E1460G |
| 4649 | G | T | 6.32E-05 | | | | | ORF1ab | nonsynonymous_A1462S |
| 469 | T | C | 6.32E-05 | | | | | ORF1ab | synonymous_Y68Y |
| 4779 | T | C | 6.32E-05 | | | | | ORF1ab | nonsynonymous_I1505T |
| 4784 | C | T | 6.32E-05 | | | | | ORF1ab | nonsynonymous_L1507F |
| 478 | C | T | 6.32E-05 | | | | | ORF1ab | synonymous_I71I |
| 4790 | G | A | 6.32E-05 | | | | | ORF1ab | nonsynonymous_G1509S |
| 4810 | C | T | 6.32E-05 | | | | | ORF1ab | synonymous_S1515S |
| 4870 | A | G | 6.32E-05 | | | | | ORF1ab | synonymous_V1535V |
| 4898 | C | T | 6.32E-05 | | | | | ORF1ab | nonsynonymous_H1545Y |
| 4910 | G | T | 6.32E-05 | | | | | ORF1ab | stopgain_E1549X |
| 4921 | C | T | 6.32E-05 | | | | | ORF1ab | synonymous_T1552T |
| 4922 | T | C | 6.32E-05 | | | | | ORF1ab | nonsynonymous_F1553L |
| 4927 | C | T | 6.32E-05 | | | | | ORF1ab | synonymous_D1554D |
| 4947 | C | T | 6.32E-05 | | | | | ORF1ab | nonsynonymous_S1561F |
| 4951 | G | A | 6.32E-05 | | | | | ORF1ab | synonymous_L1562L |
| 4955 | G | A | 6.32E-05 | | | | | ORF1ab | nonsynonymous_E1564K |
| 4956 | A | G | 6.32E-05 | | | | | ORF1ab | nonsynonymous_E1564G |
| 4962 | G | T | 6.32E-05 | | | | | ORF1ab | nonsynonymous_R1566M |
| 4987 | A | G | 6.32E-05 | | | | | ORF1ab | synonymous_V1574V |
| 4992 | A | C | 6.32E-05 | | | | | ORF1ab | nonsynonymous_N1576T |
| 533 | G | A | 6.32E-05 | | | | | ORF1ab | nonsynonymous_A90T |
| 535 | A | G | 6.32E-05 | | | | | ORF1ab | synonymous_A90A |
| 556 | C | T | 6.32E-05 | | | | | ORF1ab | synonymous_Y97Y |

| | | | | | | | | | |
|---|---|---|---|---|---|---|---|---|---|
| 5618 | A | G | 6.32E-05 | | | | | ORF1ab | nonsynonymous_I1785V |
| 5620 | A | G | 6.32E-05 | | | | | ORF1ab | nonsynonymous_I1785M |
| 5637 | A | G | 6.32E-05 | | | | | ORF1ab | nonsynonymous_K1791R |
| 5643 | C | T | 6.32E-05 | | | | | ORF1ab | nonsynonymous_A1793V |
| 5646 | C | T | 6.32E-05 | | | | | ORF1ab | nonsynonymous_T1794I |
| 5656 | A | G | 6.32E-05 | | | | | ORF1ab | synonymous_L1797L |
| 5658 | T | A | 6.32E-05 | | | | | ORF1ab | nonsynonymous_V1798E |
| 5678 | G | A | 6.32E-05 | | | | | ORF1ab | nonsynonymous_V1805I |
| 5691 | C | T | 6.32E-05 | | | | | ORF1ab | nonsynonymous_A1809V |
| 5693 | C | T | 6.32E-05 | | | | | ORF1ab | nonsynonymous_P1810S |
| 5696 | C | T | 6.32E-05 | | | | | ORF1ab | nonsynonymous_P1811S |
| 5698 | T | C | 6.32E-05 | | | | | ORF1ab | synonymous_P1811P |
| 5704 | G | A | 6.32E-05 | A | 6.32E-05 | | | ORF1ab | synonymous_Q1813Q; synonymous_Q1813Q |
| 5714 | A | G | 6.32E-05 | | | | | ORF1ab | nonsynonymous_K1817E |
| 5716 | G | T | 6.32E-05 | | | | | ORF1ab | nonsynonymous_K1817N |
| 5728 | T | C | 6.32E-05 | | | | | ORF1ab | synonymous_F1821F |
| 5742 | A | T | 6.32E-05 | | | | | ORF1ab | nonsynonymous_E1826V |
| 5796 | C | T | 6.32E-05 | | | | | ORF1ab | nonsynonymous_T1844I |
| 5797 | T | C | 6.32E-05 | | | | | ORF1ab | synonymous_T1844T |
| 5821 | A | G | 6.32E-05 | | | | | ORF1ab | synonymous_L1852L |
| 5822 | C | T | 6.32E-05 | | | | | ORF1ab | nonsynonymous_L1853F |
| 5825 | A | G | 6.32E-05 | | | | | ORF1ab | nonsynonymous_T1854A |
| 5832 | C | T | 6.32E-05 | | | | | ORF1ab | nonsynonymous_S1856F |
| 5833 | C | T | 6.32E-05 | | | | | ORF1ab | synonymous_S1856S |
| 5840 | T | C | 6.32E-05 | | | | | ORF1ab | nonsynonymous_Y1859H |
| 5842 | C | T | 6.32E-05 | | | | | ORF1ab | synonymous_Y1859Y |
| 5852 | A | G | 6.32E-05 | | | | | ORF1ab | nonsynonymous_I1863V |
| 5861 | G | A | 6.32E-05 | | | | | ORF1ab | nonsynonymous_V1866I |
| 5874 | A | G | 6.32E-05 | | | | | ORF1ab | nonsynonymous_E1870G |

| | | | | | | | | | |
|---|---|---|---|---|---|---|---|---|---|
| 5877 | A | C | 6.32E-05 | | | | | ORF1ab | nonsynonymous_N1871T |
| 5878 | C | T | 6.32E-05 | | | | | ORF1ab | synonymous_N1871N |
| 5914 | A | G | 6.32E-05 | | | | | ORF1ab | synonymous_K1883K |
| 5920 | T | C | 6.32E-05 | | | | | ORF1ab | synonymous_D1885D |
| 5923 | T | C | 6.32E-05 | | | | | ORF1ab | synonymous_G1886G |
| 5934 | C | T | 6.32E-05 | | | | | ORF1ab | nonsynonymous_T1890I |
| 5937 | A | C | 6.32E-05 | | | | | ORF1ab | nonsynonymous_E1891A |
| 5958 | A | G | 6.32E-05 | | | | | ORF1ab | nonsynonymous_N1898S |
| 5994 | A | T | 6.32E-05 | | | | | ORF1ab | nonsynonymous_Q1910L |
| 5995 | A | G | 6.32E-05 | | | | | ORF1ab | synonymous_Q1910Q |
| 5997 | C | G | 6.32E-05 | | | | | ORF1ab | nonsynonymous_P1911R |
| 6036 | G | A | 6.32E-05 | | | | | ORF1ab | nonsynonymous_S1924N |
| 6051 | A | C | 6.32E-05 | | | | | ORF1ab | nonsynonymous_K1929T |
| 6058 | A | G | 6.32E-05 | | | | | ORF1ab | synonymous_V1931V |
| 6079 | T | C | 6.32E-05 | | | | | ORF1ab | synonymous_A1938A |
| 6080 | G | T | 6.32E-05 | | | | | ORF1ab | nonsynonymous_D1939Y |
| 6096 | T | C | 6.32E-05 | | | | | ORF1ab | nonsynonymous_L1944S |
| 6101 | G | A | 6.32E-05 | | | | | ORF1ab | nonsynonymous_G1946S |
| 6128 | C | A | 6.32E-05 | A | 6.32E-05 | | | ORF1ab | nonsynonymous_L1955I; nonsynonymous_L1955I |
| 6173 | A | G | 6.32E-05 | | | | | ORF1ab | nonsynonymous_I1970V |
| 6175 | T | C | 6.32E-05 | | | | | ORF1ab | synonymous_I1970I |
| 6183 | A | G | 6.32E-05 | | | | | ORF1ab | nonsynonymous_K1973R |
| 6187 | C | T | 6.32E-05 | | | | | ORF1ab | synonymous_H1974H |
| 6197 | T | C | 6.32E-05 | | | | | ORF1ab | nonsynonymous_S1978P |
| 6236 | G | A | 6.32E-05 | | | | | ORF1ab | nonsynonymous_V1991I |
| 625 | G | T | 6.32E-05 | | | | | ORF1ab | nonsynonymous_K120N |
| 626 | G | A | 6.32E-05 | A | 6.32E-05 | | | ORF1ab | nonsynonymous_V121I; nonsynonymous_V121I |

| | | | | | | | | | |
|---|---|---|---|---|---|---|---|---|---|
| 629 | C | A | 6.32E-05 | A | 6.32E-05 | | | ORF1ab | nonsynonymous_L122I; nonsynonymous_L122I |
| 645 | G | A | 6.32E-05 | | | | | ORF1ab | nonsynonymous_G127D |
| 6500 | C | T | 6.32E-05 | | | | | ORF1ab | nonsynonymous_P2079S |
| 6502 | A | G | 6.32E-05 | | | | | ORF1ab | synonymous_P2079P |
| 654 | G | A | 6.32E-05 | | | | | ORF1ab | nonsynonymous_G130E |
| 662 | G | T | 6.32E-05 | | | | | ORF1ab | nonsynonymous_G133C |
| 664 | C | T | 6.32E-05 | | | | | ORF1ab | synonymous_G133G |
| 676 | C | T | 6.32E-05 | | | | | ORF1ab | synonymous_G137G |
| 681 | A | G | 6.32E-05 | | | | | ORF1ab | nonsynonymous_D139G |
| 709 | G | A | 6.32E-05 | | | | | ORF1ab | synonymous_E148E |
| 710 | C | T | 6.32E-05 | | | | | ORF1ab | nonsynonymous_L149F |
| 7160 | G | C | 6.32E-05 | | | | | ORF1ab | nonsynonymous_D2299H |
| 7168 | T | C | 6.32E-05 | | | | | ORF1ab | synonymous_Y2301Y |
| 7169 | C | T | 6.32E-05 | | | | | ORF1ab | nonsynonymous_P2302S |
| 7178 | G | A | 6.32E-05 | | | | | ORF1ab | nonsynonymous_E2305K |
| 7192 | T | C | 6.32E-05 | | | | | ORF1ab | synonymous_I2309I |
| 7195 | C | T | 6.32E-05 | | | | | ORF1ab | synonymous_T2310T |
| 7200 | C | T | 6.32E-05 | | | | | ORF1ab | nonsynonymous_S2312L |
| 720 | A | G | 6.32E-05 | | | | | ORF1ab | nonsynonymous_D152G |
| 7239 | C | A | 6.32E-05 | A | 6.32E-05 | | | ORF1ab | nonsynonymous_A2325E; nonsynonymous_A2325E |
| 7240 | A | G | 6.32E-05 | | | | | ORF1ab | synonymous_A2325A |
| 7367 | A | C | 6.32E-05 | | | | | ORF1ab | nonsynonymous_I2368L |
| 7389 | C | T | 6.32E-05 | | | | | ORF1ab | nonsynonymous_A2375V |
| 7397 | T | C | 6.32E-05 | | | | | ORF1ab | nonsynonymous_S2378P |
| 7401 | C | T | 6.32E-05 | | | | | ORF1ab | nonsynonymous_A2379V |
| 740 | G | A | 6.32E-05 | | | | | ORF1ab | nonsynonymous_E159K |
| 7435 | T | C | 6.32E-05 | | | | | ORF1ab | synonymous_F2390F |
| 745 | C | T | 6.32E-05 | | | | | ORF1ab | synonymous_N160N |

| | | | | | | | | | |
|---|---|---|---|---|---|---|---|---|---|
| 7468 | A | G | 6.32E-05 | | | | | ORF1ab | synonymous_V2401V |
| 7483 | A | G | 6.32E-05 | | | | | ORF1ab | synonymous_S2406S |
| 7488 | C | T | 6.32E-05 | | | | | ORF1ab | nonsynonymous_T2408I |
| 749 | A | T | 6.32E-05 | | | | | ORF1ab | nonsynonymous_N162Y |
| 7513 | T | C | 6.32E-05 | | | | | ORF1ab | synonymous_N2416N |
| 7518 | C | T | 6.32E-05 | | | | | ORF1ab | nonsynonymous_A2418V |
| 7521 | C | T | 6.32E-05 | | | | | ORF1ab | nonsynonymous_T2419I |
| 7561 | G | A | 6.32E-05 | | | | | ORF1ab | synonymous_R2432R |
| 7562 | T | C | 6.32E-05 | | | | | ORF1ab | nonsynonymous_S2433P |
| 7582 | T | C | 6.32E-05 | | | | | ORF1ab | synonymous_N2439N |
| 7603 | A | C | 6.32E-05 | | | | | ORF1ab | nonsynonymous_K2446N |
| 7606 | A | T | 6.32E-05 | | | | | ORF1ab | synonymous_L2447L |
| 7609 | C | T | 6.32E-05 | | | | | ORF1ab | synonymous_H2448H |
| 7630 | T | G | 6.32E-05 | | | | | ORF1ab | nonsynonymous_C2455W |
| 7640 | T | G | 6.32E-05 | | | | | ORF1ab | nonsynonymous_C2459G |
| 7642 | T | G | 6.32E-05 | | | | | ORF1ab | nonsynonymous_C2459W |
| 7646 | G | C | 6.32E-05 | | | | | ORF1ab | nonsynonymous_G2461R |
| 7647 | G | T | 6.32E-05 | | | | | ORF1ab | nonsynonymous_G2461V |
| 7648 | T | A | 6.32E-05 | | | | | ORF1ab | synonymous_G2461G |
| 7666 | T | A | 6.32E-05 | | | | | ORF1ab | nonsynonymous_D2467E |
| 7678 | A | C | 6.32E-05 | | | | | ORF1ab | nonsynonymous_R2471S |
| 7684 | G | A | 6.32E-05 | | | | | ORF1ab | synonymous_L2473L |
| 7777 | C | T | 6.32E-05 | | | | | ORF1ab | synonymous_Y2504Y |
| 7789 | T | C | 6.32E-05 | | | | | ORF1ab | synonymous_A2508A |
| 778 | T | C | 6.32E-05 | | | | | ORF1ab | synonymous_R171R |
| 7790 | G | T | 6.32E-05 | | | | | ORF1ab | nonsynonymous_G2509C |
| 779 | G | C | 6.32E-05 | | | | | ORF1ab | nonsynonymous_E172Q |
| 784 | C | T | 6.32E-05 | | | | | ORF1ab | synonymous_L173L |
| 806 | G | A | 6.32E-05 | | | | | ORF1ab | nonsynonymous_A181T |
| 8091 | T | A | 6.32E-05 | | | | | ORF1ab | nonsynonymous_L2609H |

| | | | | | | | | | |
|---|---|---|---|---|---|---|---|---|---|
| 8106 | C | T | 6.32E-05 | | | | | ORF1ab | nonsynonymous_A2614V |
| 810 | A | G | 6.32E-05 | | | | | ORF1ab | nonsynonymous_Y182C |
| 8115 | A | G | 6.32E-05 | | | | | ORF1ab | nonsynonymous_E2617G |
| 813 | C | A | 6.32E-05 | | | | | ORF1ab | nonsynonymous_T183N |
| 8171 | G | A | 6.32E-05 | | | | | ORF1ab | nonsynonymous_A2636T |
| 8185 | G | A | 6.32E-05 | | | | | ORF1ab | synonymous_G2640G |
| 8200 | T | C | 6.32E-05 | | | | | ORF1ab | synonymous_D2645D |
| 8201 | G | A | 6.32E-05 | | | | | ORF1ab | nonsynonymous_V2646I |
| 8214 | A | G | 6.32E-05 | | | | | ORF1ab | nonsynonymous_D2650G |
| 8264 | G | T | 6.32E-05 | | | | | ORF1ab | nonsynonymous_G2667C |
| 8281 | C | T | 6.32E-05 | | | | | ORF1ab | synonymous_N2672N |
| 8287 | G | T | 6.32E-05 | | | | | ORF1ab | nonsynonymous_M2674I |
| 8323 | T | C | 6.32E-05 | | | | | ORF1ab | synonymous_R2686R |
| 8327 | C | T | 6.32E-05 | | | | | ORF1ab | nonsynonymous_L2688F |
| 8329 | T | A | 6.32E-05 | A | 6.32E-05 | | | ORF1ab | synonymous_L2688L; synonymous_L2688L |
| 8336 | T | A | 6.32E-05 | | | | | ORF1ab | nonsynonymous_C2691S |
| 834 | T | G | 6.32E-05 | | | | | ORF1ab | nonsynonymous_F190C |
| 8380 | A | G | 6.32E-05 | | | | | ORF1ab | synonymous_K2705K |
| 8388 | A | T | 6.32E-05 | T | 6.32E-05 | | | ORF1ab | nonsynonymous_N2708I; nonsynonymous_N2708I |
| 8395 | T | C | 6.32E-05 | | | | | ORF1ab | synonymous_A2710A |
| 8443 | A | G | 6.32E-05 | | | | | ORF1ab | synonymous_R2726R |
| 8455 | T | C | 6.32E-05 | | | | | ORF1ab | synonymous_R2730R |
| 8473 | T | C | 6.32E-05 | | | | | ORF1ab | synonymous_N2736N |
| 8480 | C | T | 6.32E-05 | | | | | ORF1ab | nonsynonymous_P2739S |
| 851 | T | C | 6.32E-05 | | | | | ORF1ab | nonsynonymous_Y196H |
| 8529 | C | T | 6.32E-05 | | | | | ORF1ab | nonsynonymous_T2755I |
| 8557 | A | G | 6.32E-05 | | | | | ORF1ab | synonymous_K2764K |
| 8561 | G | A | 6.32E-05 | | | | | ORF1ab | nonsynonymous_V2766I |

| | | | | | | | | | |
|---|---|---|---|---|---|---|---|---|---|
| 8565 | A | G | 6.32E-05 | | | | | ORF1ab | nonsynonymous_N2767S |
| 8566 | T | C | 6.32E-05 | | | | | ORF1ab | synonymous_N2767N |
| 857 | C | T | 6.32E-05 | | | | | ORF1ab | nonsynonymous_L198F |
| 8595 | C | T | 6.32E-05 | | | | | ORF1ab | nonsynonymous_T2777I |
| 8603 | T | C | 6.32E-05 | | | | | ORF1ab | nonsynonymous_F2780L |
| 860 | G | A | 6.32E-05 | | | | | ORF1ab | nonsynonymous_E199K |
| 8616 | C | T | 6.32E-05 | | | | | ORF1ab | nonsynonymous_A2784V |
| 865 | C | T | 6.32E-05 | | | | | ORF1ab | synonymous_C200C |
| 871 | A | G | 6.32E-05 | | | | | ORF1ab | synonymous_K202K |
| 872 | G | T | 6.32E-05 | T | 6.32E-05 | | | ORF1ab | nonsynonymous_D203Y; nonsynonymous_D203Y |
| 873 | A | G | 6.32E-05 | | | | | ORF1ab | nonsynonymous_D203G |
| 874 | C | A | 6.32E-05 | | | | | ORF1ab | nonsynonymous_D203E |
| 878 | C | A | 6.32E-05 | | | | | ORF1ab | nonsynonymous_L205I |
| 889 | T | C | 6.32E-05 | | | | | ORF1ab | synonymous_A208A |
| 892 | T | C | 6.32E-05 | | | | | ORF1ab | synonymous_G209G |
| 8936 | G | A | 6.32E-05 | | | | | ORF1ab | nonsynonymous_A2891T |
| 8937 | C | T | 6.32E-05 | | | | | ORF1ab | nonsynonymous_A2891V |
| 8972 | A | G | 6.32E-05 | | | | | ORF1ab | nonsynonymous_I2903V |
| 897 | C | T | 6.32E-05 | | | | | ORF1ab | nonsynonymous_A211V |
| 8980 | C | T | 6.32E-05 | | | | | ORF1ab | synonymous_Y2905Y |
| 8981 | A | G | 6.32E-05 | | | | | ORF1ab | nonsynonymous_T2906A |
| 8982 | C | T | 6.32E-05 | | | | | ORF1ab | nonsynonymous_T2906I |
| 8983 | T | A | 6.32E-05 | | | | | ORF1ab | synonymous_T2906T |
| 8986 | C | T | 6.32E-05 | | | | | ORF1ab | synonymous_D2907D |
| 8998 | A | G | 6.32E-05 | | | | | ORF1ab | synonymous_S2911S |
| 9034 | A | G | 6.32E-05 | | | | | ORF1ab | synonymous_K2923K |
| 9037 | T | C | 6.32E-05 | | | | | ORF1ab | synonymous_D2924D |
| 9046 | T | C | 6.32E-05 | | | | | ORF1ab | synonymous_G2927G |
| 9049 | G | T | 6.32E-05 | | | | | ORF1ab | nonsynonymous_K2928N |

| | | | | | | | | | |
|---|---|---|---|---|---|---|---|---|---|
| 9072 | C | T | 6.32E-05 | | | | | ORF1ab | nonsynonymous_T2936I |
| 9092 | G | A | 6.32E-05 | | | | | ORF1ab | nonsynonymous_V2943I |
| 9096 | C | T | 6.32E-05 | | | | | ORF1ab | nonsynonymous_A2944V |
| 9106 | T | C | 6.32E-05 | | | | | ORF1ab | synonymous_S2947S |
| 910 | G | T | 6.32E-05 | | | | | ORF1ab | nonsynonymous_L215F |
| 9127 | T | C | 6.32E-05 | | | | | ORF1ab | synonymous_Y2954Y |
| 9131 | C | T | 6.32E-05 | | | | | ORF1ab | nonsynonymous_L2956F |
| 9142 | C | A | 6.32E-05 | | | | | ORF1ab | synonymous_G2959G |
| 914 | G | A | 6.32E-05 | | | | | ORF1ab | nonsynonymous_E217K |
| 9157 | T | C | 6.32E-05 | | | | | ORF1ab | synonymous_F2964F |
| 9159 | C | T | 6.32E-05 | | | | | ORF1ab | nonsynonymous_P2965L |
| 9171 | T | A | 6.32E-05 | | | | | ORF1ab | nonsynonymous_L2969H |
| 9178 | T | G | 6.32E-05 | | | | | ORF1ab | synonymous_G2971G |
| 9182 | G | T | 6.32E-05 | | | | | ORF1ab | nonsynonymous_V2973F |
| 9191 | G | A | 6.32E-05 | | | | | ORF1ab | nonsynonymous_V2976I |
| 9203 | G | A | 6.32E-05 | | | | | ORF1ab | nonsynonymous_D2980N |
| 9212 | T | A | 6.32E-05 | | | | | ORF1ab | nonsynonymous_Y2983N |
| 9214 | C | T | 6.32E-05 | | | | | ORF1ab | synonymous_Y2983Y |
| 9218 | A | T | 6.32E-05 | | | | | ORF1ab | nonsynonymous_R2985W |
| 9220 | G | A | 6.32E-05 | | | | | ORF1ab | synonymous_R2985R |
| 9224 | G | T | 6.32E-05 | | | | | ORF1ab | nonsynonymous_G2987C |
| 9226 | C | T | 6.32E-05 | | | | | ORF1ab | synonymous_G2987G |
| 9231 | G | C | 6.32E-05 | | | | | ORF1ab | nonsynonymous_C2989S |
| 9238 | A | G | 6.32E-05 | | | | | ORF1ab | synonymous_R2991R |
| 9244 | A | G | 6.32E-05 | | | | | ORF1ab | synonymous_E2993E |
| 9245 | G | C | 6.32E-05 | | | | | ORF1ab | nonsynonymous_A2994P |
| 929 | A | G | 6.32E-05 | | | | | ORF1ab | nonsynonymous_I222V |
| 932 | G | A | 6.32E-05 | | | | | ORF1ab | nonsynonymous_D223N |
| 937 | T | C | 6.32E-05 | | | | | ORF1ab | synonymous_T224T |
| 939 | A | G | 6.32E-05 | | | | | ORF1ab | nonsynonymous_K225R |

| | | | | | | | | | |
|---|---|---|---|---|---|---|---|---|---|
| 943 | G | A | 6.32E-05 | | | | | ORF1ab | synonymous_R226R |
| 944 | G | A | 6.32E-05 | | | | | ORF1ab | nonsynonymous_G227S |
| 9508 | T | C | 6.32E-05 | | | | | ORF1ab | synonymous_N3081N |
| 9511 | T | C | 6.32E-05 | | | | | ORF1ab | synonymous_T3082T |
| 9515 | C | T | 6.32E-05 | | | | | ORF1ab | synonymous_L3084L |
| 9521 | C | T | 6.32E-05 | | | | | ORF1ab | nonsynonymous_L3086F |
| 9562 | A | G | 6.32E-05 | | | | | ORF1ab | synonymous_S3099S |
| 9572 | G | A | 6.32E-05 | | | | | ORF1ab | nonsynonymous_G3103S |
| 9574 | T | C | 6.32E-05 | | | | | ORF1ab | synonymous_G3103G |
| 9587 | A | G | 6.32E-05 | | | | | ORF1ab | nonsynonymous_I3108V |
| 958 | C | T | 6.32E-05 | | | | | ORF1ab | synonymous_C231C |
| 959 | C | T | 6.32E-05 | | | | | ORF1ab | nonsynonymous_R232C |
| 9601 | G | A | 6.32E-05 | | | | | ORF1ab | synonymous_L3112L |
| 9603 | C | T | 6.32E-05 | | | | | ORF1ab | nonsynonymous_T3113I |
| 9624 | T | A | 6.32E-05 | | | | | ORF1ab | nonsynonymous_V3120D |
| 964 | A | G | 6.32E-05 | | | | | ORF1ab | synonymous_E233E |
| 9692 | G | A | 6.32E-05 | | | | | ORF1ab | nonsynonymous_A3143T |
| 9700 | C | T | 6.32E-05 | | | | | ORF1ab | synonymous_I3145I |
| 9701 | A | T | 6.32E-05 | | | | | ORF1ab | nonsynonymous_I3146F |
| 9704 | T | C | 6.32E-05 | | | | | ORF1ab | nonsynonymous_C3147R |
| 9705 | G | T | 6.32E-05 | | | | | ORF1ab | nonsynonymous_C3147F |
| 9707 | A | G | 6.32E-05 | | | | | ORF1ab | nonsynonymous_I3148V |
| 9708 | T | C | 6.32E-05 | | | | | ORF1ab | nonsynonymous_I3148T |
| 9710 | T | G | 6.32E-05 | | | | | ORF1ab | nonsynonymous_S3149A |
| 9711 | C | T | 6.32E-05 | | | | | ORF1ab | nonsynonymous_S3149F |
| 9738 | G | T | 6.32E-05 | | | | | ORF1ab | nonsynonymous_S3158I |
| 973 | T | C | 6.32E-05 | | | | | ORF1ab | synonymous_H236H |
| 9741 | A | C | 6.32E-05 | | | | | ORF1ab | nonsynonymous_N3159T |
| 9744 | A | G | 6.32E-05 | | | | | ORF1ab | nonsynonymous_Y3160C |
| 9746 | C | T | 6.32E-05 | | | | | ORF1ab | synonymous_L3161L |

| | | | | | | | | | |
|---|---|---|---|---|---|---|---|---|---|
| 9760 | A | G | 6.32E-05 | | | | | ORF1ab | synonymous_V3165V |
| 9763 | C | T | 6.32E-05 | | | | | ORF1ab | synonymous_V3166V |
| 9791 | G | A | 6.32E-05 | | | | | ORF1ab | nonsynonymous_E3176K |
| 9853 | T | C | 6.32E-05 | | | | | ORF1ab | synonymous_D3196D |
| 986 | T | C | 6.32E-05 | | | | | ORF1ab | nonsynonymous_Y241H |
| 9890 | G | T | 6.32E-05 | | | | | ORF1ab | nonsynonymous_A3209S |
| 9893 | C | T | 6.32E-05 | | | | | ORF1ab | nonsynonymous_L3210F |
| 990 | C | T | 6.32E-05 | | | | | ORF1ab | nonsynonymous_T242M |
| 991 | G | A | 6.32E-05 | | | | | ORF1ab | synonymous_T242T |
| 9925 | A | T | 6.32E-05 | | | | | ORF1ab | synonymous_A3220A |
| 9933 | C | T | 6.32E-05 | | | | | ORF1ab | nonsynonymous_T3223I |
| 9964 | T | C | 6.32E-05 | | | | | ORF1ab | synonymous_H3233H |
| 998 | T | G | 6.32E-05 | | | | | ORF1ab | nonsynonymous_S245A |
| 9997 | A | G | 6.32E-05 | | | | | ORF1ab | synonymous_S3244S |
