## Extended Data Table for "Worldwide tracing of mutations and the evolutionary dynamics of SARS-CoV-2": Extended Data Table 2.pdf

**Extended Data Table 2. Acknowledgement of the sharing of SARS-CoV-2 raw genome sequences in NCBI SRA database used in this study.**

| SRA Number | Location | Number of Samples | Collection Date | Submitting Lab | Authors |
|---|---|---|---|---|---|
| PRJNA613958 | Australia: Victoria, Northern Territory | 1160 | 1/24/2020-4/17/2020 | The Peter Doherty Institute for Infection and Immunity | Mark Schultz |
| PRJEB37886 | United Kingdom: Wales, Scotland, England | 400 | 3/17/2020-4/12/2020 | COVID-19 Genomics UK (COG-UK) consortium | |
| PRJNA614995 | USA: Utah | 281 | 3/3/2020-4/15/2020 | Utah Public Health Laboratory | Erin Young |
| PRJNA633948 | Australia: NSW | 204 | 1/24/2020-3/26/2020 | Institute of Clinical Pathology and Medical | Mailie Gall |
| PRJNA627662 | China: Shanghai | 112 | 1/25/2020-2/15/2020 | Ruijin Hospital Affiliated to Shanghai Jiao Tong University School of Medicine | Yun Tan |
| PRJNA625669 | India: Gujarat | 109 | 4/5/2020-5/11/2020 | Gujarat Biotechnology Research Centre | Chaitanya Joshi |
| PRJNA632475 | USA: Hamilton, MT | 47 | 3/5/2020-3/10/2020 | National Institute of Allergy and Infectious Diseases | Craig Martens |
| PRJNA614546 | USA: Los Angeles | 13 | 3/17/2020-4/5/2020 | Paragon Genomics | Vidushi Kapoor |
| PRJNA631042 | USA: Minnesota | 12 | 3/26/2020-4/5/2020 | University of Minnesota | John Garbe |
| PRJNA610428 | USA: WA | 6 | 2/27/2020-3/1/2020 | University of Washington | Pavitra |
| PRJNA627977 | USA: Washington | 2 | 2/7/2020-2/14/2020 | UW-Madison | Shelby O'Connor |
| PRJNA627354 | USA: New York | 1 | 4/2/2020-4/2/2020 | USDA National Veterinary Services Laboratories | Mary Killian |
| PRJNA612578 | USA: CA, San Diego | 1 | 3/11/2020 | The Scripps Research Institute | Karthik |
| PRJNA622817 | USA: Washington | 1 | 1/19/2020 | CDC Pathogen Discovery Team | |
| PRJNA631042 | USA | 1 | 1/19/2020 | University of Minnesota | John Garbe |
| PRJNA231221 | USA | 1 | 1/19/2020 | FDA_ARGOS | |
