## Extended Data Table for "Worldwide tracing of mutations and the evolutionary dynamics of SARS-CoV-2": Extended Data Table 3.pdf

**Extended Data Table 3. Annotations of the SNPs identified from the raw reads generated from next generation sequencing of SARS-CoV-2 samples.**

| Genome Position | Ref | Alt1 | Alt2 | Annotation |
|---|---|---|---|---|
| 61 | G | T | | five_prime_UTR |
| 66 | C | T | | five_prime_UTR |
| 71 | C | T | | five_prime_UTR |
| 78 | T | A | | five_prime_UTR |
| 84 | C | T | | five_prime_UTR |
| 98 | C | T | | five_prime_UTR |
| 100 | C | T | | five_prime_UTR |
| 103 | C | T | | five_prime_UTR |
| 106 | C | T | | five_prime_UTR |
| 110 | C | T | | five_prime_UTR |
| 139 | A | T | | five_prime_UTR |
| 140 | C | T | | five_prime_UTR |
| 149 | G | A | T | five_prime_UTR |
| 157 | C | T | | five_prime_UTR |
| 165 | G | T | | five_prime_UTR |
| 167 | G | T | | five_prime_UTR |
| 169 | A | G | | five_prime_UTR |
| 174 | G | C | | five_prime_UTR |
| 186 | C | A | | five_prime_UTR |
| 187 | A | G | | five_prime_UTR |
| 188 | G | T | | five_prime_UTR |
| 189 | G | A | | five_prime_UTR |
| 191 | T | C | | five_prime_UTR |
| 201 | T | G | | five_prime_UTR |
| 202 | T | C | | five_prime_UTR |
| 203 | C | T | | five_prime_UTR |
| 204 | G | T | | five_prime_UTR |
| 207 | C | T | | five_prime_UTR |

| | | | | | |
|---|---|---|---|---|---|
| 210 | G | T | | five_prime_UTR | |
| 211 | T | C | | five_prime_UTR | |
| 217 | C | T | | five_prime_UTR | |
| 218 | C | T | | five_prime_UTR | |
| 220 | A | T | | five_prime_UTR | |
| 221 | T | C | | five_prime_UTR | |
| 222 | C | T | | five_prime_UTR | |
| 228 | C | T | | five_prime_UTR | |
| 233 | C | T | | five_prime_UTR | |
| 238 | T | G | | five_prime_UTR | |
| 241 | C | T | | five_prime_UTR | |
| 245 | C | T | | five_prime_UTR | |
| 246 | G | T | | five_prime_UTR | |
| 247 | G | T | | five_prime_UTR | |
| 252 | G | T | | five_prime_UTR | |
| 255 | C | T | | five_prime_UTR | |
| 256 | G | A | | five_prime_UTR | |
| 259 | A | G | | five_prime_UTR | |
| 275 | C | T | | ORF1ab | nonsynonymous_L4F |
| 285 | G | A | T | ORF1ab | nonsynonymous_G7D; nonsynonymous_G7V |
| 289 | C | T | | ORF1ab | synonymous_F8F |
| 300 | C | T | | ORF1ab | nonsynonymous_T12I |
| 301 | A | C | | ORF1ab | synonymous_T12T |
| 302 | C | T | | ORF1ab | nonsynonymous_H13Y |
| 304 | C | T | | ORF1ab | synonymous_H13H |
| 311 | C | T | | ORF1ab | nonsynonymous_L16F |
| 313 | C | T | | ORF1ab | synonymous_L16L |
| 315 | G | T | | ORF1ab | nonsynonymous_S17I |
| 319 | G | A | C | ORF1ab | synonymous_L18L; nonsynonymous_L18F |

| | | | | | |
|---|---|---|---|---|---|
| 320 | C | T | | ORF1ab | nonsynonymous_P19S |
| 333 | T | C | | ORF1ab | nonsynonymous_V23A |
| 335 | C | T | | ORF1ab | nonsynonymous_R24C |
| 336 | G | T | | ORF1ab | nonsynonymous_R24L |
| 337 | C | T | | ORF1ab | synonymous_R24R |
| 344 | C | T | | ORF1ab | nonsynonymous_L27F |
| 350 | C | T | | ORF1ab | nonsynonymous_R29C |
| 351 | G | A | | ORF1ab | nonsynonymous_R29H |
| 368 | G | T | | ORF1ab | nonsynonymous_V35L |
| 370 | G | C | T | ORF1ab | synonymous_V35V;<br>synonymous_V35V |
| 374 | G | A | | ORF1ab | nonsynonymous_E37K |
| 377 | G | T | | ORF1ab | nonsynonymous_V38F |
| 379 | C | A | T | ORF1ab | synonymous_V38V;<br>synonymous_V38V |
| 380 | T | G | | ORF1ab | nonsynonymous_L39V |
| 382 | A | C | | ORF1ab | nonsynonymous_L39F |
| 388 | G | A | T | ORF1ab | synonymous_E41E;<br>nonsynonymous_E41D |
| 392 | C | T | | ORF1ab | nonsynonymous_R43C |
| 403 | T | A | | ORF1ab | synonymous_L46L |
| 409 | T | A | G | ORF1ab | nonsynonymous_D48E;<br>nonsynonymous_D48E |
| 422 | T | C | | ORF1ab | synonymous_L53L |
| 427 | A | T | | ORF1ab | synonymous_V54V |
| 428 | G | T | | ORF1ab | stopgain_E55X |
| 439 | A | T | | ORF1ab | nonsynonymous_K58N |
| 443 | G | A | | ORF1ab | nonsynonymous_V60I |
| 447 | T | C | | ORF1ab | nonsynonymous_L61S |
| 449 | C | T | | ORF1ab | nonsynonymous_P62S |
| 461 | C | T | | ORF1ab | stopgain_Q66X |

| | | | | | |
|---|---|---|---|---|---|
| 466 | C | T | | ORF1ab | synonymous_P67P |
| 467 | T | C | | ORF1ab | nonsynonymous_Y68H |
| 472 | G | T | | ORF1ab | synonymous_V69V |
| 478 | C | A | | ORF1ab | synonymous_I71I |
| 482 | C | T | | ORF1ab | nonsynonymous_R73C |
| 487 | G | T | | ORF1ab | synonymous_S74S |
| 490 | T | A | | ORF1ab | nonsynonymous_D75E |
| 492 | C | T | | ORF1ab | nonsynonymous_A76V |
| 498 | C | T | | ORF1ab | nonsynonymous_T78I |
| 500 | G | C | T | ORF1ab | nonsynonymous_A79P; nonsynonymous_A79S |
| 503 | C | T | | ORF1ab | nonsynonymous_P80S |
| 509 | G | T | | ORF1ab | nonsynonymous_G82C |
| 512 | C | T | | ORF1ab | nonsynonymous_H83Y |
| 514 | T | C | | ORF1ab | synonymous_H83H |
| 525 | A | T | | ORF1ab | nonsynonymous_E87V |
| 527 | C | T | | ORF1ab | synonymous_L88L |
| 530 | G | A | | ORF1ab | nonsynonymous_V89I |
| 533 | G | T | | ORF1ab | nonsynonymous_A90S |
| 536 | G | T | | ORF1ab | stopgain_E91X |
| 539 | C | T | | ORF1ab | nonsynonymous_L92F |
| 541 | C | T | | ORF1ab | synonymous_L92L |
| 546 | G | T | | ORF1ab | nonsynonymous_G94V |
| 547 | C | T | | ORF1ab | synonymous_G94G |
| 553 | G | T | | ORF1ab | nonsynonymous_Q96H |
| 558 | G | T | | ORF1ab | nonsynonymous_G98V |
| 564 | G | T | | ORF1ab | nonsynonymous_S100I |
| 567 | G | T | | ORF1ab | nonsynonymous_G101V |
| 569 | G | T | | ORF1ab | stopgain_E102X |
| 573 | C | G | | ORF1ab | nonsynonymous_T103R |
| 581 | G | T | | ORF1ab | nonsynonymous_V106F |

| | | | | | |
|---|---|---|---|---|---|
| 583 | C | T | | ORF1ab | synonymous_V106V |
| 591 | C | T | | ORF1ab | nonsynonymous_P109L |
| 592 | T | C | | ORF1ab | synonymous_P109P |
| 593 | C | T | | ORF1ab | nonsynonymous_H110Y |
| 596 | G | T | | ORF1ab | nonsynonymous_V111L |
| 601 | C | T | | ORF1ab | synonymous_G112G |
| 606 | T | C | | ORF1ab | nonsynonymous_I114T |
| 613 | G | A | | ORF1ab | synonymous_V116V |
| 622 | C | T | | ORF1ab | synonymous_R119R |
| 626 | G | T | | ORF1ab | nonsynonymous_V121F |
| 632 | C | T | | ORF1ab | nonsynonymous_L123F |
| 635 | C | T | | ORF1ab | nonsynonymous_R124C |
| 643 | C | T | | ORF1ab | synonymous_N126N |
| 645 | G | T | | ORF1ab | nonsynonymous_G127V |
| 648 | A | G | | ORF1ab | nonsynonymous_N128S |
| 657 | C | T | | ORF1ab | nonsynonymous_A131V |
| 660 | G | T | | ORF1ab | nonsynonymous_G132V |
| 663 | G | T | | ORF1ab | nonsynonymous_G133V |
| 664 | C | T | | ORF1ab | synonymous_G133G |
| 668 | A | T | | ORF1ab | nonsynonymous_S135C |
| 676 | C | T | | ORF1ab | synonymous_G137G |
| 678 | C | T | | ORF1ab | nonsynonymous_A138V |
| 679 | C | T | | ORF1ab | synonymous_A138A |
| 680 | G | A | T | ORF1ab | nonsynonymous_D139N; nonsynonymous_D139Y |
| 683 | C | T | | ORF1ab | synonymous_L140L |
| 684 | T | A | | ORF1ab | nonsynonymous_L140Q |
| 700 | A | T | | ORF1ab | nonsynonymous_L145F |
| 701 | G | T | | ORF1ab | nonsynonymous_G146C |
| 710 | C | T | | ORF1ab | nonsynonymous_L149F |
| 717 | C | T | | ORF1ab | nonsynonymous_T151I |

| | | | | | |
|---|---|---|---|---|---|
| 728 | G | T | | ORF1ab | stopgain_E155X |
| 740 | G | A | | ORF1ab | nonsynonymous_E159K |
| 745 | C | T | | ORF1ab | synonymous_N160N |
| 746 | T | C | G | ORF1ab | nonsynonymous_W161R;<br>nonsynonymous_W161G |
| 760 | T | C | | ORF1ab | synonymous_H165H |
| 761 | A | G | | ORF1ab | nonsynonymous_S166G |
| 771 | T | C | | ORF1ab | nonsynonymous_V169A |
| 774 | C | T | | ORF1ab | nonsynonymous_T170I |
| 775 | C | T | | ORF1ab | synonymous_T170T |
| 777 | G | A | | ORF1ab | nonsynonymous_R171H |
| 779 | G | T | | ORF1ab | stopgain_E172X |
| 782 | C | T | | ORF1ab | nonsynonymous_L173F |
| 786 | T | G | | ORF1ab | nonsynonymous_M174R |
| 787 | G | T | | ORF1ab | nonsynonymous_M174I |
| 789 | G | T | | ORF1ab | nonsynonymous_R175L |
| 791 | G | T | | ORF1ab | stopgain_E176X |
| 802 | A | C | | ORF1ab | synonymous_G179G |
| 805 | G | T | | ORF1ab | synonymous_G180G |
| 809 | T | C | | ORF1ab | nonsynonymous_Y182H |
| 815 | C | T | | ORF1ab | nonsynonymous_R184C |
| 816 | G | T | | ORF1ab | nonsynonymous_R184L |
| 817 | C | T | | ORF1ab | synonymous_R184R |
| 823 | C | T | | ORF1ab | synonymous_V186V |
| 832 | C | T | | ORF1ab | synonymous_N189N |
| 833 | T | C | | ORF1ab | nonsynonymous_F190L |
| 835 | C | T | | ORF1ab | synonymous_F190F |
| 837 | G | T | | ORF1ab | nonsynonymous_C191F |
| 839 | G | T | | ORF1ab | nonsynonymous_G192C |
| 840 | G | A | | ORF1ab | nonsynonymous_G192D |
| 841 | C | T | | ORF1ab | synonymous_G192G |

| | | | | | |
|---|---|---|---|---|---|
| 843 | C | A | T | ORF1ab | nonsynonymous_P193H; nonsynonymous_P193L |
| 846 | A | T | | ORF1ab | nonsynonymous_D194V |
| 847 | T | C | | ORF1ab | synonymous_D194D |
| 848 | G | T | | ORF1ab | nonsynonymous_G195C |
| 854 | C | T | | ORF1ab | nonsynonymous_P197S |
| 856 | T | C | | ORF1ab | synonymous_P197P |
| 859 | T | G | | ORF1ab | synonymous_L198L |
| 861 | A | C | | ORF1ab | nonsynonymous_E199A |
| 862 | G | T | | ORF1ab | nonsynonymous_E199D |
| 871 | A | G | | ORF1ab | synonymous_K202K |
| 875 | C | T | | ORF1ab | nonsynonymous_L204F |
| 878 | C | T | | ORF1ab | synonymous_L205L |
| 884 | C | T | | ORF1ab | nonsynonymous_R207C |
| 885 | G | T | | ORF1ab | nonsynonymous_R207L |
| 894 | A | C | | ORF1ab | nonsynonymous_K210T |
| 895 | A | G | | ORF1ab | synonymous_K210K |
| 896 | G | A | | ORF1ab | nonsynonymous_A211T |
| 897 | C | T | | ORF1ab | nonsynonymous_A211V |
| 900 | C | T | | ORF1ab | nonsynonymous_S212L |
| 903 | G | A | | ORF1ab | nonsynonymous_C213Y |
| 906 | C | T | | ORF1ab | nonsynonymous_T214I |
| 910 | G | T | | ORF1ab | nonsynonymous_L215F |
| 913 | C | T | | ORF1ab | synonymous_S216S |
| 914 | G | T | | ORF1ab | stopgain_E217X |
| 920 | C | T | | ORF1ab | synonymous_L219L |
| 924 | A | C | | ORF1ab | nonsynonymous_D220A |
| 925 | C | T | | ORF1ab | synonymous_D220D |
| 934 | C | T | | ORF1ab | synonymous_D223D |
| 936 | C | T | | ORF1ab | nonsynonymous_T224I |
| 945 | G | T | | ORF1ab | nonsynonymous_G227V |

| | | | | | |
|---|---|---|---|---|---|
| 946 | T | C | | ORF1ab | synonymous_G227G |
| 947 | G | T | | ORF1ab | nonsynonymous_V228L |
| 951 | A | G | | ORF1ab | nonsynonymous_Y229C |
| 952 | C | T | | ORF1ab | synonymous_Y229Y |
| 955 | C | T | | ORF1ab | synonymous_C230C |
| 957 | G | A | | ORF1ab | nonsynonymous_C231Y |
| 960 | G | A | | ORF1ab | nonsynonymous_R232H |
| 975 | A | C | | ORF1ab | nonsynonymous_E237A |
| 981 | C | T | | ORF1ab | nonsynonymous_A239V |
| 985 | G | C | | ORF1ab | nonsynonymous_W240C |
| 988 | C | T | | ORF1ab | synonymous_Y241Y |
| 990 | C | T | | ORF1ab | nonsynonymous_T242M |
| 998 | T | G | | ORF1ab | nonsynonymous_S245A |
| 1002 | A | G | | ORF1ab | nonsynonymous_E246G |
| 1006 | G | T | | ORF1ab | nonsynonymous_K247N |
| 1015 | A | G | | ORF1ab | synonymous_E250E |
| 1018 | G | T | | ORF1ab | nonsynonymous_L251F |
| 1032 | A | T | | ORF1ab | nonsynonymous_E256V |
| 1039 | A | G | | ORF1ab | synonymous_K258K |
| 1042 | G | T | | ORF1ab | nonsynonymous_L259F |
| 1044 | C | T | | ORF1ab | nonsynonymous_A260V |
| 1059 | C | T | | ORF1ab | nonsynonymous_T265I |
| 1060 | C | T | | ORF1ab | synonymous_T265T |
| 1074 | G | T | | ORF1ab | nonsynonymous_C270F |
| 1075 | T | C | | ORF1ab | synonymous_C270C |
| 1076 | C | T | | ORF1ab | nonsynonymous_P271S |
| 1079 | A | C | | ORF1ab | nonsynonymous_N272H |
| 1080 | A | T | | ORF1ab | nonsynonymous_N272I |
| 1087 | A | T | G | ORF1ab | synonymous_V274V;<br>synonymous_V274V |
| 1091 | C | T | | ORF1ab | nonsynonymous_P276S |

| | | | | | |
|---|---|---|---|---|---|
| 1093 | C | T | | ORF1ab | synonymous_P276P |
| 1102 | C | T | | ORF1ab | synonymous_S279S |
| 1113 | C | T | | ORF1ab | nonsynonymous_T283I |
| 1115 | A | G | | ORF1ab | nonsynonymous_I284V |
| 1121 | C | T | | ORF1ab | nonsynonymous_P286S |
| 1126 | G | T | | ORF1ab | nonsynonymous_R287S |
| 1134 | A | T | | ORF1ab | nonsynonymous_K290M |
| 1145 | G | A | | ORF1ab | nonsynonymous_D294N |
| 1150 | C | T | | ORF1ab | synonymous_G295G |
| 1166 | C | T | | ORF1ab | stopgain_R301X |
| 1168 | A | C | T | ORF1ab | synonymous_R301R;<br>synonymous_R301R |
| 1170 | C | T | | ORF1ab | nonsynonymous_S302F |
| 1181 | G | T | | ORF1ab | nonsynonymous_V306F |
| 1186 | G | T | | ORF1ab | synonymous_A307A |
| 1190 | C | T | | ORF1ab | nonsynonymous_P309S |
| 1191 | C | T | | ORF1ab | nonsynonymous_P309L |
| 1201 | C | T | | ORF1ab | synonymous_C312C |
| 1211 | T | G | | ORF1ab | nonsynonymous_C316G |
| 1214 | C | T | | ORF1ab | nonsynonymous_L317F |
| 1216 | T | C | | ORF1ab | synonymous_L317L |
| 1218 | C | T | | ORF1ab | nonsynonymous_S318L |
| 1221 | C | T | | ORF1ab | nonsynonymous_T319I |
| 1225 | C | T | | ORF1ab | synonymous_L320L |
| 1235 | G | T | | ORF1ab | nonsynonymous_D324Y |
| 1238 | C | T | | ORF1ab | nonsynonymous_H325Y |
| 1244 | G | T | | ORF1ab | nonsynonymous_G327C |
| 1259 | C | T | | ORF1ab | stopgain_Q332X |
| 1263 | C | T | | ORF1ab | nonsynonymous_T333M |
| 1266 | G | T | | ORF1ab | nonsynonymous_G334V |
| 1278 | A | G | | ORF1ab | nonsynonymous_K338R |

| | | | | | |
|---|---|---|---|---|---|
| 1281 | C | T | | ORF1ab | nonsynonymous_A339V |
| 1288 | C | T | | ORF1ab | synonymous_C341C |
| 1289 | G | A | | ORF1ab | nonsynonymous_E342K |
| 1299 | G | A | | ORF1ab | nonsynonymous_G345D |
| 1307 | A | C | | ORF1ab | nonsynonymous_N348H |
| 1308 | A | G | | ORF1ab | nonsynonymous_N348S |
| 1312 | G | T | | ORF1ab | nonsynonymous_L349F |
| 1314 | C | T | | ORF1ab | nonsynonymous_T350I |
| 1315 | T | A | | ORF1ab | synonymous_T350T |
| 1319 | G | A | | ORF1ab | nonsynonymous_E352K |
| 1322 | G | T | | ORF1ab | nonsynonymous_G353C |
| 1337 | G | T | | ORF1ab | nonsynonymous_G358C |
| 1338 | G | T | | ORF1ab | nonsynonymous_G358V |
| 1339 | T | C | | ORF1ab | synonymous_G358G |
| 1346 | C | T | | ORF1ab | nonsynonymous_P361S |
| 1347 | C | T | | ORF1ab | nonsynonymous_P361L |
| 1349 | C | T | | ORF1ab | stopgain_Q362X |
| 1367 | A | C | | ORF1ab | nonsynonymous_I368L |
| 1368 | T | C | | ORF1ab | nonsynonymous_I368T |
| 1383 | G | A | | ORF1ab | nonsynonymous_C373Y |
| 1387 | C | T | | ORF1ab | synonymous_H374H |
| 1392 | C | T | | ORF1ab | nonsynonymous_S376L |
| 1397 | G | A | | ORF1ab | nonsynonymous_V378I |
| 1403 | C | T | | ORF1ab | nonsynonymous_P380S |
| 1404 | C | A | T | ORF1ab | nonsynonymous_P380H; nonsynonymous_P380L |
| 1406 | G | T | | ORF1ab | stopgain_E381X |
| 1407 | A | G | | ORF1ab | nonsynonymous_E381G |
| 1409 | C | T | | ORF1ab | nonsynonymous_H382Y |
| 1418 | G | T | | ORF1ab | nonsynonymous_A385S |
| 1419 | C | T | | ORF1ab | nonsynonymous_A385V |

| | | | | | |
|---|---|---|---|---|---|
| 1420 | C | T | | ORF1ab | synonymous_A385A |
| 1421 | G | A | | ORF1ab | nonsynonymous_E386K |
| 1427 | C | T | | ORF1ab | nonsynonymous_H388Y |
| 1430 | A | G | | ORF1ab | nonsynonymous_N389D |
| 1435 | A | C | | ORF1ab | nonsynonymous_E390D |
| 1440 | G | A | T | ORF1ab | nonsynonymous_G392D; nonsynonymous_G392V |
| 1449 | C | T | | ORF1ab | nonsynonymous_T395I |
| 1457 | C | T | | ORF1ab | nonsynonymous_R398C |
| 1463 | G | T | | ORF1ab | nonsynonymous_G400C |
| 1466 | G | T | | ORF1ab | nonsynonymous_G401C |
| 1469 | C | T | | ORF1ab | nonsynonymous_R402C |
| 1471 | C | T | | ORF1ab | synonymous_R402R |
| 1472 | A | G | | ORF1ab | nonsynonymous_T403A |
| 1473 | C | T | | ORF1ab | nonsynonymous_T403I |
| 1478 | G | C | | ORF1ab | nonsynonymous_A405P |
| 1479 | C | T | | ORF1ab | nonsynonymous_A405V |
| 1480 | C | T | | ORF1ab | synonymous_A405A |
| 1483 | T | A | | ORF1ab | nonsynonymous_F406L |
| 1488 | G | T | | ORF1ab | nonsynonymous_G408V |
| 1489 | C | T | | ORF1ab | synonymous_G408G |
| 1491 | G | T | | ORF1ab | nonsynonymous_C409F |
| 1495 | G | T | | ORF1ab | synonymous_V410V |
| 1511 | T | C | | ORF1ab | nonsynonymous_C416R |
| 1512 | G | T | | ORF1ab | nonsynonymous_C416F |
| 1514 | C | T | | ORF1ab | nonsynonymous_H417Y |
| 1515 | A | G | | ORF1ab | nonsynonymous_H417R |
| 1519 | C | T | | ORF1ab | synonymous_N418N |
| 1538 | C | T | | ORF1ab | nonsynonymous_P425S |
| 1539 | C | T | | ORF1ab | nonsynonymous_P425L |

| | | | | | |
|---|---|---|---|---|---|
| 1547 | A | T | G | ORF1ab | nonsynonymous_S428C; nonsynonymous_S428G |
| 1549 | C | T | | ORF1ab | synonymous_S428S |
| 1560 | G | T | | ORF1ab | nonsynonymous_G432V |
| 1567 | C | T | | ORF1ab | synonymous_N434N |
| 1568 | C | T | | ORF1ab | nonsynonymous_H435Y |
| 1570 | T | C | | ORF1ab | synonymous_H435H |
| 1572 | C | T | | ORF1ab | nonsynonymous_T436I |
| 1574 | G | T | | ORF1ab | nonsynonymous_G437C |
| 1580 | G | T | | ORF1ab | nonsynonymous_V439F |
| 1587 | A | G | | ORF1ab | nonsynonymous_E441G |
| 1589 | G | T | | ORF1ab | nonsynonymous_G442C |
| 1592 | T | C | | ORF1ab | nonsynonymous_S443P |
| 1594 | C | T | | ORF1ab | synonymous_S443S |
| 1595 | G | T | | ORF1ab | stopgain_E444X |
| 1610 | A | T | | ORF1ab | nonsynonymous_N449Y |
| 1646 | A | G | | ORF1ab | nonsynonymous_I461V |
| 1651 | T | C | | ORF1ab | synonymous_N462N |
| 1658 | G | T | | ORF1ab | nonsynonymous_G465C |
| 1670 | C | T | | ORF1ab | nonsynonymous_L469F |
| 1676 | G | T | | ORF1ab | stopgain_E471X |
| 1681 | G | T | | ORF1ab | nonsynonymous_E472D |
| 1683 | T | C | | ORF1ab | nonsynonymous_I473T |
| 1702 | T | C | | ORF1ab | synonymous_S479S |
| 1707 | C | T | | ORF1ab | nonsynonymous_S481F |
| 1711 | T | C | | ORF1ab | synonymous_A482A |
| 1713 | C | T | | ORF1ab | nonsynonymous_S483F |
| 1721 | G | T | | ORF1ab | nonsynonymous_A486S |
| 1722 | C | T | | ORF1ab | nonsynonymous_A486V |
| 1727 | G | T | | ORF1ab | nonsynonymous_V488L |
| 1730 | G | A | | ORF1ab | nonsynonymous_E489K |

| | | | | | |
|---|---|---|---|---|---|
| 1734 | C | T | | ORF1ab | nonsynonymous_T490I |
| 1738 | G | A | | ORF1ab | synonymous_V491V |
| 1745 | T | C | | ORF1ab | synonymous_L494L |
| 1748 | G | T | | ORF1ab | nonsynonymous_D495Y |
| 1752 | A | T | | ORF1ab | nonsynonymous_Y496F |
| 1757 | G | C | | ORF1ab | nonsynonymous_A498P |
| 1762 | C | T | | ORF1ab | synonymous_F499F |
| 1779 | C | T | | ORF1ab | nonsynonymous_S505F |
| 1783 | T | C | | ORF1ab | synonymous_C506C |
| 1799 | A | G | | ORF1ab | nonsynonymous_T512A |
| 1801 | A | G | | ORF1ab | synonymous_T512T |
| 1805 | G | A | | ORF1ab | nonsynonymous_G514R |
| 1806 | G | A | | ORF1ab | nonsynonymous_G514E |
| 1812 | C | T | | ORF1ab | nonsynonymous_A516V |
| 1819 | A | G | | ORF1ab | synonymous_K518K |
| 1820 | G | A | | ORF1ab | nonsynonymous_G519S |
| 1841 | C | T | | ORF1ab | stopgain_Q526X |
| 1843 | G | A | | ORF1ab | synonymous_Q526Q |
| 1848 | C | T | | ORF1ab | nonsynonymous_S528L |
| 1853 | C | T | | ORF1ab | synonymous_L530L |
| 1857 | G | T | | ORF1ab | nonsynonymous_S531I |
| 1862 | C | T | | ORF1ab | nonsynonymous_L533F |
| 1875 | C | T | | ORF1ab | nonsynonymous_A537V |
| 1878 | C | T | | ORF1ab | nonsynonymous_S538L |
| 1880 | G | T | | ORF1ab | stopgain_E539X |
| 1882 | G | T | | ORF1ab | nonsynonymous_E539D |
| 1884 | C | T | | ORF1ab | nonsynonymous_A540V |
| 1893 | T | C | | ORF1ab | nonsynonymous_V543A |
| 1897 | A | T | | ORF1ab | synonymous_V544V |
| 1904 | A | G | | ORF1ab | nonsynonymous_I547V |
| 1909 | C | T | | ORF1ab | synonymous_F548F |
| 1911 | C | T | | ORF1ab | nonsynonymous_S549F |

| | | | | | |
|---|---|---|---|---|---|
| 1912 | C | T | | ORF1ab | synonymous_S549S |
| 1914 | G | T | | ORF1ab | nonsynonymous_R550L |
| 1917 | C | T | | ORF1ab | nonsynonymous_T551I |
| 1918 | T | C | | ORF1ab | synonymous_T551T |
| 1919 | C | T | | ORF1ab | nonsynonymous_L552F |
| 1922 | G | T | | ORF1ab | stopgain_E553X |
| 1929 | C | T | | ORF1ab | nonsynonymous_A555V |
| 1930 | T | G | | ORF1ab | synonymous_A555A |
| 1938 | C | T | | ORF1ab | nonsynonymous_S558F |
| 1943 | C | T | | ORF1ab | nonsynonymous_R560C |
| 1945 | T | C | | ORF1ab | synonymous_R560R |
| 1946 | G | T | | ORF1ab | nonsynonymous_V561F |
| 1948 | T | C | | ORF1ab | synonymous_V561V |
| 1957 | G | T | | ORF1ab | nonsynonymous_K564N |
| 1958 | G | A | | ORF1ab | nonsynonymous_A565T |
| 1961 | G | T | | ORF1ab | nonsynonymous_A566S |
| 1962 | C | T | | ORF1ab | nonsynonymous_A566V |
| 1963 | T | C | | ORF1ab | synonymous_A566A |
| 1971 | T | C | | ORF1ab | nonsynonymous_I569T |
| 1973 | C | T | | ORF1ab | synonymous_L570L |
| 1975 | A | T | | ORF1ab | synonymous_L570L |
| 1979 | G | T | | ORF1ab | stopgain_G572X |
| 1980 | G | T | | ORF1ab | nonsynonymous_G572V |
| 1988 | C | T | | ORF1ab | stopgain_Q575X |
| 1995 | C | T | | ORF1ab | nonsynonymous_S577L |
| 1997 | C | T | | ORF1ab | synonymous_L578L |
| 2001 | G | C | T | ORF1ab | nonsynonymous_R579T; nonsynonymous_R579I |
| 2006 | A | T | | ORF1ab | nonsynonymous_I581F |
| 2007 | T | G | | ORF1ab | nonsynonymous_I581S |
| 2018 | A | G | | ORF1ab | nonsynonymous_M585V |

| | | | | | |
|---|---|---|---|---|---|
| 2019 | T | C | | ORF1ab | nonsynonymous_M585T |
| 2035 | G | T | | ORF1ab | nonsynonymous_L590F |
| 2036 | G | C | T | ORF1ab | nonsynonymous_A591P;<br>nonsynonymous_A591S |
| 2037 | C | T | | ORF1ab | nonsynonymous_A591V |
| 2043 | A | T | | ORF1ab | nonsynonymous_N593I |
| 2047 | T | C | | ORF1ab | synonymous_N594N |
| 2061 | C | T | | ORF1ab | nonsynonymous_A599V |
| 2062 | C | T | | ORF1ab | synonymous_A599A |
| 2064 | A | G | | ORF1ab | nonsynonymous_Y600C |
| 2065 | C | T | | ORF1ab | synonymous_Y600Y |
| 2080 | T | C | | ORF1ab | synonymous_V605V |
| 2084 | C | T | | ORF1ab | stopgain_Q607X |
| 2091 | C | T | | ORF1ab | nonsynonymous_T609I |
| 2094 | C | T | | ORF1ab | nonsynonymous_S610L |
| 2098 | G | T | | ORF1ab | nonsynonymous_Q611H |
| 2101 | G | T | | ORF1ab | nonsynonymous_W612C |
| 2102 | C | T | | ORF1ab | synonymous_L613L |
| 2106 | C | T | | ORF1ab | nonsynonymous_T614I |
| 2110 | C | T | | ORF1ab | synonymous_N615N |
| 2113 | C | T | | ORF1ab | synonymous_I616I |
| 2114 | T | A | | ORF1ab | nonsynonymous_F617I |
| 2141 | C | T | | ORF1ab | nonsynonymous_P626S |
| 2144 | G | T | | ORF1ab | nonsynonymous_V627F |
| 2145 | T | C | | ORF1ab | nonsynonymous_V627A |
| 2150 | G | A | | ORF1ab | nonsynonymous_D629N |
| 2156 | C | T | | ORF1ab | nonsynonymous_L631F |
| 2159 | G | T | | ORF1ab | stopgain_E632X |
| 2160 | A | T | | ORF1ab | nonsynonymous_E632V |
| 2164 | G | T | | ORF1ab | nonsynonymous_E633D |
| 2166 | A | T | | ORF1ab | nonsynonymous_K634M |

| | | | | | |
|---|---|---|---|---|---|
| 2168 | T | G | | ORF1ab | nonsynonymous_F635V |
| 2182 | A | G | | ORF1ab | synonymous_V639V |
| 2189 | C | T | | ORF1ab | nonsynonymous_L642F |
| 2196 | A | C | | ORF1ab | nonsynonymous_D644A |
| 2204 | G | A | | ORF1ab | nonsynonymous_E647K |
| 2215 | A | G | | ORF1ab | synonymous_K650K |
| 2217 | T | C | | ORF1ab | nonsynonymous_F651S |
| 2237 | G | A | | ORF1ab | nonsynonymous_E658K |
| 2240 | A | G | | ORF1ab | nonsynonymous_I659V |
| 2247 | G | T | | ORF1ab | nonsynonymous_G661V |
| 2248 | T | A | | ORF1ab | synonymous_G661G |
| 2250 | G | T | | ORF1ab | nonsynonymous_G662V |
| 2258 | G | T | | ORF1ab | nonsynonymous_V665F |
| 2281 | G | T | | ORF1ab | nonsynonymous_K672N |
| 2283 | A | C | | ORF1ab | nonsynonymous_E673A |
| 2288 | G | T | | ORF1ab | nonsynonymous_V675F |
| 2293 | G | T | | ORF1ab | nonsynonymous_Q676H |
| 2294 | A | C | | ORF1ab | nonsynonymous_T677P |
| 2299 | C | T | | ORF1ab | synonymous_F678F |
| 2304 | A | T | | ORF1ab | nonsynonymous_K680M |
| 2309 | G | T | | ORF1ab | nonsynonymous_V682L |
| 2318 | T | G | | ORF1ab | nonsynonymous_F685V |
| 2327 | T | C | | ORF1ab | synonymous_L688L |
| 2334 | C | T | | ORF1ab | nonsynonymous_A690V |
| 2337 | A | C | | ORF1ab | nonsynonymous_D691A |
| 2338 | C | T | | ORF1ab | synonymous_D691D |
| 2354 | G | T | | ORF1ab | stopgain_G697X |
| 2360 | A | T | | ORF1ab | stopgain_K699X |
| 2363 | C | T | | ORF1ab | nonsynonymous_L700F |
| 2366 | A | T | | ORF1ab | stopgain_K701X |
| 2370 | C | T | | ORF1ab | nonsynonymous_A702V |
| 2381 | G | T | | ORF1ab | nonsynonymous_G706C |

| | | | | | |
|---|---|---|---|---|---|
| 2388 | C | T | | ORF1ab | nonsynonymous_T708I |
| 2389 | A | T | | ORF1ab | synonymous_T708T |
| 2393 | G | A | | ORF1ab | nonsynonymous_V710I |
| 2395 | C | T | | ORF1ab | synonymous_V710V |
| 2397 | C | T | | ORF1ab | nonsynonymous_T711M |
| 2398 | G | A | | ORF1ab | synonymous_T711T |
| 2401 | C | A | | ORF1ab | nonsynonymous_H712Q |
| 2403 | C | T | | ORF1ab | nonsynonymous_S713L |
| 2404 | A | C | | ORF1ab | synonymous_S713S |
| 2409 | G | C | | ORF1ab | nonsynonymous_G715A |
| 2415 | A | G | | ORF1ab | nonsynonymous_Y717C |
| 2416 | C | T | | ORF1ab | synonymous_Y717Y |
| 2423 | T | G | | ORF1ab | nonsynonymous_C720G |
| 2434 | C | T | | ORF1ab | synonymous_S723S |
| 2444 | A | C | | ORF1ab | nonsynonymous_T727P |
| 2445 | C | T | | ORF1ab | nonsynonymous_T727I |
| 2446 | T | C | | ORF1ab | synonymous_T727T |
| 2447 | G | T | | ORF1ab | nonsynonymous_G728C |
| 2448 | G | A | | ORF1ab | nonsynonymous_G728D |
| 2455 | C | T | | ORF1ab | synonymous_L730L |
| 2459 | C | T | | ORF1ab | nonsynonymous_P732S |
| 2466 | A | G | | ORF1ab | nonsynonymous_K734R |
| 2468 | G | C | | ORF1ab | nonsynonymous_A735P |
| 2470 | C | T | | ORF1ab | synonymous_A735A |
| 2476 | A | G | | ORF1ab | synonymous_K737K |
| 2480 | A | G | | ORF1ab | nonsynonymous_I739V |
| 2485 | C | T | | ORF1ab | synonymous_I740I |
| 2489 | T | C | | ORF1ab | synonymous_L742L |
| 2492 | G | T | | ORF1ab | stopgain_E743X |
| 2495 | G | A | | ORF1ab | nonsynonymous_G744R |
| 2498 | G | A | | ORF1ab | nonsynonymous_E745K |
| 2499 | A | G | | ORF1ab | nonsynonymous_E745G |

| | | | | | |
|---|---|---|---|---|---|
| 2502 | C | T | | ORF1ab | nonsynonymous_T746I |
| 2507 | C | T | | ORF1ab | nonsynonymous_P748S |
| 2508 | C | T | | ORF1ab | nonsynonymous_P748L |
| 2509 | C | T | | ORF1ab | synonymous_P748P |
| 2510 | A | G | | ORF1ab | nonsynonymous_T749A |
| 2511 | C | T | | ORF1ab | nonsynonymous_T749I |
| 2518 | G | T | | ORF1ab | synonymous_V751V |
| 2523 | C | T | | ORF1ab | nonsynonymous_T753I |
| 2524 | A | C | T | ORF1ab | synonymous_T753T; synonymous_T753T |
| 2528 | G | A | | ORF1ab | nonsynonymous_E755K |
| 2534 | G | T | | ORF1ab | nonsynonymous_V757F |
| 2536 | C | T | | ORF1ab | synonymous_V757V |
| 2552 | T | C | | ORF1ab | synonymous_L763L |
| 2555 | C | T | | ORF1ab | stopgain_Q764X |
| 2558 | C | T | | ORF1ab | nonsynonymous_P765S |
| 2561 | T | C | | ORF1ab | synonymous_L766L |
| 2564 | G | T | | ORF1ab | stopgain_E767X |
| 2567 | C | T | | ORF1ab | stopgain_Q768X |
| 2570 | C | T | | ORF1ab | nonsynonymous_P769S |
| 2571 | C | T | | ORF1ab | nonsynonymous_P769L |
| 2585 | G | T | | ORF1ab | nonsynonymous_V774F |
| 2595 | C | T | | ORF1ab | nonsynonymous_P777L |
| 2604 | G | A | T | ORF1ab | nonsynonymous_G780D; nonsynonymous_G780V |
| 2606 | A | T | | ORF1ab | nonsynonymous_T781S |
| 2612 | G | T | | ORF1ab | nonsynonymous_V783F |
| 2623 | C | G | | ORF1ab | nonsynonymous_N786K |
| 2632 | G | T | | ORF1ab | nonsynonymous_M789I |
| 2635 | G | A | | ORF1ab | synonymous_L790L |

| | | | | | |
|---|---|---|---|---|---|
| 2638 | C | A | G | ORF1ab | synonymous_L791L; synonymous_L791L |
| 2639 | G | A | | ORF1ab | nonsynonymous_E792K |
| 2644 | C | T | | ORF1ab | synonymous_I793I |
| 2648 | G | T | | ORF1ab | nonsynonymous_D795Y |
| 2662 | C | T | | ORF1ab | synonymous_Y799Y |
| 2669 | C | A | T | ORF1ab | nonsynonymous_L802I; nonsynonymous_L802F |
| 2673 | C | T | | ORF1ab | nonsynonymous_A803V |
| 2675 | C | T | | ORF1ab | nonsynonymous_P804S |
| 2676 | C | T | | ORF1ab | nonsynonymous_P804L |
| 2691 | C | A | T | ORF1ab | nonsynonymous_T809K; nonsynonymous_T809I |
| 2695 | C | T | | ORF1ab | synonymous_N810N |
| 2698 | T | C | | ORF1ab | synonymous_N811N |
| 2705 | A | G | | ORF1ab | nonsynonymous_T814A |
| 2706 | C | T | | ORF1ab | nonsynonymous_T814I |
| 2710 | C | T | G | ORF1ab | synonymous_L815L; synonymous_L815L |
| 2716 | C | T | | ORF1ab | synonymous_G817G |
| 2718 | G | T | | ORF1ab | nonsynonymous_G818V |
| 2737 | T | C | | ORF1ab | synonymous_T824T |
| 2744 | G | C | | ORF1ab | nonsynonymous_D827H |
| 2756 | A | G | | ORF1ab | nonsynonymous_I831V |
| 2759 | G | A | T | ORF1ab | nonsynonymous_E832K; stopgain_E832X |
| 2765 | C | T | | ORF1ab | stopgain_Q834X |
| 2768 | G | T | | ORF1ab | nonsynonymous_G835C |
| 2769 | G | C | T | ORF1ab | nonsynonymous_G835A; nonsynonymous_G835V |
| 2773 | C | T | | ORF1ab | synonymous_Y836Y |

| | | | | | |
|---|---|---|---|---|---|
| 2786 | A | G | | ORF1ab | nonsynonymous_I841V |
| 2788 | C | T | | ORF1ab | synonymous_I841I |
| 2798 | C | T | | ORF1ab | nonsynonymous_L845F |
| 2802 | A | T | | ORF1ab | nonsynonymous_D846V |
| 2809 | G | T | | ORF1ab | nonsynonymous_R848S |
| 2822 | C | T | | ORF1ab | nonsynonymous_L853F |
| 2832 | A | G | | ORF1ab | nonsynonymous_K856R |
| 2836 | C | T | | ORF1ab | synonymous_C857C |
| 2861 | A | G | | ORF1ab | nonsynonymous_T866A |
| 2868 | T | A | | ORF1ab | nonsynonymous_V868E |
| 2872 | T | C | | ORF1ab | synonymous_N869N |
| 2891 | G | A | | ORF1ab | nonsynonymous_A876T |
| 2910 | C | T | | ORF1ab | nonsynonymous_T882I |
| 2913 | T | C | | ORF1ab | nonsynonymous_L883S |
| 2914 | G | A | | ORF1ab | synonymous_L883L |
| 2919 | C | T | | ORF1ab | nonsynonymous_P885L |
| 2930 | T | C | | ORF1ab | synonymous_L889L |
| 2937 | C | T | | ORF1ab | nonsynonymous_T891I |
| 2939 | C | T | | ORF1ab | nonsynonymous_P892S |
| 2942 | C | T | | ORF1ab | synonymous_L893L |
| 2945 | G | A | | ORF1ab | nonsynonymous_G894S |
| 2965 | G | T | | ORF1ab | nonsynonymous_W900C |
| 2970 | T | G | | ORF1ab | nonsynonymous_M902R |
| 2972 | G | T | | ORF1ab | nonsynonymous_A903S |
| 2995 | G | A | | ORF1ab | synonymous_E910E |
| 3004 | G | T | | ORF1ab | nonsynonymous_E913D |
| 3006 | T | C | | ORF1ab | nonsynonymous_F914S |
| 3014 | G | A | | ORF1ab | nonsynonymous_A917T |
| 3025 | G | A | T | ORF1ab | nonsynonymous_M920I;<br>nonsynonymous_M920I |
| 3027 | A | G | | ORF1ab | nonsynonymous_Y921C |

| | | | | | |
|---|---|---|---|---|---|
| 3037 | C | A | T | ORF1ab | nonsynonymous_F924L; synonymous_F924F |
| 3040 | C | T | | ORF1ab | synonymous_Y925Y |
| 3041 | C | T | | ORF1ab | nonsynonymous_P926S |
| 3043 | T | C | | ORF1ab | synonymous_P926P |
| 3045 | C | T | | ORF1ab | nonsynonymous_P927L |
| 3046 | A | G | | ORF1ab | synonymous_P927P |
| 3047 | G | T | | ORF1ab | nonsynonymous_D928Y |
| 3058 | A | T | | ORF1ab | nonsynonymous_E931D |
| 3061 | A | G | | ORF1ab | synonymous_E932E |
| 3079 | A | G | | ORF1ab | synonymous_E938E |
| 3085 | G | T | | ORF1ab | nonsynonymous_E940D |
| 3088 | T | C | | ORF1ab | synonymous_F941F |
| 3090 | A | T | | ORF1ab | nonsynonymous_E942V |
| 3096 | C | T | | ORF1ab | nonsynonymous_S944L |
| 3099 | C | T | | ORF1ab | nonsynonymous_T945I |
| 3101 | C | A | | ORF1ab | nonsynonymous_Q946K |
| 3106 | T | C | | ORF1ab | synonymous_Y947Y |
| 3109 | G | A | | ORF1ab | synonymous_E948E |
| 3117 | C | T | | ORF1ab | nonsynonymous_T951I |
| 3130 | C | T | | ORF1ab | synonymous_Y955Y |
| 3132 | A | G | | ORF1ab | nonsynonymous_Q956R |
| 3140 | C | T | | ORF1ab | nonsynonymous_P959S |
| 3142 | T | C | | ORF1ab | synonymous_P959P |
| 3146 | G | A | | ORF1ab | nonsynonymous_E961K |
| 3150 | T | C | | ORF1ab | nonsynonymous_F962S |
| 3157 | C | T | | ORF1ab | synonymous_A964A |
| 3160 | T | C | | ORF1ab | synonymous_T965T |
| 3164 | G | T | | ORF1ab | nonsynonymous_A967S |
| 3166 | T | C | | ORF1ab | synonymous_A967A |
| 3167 | G | A | | ORF1ab | nonsynonymous_A968T |

| | | | | | |
|---|---|---|---|---|---|
| 3168 | C | T | | ORF1ab | nonsynonymous_A968V |
| 3175 | A | C | | ORF1ab | nonsynonymous_Q970H |
| 3176 | C | T | | ORF1ab | nonsynonymous_P971S |
| 3177 | C | T | | ORF1ab | nonsynonymous_P971L |
| 3179 | G | T | | ORF1ab | stopgain_E972X |
| 3180 | A | G | | ORF1ab | nonsynonymous_E972G |
| 3191 | G | T | | ORF1ab | stopgain_E976X |
| 3194 | G | T | | ORF1ab | stopgain_E977X |
| 3209 | G | A | | ORF1ab | nonsynonymous_D982N |
| 3210 | A | T | | ORF1ab | nonsynonymous_D982V |
| 3216 | G | T | | ORF1ab | nonsynonymous_S984I |
| 3230 | G | C | T | ORF1ab | nonsynonymous_G989R; nonsynonymous_G989C |
| 3232 | T | C | | ORF1ab | synonymous_G989G |
| 3236 | C | T | | ORF1ab | stopgain_Q991X |
| 3237 | A | G | | ORF1ab | nonsynonymous_Q991R |
| 3238 | A | T | | ORF1ab | nonsynonymous_Q991H |
| 3240 | A | T | | ORF1ab | nonsynonymous_D992V |
| 3242 | G | A | | ORF1ab | nonsynonymous_G993S |
| 3248 | G | A | | ORF1ab | nonsynonymous_E995K |
| 3250 | G | C | | ORF1ab | nonsynonymous_E995D |
| 3251 | G | T | | ORF1ab | nonsynonymous_D996Y |
| 3253 | C | T | | ORF1ab | synonymous_D996D |
| 3259 | G | A | T | ORF1ab | synonymous_Q998Q; nonsynonymous_Q998H |
| 3261 | C | T | | ORF1ab | nonsynonymous_T999I |
| 3264 | C | T | | ORF1ab | nonsynonymous_T1000I |
| 3270 | T | C | | ORF1ab | nonsynonymous_I1002T |
| 3279 | T | C | | ORF1ab | nonsynonymous_I1005T |
| 3295 | T | C | | ORF1ab | synonymous_P1010P |
| 3299 | T | C | | ORF1ab | synonymous_L1012L |

| | | | | | |
|---|---|---|---|---|---|
| 3306 | T | C | | ORF1ab | nonsynonymous_M1014T |
| 3307 | G | A | | ORF1ab | nonsynonymous_M1014I |
| 3313 | T | C | | ORF1ab | synonymous_L1016L |
| 3318 | C | T | | ORF1ab | nonsynonymous_P1018L |
| 3330 | C | T | | ORF1ab | nonsynonymous_T1022I |
| 3331 | T | C | | ORF1ab | synonymous_T1022T |
| 3342 | A | G | | ORF1ab | nonsynonymous_N1026S |
| 3351 | G | T | | ORF1ab | nonsynonymous_S1029I |
| 3356 | T | C | | ORF1ab | nonsynonymous_Y1031H |
| 3365 | C | T | G | ORF1ab | nonsynonymous_L1034F; nonsynonymous_L1034V |
| 3373 | C | A | | ORF1ab | nonsynonymous_D1036E |
| 3383 | A | T | | ORF1ab | nonsynonymous_I1040F |
| 3384 | T | C | | ORF1ab | nonsynonymous_I1040T |
| 3385 | T | C | | ORF1ab | synonymous_I1040I |
| 3393 | C | T | | ORF1ab | nonsynonymous_A1043V |
| 3403 | G | A | T | ORF1ab | synonymous_V1046V; synonymous_V1046V |
| 3405 | A | C | | ORF1ab | nonsynonymous_E1047A |
| 3406 | A | C | | ORF1ab | nonsynonymous_E1047D |
| 3407 | G | T | | ORF1ab | stopgain_E1048X |
| 3411 | C | T | | ORF1ab | nonsynonymous_A1049V |
| 3419 | G | A | | ORF1ab | nonsynonymous_V1052I |
| 3425 | C | T | | ORF1ab | nonsynonymous_P1054S |
| 3431 | G | T | | ORF1ab | nonsynonymous_V1056L |
| 3447 | C | A | | ORF1ab | nonsynonymous_A1061D |
| 3452 | G | T | | ORF1ab | nonsynonymous_V1063F |
| 3455 | T | C | | ORF1ab | nonsynonymous_Y1064H |
| 3457 | C | T | | ORF1ab | synonymous_Y1064Y |
| 3458 | C | A | | ORF1ab | nonsynonymous_L1065I |
| 3473 | G | T | | ORF1ab | nonsynonymous_G1070C |

| | | | | | |
|---|---|---|---|---|---|
| 3478 | T | C | | ORF1ab | synonymous_V1071V |
| 3483 | G | T | | ORF1ab | nonsynonymous_G1073V |
| 3500 | A | C | | ORF1ab | nonsynonymous_T1079P |
| 3502 | T | C | | ORF1ab | synonymous_T1079T |
| 3504 | A | G | | ORF1ab | nonsynonymous_N1080S |
| 3505 | C | T | | ORF1ab | synonymous_N1080N |
| 3508 | T | C | | ORF1ab | synonymous_N1081N |
| 3513 | T | C | | ORF1ab | nonsynonymous_M1083T |
| 3517 | A | G | | ORF1ab | synonymous_Q1084Q |
| 3518 | G | T | | ORF1ab | nonsynonymous_V1085F |
| 3520 | T | C | | ORF1ab | synonymous_V1085V |
| 3521 | G | A | | ORF1ab | nonsynonymous_E1086K |
| 3551 | C | A | T | ORF1ab | nonsynonymous_P1096T;<br>nonsynonymous_P1096S |
| 3560 | G | A | | ORF1ab | nonsynonymous_V1099M |
| 3564 | G | T | | ORF1ab | nonsynonymous_G1100V |
| 3583 | C | T | | ORF1ab | synonymous_S1106S |
| 3587 | C | T | | ORF1ab | nonsynonymous_H1108Y |
| 3589 | C | T | | ORF1ab | synonymous_H1108H |
| 3593 | C | T | | ORF1ab | nonsynonymous_L1110F |
| 3597 | C | T | | ORF1ab | nonsynonymous_A1111V |
| 3602 | C | T | | ORF1ab | nonsynonymous_H1113Y |
| 3604 | C | T | | ORF1ab | synonymous_H1113H |
| 3618 | T | A | C | ORF1ab | nonsynonymous_V1118D;<br>nonsynonymous_V1118A |
| 3623 | C | T | | ORF1ab | nonsynonymous_P1120S |
| 3625 | A | C | | ORF1ab | synonymous_P1120P |
| 3634 | C | T | | ORF1ab | synonymous_N1123N |
| 3638 | G | C | T | ORF1ab | nonsynonymous_G1125R;<br>nonsynonymous_G1125C |
| 3639 | G | A | | ORF1ab | nonsynonymous_G1125D |

| | | | | | |
|---|---|---|---|---|---|
| 3646 | C | T | | ORF1ab | synonymous_D1127D |
| 3652 | A | T | | ORF1ab | nonsynonymous_Q1129H |
| 3653 | C | T | | ORF1ab | nonsynonymous_L1130F |
| 3656 | C | T | | ORF1ab | nonsynonymous_L1131F |
| 3661 | G | T | | ORF1ab | nonsynonymous_K1132N |
| 3671 | G | T | | ORF1ab | stopgain_E1136X |
| 3675 | A | T | | ORF1ab | nonsynonymous_N1137I |
| 3679 | T | C | | ORF1ab | synonymous_F1138F |
| 3689 | G | T | | ORF1ab | stopgain_E1142X |
| 3690 | A | T | | ORF1ab | nonsynonymous_E1142V |
| 3695 | C | T | | ORF1ab | synonymous_L1144L |
| 3698 | C | T | | ORF1ab | nonsynonymous_L1145F |
| 3707 | T | C | | ORF1ab | synonymous_L1148L |
| 3710 | T | C | | ORF1ab | synonymous_L1149L |
| 3713 | T | C | | ORF1ab | nonsynonymous_S1150P |
| 3714 | C | T | | ORF1ab | nonsynonymous_S1150L |
| 3717 | C | T | | ORF1ab | nonsynonymous_A1151V |
| 3720 | G | T | | ORF1ab | nonsynonymous_G1152V |
| 3727 | T | C | | ORF1ab | synonymous_F1154F |
| 3728 | G | T | | ORF1ab | nonsynonymous_G1155C |
| 3731 | G | T | | ORF1ab | nonsynonymous_A1156S |
| 3732 | C | T | | ORF1ab | nonsynonymous_A1156V |
| 3736 | C | T | | ORF1ab | synonymous_D1157D |
| 3737 | C | T | | ORF1ab | nonsynonymous_P1158S |
| 3738 | C | T | | ORF1ab | nonsynonymous_P1158L |
| 3743 | C | T | | ORF1ab | nonsynonymous_H1160Y |
| 3753 | G | T | | ORF1ab | nonsynonymous_R1163I |
| 3768 | C | T | | ORF1ab | nonsynonymous_T1168I |
| 3770 | G | T | | ORF1ab | nonsynonymous_V1169F |
| 3773 | C | T | | ORF1ab | nonsynonymous_R1170C |
| 3777 | C | A | | ORF1ab | nonsynonymous_T1171K |
| 3782 | G | T | | ORF1ab | nonsynonymous_V1173F |

| | | | | | |
|---|---|---|---|---|---|
| 3784 | C | A | T | ORF1ab | synonymous_V1173V; synonymous_V1173V |
| 3787 | C | T | | ORF1ab | synonymous_Y1174Y |
| 3792 | C | T | | ORF1ab | nonsynonymous_A1176V |
| 3796 | C | T | | ORF1ab | synonymous_V1177V |
| 3797 | T | C | G | ORF1ab | nonsynonymous_F1178L; nonsynonymous_F1178V |
| 3808 | T | C | | ORF1ab | synonymous_N1181N |
| 3815 | G | T | | ORF1ab | nonsynonymous_D1184Y |
| 3817 | C | T | | ORF1ab | synonymous_D1184D |
| 3819 | A | G | | ORF1ab | nonsynonymous_K1185R |
| 3821 | C | T | | ORF1ab | nonsynonymous_L1186F |
| 3830 | A | G | | ORF1ab | nonsynonymous_S1189G |
| 3831 | G | A | | ORF1ab | nonsynonymous_S1189N |
| 3839 | G | T | | ORF1ab | stopgain_E1192X |
| 3841 | A | T | | ORF1ab | nonsynonymous_E1192D |
| 3868 | A | G | | ORF1ab | synonymous_Q1201Q |
| 3870 | A | G | | ORF1ab | nonsynonymous_K1202R |
| 3874 | C | A | T | ORF1ab | synonymous_I1203I; synonymous_I1203I |
| 3875 | G | A | | ORF1ab | nonsynonymous_A1204T |
| 3877 | T | G | | ORF1ab | synonymous_A1204A |
| 3883 | T | C | | ORF1ab | synonymous_I1206I |
| 3885 | C | A | T | ORF1ab | nonsynonymous_P1207H; nonsynonymous_P1207L |
| 3892 | G | T | | ORF1ab | nonsynonymous_E1209D |
| 3894 | A | G | | ORF1ab | nonsynonymous_E1210G |
| 3896 | G | T | | ORF1ab | nonsynonymous_V1211F |
| 3902 | C | T | | ORF1ab | nonsynonymous_P1213S |
| 3908 | A | T | | ORF1ab | nonsynonymous_I1215L |
| 3919 | T | C | | ORF1ab | synonymous_S1218S |

| | | | | | |
|---|---|---|---|---|---|
| 3923 | C | T | | ORF1ab | nonsynonymous_P1220S |
| 3924 | C | T | | ORF1ab | nonsynonymous_P1220L |
| 3932 | G | C | | ORF1ab | nonsynonymous_E1223Q |
| 3935 | C | T | | ORF1ab | stopgain_Q1224X |
| 3940 | A | G | | ORF1ab | synonymous_R1225R |
| 3946 | A | G | | ORF1ab | synonymous_Q1227Q |
| 3948 | A | G | | ORF1ab | nonsynonymous_D1228G |
| 3950 | G | A | | ORF1ab | nonsynonymous_D1229N |
| 3951 | A | T | | ORF1ab | nonsynonymous_D1229V |
| 3955 | G | T | | ORF1ab | nonsynonymous_K1230N |
| 3961 | C | T | | ORF1ab | synonymous_I1232I |
| 3962 | A | G | | ORF1ab | nonsynonymous_K1233E |
| 3971 | G | T | | ORF1ab | nonsynonymous_V1236F |
| 3977 | G | T | | ORF1ab | stopgain_E1238X |
| 3989 | A | G | | ORF1ab | nonsynonymous_T1242A |
| 3991 | T | A | | ORF1ab | synonymous_T1242T |
| 3996 | A | C | | ORF1ab | nonsynonymous_E1244A |
| 4002 | C | T | | ORF1ab | nonsynonymous_T1246I |
| 4009 | C | T | | ORF1ab | synonymous_F1248F |
| 4015 | A | G | | ORF1ab | synonymous_T1250T |
| 4017 | A | G | | ORF1ab | nonsynonymous_E1251G |
| 4021 | C | T | | ORF1ab | synonymous_N1252N |
| 4027 | A | G | | ORF1ab | synonymous_L1254L |
| 4028 | C | T | | ORF1ab | nonsynonymous_L1255F |
| 4048 | C | T | | ORF1ab | synonymous_G1261G |
| 4049 | A | G | | ORF1ab | nonsynonymous_N1262D |
| 4052 | C | T | | ORF1ab | nonsynonymous_L1263F |
| 4055 | C | T | | ORF1ab | nonsynonymous_H1264Y |
| 4057 | T | C | | ORF1ab | synonymous_H1264H |
| 4059 | C | T | | ORF1ab | nonsynonymous_P1265L |
| 4065 | C | T | | ORF1ab | nonsynonymous_S1267F |
| 4071 | C | T | | ORF1ab | nonsynonymous_T1269I |

| | | | | | |
|---|---|---|---|---|---|
| 4073 | C | T | | ORF1ab | nonsynonymous_L1270F |
| 4082 | G | T | | ORF1ab | nonsynonymous_D1273Y |
| 4084 | C | T | | ORF1ab | synonymous_D1273D |
| 4101 | T | A | | ORF1ab | stopgain_L1279X |
| 4106 | A | G | | ORF1ab | nonsynonymous_K1281E |
| 4113 | C | T | | ORF1ab | nonsynonymous_A1283V |
| 4139 | C | T | | ORF1ab | stopgain_Q1292X |
| 4145 | G | T | | ORF1ab | nonsynonymous_G1294C |
| 4158 | C | T | | ORF1ab | nonsynonymous_A1298V |
| 4162 | G | T | | ORF1ab | synonymous_V1299V |
| 4165 | T | C | | ORF1ab | synonymous_V1300V |
| 4173 | C | T | | ORF1ab | nonsynonymous_T1303I |
| 4175 | A | G | | ORF1ab | nonsynonymous_K1304E |
| 4180 | G | A | | ORF1ab | synonymous_K1305K |
| 4185 | G | T | | ORF1ab | nonsynonymous_G1307V |
| 4194 | C | T | | ORF1ab | nonsynonymous_T1310I |
| 4202 | C | T | | ORF1ab | synonymous_L1313L |
| 4207 | G | A | T | ORF1ab | synonymous_A1314A;<br>synonymous_A1314A |
| 4220 | A | T | | ORF1ab | stopgain_K1319X |
| 4230 | C | T | | ORF1ab | nonsynonymous_T1322I |
| 4255 | G | A | | ORF1ab | synonymous_P1330P |
| 4257 | G | T | | ORF1ab | nonsynonymous_G1331V |
| 4262 | G | C | | ORF1ab | nonsynonymous_G1333R |
| 4263 | G | T | | ORF1ab | nonsynonymous_G1333V |
| 4272 | G | C | | ORF1ab | nonsynonymous_G1336A |
| 4276 | C | A | | ORF1ab | stopgain_Y1337X |
| 4278 | C | T | | ORF1ab | nonsynonymous_T1338I |
| 4279 | T | C | | ORF1ab | synonymous_T1338T |
| 4281 | T | C | | ORF1ab | nonsynonymous_V1339A |
| 4286 | G | T | | ORF1ab | stopgain_E1341X |

| | | | | | |
|---|---|---|---|---|---|
| 4290 | C | A | | ORF1ab | nonsynonymous_A1342E |
| 4298 | G | T | | ORF1ab | nonsynonymous_V1345L |
| 4300 | G | T | | ORF1ab | synonymous_V1345V |
| 4301 | C | T | | ORF1ab | nonsynonymous_L1346F |
| 4309 | G | A | T | ORF1ab | synonymous_K1348K;<br>nonsynonymous_K1348N |
| 4316 | A | G | | ORF1ab | nonsynonymous_S1351G |
| 4321 | C | T | | ORF1ab | synonymous_A1352A |
| 4322 | T | C | | ORF1ab | nonsynonymous_F1353L |
| 4331 | C | T | | ORF1ab | synonymous_L1356L |
| 4339 | T | C | | ORF1ab | synonymous_S1358S |
| 4354 | G | T | | ORF1ab | nonsynonymous_E1363D |
| 4358 | C | T | | ORF1ab | stopgain_Q1365X |
| 4364 | A | T | | ORF1ab | nonsynonymous_I1367F |
| 4367 | C | T | | ORF1ab | nonsynonymous_L1368F |
| 4390 | G | T | | ORF1ab | nonsynonymous_L1375F |
| 4400 | C | T | | ORF1ab | nonsynonymous_L1379F |
| 4406 | C | T | | ORF1ab | nonsynonymous_H1381Y |
| 4412 | G | A | T | ORF1ab | nonsynonymous_E1383K;<br>stopgain_E1383X |
| 4415 | G | A | | ORF1ab | nonsynonymous_E1384K |
| 4419 | C | T | | ORF1ab | nonsynonymous_T1385I |
| 4421 | C | T | | ORF1ab | nonsynonymous_R1386C |
| 4423 | C | T | | ORF1ab | synonymous_R1386R |
| 4433 | C | T | | ORF1ab | nonsynonymous_P1390S |
| 4434 | C | T | | ORF1ab | nonsynonymous_P1390L |
| 4438 | C | T | | ORF1ab | synonymous_V1391V |
| 4441 | T | G | | ORF1ab | nonsynonymous_C1392W |
| 4444 | G | T | | ORF1ab | synonymous_V1393V |
| 4455 | C | T | | ORF1ab | nonsynonymous_A1397V |
| 4456 | C | T | | ORF1ab | synonymous_A1397A |

| | | | | | |
|---|---|---|---|---|---|
| 4464 | C | T | | ORF1ab | nonsynonymous_S1400L |
| 4467 | C | T | | ORF1ab | nonsynonymous_T1401I |
| 4475 | C | T | | ORF1ab | nonsynonymous_R1404C |
| 4485 | A | G | | ORF1ab | nonsynonymous_K1407R |
| 4490 | A | G | | ORF1ab | nonsynonymous_I1409V |
| 4499 | C | T | | ORF1ab | stopgain_Q1412X |
| 4500 | A | G | | ORF1ab | nonsynonymous_Q1412R |
| 4504 | G | A | | ORF1ab | synonymous_E1413E |
| 4505 | G | T | | ORF1ab | nonsynonymous_G1414C |
| 4510 | G | T | | ORF1ab | synonymous_V1415V |
| 4511 | G | T | | ORF1ab | nonsynonymous_V1416F |
| 4518 | A | G | | ORF1ab | nonsynonymous_Y1418C |
| 4520 | G | T | | ORF1ab | nonsynonymous_G1419C |
| 4521 | G | T | | ORF1ab | nonsynonymous_G1419V |
| 4527 | G | T | | ORF1ab | nonsynonymous_R1421I |
| 4534 | C | T | | ORF1ab | synonymous_Y1423Y |
| 4535 | T | C | | ORF1ab | nonsynonymous_F1424L |
| 4540 | C | T | | ORF1ab | synonymous_Y1425Y |
| 4542 | C | T | | ORF1ab | nonsynonymous_T1426I |
| 4550 | A | G | | ORF1ab | nonsynonymous_T1429A |
| 4551 | C | T | | ORF1ab | nonsynonymous_T1429I |
| 4554 | C | T | | ORF1ab | nonsynonymous_T1430I |
| 4555 | T | C | | ORF1ab | synonymous_T1430T |
| 4560 | C | T | | ORF1ab | nonsynonymous_A1432V |
| 4563 | C | T | | ORF1ab | nonsynonymous_S1433L |
| 4585 | T | C | | ORF1ab | synonymous_D1440D |
| 4586 | C | T | | ORF1ab | synonymous_L1441L |
| 4587 | T | A | | ORF1ab | nonsynonymous_L1441Q |
| 4596 | C | T | | ORF1ab | nonsynonymous_T1444I |
| 4597 | T | C | | ORF1ab | synonymous_T1444T |
| 4598 | C | T | | ORF1ab | nonsynonymous_L1445F |
| 4618 | C | T | | ORF1ab | synonymous_G1451G |

| | | | | | |
|---|---|---|---|---|---|
| 4622 | G | T | | ORF1ab | nonsynonymous_V1453L |
| 4628 | C | T | | ORF1ab | nonsynonymous_H1455Y |
| 4629 | A | T | | ORF1ab | nonsynonymous_H1455L |
| 4631 | G | T | | ORF1ab | nonsynonymous_G1456C |
| 4633 | C | T | | ORF1ab | synonymous_G1456G |
| 4646 | G | T | | ORF1ab | stopgain_E1461X |
| 4650 | C | T | | ORF1ab | nonsynonymous_A1462V |
| 4655 | C | T | | ORF1ab | nonsynonymous_R1464W |
| 4656 | G | T | | ORF1ab | nonsynonymous_R1464L |
| 4657 | G | A | | ORF1ab | synonymous_R1464R |
| 4668 | C | T | | ORF1ab | nonsynonymous_S1468F |
| 4675 | A | T | G | ORF1ab | nonsynonymous_K1470N;<br>synonymous_K1470K |
| 4676 | G | C | | ORF1ab | nonsynonymous_V1471L |
| 4678 | G | A | | ORF1ab | synonymous_V1471V |
| 4682 | G | A | T | ORF1ab | nonsynonymous_A1473T;<br>nonsynonymous_A1473S |
| 4683 | C | T | | ORF1ab | nonsynonymous_A1473V |
| 4698 | C | T | | ORF1ab | nonsynonymous_S1478F |
| 4700 | T | C | | ORF1ab | nonsynonymous_S1479P |
| 4703 | C | T | | ORF1ab | nonsynonymous_P1480S |
| 4710 | C | T | | ORF1ab | nonsynonymous_A1482V |
| 4721 | T | C | | ORF1ab | nonsynonymous_Y1486H |
| 4728 | G | A | T | ORF1ab | nonsynonymous_G1488D;<br>nonsynonymous_G1488V |
| 4733 | C | A | | ORF1ab | nonsynonymous_L1490I |
| 4738 | T | C | | ORF1ab | synonymous_T1491T |
| 4743 | C | T | | ORF1ab | nonsynonymous_S1493F |
| 4746 | C | T | | ORF1ab | nonsynonymous_S1494F |
| 4752 | C | T | | ORF1ab | nonsynonymous_T1496I |
| 4754 | C | T | | ORF1ab | nonsynonymous_P1497S |

| | | | | | |
|---|---|---|---|---|---|
| 4755 | C | T | | ORF1ab | nonsynonymous_P1497L |
| 4763 | C | T | | ORF1ab | nonsynonymous_H1500Y |
| 4776 | C | T | | ORF1ab | nonsynonymous_T1504I |
| 4777 | C | T | | ORF1ab | synonymous_T1504T |
| 4791 | G | T | | ORF1ab | nonsynonymous_G1509V |
| 4795 | C | T | | ORF1ab | synonymous_S1510S |
| 4809 | C | T | | ORF1ab | nonsynonymous_S1515F |
| 4824 | C | T | | ORF1ab | nonsynonymous_S1520F |
| 4827 | C | T | | ORF1ab | nonsynonymous_T1521I |
| 4832 | C | T | | ORF1ab | synonymous_L1523L |
| 4841 | G | A | | ORF1ab | nonsynonymous_E1526K |
| 4851 | A | C | | ORF1ab | nonsynonymous_K1529T |
| 4854 | G | T | | ORF1ab | nonsynonymous_R1530I |
| 4857 | G | T | | ORF1ab | nonsynonymous_G1531V |
| 4866 | G | T | | ORF1ab | nonsynonymous_S1534I |
| 4886 | C | T | | ORF1ab | nonsynonymous_P1541S |
| 4890 | C | T | | ORF1ab | nonsynonymous_T1542I |
| 4893 | C | T | | ORF1ab | nonsynonymous_T1543I |
| 4900 | C | T | | ORF1ab | synonymous_H1545H |
| 4901 | C | T | | ORF1ab | synonymous_L1546L |
| 4902 | T | C | | ORF1ab | nonsynonymous_L1546P |
| 4904 | G | T | | ORF1ab | nonsynonymous_D1547Y |
| 4908 | G | T | | ORF1ab | nonsynonymous_G1548V |
| 4913 | G | T | | ORF1ab | nonsynonymous_V1550F |
| 4918 | C | T | | ORF1ab | synonymous_I1551I |
| 4920 | C | T | | ORF1ab | nonsynonymous_T1552I |
| 4921 | C | T | | ORF1ab | synonymous_T1552T |
| 4943 | C | T | | ORF1ab | nonsynonymous_L1560F |
| 4951 | G | A | | ORF1ab | synonymous_L1562L |
| 4952 | A | C | | ORF1ab | synonymous_R1563R |
| 4960 | G | A | | ORF1ab | synonymous_V1565V |
| 4963 | G | T | | ORF1ab | nonsynonymous_R1566S |

| | | | | | |
|---|---|---|---|---|---|
| 4983 | C | T | | ORF1ab | nonsynonymous_T1573I |
| 4987 | A | G | | ORF1ab | synonymous_V1574V |
| 4988 | G | T | | ORF1ab | nonsynonymous_D1575Y |
| 4993 | C | T | | ORF1ab | synonymous_N1576N |
| 5000 | C | A | | ORF1ab | nonsynonymous_L1579I |
| 5002 | C | T | | ORF1ab | synonymous_L1579L |
| 5007 | C | T | | ORF1ab | nonsynonymous_T1581M |
| 5011 | A | G | | ORF1ab | synonymous_Q1582Q |
| 5012 | G | T | | ORF1ab | nonsynonymous_V1583F |
| 5013 | T | C | | ORF1ab | nonsynonymous_V1583A |
| 5015 | G | T | | ORF1ab | nonsynonymous_V1584L |
| 5018 | G | T | | ORF1ab | nonsynonymous_D1585Y |
| 5020 | C | T | | ORF1ab | synonymous_D1585D |
| 5023 | G | T | | ORF1ab | nonsynonymous_M1586I |
| 5025 | C | T | | ORF1ab | nonsynonymous_S1587L |
| 5026 | A | T | G | ORF1ab | synonymous_S1587S; synonymous_S1587S |
| 5035 | T | C | | ORF1ab | synonymous_Y1590Y |
| 5036 | G | T | | ORF1ab | stopgain_G1591X |
| 5039 | C | A | | ORF1ab | nonsynonymous_Q1592K |
| 5055 | C | T | | ORF1ab | nonsynonymous_T1597I |
| 5062 | G | A | | ORF1ab | synonymous_L1599L |
| 5075 | G | A | | ORF1ab | nonsynonymous_V1604I |
| 5084 | A | G | | ORF1ab | nonsynonymous_I1607V |
| 5100 | C | T | | ORF1ab | nonsynonymous_S1612L |
| 5105 | G | T | | ORF1ab | stopgain_E1614X |
| 5115 | C | T | | ORF1ab | nonsynonymous_T1617I |
| 5116 | A | T | | ORF1ab | synonymous_T1617T |
| 5123 | G | T | | ORF1ab | nonsynonymous_V1620F |
| 5129 | C | T | | ORF1ab | nonsynonymous_P1622S |
| 5130 | C | T | | ORF1ab | nonsynonymous_P1622L |

| | | | | | |
|---|---|---|---|---|---|
| 5136 | A | T | | ORF1ab | nonsynonymous_D1624V |
| 5138 | G | T | | ORF1ab | nonsynonymous_D1625Y |
| 5140 | C | T | | ORF1ab | synonymous_D1625D |
| 5142 | C | T | | ORF1ab | nonsynonymous_T1626I |
| 5144 | C | T | | ORF1ab | synonymous_L1627L |
| 5147 | C | T | | ORF1ab | nonsynonymous_R1628C |
| 5155 | G | T | | ORF1ab | nonsynonymous_E1630D |
| 5156 | G | C | | ORF1ab | nonsynonymous_A1631P |
| 5157 | C | T | | ORF1ab | nonsynonymous_A1631V |
| 5167 | C | T | | ORF1ab | synonymous_Y1634Y |
| 5170 | C | T | | ORF1ab | synonymous_Y1635Y |
| 5173 | C | T | | ORF1ab | synonymous_H1636H |
| 5183 | C | T | | ORF1ab | nonsynonymous_P1640S |
| 5184 | C | T | | ORF1ab | nonsynonymous_P1640L |
| 5189 | T | C | | ORF1ab | nonsynonymous_F1642L |
| 5192 | C | T | | ORF1ab | synonymous_L1643L |
| 5193 | T | G | | ORF1ab | nonsynonymous_L1643R |
| 5196 | G | T | | ORF1ab | nonsynonymous_G1644V |
| 5200 | G | C | T | ORF1ab | nonsynonymous_R1645S; nonsynonymous_R1645S |
| 5206 | G | A | T | ORF1ab | nonsynonymous_M1647I; nonsynonymous_M1647I |
| 5211 | C | T | | ORF1ab | nonsynonymous_A1649V |
| 5212 | A | G | | ORF1ab | synonymous_A1649A |
| 5219 | C | T | | ORF1ab | nonsynonymous_H1652Y |
| 5221 | C | T | | ORF1ab | synonymous_H1652H |
| 5223 | C | T | | ORF1ab | nonsynonymous_T1653I |
| 5229 | A | T | | ORF1ab | nonsynonymous_K1655M |
| 5230 | G | T | | ORF1ab | nonsynonymous_K1655N |
| 5231 | T | C | | ORF1ab | nonsynonymous_W1656R |
| 5232 | G | A | | ORF1ab | stopgain_W1656X |

| | | | | | |
|---|---|---|---|---|---|
| 5233 | G | A | T | ORF1ab | stopgain_W1656X;<br>nonsynonymous_W1656C |
| 5239 | C | T | | ORF1ab | synonymous_Y1658Y |
| 5241 | C | T | | ORF1ab | nonsynonymous_P1659L |
| 5253 | G | T | | ORF1ab | nonsynonymous_G1663V |
| 5259 | C | A | | ORF1ab | nonsynonymous_T1665N |
| 5284 | C | T | | ORF1ab | synonymous_N1673N |
| 5294 | G | T | | ORF1ab | nonsynonymous_A1677S |
| 5301 | C | A | | ORF1ab | nonsynonymous_A1679E |
| 5312 | C | T | | ORF1ab | nonsynonymous_L1683F |
| 5340 | C | T | | ORF1ab | nonsynonymous_P1692L |
| 5348 | C | T | | ORF1ab | synonymous_L1695L |
| 5350 | A | G | | ORF1ab | synonymous_L1695L |
| 5351 | C | T | | ORF1ab | stopgain_Q1696X |
| 5356 | T | C | | ORF1ab | synonymous_D1697D |
| 5365 | C | T | | ORF1ab | synonymous_Y1700Y |
| 5375 | G | T | | ORF1ab | nonsynonymous_A1704S |
| 5378 | G | T | | ORF1ab | nonsynonymous_G1705C |
| 5385 | C | T | | ORF1ab | nonsynonymous_A1707V |
| 5387 | G | T | | ORF1ab | nonsynonymous_A1708S |
| 5388 | C | T | | ORF1ab | nonsynonymous_A1708V |
| 5391 | A | G | | ORF1ab | nonsynonymous_N1709S |
| 5392 | C | T | | ORF1ab | synonymous_N1709N |
| 5406 | T | C | | ORF1ab | nonsynonymous_I1714T |
| 5407 | C | T | | ORF1ab | synonymous_I1714I |
| 5410 | A | G | | ORF1ab | synonymous_L1715L |
| 5412 | C | G | | ORF1ab | nonsynonymous_A1716G |
| 5416 | C | T | | ORF1ab | synonymous_Y1717Y |
| 5423 | A | T | | ORF1ab | stopgain_K1720X |
| 5425 | G | T | | ORF1ab | nonsynonymous_K1720N |
| 5432 | G | A | | ORF1ab | nonsynonymous_G1723S |

| | | | | | |
|---|---|---|---|---|---|
| 5433 | G | T | | ORF1ab | nonsynonymous_G1723V |
| 5436 | A | G | | ORF1ab | nonsynonymous_E1724G |
| 5462 | A | C | G | ORF1ab | nonsynonymous_S1733R; nonsynonymous_S1733G |
| 5466 | A | G | | ORF1ab | nonsynonymous_Y1734C |
| 5467 | C | T | | ORF1ab | synonymous_Y1734Y |
| 5477 | C | T | | ORF1ab | nonsynonymous_H1738Y |
| 5481 | C | T | G | ORF1ab | nonsynonymous_A1739V; nonsynonymous_A1739G |
| 5482 | C | T | | ORF1ab | synonymous_A1739A |
| 5493 | C | T | | ORF1ab | nonsynonymous_S1743F |
| 5497 | C | T | | ORF1ab | synonymous_C1744C |
| 5505 | T | G | | ORF1ab | nonsynonymous_V1747G |
| 5506 | C | T | | ORF1ab | synonymous_V1747V |
| 5508 | T | C | | ORF1ab | nonsynonymous_L1748S |
| 5512 | C | T | | ORF1ab | synonymous_N1749N |
| 5526 | C | T | | ORF1ab | nonsynonymous_T1754I |
| 5534 | C | T | | ORF1ab | stopgain_Q1757X |
| 5537 | C | T | | ORF1ab | stopgain_Q1758X |
| 5538 | A | T | | ORF1ab | nonsynonymous_Q1758L |
| 5542 | G | T | | ORF1ab | nonsynonymous_Q1759H |
| 5544 | C | T | | ORF1ab | nonsynonymous_T1760I |
| 5548 | C | T | | ORF1ab | synonymous_T1761T |
| 5551 | T | C | | ORF1ab | synonymous_L1762L |
| 5555 | G | A | T | ORF1ab | nonsynonymous_G1764S; nonsynonymous_G1764C |
| 5558 | G | T | | ORF1ab | nonsynonymous_V1765L |
| 5561 | G | T | | ORF1ab | stopgain_E1766X |
| 5567 | G | T | | ORF1ab | nonsynonymous_V1768F |
| 5570 | A | G | | ORF1ab | nonsynonymous_M1769V |
| 5572 | G | T | | ORF1ab | nonsynonymous_M1769I |

| | | | | | |
|---|---|---|---|---|---|
| 5586 | T | C | | ORF1ab | nonsynonymous_L1774P |
| 5597 | C | T | | ORF1ab | stopgain_Q1778X |
| 5599 | A | G | | ORF1ab | synonymous_Q1778Q |
| 5600 | T | C | | ORF1ab | nonsynonymous_F1779L |
| 5606 | A | G | | ORF1ab | nonsynonymous_K1781E |
| 5612 | G | T | | ORF1ab | nonsynonymous_V1783F |
| 5617 | G | T | | ORF1ab | nonsynonymous_Q1784H |
| 5618 | A | T | | ORF1ab | nonsynonymous_I1785L |
| 5622 | C | T | | ORF1ab | nonsynonymous_P1786L |
| 5628 | C | T | | ORF1ab | nonsynonymous_T1788M |
| 5632 | T | C | | ORF1ab | synonymous_C1789C |
| 5634 | G | T | | ORF1ab | nonsynonymous_G1790V |
| 5642 | G | C | T | ORF1ab | nonsynonymous_A1793P; nonsynonymous_A1793S |
| 5643 | C | T | | ORF1ab | nonsynonymous_A1793V |
| 5645 | A | C | | ORF1ab | nonsynonymous_T1794P |
| 5646 | C | T | | ORF1ab | nonsynonymous_T1794I |
| 5654 | C | T | | ORF1ab | synonymous_L1797L |
| 5666 | G | A | C | ORF1ab | nonsynonymous_E1801K; nonsynonymous_E1801Q |
| 5668 | G | A | | ORF1ab | synonymous_E1801E |
| 5670 | C | T | | ORF1ab | nonsynonymous_S1802L |
| 5674 | T | C | | ORF1ab | synonymous_P1803P |
| 5677 | T | C | | ORF1ab | synonymous_F1804F |
| 5686 | G | C | | ORF1ab | nonsynonymous_M1807I |
| 5688 | C | T | | ORF1ab | nonsynonymous_S1808L |
| 5691 | C | T | | ORF1ab | nonsynonymous_A1809V |
| 5693 | C | T | | ORF1ab | nonsynonymous_P1810S |
| 5697 | C | T | | ORF1ab | nonsynonymous_P1811L |
| 5700 | C | A | | ORF1ab | nonsynonymous_A1812D |
| 5709 | A | T | | ORF1ab | nonsynonymous_E1815V |

| | | | | | |
|---|---|---|---|---|---|
| 5715 | A | G | | ORF1ab | nonsynonymous_K1817R |
| 5717 | C | T | | ORF1ab | nonsynonymous_H1818Y |
| 5721 | G | T | | ORF1ab | nonsynonymous_G1819V |
| 5730 | C | T | | ORF1ab | nonsynonymous_T1822I |
| 5734 | T | C | | ORF1ab | synonymous_C1823C |
| 5735 | G | C | | ORF1ab | nonsynonymous_A1824P |
| 5742 | A | T | | ORF1ab | nonsynonymous_E1826V |
| 5743 | G | C | | ORF1ab | nonsynonymous_E1826D |
| 5744 | T | C | | ORF1ab | nonsynonymous_Y1827H |
| 5746 | C | T | | ORF1ab | synonymous_Y1827Y |
| 5750 | G | A | T | ORF1ab | nonsynonymous_G1829S; nonsynonymous_G1829C |
| 5765 | G | A | | ORF1ab | nonsynonymous_G1834S |
| 5766 | G | C | | ORF1ab | nonsynonymous_G1834A |
| 5777 | C | T | | ORF1ab | nonsynonymous_H1838Y |
| 5784 | C | T | | ORF1ab | nonsynonymous_T1840I |
| 5792 | G | A | | ORF1ab | nonsynonymous_E1843K |
| 5796 | C | T | | ORF1ab | nonsynonymous_T1844I |
| 5810 | G | T | | ORF1ab | nonsynonymous_D1849Y |
| 5812 | C | T | | ORF1ab | synonymous_D1849D |
| 5813 | G | A | | ORF1ab | nonsynonymous_G1850S |
| 5817 | C | T | | ORF1ab | nonsynonymous_A1851V |
| 5820 | T | C | | ORF1ab | nonsynonymous_L1852S |
| 5821 | A | G | | ORF1ab | synonymous_L1852L |
| 5822 | C | T | | ORF1ab | nonsynonymous_L1853F |
| 5825 | A | G | | ORF1ab | nonsynonymous_T1854A |
| 5826 | C | T | | ORF1ab | nonsynonymous_T1854I |
| 5830 | G | T | | ORF1ab | nonsynonymous_K1855N |
| 5833 | C | T | | ORF1ab | synonymous_S1856S |
| 5834 | T | C | | ORF1ab | nonsynonymous_S1857P |
| 5835 | C | T | | ORF1ab | nonsynonymous_S1857L |

| | | | | | |
|---|---|---|---|---|---|
| 5842 | C | A | T | ORF1ab | stopgain_Y1859X;<br>synonymous_Y1859Y |
| 5847 | G | T | | ORF1ab | nonsynonymous_G1861V |
| 5849 | C | T | | ORF1ab | nonsynonymous_P1862S |
| 5850 | C | T | | ORF1ab | nonsynonymous_P1862L |
| 5856 | C | T | | ORF1ab | nonsynonymous_T1864M |
| 5869 | C | T | | ORF1ab | synonymous_Y1868Y |
| 5870 | A | T | | ORF1ab | stopgain_K1869X |
| 5878 | C | T | | ORF1ab | synonymous_N1871N |
| 5880 | G | T | | ORF1ab | nonsynonymous_S1872I |
| 5884 | C | T | | ORF1ab | synonymous_Y1873Y |
| 5907 | C | T | | ORF1ab | nonsynonymous_T1881I |
| 5945 | C | T | | ORF1ab | nonsynonymous_P1894S |
| 5956 | C | T | | ORF1ab | synonymous_D1897D |
| 5960 | T | C | | ORF1ab | nonsynonymous_Y1899H |
| 5964 | A | T | | ORF1ab | nonsynonymous_Y1900F |
| 5969 | A | C | | ORF1ab | nonsynonymous_K1902Q |
| 5977 | T | C | | ORF1ab | synonymous_N1904N |
| 5986 | C | T | | ORF1ab | synonymous_F1907F |
| 5988 | C | G | | ORF1ab | nonsynonymous_T1908R |
| 5992 | G | C | | ORF1ab | nonsynonymous_E1909D |
| 5997 | C | T | | ORF1ab | nonsynonymous_P1911L |
| 6011 | C | T | | ORF1ab | nonsynonymous_P1916S |
| 6012 | C | A | | ORF1ab | nonsynonymous_P1916Q |
| 6015 | A | C | | ORF1ab | nonsynonymous_N1917T |
| 6016 | C | T | | ORF1ab | synonymous_N1917N |
| 6026 | C | T | | ORF1ab | nonsynonymous_P1921S |
| 6027 | C | T | | ORF1ab | nonsynonymous_P1921L |
| 6035 | A | G | | ORF1ab | nonsynonymous_S1924G |
| 6037 | C | T | | ORF1ab | synonymous_S1924S |
| 6038 | T | G | | ORF1ab | nonsynonymous_F1925V |

| | | | | | |
|---|---|---|---|---|---|
| 6040 | C | T | G | ORF1ab | synonymous_F1925F;<br>nonsynonymous_F1925L |
| 6063 | A | G | | ORF1ab | nonsynonymous_D1933G |
| 6064 | T | C | | ORF1ab | synonymous_D1933D |
| 6075 | T | C | | ORF1ab | nonsynonymous_F1937S |
| 6078 | C | T | | ORF1ab | nonsynonymous_A1938V |
| 6079 | T | C | | ORF1ab | synonymous_A1938A |
| 6081 | A | G | | ORF1ab | nonsynonymous_D1939G |
| 6094 | G | T | | ORF1ab | nonsynonymous_Q1943H |
| 6097 | A | G | | ORF1ab | synonymous_L1944L |
| 6099 | C | T | | ORF1ab | nonsynonymous_T1945I |
| 6101 | G | A | | ORF1ab | nonsynonymous_G1946S |
| 6105 | A | T | | ORF1ab | nonsynonymous_Y1947F |
| 6109 | G | A | | ORF1ab | synonymous_K1948K |
| 6110 | A | T | | ORF1ab | stopgain_K1949X |
| 6117 | C | T | | ORF1ab | nonsynonymous_A1951V |
| 6127 | G | A | | ORF1ab | synonymous_E1954E |
| 6128 | C | T | | ORF1ab | nonsynonymous_L1955F |
| 6133 | A | G | | ORF1ab | synonymous_K1956K |
| 6138 | C | T | | ORF1ab | nonsynonymous_T1958I |
| 6145 | C | T | | ORF1ab | synonymous_F1960F |
| 6147 | C | T | | ORF1ab | nonsynonymous_P1961L |
| 6151 | C | T | | ORF1ab | synonymous_D1962D |
| 6163 | T | C | G | ORF1ab | synonymous_D1966D;<br>nonsynonymous_D1966E |
| 6170 | G | T | | ORF1ab | nonsynonymous_A1969S |
| 6172 | T | C | | ORF1ab | synonymous_A1969A |
| 6183 | A | G | | ORF1ab | nonsynonymous_K1973R |
| 6187 | C | T | | ORF1ab | synonymous_H1974H |
| 6195 | C | T | | ORF1ab | nonsynonymous_P1977L |
| 6198 | C | T | | ORF1ab | nonsynonymous_S1978F |

| | | | | | |
|---|---|---|---|---|---|
| 6205 | G | A | | ORF1ab | synonymous_K1980K |
| 6213 | C | T | | ORF1ab | nonsynonymous_A1983V |
| 6224 | C | T | | ORF1ab | nonsynonymous_H1987Y |
| 6228 | A | G | | ORF1ab | nonsynonymous_K1988R |
| 6230 | C | T | | ORF1ab | nonsynonymous_P1989S |
| 6231 | C | T | | ORF1ab | nonsynonymous_P1989L |
| 6235 | T | G | | ORF1ab | nonsynonymous_I1990M |
| 6236 | G | A | T | ORF1ab | nonsynonymous_V1991I; nonsynonymous_V1991F |
| 6242 | C | T | | ORF1ab | nonsynonymous_H1993Y |
| 6245 | G | T | | ORF1ab | nonsynonymous_V1994F |
| 6250 | C | T | | ORF1ab | synonymous_N1995N |
| 6251 | A | G | | ORF1ab | nonsynonymous_N1996D |
| 6260 | A | G | | ORF1ab | nonsynonymous_N1999D |
| 6267 | C | T | | ORF1ab | nonsynonymous_A2001V |
| 6282 | A | G | | ORF1ab | nonsynonymous_N2006S |
| 6285 | C | T | | ORF1ab | nonsynonymous_T2007I |
| 6286 | C | T | | ORF1ab | synonymous_T2007T |
| 6294 | T | C | | ORF1ab | nonsynonymous_I2010T |
| 6303 | T | C | | ORF1ab | nonsynonymous_L2013P |
| 6307 | G | A | | ORF1ab | stopgain_W2014X |
| 6308 | A | C | | ORF1ab | nonsynonymous_S2015R |
| 6310 | C | A | T | ORF1ab | nonsynonymous_S2015R; synonymous_S2015S |
| 6312 | C | A | | ORF1ab | nonsynonymous_T2016K |
| 6317 | C | T | | ORF1ab | nonsynonymous_P2018S |
| 6318 | C | T | | ORF1ab | nonsynonymous_P2018L |
| 6327 | C | T | | ORF1ab | nonsynonymous_T2021I |
| 6335 | T | C | | ORF1ab | nonsynonymous_S2024P |
| 6336 | C | T | | ORF1ab | nonsynonymous_S2024L |
| 6344 | G | T | | ORF1ab | nonsynonymous_V2027L |

| | | | | | |
|---|---|---|---|---|---|
| 6350 | A | G | | ORF1ab | nonsynonymous_K2029E |
| 6352 | G | T | | ORF1ab | nonsynonymous_K2029N |
| 6354 | C | T | | ORF1ab | nonsynonymous_S2030L |
| 6359 | G | T | | ORF1ab | nonsynonymous_D2032Y |
| 6362 | G | T | | ORF1ab | nonsynonymous_A2033S |
| 6363 | C | T | | ORF1ab | nonsynonymous_A2033V |
| 6364 | G | A | | ORF1ab | synonymous_A2033A |
| 6371 | A | G | | ORF1ab | nonsynonymous_M2036V |
| 6384 | C | T | | ORF1ab | nonsynonymous_A2040V |
| 6385 | C | T | | ORF1ab | synonymous_A2040A |
| 6388 | C | T | | ORF1ab | synonymous_C2041C |
| 6393 | A | T | G | ORF1ab | nonsynonymous_D2043V; nonsynonymous_D2043G |
| 6395 | C | T | | ORF1ab | synonymous_L2044L |
| 6396 | T | A | | ORF1ab | nonsynonymous_L2044Q |
| 6401 | C | T | | ORF1ab | nonsynonymous_P2046S |
| 6402 | C | T | | ORF1ab | nonsynonymous_P2046L |
| 6406 | C | T | | ORF1ab | synonymous_V2047V |
| 6410 | G | A | | ORF1ab | nonsynonymous_E2049K |
| 6427 | T | C | | ORF1ab | synonymous_N2054N |
| 6428 | C | T | | ORF1ab | nonsynonymous_P2055S |
| 6429 | C | T | | ORF1ab | nonsynonymous_P2055L |
| 6432 | C | T | | ORF1ab | nonsynonymous_T2056I |
| 6437 | C | T | | ORF1ab | stopgain_Q2058X |
| 6438 | A | G | | ORF1ab | nonsynonymous_Q2058R |
| 6439 | G | A | | ORF1ab | synonymous_Q2058Q |
| 6440 | A | G | | ORF1ab | nonsynonymous_K2059E |
| 6441 | A | G | | ORF1ab | nonsynonymous_K2059R |
| 6446 | G | A | T | ORF1ab | nonsynonymous_V2061I; nonsynonymous_V2061F |
| 6449 | C | T | | ORF1ab | nonsynonymous_L2062F |

| | | | | | |
|---|---|---|---|---|---|
| 6463 | G | T | | ORF1ab | synonymous_V2066V |
| 6464 | A | G | | ORF1ab | nonsynonymous_K2067E |
| 6468 | C | T | | ORF1ab | nonsynonymous_T2068I |
| 6483 | G | T | | ORF1ab | nonsynonymous_G2073V |
| 6487 | C | T | | ORF1ab | synonymous_D2074D |
| 6493 | A | G | | ORF1ab | nonsynonymous_I2076M |
| 6501 | C | T | | ORF1ab | nonsynonymous_P2079L |
| 6504 | C | T | | ORF1ab | nonsynonymous_A2080V |
| 6509 | A | T | | ORF1ab | nonsynonymous_N2082Y |
| 6512 | A | C | | ORF1ab | nonsynonymous_S2083R |
| 6513 | G | T | | ORF1ab | nonsynonymous_S2083I |
| 6525 | C | T | | ORF1ab | nonsynonymous_T2087I |
| 6532 | G | T | | ORF1ab | nonsynonymous_E2089D |
| 6537 | G | T | | ORF1ab | nonsynonymous_G2091V |
| 6543 | C | T | | ORF1ab | nonsynonymous_T2093I |
| 6553 | G | C | | ORF1ab | nonsynonymous_M2096I |
| 6554 | G | A | | ORF1ab | nonsynonymous_A2097T |
| 6555 | C | T | | ORF1ab | nonsynonymous_A2097V |
| 6573 | C | T | | ORF1ab | nonsynonymous_S2103F |
| 6576 | G | T | | ORF1ab | nonsynonymous_S2104I |
| 6578 | C | T | | ORF1ab | nonsynonymous_L2105F |
| 6582 | C | T | | ORF1ab | nonsynonymous_T2106I |
| 6589 | G | A | | ORF1ab | synonymous_K2108K |
| 6608 | A | C | | ORF1ab | synonymous_R2115R |
| 6616 | A | G | | ORF1ab | synonymous_L2117L |
| 6627 | C | T | | ORF1ab | nonsynonymous_T2121I |
| 6632 | G | T | | ORF1ab | nonsynonymous_A2123S |
| 6633 | C | T | | ORF1ab | nonsynonymous_A2123V |
| 6637 | T | C | | ORF1ab | synonymous_T2124T |
| 6638 | C | T | | ORF1ab | nonsynonymous_H2125Y |
| 6648 | C | T | | ORF1ab | nonsynonymous_A2128V |
| 6653 | G | A | | ORF1ab | nonsynonymous_V2130I |

| | | | | | |
|---|---|---|---|---|---|
| 6669 | G | T | | ORF1ab | nonsynonymous_W2135L |
| 6673 | T | C | | ORF1ab | synonymous_D2136D |
| 6681 | C | T | | ORF1ab | nonsynonymous_A2139V |
| 6690 | C | T | | ORF1ab | nonsynonymous_A2142V |
| 6695 | C | A | | ORF1ab | nonsynonymous_P2144T |
| 6696 | C | T | | ORF1ab | nonsynonymous_P2144L |
| 6701 | C | T | | ORF1ab | nonsynonymous_L2146F |
| 6702 | T | C | | ORF1ab | nonsynonymous_L2146P |
| 6705 | A | G | | ORF1ab | nonsynonymous_N2147S |
| 6706 | C | T | | ORF1ab | synonymous_N2147N |
| 6711 | T | C | | ORF1ab | nonsynonymous_V2149A |
| 6712 | T | C | | ORF1ab | synonymous_V2149V |
| 6720 | C | T | | ORF1ab | nonsynonymous_T2152I |
| 6723 | C | T | | ORF1ab | nonsynonymous_T2153I |
| 6726 | C | T | | ORF1ab | nonsynonymous_T2154I |
| 6730 | C | T | | ORF1ab | synonymous_N2155N |
| 6733 | A | T | | ORF1ab | synonymous_I2156I |
| 6738 | C | T | | ORF1ab | nonsynonymous_T2158I |
| 6747 | T | A | | ORF1ab | stopgain_L2161X |
| 6749 | A | G | | ORF1ab | nonsynonymous_N2162D |
| 6752 | C | T | | ORF1ab | nonsynonymous_R2163C |
| 6755 | G | T | | ORF1ab | nonsynonymous_V2164F |
| 6757 | T | C | | ORF1ab | synonymous_V2164V |
| 6762 | C | T | | ORF1ab | nonsynonymous_T2166I |
| 6773 | C | T | | ORF1ab | nonsynonymous_P2170S |
| 6774 | C | T | | ORF1ab | nonsynonymous_P2170L |
| 6781 | C | T | | ORF1ab | synonymous_F2172F |
| 6786 | C | T | | ORF1ab | nonsynonymous_T2174I |
| 6804 | G | T | | ORF1ab | nonsynonymous_C2180F |
| 6808 | T | C | | ORF1ab | synonymous_T2181T |
| 6810 | T | C | | ORF1ab | nonsynonymous_F2182S |
| 6813 | C | T | | ORF1ab | nonsynonymous_T2183I |

| | | | | | |
|---|---|---|---|---|---|
| 6819 | G | T | | ORF1ab | nonsynonymous_S2185I |
| 6825 | A | C | | ORF1ab | nonsynonymous_N2187T |
| 6842 | T | C | | ORF1ab | nonsynonymous_S2193P |
| 6843 | C | T | | ORF1ab | nonsynonymous_S2193F |
| 6852 | C | T | | ORF1ab | nonsynonymous_T2196I |
| 6855 | C | T | | ORF1ab | nonsynonymous_T2197I |
| 6872 | G | A | | ORF1ab | nonsynonymous_V2203I |
| 6883 | C | T | | ORF1ab | synonymous_V2206V |
| 6893 | T | C | | ORF1ab | nonsynonymous_C2210R |
| 6895 | T | C | | ORF1ab | synonymous_C2210C |
| 6896 | C | T | | ORF1ab | synonymous_L2211L |
| 6902 | G | A | | ORF1ab | nonsynonymous_A2213T |
| 6912 | A | T | | ORF1ab | nonsynonymous_N2216I |
| 6924 | C | T | | ORF1ab | nonsynonymous_S2220L |
| 6926 | C | T | | ORF1ab | nonsynonymous_P2221S |
| 6927 | C | T | | ORF1ab | nonsynonymous_P2221L |
| 6936 | C | A | T | ORF1ab | nonsynonymous_S2224Y;<br>nonsynonymous_S2224F |
| 6937 | T | A | | ORF1ab | synonymous_S2224S |
| 6941 | C | T | | ORF1ab | synonymous_L2226L |
| 6944 | A | G | | ORF1ab | nonsynonymous_I2227V |
| 6958 | T | C | | ORF1ab | synonymous_I2231I |
| 6971 | T | C | | ORF1ab | synonymous_L2236L |
| 6975 | G | T | | ORF1ab | nonsynonymous_S2237I |
| 6976 | T | G | | ORF1ab | nonsynonymous_S2237R |
| 6982 | C | T | | ORF1ab | synonymous_C2239C |
| 6987 | G | T | | ORF1ab | nonsynonymous_G2241V |
| 6989 | T | C | | ORF1ab | nonsynonymous_S2242P |
| 6990 | C | T | | ORF1ab | nonsynonymous_S2242F |
| 6993 | T | A | | ORF1ab | stopgain_L2243X |
| 7005 | C | T | | ORF1ab | nonsynonymous_T2247I |

| | | | | | |
|---|---|---|---|---|---|
| 7008 | C | T | | ORF1ab | nonsynonymous_A2248V |
| 7010 | G | C | T | ORF1ab | nonsynonymous_A2249P; nonsynonymous_A2249S |
| 7011 | C | T | | ORF1ab | nonsynonymous_A2249V |
| 7017 | G | C | T | ORF1ab | nonsynonymous_G2251A; nonsynonymous_G2251V |
| 7024 | A | T | | ORF1ab | nonsynonymous_L2253F |
| 7029 | C | T | | ORF1ab | nonsynonymous_S2255F |
| 7038 | G | T | | ORF1ab | nonsynonymous_G2258V |
| 7042 | G | T | | ORF1ab | nonsynonymous_M2259I |
| 7066 | A | G | | ORF1ab | synonymous_R2267R |
| 7067 | G | T | | ORF1ab | stopgain_E2268X |
| 7069 | A | G | | ORF1ab | synonymous_E2268E |
| 7070 | G | A | | ORF1ab | nonsynonymous_G2269S |
| 7071 | G | A | T | ORF1ab | nonsynonymous_G2269D; nonsynonymous_G2269V |
| 7072 | C | T | | ORF1ab | synonymous_G2269G |
| 7081 | C | T | | ORF1ab | synonymous_N2272N |
| 7083 | C | T | | ORF1ab | nonsynonymous_S2273F |
| 7086 | C | T | | ORF1ab | nonsynonymous_T2274I |
| 7089 | A | T | | ORF1ab | nonsynonymous_N2275I |
| 7093 | C | T | | ORF1ab | synonymous_V2276V |
| 7108 | C | T | | ORF1ab | synonymous_Y2281Y |
| 7113 | C | T | | ORF1ab | nonsynonymous_T2283I |
| 7119 | C | T | | ORF1ab | nonsynonymous_S2285F |
| 7122 | T | C | | ORF1ab | nonsynonymous_I2286T |
| 7128 | G | A | | ORF1ab | nonsynonymous_C2288Y |
| 7165 | C | T | | ORF1ab | synonymous_T2300T |
| 7170 | C | T | | ORF1ab | nonsynonymous_P2302L |
| 7173 | C | T | | ORF1ab | nonsynonymous_S2303F |
| 7177 | A | T | | ORF1ab | nonsynonymous_L2304F |

| | | | | | |
|---|---|---|---|---|---|
| 7178 | G | A | | ORF1ab | nonsynonymous_E2305K |
| 7187 | C | T | | ORF1ab | stopgain_Q2308X |
| 7191 | T | C | | ORF1ab | nonsynonymous_I2309T |
| 7194 | C | T | | ORF1ab | nonsynonymous_T2310I |
| 7200 | C | T | | ORF1ab | nonsynonymous_S2312L |
| 7203 | C | T | | ORF1ab | nonsynonymous_S2313F |
| 7204 | T | C | | ORF1ab | synonymous_S2313S |
| 7214 | G | T | | ORF1ab | nonsynonymous_D2317Y |
| 7225 | T | C | | ORF1ab | synonymous_A2320A |
| 7229 | G | T | | ORF1ab | nonsynonymous_G2322C |
| 7231 | C | T | | ORF1ab | synonymous_G2322G |
| 7239 | C | T | | ORF1ab | nonsynonymous_A2325V |
| 7246 | G | T | | ORF1ab | nonsynonymous_W2327C |
| 7262 | C | T | | ORF1ab | nonsynonymous_L2333F |
| 7263 | T | C | | ORF1ab | nonsynonymous_L2333P |
| 7267 | C | T | | ORF1ab | synonymous_F2334F |
| 7269 | C | T | | ORF1ab | nonsynonymous_T2335I |
| 7272 | G | T | | ORF1ab | nonsynonymous_R2336M |
| 7273 | G | C | T | ORF1ab | nonsynonymous_R2336S; nonsynonymous_R2336S |
| 7295 | G | C | T | ORF1ab | nonsynonymous_A2344P; nonsynonymous_A2344S |
| 7296 | C | T | | ORF1ab | nonsynonymous_A2344V |
| 7303 | C | T | | ORF1ab | synonymous_I2346I |
| 7306 | G | T | | ORF1ab | nonsynonymous_M2347I |
| 7307 | C | T | | ORF1ab | stopgain_Q2348X |
| 7334 | C | T | | ORF1ab | nonsynonymous_H2357Y |
| 7355 | C | T | | ORF1ab | nonsynonymous_L2364F |
| 7362 | G | T | | ORF1ab | nonsynonymous_W2366L |
| 7363 | G | T | | ORF1ab | nonsynonymous_W2366C |
| 7377 | T | A | | ORF1ab | nonsynonymous_L2371H |

| | | | | | |
|---|---|---|---|---|---|
| 7386 | T | C | | ORF1ab | nonsynonymous_M2374T |
| 7390 | C | T | | ORF1ab | synonymous_A2375A |
| 7394 | A | T | | ORF1ab | nonsynonymous_I2377F |
| 7406 | G | A | | ORF1ab | nonsynonymous_V2381I |
| 7410 | G | A | | ORF1ab | nonsynonymous_R2382K |
| 7420 | C | T | | ORF1ab | synonymous_I2385I |
| 7423 | C | T | | ORF1ab | synonymous_F2386F |
| 7425 | T | C | | ORF1ab | nonsynonymous_F2387S |
| 7432 | A | T | | ORF1ab | synonymous_S2389S |
| 7438 | T | C | | ORF1ab | synonymous_Y2391Y |
| 7447 | G | A | | ORF1ab | stopgain_W2394X |
| 7463 | G | C | | ORF1ab | nonsynonymous_V2400L |
| 7464 | T | G | | ORF1ab | nonsynonymous_V2400G |
| 7465 | T | C | | ORF1ab | synonymous_V2400V |
| 7471 | C | T | | ORF1ab | synonymous_D2402D |
| 7479 | A | G | | ORF1ab | nonsynonymous_N2405S |
| 7483 | A | T | | ORF1ab | synonymous_S2406S |
| 7488 | C | T | | ORF1ab | nonsynonymous_T2408I |
| 7512 | A | G | | ORF1ab | nonsynonymous_N2416S |
| 7514 | A | C | | ORF1ab | synonymous_R2417R |
| 7515 | G | T | | ORF1ab | nonsynonymous_R2417I |
| 7517 | G | T | | ORF1ab | nonsynonymous_A2418S |
| 7518 | C | T | | ORF1ab | nonsynonymous_A2418V |
| 7521 | C | T | | ORF1ab | nonsynonymous_T2419I |
| 7524 | G | T | | ORF1ab | nonsynonymous_R2420I |
| 7528 | C | T | | ORF1ab | synonymous_V2421V |
| 7533 | G | T | | ORF1ab | nonsynonymous_C2423F |
| 7534 | T | C | | ORF1ab | synonymous_C2423C |
| 7536 | C | T | | ORF1ab | nonsynonymous_T2424I |
| 7540 | T | C | | ORF1ab | synonymous_T2425T |
| 7544 | G | T | | ORF1ab | nonsynonymous_V2427F |
| 7550 | G | T | | ORF1ab | nonsynonymous_G2429C |

| | | | | | |
|---|---|---|---|---|---|
| 7553 | G | A | | ORF1ab | nonsynonymous_V2430I |
| 7558 | A | T | | ORF1ab | nonsynonymous_R2431S |
| 7561 | G | T | | ORF1ab | nonsynonymous_R2432S |
| 7563 | C | T | | ORF1ab | nonsynonymous_S2433F |
| 7564 | C | T | | ORF1ab | synonymous_S2433S |
| 7571 | G | T | | ORF1ab | nonsynonymous_V2436F |
| 7579 | T | A | | ORF1ab | synonymous_A2438A |
| 7580 | A | G | | ORF1ab | nonsynonymous_N2439D |
| 7582 | T | C | | ORF1ab | synonymous_N2439N |
| 7585 | A | C | | ORF1ab | synonymous_G2440G |
| 7586 | G | T | | ORF1ab | nonsynonymous_G2441C |
| 7595 | T | C | | ORF1ab | nonsynonymous_F2444L |
| 7597 | T | C | | ORF1ab | synonymous_F2444F |
| 7600 | C | T | | ORF1ab | synonymous_C2445C |
| 7607 | C | T | | ORF1ab | nonsynonymous_H2448Y |
| 7615 | G | A | | ORF1ab | stopgain_W2450X |
| 7616 | A | T | | ORF1ab | nonsynonymous_N2451Y |
| 7626 | A | G | | ORF1ab | nonsynonymous_N2454S |
| 7630 | T | C | | ORF1ab | synonymous_C2455C |
| 7634 | A | G | | ORF1ab | nonsynonymous_T2457A |
| 7660 | T | C | | ORF1ab | synonymous_I2465I |
| 7674 | C | T | | ORF1ab | nonsynonymous_A2470V |
| 7679 | G | A | | ORF1ab | nonsynonymous_D2472N |
| 7688 | C | T | | ORF1ab | synonymous_L2475L |
| 7691 | C | T | | ORF1ab | stopgain_Q2476X |
| 7696 | T | A | | ORF1ab | nonsynonymous_F2477L |
| 7703 | C | T | | ORF1ab | nonsynonymous_P2480S |
| 7704 | C | T | | ORF1ab | nonsynonymous_P2480L |
| 7711 | T | C | | ORF1ab | synonymous_N2482N |
| 7713 | C | T | | ORF1ab | nonsynonymous_P2483L |
| 7720 | C | T | | ORF1ab | synonymous_D2485D |
| 7725 | C | T | | ORF1ab | nonsynonymous_S2487F |

| | | | | | |
|---|---|---|---|---|---|
| 7728 | C | T | | ORF1ab | nonsynonymous_S2488F |
| 7729 | T | C | | ORF1ab | synonymous_S2488S |
| 7732 | C | T | | ORF1ab | synonymous_Y2489Y |
| 7735 | C | A | | ORF1ab | synonymous_I2490I |
| 7749 | C | T | | ORF1ab | nonsynonymous_T2495I |
| 7761 | G | T | | ORF1ab | nonsynonymous_G2499V |
| 7764 | C | T | | ORF1ab | nonsynonymous_S2500F |
| 7765 | C | T | G | ORF1ab | synonymous_S2500S;<br>synonymous_S2500S |
| 7769 | C | T | | ORF1ab | nonsynonymous_H2502Y |
| 7784 | A | T | | ORF1ab | stopgain_K2507X |
| 7786 | A | T | | ORF1ab | nonsynonymous_K2507N |
| 7787 | G | T | | ORF1ab | nonsynonymous_A2508S |
| 7788 | C | T | | ORF1ab | nonsynonymous_A2508V |
| 7791 | G | T | | ORF1ab | nonsynonymous_G2509V |
| 7793 | C | A | | ORF1ab | nonsynonymous_Q2510K |
| 7798 | G | T | | ORF1ab | nonsynonymous_K2511N |
| 7800 | C | A | | ORF1ab | nonsynonymous_T2512N |
| 7811 | C | T | | ORF1ab | nonsynonymous_H2516Y |
| 7815 | C | T | | ORF1ab | nonsynonymous_S2517F |
| 7816 | T | C | | ORF1ab | synonymous_S2517S |
| 7817 | C | T | | ORF1ab | nonsynonymous_L2518F |
| 7818 | T | C | | ORF1ab | nonsynonymous_L2518P |
| 7820 | T | C | | ORF1ab | nonsynonymous_S2519P |
| 7822 | T | A | | ORF1ab | synonymous_S2519S |
| 7826 | T | C | | ORF1ab | nonsynonymous_F2521L |
| 7834 | C | T | | ORF1ab | synonymous_N2523N |
| 7843 | C | T | | ORF1ab | synonymous_N2526N |
| 7860 | C | T | | ORF1ab | nonsynonymous_T2532I |
| 7865 | G | A | | ORF1ab | nonsynonymous_G2534S |
| 7869 | C | T | | ORF1ab | nonsynonymous_S2535L |

| | | | | | |
|---|---|---|---|---|---|
| 7870 | A | G | | ORF1ab | synonymous_S2535S |
| 7887 | T | C | | ORF1ab | nonsynonymous_I2541T |
| 7905 | C | T | | ORF1ab | nonsynonymous_S2547L |
| 7919 | T | C | | ORF1ab | nonsynonymous_S2552P |
| 7924 | T | C | | ORF1ab | synonymous_S2553S |
| 7926 | C | T | | ORF1ab | nonsynonymous_A2554V |
| 7932 | C | T | | ORF1ab | nonsynonymous_S2556L |
| 7936 | G | T | | ORF1ab | synonymous_A2557A |
| 7938 | C | T | | ORF1ab | nonsynonymous_S2558F |
| 7942 | T | C | | ORF1ab | synonymous_V2559V |
| 7944 | A | G | | ORF1ab | nonsynonymous_Y2560C |
| 7945 | C | T | | ORF1ab | synonymous_Y2560Y |
| 7952 | C | T | | ORF1ab | stopgain_Q2563X |
| 7955 | C | T | | ORF1ab | nonsynonymous_L2564F |
| 7967 | C | T | | ORF1ab | nonsynonymous_P2568S |
| 7990 | A | T | | ORF1ab | synonymous_A2575A |
| 7997 | T | C | | ORF1ab | nonsynonymous_S2578P |
| 8016 | C | T | | ORF1ab | nonsynonymous_A2584V |
| 8018 | G | T | | ORF1ab | stopgain_E2585X |
| 8025 | C | T | | ORF1ab | nonsynonymous_A2587V |
| 8030 | A | T | | ORF1ab | stopgain_K2589X |
| 8031 | A | G | | ORF1ab | nonsynonymous_K2589R |
| 8039 | G | A | | ORF1ab | nonsynonymous_D2592N |
| 8043 | C | T | | ORF1ab | nonsynonymous_A2593V |
| 8047 | C | T | | ORF1ab | synonymous_Y2594Y |
| 8049 | T | C | | ORF1ab | nonsynonymous_V2595A |
| 8054 | A | T | | ORF1ab | nonsynonymous_T2597S |
| 8062 | A | T | | ORF1ab | synonymous_S2599S |
| 8064 | C | T | | ORF1ab | nonsynonymous_S2600L |
| 8068 | T | C | | ORF1ab | synonymous_T2601T |

| | | | | | |
|---|---|---|---|---|---|
| 8075 | G | C | T | ORF1ab | nonsynonymous_V2604L; nonsynonymous_V2604L |
| 8078 | C | T | G | ORF1ab | nonsynonymous_P2605S; nonsynonymous_P2605A |
| 8082 | T | C | | ORF1ab | nonsynonymous_M2606T |
| 8083 | G | A | | ORF1ab | nonsynonymous_M2606I |
| 8084 | G | A | | ORF1ab | nonsynonymous_E2607K |
| 8090 | C | T | | ORF1ab | nonsynonymous_L2609F |
| 8102 | G | T | | ORF1ab | nonsynonymous_V2613F |
| 8105 | G | T | | ORF1ab | nonsynonymous_A2614S |
| 8109 | C | T | | ORF1ab | nonsynonymous_T2615I |
| 8118 | C | T | | ORF1ab | nonsynonymous_A2618V |
| 8120 | G | C | | ORF1ab | nonsynonymous_E2619Q |
| 8123 | C | T | | ORF1ab | nonsynonymous_L2620F |
| 8126 | G | T | | ORF1ab | nonsynonymous_A2621S |
| 8131 | G | A | | ORF1ab | synonymous_K2622K |
| 8139 | C | T | | ORF1ab | nonsynonymous_S2625F |
| 8140 | C | T | | ORF1ab | synonymous_S2625S |
| 8151 | T | C | | ORF1ab | nonsynonymous_V2629A |
| 8156 | T | C | | ORF1ab | nonsynonymous_S2631P |
| 8174 | G | T | | ORF1ab | nonsynonymous_A2637S |
| 8175 | C | T | | ORF1ab | nonsynonymous_A2637V |
| 8177 | C | T | | ORF1ab | nonsynonymous_R2638W |
| 8180 | C | T | | ORF1ab | stopgain_Q2639X |
| 8182 | A | G | | ORF1ab | synonymous_Q2639Q |
| 8185 | G | A | T | ORF1ab | synonymous_G2640G; synonymous_G2640G |
| 8190 | T | C | | ORF1ab | nonsynonymous_V2642A |
| 8196 | C | T | | ORF1ab | nonsynonymous_S2644L |
| 8206 | A | C | | ORF1ab | nonsynonymous_E2647D |
| 8207 | A | T | | ORF1ab | nonsynonymous_T2648S |

| | | | | | |
|---|---|---|---|---|---|
| 8208 | C | T | | ORF1ab | nonsynonymous_T2648I |
| 8214 | A | G | | ORF1ab | nonsynonymous_D2650G |
| 8228 | C | T | | ORF1ab | nonsynonymous_L2655F |
| 8238 | C | T | | ORF1ab | nonsynonymous_S2658L |
| 8240 | C | T | | ORF1ab | nonsynonymous_H2659Y |
| 8243 | C | T | G | ORF1ab | stopgain_Q2660X; nonsynonymous_Q2660E |
| 8245 | A | G | | ORF1ab | synonymous_Q2660Q |
| 8247 | C | T | | ORF1ab | nonsynonymous_S2661F |
| 8251 | C | T | | ORF1ab | synonymous_D2662D |
| 8261 | A | C | | ORF1ab | nonsynonymous_T2666P |
| 8262 | C | T | | ORF1ab | nonsynonymous_T2666I |
| 8275 | T | C | | ORF1ab | synonymous_C2670C |
| 8281 | C | T | | ORF1ab | synonymous_N2672N |
| 8288 | C | T | | ORF1ab | nonsynonymous_L2675F |
| 8290 | C | T | | ORF1ab | synonymous_L2675L |
| 8291 | A | T | | ORF1ab | nonsynonymous_T2676S |
| 8293 | C | A | | ORF1ab | synonymous_T2676T |
| 8296 | T | C | | ORF1ab | synonymous_Y2677Y |
| 8298 | A | G | | ORF1ab | nonsynonymous_N2678S |
| 8299 | C | T | | ORF1ab | synonymous_N2678N |
| 8318 | C | T | | ORF1ab | nonsynonymous_P2685S |
| 8319 | C | T | | ORF1ab | nonsynonymous_P2685L |
| 8320 | C | T | | ORF1ab | synonymous_P2685P |
| 8331 | G | T | | ORF1ab | nonsynonymous_G2689V |
| 8335 | T | C | | ORF1ab | synonymous_A2690A |
| 8344 | C | T | | ORF1ab | synonymous_D2693D |
| 8349 | G | T | | ORF1ab | nonsynonymous_S2695I |
| 8350 | T | C | | ORF1ab | synonymous_S2695S |
| 8351 | G | T | | ORF1ab | nonsynonymous_A2696S |
| 8352 | C | T | | ORF1ab | nonsynonymous_A2696V |

| | | | | | |
|---|---|---|---|---|---|
| 8367 | C | T | | ORF1ab | nonsynonymous_A2701V |
| 8371 | G | A | T | ORF1ab | synonymous_Q2702Q; nonsynonymous_Q2702H |
| 8389 | C | T | | ORF1ab | synonymous_N2708N |
| 8395 | T | C | | ORF1ab | synonymous_A2710A |
| 8403 | G | T | | ORF1ab | nonsynonymous_W2713L |
| 8415 | A | T | | ORF1ab | nonsynonymous_D2717V |
| 8419 | C | T | | ORF1ab | synonymous_F2718F |
| 8430 | C | T | | ORF1ab | nonsynonymous_S2722F |
| 8433 | A | C | G | ORF1ab | nonsynonymous_E2723A; nonsynonymous_E2723G |
| 8447 | C | T | | ORF1ab | stopgain_Q2728X |
| 8459 | G | T | | ORF1ab | nonsynonymous_A2732S |
| 8460 | C | T | | ORF1ab | nonsynonymous_A2732V |
| 8463 | C | T | | ORF1ab | nonsynonymous_A2733V |
| 8476 | C | T | | ORF1ab | synonymous_N2737N |
| 8480 | C | T | | ORF1ab | nonsynonymous_P2739S |
| 8492 | A | T | | ORF1ab | nonsynonymous_T2743S |
| 8495 | T | C | | ORF1ab | nonsynonymous_C2744R |
| 8496 | G | A | | ORF1ab | nonsynonymous_C2744Y |
| 8503 | T | C | | ORF1ab | synonymous_T2746T |
| 8506 | T | C | | ORF1ab | synonymous_T2747T |
| 8510 | C | T | | ORF1ab | stopgain_Q2749X |
| 8527 | A | G | | ORF1ab | synonymous_V2754V |
| 8529 | C | T | | ORF1ab | nonsynonymous_T2755I |
| 8532 | C | T | | ORF1ab | nonsynonymous_T2756I |
| 8533 | A | G | | ORF1ab | synonymous_T2756T |
| 8541 | C | T | | ORF1ab | nonsynonymous_A2759V |
| 8550 | G | T | | ORF1ab | nonsynonymous_G2762V |
| 8552 | G | T | | ORF1ab | nonsynonymous_G2763C |
| 8579 | C | T | | ORF1ab | stopgain_Q2772X |

| | | | | | |
|---|---|---|---|---|---|
| 8599 | T | C | | ORF1ab | synonymous_L2778L |
| 8600 | G | C | | ORF1ab | nonsynonymous_V2779L |
| 8605 | C | T | | ORF1ab | synonymous_F2780F |
| 8606 | C | T | | ORF1ab | nonsynonymous_L2781F |
| 8625 | T | C | | ORF1ab | nonsynonymous_F2787S |
| 8626 | C | T | | ORF1ab | synonymous_F2787F |
| 8645 | C | T | | ORF1ab | nonsynonymous_H2794Y |
| 8653 | G | T | | ORF1ab | nonsynonymous_M2796I |
| 8660 | C | T | | ORF1ab | nonsynonymous_H2799Y |
| 8664 | C | T | | ORF1ab | nonsynonymous_T2800I |
| 8676 | G | T | | ORF1ab | nonsynonymous_S2804I |
| 8683 | C | T | | ORF1ab | synonymous_I2806I |
| 8688 | G | T | | ORF1ab | nonsynonymous_G2808V |
| 8690 | T | C | | ORF1ab | nonsynonymous_Y2809H |
| 8696 | G | T | | ORF1ab | nonsynonymous_A2811S |
| 8713 | C | A | | ORF1ab | synonymous_V2816V |
| 8721 | A | G | | ORF1ab | nonsynonymous_D2819G |
| 8727 | C | T | | ORF1ab | nonsynonymous_A2821V |
| 8728 | A | G | | ORF1ab | synonymous_A2821A |
| 8733 | C | T | | ORF1ab | nonsynonymous_T2823I |
| 8739 | C | T | | ORF1ab | nonsynonymous_T2825I |
| 8748 | C | T | | ORF1ab | nonsynonymous_A2828V |
| 8751 | A | G | | ORF1ab | nonsynonymous_N2829S |
| 8756 | C | T | | ORF1ab | nonsynonymous_H2831Y |
| 8759 | G | T | | ORF1ab | nonsynonymous_A2832S |
| 8770 | C | T | | ORF1ab | synonymous_D2835D |
| 8782 | C | T | | ORF1ab | synonymous_S2839S |
| 8786 | C | T | | ORF1ab | nonsynonymous_R2841C |
| 8787 | G | T | | ORF1ab | nonsynonymous_R2841L |
| 8790 | G | T | | ORF1ab | nonsynonymous_G2842V |
| 8802 | C | T | | ORF1ab | nonsynonymous_T2846I |
| 8809 | C | T | | ORF1ab | synonymous_D2848D |

| | | | | | |
|---|---|---|---|---|---|
| 8815 | T | C | | ORF1ab | synonymous_A2850A |
| 8819 | C | T | | ORF1ab | nonsynonymous_P2852S |
| 8825 | A | T | G | ORF1ab | nonsynonymous_I2854F;<br>nonsynonymous_I2854V |
| 8849 | G | A | | ORF1ab | nonsynonymous_V2862M |
| 8852 | G | A | | ORF1ab | nonsynonymous_G2863S |
| 8873 | C | T | | ORF1ab | nonsynonymous_P2870S |
| 8879 | A | G | | ORF1ab | nonsynonymous_T2872A |
| 8884 | A | T | | ORF1ab | synonymous_I2873I |
| 8889 | G | T | | ORF1ab | nonsynonymous_R2875L |
| 8892 | C | T | | ORF1ab | nonsynonymous_T2876I |
| 8895 | C | T | | ORF1ab | nonsynonymous_T2877I |
| 8917 | C | T | | ORF1ab | synonymous_F2884F |
| 8919 | T | C | | ORF1ab | nonsynonymous_L2885S |
| 8921 | C | T | | ORF1ab | nonsynonymous_P2886S |
| 8922 | C | T | | ORF1ab | nonsynonymous_P2886L |
| 8934 | G | T | | ORF1ab | nonsynonymous_S2890I |
| 8947 | C | T | | ORF1ab | synonymous_N2894N |
| 8950 | C | T | | ORF1ab | synonymous_I2895I |
| 8952 | G | T | | ORF1ab | nonsynonymous_C2896F |
| 8960 | C | T | | ORF1ab | nonsynonymous_P2899S |
| 8963 | T | C | | ORF1ab | nonsynonymous_S2900P |
| 8964 | C | T | | ORF1ab | nonsynonymous_S2900L |
| 8991 | C | T | | ORF1ab | nonsynonymous_A2909V |
| 8995 | A | T | | ORF1ab | synonymous_T2910T |
| 9005 | G | T | | ORF1ab | nonsynonymous_V2914F |
| 9015 | C | T | | ORF1ab | nonsynonymous_A2917V |
| 9018 | A | G | | ORF1ab | nonsynonymous_E2918G |
| 9034 | A | G | | ORF1ab | synonymous_K2923K |
| 9037 | T | C | | ORF1ab | synonymous_D2924D |

| | | | | | |
|---|---|---|---|---|---|
| 9052 | A | T | G | ORF1ab | synonymous_P2929P; synonymous_P2929P |
| 9053 | G | T | | ORF1ab | nonsynonymous_V2930L |
| 9054 | T | A | | ORF1ab | nonsynonymous_V2930E |
| 9073 | C | T | | ORF1ab | synonymous_T2936T |
| 9079 | A | C | | ORF1ab | synonymous_V2938V |
| 9086 | G | T | | ORF1ab | nonsynonymous_G2941C |
| 9087 | G | T | | ORF1ab | nonsynonymous_G2941V |
| 9090 | C | T | | ORF1ab | nonsynonymous_S2942F |
| 9092 | G | A | | ORF1ab | nonsynonymous_V2943I |
| 9096 | C | T | | ORF1ab | nonsynonymous_A2944V |
| 9104 | A | G | | ORF1ab | nonsynonymous_S2947G |
| 9112 | C | T | | ORF1ab | synonymous_R2949R |
| 9113 | C | T | | ORF1ab | nonsynonymous_P2950S |
| 9119 | A | G | | ORF1ab | nonsynonymous_T2952A |
| 9122 | C | T | | ORF1ab | nonsynonymous_R2953C |
| 9128 | G | T | | ORF1ab | nonsynonymous_V2955L |
| 9130 | G | T | | ORF1ab | synonymous_V2955V |
| 9133 | C | T | | ORF1ab | synonymous_L2956L |
| 9135 | T | G | | ORF1ab | nonsynonymous_M2957R |
| 9136 | G | C | | ORF1ab | nonsynonymous_M2957I |
| 9137 | G | T | | ORF1ab | nonsynonymous_D2958Y |
| 9141 | G | T | | ORF1ab | nonsynonymous_G2959V |
| 9146 | A | C | | ORF1ab | nonsynonymous_I2961L |
| 9152 | C | T | | ORF1ab | stopgain_Q2963X |
| 9158 | C | T | | ORF1ab | nonsynonymous_P2965S |
| 9163 | C | T | | ORF1ab | synonymous_N2966N |
| 9165 | C | T | | ORF1ab | nonsynonymous_T2967I |
| 9166 | C | T | | ORF1ab | synonymous_T2967T |
| 9167 | T | C | | ORF1ab | nonsynonymous_Y2968H |
| 9170 | C | T | | ORF1ab | nonsynonymous_L2969F |

| | | | | | |
|---|---|---|---|---|---|
| 9172 | T | C | | ORF1ab | synonymous_L2969L |
| 9175 | A | C | | ORF1ab | nonsynonymous_E2970D |
| 9176 | G | T | | ORF1ab | nonsynonymous_G2971C |
| 9180 | C | T | | ORF1ab | nonsynonymous_S2972F |
| 9190 | G | A | C | ORF1ab | synonymous_V2975V; synonymous_V2975V |
| 9191 | G | T | | ORF1ab | nonsynonymous_V2976L |
| 9198 | C | T | | ORF1ab | nonsynonymous_T2978I |
| 9207 | C | T | | ORF1ab | nonsynonymous_S2981F |
| 9208 | T | A | | ORF1ab | synonymous_S2981S |
| 9214 | C | T | | ORF1ab | synonymous_Y2983Y |
| 9215 | T | G | | ORF1ab | nonsynonymous_C2984G |
| 9220 | G | T | | ORF1ab | nonsynonymous_R2985S |
| 9221 | C | T | | ORF1ab | nonsynonymous_H2986Y |
| 9223 | C | T | | ORF1ab | synonymous_H2986H |
| 9228 | C | T | | ORF1ab | nonsynonymous_T2988I |
| 9231 | G | T | | ORF1ab | nonsynonymous_C2989F |
| 9240 | C | T | | ORF1ab | nonsynonymous_S2992L |
| 9241 | A | G | | ORF1ab | synonymous_S2992S |
| 9242 | G | T | | ORF1ab | stopgain_E2993X |
| 9244 | A | G | | ORF1ab | synonymous_E2993E |
| 9245 | G | T | | ORF1ab | nonsynonymous_A2994S |
| 9246 | C | T | | ORF1ab | nonsynonymous_A2994V |
| 9249 | G | T | | ORF1ab | nonsynonymous_G2995V |
| 9251 | G | T | | ORF1ab | nonsynonymous_V2996F |
| 9258 | T | C | | ORF1ab | nonsynonymous_V2998A |
| 9274 | A | G | | ORF1ab | synonymous_R3003R |
| 9286 | C | T | | ORF1ab | synonymous_N3007N |
| 9290 | G | T | | ORF1ab | nonsynonymous_D3009Y |
| 9307 | A | G | | ORF1ab | synonymous_L3014L |
| 9319 | C | T | | ORF1ab | synonymous_F3018F |

| | | | | | |
|---|---|---|---|---|---|
| 9333 | C | T | | ORF1ab | nonsynonymous_A3023V |
| 9335 | G | T | | ORF1ab | nonsynonymous_V3024L |
| 9344 | C | T | | ORF1ab | nonsynonymous_L3027F |
| 9354 | T | C | | ORF1ab | nonsynonymous_M3030T |
| 9355 | G | T | | ORF1ab | nonsynonymous_M3030I |
| 9360 | C | T | | ORF1ab | nonsynonymous_T3032I |
| 9362 | C | T | | ORF1ab | nonsynonymous_P3033S |
| 9371 | C | T | | ORF1ab | stopgain_Q3036X |
| 9374 | C | T | | ORF1ab | nonsynonymous_P3037S |
| 9375 | C | T | | ORF1ab | nonsynonymous_P3037L |
| 9384 | C | T | | ORF1ab | nonsynonymous_A3040V |
| 9389 | G | T | | ORF1ab | nonsynonymous_D3042Y |
| 9391 | C | T | | ORF1ab | synonymous_D3042D |
| 9399 | C | T | | ORF1ab | nonsynonymous_A3045V |
| 9402 | C | T | | ORF1ab | nonsynonymous_S3046F |
| 9421 | T | C | | ORF1ab | synonymous_I3052I |
| 9426 | C | T | | ORF1ab | nonsynonymous_A3054V |
| 9430 | C | T | | ORF1ab | synonymous_I3055I |
| 9431 | G | C | T | ORF1ab | nonsynonymous_V3056L;<br>nonsynonymous_V3056L |
| 9433 | A | G | | ORF1ab | synonymous_V3056V |
| 9438 | C | T | | ORF1ab | nonsynonymous_T3058I |
| 9442 | C | T | | ORF1ab | synonymous_C3059C |
| 9443 | C | T | | ORF1ab | nonsynonymous_L3060F |
| 9448 | C | T | | ORF1ab | synonymous_A3061A |
| 9471 | G | T | | ORF1ab | nonsynonymous_R3069I |
| 9474 | C | T | | ORF1ab | nonsynonymous_A3070V |
| 9477 | T | A | | ORF1ab | nonsynonymous_F3071Y |
| 9479 | G | T | | ORF1ab | nonsynonymous_G3072C |
| 9483 | A | G | | ORF1ab | nonsynonymous_E3073G |
| 9491 | C | T | | ORF1ab | nonsynonymous_H3076Y |

| | | | | | |
|---|---|---|---|---|---|
| 9500 | G | T | | ORF1ab | nonsynonymous_A3079S |
| 9502 | C | T | | ORF1ab | synonymous_A3079A |
| 9503 | T | C | | ORF1ab | nonsynonymous_F3080L |
| 9514 | A | G | | ORF1ab | synonymous_L3083L |
| 9519 | T | C | G | ORF1ab | nonsynonymous_F3085S; nonsynonymous_F3085C |
| 9528 | C | T | | ORF1ab | nonsynonymous_S3088L |
| 9534 | C | T | | ORF1ab | nonsynonymous_T3090I |
| 9551 | C | T | | ORF1ab | nonsynonymous_P3096S |
| 9565 | C | T | | ORF1ab | synonymous_F3100F |
| 9571 | T | C | | ORF1ab | synonymous_P3102P |
| 9574 | T | C | | ORF1ab | synonymous_G3103G |
| 9598 | C | T | | ORF1ab | synonymous_Y3111Y |
| 9604 | A | T | | ORF1ab | synonymous_T3113T |
| 9611 | C | T | | ORF1ab | nonsynonymous_L3116F |
| 9613 | T | A | | ORF1ab | synonymous_L3116L |
| 9614 | A | T | | ORF1ab | nonsynonymous_T3117S |
| 9621 | A | T | | ORF1ab | nonsynonymous_D3119V |
| 9626 | T | C | | ORF1ab | nonsynonymous_S3121P |
| 9627 | C | T | | ORF1ab | nonsynonymous_S3121F |
| 9630 | T | C | | ORF1ab | nonsynonymous_F3122S |
| 9635 | G | A | | ORF1ab | nonsynonymous_A3124T |
| 9649 | G | T | | ORF1ab | nonsynonymous_W3128C |
| 9652 | G | T | | ORF1ab | nonsynonymous_M3129I |
| 9663 | C | T | | ORF1ab | nonsynonymous_T3133I |
| 9666 | C | T | | ORF1ab | nonsynonymous_P3134L |
| 9676 | T | C | | ORF1ab | synonymous_P3137P |
| 9682 | G | C | | ORF1ab | nonsynonymous_W3139C |
| 9683 | A | G | | ORF1ab | nonsynonymous_I3140V |
| 9693 | C | T | | ORF1ab | nonsynonymous_A3143V |
| 9700 | C | T | | ORF1ab | synonymous_I3145I |

| | | | | | |
|---|---|---|---|---|---|
| 9707 | A | G | | ORF1ab | nonsynonymous_I3148V |
| 9710 | T | C | | ORF1ab | nonsynonymous_S3149P |
| 9714 | C | T | | ORF1ab | nonsynonymous_T3150I |
| 9717 | A | G | | ORF1ab | nonsynonymous_K3151R |
| 9721 | T | C | | ORF1ab | synonymous_H3152H |
| 9724 | C | T | | ORF1ab | synonymous_F3153F |
| 9734 | T | C | | ORF1ab | nonsynonymous_F3157L |
| 9743 | T | C | | ORF1ab | nonsynonymous_Y3160H |
| 9745 | C | T | | ORF1ab | synonymous_Y3160Y |
| 9747 | T | C | | ORF1ab | nonsynonymous_L3161P |
| 9748 | A | T | | ORF1ab | synonymous_L3161L |
| 9749 | A | G | | ORF1ab | nonsynonymous_K3162E |
| 9755 | C | T | | ORF1ab | nonsynonymous_R3164C |
| 9762 | T | C | | ORF1ab | nonsynonymous_V3166A |
| 9763 | C | T | | ORF1ab | synonymous_V3166V |
| 9768 | A | G | | ORF1ab | nonsynonymous_N3168S |
| 9772 | T | C | | ORF1ab | synonymous_G3169G |
| 9777 | C | T | | ORF1ab | nonsynonymous_S3171F |
| 9785 | A | T | | ORF1ab | nonsynonymous_T3174S |
| 9787 | T | C | | ORF1ab | synonymous_T3174T |
| 9792 | A | T | | ORF1ab | nonsynonymous_E3176V |
| 9794 | G | T | | ORF1ab | stopgain_E3177X |
| 9798 | C | T | | ORF1ab | nonsynonymous_A3178V |
| 9801 | C | T | | ORF1ab | nonsynonymous_A3179V |
| 9802 | G | T | | ORF1ab | synonymous_A3179A |
| 9803 | C | T | | ORF1ab | synonymous_L3180L |
| 9804 | T | G | | ORF1ab | nonsynonymous_L3180R |
| 9805 | G | C | T | ORF1ab | synonymous_L3180L;<br>synonymous_L3180L |
| 9810 | C | T | | ORF1ab | nonsynonymous_T3182I |
| 9857 | C | T | | ORF1ab | synonymous_L3198L |

| | | | | | |
|---|---|---|---|---|---|
| 9863 | C | T | | ORF1ab | nonsynonymous_P3200S |
| 9864 | C | T | | ORF1ab | nonsynonymous_P3200L |
| 9866 | C | T | | ORF1ab | nonsynonymous_L3201F |
| 9873 | A | T | | ORF1ab | nonsynonymous_Q3203L |
| 9890 | G | T | | ORF1ab | nonsynonymous_A3209S |
| 9891 | C | T | | ORF1ab | nonsynonymous_A3209V |
| 9893 | C | T | | ORF1ab | nonsynonymous_L3210F |
| 9895 | T | C | | ORF1ab | synonymous_L3210L |
| 9923 | G | A | | ORF1ab | nonsynonymous_A3220T |
| 9924 | C | T | | ORF1ab | nonsynonymous_A3220V |
| 9933 | C | T | | ORF1ab | nonsynonymous_T3223I |
| 9940 | C | T | | ORF1ab | synonymous_S3225S |
| 9946 | A | T | | ORF1ab | nonsynonymous_R3227S |
| 9962 | C | T | | ORF1ab | nonsynonymous_H3233Y |
| 9967 | C | T | | ORF1ab | synonymous_L3234L |
| 9968 | G | A | | ORF1ab | nonsynonymous_A3235T |
| 9974 | G | T | | ORF1ab | nonsynonymous_A3237S |
| 9975 | C | T | | ORF1ab | nonsynonymous_A3237V |
| 9977 | C | T | | ORF1ab | nonsynonymous_L3238F |
| 9979 | C | T | | ORF1ab | synonymous_L3238L |
| 9990 | G | T | | ORF1ab | nonsynonymous_S3242I |
| 9996 | C | T | | ORF1ab | nonsynonymous_S3244L |
| 9997 | A | T | | ORF1ab | synonymous_S3244S |
| 10002 | C | T | | ORF1ab | nonsynonymous_S3246F |
| 10010 | C | T | | ORF1ab | nonsynonymous_L3249F |
| 10019 | C | T | | ORF1ab | nonsynonymous_P3252S |
| 10025 | C | T | | ORF1ab | stopgain_Q3254X |
| 10029 | C | T | | ORF1ab | nonsynonymous_T3255I |
| 10030 | C | T | | ORF1ab | synonymous_T3255T |
| 10032 | C | T | | ORF1ab | nonsynonymous_S3256F |
| 10036 | C | T | | ORF1ab | synonymous_I3257I |
| 10038 | C | T | | ORF1ab | nonsynonymous_T3258I |

| | | | | | |
|---|---|---|---|---|---|
| 10043 | G | T | | ORF1ab | nonsynonymous_A3260S |
| 10044 | C | T | | ORF1ab | nonsynonymous_A3260V |
| 10046 | G | A | | ORF1ab | nonsynonymous_V3261I |
| 10052 | C | T | | ORF1ab | stopgain_Q3263X |
| 10054 | G | T | | ORF1ab | nonsynonymous_Q3263H |
| 10057 | T | G | | ORF1ab | nonsynonymous_S3264R |
| 10059 | G | T | | ORF1ab | nonsynonymous_G3265V |
| 10073 | G | A | | ORF1ab | nonsynonymous_A3270T |
| 10074 | C | T | | ORF1ab | nonsynonymous_A3270V |
| 10078 | C | T | | ORF1ab | synonymous_F3271F |
| 10079 | C | T | | ORF1ab | nonsynonymous_P3272S |
| 10080 | C | T | | ORF1ab | nonsynonymous_P3272L |
| 10081 | A | G | | ORF1ab | synonymous_P3272P |
| 10083 | C | T | | ORF1ab | nonsynonymous_S3273F |
| 10097 | G | A | | ORF1ab | nonsynonymous_G3278S |
| 10098 | G | T | | ORF1ab | nonsynonymous_G3278V |
| 10101 | G | T | | ORF1ab | nonsynonymous_C3279F |
| 10111 | A | T | | ORF1ab | nonsynonymous_Q3282H |
| 10114 | A | G | | ORF1ab | synonymous_V3283V |
| 10124 | A | G | | ORF1ab | nonsynonymous_T3287A |
| 10125 | C | T | | ORF1ab | nonsynonymous_T3287I |
| 10134 | T | G | | ORF1ab | nonsynonymous_L3290R |
| 10138 | C | T | | ORF1ab | synonymous_N3291N |
| 10139 | G | T | | ORF1ab | nonsynonymous_G3292C |
| 10143 | T | G | | ORF1ab | nonsynonymous_L3293R |
| 10162 | T | C | | ORF1ab | synonymous_V3299V |
| 10167 | G | T | | ORF1ab | nonsynonymous_C3301F |
| 10187 | A | C | | ORF1ab | nonsynonymous_T3308P |
| 10193 | G | T | | ORF1ab | stopgain_E3310X |
| 10196 | G | T | | ORF1ab | nonsynonymous_D3311Y |
| 10198 | C | T | | ORF1ab | synonymous_D3311D |
| 10201 | G | T | | ORF1ab | nonsynonymous_M3312I |

| | | | | | |
|---|---|---|---|---|---|
| 10228 | C | T | | ORF1ab | synonymous_L3321L |
| 10240 | T | A | | ORF1ab | synonymous_S3325S |
| 10252 | C | T | | ORF1ab | synonymous_F3329F |
| 10256 | G | T | | ORF1ab | nonsynonymous_V3331L |
| 10258 | A | C | | ORF1ab | synonymous_V3331V |
| 10260 | A | G | | ORF1ab | nonsynonymous_Q3332R |
| 10263 | C | T | | ORF1ab | nonsynonymous_A3333V |
| 10264 | T | C | | ORF1ab | synonymous_A3333A |
| 10265 | G | A | | ORF1ab | nonsynonymous_G3334S |
| 10266 | G | T | | ORF1ab | nonsynonymous_G3334V |
| 10271 | G | T | | ORF1ab | nonsynonymous_V3336F |
| 10277 | C | T | | ORF1ab | nonsynonymous_L3338F |
| 10279 | C | T | | ORF1ab | synonymous_L3338L |
| 10281 | G | T | | ORF1ab | nonsynonymous_R3339M |
| 10282 | G | A | | ORF1ab | synonymous_R3339R |
| 10296 | C | T | | ORF1ab | nonsynonymous_S3344F |
| 10301 | C | A | | ORF1ab | nonsynonymous_Q3346K |
| 10308 | G | T | | ORF1ab | nonsynonymous_C3348F |
| 10309 | T | C | | ORF1ab | synonymous_C3348C |
| 10310 | G | A | | ORF1ab | nonsynonymous_V3349I |
| 10311 | T | A | | ORF1ab | nonsynonymous_V3349E |
| 10313 | C | T | | ORF1ab | nonsynonymous_L3350F |
| 10319 | C | T | | ORF1ab | nonsynonymous_L3352F |
| 10323 | A | G | | ORF1ab | nonsynonymous_K3353R |
| 10324 | G | T | | ORF1ab | nonsynonymous_K3353N |
| 10325 | G | T | | ORF1ab | nonsynonymous_V3354F |
| 10326 | T | C | | ORF1ab | nonsynonymous_V3354A |
| 10335 | C | T | | ORF1ab | nonsynonymous_A3357V |
| 10336 | C | T | | ORF1ab | synonymous_A3357A |
| 10347 | C | T | | ORF1ab | nonsynonymous_T3361I |
| 10350 | C | T | | ORF1ab | nonsynonymous_P3362L |

| | | | | | |
|---|---|---|---|---|---|
| 10353 | A | C | T | ORF1ab | nonsynonymous_K3363T; nonsynonymous_K3363M |
| 10360 | G | T | | ORF1ab | nonsynonymous_K3365N |
| 10366 | T | C | | ORF1ab | synonymous_V3367V |
| 10368 | G | A | T | ORF1ab | nonsynonymous_R3368H; nonsynonymous_R3368L |
| 10373 | C | T | G | ORF1ab | stopgain_Q3370X; nonsynonymous_Q3370E |
| 10375 | A | T | | ORF1ab | nonsynonymous_Q3370H |
| 10376 | C | T | | ORF1ab | nonsynonymous_P3371S |
| 10378 | A | T | | ORF1ab | synonymous_P3371P |
| 10379 | G | A | T | ORF1ab | nonsynonymous_G3372R; stopgain_G3372X |
| 10381 | A | T | | ORF1ab | synonymous_G3372G |
| 10384 | G | T | | ORF1ab | nonsynonymous_Q3373H |
| 10387 | T | G | | ORF1ab | synonymous_T3374T |
| 10390 | T | C | | ORF1ab | synonymous_F3375F |
| 10396 | G | A | | ORF1ab | synonymous_V3377V |
| 10401 | C | T | | ORF1ab | nonsynonymous_A3379V |
| 10402 | T | C | | ORF1ab | synonymous_A3379A |
| 10408 | C | T | | ORF1ab | synonymous_Y3381Y |
| 10419 | C | T | | ORF1ab | nonsynonymous_P3385L |
| 10422 | C | T | | ORF1ab | nonsynonymous_S3386F |
| 10425 | G | T | | ORF1ab | nonsynonymous_G3387V |
| 10429 | T | C | | ORF1ab | synonymous_V3388V |
| 10433 | C | T | | ORF1ab | stopgain_Q3390X |
| 10437 | G | C | | ORF1ab | nonsynonymous_C3391S |
| 10440 | C | T | | ORF1ab | nonsynonymous_A3392V |
| 10447 | G | T | | ORF1ab | nonsynonymous_R3394S |
| 10454 | T | C | | ORF1ab | nonsynonymous_F3397L |
| 10456 | C | T | | ORF1ab | synonymous_F3397F |

| | | | | | |
|---|---|---|---|---|---|
| 10458 | C | T | | ORF1ab | nonsynonymous_T3398I |
| 10466 | G | T | | ORF1ab | nonsynonymous_G3401C |
| 10478 | A | C | | ORF1ab | nonsynonymous_N3405H |
| 10479 | A | T | | ORF1ab | nonsynonymous_N3405I |
| 10480 | T | C | | ORF1ab | synonymous_N3405N |
| 10500 | G | T | | ORF1ab | nonsynonymous_G3412V |
| 10507 | C | T | | ORF1ab | synonymous_N3414N |
| 10512 | A | T | | ORF1ab | nonsynonymous_D3416V |
| 10521 | G | T | | ORF1ab | nonsynonymous_C3419F |
| 10527 | C | T | | ORF1ab | nonsynonymous_S3421F |
| 10528 | T | C | | ORF1ab | synonymous_S3421S |
| 10532 | T | C | | ORF1ab | nonsynonymous_C3423R |
| 10535 | T | C | | ORF1ab | nonsynonymous_Y3424H |
| 10537 | C | T | | ORF1ab | synonymous_Y3424Y |
| 10543 | C | T | | ORF1ab | synonymous_H3426H |
| 10547 | A | G | | ORF1ab | nonsynonymous_M3428V |
| 10548 | T | C | | ORF1ab | nonsynonymous_M3428T |
| 10549 | G | T | | ORF1ab | nonsynonymous_M3428I |
| 10556 | C | T | | ORF1ab | nonsynonymous_P3431S |
| 10568 | C | T | | ORF1ab | nonsynonymous_H3435Y |
| 10569 | A | T | | ORF1ab | nonsynonymous_H3435L |
| 10571 | G | T | | ORF1ab | nonsynonymous_A3436S |
| 10575 | G | A | | ORF1ab | nonsynonymous_G3437D |
| 10580 | G | C | T | ORF1ab | nonsynonymous_D3439H; nonsynonymous_D3439Y |
| 10581 | A | G | | ORF1ab | nonsynonymous_D3439G |
| 10582 | C | T | | ORF1ab | synonymous_D3439D |
| 10586 | G | T | | ORF1ab | stopgain_E3441X |
| 10600 | T | C | | ORF1ab | synonymous_Y3445Y |
| 10605 | C | T | | ORF1ab | nonsynonymous_P3447L |
| 10615 | C | T | | ORF1ab | synonymous_D3450D |

| | | | | | |
|---|---|---|---|---|---|
| 10619 | C | A | | ORF1ab | nonsynonymous_Q3452K |
| 10623 | C | T | | ORF1ab | nonsynonymous_T3453I |
| 10626 | C | T | G | ORF1ab | nonsynonymous_A3454V; nonsynonymous_A3454G |
| 10628 | C | T | | ORF1ab | stopgain_Q3455X |
| 10630 | A | T | | ORF1ab | nonsynonymous_Q3455H |
| 10631 | G | A | | ORF1ab | nonsynonymous_A3456T |
| 10641 | C | T | | ORF1ab | nonsynonymous_T3459M |
| 10643 | G | A | | ORF1ab | nonsynonymous_D3460N |
| 10649 | A | T | | ORF1ab | nonsynonymous_T3462S |
| 10657 | A | C | | ORF1ab | synonymous_T3464T |
| 10675 | G | C | T | ORF1ab | nonsynonymous_W3470C; nonsynonymous_W3470C |
| 10700 | G | T | | ORF1ab | nonsynonymous_D3479Y |
| 10714 | C | T | | ORF1ab | synonymous_L3483L |
| 10717 | T | C | | ORF1ab | synonymous_N3484N |
| 10718 | C | T | G | ORF1ab | stopgain_R3485X; nonsynonymous_R3485G |
| 10720 | A | T | | ORF1ab | synonymous_R3485R |
| 10722 | T | C | | ORF1ab | nonsynonymous_F3486S |
| 10741 | C | T | | ORF1ab | synonymous_D3492D |
| 10748 | C | A | | ORF1ab | nonsynonymous_L3495I |
| 10750 | T | C | | ORF1ab | synonymous_L3495L |
| 10754 | G | T | | ORF1ab | nonsynonymous_A3497S |
| 10755 | C | T | | ORF1ab | nonsynonymous_A3497V |
| 10770 | A | G | | ORF1ab | nonsynonymous_Y3502C |
| 10771 | T | C | | ORF1ab | synonymous_Y3502Y |
| 10776 | C | A | T | ORF1ab | nonsynonymous_P3504H; nonsynonymous_P3504L |
| 10784 | C | A | | ORF1ab | nonsynonymous_Q3507K |
| 10789 | C | T | | ORF1ab | synonymous_D3508D |

| | | | | | |
|---|---|---|---|---|---|
| 10793 | G | A | T | ORF1ab | nonsynonymous_V3510I; nonsynonymous_V3510F |
| 10798 | C | A | | ORF1ab | nonsynonymous_D3511E |
| 10805 | G | T | | ORF1ab | stopgain_G3514X |
| 10806 | G | A | | ORF1ab | nonsynonymous_G3514E |
| 10824 | C | T | | ORF1ab | nonsynonymous_T3520I |
| 10828 | A | G | | ORF1ab | synonymous_G3521G |
| 10830 | T | C | | ORF1ab | nonsynonymous_I3522T |
| 10833 | C | T | | ORF1ab | nonsynonymous_A3523V |
| 10835 | G | T | | ORF1ab | nonsynonymous_V3524F |
| 10842 | A | G | | ORF1ab | nonsynonymous_D3526G |
| 10851 | C | T | | ORF1ab | nonsynonymous_A3529V |
| 10856 | T | C | | ORF1ab | synonymous_L3531L |
| 10870 | G | T | | ORF1ab | synonymous_L3535L |
| 10871 | C | T | | ORF1ab | stopgain_Q3536X |
| 10873 | A | G | | ORF1ab | synonymous_Q3536Q |
| 10881 | T | C | | ORF1ab | nonsynonymous_M3539T |
| 10886 | G | T | | ORF1ab | stopgain_G3541X |
| 10889 | C | T | | ORF1ab | nonsynonymous_R3542C |
| 10904 | A | G | | ORF1ab | nonsynonymous_S3547G |
| 10907 | G | T | | ORF1ab | nonsynonymous_A3548S |
| 10912 | A | G | | ORF1ab | synonymous_L3549L |
| 10929 | C | T | | ORF1ab | nonsynonymous_T3555I |
| 10930 | A | G | | ORF1ab | synonymous_T3555T |
| 10931 | C | T | | ORF1ab | nonsynonymous_P3556S |
| 10936 | T | C | | ORF1ab | synonymous_F3557F |
| 10943 | G | C | | ORF1ab | nonsynonymous_V3560L |
| 10948 | A | G | | ORF1ab | synonymous_R3561R |
| 10949 | C | T | | ORF1ab | stopgain_Q3562X |
| 10956 | C | T | | ORF1ab | nonsynonymous_S3564L |
| 10965 | C | T | | ORF1ab | nonsynonymous_T3567I |

| | | | | | |
|---|---|---|---|---|---|
| 10966 | T | C | | ORF1ab | synonymous_T3567T |
| 10969 | C | T | | ORF1ab | synonymous_F3568F |
| 10970 | C | T | | ORF1ab | stopgain_Q3569X |
| 10971 | A | C | | ORF1ab | nonsynonymous_Q3569P |
| 10973 | A | G | | ORF1ab | nonsynonymous_S3570G |
| 10978 | A | G | | ORF1ab | synonymous_A3571A |
| 10979 | G | T | | ORF1ab | nonsynonymous_V3572L |
| 10986 | G | T | | ORF1ab | nonsynonymous_R3574I |
| 10993 | C | T | | ORF1ab | synonymous_I3576I |
| 10995 | A | T | | ORF1ab | nonsynonymous_K3577M |
| 10997 | G | T | | ORF1ab | nonsynonymous_G3578C |
| 11001 | C | T | | ORF1ab | nonsynonymous_T3579I |
| 11003 | C | T | | ORF1ab | nonsynonymous_H3580Y |
| 11005 | C | T | | ORF1ab | synonymous_H3580H |
| 11006 | C | T | | ORF1ab | nonsynonymous_H3581Y |
| 11014 | G | T | | ORF1ab | nonsynonymous_L3583F |
| 11016 | T | C | | ORF1ab | nonsynonymous_L3584S |
| 11022 | C | T | | ORF1ab | nonsynonymous_T3586I |
| 11032 | T | C | | ORF1ab | synonymous_T3589T |
| 11036 | C | T | | ORF1ab | nonsynonymous_L3591F |
| 11040 | T | C | | ORF1ab | nonsynonymous_L3592S |
| 11047 | A | T | | ORF1ab | nonsynonymous_L3594F |
| 11050 | C | T | | ORF1ab | synonymous_V3595V |
| 11056 | T | C | | ORF1ab | synonymous_S3597S |
| 11058 | C | T | | ORF1ab | nonsynonymous_T3598I |
| 11060 | C | T | | ORF1ab | stopgain_Q3599X |
| 11065 | G | C | | ORF1ab | nonsynonymous_W3600C |
| 11066 | T | C | | ORF1ab | nonsynonymous_S3601P |
| 11071 | G | C | | ORF1ab | nonsynonymous_L3602F |
| 11074 | C | T | | ORF1ab | synonymous_F3603F |
| 11075 | T | C | | ORF1ab | nonsynonymous_F3604L |
| 11076 | T | C | | ORF1ab | nonsynonymous_F3604S |

| | | | | | |
|---|---|---|---|---|---|
| 11083 | G | T | | ORF1ab | nonsynonymous_L3606F |
| 11085 | A | C | | ORF1ab | nonsynonymous_Y3607S |
| 11086 | T | A | | ORF1ab | stopgain_Y3607X |
| 11088 | A | G | | ORF1ab | nonsynonymous_E3608G |
| 11094 | C | T | | ORF1ab | nonsynonymous_A3610V |
| 11095 | C | T | | ORF1ab | synonymous_A3610A |
| 11096 | T | C | | ORF1ab | nonsynonymous_F3611L |
| 11104 | T | C | | ORF1ab | synonymous_P3613P |
| 11107 | T | C | | ORF1ab | synonymous_F3614F |
| 11109 | C | T | | ORF1ab | nonsynonymous_A3615V |
| 11113 | G | T | | ORF1ab | nonsynonymous_M3616I |
| 11114 | G | A | | ORF1ab | nonsynonymous_G3617S |
| 11120 | A | T | | ORF1ab | nonsynonymous_I3619F |
| 11131 | T | A | | ORF1ab | synonymous_S3622S |
| 11133 | C | T | | ORF1ab | nonsynonymous_A3623V |
| 11139 | C | T | | ORF1ab | nonsynonymous_A3625V |
| 11149 | T | C | | ORF1ab | synonymous_F3628F |
| 11162 | C | T | | ORF1ab | nonsynonymous_H3633Y |
| 11166 | C | T | | ORF1ab | nonsynonymous_A3634V |
| 11167 | A | T | | ORF1ab | synonymous_A3634A |
| 11182 | T | C | | ORF1ab | synonymous_F3639F |
| 11189 | C | T | | ORF1ab | nonsynonymous_P3642S |
| 11195 | C | T | | ORF1ab | nonsynonymous_L3644F |
| 11198 | G | T | | ORF1ab | nonsynonymous_A3645S |
| 11199 | C | T | | ORF1ab | nonsynonymous_A3645V |
| 11200 | C | T | | ORF1ab | synonymous_A3645A |
| 11202 | C | T | G | ORF1ab | nonsynonymous_T3646I;<br>nonsynonymous_T3646S |
| 11214 | T | A | | ORF1ab | nonsynonymous_F3650Y |
| 11217 | A | C | | ORF1ab | nonsynonymous_N3651T |
| 11218 | T | C | | ORF1ab | synonymous_N3651N |

| | | | | | |
|---|---|---|---|---|---|
| 11221 | G | A | | ORF1ab | nonsynonymous_M3652I |
| 11224 | C | T | | ORF1ab | synonymous_V3653V |
| 11226 | A | C | | ORF1ab | nonsynonymous_Y3654S |
| 11229 | T | G | | ORF1ab | nonsynonymous_M3655R |
| 11231 | C | T | | ORF1ab | nonsynonymous_P3656S |
| 11232 | C | T | | ORF1ab | nonsynonymous_P3656L |
| 11246 | A | G | | ORF1ab | nonsynonymous_M3661V |
| 11253 | T | C | | ORF1ab | nonsynonymous_I3663T |
| 11254 | T | A | | ORF1ab | synonymous_I3663I |
| 11258 | A | G | | ORF1ab | nonsynonymous_T3665A |
| 11280 | C | T | | ORF1ab | nonsynonymous_T3672I |
| 11283 | G | T | | ORF1ab | nonsynonymous_S3673I |
| 11284 | T | C | | ORF1ab | synonymous_S3673S |
| 11286 | T | G | | ORF1ab | nonsynonymous_L3674W |
| 11288 | T | C | | ORF1ab | nonsynonymous_S3675P |
| 11289 | C | T | | ORF1ab | nonsynonymous_S3675F |
| 11300 | C | T | | ORF1ab | synonymous_L3679L |
| 11306 | G | A | | ORF1ab | nonsynonymous_D3681N |
| 11308 | C | T | | ORF1ab | synonymous_D3681D |
| 11339 | C | T | | ORF1ab | synonymous_L3692L |
| 11355 | C | T | | ORF1ab | nonsynonymous_A3697V |
| 11356 | A | G | | ORF1ab | synonymous_A3697A |
| 11364 | T | C | G | ORF1ab | nonsynonymous_V3700A;<br>nonsynonymous_V3700G |
| 11365 | G | T | | ORF1ab | synonymous_V3700V |
| 11375 | G | A | | ORF1ab | nonsynonymous_G3704S |
| 11376 | G | T | | ORF1ab | nonsynonymous_G3704V |
| 11377 | T | C | | ORF1ab | synonymous_G3704G |
| 11392 | G | T | | ORF1ab | nonsynonymous_W3709C |
| 11405 | G | A | | ORF1ab | nonsynonymous_V3714I |
| 11410 | G | A | | ORF1ab | synonymous_L3715L |

| | | | | | |
|---|---|---|---|---|---|
| 11413 | A | G | | ORF1ab | synonymous_T3716T |
| 11416 | C | T | | ORF1ab | synonymous_L3717L |
| 11417 | G | T | | ORF1ab | nonsynonymous_V3718F |
| 11418 | T | C | | ORF1ab | nonsynonymous_V3718A |
| 11422 | T | A | | ORF1ab | stopgain_Y3719X |
| 11430 | A | G | | ORF1ab | nonsynonymous_Y3722C |
| 11433 | A | C | | ORF1ab | nonsynonymous_Y3723S |
| 11443 | T | C | | ORF1ab | synonymous_A3726A |
| 11448 | A | G | | ORF1ab | nonsynonymous_D3728G |
| 11451 | A | G | | ORF1ab | nonsynonymous_Q3729R |
| 11455 | C | T | | ORF1ab | synonymous_A3730A |
| 11471 | C | T | | ORF1ab | nonsynonymous_L3736F |
| 11479 | C | T | | ORF1ab | synonymous_I3738I |
| 11486 | A | T | | ORF1ab | nonsynonymous_T3741S |
| 11487 | C | T | | ORF1ab | nonsynonymous_T3741I |
| 11489 | T | C | | ORF1ab | nonsynonymous_S3742P |
| 11499 | C | T | | ORF1ab | nonsynonymous_S3745L |
| 11502 | G | T | | ORF1ab | nonsynonymous_G3746V |
| 11505 | T | C | | ORF1ab | nonsynonymous_V3747A |
| 11514 | C | T | | ORF1ab | nonsynonymous_T3750I |
| 11515 | T | A | | ORF1ab | synonymous_T3750T |
| 11522 | T | A | | ORF1ab | nonsynonymous_F3753I |
| 11526 | T | G | | ORF1ab | nonsynonymous_L3754W |
| 11530 | C | T | | ORF1ab | synonymous_A3755A |
| 11534 | G | A | | ORF1ab | nonsynonymous_G3757S |
| 11535 | G | T | | ORF1ab | nonsynonymous_G3757V |
| 11537 | A | G | | ORF1ab | nonsynonymous_I3758V |
| 11539 | T | C | G | ORF1ab | synonymous_I3758I;<br>nonsynonymous_I3758M |
| 11540 | G | T | | ORF1ab | nonsynonymous_V3759F |
| 11541 | T | G | | ORF1ab | nonsynonymous_V3759G |

| | | | | | |
|---|---|---|---|---|---|
| 11545 | T | C | | ORF1ab | synonymous_F3760F |
| 11563 | C | T | | ORF1ab | synonymous_C3766C |
| 11564 | C | T | | ORF1ab | nonsynonymous_P3767S |
| 11565 | C | T | | ORF1ab | nonsynonymous_P3767L |
| 11571 | T | C | | ORF1ab | nonsynonymous_F3769S |
| 11572 | C | T | | ORF1ab | synonymous_F3769F |
| 11575 | C | T | | ORF1ab | synonymous_F3770F |
| 11594 | C | T | | ORF1ab | stopgain_Q3777X |
| 11595 | A | G | | ORF1ab | nonsynonymous_Q3777R |
| 11596 | G | T | | ORF1ab | nonsynonymous_Q3777H |
| 11612 | T | A | | ORF1ab | nonsynonymous_Y3783N |
| 11620 | C | A | | ORF1ab | nonsynonymous_F3785L |
| 11634 | G | T | | ORF1ab | nonsynonymous_C3790F |
| 11638 | T | C | | ORF1ab | synonymous_T3791T |
| 11647 | T | C | | ORF1ab | synonymous_F3794F |
| 11650 | C | T | | ORF1ab | synonymous_G3795G |
| 11651 | C | T | | ORF1ab | nonsynonymous_L3796F |
| 11655 | T | C | | ORF1ab | nonsynonymous_F3797S |
| 11658 | G | T | | ORF1ab | nonsynonymous_C3798F |
| 11662 | A | T | | ORF1ab | nonsynonymous_L3799F |
| 11663 | C | T | | ORF1ab | nonsynonymous_L3800F |
| 11668 | C | T | | ORF1ab | synonymous_N3801N |
| 11669 | C | A | T | ORF1ab | nonsynonymous_R3802S;<br>nonsynonymous_R3802C |
| 11671 | C | T | | ORF1ab | synonymous_R3802R |
| 11674 | C | T | | ORF1ab | synonymous_Y3803Y |
| 11677 | T | C | | ORF1ab | synonymous_F3804F |
| 11687 | C | T | | ORF1ab | nonsynonymous_L3808F |
| 11693 | G | T | | ORF1ab | nonsynonymous_V3810F |
| 11699 | G | C | | ORF1ab | nonsynonymous_D3812H |
| 11704 | C | T | | ORF1ab | synonymous_Y3813Y |

| | | | | | |
|---|---|---|---|---|---|
| 11719 | G | T | | ORF1ab | nonsynonymous_Q3818H |
| 11731 | T | C | | ORF1ab | synonymous_Y3822Y |
| 11737 | T | C | | ORF1ab | synonymous_N3824N |
| 11741 | C | T | | ORF1ab | stopgain_Q3826X |
| 11747 | C | T | | ORF1ab | synonymous_L3828L |
| 11750 | C | T | | ORF1ab | nonsynonymous_L3829F |
| 11756 | C | T | | ORF1ab | nonsynonymous_P3831S |
| 11757 | C | T | | ORF1ab | nonsynonymous_P3831L |
| 11758 | C | T | | ORF1ab | synonymous_P3831P |
| 11761 | G | A | | ORF1ab | synonymous_K3832K |
| 11765 | A | T | | ORF1ab | nonsynonymous_S3834C |
| 11771 | G | T | | ORF1ab | nonsynonymous_D3836Y |
| 11775 | C | T | | ORF1ab | nonsynonymous_A3837V |
| 11779 | C | T | | ORF1ab | synonymous_F3838F |
| 11782 | A | G | | ORF1ab | synonymous_K3839K |
| 11801 | G | A | | ORF1ab | nonsynonymous_G3846S |
| 11812 | C | A | | ORF1ab | synonymous_G3849G |
| 11817 | C | T | | ORF1ab | nonsynonymous_P3851L |
| 11824 | C | T | | ORF1ab | synonymous_I3853I |
| 11827 | A | G | | ORF1ab | synonymous_K3854K |
| 11830 | A | T | | ORF1ab | synonymous_V3855V |
| 11832 | C | T | | ORF1ab | nonsynonymous_A3856V |
| 11833 | C | T | | ORF1ab | synonymous_A3856A |
| 11837 | G | T | | ORF1ab | nonsynonymous_V3858L |
| 11840 | C | T | G | ORF1ab | stopgain_Q3859X; nonsynonymous_Q3859E |
| 11862 | A | T | | ORF1ab | nonsynonymous_K3866M |
| 11863 | G | T | | ORF1ab | nonsynonymous_K3866N |
| 11866 | C | T | | ORF1ab | synonymous_C3867C |
| 11893 | G | T | | ORF1ab | nonsynonymous_L3876F |
| 11900 | C | T | | ORF1ab | nonsynonymous_L3879F |

| | | | | | |
|---|---|---|---|---|---|
| 11913 | C | T | | ORF1ab | nonsynonymous_S3883L |
| 11916 | C | T | | ORF1ab | nonsynonymous_S3884L |
| 11919 | C | T | | ORF1ab | nonsynonymous_S3885F |
| 11933 | C | T | | ORF1ab | stopgain_Q3890X |
| 11940 | T | C | | ORF1ab | nonsynonymous_V3892A |
| 11941 | C | T | | ORF1ab | synonymous_V3892V |
| 11948 | C | T | | ORF1ab | nonsynonymous_H3895Y |
| 11950 | C | T | | ORF1ab | synonymous_H3895H |
| 11955 | A | C | | ORF1ab | nonsynonymous_D3897A |
| 11956 | C | T | | ORF1ab | synonymous_D3897D |
| 11959 | T | C | | ORF1ab | synonymous_I3898I |
| 11962 | C | A | T | ORF1ab | synonymous_L3899L; synonymous_L3899L |
| 11967 | C | T | | ORF1ab | nonsynonymous_A3901V |
| 11976 | C | T | | ORF1ab | nonsynonymous_T3904I |
| 11984 | G | T | | ORF1ab | nonsynonymous_A3907S |
| 11991 | A | G | | ORF1ab | nonsynonymous_E3909G |
| 12008 | C | T | | ORF1ab | nonsynonymous_L3915F |
| 12010 | T | A | | ORF1ab | synonymous_L3915L |
| 12021 | T | C | | ORF1ab | nonsynonymous_L3919P |
| 12025 | C | T | | ORF1ab | synonymous_S3920S |
| 12027 | T | C | | ORF1ab | nonsynonymous_M3921T |
| 12029 | C | T | | ORF1ab | stopgain_Q3922X |
| 12031 | G | T | | ORF1ab | nonsynonymous_Q3922H |
| 12033 | G | T | | ORF1ab | nonsynonymous_G3923V |
| 12035 | G | C | | ORF1ab | nonsynonymous_A3924P |
| 12041 | G | T | | ORF1ab | nonsynonymous_D3926Y |
| 12049 | C | T | | ORF1ab | synonymous_N3928N |
| 12050 | A | T | | ORF1ab | stopgain_K3929X |
| 12052 | G | T | | ORF1ab | nonsynonymous_K3929N |
| 12053 | C | T | | ORF1ab | nonsynonymous_L3930F |

| | | | | | |
|---|---|---|---|---|---|
| 12062 | G | A | | ORF1ab | nonsynonymous_E3933K |
| 12066 | T | C | | ORF1ab | nonsynonymous_M3934T |
| 12068 | C | T | | ORF1ab | synonymous_L3935L |
| 12071 | G | A | | ORF1ab | nonsynonymous_D3936N |
| 12073 | C | T | | ORF1ab | synonymous_D3936D |
| 12076 | C | T | | ORF1ab | synonymous_N3937N |
| 12081 | C | T | | ORF1ab | nonsynonymous_A3939V |
| 12084 | C | T | | ORF1ab | nonsynonymous_T3940I |
| 12088 | A | C | | ORF1ab | nonsynonymous_L3941F |
| 12091 | A | T | | ORF1ab | nonsynonymous_Q3942H |
| 12092 | G | T | | ORF1ab | nonsynonymous_A3943S |
| 12099 | C | T | | ORF1ab | nonsynonymous_A3945V |
| 12104 | G | T | | ORF1ab | stopgain_E3947X |
| 12106 | G | T | | ORF1ab | nonsynonymous_E3947D |
| 12112 | T | C | | ORF1ab | synonymous_S3949S |
| 12115 | C | T | | ORF1ab | synonymous_S3950S |
| 12116 | C | T | | ORF1ab | nonsynonymous_L3951F |
| 12119 | C | T | | ORF1ab | nonsynonymous_P3952S |
| 12136 | T | C | | ORF1ab | synonymous_F3957F |
| 12141 | C | T | | ORF1ab | nonsynonymous_T3959I |
| 12148 | A | G | | ORF1ab | synonymous_Q3961Q |
| 12149 | G | A | | ORF1ab | nonsynonymous_E3962K |
| 12158 | G | T | | ORF1ab | stopgain_E3965X |
| 12164 | G | A | T | ORF1ab | nonsynonymous_A3967T;<br>nonsynonymous_A3967S |
| 12165 | C | T | | ORF1ab | nonsynonymous_A3967V |
| 12167 | G | A | | ORF1ab | nonsynonymous_V3968I |
| 12171 | C | T | | ORF1ab | nonsynonymous_A3969V |
| 12191 | G | T | | ORF1ab | nonsynonymous_V3976F |
| 12202 | G | T | | ORF1ab | nonsynonymous_K3979N |
| 12212 | T | C | | ORF1ab | nonsynonymous_S3983P |

| | | | | | |
|---|---|---|---|---|---|
| 12228 | A | G | | ORF1ab | nonsynonymous_K3988R |
| 12233 | G | A | | ORF1ab | nonsynonymous_E3990K |
| 12239 | G | C | | ORF1ab | nonsynonymous_D3992H |
| 12242 | C | T | | ORF1ab | nonsynonymous_R3993C |
| 12243 | G | T | | ORF1ab | nonsynonymous_R3993L |
| 12248 | G | T | | ORF1ab | nonsynonymous_A3995S |
| 12249 | C | T | | ORF1ab | nonsynonymous_A3995V |
| 12253 | C | T | | ORF1ab | synonymous_A3996A |
| 12254 | A | G | | ORF1ab | nonsynonymous_M3997V |
| 12257 | C | T | | ORF1ab | stopgain_Q3998X |
| 12260 | C | T | | ORF1ab | nonsynonymous_R3999C |
| 12261 | G | A | | ORF1ab | nonsynonymous_R3999H |
| 12268 | G | C | | ORF1ab | nonsynonymous_L4001F |
| 12269 | G | A | | ORF1ab | nonsynonymous_E4002K |
| 12273 | A | T | | ORF1ab | nonsynonymous_K4003M |
| 12274 | G | A | T | ORF1ab | synonymous_K4003K;<br>nonsynonymous_K4003N |
| 12280 | T | C | | ORF1ab | synonymous_A4005A |
| 12287 | G | A | | ORF1ab | nonsynonymous_A4008T |
| 12293 | A | C | | ORF1ab | nonsynonymous_T4010P |
| 12294 | C | T | | ORF1ab | nonsynonymous_T4010I |
| 12298 | A | G | | ORF1ab | synonymous_Q4011Q |
| 12302 | T | C | | ORF1ab | nonsynonymous_Y4013H |
| 12307 | A | C | | ORF1ab | nonsynonymous_K4014N |
| 12310 | G | T | | ORF1ab | nonsynonymous_Q4015H |
| 12312 | C | T | | ORF1ab | nonsynonymous_A4016V |
| 12322 | G | A | T | ORF1ab | synonymous_E4019E;<br>nonsynonymous_E4019D |
| 12323 | G | T | | ORF1ab | nonsynonymous_D4020Y |
| 12325 | C | T | | ORF1ab | synonymous_D4020D |

| | | | | | |
|---|---|---|---|---|---|
| 12331 | G | A | T | ORF1ab | synonymous_R4022R; nonsynonymous_R4022S |
| 12333 | C | T | | ORF1ab | nonsynonymous_A4023V |
| 12334 | A | T | | ORF1ab | synonymous_A4023A |
| 12347 | G | T | | ORF1ab | nonsynonymous_A4028S |
| 12348 | C | T | | ORF1ab | nonsynonymous_A4028V |
| 12353 | C | T | | ORF1ab | stopgain_Q4030X |
| 12354 | A | G | | ORF1ab | nonsynonymous_Q4030R |
| 12357 | C | T | | ORF1ab | nonsynonymous_T4031I |
| 12362 | C | T | | ORF1ab | nonsynonymous_L4033F |
| 12364 | T | C | | ORF1ab | synonymous_L4033L |
| 12367 | C | T | | ORF1ab | synonymous_F4034F |
| 12369 | C | T | | ORF1ab | nonsynonymous_T4035I |
| 12374 | C | T | | ORF1ab | nonsynonymous_L4037F |
| 12377 | A | T | | ORF1ab | stopgain_R4038X |
| 12379 | A | T | | ORF1ab | nonsynonymous_R4038S |
| 12382 | G | A | | ORF1ab | synonymous_K4039K |
| 12390 | A | G | | ORF1ab | nonsynonymous_N4042S |
| 12397 | A | T | | ORF1ab | synonymous_A4044A |
| 12398 | C | T | | ORF1ab | nonsynonymous_L4045F |
| 12401 | A | G | | ORF1ab | nonsynonymous_N4046D |
| 12406 | C | T | | ORF1ab | synonymous_N4047N |
| 12412 | C | T | | ORF1ab | synonymous_I4049I |
| 12414 | A | T | | ORF1ab | nonsynonymous_N4050I |
| 12415 | C | T | | ORF1ab | synonymous_N4050N |
| 12416 | A | T | G | ORF1ab | nonsynonymous_N4051Y; nonsynonymous_N4051D |
| 12420 | C | T | | ORF1ab | nonsynonymous_A4052V |
| 12438 | C | T | | ORF1ab | nonsynonymous_P4058L |
| 12439 | C | T | | ORF1ab | synonymous_P4058P |
| 12453 | C | T | | ORF1ab | nonsynonymous_P4063L |

| | | | | | |
|---|---|---|---|---|---|
| 12455 | C | T | | ORF1ab | nonsynonymous_L4064F |
| 12456 | T | C | | ORF1ab | nonsynonymous_L4064P |
| 12465 | C | T | | ORF1ab | nonsynonymous_A4067V |
| 12467 | G | A | | ORF1ab | nonsynonymous_A4068T |
| 12468 | C | T | | ORF1ab | nonsynonymous_A4068V |
| 12473 | C | T | | ORF1ab | synonymous_L4070L |
| 12478 | G | A | | ORF1ab | nonsynonymous_M4071I |
| 12486 | T | C | | ORF1ab | nonsynonymous_I4074T |
| 12490 | A | T | | ORF1ab | synonymous_P4075P |
| 12491 | G | T | | ORF1ab | nonsynonymous_D4076Y |
| 12501 | C | T | | ORF1ab | nonsynonymous_T4079I |
| 12505 | T | C | | ORF1ab | synonymous_Y4080Y |
| 12507 | A | G | | ORF1ab | nonsynonymous_K4081R |
| 12513 | C | T | | ORF1ab | nonsynonymous_T4083M |
| 12516 | G | C | | ORF1ab | nonsynonymous_C4084S |
| 12525 | C | T | | ORF1ab | nonsynonymous_T4087I |
| 12528 | C | T | | ORF1ab | nonsynonymous_T4088I |
| 12534 | C | T | | ORF1ab | nonsynonymous_T4090I |
| 12546 | C | T | | ORF1ab | nonsynonymous_A4094V |
| 12551 | T | G | | ORF1ab | nonsynonymous_W4096G |
| 12554 | G | A | | ORF1ab | nonsynonymous_E4097K |
| 12571 | A | G | | ORF1ab | synonymous_V4102V |
| 12578 | G | T | | ORF1ab | nonsynonymous_D4105Y |
| 12579 | A | T | | ORF1ab | nonsynonymous_D4105V |
| 12589 | T | C | | ORF1ab | synonymous_I4108I |
| 12598 | T | C | | ORF1ab | synonymous_L4111L |
| 12600 | G | T | | ORF1ab | nonsynonymous_S4112I |
| 12602 | G | A | | ORF1ab | nonsynonymous_E4113K |
| 12614 | G | T | | ORF1ab | nonsynonymous_D4117Y |
| 12616 | C | T | | ORF1ab | synonymous_D4117D |
| 12619 | T | C | | ORF1ab | synonymous_N4118N |
| 12621 | C | T | | ORF1ab | nonsynonymous_S4119L |

| | | | | | |
|---|---|---|---|---|---|
| 12623 | C | T | | ORF1ab | nonsynonymous_P4120S |
| 12630 | T | C | | ORF1ab | nonsynonymous_L4122S |
| 12647 | G | A | | ORF1ab | nonsynonymous_V4128I |
| 12654 | C | T | | ORF1ab | nonsynonymous_A4130V |
| 12660 | G | A | | ORF1ab | nonsynonymous_R4132K |
| 12661 | G | T | | ORF1ab | nonsynonymous_R4132S |
| 12664 | C | A | T | ORF1ab | synonymous_A4133A; synonymous_A4133A |
| 12666 | A | C | | ORF1ab | nonsynonymous_N4134T |
| 12667 | T | A | C | ORF1ab | nonsynonymous_N4134K; synonymous_N4134N |
| 12668 | T | C | | ORF1ab | nonsynonymous_S4135P |
| 12669 | C | T | G | ORF1ab | nonsynonymous_S4135F; nonsynonymous_S4135C |
| 12672 | C | T | | ORF1ab | nonsynonymous_A4136V |
| 12676 | C | T | | ORF1ab | synonymous_V4137V |
| 12685 | G | T | | ORF1ab | nonsynonymous_Q4140H |
| 12694 | G | T | | ORF1ab | nonsynonymous_E4143D |
| 12710 | C | T | | ORF1ab | synonymous_L4149L |
| 12713 | C | T | | ORF1ab | stopgain_R4150X |
| 12716 | C | T | | ORF1ab | stopgain_Q4151X |
| 12717 | A | T | | ORF1ab | nonsynonymous_Q4151L |
| 12719 | A | G | | ORF1ab | nonsynonymous_M4152V |
| 12729 | C | T | | ORF1ab | nonsynonymous_A4155V |
| 12732 | C | T | | ORF1ab | nonsynonymous_A4156V |
| 12737 | A | C | | ORF1ab | nonsynonymous_T4158P |
| 12738 | C | T | | ORF1ab | nonsynonymous_T4158I |
| 12741 | C | T | | ORF1ab | nonsynonymous_T4159I |
| 12750 | C | T | | ORF1ab | nonsynonymous_A4162V |
| 12753 | G | C | | ORF1ab | nonsynonymous_C4163S |
| 12754 | C | T | | ORF1ab | synonymous_C4163C |

| | | | | | |
|---|---|---|---|---|---|
| 12757 | T | C | | ORF1ab | synonymous_T4164T |
| 12768 | C | T | | ORF1ab | nonsynonymous_A4168V |
| 12769 | G | T | | ORF1ab | synonymous_A4168A |
| 12778 | C | T | | ORF1ab | synonymous_Y4171Y |
| 12781 | C | T | | ORF1ab | synonymous_Y4172Y |
| 12784 | C | T | | ORF1ab | synonymous_N4173N |
| 12789 | C | T | | ORF1ab | nonsynonymous_T4175I |
| 12791 | A | T | | ORF1ab | stopgain_K4176X |
| 12793 | G | T | | ORF1ab | nonsynonymous_K4176N |
| 12796 | A | T | | ORF1ab | synonymous_G4177G |
| 12800 | A | C | | ORF1ab | synonymous_R4179R |
| 12802 | G | T | | ORF1ab | nonsynonymous_R4179S |
| 12806 | G | A | | ORF1ab | nonsynonymous_V4181I |
| 12808 | A | T | | ORF1ab | synonymous_V4181V |
| 12822 | C | T | | ORF1ab | nonsynonymous_S4186F |
| 12824 | G | A | | ORF1ab | nonsynonymous_D4187N |
| 12832 | G | A | | ORF1ab | synonymous_Q4189Q |
| 12833 | G | T | | ORF1ab | nonsynonymous_D4190Y |
| 12838 | G | A | | ORF1ab | synonymous_L4191L |
| 12844 | G | T | | ORF1ab | nonsynonymous_W4193C |
| 12845 | G | A | | ORF1ab | nonsynonymous_A4194T |
| 12850 | A | T | | ORF1ab | nonsynonymous_R4195S |
| 12854 | C | T | | ORF1ab | nonsynonymous_P4197S |
| 12855 | C | T | | ORF1ab | nonsynonymous_P4197L |
| 12863 | G | T | | ORF1ab | nonsynonymous_D4200Y |
| 12868 | A | C | | ORF1ab | synonymous_G4201G |
| 12875 | A | G | | ORF1ab | nonsynonymous_T4204A |
| 12884 | A | T | | ORF1ab | nonsynonymous_T4207S |
| 12885 | C | T | | ORF1ab | nonsynonymous_T4207I |
| 12886 | A | C | | ORF1ab | synonymous_T4207T |
| 12890 | C | T | | ORF1ab | synonymous_L4209L |
| 12896 | C | T | | ORF1ab | nonsynonymous_P4211S |

| | | | | | |
|---|---|---|---|---|---|
| 12899 | C | T | | ORF1ab | nonsynonymous_P4212S |
| 12900 | C | T | | ORF1ab | nonsynonymous_P4212L |
| 12903 | G | T | | ORF1ab | nonsynonymous_C4213F |
| 12907 | G | T | | ORF1ab | nonsynonymous_R4214S |
| 12915 | C | T | | ORF1ab | nonsynonymous_T4217I |
| 12920 | A | C | | ORF1ab | nonsynonymous_T4219P |
| 12921 | C | T | | ORF1ab | nonsynonymous_T4219I |
| 12924 | C | T | | ORF1ab | nonsynonymous_P4220L |
| 12928 | A | G | | ORF1ab | synonymous_K4221K |
| 12929 | G | T | | ORF1ab | nonsynonymous_G4222C |
| 12930 | G | T | | ORF1ab | nonsynonymous_G4222V |
| 12933 | C | T | | ORF1ab | nonsynonymous_P4223L |
| 12937 | A | T | | ORF1ab | nonsynonymous_K4224N |
| 12938 | G | C | | ORF1ab | nonsynonymous_V4225L |
| 12952 | C | T | | ORF1ab | synonymous_Y4229Y |
| 12963 | G | C | | ORF1ab | nonsynonymous_G4233A |
| 12966 | T | A | | ORF1ab | stopgain_L4234X |
| 12970 | C | T | | ORF1ab | synonymous_N4235N |
| 12973 | C | T | | ORF1ab | synonymous_N4236N |
| 12989 | G | T | | ORF1ab | nonsynonymous_V4242L |
| 13004 | G | T | | ORF1ab | nonsynonymous_A4247S |
| 13005 | C | T | | ORF1ab | nonsynonymous_A4247V |
| 13006 | T | C | | ORF1ab | synonymous_A4247A |
| 13009 | C | T | | ORF1ab | synonymous_A4248A |
| 13011 | C | T | | ORF1ab | nonsynonymous_T4249I |
| 13013 | G | T | | ORF1ab | nonsynonymous_V4250L |
| 13018 | T | C | | ORF1ab | synonymous_R4251R |
| 13019 | C | T | | ORF1ab | synonymous_L4252L |
| 13022 | C | T | | ORF1ab | stopgain_Q4253X |
| 13029 | G | T | | ORF1ab | nonsynonymous_G4255V |
| 13035 | C | T | | ORF1ab | nonsynonymous_A4257V |
| 13040 | G | T | | ORF1ab | stopgain_E4259X |

| | | | | | |
|---|---|---|---|---|---|
| 13043 | G | T | | ORF1ab | nonsynonymous_V4260L |
| 13047 | C | T | | ORF1ab | nonsynonymous_P4261L |
| 13050 | C | T | | ORF1ab | nonsynonymous_A4262V |
| 13059 | C | T | | ORF1ab | nonsynonymous_T4265I |
| 13066 | A | T | | ORF1ab | nonsynonymous_L4267F |
| 13082 | G | C | | ORF1ab | nonsynonymous_A4273P |
| 13083 | C | T | | ORF1ab | nonsynonymous_A4273V |
| 13089 | A | C | | ORF1ab | nonsynonymous_D4275A |
| 13090 | T | C | | ORF1ab | synonymous_D4275D |
| 13092 | C | T | | ORF1ab | nonsynonymous_A4276V |
| 13093 | T | C | | ORF1ab | synonymous_A4276A |
| 13115 | C | T | | ORF1ab | synonymous_L4284L |
| 13118 | G | T | | ORF1ab | nonsynonymous_A4285S |
| 13119 | C | T | | ORF1ab | nonsynonymous_A4285V |
| 13123 | T | C | | ORF1ab | synonymous_S4286S |
| 13133 | C | T | | ORF1ab | nonsynonymous_P4290S |
| 13134 | C | A | T | ORF1ab | nonsynonymous_P4290Q; nonsynonymous_P4290L |
| 13137 | T | C | | ORF1ab | nonsynonymous_I4291T |
| 13138 | C | T | | ORF1ab | synonymous_I4291I |
| 13140 | C | T | | ORF1ab | nonsynonymous_T4292I |
| 13148 | G | A | | ORF1ab | nonsynonymous_V4295I |
| 13157 | T | G | | ORF1ab | nonsynonymous_L4298V |
| 13161 | G | T | | ORF1ab | nonsynonymous_C4299F |
| 13166 | C | T | | ORF1ab | nonsynonymous_H4301Y |
| 13168 | C | T | | ORF1ab | synonymous_H4301H |
| 13175 | A | G | | ORF1ab | nonsynonymous_T4304A |
| 13176 | C | T | | ORF1ab | nonsynonymous_T4304I |
| 13188 | T | C | | ORF1ab | nonsynonymous_I4308T |
| 13191 | C | T | | ORF1ab | nonsynonymous_T4309I |
| 13193 | G | T | | ORF1ab | nonsynonymous_V4310F |

| | | | | | |
|---|---|---|---|---|---|
| 13197 | C | T | | ORF1ab | nonsynonymous_T4311I |
| 13199 | C | T | | ORF1ab | nonsynonymous_P4312S |
| 13201 | G | T | | ORF1ab | synonymous_P4312P |
| 13210 | T | C | | ORF1ab | synonymous_N4315N |
| 13212 | T | C | | ORF1ab | nonsynonymous_M4316T |
| 13216 | T | G | | ORF1ab | nonsynonymous_D4317E |
| 13228 | T | C | | ORF1ab | synonymous_F4321F |
| 13239 | C | T | | ORF1ab | nonsynonymous_S4325L |
| 13240 | G | T | | ORF1ab | synonymous_S4325S |
| 13244 | T | G | | ORF1ab | nonsynonymous_C4327G |
| 13248 | T | G | | ORF1ab | nonsynonymous_L4328R |
| 13253 | T | C | | ORF1ab | nonsynonymous_C4330R |
| 13254 | G | T | | ORF1ab | nonsynonymous_C4330F |
| 13256 | C | T | | ORF1ab | nonsynonymous_R4331C |
| 13259 | T | C | | ORF1ab | nonsynonymous_C4332R |
| 13262 | C | T | | ORF1ab | nonsynonymous_H4333Y |
| 13264 | C | T | | ORF1ab | synonymous_H4333H |
| 13270 | T | C | | ORF1ab | synonymous_D4335D |
| 13274 | C | T | | ORF1ab | nonsynonymous_P4337S |
| 13280 | C | T | | ORF1ab | nonsynonymous_P4339S |
| 13289 | T | A | | ORF1ab | nonsynonymous_F4342I |
| 13299 | T | A | | ORF1ab | stopgain_L4345X |
| 13305 | G | T | | ORF1ab | nonsynonymous_G4347V |
| 13307 | A | T | | ORF1ab | stopgain_K4348X |
| 13314 | T | C | | ORF1ab | nonsynonymous_V4350A |
| 13316 | C | T | | ORF1ab | stopgain_Q4351X |
| 13318 | A | G | | ORF1ab | synonymous_Q4351Q |
| 13322 | C | T | | ORF1ab | nonsynonymous_P4353S |
| 13323 | C | A | T | ORF1ab | nonsynonymous_P4353H; nonsynonymous_P4353L |
| 13326 | C | T | | ORF1ab | nonsynonymous_T4354I |

| | | | | | |
|---|---|---|---|---|---|
| 13329 | C | T | | ORF1ab | nonsynonymous_T4355I |
| 13331 | T | G | | ORF1ab | nonsynonymous_C4356G |
| 13335 | C | T | | ORF1ab | nonsynonymous_A4357V |
| 13344 | C | T | | ORF1ab | nonsynonymous_P4360L |
| 13345 | T | C | | ORF1ab | synonymous_P4360P |
| 13346 | G | T | | ORF1ab | nonsynonymous_V4361L |
| 13348 | G | T | | ORF1ab | synonymous_V4361V |
| 13356 | C | T | | ORF1ab | nonsynonymous_T4364I |
| 13358 | C | T | | ORF1ab | nonsynonymous_L4365F |
| 13366 | C | A | T | ORF1ab | nonsynonymous_N4367K; synonymous_N4367N |
| 13391 | T | C | | ORF1ab | nonsynonymous_W4376R |
| 13394 | A | G | | ORF1ab | nonsynonymous_K4377E |
| 13409 | A | T | | ORF1ab | nonsynonymous_S4382C |
| 13413 | G | C | | ORF1ab | nonsynonymous_C4383S |
| 13421 | C | T | | ORF1ab | nonsynonymous_L4386F |
| 13422 | T | C | | ORF1ab | nonsynonymous_L4386P |
| 13432 | C | T | | ORF1ab | synonymous_P4389P |
| 13437 | T | A | | ORF1ab | nonsynonymous_L4391H |
| 13443 | C | T | | ORF1ab | nonsynonymous_S4393L |
| 13458 | C | T | | ORF1ab | nonsynonymous_S4398L |
| 13460 | T | C | | ORF1ab | nonsynonymous_F4399L |
| 13464 | T | A | | ORF1ab | stopgain_L4400X |
| 13471 | G | T | | ORF1ab | nonsynonymous_V4403F |
| 13476 | C | T | | ORF1ab | nonsynonymous_A4404V |
| 13487 | C | T | | ORF1ab | nonsynonymous_A4408V |
| 13490 | C | T | | ORF1ab | nonsynonymous_A4409V |
| 13491 | C | T | | ORF1ab | synonymous_A4409A |
| 13495 | C | T | | ORF1ab | nonsynonymous_L4411F |
| 13501 | C | T | | ORF1ab | nonsynonymous_P4413S |
| 13506 | C | T | | ORF1ab | synonymous_C4414C |

| | | | | | |
|---|---|---|---|---|---|
| 13509 | C | T | | ORF1ab | synonymous_G4415G |
| 13511 | C | T | | ORF1ab | nonsynonymous_T4416I |
| 13512 | A | G | | ORF1ab | synonymous_T4416T |
| 13517 | C | T | | ORF1ab | nonsynonymous_T4418I |
| 13536 | C | T | | ORF1ab | synonymous_Y4424Y |
| 13537 | A | G | | ORF1ab | nonsynonymous_R4425G |
| 13545 | T | C | | ORF1ab | synonymous_F4427F |
| 13548 | C | T | | ORF1ab | synonymous_D4428D |
| 13551 | C | A | T | ORF1ab | synonymous_I4429I; synonymous_I4429I |
| 13568 | C | T | | ORF1ab | nonsynonymous_A4435V |
| 13569 | T | A | | ORF1ab | synonymous_A4435A |
| 13571 | G | T | | ORF1ab | nonsynonymous_G4436V |
| 13576 | G | A | | ORF1ab | nonsynonymous_A4438T |
| 13578 | T | A | | ORF1ab | synonymous_A4438A |
| 13612 | G | A | | ORF1ab | nonsynonymous_E4450K |
| 13613 | A | G | | ORF1ab | nonsynonymous_E4450G |
| 13617 | G | T | | ORF1ab | nonsynonymous_K4451N |
| 13623 | A | G | | ORF1ab | synonymous_E4453E |
| 13624 | G | A | T | ORF1ab | nonsynonymous_D4454N; nonsynonymous_D4454Y |
| 13625 | A | T | | ORF1ab | nonsynonymous_D4454V |
| 13627 | G | T | | ORF1ab | nonsynonymous_D4455Y |
| 13629 | C | T | | ORF1ab | synonymous_D4455D |
| 13631 | A | G | | ORF1ab | nonsynonymous_N4456S |
| 13638 | T | A | C | ORF1ab | synonymous_I4458I; synonymous_I4458I |
| 13640 | A | T | | ORF1ab | nonsynonymous_D4459V |
| 13643 | C | T | | ORF1ab | nonsynonymous_S4460F |
| 13647 | C | T | | ORF1ab | synonymous_Y4461Y |
| 13660 | A | G | | ORF1ab | nonsynonymous_R4466G |

| | | | | | |
|---|---|---|---|---|---|
| 13663 | C | T | | ORF1ab | nonsynonymous_H4467Y |
| 13665 | C | T | | ORF1ab | synonymous_H4467H |
| 13666 | A | C | T | ORF1ab | nonsynonymous_T4468P;<br>nonsynonymous_T4468S |
| 13677 | C | T | | ORF1ab | synonymous_N4471N |
| 13694 | C | T | | ORF1ab | nonsynonymous_T4477I |
| 13701 | T | C | | ORF1ab | synonymous_Y4479Y |
| 13708 | C | T | | ORF1ab | nonsynonymous_L4482F |
| 13721 | C | T | | ORF1ab | nonsynonymous_P4486L |
| 13724 | C | T | | ORF1ab | nonsynonymous_A4487V |
| 13730 | C | T | | ORF1ab | nonsynonymous_A4489V |
| 13738 | G | T | | ORF1ab | nonsynonymous_D4492Y |
| 13740 | C | T | | ORF1ab | synonymous_D4492D |
| 13743 | C | T | | ORF1ab | synonymous_F4493F |
| 13760 | A | C | | ORF1ab | nonsynonymous_D4499A |
| 13775 | C | T | | ORF1ab | nonsynonymous_P4504L |
| 13777 | C | T | | ORF1ab | nonsynonymous_H4505Y |
| 13779 | T | C | | ORF1ab | synonymous_H4505H |
| 13780 | A | T | G | ORF1ab | nonsynonymous_I4506L;<br>nonsynonymous_I4506V |
| 13784 | C | T | | ORF1ab | nonsynonymous_S4507L |
| 13799 | C | T | | ORF1ab | nonsynonymous_T4512I |
| 13819 | C | A | | ORF1ab | nonsynonymous_L4519I |
| 13822 | G | A | | ORF1ab | nonsynonymous_V4520I |
| 13824 | C | T | | ORF1ab | synonymous_V4520V |
| 13835 | G | T | | ORF1ab | nonsynonymous_R4524M |
| 13850 | G | T | | ORF1ab | nonsynonymous_G4529V |
| 13857 | T | C | | ORF1ab | synonymous_C4531C |
| 13858 | G | T | | ORF1ab | nonsynonymous_D4532Y |
| 13859 | A | G | | ORF1ab | nonsynonymous_D4532G |
| 13862 | C | T | | ORF1ab | nonsynonymous_T4533I |

| | | | | | |
|---|---|---|---|---|---|
| 13870 | G | A | | ORF1ab | nonsynonymous_E4536K |
| 13882 | A | C | | ORF1ab | nonsynonymous_T4540P |
| 13885 | T | C | | ORF1ab | nonsynonymous_Y4541H |
| 13890 | T | C | | ORF1ab | synonymous_N4542N |
| 13896 | T | C | | ORF1ab | synonymous_C4544C |
| 13897 | G | C | | ORF1ab | nonsynonymous_D4545H |
| 13900 | G | T | | ORF1ab | nonsynonymous_D4546Y |
| 13901 | A | G | | ORF1ab | nonsynonymous_D4546G |
| 13911 | C | A | | ORF1ab | nonsynonymous_F4549L |
| 13914 | T | C | | ORF1ab | synonymous_N4550N |
| 13917 | A | G | | ORF1ab | synonymous_K4551K |
| 13920 | G | T | | ORF1ab | nonsynonymous_K4552N |
| 13923 | C | T | | ORF1ab | synonymous_D4553D |
| 13927 | T | G | | ORF1ab | nonsynonymous_Y4555D |
| 13928 | A | C | | ORF1ab | nonsynonymous_Y4555S |
| 13931 | A | T | | ORF1ab | nonsynonymous_D4556V |
| 13936 | G | T | | ORF1ab | nonsynonymous_V4558L |
| 13940 | A | G | | ORF1ab | nonsynonymous_E4559G |
| 13943 | A | G | | ORF1ab | nonsynonymous_N4560S |
| 13951 | A | T | | ORF1ab | nonsynonymous_I4563L |
| 13957 | C | T | | ORF1ab | nonsynonymous_R4565C |
| 13966 | G | A | | ORF1ab | nonsynonymous_A4568T |
| 13981 | C | T | | ORF1ab | nonsynonymous_R4573C |
| 13994 | C | T | | ORF1ab | nonsynonymous_A4577V |
| 13998 | G | T | | ORF1ab | nonsynonymous_L4578F |
| 14004 | A | G | | ORF1ab | synonymous_K4580K |
| 14007 | A | G | | ORF1ab | synonymous_T4581T |
| 14008 | G | A | | ORF1ab | nonsynonymous_V4582I |
| 14015 | T | C | | ORF1ab | nonsynonymous_F4584S |
| 14028 | G | T | | ORF1ab | nonsynonymous_M4588I |
| 14044 | G | T | | ORF1ab | nonsynonymous_V4594F |
| 14047 | G | A | | ORF1ab | nonsynonymous_G4595S |

| | | | | | |
|---|---|---|---|---|---|
| 14055 | G | T | | ORF1ab | synonymous_L4597L |
| 14069 | A | G | | ORF1ab | nonsynonymous_Q4602R |
| 14071 | G | A | | ORF1ab | nonsynonymous_D4603N |
| 14076 | C | T | | ORF1ab | synonymous_L4604L |
| 14081 | G | C | | ORF1ab | nonsynonymous_G4606A |
| 14087 | G | T | | ORF1ab | nonsynonymous_W4608L |
| 14089 | T | C | | ORF1ab | nonsynonymous_Y4609H |
| 14104 | T | A | | ORF1ab | nonsynonymous_F4614I |
| 14115 | C | T | | ORF1ab | synonymous_T4617T |
| 14119 | C | T | | ORF1ab | nonsynonymous_P4619S |
| 14120 | C | T | | ORF1ab | nonsynonymous_P4619L |
| 14122 | G | T | | ORF1ab | nonsynonymous_G4620C |
| 14125 | A | C | | ORF1ab | nonsynonymous_S4621R |
| 14126 | G | A | T | ORF1ab | nonsynonymous_S4621N; nonsynonymous_S4621I |
| 14132 | T | C | | ORF1ab | nonsynonymous_V4623A |
| 14144 | A | T | | ORF1ab | nonsynonymous_D4627V |
| 14158 | T | C | | ORF1ab | synonymous_L4632L |
| 14160 | G | T | | ORF1ab | nonsynonymous_L4632F |
| 14167 | C | T | | ORF1ab | nonsynonymous_P4635S |
| 14174 | T | C | | ORF1ab | nonsynonymous_L4637S |
| 14177 | C | T | | ORF1ab | nonsynonymous_T4638I |
| 14183 | C | T | | ORF1ab | nonsynonymous_T4640I |
| 14184 | C | T | | ORF1ab | synonymous_T4640T |
| 14187 | G | A | | ORF1ab | synonymous_R4641R |
| 14188 | G | C | T | ORF1ab | nonsynonymous_A4642P; nonsynonymous_A4642S |
| 14195 | C | A | T | ORF1ab | nonsynonymous_T4644N; nonsynonymous_T4644I |
| 14197 | G | T | | ORF1ab | nonsynonymous_A4645S |
| 14206 | C | T | | ORF1ab | nonsynonymous_H4648Y |

| | | | | | |
|---|---|---|---|---|---|
| 14216 | C | T | | ORF1ab | nonsynonymous_T4651I |
| 14224 | A | G | | ORF1ab | nonsynonymous_T4654A |
| 14225 | C | A | | ORF1ab | nonsynonymous_T4654K |
| 14233 | T | A | | ORF1ab | nonsynonymous_Y4657N |
| 14235 | C | T | | ORF1ab | synonymous_Y4657Y |
| 14238 | T | C | | ORF1ab | synonymous_I4658I |
| 14240 | A | T | | ORF1ab | nonsynonymous_K4659M |
| 14241 | G | T | | ORF1ab | nonsynonymous_K4659N |
| 14245 | G | C | | ORF1ab | nonsynonymous_D4661H |
| 14262 | C | T | | ORF1ab | synonymous_D4666D |
| 14272 | G | T | | ORF1ab | stopgain_E4670X |
| 14274 | G | C | T | ORF1ab | nonsynonymous_E4670D; nonsynonymous_E4670D |
| 14277 | G | T | | ORF1ab | nonsynonymous_R4671S |
| 14278 | T | A | | ORF1ab | nonsynonymous_L4672I |
| 14279 | T | A | | ORF1ab | stopgain_L4672X |
| 14290 | G | T | | ORF1ab | nonsynonymous_D4676Y |
| 14307 | T | C | | ORF1ab | synonymous_Y4681Y |
| 14313 | T | C | | ORF1ab | synonymous_D4683D |
| 14314 | C | T | | ORF1ab | stopgain_Q4684X |
| 14317 | A | G | | ORF1ab | nonsynonymous_T4685A |
| 14318 | C | T | | ORF1ab | nonsynonymous_T4685I |
| 14320 | T | C | | ORF1ab | nonsynonymous_Y4686H |
| 14322 | C | T | | ORF1ab | synonymous_Y4686Y |
| 14323 | C | T | G | ORF1ab | nonsynonymous_H4687Y; nonsynonymous_H4687D |
| 14326 | C | T | | ORF1ab | nonsynonymous_P4688S |
| 14327 | C | A | | ORF1ab | nonsynonymous_P4688Q |
| 14328 | A | G | | ORF1ab | synonymous_P4688P |
| 14331 | T | A | | ORF1ab | nonsynonymous_N4689K |
| 14332 | T | A | | ORF1ab | nonsynonymous_C4690S |

| | | | | | |
|---|---|---|---|---|---|
| 14346 | G | T | | ORF1ab | nonsynonymous_L4694F |
| 14347 | G | T | | ORF1ab | nonsynonymous_D4695Y |
| 14352 | C | T | | ORF1ab | synonymous_D4696D |
| 14354 | G | T | | ORF1ab | nonsynonymous_R4697I |
| 14358 | C | T | | ORF1ab | synonymous_C4698C |
| 14364 | G | T | | ORF1ab | synonymous_L4700L |
| 14365 | C | T | | ORF1ab | nonsynonymous_H4701Y |
| 14369 | G | T | | ORF1ab | nonsynonymous_C4702F |
| 14372 | C | T | | ORF1ab | nonsynonymous_A4703V |
| 14375 | A | G | | ORF1ab | nonsynonymous_N4704S |
| 14382 | T | C | | ORF1ab | synonymous_N4706N |
| 14383 | G | A | T | ORF1ab | nonsynonymous_V4707I; nonsynonymous_V4707F |
| 14391 | C | T | | ORF1ab | synonymous_F4709F |
| 14393 | C | T | | ORF1ab | nonsynonymous_S4710F |
| 14396 | C | T | | ORF1ab | nonsynonymous_T4711I |
| 14404 | C | T | | ORF1ab | nonsynonymous_P4714S |
| 14405 | C | T | | ORF1ab | nonsynonymous_P4714L |
| 14406 | A | T | | ORF1ab | synonymous_P4714P |
| 14408 | C | T | | ORF1ab | nonsynonymous_P4715L |
| 14411 | C | T | | ORF1ab | nonsynonymous_T4716I |
| 14412 | A | C | | ORF1ab | synonymous_T4716T |
| 14414 | G | T | | ORF1ab | nonsynonymous_S4717I |
| 14418 | T | G | | ORF1ab | nonsynonymous_F4718L |
| 14419 | G | C | T | ORF1ab | nonsynonymous_G4719R; stopgain_G4719X |
| 14424 | A | G | | ORF1ab | synonymous_P4720P |
| 14425 | C | A | T | ORF1ab | nonsynonymous_L4721I; synonymous_L4721L |
| 14428 | G | T | | ORF1ab | nonsynonymous_V4722L |
| 14432 | G | A | | ORF1ab | nonsynonymous_R4723K |

| | | | | | |
|---|---|---|---|---|---|
| 14436 | A | T | | ORF1ab | nonsynonymous_K4724N |
| 14458 | T | G | | ORF1ab | nonsynonymous_F4732V |
| 14461 | G | T | | ORF1ab | nonsynonymous_V4733L |
| 14466 | T | G | | ORF1ab | synonymous_V4734V |
| 14468 | C | T | | ORF1ab | nonsynonymous_S4735L |
| 14475 | A | G | | ORF1ab | synonymous_G4737G |
| 14477 | A | C | | ORF1ab | nonsynonymous_Y4738S |
| 14478 | C | T | | ORF1ab | synonymous_Y4738Y |
| 14484 | C | T | | ORF1ab | synonymous_F4740F |
| 14486 | G | T | | ORF1ab | nonsynonymous_R4741I |
| 14488 | G | C | | ORF1ab | nonsynonymous_E4742Q |
| 14495 | G | T | | ORF1ab | nonsynonymous_G4744V |
| 14500 | G | T | | ORF1ab | nonsynonymous_V4746L |
| 14507 | A | G | | ORF1ab | nonsynonymous_N4748S |
| 14520 | C | T | | ORF1ab | synonymous_N4752N |
| 14528 | G | T | | ORF1ab | nonsynonymous_S4755I |
| 14536 | C | T | | ORF1ab | nonsynonymous_L4758F |
| 14548 | G | A | | ORF1ab | nonsynonymous_E4762K |
| 14555 | T | C | | ORF1ab | nonsynonymous_L4764P |
| 14571 | C | T | | ORF1ab | synonymous_D4769D |
| 14573 | C | T | | ORF1ab | nonsynonymous_P4770L |
| 14583 | C | T | | ORF1ab | synonymous_H4773H |
| 14584 | G | T | | ORF1ab | nonsynonymous_A4774S |
| 14585 | C | T | | ORF1ab | nonsynonymous_A4774V |
| 14611 | A | T | | ORF1ab | stopgain_K4783X |
| 14614 | C | T | | ORF1ab | nonsynonymous_R4784C |
| 14615 | G | C | | ORF1ab | nonsynonymous_R4784P |
| 14625 | C | T | | ORF1ab | synonymous_C4787C |
| 14636 | C | T | | ORF1ab | nonsynonymous_A4791V |
| 14644 | A | C | | ORF1ab | nonsynonymous_T4794P |
| 14649 | C | T | | ORF1ab | synonymous_N4795N |
| 14657 | C | T | | ORF1ab | nonsynonymous_A4798V |

| | | | | | |
|---|---|---|---|---|---|
| 14659 | T | A | | ORF1ab | nonsynonymous_F4799I |
| 14660 | T | C | | ORF1ab | nonsynonymous_F4799S |
| 14670 | C | T | | ORF1ab | synonymous_V4802V |
| 14673 | A | G | | ORF1ab | synonymous_K4803K |
| 14675 | C | T | | ORF1ab | nonsynonymous_P4804L |
| 14676 | C | T | | ORF1ab | synonymous_P4804P |
| 14679 | T | C | | ORF1ab | synonymous_G4805G |
| 14688 | C | T | | ORF1ab | synonymous_N4808N |
| 14700 | T | C | | ORF1ab | synonymous_Y4812Y |
| 14708 | C | T | | ORF1ab | nonsynonymous_A4815V |
| 14712 | G | A | T | ORF1ab | synonymous_V4816V; synonymous_V4816V |
| 14714 | C | T | | ORF1ab | nonsynonymous_S4817F |
| 14718 | G | T | | ORF1ab | nonsynonymous_K4818N |
| 14719 | G | T | | ORF1ab | nonsynonymous_G4819C |
| 14720 | G | T | | ORF1ab | nonsynonymous_G4819V |
| 14724 | C | T | | ORF1ab | synonymous_F4820F |
| 14741 | C | T | | ORF1ab | nonsynonymous_S4826F |
| 14746 | G | A | C | ORF1ab | nonsynonymous_E4828K; nonsynonymous_E4828Q |
| 14747 | A | G | | ORF1ab | nonsynonymous_E4828G |
| 14768 | C | T | | ORF1ab | nonsynonymous_A4835V |
| 14773 | G | T | | ORF1ab | nonsynonymous_D4837Y |
| 14786 | C | T | | ORF1ab | nonsynonymous_A4841V |
| 14790 | C | T | | ORF1ab | synonymous_I4842I |
| 14792 | G | T | | ORF1ab | nonsynonymous_S4843I |
| 14793 | C | T | | ORF1ab | synonymous_S4843S |
| 14802 | C | T | | ORF1ab | synonymous_D4846D |
| 14805 | C | T | | ORF1ab | synonymous_Y4847Y |
| 14818 | C | T | | ORF1ab | synonymous_L4852L |
| 14823 | A | T | | ORF1ab | synonymous_P4853P |

| | | | | | |
|---|---|---|---|---|---|
| 14824 | A | T | | ORF1ab | nonsynonymous_T4854S |
| 14825 | C | T | | ORF1ab | nonsynonymous_T4854I |
| 14829 | G | T | | ORF1ab | nonsynonymous_M4855I |
| 14853 | T | C | | ORF1ab | synonymous_F4863F |
| 14865 | T | A | | ORF1ab | synonymous_V4867V |
| 14870 | A | G | | ORF1ab | nonsynonymous_D4869G |
| 14877 | C | T | | ORF1ab | synonymous_Y4871Y |
| 14889 | C | T | | ORF1ab | synonymous_Y4875Y |
| 14898 | C | T | | ORF1ab | synonymous_G4878G |
| 14908 | G | T | | ORF1ab | nonsynonymous_A4882S |
| 14913 | C | T | | ORF1ab | synonymous_N4883N |
| 14932 | C | T | | ORF1ab | synonymous_L4890L |
| 14940 | A | G | | ORF1ab | synonymous_K4892K |
| 14945 | C | T | | ORF1ab | nonsynonymous_A4894V |
| 14953 | C | T | | ORF1ab | nonsynonymous_P4897S |
| 14961 | T | A | | ORF1ab | nonsynonymous_N4899K |
| 14962 | A | T | | ORF1ab | stopgain_K4900X |
| 14968 | G | C | | ORF1ab | nonsynonymous_G4902R |
| 14969 | G | T | | ORF1ab | nonsynonymous_G4902V |
| 14974 | G | T | | ORF1ab | nonsynonymous_A4904S |
| 14975 | C | T | | ORF1ab | nonsynonymous_A4904V |
| 14985 | T | C | | ORF1ab | synonymous_Y4907Y |
| 15004 | G | T | | ORF1ab | stopgain_E4914X |
| 15010 | C | T | | ORF1ab | stopgain_Q4916X |
| 15017 | C | T | | ORF1ab | nonsynonymous_A4918V |
| 15021 | T | C | | ORF1ab | synonymous_L4919L |
| 15024 | C | T | | ORF1ab | synonymous_F4920F |
| 15025 | G | T | | ORF1ab | nonsynonymous_A4921S |
| 15026 | C | T | | ORF1ab | nonsynonymous_A4921V |
| 15048 | C | T | | ORF1ab | synonymous_I4928I |
| 15050 | C | T | | ORF1ab | nonsynonymous_P4929L |
| 15053 | C | T | | ORF1ab | nonsynonymous_T4930I |

| | | | | | |
|---|---|---|---|---|---|
| 15059 | C | T | | ORF1ab | nonsynonymous_T4932I |
| 15075 | G | T | | ORF1ab | nonsynonymous_K4937N |
| 15080 | C | T | | ORF1ab | nonsynonymous_A4939V |
| 15093 | G | A | | ORF1ab | synonymous_K4943K |
| 15095 | A | C | | ORF1ab | nonsynonymous_N4944T |
| 15101 | C | T | | ORF1ab | nonsynonymous_A4946V |
| 15103 | C | T | | ORF1ab | nonsynonymous_R4947C |
| 15112 | G | T | | ORF1ab | nonsynonymous_A4950S |
| 15115 | G | T | | ORF1ab | nonsynonymous_G4951C |
| 15116 | G | T | | ORF1ab | nonsynonymous_G4951V |
| 15121 | T | C | | ORF1ab | nonsynonymous_S4953P |
| 15128 | G | T | | ORF1ab | nonsynonymous_C4955F |
| 15138 | G | T | | ORF1ab | nonsynonymous_M4958I |
| 15140 | C | T | | ORF1ab | nonsynonymous_T4959I |
| 15141 | C | A | T | ORF1ab | synonymous_T4959T;<br>synonymous_T4959T |
| 15144 | T | C | | ORF1ab | synonymous_N4960N |
| 15145 | A | G | | ORF1ab | nonsynonymous_R4961G |
| 15150 | G | T | | ORF1ab | nonsynonymous_Q4962H |
| 15154 | C | T | | ORF1ab | nonsynonymous_H4964Y |
| 15157 | C | A | | ORF1ab | nonsynonymous_Q4965K |
| 15166 | T | C | | ORF1ab | synonymous_L4968L |
| 15173 | C | T | | ORF1ab | nonsynonymous_S4970L |
| 15175 | A | G | | ORF1ab | nonsynonymous_I4971V |
| 15185 | C | T | | ORF1ab | nonsynonymous_T4974I |
| 15193 | G | T | | ORF1ab | nonsynonymous_A4977S |
| 15199 | G | T | | ORF1ab | nonsynonymous_V4979L |
| 15202 | G | T | | ORF1ab | nonsynonymous_V4980L |
| 15205 | A | G | | ORF1ab | nonsynonymous_I4981V |
| 15209 | G | T | | ORF1ab | nonsynonymous_G4982V |
| 15216 | C | T | | ORF1ab | synonymous_S4984S |

| | | | | | |
|---|---|---|---|---|---|
| 15222 | C | T | | ORF1ab | synonymous_F4986F |
| 15233 | G | T | | ORF1ab | nonsynonymous_W4990L |
| 15235 | C | T | | ORF1ab | nonsynonymous_H4991Y |
| 15237 | C | T | | ORF1ab | synonymous_H4991H |
| 15254 | T | C | | ORF1ab | nonsynonymous_V4997A |
| 15265 | G | T | | ORF1ab | nonsynonymous_V5001L |
| 15268 | G | A | | ORF1ab | nonsynonymous_E5002K |
| 15277 | C | T | | ORF1ab | nonsynonymous_H5005Y |
| 15279 | C | T | | ORF1ab | synonymous_H5005H |
| 15285 | G | T | | ORF1ab | nonsynonymous_M5007I |
| 15315 | C | T | | ORF1ab | synonymous_A5017A |
| 15316 | A | G | | ORF1ab | nonsynonymous_M5018V |
| 15319 | C | T | | ORF1ab | nonsynonymous_P5019S |
| 15324 | C | T | | ORF1ab | synonymous_N5020N |
| 15328 | C | T | | ORF1ab | nonsynonymous_L5022F |
| 15332 | G | A | | ORF1ab | nonsynonymous_R5023K |
| 15344 | C | T | | ORF1ab | nonsynonymous_S5027L |
| 15346 | C | T | | ORF1ab | nonsynonymous_L5028F |
| 15352 | C | T | | ORF1ab | nonsynonymous_L5030F |
| 15356 | C | T | | ORF1ab | nonsynonymous_A5031V |
| 15357 | T | C | | ORF1ab | synonymous_A5031A |
| 15359 | G | A | T | ORF1ab | nonsynonymous_R5032H; nonsynonymous_R5032L |
| 15364 | C | T | | ORF1ab | nonsynonymous_H5034Y |
| 15368 | C | G | | ORF1ab | nonsynonymous_T5035R |
| 15371 | C | T | | ORF1ab | nonsynonymous_T5036M |
| 15372 | G | T | | ORF1ab | synonymous_T5036T |
| 15374 | G | A | T | ORF1ab | nonsynonymous_C5037Y; nonsynonymous_C5037F |
| 15380 | G | T | | ORF1ab | nonsynonymous_S5039I |
| 15381 | C | T | | ORF1ab | synonymous_S5039S |

| | | | | | |
|---|---|---|---|---|---|
| 15391 | C | T | | ORF1ab | nonsynonymous_R5043C |
| 15407 | C | T | | ORF1ab | nonsynonymous_A5048V |
| 15418 | G | T | | ORF1ab | nonsynonymous_A5052S |
| 15419 | C | T | | ORF1ab | nonsynonymous_A5052V |
| 15433 | G | A | | ORF1ab | nonsynonymous_E5057K |
| 15438 | G | T | | ORF1ab | nonsynonymous_M5058I |
| 15439 | G | A | | ORF1ab | nonsynonymous_V5059I |
| 15441 | C | T | | ORF1ab | synonymous_V5059V |
| 15444 | G | A | T | ORF1ab | nonsynonymous_M5060I; nonsynonymous_M5060I |
| 15446 | G | T | | ORF1ab | nonsynonymous_C5061F |
| 15470 | C | G | | ORF1ab | nonsynonymous_P5069R |
| 15480 | C | T | | ORF1ab | synonymous_T5072T |
| 15509 | C | T | | ORF1ab | nonsynonymous_A5082V |
| 15511 | A | G | | ORF1ab | nonsynonymous_N5083D |
| 15515 | G | T | | ORF1ab | nonsynonymous_S5084I |
| 15517 | G | T | | ORF1ab | nonsynonymous_V5085F |
| 15533 | A | C | | ORF1ab | nonsynonymous_Q5090P |
| 15540 | C | T | | ORF1ab | synonymous_V5092V |
| 15543 | G | T | | ORF1ab | synonymous_T5093T |
| 15554 | A | G | | ORF1ab | nonsynonymous_N5097S |
| 15564 | A | G | | ORF1ab | synonymous_L5100L |
| 15571 | G | C | | ORF1ab | nonsynonymous_D5103H |
| 15575 | G | T | | ORF1ab | nonsynonymous_G5104V |
| 15579 | C | T | | ORF1ab | synonymous_N5105N |
| 15588 | C | T | | ORF1ab | synonymous_A5108A |
| 15591 | T | C | | ORF1ab | synonymous_D5109D |
| 15597 | T | C | | ORF1ab | synonymous_Y5111Y |
| 15613 | C | T | | ORF1ab | nonsynonymous_H5117Y |
| 15639 | A | G | | ORF1ab | synonymous_R5125R |
| 15649 | G | T | | ORF1ab | nonsynonymous_V5129F |

| | | | | | |
|---|---|---|---|---|---|
| 15653 | A | T | | ORF1ab | nonsynonymous_D5130V |
| 15654 | C | T | | ORF1ab | synonymous_D5130D |
| 15656 | C | T | | ORF1ab | nonsynonymous_T5131I |
| 15682 | T | A | C | ORF1ab | nonsynonymous_Y5140N; nonsynonymous_Y5140H |
| 15685 | T | C | | ORF1ab | synonymous_L5141L |
| 15695 | A | T | | ORF1ab | nonsynonymous_H5144L |
| 15701 | C | T | | ORF1ab | nonsynonymous_S5146L |
| 15708 | G | C | T | ORF1ab | nonsynonymous_M5148I; nonsynonymous_M5148I |
| 15711 | A | T | | ORF1ab | synonymous_I5149I |
| 15714 | C | T | | ORF1ab | synonymous_L5150L |
| 15717 | T | C | | ORF1ab | synonymous_S5151S |
| 15720 | C | T | | ORF1ab | synonymous_D5152D |
| 15721 | G | T | | ORF1ab | nonsynonymous_D5153Y |
| 15732 | G | T | | ORF1ab | synonymous_V5156V |
| 15734 | G | C | | ORF1ab | nonsynonymous_C5157S |
| 15738 | C | T | | ORF1ab | synonymous_F5158F |
| 15743 | G | T | | ORF1ab | nonsynonymous_S5160I |
| 15744 | C | T | | ORF1ab | synonymous_S5160S |
| 15746 | C | T | | ORF1ab | nonsynonymous_T5161I |
| 15756 | T | C | | ORF1ab | synonymous_S5164S |
| 15760 | G | A | | ORF1ab | nonsynonymous_G5166S |
| 15761 | G | T | | ORF1ab | nonsynonymous_G5166V |
| 15763 | C | T | | ORF1ab | synonymous_L5167L |
| 15769 | G | T | | ORF1ab | nonsynonymous_A5169S |
| 15770 | C | T | | ORF1ab | nonsynonymous_A5169V |
| 15771 | T | C | | ORF1ab | synonymous_A5169A |
| 15777 | A | G | | ORF1ab | nonsynonymous_I5171M |
| 15786 | T | C | | ORF1ab | synonymous_F5174F |
| 15791 | C | T | | ORF1ab | nonsynonymous_S5176L |

| | | | | | |
|---|---|---|---|---|---|
| 15792 | A | G | | ORF1ab | synonymous_S5176S |
| 15810 | C | T | | ORF1ab | synonymous_N5182N |
| 15824 | C | T | | ORF1ab | nonsynonymous_S5187F |
| 15826 | G | T | | ORF1ab | stopgain_E5188X |
| 15831 | A | G | | ORF1ab | synonymous_A5189A |
| 15840 | G | T | | ORF1ab | nonsynonymous_W5192C |
| 15842 | C | T | | ORF1ab | nonsynonymous_T5193I |
| 15844 | G | T | | ORF1ab | stopgain_E5194X |
| 15849 | T | A | | ORF1ab | synonymous_T5195T |
| 15852 | C | T | | ORF1ab | synonymous_D5196D |
| 15857 | C | T | | ORF1ab | nonsynonymous_T5198I |
| 15866 | C | T | | ORF1ab | nonsynonymous_P5201L |
| 15868 | C | T | | ORF1ab | nonsynonymous_H5202Y |
| 15880 | T | C | | ORF1ab | nonsynonymous_S5206P |
| 15890 | C | T | | ORF1ab | nonsynonymous_T5209I |
| 15894 | G | C | | ORF1ab | nonsynonymous_M5210I |
| 15898 | G | T | | ORF1ab | nonsynonymous_V5212F |
| 15900 | T | C | | ORF1ab | synonymous_V5212V |
| 15903 | A | G | | ORF1ab | synonymous_K5213K |
| 15910 | G | T | | ORF1ab | nonsynonymous_D5216Y |
| 15915 | T | C | | ORF1ab | synonymous_D5217D |
| 15933 | C | T | | ORF1ab | synonymous_Y5223Y |
| 15934 | C | T | | ORF1ab | nonsynonymous_P5224S |
| 15936 | A | T | | ORF1ab | synonymous_P5224P |
| 15941 | C | T | | ORF1ab | nonsynonymous_P5226L |
| 15944 | C | T | | ORF1ab | nonsynonymous_S5227L |
| 15957 | G | T | | ORF1ab | synonymous_G5231G |
| 15958 | G | T | | ORF1ab | nonsynonymous_A5232S |
| 15959 | C | T | | ORF1ab | nonsynonymous_A5232V |
| 15960 | C | T | | ORF1ab | synonymous_A5232A |
| 15963 | C | T | | ORF1ab | synonymous_G5233G |
| 15969 | T | G | | ORF1ab | nonsynonymous_F5235L |

| | | | | | |
|---|---|---|---|---|---|
| 15981 | C | T | | ORF1ab | synonymous_I5239I |
| 15982 | G | T | | ORF1ab | nonsynonymous_V5240L |
| 16000 | C | T | | ORF1ab | nonsynonymous_L5246F |
| 16006 | A | G | | ORF1ab | nonsynonymous_I5248V |
| 16010 | A | T | | ORF1ab | nonsynonymous_E5249V |
| 16011 | A | G | | ORF1ab | synonymous_E5249E |
| 16017 | C | T | | ORF1ab | synonymous_F5251F |
| 16022 | C | T | | ORF1ab | nonsynonymous_S5253F |
| 16026 | A | G | | ORF1ab | synonymous_L5254L |
| 16028 | C | T | | ORF1ab | nonsynonymous_A5255V |
| 16037 | C | T | | ORF1ab | nonsynonymous_A5258V |
| 16041 | C | T | | ORF1ab | synonymous_Y5259Y |
| 16056 | T | C | | ORF1ab | synonymous_H5264H |
| 16057 | C | T | | ORF1ab | nonsynonymous_P5265S |
| 16059 | T | A | | ORF1ab | synonymous_P5265P |
| 16063 | C | T | | ORF1ab | stopgain_Q5267X |
| 16068 | G | T | | ORF1ab | nonsynonymous_E5268D |
| 16073 | C | T | | ORF1ab | nonsynonymous_A5270V |
| 16076 | A | G | | ORF1ab | nonsynonymous_D5271G |
| 16078 | G | A | | ORF1ab | nonsynonymous_V5272I |
| 16080 | C | T | | ORF1ab | synonymous_V5272V |
| 16096 | C | T | | ORF1ab | stopgain_Q5278X |
| 16098 | A | T | | ORF1ab | nonsynonymous_Q5278H |
| 16103 | T | A | | ORF1ab | nonsynonymous_I5280K |
| 16110 | G | T | | ORF1ab | nonsynonymous_K5282N |
| 16111 | C | T | | ORF1ab | synonymous_L5283L |
| 16122 | G | T | | ORF1ab | nonsynonymous_E5286D |
| 16143 | C | T | | ORF1ab | synonymous_D5293D |
| 16152 | T | C | | ORF1ab | synonymous_S5296S |
| 16154 | T | G | | ORF1ab | nonsynonymous_V5297G |
| 16163 | C | T | | ORF1ab | nonsynonymous_T5300I |
| 16171 | A | C | | ORF1ab | nonsynonymous_N5303H |

| | | | | | |
|---|---|---|---|---|---|
| 16188 | G | T | | ORF1ab | nonsynonymous_W5308C |
| 16189 | G | A | | ORF1ab | nonsynonymous_E5309K |
| 16192 | C | T | | ORF1ab | nonsynonymous_P5310S |
| 16196 | A | T | | ORF1ab | nonsynonymous_E5311V |
| 16206 | G | T | | ORF1ab | nonsynonymous_E5314D |
| 16207 | G | C | | ORF1ab | nonsynonymous_A5315P |
| 16208 | C | T | | ORF1ab | nonsynonymous_A5315V |
| 16212 | G | T | | ORF1ab | nonsynonymous_M5316I |
| 16217 | C | T | | ORF1ab | nonsynonymous_T5318I |
| 16221 | G | A | T | ORF1ab | synonymous_P5319P; synonymous_P5319P |
| 16222 | C | T | | ORF1ab | nonsynonymous_H5320Y |
| 16229 | T | C | | ORF1ab | nonsynonymous_V5322A |
| 16232 | T | C | | ORF1ab | nonsynonymous_L5323S |
| 16234 | C | T | | ORF1ab | stopgain_Q5324X |
| 16236 | G | A | | ORF1ab | synonymous_Q5324Q |
| 16242 | T | A | | ORF1ab | synonymous_V5326V |
| 16247 | C | T | | ORF1ab | nonsynonymous_A5328V |
| 16255 | C | T | | ORF1ab | nonsynonymous_L5331F |
| 16259 | G | T | | ORF1ab | nonsynonymous_C5332F |
| 16260 | C | T | | ORF1ab | synonymous_C5332C |
| 16262 | A | T | | ORF1ab | nonsynonymous_N5333I |
| 16263 | T | C | | ORF1ab | synonymous_N5333N |
| 16265 | C | T | | ORF1ab | nonsynonymous_S5334L |
| 16273 | T | C | | ORF1ab | nonsynonymous_S5337P |
| 16276 | T | G | | ORF1ab | nonsynonymous_L5338V |
| 16285 | G | T | | ORF1ab | nonsynonymous_G5341C |
| 16289 | C | T | | ORF1ab | nonsynonymous_A5342V |
| 16293 | C | T | | ORF1ab | synonymous_C5343C |
| 16295 | T | C | | ORF1ab | nonsynonymous_I5344T |
| 16302 | A | G | | ORF1ab | synonymous_R5346R |

| | | | | | |
|---|---|---|---|---|---|
| 16303 | C | T | | ORF1ab | nonsynonymous_P5347S |
| 16305 | A | T | | ORF1ab | synonymous_P5347P |
| 16308 | C | T | | ORF1ab | synonymous_F5348F |
| 16319 | A | G | | ORF1ab | nonsynonymous_K5352R |
| 16323 | C | T | | ORF1ab | synonymous_C5353C |
| 16325 | G | C | T | ORF1ab | nonsynonymous_C5354S; nonsynonymous_C5354F |
| 16329 | C | T | | ORF1ab | synonymous_Y5355Y |
| 16330 | G | T | | ORF1ab | nonsynonymous_D5356Y |
| 16332 | C | T | | ORF1ab | synonymous_D5356D |
| 16338 | C | T | | ORF1ab | synonymous_V5358V |
| 16341 | A | G | | ORF1ab | nonsynonymous_I5359M |
| 16343 | C | T | | ORF1ab | nonsynonymous_S5360L |
| 16362 | C | T | | ORF1ab | synonymous_V5366V |
| 16375 | C | T | | ORF1ab | nonsynonymous_P5371S |
| 16381 | G | T | | ORF1ab | nonsynonymous_V5373F |
| 16386 | C | T | | ORF1ab | synonymous_C5374C |
| 16393 | C | T | | ORF1ab | nonsynonymous_P5377S |
| 16407 | C | T | | ORF1ab | synonymous_V5381V |
| 16416 | G | T | | ORF1ab | synonymous_V5384V |
| 16418 | C | T | | ORF1ab | nonsynonymous_T5385I |
| 16423 | C | T | | ORF1ab | nonsynonymous_L5387F |
| 16428 | C | T | | ORF1ab | synonymous_Y5388Y |
| 16432 | G | T | | ORF1ab | stopgain_G5390X |
| 16436 | G | T | | ORF1ab | nonsynonymous_G5391V |
| 16442 | G | C | | ORF1ab | nonsynonymous_S5393T |
| 16457 | C | T | | ORF1ab | nonsynonymous_S5398L |
| 16460 | A | T | | ORF1ab | nonsynonymous_H5399L |
| 16466 | C | T | | ORF1ab | nonsynonymous_P5401L |
| 16467 | A | G | | ORF1ab | synonymous_P5401P |

| | | | | | |
|---|---|---|---|---|---|
| 16468 | C | A | T | ORF1ab | nonsynonymous_P5402T;<br>nonsynonymous_P5402S |
| 16470 | C | T | | ORF1ab | synonymous_P5402P |
| 16474 | A | G | | ORF1ab | nonsynonymous_S5404G |
| 16501 | G | T | | ORF1ab | nonsynonymous_V5413F |
| 16508 | G | T | | ORF1ab | nonsynonymous_G5415V |
| 16523 | C | T | | ORF1ab | nonsynonymous_T5420I |
| 16535 | G | T | | ORF1ab | nonsynonymous_S5424I |
| 16551 | C | T | | ORF1ab | synonymous_D5429D |
| 16566 | A | T | | ORF1ab | synonymous_A5434A |
| 16568 | C | T | | ORF1ab | nonsynonymous_T5435I |
| 16571 | G | T | | ORF1ab | nonsynonymous_C5436F |
| 16574 | A | G | | ORF1ab | nonsynonymous_D5437G |
| 16575 | C | T | | ORF1ab | synonymous_D5437D |
| 16579 | A | T | | ORF1ab | nonsynonymous_T5439S |
| 16580 | C | T | | ORF1ab | nonsynonymous_T5439I |
| 16588 | G | T | | ORF1ab | nonsynonymous_G5442C |
| 16590 | T | C | | ORF1ab | synonymous_G5442G |
| 16592 | A | G | | ORF1ab | nonsynonymous_D5443G |
| 16596 | C | T | | ORF1ab | synonymous_Y5444Y |
| 16601 | T | C | | ORF1ab | nonsynonymous_L5446S |
| 16603 | G | T | | ORF1ab | nonsynonymous_A5447S |
| 16604 | C | T | | ORF1ab | nonsynonymous_A5447V |
| 16605 | T | A | C | ORF1ab | synonymous_A5447A;<br>synonymous_A5447A |
| 16606 | A | C | | ORF1ab | nonsynonymous_N5448H |
| 16610 | C | T | | ORF1ab | nonsynonymous_T5449I |
| 16613 | G | A | T | ORF1ab | nonsynonymous_C5450Y;<br>nonsynonymous_C5450F |
| 16619 | A | G | | ORF1ab | nonsynonymous_E5452G |

| | | | | | |
|---|---|---|---|---|---|
| 16621 | A | C | G | ORF1ab | synonymous_R5453R; nonsynonymous_R5453G |
| 16622 | G | A | | ORF1ab | nonsynonymous_R5453K |
| 16624 | C | T | | ORF1ab | nonsynonymous_L5454F |
| 16625 | T | C | | ORF1ab | nonsynonymous_L5454P |
| 16626 | C | T | | ORF1ab | synonymous_L5454L |
| 16636 | G | T | | ORF1ab | nonsynonymous_A5458S |
| 16637 | C | T | | ORF1ab | nonsynonymous_A5458V |
| 16640 | C | T | | ORF1ab | nonsynonymous_A5459V |
| 16641 | A | T | | ORF1ab | synonymous_A5459A |
| 16642 | G | A | | ORF1ab | nonsynonymous_E5460K |
| 16646 | C | T | | ORF1ab | nonsynonymous_T5461M |
| 16648 | C | T | | ORF1ab | nonsynonymous_L5462F |
| 16658 | C | A | T | ORF1ab | nonsynonymous_T5465N; nonsynonymous_T5465I |
| 16662 | G | A | C | ORF1ab | synonymous_E5466E; nonsynonymous_E5466D |
| 16663 | G | C | T | ORF1ab | nonsynonymous_E5467Q; stopgain_E5467X |
| 16665 | G | T | | ORF1ab | nonsynonymous_E5467D |
| 16669 | T | G | | ORF1ab | nonsynonymous_F5469V |
| 16670 | T | C | | ORF1ab | nonsynonymous_F5469S |
| 16671 | T | C | | ORF1ab | synonymous_F5469F |
| 16685 | G | C | | ORF1ab | nonsynonymous_G5474A |
| 16687 | A | G | | ORF1ab | nonsynonymous_I5475V |
| 16688 | T | C | | ORF1ab | nonsynonymous_I5475T |
| 16690 | G | T | | ORF1ab | nonsynonymous_A5476S |
| 16691 | C | T | | ORF1ab | nonsynonymous_A5476V |
| 16694 | C | T | | ORF1ab | nonsynonymous_T5477I |
| 16695 | T | C | | ORF1ab | synonymous_T5477T |
| 16699 | C | T | | ORF1ab | nonsynonymous_R5479C |

| | | | | | |
|---|---|---|---|---|---|
| 16700 | G | T | | ORF1ab | nonsynonymous_R5479L |
| 16707 | G | T | | ORF1ab | synonymous_V5481V |
| 16708 | C | T | | ORF1ab | synonymous_L5482L |
| 16712 | C | T | | ORF1ab | nonsynonymous_S5483F |
| 16729 | C | A | T | ORF1ab | nonsynonymous_L5489I;<br>nonsynonymous_L5489F |
| 16733 | C | T | | ORF1ab | nonsynonymous_S5490L |
| 16736 | G | T | | ORF1ab | nonsynonymous_W5491L |
| 16738 | G | T | | ORF1ab | stopgain_E5492X |
| 16741 | G | T | | ORF1ab | nonsynonymous_V5493F |
| 16745 | G | T | | ORF1ab | nonsynonymous_G5494V |
| 16749 | A | C | | ORF1ab | nonsynonymous_K5495N |
| 16750 | C | T | | ORF1ab | nonsynonymous_P5496S |
| 16751 | C | T | | ORF1ab | nonsynonymous_P5496L |
| 16760 | C | T | | ORF1ab | nonsynonymous_P5499L |
| 16767 | C | T | | ORF1ab | synonymous_N5501N |
| 16768 | C | T | | ORF1ab | stopgain_R5502X |
| 16779 | C | T | | ORF1ab | synonymous_V5505V |
| 16787 | G | T | | ORF1ab | nonsynonymous_G5508V |
| 16788 | T | A | | ORF1ab | synonymous_G5508G |
| 16792 | C | T | | ORF1ab | nonsynonymous_R5510C |
| 16793 | G | A | | ORF1ab | nonsynonymous_R5510H |
| 16795 | G | T | | ORF1ab | nonsynonymous_V5511L |
| 16830 | C | T | | ORF1ab | synonymous_Y5522Y |
| 16832 | C | T | | ORF1ab | nonsynonymous_T5523I |
| 16836 | T | C | | ORF1ab | synonymous_F5524F |
| 16837 | G | T | | ORF1ab | stopgain_E5525X |
| 16842 | A | T | | ORF1ab | nonsynonymous_K5526N |
| 16848 | C | T | | ORF1ab | synonymous_D5528D |
| 16852 | G | T | | ORF1ab | nonsynonymous_G5530C |
| 16853 | G | T | | ORF1ab | nonsynonymous_G5530V |

| | | | | | |
|---|---|---|---|---|---|
| 16855 | G | T | | ORF1ab | nonsynonymous_D5531Y |
| 16858 | G | A | T | ORF1ab | nonsynonymous_A5532T;<br>nonsynonymous_A5532S |
| 16859 | C | G | | ORF1ab | nonsynonymous_A5532G |
| 16877 | C | T | | ORF1ab | nonsynonymous_T5538I |
| 16881 | A | T | | ORF1ab | synonymous_T5539T |
| 16887 | C | T | | ORF1ab | synonymous_Y5541Y |
| 16903 | G | T | | ORF1ab | nonsynonymous_D5547Y |
| 16919 | C | T | | ORF1ab | nonsynonymous_T5552I |
| 16924 | C | T | | ORF1ab | nonsynonymous_H5554Y |
| 16942 | A | G | | ORF1ab | nonsynonymous_S5560G |
| 16943 | G | T | | ORF1ab | nonsynonymous_S5560I |
| 16949 | C | T | | ORF1ab | nonsynonymous_P5562L |
| 16963 | C | T | | ORF1ab | stopgain_Q5567X |
| 16966 | G | T | | ORF1ab | stopgain_E5568X |
| 16968 | G | T | | ORF1ab | nonsynonymous_E5568D |
| 16971 | C | T | | ORF1ab | synonymous_H5569H |
| 16975 | G | C | T | ORF1ab | nonsynonymous_V5571L;<br>nonsynonymous_V5571F |
| 16988 | G | T | | ORF1ab | nonsynonymous_G5575V |
| 16989 | C | T | | ORF1ab | synonymous_G5575G |
| 16992 | A | G | | ORF1ab | synonymous_L5576L |
| 16993 | T | C | | ORF1ab | nonsynonymous_Y5577H |
| 16995 | C | T | | ORF1ab | synonymous_Y5577Y |
| 17004 | C | T | | ORF1ab | synonymous_L5580L |
| 17007 | T | C | | ORF1ab | synonymous_N5581N |
| 17010 | C | A | T | ORF1ab | synonymous_I5582I;<br>synonymous_I5582I |
| 17012 | C | T | | ORF1ab | nonsynonymous_S5583L |
| 17027 | G | T | | ORF1ab | nonsynonymous_S5588I |
| 17028 | C | T | | ORF1ab | synonymous_S5588S |

| | | | | | |
|---|---|---|---|---|---|
| 17032 | G | T | | ORF1ab | nonsynonymous_V5590F |
| 17035 | G | T | | ORF1ab | nonsynonymous_A5591S |
| 17049 | G | C | | ORF1ab | nonsynonymous_K5595N |
| 17050 | G | T | | ORF1ab | nonsynonymous_V5596F |
| 17058 | G | A | | ORF1ab | nonsynonymous_M5598I |
| 17066 | A | T | | ORF1ab | nonsynonymous_Y5601F |
| 17072 | C | T | | ORF1ab | nonsynonymous_T5603I |
| 17084 | C | T | | ORF1ab | nonsynonymous_P5607L |
| 17096 | G | T | | ORF1ab | nonsynonymous_G5611V |
| 17104 | C | T | | ORF1ab | nonsynonymous_H5614Y |
| 17122 | G | T | | ORF1ab | nonsynonymous_A5620S |
| 17125 | C | T | | ORF1ab | nonsynonymous_L5621F |
| 17126 | T | C | | ORF1ab | nonsynonymous_L5621P |
| 17135 | C | T | | ORF1ab | nonsynonymous_P5624L |
| 17159 | C | T | | ORF1ab | nonsynonymous_A5632V |
| 17167 | C | T | | ORF1ab | nonsynonymous_H5635Y |
| 17171 | C | T | | ORF1ab | nonsynonymous_A5636V |
| 17172 | C | A | T | ORF1ab | synonymous_A5636A;<br>synonymous_A5636A |
| 17180 | A | C | T | ORF1ab | nonsynonymous_D5639A;<br>nonsynonymous_D5639V |
| 17190 | T | C | | ORF1ab | synonymous_C5642C |
| 17193 | G | T | | ORF1ab | nonsynonymous_E5643D |
| 17196 | G | A | | ORF1ab | synonymous_K5644K |
| 17212 | C | T | | ORF1ab | nonsynonymous_P5650S |
| 17220 | T | A | | ORF1ab | nonsynonymous_D5652E |
| 17239 | C | T | | ORF1ab | nonsynonymous_P5659S |
| 17242 | G | T | | ORF1ab | nonsynonymous_A5660S |
| 17245 | C | T | | ORF1ab | nonsynonymous_R5661C |
| 17247 | T | C | | ORF1ab | synonymous_R5661R |
| 17249 | C | T | | ORF1ab | nonsynonymous_A5662V |

| | | | | | |
|---|---|---|---|---|---|
| 17251 | C | T | | ORF1ab | nonsynonymous_R5663C |
| 17254 | G | T | | ORF1ab | nonsynonymous_V5664L |
| 17259 | G | A | T | ORF1ab | synonymous_E5665E;<br>nonsynonymous_E5665D |
| 17261 | G | T | | ORF1ab | nonsynonymous_C5666F |
| 17277 | A | T | | ORF1ab | nonsynonymous_K5671N |
| 17278 | G | T | | ORF1ab | nonsynonymous_V5672L |
| 17285 | C | T | | ORF1ab | nonsynonymous_S5674L |
| 17288 | C | T | | ORF1ab | nonsynonymous_T5675I |
| 17303 | T | C | | ORF1ab | nonsynonymous_V5680A |
| 17304 | C | T | | ORF1ab | synonymous_V5680V |
| 17326 | C | A | T | ORF1ab | nonsynonymous_P5688T;<br>nonsynonymous_P5688S |
| 17327 | C | T | | ORF1ab | nonsynonymous_P5688L |
| 17333 | C | T | | ORF1ab | nonsynonymous_T5690M |
| 17336 | C | T | | ORF1ab | nonsynonymous_T5691I |
| 17338 | G | T | | ORF1ab | nonsynonymous_A5692S |
| 17339 | C | T | | ORF1ab | nonsynonymous_A5692V |
| 17352 | C | T | | ORF1ab | synonymous_V5696V |
| 17366 | C | T | | ORF1ab | nonsynonymous_S5701L |
| 17367 | A | G | | ORF1ab | synonymous_S5701S |
| 17372 | C | T | | ORF1ab | nonsynonymous_A5703V |
| 17373 | C | T | | ORF1ab | synonymous_A5703A |
| 17375 | C | T | | ORF1ab | nonsynonymous_T5704I |
| 17380 | T | C | | ORF1ab | nonsynonymous_Y5706H |
| 17383 | G | A | | ORF1ab | nonsynonymous_D5707N |
| 17392 | G | T | | ORF1ab | nonsynonymous_V5710F |
| 17397 | C | T | | ORF1ab | synonymous_V5711V |
| 17403 | C | T | | ORF1ab | synonymous_A5713A |
| 17410 | C | T | | ORF1ab | nonsynonymous_R5716C |
| 17415 | T | C | | ORF1ab | synonymous_A5717A |

| | | | | | |
|---|---|---|---|---|---|
| 17421 | C | T | | ORF1ab | synonymous_H5719H |
| 17452 | C | T | | ORF1ab | nonsynonymous_P5730S |
| 17454 | T | C | | ORF1ab | synonymous_P5730P |
| 17460 | A | C | G | ORF1ab | synonymous_P5732P;<br>synonymous_P5732P |
| 17461 | C | T | | ORF1ab | nonsynonymous_R5733C |
| 17470 | C | T | | ORF1ab | synonymous_L5736L |
| 17471 | T | C | | ORF1ab | nonsynonymous_L5736P |
| 17477 | A | G | | ORF1ab | nonsynonymous_K5738R |
| 17488 | G | T | | ORF1ab | stopgain_E5742X |
| 17491 | C | T | | ORF1ab | nonsynonymous_P5743S |
| 17492 | C | T | | ORF1ab | nonsynonymous_P5743L |
| 17502 | C | T | | ORF1ab | synonymous_F5746F |
| 17504 | A | T | | ORF1ab | nonsynonymous_N5747I |
| 17522 | T | C | | ORF1ab | nonsynonymous_M5753T |
| 17523 | G | A | T | ORF1ab | nonsynonymous_M5753I;<br>nonsynonymous_M5753I |
| 17536 | C | T | | ORF1ab | nonsynonymous_P5758S |
| 17548 | C | A | T | ORF1ab | nonsynonymous_L5762I;<br>nonsynonymous_L5762F |
| 17552 | G | A | | ORF1ab | nonsynonymous_G5763E |
| 17555 | C | T | | ORF1ab | nonsynonymous_T5764I |
| 17562 | G | T | | ORF1ab | synonymous_R5766R |
| 17563 | C | T | | ORF1ab | nonsynonymous_R5767C |
| 17564 | G | T | | ORF1ab | nonsynonymous_R5767L |
| 17569 | C | T | | ORF1ab | nonsynonymous_P5769S |
| 17573 | C | T | | ORF1ab | nonsynonymous_A5770V |
| 17574 | T | A | | ORF1ab | synonymous_A5770A |
| 17575 | G | A | | ORF1ab | nonsynonymous_E5771K |
| 17585 | A | T | | ORF1ab | nonsynonymous_D5774V |

| | | | | | |
|---|---|---|---|---|---|
| 17588 | C | T | G | ORF1ab | nonsynonymous_T5775I; nonsynonymous_T5775S |
| 17589 | T | A | | ORF1ab | synonymous_T5775T |
| 17590 | G | T | | ORF1ab | nonsynonymous_V5776L |
| 17592 | G | T | | ORF1ab | synonymous_V5776V |
| 17596 | G | A | | ORF1ab | nonsynonymous_A5778T |
| 17598 | T | C | | ORF1ab | synonymous_A5778A |
| 17615 | A | G | | ORF1ab | nonsynonymous_K5784R |
| 17623 | G | A | T | ORF1ab | nonsynonymous_A5787T; nonsynonymous_A5787S |
| 17624 | C | T | | ORF1ab | nonsynonymous_A5787V |
| 17632 | G | A | | ORF1ab | nonsynonymous_D5790N |
| 17634 | C | T | | ORF1ab | synonymous_D5790D |
| 17639 | C | T | | ORF1ab | nonsynonymous_S5792L |
| 17642 | C | T | | ORF1ab | nonsynonymous_A5793V |
| 17648 | G | A | | ORF1ab | nonsynonymous_C5795Y |
| 17649 | C | T | | ORF1ab | synonymous_C5795C |
| 17656 | A | G | | ORF1ab | nonsynonymous_M5798V |
| 17668 | G | T | | ORF1ab | nonsynonymous_G5802C |
| 17669 | G | T | | ORF1ab | nonsynonymous_G5802V |
| 17671 | G | T | | ORF1ab | nonsynonymous_V5803F |
| 17676 | C | T | | ORF1ab | synonymous_I5804I |
| 17678 | C | T | | ORF1ab | nonsynonymous_T5805M |
| 17688 | T | C | | ORF1ab | synonymous_V5808V |
| 17689 | T | C | | ORF1ab | nonsynonymous_S5809P |
| 17690 | C | T | | ORF1ab | nonsynonymous_S5809L |
| 17692 | T | C | | ORF1ab | nonsynonymous_S5810P |
| 17693 | C | T | | ORF1ab | nonsynonymous_S5810F |
| 17708 | C | T | | ORF1ab | nonsynonymous_P5815L |
| 17716 | G | T | | ORF1ab | nonsynonymous_G5818C |

| | | | | | |
|---|---|---|---|---|---|
| 17729 | A | C | G | ORF1ab | nonsynonymous_E5822A; nonsynonymous_E5822G |
| 17730 | A | T | | ORF1ab | nonsynonymous_E5822D |
| 17733 | C | T | G | ORF1ab | synonymous_F5823F; nonsynonymous_F5823L |
| 17734 | C | T | | ORF1ab | nonsynonymous_L5824F |
| 17738 | C | T | | ORF1ab | nonsynonymous_T5825I |
| 17747 | C | T | | ORF1ab | nonsynonymous_P5828L |
| 17753 | G | A | | ORF1ab | stopgain_W5830X |
| 17761 | G | T | | ORF1ab | nonsynonymous_A5833S |
| 17762 | C | T | | ORF1ab | nonsynonymous_A5833V |
| 17764 | G | T | | ORF1ab | nonsynonymous_V5834F |
| 17765 | T | C | | ORF1ab | nonsynonymous_V5834A |
| 17766 | C | T | | ORF1ab | synonymous_V5834V |
| 17771 | T | C | | ORF1ab | nonsynonymous_I5836T |
| 17773 | T | C | | ORF1ab | nonsynonymous_S5837P |
| 17776 | C | T | | ORF1ab | nonsynonymous_P5838S |
| 17783 | A | T | | ORF1ab | nonsynonymous_N5840I |
| 17790 | G | T | | ORF1ab | nonsynonymous_Q5842H |
| 17795 | C | T | | ORF1ab | nonsynonymous_A5844V |
| 17796 | T | C | | ORF1ab | synonymous_A5844A |
| 17799 | A | T | G | ORF1ab | synonymous_V5845V; synonymous_V5845V |
| 17802 | C | T | | ORF1ab | synonymous_A5846A |
| 17808 | G | A | | ORF1ab | synonymous_K5848K |
| 17816 | G | T | | ORF1ab | nonsynonymous_G5851V |
| 17825 | C | T | | ORF1ab | nonsynonymous_T5854I |
| 17829 | A | T | | ORF1ab | nonsynonymous_Q5855H |
| 17831 | C | T | | ORF1ab | nonsynonymous_T5856I |
| 17836 | G | A | | ORF1ab | nonsynonymous_D5858N |
| 17837 | A | G | | ORF1ab | nonsynonymous_D5858G |

| | | | | | |
|---|---|---|---|---|---|
| 17843 | C | T | | ORF1ab | nonsynonymous_S5860L |
| 17848 | G | C | T | ORF1ab | nonsynonymous_G5862R; nonsynonymous_G5862C |
| 17849 | G | T | | ORF1ab | nonsynonymous_G5862V |
| 17850 | C | T | | ORF1ab | synonymous_G5862G |
| 17855 | A | G | | ORF1ab | nonsynonymous_E5864G |
| 17858 | A | G | | ORF1ab | nonsynonymous_Y5865C |
| 17863 | T | A | | ORF1ab | nonsynonymous_Y5867N |
| 17864 | A | T | | ORF1ab | nonsynonymous_Y5867F |
| 17865 | T | A | C | ORF1ab | stopgain_Y5867X; synonymous_Y5867Y |
| 17874 | C | T | | ORF1ab | synonymous_F5870F |
| 17876 | C | T | | ORF1ab | nonsynonymous_T5871I |
| 17891 | C | T | | ORF1ab | nonsynonymous_T5876I |
| 17894 | C | T | | ORF1ab | nonsynonymous_A5877V |
| 17898 | C | T | | ORF1ab | synonymous_H5878H |
| 17900 | C | T | | ORF1ab | nonsynonymous_S5879F |
| 17906 | A | G | | ORF1ab | nonsynonymous_N5881S |
| 17915 | G | T | | ORF1ab | nonsynonymous_R5884I |
| 17919 | T | A | | ORF1ab | nonsynonymous_F5885L |
| 17921 | A | G | | ORF1ab | nonsynonymous_N5886S |
| 17926 | G | A | | ORF1ab | nonsynonymous_A5888T |
| 17935 | A | G | | ORF1ab | nonsynonymous_R5891G |
| 17940 | A | T | G | ORF1ab | synonymous_A5892A; synonymous_A5892A |
| 17944 | G | T | | ORF1ab | nonsynonymous_V5894L |
| 17949 | C | T | | ORF1ab | synonymous_G5895G |
| 17951 | T | C | | ORF1ab | nonsynonymous_I5896T |
| 17958 | C | T | | ORF1ab | synonymous_C5898C |
| 17973 | A | G | | ORF1ab | synonymous_R5903R |
| 17985 | C | T | | ORF1ab | synonymous_D5907D |

| | | | | | |
|---|---|---|---|---|---|
| 17993 | A | G | | ORF1ab | nonsynonymous_Q5910R |
| 17995 | T | A | | ORF1ab | nonsynonymous_F5911I |
| 17999 | C | T | | ORF1ab | nonsynonymous_T5912I |
| 18004 | C | T | | ORF1ab | nonsynonymous_L5914F |
| 18006 | T | C | | ORF1ab | synonymous_L5914L |
| 18008 | A | G | | ORF1ab | nonsynonymous_E5915G |
| 18009 | A | G | | ORF1ab | synonymous_E5915E |
| 18012 | T | C | | ORF1ab | synonymous_I5916I |
| 18017 | G | T | | ORF1ab | nonsynonymous_R5918L |
| 18020 | G | T | | ORF1ab | nonsynonymous_R5919M |
| 18022 | A | G | | ORF1ab | nonsynonymous_N5920D |
| 18029 | C | T | | ORF1ab | nonsynonymous_A5922V |
| 18060 | C | T | | ORF1ab | synonymous_L5932L |
| 18071 | G | T | | ORF1ab | nonsynonymous_C5936F |
| 18078 | G | C | | ORF1ab | nonsynonymous_K5938N |
| 18082 | A | G | | ORF1ab | nonsynonymous_I5940V |
| 18086 | C | T | | ORF1ab | nonsynonymous_T5941I |
| 18088 | G | T | | ORF1ab | nonsynonymous_G5942W |
| 18090 | G | T | | ORF1ab | synonymous_G5942G |
| 18094 | C | T | | ORF1ab | nonsynonymous_H5944Y |
| 18096 | T | C | | ORF1ab | synonymous_H5944H |
| 18098 | C | T | | ORF1ab | nonsynonymous_P5945L |
| 18101 | C | T | | ORF1ab | nonsynonymous_T5946I |
| 18103 | C | T | | ORF1ab | stopgain_Q5947X |
| 18105 | G | T | | ORF1ab | nonsynonymous_Q5947H |
| 18106 | G | T | | ORF1ab | nonsynonymous_A5948S |
| 18107 | C | T | | ORF1ab | nonsynonymous_A5948V |
| 18109 | C | T | | ORF1ab | nonsynonymous_P5949S |
| 18110 | C | T | | ORF1ab | nonsynonymous_P5949L |
| 18111 | T | C | | ORF1ab | synonymous_P5949P |
| 18120 | C | T | | ORF1ab | synonymous_L5952L |
| 18122 | G | T | | ORF1ab | nonsynonymous_S5953I |

| | | | | | |
|---|---|---|---|---|---|
| 18131 | C | T | | ORF1ab | nonsynonymous_T5956I |
| 18134 | A | G | | ORF1ab | nonsynonymous_K5957R |
| 18138 | C | A | T | ORF1ab | nonsynonymous_F5958L;<br>synonymous_F5958F |
| 18141 | A | G | | ORF1ab | synonymous_K5959K |
| 18149 | G | T | | ORF1ab | nonsynonymous_G5962V |
| 18161 | A | G | | ORF1ab | nonsynonymous_D5966G |
| 18162 | C | T | | ORF1ab | synonymous_D5966D |
| 18163 | A | C | | ORF1ab | nonsynonymous_I5967L |
| 18166 | C | T | | ORF1ab | nonsynonymous_P5968S |
| 18167 | C | T | | ORF1ab | nonsynonymous_P5968L |
| 18170 | G | T | | ORF1ab | nonsynonymous_G5969V |
| 18171 | C | T | | ORF1ab | synonymous_G5969G |
| 18175 | C | T | | ORF1ab | nonsynonymous_P5971S |
| 18176 | C | T | | ORF1ab | nonsynonymous_P5971L |
| 18180 | G | A | | ORF1ab | synonymous_K5972K |
| 18181 | G | T | | ORF1ab | nonsynonymous_D5973Y |
| 18183 | C | T | | ORF1ab | synonymous_D5973D |
| 18186 | G | T | | ORF1ab | nonsynonymous_M5974I |
| 18189 | C | T | | ORF1ab | synonymous_T5975T |
| 18197 | G | A | | ORF1ab | nonsynonymous_R5978K |
| 18213 | G | T | | ORF1ab | nonsynonymous_M5983I |
| 18214 | G | T | | ORF1ab | nonsynonymous_G5984C |
| 18215 | G | T | | ORF1ab | nonsynonymous_G5984V |
| 18220 | A | T | | ORF1ab | stopgain_K5986X |
| 18222 | A | T | | ORF1ab | nonsynonymous_K5986N |
| 18223 | A | T | | ORF1ab | nonsynonymous_M5987L |
| 18225 | G | A | | ORF1ab | nonsynonymous_M5987I |
| 18231 | T | C | | ORF1ab | synonymous_Y5989Y |
| 18232 | C | T | | ORF1ab | stopgain_Q5990X |
| 18237 | T | C | | ORF1ab | synonymous_V5991V |

| | | | | | |
|---|---|---|---|---|---|
| 18239 | A | T | | ORF1ab | nonsynonymous_N5992I |
| 18247 | C | T | | ORF1ab | nonsynonymous_P5995S |
| 18251 | A | G | | ORF1ab | nonsynonymous_N5996S |
| 18252 | C | T | | ORF1ab | synonymous_N5996N |
| 18255 | G | T | | ORF1ab | nonsynonymous_M5997I |
| 18261 | C | T | | ORF1ab | synonymous_I5999I |
| 18263 | C | T | | ORF1ab | nonsynonymous_T6000I |
| 18264 | C | A | T | ORF1ab | synonymous_T6000T;<br>synonymous_T6000T |
| 18287 | T | C | | ORF1ab | nonsynonymous_V6008A |
| 18290 | G | T | | ORF1ab | nonsynonymous_R6009L |
| 18292 | G | A | | ORF1ab | nonsynonymous_A6010T |
| 18297 | G | T | | ORF1ab | nonsynonymous_W6011C |
| 18303 | C | T | | ORF1ab | synonymous_G6013G |
| 18306 | C | T | | ORF1ab | synonymous_F6014F |
| 18312 | C | T | | ORF1ab | synonymous_V6016V |
| 18317 | G | T | | ORF1ab | nonsynonymous_G6018V |
| 18318 | G | T | | ORF1ab | synonymous_G6018G |
| 18326 | C | T | | ORF1ab | nonsynonymous_A6021V |
| 18328 | A | G | | ORF1ab | nonsynonymous_T6022A |
| 18329 | C | T | | ORF1ab | nonsynonymous_T6022I |
| 18331 | A | T | | ORF1ab | stopgain_R6023X |
| 18333 | A | T | | ORF1ab | nonsynonymous_R6023S |
| 18334 | G | A | | ORF1ab | nonsynonymous_E6024K |
| 18341 | T | C | | ORF1ab | nonsynonymous_V6026A |
| 18348 | C | T | | ORF1ab | synonymous_T6028T |
| 18351 | T | C | | ORF1ab | synonymous_N6029N |
| 18355 | C | T | | ORF1ab | nonsynonymous_P6031S |
| 18364 | C | T | | ORF1ab | synonymous_L6034L |
| 18368 | G | T | | ORF1ab | nonsynonymous_G6035V |
| 18374 | C | T | | ORF1ab | nonsynonymous_S6037F |

| | | | | | |
|---|---|---|---|---|---|
| 18380 | G | T | | ORF1ab | nonsynonymous_G6039V |
| 18381 | T | C | | ORF1ab | synonymous_G6039G |
| 18387 | C | T | | ORF1ab | synonymous_N6041N |
| 18391 | G | T | | ORF1ab | nonsynonymous_V6043F |
| 18398 | T | C | | ORF1ab | nonsynonymous_V6045A |
| 18399 | A | T | | ORF1ab | synonymous_V6045V |
| 18400 | C | T | G | ORF1ab | nonsynonymous_P6046S; nonsynonymous_P6046A |
| 18404 | C | T | | ORF1ab | nonsynonymous_T6047I |
| 18407 | G | T | | ORF1ab | nonsynonymous_G6048V |
| 18419 | C | T | | ORF1ab | nonsynonymous_T6052I |
| 18421 | C | T | | ORF1ab | nonsynonymous_P6053S |
| 18428 | A | T | | ORF1ab | nonsynonymous_N6055I |
| 18431 | C | T | | ORF1ab | nonsynonymous_T6056I |
| 18433 | G | T | | ORF1ab | nonsynonymous_D6057Y |
| 18435 | T | C | | ORF1ab | synonymous_D6057D |
| 18441 | C | T | | ORF1ab | synonymous_S6059S |
| 18443 | G | T | | ORF1ab | nonsynonymous_R6060I |
| 18444 | A | T | | ORF1ab | nonsynonymous_R6060S |
| 18448 | A | T | | ORF1ab | nonsynonymous_S6062C |
| 18449 | G | T | | ORF1ab | nonsynonymous_S6062I |
| 18452 | C | T | | ORF1ab | nonsynonymous_A6063V |
| 18457 | C | T | | ORF1ab | nonsynonymous_P6065S |
| 18458 | C | T | | ORF1ab | nonsynonymous_P6065L |
| 18461 | C | T | | ORF1ab | nonsynonymous_P6066L |
| 18462 | G | A | | ORF1ab | synonymous_P6066P |
| 18463 | C | T | | ORF1ab | nonsynonymous_P6067S |
| 18471 | T | C | | ORF1ab | synonymous_D6069D |
| 18472 | C | T | | ORF1ab | stopgain_Q6070X |
| 18473 | A | T | | ORF1ab | nonsynonymous_Q6070L |
| 18475 | T | A | | ORF1ab | nonsynonymous_F6071I |

| | | | | | |
|---|---|---|---|---|---|
| 18477 | T | C | | ORF1ab | synonymous_F6071F |
| 18481 | C | T | | ORF1ab | nonsynonymous_H6073Y |
| 18483 | C | T | | ORF1ab | synonymous_H6073H |
| 18484 | C | T | | ORF1ab | nonsynonymous_L6074F |
| 18485 | T | C | | ORF1ab | nonsynonymous_L6074P |
| 18486 | C | T | | ORF1ab | synonymous_L6074L |
| 18489 | A | C | | ORF1ab | synonymous_I6075I |
| 18501 | C | T | | ORF1ab | synonymous_Y6079Y |
| 18507 | A | G | | ORF1ab | synonymous_G6081G |
| 18508 | C | T | | ORF1ab | nonsynonymous_L6082F |
| 18511 | C | T | G | ORF1ab | nonsynonymous_P6083S;<br>nonsynonymous_P6083A |
| 18516 | G | A | | ORF1ab | stopgain_W6084X |
| 18526 | C | T | | ORF1ab | nonsynonymous_R6088C |
| 18527 | G | T | | ORF1ab | nonsynonymous_R6088L |
| 18536 | T | C | | ORF1ab | nonsynonymous_I6091T |
| 18541 | C | T | | ORF1ab | stopgain_Q6093X |
| 18546 | G | T | | ORF1ab | nonsynonymous_M6094I |
| 18551 | G | T | | ORF1ab | nonsynonymous_S6096I |
| 18552 | T | C | G | ORF1ab | synonymous_S6096S;<br>nonsynonymous_S6096R |
| 18555 | C | T | | ORF1ab | synonymous_D6097D |
| 18559 | C | T | | ORF1ab | nonsynonymous_L6099F |
| 18561 | T | A | | ORF1ab | synonymous_L6099L |
| 18568 | C | T | G | ORF1ab | nonsynonymous_L6102F;<br>nonsynonymous_L6102V |
| 18572 | C | T | | ORF1ab | nonsynonymous_S6103F |
| 18580 | G | T | | ORF1ab | nonsynonymous_V6106F |
| 18582 | C | T | | ORF1ab | synonymous_V6106V |
| 18583 | G | A | T | ORF1ab | nonsynonymous_V6107I;<br>nonsynonymous_V6107L |

| | | | | | |
|---|---|---|---|---|---|
| 18584 | T | C | | ORF1ab | nonsynonymous_V6107A |
| 18589 | G | T | | ORF1ab | nonsynonymous_V6109F |
| 18617 | C | G | | ORF1ab | nonsynonymous_T6118R |
| 18629 | A | T | | ORF1ab | nonsynonymous_Y6122F |
| 18634 | G | T | | ORF1ab | nonsynonymous_V6124L |
| 18636 | G | A | T | ORF1ab | synonymous_V6124V;<br>synonymous_V6124V |
| 18639 | A | G | | ORF1ab | synonymous_K6125K |
| 18643 | G | T | | ORF1ab | stopgain_G6127X |
| 18647 | C | T | | ORF1ab | nonsynonymous_P6128L |
| 18651 | G | T | | ORF1ab | nonsynonymous_E6129D |
| 18657 | C | T | | ORF1ab | synonymous_T6131T |
| 18664 | C | T | | ORF1ab | synonymous_L6134L |
| 18676 | C | T | | ORF1ab | nonsynonymous_R6138C |
| 18679 | G | C | | ORF1ab | nonsynonymous_A6139P |
| 18680 | C | T | | ORF1ab | nonsynonymous_A6139V |
| 18685 | T | A | | ORF1ab | nonsynonymous_C6141S |
| 18687 | C | T | | ORF1ab | synonymous_C6141C |
| 18688 | T | C | | ORF1ab | nonsynonymous_F6142L |
| 18689 | T | C | | ORF1ab | nonsynonymous_F6142S |
| 18691 | T | C | | ORF1ab | nonsynonymous_S6143P |
| 18692 | C | T | | ORF1ab | nonsynonymous_S6143F |
| 18695 | C | T | | ORF1ab | nonsynonymous_T6144I |
| 18697 | G | A | | ORF1ab | nonsynonymous_A6145T |
| 18703 | G | T | | ORF1ab | nonsynonymous_D6147Y |
| 18707 | C | T | | ORF1ab | nonsynonymous_T6148I |
| 18710 | A | G | | ORF1ab | nonsynonymous_Y6149C |
| 18714 | C | T | | ORF1ab | synonymous_A6150A |
| 18736 | T | C | | ORF1ab | nonsynonymous_F6158L |
| 18744 | C | T | | ORF1ab | synonymous_Y6160Y |
| 18745 | G | T | | ORF1ab | nonsynonymous_V6161F |

| | | | | | |
|---|---|---|---|---|---|
| 18747 | C | T | | ORF1ab | synonymous_V6161V |
| 18756 | G | T | | ORF1ab | synonymous_P6164P |
| 18768 | T | C | | ORF1ab | synonymous_D6168D |
| 18775 | C | T | | ORF1ab | stopgain_Q6171X |
| 18788 | C | T | | ORF1ab | nonsynonymous_T6175I |
| 18789 | A | C | | ORF1ab | synonymous_T6175T |
| 18795 | C | T | | ORF1ab | synonymous_N6177N |
| 18803 | G | T | | ORF1ab | nonsynonymous_S6180I |
| 18804 | C | A | T | ORF1ab | nonsynonymous_S6180R; synonymous_S6180S |
| 18807 | C | T | | ORF1ab | synonymous_N6181N |
| 18814 | C | T | | ORF1ab | synonymous_L6184L |
| 18816 | G | C | | ORF1ab | synonymous_L6184L |
| 18817 | T | A | | ORF1ab | nonsynonymous_Y6185N |
| 18828 | C | T | | ORF1ab | synonymous_V6188V |
| 18835 | A | C | | ORF1ab | nonsynonymous_N6191H |
| 18843 | T | C | | ORF1ab | synonymous_H6193H |
| 18847 | G | T | | ORF1ab | nonsynonymous_A6195S |
| 18857 | A | G | | ORF1ab | nonsynonymous_D6198G |
| 18859 | G | T | | ORF1ab | nonsynonymous_A6199S |
| 18873 | G | T | | ORF1ab | nonsynonymous_R6203S |
| 18877 | C | T | | ORF1ab | synonymous_L6205L |
| 18880 | G | A | | ORF1ab | nonsynonymous_A6206T |
| 18881 | C | T | | ORF1ab | nonsynonymous_A6206V |
| 18883 | G | A | T | ORF1ab | nonsynonymous_V6207I; nonsynonymous_V6207F |
| 18885 | C | T | | ORF1ab | synonymous_V6207V |
| 18904 | C | T | | ORF1ab | nonsynonymous_R6214C |
| 18909 | T | C | | ORF1ab | synonymous_V6215V |
| 18922 | G | T | | ORF1ab | stopgain_E6220X |
| 18923 | A | G | | ORF1ab | nonsynonymous_E6220G |

| | | | | | |
|---|---|---|---|---|---|
| 18928 | C | T | | ORF1ab | nonsynonymous_P6222S |
| 18929 | C | T | | ORF1ab | nonsynonymous_P6222L |
| 18946 | C | T | | ORF1ab | synonymous_L6228L |
| 18959 | C | T | | ORF1ab | nonsynonymous_A6232V |
| 18969 | A | G | | ORF1ab | synonymous_R6235R |
| 18985 | G | T | | ORF1ab | nonsynonymous_V6241F |
| 18988 | G | T | | ORF1ab | nonsynonymous_V6242F |
| 18998 | C | T | | ORF1ab | nonsynonymous_A6245V |
| 19001 | T | C | | ORF1ab | nonsynonymous_L6246S |
| 19006 | G | A | T | ORF1ab | nonsynonymous_A6248T; nonsynonymous_A6248S |
| 19009 | G | T | | ORF1ab | nonsynonymous_D6249Y |
| 19011 | C | T | | ORF1ab | synonymous_D6249D |
| 19016 | T | C | | ORF1ab | nonsynonymous_F6251S |
| 19017 | C | T | | ORF1ab | synonymous_F6251F |
| 19018 | C | T | | ORF1ab | nonsynonymous_P6252S |
| 19021 | G | T | | ORF1ab | nonsynonymous_V6253F |
| 19024 | C | T | | ORF1ab | nonsynonymous_L6254F |
| 19029 | C | G | | ORF1ab | nonsynonymous_H6255Q |
| 19035 | T | A | | ORF1ab | synonymous_I6257I |
| 19043 | C | T | | ORF1ab | nonsynonymous_P6260L |
| 19044 | T | C | | ORF1ab | synonymous_P6260P |
| 19045 | A | T | | ORF1ab | stopgain_K6261X |
| 19049 | C | T | | ORF1ab | nonsynonymous_A6262V |
| 19052 | T | C | | ORF1ab | nonsynonymous_I6263T |
| 19056 | G | C | | ORF1ab | nonsynonymous_K6264N |
| 19064 | C | T | | ORF1ab | nonsynonymous_P6267L |
| 19066 | C | T | | ORF1ab | stopgain_Q6268X |
| 19070 | C | A | | ORF1ab | nonsynonymous_A6269D |
| 19071 | T | A | | ORF1ab | synonymous_A6269A |
| 19072 | G | T | | ORF1ab | nonsynonymous_D6270Y |

| | | | | | |
|---|---|---|---|---|---|
| 19073 | A | G | | ORF1ab | nonsynonymous_D6270G |
| 19078 | G | A | | ORF1ab | nonsynonymous_E6272K |
| 19083 | G | A | | ORF1ab | stopgain_W6273X |
| 19084 | A | T | | ORF1ab | stopgain_K6274X |
| 19086 | G | T | | ORF1ab | nonsynonymous_K6274N |
| 19089 | C | T | | ORF1ab | synonymous_F6275F |
| 19097 | C | T | | ORF1ab | nonsynonymous_A6278V |
| 19099 | C | T | | ORF1ab | stopgain_Q6279X |
| 19102 | C | T | | ORF1ab | nonsynonymous_P6280S |
| 19109 | G | T | | ORF1ab | nonsynonymous_S6282I |
| 19113 | C | T | | ORF1ab | synonymous_D6283D |
| 19118 | C | T | | ORF1ab | nonsynonymous_A6285V |
| 19135 | T | C | | ORF1ab | synonymous_L6291L |
| 19138 | T | C | G | ORF1ab | nonsynonymous_F6292L; nonsynonymous_F6292V |
| 19143 | T | C | | ORF1ab | synonymous_Y6293Y |
| 19148 | A | G | | ORF1ab | nonsynonymous_Y6295C |
| 19151 | C | T | | ORF1ab | nonsynonymous_A6296V |
| 19152 | C | T | | ORF1ab | synonymous_A6296A |
| 19156 | C | T | | ORF1ab | nonsynonymous_H6298Y |
| 19164 | C | T | | ORF1ab | synonymous_D6300D |
| 19169 | T | C | | ORF1ab | nonsynonymous_F6302S |
| 19170 | C | T | | ORF1ab | synonymous_F6302F |
| 19172 | C | T | | ORF1ab | nonsynonymous_T6303I |
| 19175 | A | G | | ORF1ab | nonsynonymous_D6304G |
| 19176 | T | G | | ORF1ab | nonsynonymous_D6304E |
| 19178 | G | T | | ORF1ab | nonsynonymous_G6305V |
| 19181 | T | A | | ORF1ab | nonsynonymous_V6306E |
| 19185 | C | T | | ORF1ab | synonymous_C6307C |
| 19186 | C | T | | ORF1ab | synonymous_L6308L |
| 19196 | A | G | | ORF1ab | nonsynonymous_N6311S |

| | | | | | |
|---|---|---|---|---|---|
| 19202 | A | T | | ORF1ab | nonsynonymous_N6313I |
| 19209 | T | C | | ORF1ab | synonymous_D6315D |
| 19211 | G | T | | ORF1ab | nonsynonymous_R6316I |
| 19217 | C | T | | ORF1ab | nonsynonymous_P6318L |
| 19219 | G | C | T | ORF1ab | nonsynonymous_A6319P; nonsynonymous_A6319S |
| 19220 | C | T | | ORF1ab | nonsynonymous_A6319V |
| 19226 | C | T | | ORF1ab | nonsynonymous_S6321F |
| 19235 | G | T | | ORF1ab | nonsynonymous_C6324F |
| 19238 | G | T | | ORF1ab | nonsynonymous_R6325I |
| 19245 | C | T | | ORF1ab | synonymous_D6327D |
| 19247 | C | T | | ORF1ab | nonsynonymous_T6328I |
| 19252 | G | C | T | ORF1ab | nonsynonymous_V6330L; nonsynonymous_V6330L |
| 19254 | G | T | | ORF1ab | synonymous_V6330V |
| 19255 | C | T | | ORF1ab | synonymous_L6331L |
| 19259 | C | T | | ORF1ab | nonsynonymous_S6332F |
| 19263 | C | T | | ORF1ab | synonymous_N6333N |
| 19264 | C | T | | ORF1ab | nonsynonymous_L6334F |
| 19268 | A | G | | ORF1ab | nonsynonymous_N6335S |
| 19269 | C | T | | ORF1ab | synonymous_N6335N |
| 19272 | G | A | | ORF1ab | synonymous_L6336L |
| 19273 | C | T | | ORF1ab | nonsynonymous_P6337S |
| 19274 | C | T | | ORF1ab | nonsynonymous_P6337L |
| 19275 | T | A | | ORF1ab | synonymous_P6337P |
| 19313 | C | T | | ORF1ab | nonsynonymous_A6350V |
| 19322 | C | T | | ORF1ab | nonsynonymous_T6353I |
| 19329 | T | C | | ORF1ab | synonymous_A6355A |
| 19332 | T | A | | ORF1ab | nonsynonymous_F6356L |
| 19355 | T | A | | ORF1ab | stopgain_L6364X |
| 19360 | C | T | | ORF1ab | stopgain_Q6366X |

| | | | | | |
|---|---|---|---|---|---|
| 19366 | C | T | | ORF1ab | nonsynonymous_P6368S |
| 19367 | C | T | | ORF1ab | nonsynonymous_P6368L |
| 19374 | C | T | | ORF1ab | synonymous_F6370F |
| 19386 | C | T | | ORF1ab | synonymous_D6374D |
| 19391 | C | T | G | ORF1ab | nonsynonymous_P6376L; nonsynonymous_P6376R |
| 19396 | G | T | | ORF1ab | stopgain_E6378X |
| 19406 | G | A | | ORF1ab | nonsynonymous_G6381E |
| 19417 | G | T | | ORF1ab | nonsynonymous_V6385L |
| 19420 | T | C | | ORF1ab | nonsynonymous_S6386P |
| 19446 | G | T | | ORF1ab | nonsynonymous_K6394N |
| 19454 | C | T | | ORF1ab | nonsynonymous_T6397M |
| 19470 | C | T | | ORF1ab | synonymous_C6402C |
| 19478 | G | T | | ORF1ab | nonsynonymous_G6405V |
| 19480 | G | T | | ORF1ab | nonsynonymous_G6406C |
| 19488 | C | T | | ORF1ab | synonymous_V6408V |
| 19491 | T | C | | ORF1ab | synonymous_C6409C |
| 19499 | A | T | | ORF1ab | nonsynonymous_H6412L |
| 19502 | C | T | | ORF1ab | nonsynonymous_A6413V |
| 19509 | G | A | | ORF1ab | synonymous_E6415E |
| 19512 | C | T | | ORF1ab | synonymous_Y6416Y |
| 19518 | G | T | | ORF1ab | nonsynonymous_L6418F |
| 19522 | C | T | | ORF1ab | nonsynonymous_L6420F |
| 19524 | C | T | | ORF1ab | synonymous_L6420L |
| 19542 | G | T | | ORF1ab | nonsynonymous_M6426I |
| 19547 | C | T | | ORF1ab | nonsynonymous_S6428L |
| 19553 | G | T | | ORF1ab | nonsynonymous_G6430V |
| 19576 | C | T | | ORF1ab | stopgain_Q6438X |
| 19583 | A | C | | ORF1ab | nonsynonymous_D6440A |
| 19593 | C | T | | ORF1ab | synonymous_N6443N |
| 19599 | G | T | | ORF1ab | nonsynonymous_W6445C |

| | | | | | |
|---|---|---|---|---|---|
| 19607 | T | C | | ORF1ab | nonsynonymous_F6448S |
| 19610 | C | T | | ORF1ab | nonsynonymous_T6449I |
| 19621 | A | G | | ORF1ab | nonsynonymous_S6453G |
| 19623 | T | C | | ORF1ab | synonymous_S6453S |
| 19627 | G | A | | ORF1ab | nonsynonymous_E6455K |
| 19637 | C | T | | ORF1ab | nonsynonymous_A6458V |
| 19639 | T | C | | ORF1ab | nonsynonymous_F6459L |
| 19644 | T | C | | ORF1ab | synonymous_N6460N |
| 19648 | G | T | | ORF1ab | nonsynonymous_V6462L |
| 19649 | T | C | | ORF1ab | nonsynonymous_V6462A |
| 19653 | T | C | | ORF1ab | synonymous_N6463N |
| 19656 | G | T | | ORF1ab | nonsynonymous_K6464N |
| 19662 | C | T | | ORF1ab | synonymous_H6466H |
| 19677 | G | T | | ORF1ab | nonsynonymous_Q6471H |
| 19684 | G | T | | ORF1ab | nonsynonymous_V6474L |
| 19689 | A | T | | ORF1ab | synonymous_P6475P |
| 19695 | T | G | | ORF1ab | synonymous_S6477S |
| 19697 | T | C | | ORF1ab | nonsynonymous_I6478T |
| 19713 | T | C | | ORF1ab | synonymous_V6483V |
| 19716 | C | T | | ORF1ab | synonymous_Y6484Y |
| 19718 | C | A | | ORF1ab | nonsynonymous_T6485K |
| 19741 | G | A | | ORF1ab | nonsynonymous_E6493K |
| 19744 | T | C | | ORF1ab | synonymous_L6494L |
| 19763 | C | T | | ORF1ab | nonsynonymous_T6500I |
| 19783 | T | C | | ORF1ab | nonsynonymous_F6507L |
| 19788 | G | T | | ORF1ab | nonsynonymous_E6508D |
| 19794 | G | T | | ORF1ab | nonsynonymous_W6510C |
| 19797 | T | C | | ORF1ab | synonymous_A6511A |
| 19800 | G | T | | ORF1ab | nonsynonymous_K6512N |
| 19801 | C | T | | ORF1ab | nonsynonymous_R6513C |
| 19803 | C | T | | ORF1ab | synonymous_R6513R |
| 19807 | A | G | | ORF1ab | nonsynonymous_I6515V |

| | | | | | |
|---|---|---|---|---|---|
| 19809 | T | A | | ORF1ab | synonymous_I6515I |
| 19813 | C | T | | ORF1ab | nonsynonymous_P6517S |
| 19814 | C | T | | ORF1ab | nonsynonymous_P6517L |
| 19819 | C | T | | ORF1ab | nonsynonymous_P6519S |
| 19836 | C | T | | ORF1ab | synonymous_L6524L |
| 19839 | T | C | | ORF1ab | synonymous_N6525N |
| 19845 | G | T | | ORF1ab | nonsynonymous_L6527F |
| 19846 | G | T | | ORF1ab | nonsynonymous_G6528C |
| 19854 | C | T | | ORF1ab | synonymous_D6530D |
| 19862 | C | T | | ORF1ab | nonsynonymous_A6533V |
| 19866 | T | C | | ORF1ab | synonymous_N6534N |
| 19868 | C | T | | ORF1ab | nonsynonymous_T6535I |
| 19870 | G | T | | ORF1ab | nonsynonymous_V6536L |
| 19872 | G | T | | ORF1ab | synonymous_V6536V |
| 19875 | C | T | | ORF1ab | synonymous_I6537I |
| 19876 | T | C | | ORF1ab | nonsynonymous_W6538R |
| 19877 | G | T | | ORF1ab | nonsynonymous_W6538L |
| 19881 | C | T | | ORF1ab | synonymous_D6539D |
| 19884 | C | T | | ORF1ab | synonymous_Y6540Y |
| 19889 | G | T | | ORF1ab | nonsynonymous_R6542I |
| 19893 | T | C | | ORF1ab | synonymous_D6543D |
| 19898 | C | T | | ORF1ab | nonsynonymous_P6545L |
| 19900 | G | T | | ORF1ab | nonsynonymous_A6546S |
| 19910 | C | A | T | ORF1ab | nonsynonymous_S6549Y; nonsynonymous_S6549F |
| 19947 | G | C | | ORF1ab | nonsynonymous_K6561N |
| 19951 | C | T | | ORF1ab | nonsynonymous_P6563S |
| 19957 | G | A | | ORF1ab | nonsynonymous_E6565K |
| 19961 | C | A | T | ORF1ab | nonsynonymous_T6566K; nonsynonymous_T6566M |
| 19970 | C | T | | ORF1ab | nonsynonymous_A6569V |

| | | | | | |
|---|---|---|---|---|---|
| 19974 | A | G | | ORF1ab | synonymous_P6570P |
| 19983 | C | T | | ORF1ab | synonymous_V6573V |
| 20001 | T | G | | ORF1ab | synonymous_V6579V |
| 20002 | G | A | | ORF1ab | nonsynonymous_D6580N |
| 20016 | C | T | | ORF1ab | synonymous_D6584D |
| 20030 | C | T | | ORF1ab | nonsynonymous_A6589V |
| 20031 | C | A | T | ORF1ab | synonymous_A6589A;<br>synonymous_A6589A |
| 20032 | C | T | | ORF1ab | nonsynonymous_R6590C |
| 20036 | A | T | | ORF1ab | nonsynonymous_N6591I |
| 20044 | C | T | | ORF1ab | nonsynonymous_L6594F |
| 20056 | G | A | | ORF1ab | nonsynonymous_G6598S |
| 20057 | G | T | | ORF1ab | nonsynonymous_G6598V |
| 20061 | T | A | | ORF1ab | nonsynonymous_S6599R |
| 20063 | T | C | | ORF1ab | nonsynonymous_V6600A |
| 20064 | T | C | | ORF1ab | synonymous_V6600V |
| 20072 | T | C | | ORF1ab | nonsynonymous_L6603S |
| 20078 | C | T | | ORF1ab | nonsynonymous_P6605L |
| 20081 | C | T | | ORF1ab | nonsynonymous_S6606F |
| 20087 | G | T | | ORF1ab | nonsynonymous_G6608V |
| 20092 | A | G | | ORF1ab | nonsynonymous_K6610E |
| 20098 | G | A | | ORF1ab | nonsynonymous_A6612T |
| 20099 | C | T | | ORF1ab | nonsynonymous_A6612V |
| 20104 | C | T | | ORF1ab | nonsynonymous_L6614F |
| 20113 | G | T | | ORF1ab | nonsynonymous_V6617F |
| 20116 | A | G | | ORF1ab | nonsynonymous_T6618A |
| 20126 | G | T | | ORF1ab | nonsynonymous_G6621V |
| 20128 | G | T | | ORF1ab | stopgain_E6622X |
| 20132 | C | T | | ORF1ab | nonsynonymous_A6623V |
| 20134 | G | T | | ORF1ab | nonsynonymous_V6624L |
| 20135 | T | A | | ORF1ab | nonsynonymous_V6624E |

| | | | | | |
|---|---|---|---|---|---|
| 20141 | C | T | | ORF1ab | nonsynonymous_T6626I |
| 20143 | C | T | | ORF1ab | stopgain_Q6627X |
| 20144 | A | T | | ORF1ab | nonsynonymous_Q6627L |
| 20148 | C | A | T | ORF1ab | nonsynonymous_F6628L;<br>synonymous_F6628F |
| 20150 | A | G | | ORF1ab | nonsynonymous_N6629S |
| 20151 | T | C | | ORF1ab | synonymous_N6629N |
| 20156 | A | T | | ORF1ab | nonsynonymous_Y6631F |
| 20165 | T | C | | ORF1ab | nonsynonymous_V6634A |
| 20176 | G | T | | ORF1ab | nonsynonymous_V6638F |
| 20177 | T | C | | ORF1ab | nonsynonymous_V6638A |
| 20178 | C | T | | ORF1ab | synonymous_V6638V |
| 20180 | A | G | | ORF1ab | nonsynonymous_Q6639R |
| 20199 | C | T | | ORF1ab | synonymous_Y6645Y |
| 20204 | C | T | | ORF1ab | nonsynonymous_T6647I |
| 20210 | G | T | | ORF1ab | nonsynonymous_S6649I |
| 20211 | T | C | | ORF1ab | synonymous_S6649S |
| 20212 | A | T | | ORF1ab | stopgain_R6650X |
| 20221 | C | T | | ORF1ab | stopgain_Q6653X |
| 20224 | G | C | | ORF1ab | nonsynonymous_E6654Q |
| 20227 | T | C | | ORF1ab | nonsynonymous_F6655L |
| 20230 | A | T | | ORF1ab | stopgain_K6656X |
| 20231 | A | G | | ORF1ab | nonsynonymous_K6656R |
| 20233 | C | T | | ORF1ab | nonsynonymous_P6657S |
| 20242 | C | T | | ORF1ab | stopgain_Q6660X |
| 20247 | G | T | | ORF1ab | nonsynonymous_M6661I |
| 20259 | C | T | | ORF1ab | synonymous_F6665F |
| 20268 | A | G | | ORF1ab | synonymous_L6668L |
| 20280 | A | T | | ORF1ab | nonsynonymous_E6672D |
| 20281 | T | C | | ORF1ab | nonsynonymous_F6673L |
| 20283 | C | T | | ORF1ab | synonymous_F6673F |

| | | | | | |
|---|---|---|---|---|---|
| 20287 | G | C | | ORF1ab | nonsynonymous_E6675Q |
| 20320 | C | T | | ORF1ab | nonsynonymous_H6686Y |
| 20335 | G | A | | ORF1ab | nonsynonymous_D6691N |
| 20344 | C | T | | ORF1ab | nonsynonymous_H6694Y |
| 20349 | T | G | | ORF1ab | nonsynonymous_S6695R |
| 20350 | C | T | | ORF1ab | stopgain_Q6696X |
| 20352 | G | C | | ORF1ab | nonsynonymous_Q6696H |
| 20356 | G | T | | ORF1ab | nonsynonymous_G6698C |
| 20367 | T | C | | ORF1ab | synonymous_H6701H |
| 20383 | G | A | | ORF1ab | nonsynonymous_A6707T |
| 20384 | C | T | | ORF1ab | nonsynonymous_A6707V |
| 20389 | C | T | | ORF1ab | nonsynonymous_R6709C |
| 20402 | C | T | | ORF1ab | nonsynonymous_S6713L |
| 20405 | C | T | | ORF1ab | nonsynonymous_P6714L |
| 20410 | G | A | | ORF1ab | nonsynonymous_E6716K |
| 20418 | A | C | | ORF1ab | nonsynonymous_E6718D |
| 20429 | C | T | | ORF1ab | nonsynonymous_P6722L |
| 20433 | G | T | | ORF1ab | nonsynonymous_M6723I |
| 20436 | C | T | | ORF1ab | synonymous_D6724D |
| 20438 | G | T | | ORF1ab | nonsynonymous_S6725I |
| 20451 | C | T | | ORF1ab | synonymous_N6729N |
| 20459 | T | C | | ORF1ab | nonsynonymous_I6732T |
| 20464 | G | T | | ORF1ab | nonsynonymous_D6734Y |
| 20468 | C | T | | ORF1ab | nonsynonymous_A6735V |
| 20470 | C | T | | ORF1ab | stopgain_Q6736X |
| 20474 | C | T | | ORF1ab | nonsynonymous_T6737I |
| 20475 | A | G | | ORF1ab | synonymous_T6737T |
| 20476 | G | T | | ORF1ab | nonsynonymous_G6738C |
| 20480 | C | T | | ORF1ab | nonsynonymous_S6739L |
| 20481 | A | G | | ORF1ab | synonymous_S6739S |
| 20483 | C | T | | ORF1ab | nonsynonymous_S6740F |
| 20486 | A | C | | ORF1ab | nonsynonymous_K6741T |

| | | | | | |
|---|---|---|---|---|---|
| 20487 | G | T | | ORF1ab | nonsynonymous_K6741N |
| 20494 | T | C | | ORF1ab | nonsynonymous_C6744R |
| 20515 | C | T | | ORF1ab | nonsynonymous_L6751F |
| 20527 | G | T | | ORF1ab | nonsynonymous_V6755F |
| 20530 | G | C | | ORF1ab | nonsynonymous_E6756Q |
| 20545 | C | T | | ORF1ab | stopgain_Q6761X |
| 20564 | C | T | | ORF1ab | nonsynonymous_S6767F |
| 20568 | G | A | T | ORF1ab | synonymous_K6768K;<br>nonsynonymous_K6768N |
| 20569 | G | T | | ORF1ab | nonsynonymous_V6769F |
| 20578 | G | T | | ORF1ab | nonsynonymous_V6772L |
| 20580 | G | T | | ORF1ab | synonymous_V6772V |
| 20582 | C | T | | ORF1ab | nonsynonymous_T6773I |
| 20586 | T | G | | ORF1ab | nonsynonymous_I6774M |
| 20592 | T | C | | ORF1ab | synonymous_Y6776Y |
| 20594 | C | T | | ORF1ab | nonsynonymous_T6777I |
| 20596 | G | A | | ORF1ab | nonsynonymous_E6778K |
| 20603 | C | T | | ORF1ab | nonsynonymous_S6780L |
| 20604 | A | G | | ORF1ab | synonymous_S6780S |
| 20613 | T | C | | ORF1ab | synonymous_L6783L |
| 20627 | G | T | | ORF1ab | nonsynonymous_G6788V |
| 20628 | C | T | | ORF1ab | synonymous_G6788G |
| 20636 | A | G | | ORF1ab | nonsynonymous_E6791G |
| 20639 | C | T | | ORF1ab | nonsynonymous_T6792I |
| 20647 | C | A | | ORF1ab | nonsynonymous_P6795T |
| 20656 | C | T | | ORF1ab | stopgain_Q6798X |
| 20663 | G | A | T | ORF1ab | nonsynonymous_S6800N;<br>nonsynonymous_S6800I |
| 20667 | A | T | G | ORF1ab | nonsynonymous_Q6801H;<br>synonymous_Q6801Q |
| 20670 | G | T | | ORF1ab | synonymous_A6802A |

| | | | | | |
|---|---|---|---|---|---|
| 20677 | C | T | | ORF1ab | nonsynonymous_P6805S |
| 20678 | C | G | | ORF1ab | nonsynonymous_P6805R |
| 20679 | G | C | | ORF1ab | synonymous_P6805P |
| 20681 | G | T | | ORF1ab | nonsynonymous_G6806V |
| 20683 | G | T | | ORF1ab | nonsynonymous_V6807F |
| 20687 | C | T | | ORF1ab | nonsynonymous_A6808V |
| 20692 | C | T | | ORF1ab | nonsynonymous_P6810S |
| 20697 | T | C | | ORF1ab | synonymous_N6811N |
| 20698 | C | T | | ORF1ab | nonsynonymous_L6812F |
| 20703 | C | T | | ORF1ab | synonymous_Y6813Y |
| 20713 | A | T | | ORF1ab | stopgain_R6817X |
| 20720 | T | C | | ORF1ab | nonsynonymous_L6819P |
| 20731 | T | C | | ORF1ab | nonsynonymous_C6823R |
| 20736 | C | T | | ORF1ab | synonymous_D6824D |
| 20740 | C | T | | ORF1ab | stopgain_Q6826X |
| 20753 | A | G | | ORF1ab | nonsynonymous_D6830G |
| 20755 | A | C | | ORF1ab | nonsynonymous_S6831R |
| 20756 | G | T | | ORF1ab | nonsynonymous_S6831I |
| 20758 | G | T | | ORF1ab | nonsynonymous_A6832S |
| 20759 | C | T | | ORF1ab | nonsynonymous_A6832V |
| 20762 | C | T | | ORF1ab | nonsynonymous_T6833I |
| 20768 | C | T | | ORF1ab | nonsynonymous_P6835L |
| 20770 | A | G | | ORF1ab | nonsynonymous_K6836E |
| 20773 | G | A | | ORF1ab | nonsynonymous_G6837S |
| 20775 | C | T | | ORF1ab | synonymous_G6837G |
| 20781 | G | T | | ORF1ab | nonsynonymous_M6839I |
| 20787 | T | C | | ORF1ab | synonymous_N6841N |
| 20790 | C | T | | ORF1ab | synonymous_V6842V |
| 20792 | C | T | | ORF1ab | nonsynonymous_A6843V |
| 20810 | G | T | | ORF1ab | nonsynonymous_C6849F |
| 20818 | T | G | | ORF1ab | nonsynonymous_L6852V |
| 20820 | A | T | | ORF1ab | nonsynonymous_L6852F |

| | | | | | |
|---|---|---|---|---|---|
| 20823 | C | T | | ORF1ab | synonymous_N6853N |
| 20844 | C | A | T | ORF1ab | synonymous_P6860P; synonymous_P6860P |
| 20845 | T | C | | ORF1ab | nonsynonymous_Y6861H |
| 20846 | A | T | | ORF1ab | nonsynonymous_Y6861F |
| 20849 | A | C | | ORF1ab | nonsynonymous_N6862T |
| 20869 | G | T | | ORF1ab | nonsynonymous_G6869C |
| 20875 | G | T | | ORF1ab | nonsynonymous_G6871C |
| 20879 | C | T | | ORF1ab | nonsynonymous_S6872F |
| 20887 | G | A | | ORF1ab | nonsynonymous_G6875R |
| 20890 | G | A | | ORF1ab | nonsynonymous_V6876I |
| 20910 | T | C | | ORF1ab | synonymous_V6882V |
| 20911 | T | C | | ORF1ab | synonymous_L6883L |
| 20916 | A | G | | ORF1ab | synonymous_R6884R |
| 20922 | G | T | | ORF1ab | nonsynonymous_W6886C |
| 20925 | G | C | | ORF1ab | nonsynonymous_L6887F |
| 20930 | C | T | | ORF1ab | nonsynonymous_T6889M |
| 20931 | G | T | | ORF1ab | synonymous_T6889T |
| 20933 | G | T | | ORF1ab | nonsynonymous_G6890V |
| 20935 | A | G | | ORF1ab | nonsynonymous_T6891A |
| 20936 | C | A | T | ORF1ab | nonsynonymous_T6891K; nonsynonymous_T6891M |
| 20938 | C | T | | ORF1ab | synonymous_L6892L |
| 20940 | G | A | | ORF1ab | synonymous_L6892L |
| 20941 | C | T | | ORF1ab | nonsynonymous_L6893F |
| 20944 | G | T | | ORF1ab | nonsynonymous_V6894F |
| 20946 | C | T | | ORF1ab | synonymous_V6894V |
| 20947 | G | T | | ORF1ab | nonsynonymous_D6895Y |
| 20951 | C | T | | ORF1ab | nonsynonymous_S6896L |
| 20954 | A | C | | ORF1ab | nonsynonymous_D6897A |
| 20956 | C | T | | ORF1ab | nonsynonymous_L6898F |

| | | | | | |
|---|---|---|---|---|---|
| 20966 | T | C | | ORF1ab | nonsynonymous_F6901S |
| 20978 | C | T | | ORF1ab | nonsynonymous_A6905V |
| 20987 | C | T | | ORF1ab | nonsynonymous_T6908I |
| 20991 | G | C | | ORF1ab | nonsynonymous_L6909F |
| 20994 | T | G | | ORF1ab | nonsynonymous_I6910M |
| 20995 | G | A | T | ORF1ab | nonsynonymous_G6911S; nonsynonymous_G6911C |
| 21000 | T | C | | ORF1ab | synonymous_D6912D |
| 21004 | G | T | | ORF1ab | nonsynonymous_A6914S |
| 21005 | C | T | | ORF1ab | nonsynonymous_A6914V |
| 21008 | C | T | | ORF1ab | nonsynonymous_T6915I |
| 21009 | T | C | | ORF1ab | synonymous_T6915T |
| 21029 | G | A | | ORF1ab | stopgain_W6922X |
| 21034 | C | T | | ORF1ab | nonsynonymous_L6924F |
| 21036 | C | T | | ORF1ab | synonymous_L6924L |
| 21042 | T | C | | ORF1ab | synonymous_I6926I |
| 21049 | A | T | | ORF1ab | nonsynonymous_M6929L |
| 21057 | C | T | | ORF1ab | synonymous_D6931D |
| 21058 | C | T | | ORF1ab | nonsynonymous_P6932S |
| 21065 | C | T | | ORF1ab | nonsynonymous_T6934I |
| 21077 | C | T | | ORF1ab | nonsynonymous_T6938I |
| 21082 | G | A | | ORF1ab | nonsynonymous_E6940K |
| 21084 | A | G | | ORF1ab | synonymous_E6940E |
| 21090 | C | T | | ORF1ab | synonymous_D6942D |
| 21096 | A | T | | ORF1ab | nonsynonymous_K6944N |
| 21098 | A | T | | ORF1ab | nonsynonymous_E6945V |
| 21099 | G | C | T | ORF1ab | nonsynonymous_E6945D; nonsynonymous_E6945D |
| 21100 | G | T | | ORF1ab | nonsynonymous_G6946C |
| 21101 | G | T | | ORF1ab | nonsynonymous_G6946V |
| 21110 | C | T | | ORF1ab | nonsynonymous_T6949I |

| | | | | | |
|---|---|---|---|---|---|
| 21114 | C | T | | ORF1ab | synonymous_Y6950Y |
| 21118 | T | G | | ORF1ab | nonsynonymous_C6952G |
| 21122 | G | T | | ORF1ab | nonsynonymous_G6953V |
| 21123 | G | A | T | ORF1ab | synonymous_G6953G; synonymous_G6953G |
| 21131 | A | G | | ORF1ab | nonsynonymous_Q6956R |
| 21137 | A | G | | ORF1ab | nonsynonymous_K6958R |
| 21139 | C | T | | ORF1ab | synonymous_L6959L |
| 21143 | C | T | | ORF1ab | nonsynonymous_A6960V |
| 21145 | C | T | | ORF1ab | nonsynonymous_L6961F |
| 21147 | T | C | G | ORF1ab | synonymous_L6961L; synonymous_L6961L |
| 21182 | C | T | | ORF1ab | nonsynonymous_S6973F |
| 21196 | C | A | T | ORF1ab | nonsynonymous_L6978I; nonsynonymous_L6978F |
| 21198 | T | C | | ORF1ab | synonymous_L6978L |
| 21207 | C | T | | ORF1ab | synonymous_L6981L |
| 21214 | C | T | | ORF1ab | nonsynonymous_H6984Y |
| 21219 | C | T | | ORF1ab | synonymous_F6985F |
| 21225 | G | T | | ORF1ab | nonsynonymous_W6987C |
| 21260 | C | T | | ORF1ab | nonsynonymous_S6999L |
| 21267 | A | G | | ORF1ab | synonymous_E7001E |
| 21281 | G | T | | ORF1ab | nonsynonymous_G7006V |
| 21301 | C | A | | ORF1ab | nonsynonymous_P7013T |
| 21305 | G | T | | ORF1ab | nonsynonymous_R7014L |
| 21306 | C | T | | ORF1ab | synonymous_R7014R |
| 21320 | G | T | | ORF1ab | nonsynonymous_G7019V |
| 21331 | C | T | | ORF1ab | nonsynonymous_H7023Y |
| 21334 | G | T | | ORF1ab | nonsynonymous_A7024S |
| 21364 | C | T | | ORF1ab | nonsynonymous_P7034S |
| 21365 | C | T | | ORF1ab | nonsynonymous_P7034L |

| | | | | | |
|---|---|---|---|---|---|
| 21394 | G | T | | ORF1ab | nonsynonymous_D7044Y |
| 21402 | T | A | | ORF1ab | nonsynonymous_S7046R |
| 21408 | T | C | | ORF1ab | synonymous_F7048F |
| 21428 | C | T | | ORF1ab | nonsynonymous_T7055I |
| 21430 | G | A | | ORF1ab | nonsynonymous_A7056T |
| 21440 | C | T | | ORF1ab | nonsynonymous_S7059F |
| 21448 | G | T | | ORF1ab | stopgain_E7062X |
| 21452 | G | A | | ORF1ab | nonsynonymous_G7063D |
| 21454 | C | A | | ORF1ab | nonsynonymous_Q7064K |
| 21489 | A | T | | ORF1ab | nonsynonymous_K7075N |
| 21497 | T | C | | ORF1ab | nonsynonymous_L7078P |
| 21511 | A | G | | ORF1ab | nonsynonymous_N7083D |
| 21515 | A | G | | ORF1ab | nonsynonymous_N7084S |
| 21516 | C | T | | ORF1ab | synonymous_N7084N |
| 21533 | G | T | | ORF1ab | nonsynonymous_S7090I |
| 21538 | G | T | | ORF1ab | nonsynonymous_V7092F |
| 21543 | T | C | | ORF1ab | synonymous_L7093L |
| 21549 | C | T | | ORF1ab | synonymous_N7095N |
| 21552 | C | T | | ORF1ab | synonymous_N7096N |
| 21561 | C | T | | intergenic | |
| 21568 | T | A | | S | nonsynonymous_F2L |
| 21575 | C | T | | S | nonsynonymous_L5F |
| 21586 | G | T | | S | nonsynonymous_L8F |
| 21587 | C | T | | S | nonsynonymous_P9S |
| 21588 | C | T | | S | nonsynonymous_P9L |
| 21590 | C | T | | S | synonymous_L10L |
| 21595 | C | T | | S | synonymous_V11V |
| 21596 | T | C | | S | nonsynonymous_S12P |
| 21600 | G | T | | S | nonsynonymous_S13I |
| 21604 | G | T | | S | nonsynonymous_Q14H |
| 21606 | G | T | | S | nonsynonymous_C15F |
| 21614 | C | T | | S | nonsynonymous_L18F |

| | | | | | |
|---|---|---|---|---|---|
| 21624 | G | A | T | S | nonsynonymous_R21K; nonsynonymous_R21I |
| 21628 | T | G | | S | synonymous_T22T |
| 21630 | A | G | | S | nonsynonymous_Q23R |
| 21632 | T | C | | S | synonymous_L24L |
| 21637 | C | T | | S | synonymous_P25P |
| 21641 | G | T | | S | nonsynonymous_A27S |
| 21642 | C | T | | S | nonsynonymous_A27V |
| 21648 | C | T | | S | nonsynonymous_T29I |
| 21649 | T | C | | S | synonymous_T29T |
| 21654 | C | T | | S | nonsynonymous_S31F |
| 21658 | C | T | | S | synonymous_F32F |
| 21662 | C | T | | S | nonsynonymous_R34C |
| 21663 | G | T | | S | nonsynonymous_R34L |
| 21667 | T | C | | S | synonymous_G35G |
| 21691 | C | T | | S | synonymous_F43F |
| 21697 | C | T | G | S | synonymous_S45S; synonymous_S45S |
| 21707 | C | T | | S | nonsynonymous_H49Y |
| 21709 | T | C | | S | synonymous_H49H |
| 21711 | C | T | | S | nonsynonymous_S50L |
| 21714 | C | T | | S | nonsynonymous_T51I |
| 21715 | T | C | | S | synonymous_T51T |
| 21716 | C | T | | S | stopgain_Q52X |
| 21721 | C | T | | S | synonymous_D53D |
| 21724 | G | T | | S | nonsynonymous_L54F |
| 21728 | T | C | | S | synonymous_L56L |
| 21731 | C | T | | S | nonsynonymous_P57S |
| 21732 | C | T | | S | nonsynonymous_P57L |
| 21734 | T | C | | S | nonsynonymous_F58L |
| 21742 | C | T | | S | synonymous_S60S |

| | | | | | |
|---|---|---|---|---|---|
| 21746 | G | T | | S | nonsynonymous_V62F |
| 21757 | C | T | | S | synonymous_F65F |
| 21761 | G | T | | S | nonsynonymous_A67S |
| 21762 | C | T | | S | nonsynonymous_A67V |
| 21770 | G | T | | S | nonsynonymous_V70F |
| 21772 | C | T | | S | synonymous_V70V |
| 21781 | C | A | T | S | synonymous_T73T;<br>synonymous_T73T |
| 21785 | G | T | | S | nonsynonymous_G75C |
| 21786 | G | A | T | S | nonsynonymous_G75D;<br>nonsynonymous_G75V |
| 21789 | C | T | | S | nonsynonymous_T76I |
| 21795 | G | A | T | S | nonsynonymous_R78K;<br>nonsynonymous_R78M |
| 21796 | G | C | | S | nonsynonymous_R78S |
| 21802 | T | C | | S | synonymous_D80D |
| 21807 | C | T | | S | nonsynonymous_P82L |
| 21811 | C | T | | S | synonymous_V83V |
| 21815 | C | T | | S | nonsynonymous_P85S |
| 21816 | C | T | | S | nonsynonymous_P85L |
| 21830 | G | T | | S | nonsynonymous_V90F |
| 21831 | T | A | | S | nonsynonymous_V90D |
| 21840 | C | T | | S | nonsynonymous_A93V |
| 21844 | C | T | | S | synonymous_S94S |
| 21846 | C | T | | S | nonsynonymous_T95I |
| 21849 | A | G | | S | nonsynonymous_E96G |
| 21853 | G | A | | S | synonymous_K97K |
| 21855 | C | T | | S | nonsynonymous_S98F |
| 21859 | C | T | | S | synonymous_N99N |
| 21872 | T | C | | S | nonsynonymous_W104R |
| 21875 | A | T | | S | nonsynonymous_I105F |

| | | | | | |
|---|---|---|---|---|---|
| 21879 | T | C | | S | nonsynonymous_F106S |
| 21881 | G | T | | S | nonsynonymous_G107C |
| 21890 | T | C | | S | synonymous_L110L |
| 21905 | C | T | | S | stopgain_Q115X |
| 21908 | T | C | | S | nonsynonymous_S116P |
| 21911 | C | T | | S | synonymous_L117L |
| 21930 | C | T | | S | nonsynonymous_A123V |
| 21931 | T | C | | S | synonymous_A123A |
| 21944 | A | T | | S | nonsynonymous_I128F |
| 21952 | C | T | | S | synonymous_V130V |
| 21953 | T | G | | S | nonsynonymous_C131G |
| 21956 | G | A | | S | nonsynonymous_E132K |
| 21961 | T | C | | S | synonymous_F133F |
| 21974 | G | C | T | S | nonsynonymous_D138H; nonsynonymous_D138Y |
| 21977 | C | T | | S | nonsynonymous_P139S |
| 21978 | C | T | | S | nonsynonymous_P139L |
| 21980 | T | G | | S | nonsynonymous_F140V |
| 21987 | G | C | | S | nonsynonymous_G142A |
| 21995 | T | C | | S | nonsynonymous_Y145H |
| 21998 | C | T | | S | nonsynonymous_H146Y |
| 22000 | C | T | | S | synonymous_H146H |
| 22003 | A | G | | S | synonymous_K147K |
| 22006 | C | T | | S | synonymous_N148N |
| 22009 | C | T | | S | synonymous_N149N |
| 22010 | A | G | | S | nonsynonymous_K150E |
| 22014 | G | C | | S | nonsynonymous_S151T |
| 22016 | T | G | | S | nonsynonymous_W152G |
| 22017 | G | T | | S | nonsynonymous_W152L |
| 22018 | G | T | | S | nonsynonymous_W152C |
| 22026 | G | T | | S | nonsynonymous_S155I |

| | | | | | |
|---|---|---|---|---|---|
| 22028 | G | T | | S | stopgain_E156X |
| 22030 | G | T | | S | nonsynonymous_E156D |
| 22044 | C | T | | S | nonsynonymous_S161F |
| 22047 | G | T | | S | nonsynonymous_S162I |
| 22053 | A | C | | S | nonsynonymous_N164T |
| 22062 | C | T | | S | nonsynonymous_T167I |
| 22075 | C | T | | S | synonymous_V171V |
| 22081 | G | A | | S | synonymous_Q173Q |
| 22083 | C | T | | S | nonsynonymous_P174L |
| 22084 | T | C | | S | synonymous_P174P |
| 22086 | T | C | | S | nonsynonymous_F175S |
| 22088 | C | A | T | S | nonsynonymous_L176I; nonsynonymous_L176F |
| 22093 | G | A | T | S | nonsynonymous_M177I; nonsynonymous_M177I |
| 22099 | T | C | | S | synonymous_L179L |
| 22104 | G | A | | S | nonsynonymous_G181E |
| 22120 | C | T | | S | synonymous_F186F |
| 22122 | A | T | | S | nonsynonymous_K187I |
| 22127 | C | T | | S | nonsynonymous_L189F |
| 22133 | G | A | | S | nonsynonymous_E191K |
| 22141 | G | A | | S | synonymous_V193V |
| 22150 | T | A | | S | nonsynonymous_N196K |
| 22153 | T | C | | S | synonymous_I197I |
| 22176 | C | T | | S | nonsynonymous_S205F |
| 22185 | C | T | | S | nonsynonymous_T208M |
| 22187 | C | T | | S | nonsynonymous_P209S |
| 22202 | C | T | | S | nonsynonymous_R214C |
| 22203 | G | T | | S | nonsynonymous_R214L |
| 22206 | A | G | | S | nonsynonymous_D215G |

| | | | | | |
|---|---|---|---|---|---|
| 22208 | C | A | T | S | nonsynonymous_L216I; nonsynonymous_L216F |
| 22213 | T | C | | S | synonymous_P217P |
| 22214 | C | T | | S | stopgain_Q218X |
| 22218 | G | T | | S | nonsynonymous_G219V |
| 22224 | C | T | | S | nonsynonymous_S221L |
| 22225 | G | A | T | S | synonymous_S221S; synonymous_S221S |
| 22227 | C | T | | S | nonsynonymous_A222V |
| 22235 | C | T | | S | nonsynonymous_P225S |
| 22256 | G | T | | S | nonsynonymous_G232C |
| 22257 | G | T | | S | nonsynonymous_G232V |
| 22264 | C | T | | S | synonymous_N234N |
| 22268 | A | G | | S | nonsynonymous_T236A |
| 22277 | C | A | | S | nonsynonymous_Q239K |
| 22279 | A | T | | S | nonsynonymous_Q239H |
| 22281 | C | T | | S | nonsynonymous_T240I |
| 22286 | C | T | | S | nonsynonymous_L242F |
| 22288 | T | C | | S | synonymous_L242L |
| 22289 | G | T | | S | nonsynonymous_A243S |
| 22290 | C | T | | S | nonsynonymous_A243V |
| 22295 | C | T | | S | nonsynonymous_H245Y |
| 22311 | C | T | | S | nonsynonymous_T250I |
| 22320 | A | G | | S | nonsynonymous_D253G |
| 22323 | C | T | | S | nonsynonymous_S254F |
| 22326 | C | T | | S | nonsynonymous_S255F |
| 22332 | G | T | | S | nonsynonymous_G257V |
| 22335 | G | T | | S | nonsynonymous_W258L |
| 22346 | G | A | | S | nonsynonymous_A262T |
| 22349 | G | T | | S | nonsynonymous_A263S |
| 22356 | A | G | | S | nonsynonymous_Y265C |

| | | | | | |
|---|---|---|---|---|---|
| 22373 | C | T | | S | stopgain_Q271X |
| 22380 | G | C | | S | nonsynonymous_R273T |
| 22424 | G | T | | S | nonsynonymous_A288S |
| 22444 | C | T | | S | synonymous_D294D |
| 22446 | C | T | | S | nonsynonymous_P295L |
| 22452 | C | T | | S | nonsynonymous_S297L |
| 22454 | G | A | | S | nonsynonymous_E298K |
| 22460 | A | G | | S | nonsynonymous_K300E |
| 22468 | G | T | | S | synonymous_T302T |
| 22469 | T | C | | S | synonymous_L303L |
| 22480 | C | T | | S | synonymous_F306F |
| 22482 | C | T | | S | nonsynonymous_T307I |
| 22487 | G | A | C | S | nonsynonymous_E309K;<br>nonsynonymous_E309Q |
| 22506 | C | T | | S | nonsynonymous_T315I |
| 22509 | C | T | | S | nonsynonymous_S316F |
| 22514 | T | C | | S | nonsynonymous_F318L |
| 22523 | C | T | | S | stopgain_Q321X |
| 22526 | C | T | | S | nonsynonymous_P322S |
| 22534 | A | G | | S | synonymous_E324E |
| 22570 | C | T | | S | synonymous_C336C |
| 22574 | T | C | | S | nonsynonymous_F338L |
| 22590 | A | T | | S | nonsynonymous_N343I |
| 22591 | C | T | | S | synonymous_N343N |
| 22596 | C | T | | S | nonsynonymous_T345I |
| 22606 | A | T | | S | synonymous_A348A |
| 22608 | C | T | | S | nonsynonymous_S349F |
| 22612 | T | A | | S | synonymous_V350V |
| 22617 | C | T | | S | nonsynonymous_A352V |
| 22619 | T | C | | S | nonsynonymous_W353R |

| | | | | | |
|---|---|---|---|---|---|
| 22624 | C | T | G | S | synonymous_N354N; nonsynonymous_N354K |
| 22638 | G | A | | S | nonsynonymous_S359N |
| 22655 | T | C | | S | nonsynonymous_Y365H |
| 22659 | C | T | | S | nonsynonymous_S366F |
| 22661 | G | T | | S | nonsynonymous_V367F |
| 22669 | T | A | | S | stopgain_Y369X |
| 22675 | C | T | | S | synonymous_S371S |
| 22678 | A | G | | S | synonymous_A372A |
| 22680 | C | T | | S | nonsynonymous_S373L |
| 22681 | A | T | | S | synonymous_S373S |
| 22685 | T | C | | S | nonsynonymous_S375P |
| 22689 | C | T | | S | nonsynonymous_T376I |
| 22691 | T | C | | S | nonsynonymous_F377L |
| 22695 | A | G | | S | nonsynonymous_K378R |
| 22698 | G | T | | S | nonsynonymous_C379F |
| 22700 | T | C | | S | nonsynonymous_Y380H |
| 22708 | G | T | | S | synonymous_V382V |
| 22711 | T | C | | S | synonymous_S383S |
| 22712 | C | T | | S | nonsynonymous_P384S |
| 22713 | C | T | | S | nonsynonymous_P384L |
| 22716 | C | T | | S | nonsynonymous_T385I |
| 22730 | C | T | | S | nonsynonymous_L390F |
| 22735 | C | T | | S | synonymous_C391C |
| 22737 | T | C | | S | nonsynonymous_F392S |
| 22754 | G | T | | S | nonsynonymous_D398Y |
| 22787 | C | T | | S | stopgain_Q409X |
| 22793 | G | T | | S | nonsynonymous_A411S |
| 22797 | C | A | | S | nonsynonymous_P412Q |
| 22798 | A | G | | S | synonymous_P412P |
| 22802 | C | T | | S | stopgain_Q414X |

| | | | | | |
|---|---|---|---|---|---|
| 22809 | G | A | | S | nonsynonymous_G416E |
| 22823 | T | C | | S | nonsynonymous_Y421H |
| 22858 | C | T | | S | synonymous_C432C |
| 22859 | G | T | | S | nonsynonymous_V433F |
| 22865 | G | T | | S | nonsynonymous_A435S |
| 22866 | C | T | | S | nonsynonymous_A435V |
| 22875 | C | T | | S | nonsynonymous_S438F |
| 22879 | C | A | | S | nonsynonymous_N439K |
| 22886 | G | T | | S | nonsynonymous_D442Y |
| 22890 | C | T | G | S | nonsynonymous_S443F;<br>nonsynonymous_S443C |
| 22899 | G | T | | S | nonsynonymous_G446V |
| 22915 | C | T | | S | synonymous_Y451Y |
| 22918 | G | T | | S | synonymous_L452L |
| 22927 | G | T | | S | nonsynonymous_L455F |
| 22931 | A | T | | S | nonsynonymous_R457W |
| 22939 | T | C | | S | synonymous_S459S |
| 22942 | T | C | | S | synonymous_N460N |
| 22949 | C | T | | S | nonsynonymous_P463S |
| 22950 | C | T | | S | nonsynonymous_P463L |
| 22951 | T | C | | S | synonymous_P463P |
| 22960 | A | G | | S | synonymous_R466R |
| 22965 | T | C | | S | nonsynonymous_I468T |
| 22973 | G | C | | S | nonsynonymous_E471Q |
| 22981 | T | C | | S | synonymous_Y473Y |
| 22982 | C | T | | S | stopgain_Q474X |
| 22983 | A | G | | S | nonsynonymous_Q474R |
| 22986 | C | T | | S | nonsynonymous_A475V |
| 22987 | C | T | | S | synonymous_A475A |
| 22992 | G | C | | S | nonsynonymous_S477T |
| 22995 | C | T | | S | nonsynonymous_T478I |

| | | | | | |
|---|---|---|---|---|---|
| 22997 | C | T | | S | nonsynonymous_P479S |
| 23010 | T | C | | S | nonsynonymous_V483A |
| 23011 | T | C | | S | synonymous_V483V |
| 23016 | G | T | | S | nonsynonymous_G485V |
| 23029 | C | T | | S | synonymous_Y489Y |
| 23030 | T | C | | S | nonsynonymous_F490L |
| 23043 | C | T | | S | nonsynonymous_S494L |
| 23054 | C | T | | S | stopgain_Q498X |
| 23057 | C | A | T | S | nonsynonymous_P499T; nonsynonymous_P499S |
| 23066 | G | T | | S | nonsynonymous_G502C |
| 23081 | C | T | | S | nonsynonymous_P507S |
| 23083 | A | G | | S | synonymous_P507P |
| 23086 | C | T | | S | synonymous_Y508Y |
| 23090 | G | T | | S | nonsynonymous_V510L |
| 23099 | C | T | | S | nonsynonymous_L513F |
| 23104 | T | C | | S | synonymous_S514S |
| 23117 | C | T | | S | nonsynonymous_H519Y |
| 23118 | A | T | | S | nonsynonymous_H519L |
| 23121 | C | T | | S | nonsynonymous_A520V |
| 23122 | A | G | | S | synonymous_A520A |
| 23124 | C | T | | S | nonsynonymous_P521L |
| 23126 | G | T | | S | nonsynonymous_A522S |
| 23127 | C | T | | S | nonsynonymous_A522V |
| 23128 | A | G | | S | synonymous_A522A |
| 23130 | C | T | | S | nonsynonymous_T523I |
| 23155 | T | A | | S | synonymous_T531T |
| 23157 | A | T | | S | nonsynonymous_N532I |
| 23162 | G | T | | S | nonsynonymous_V534F |
| 23185 | C | T | | S | synonymous_F541F |
| 23196 | G | T | | S | nonsynonymous_G545V |

| | | | | | |
|---|---|---|---|---|---|
| 23202 | C | T | | S | nonsynonymous_T547I |
| 23205 | G | T | | S | nonsynonymous_G548V |
| 23206 | C | T | | S | synonymous_G548G |
| 23211 | G | T | | S | nonsynonymous_G550V |
| 23216 | C | T | | S | nonsynonymous_L552F |
| 23226 | C | T | | S | nonsynonymous_S555F |
| 23228 | A | T | | S | nonsynonymous_N556Y |
| 23236 | G | T | | S | nonsynonymous_K558N |
| 23243 | C | T | | S | nonsynonymous_P561S |
| 23244 | C | T | | S | nonsynonymous_P561L |
| 23248 | C | T | | S | synonymous_F562F |
| 23256 | T | C | | S | nonsynonymous_F565S |
| 23260 | C | T | | S | synonymous_G566G |
| 23271 | C | T | | S | nonsynonymous_A570V |
| 23277 | C | T | | S | nonsynonymous_T572I |
| 23280 | C | T | | S | nonsynonymous_T573I |
| 23285 | G | C | T | S | nonsynonymous_A575P; nonsynonymous_A575S |
| 23287 | T | C | | S | synonymous_A575A |
| 23288 | G | T | | S | nonsynonymous_V576F |
| 23289 | T | C | | S | nonsynonymous_V576A |
| 23290 | C | T | | S | synonymous_V576V |
| 23291 | C | T | | S | nonsynonymous_R577C |
| 23295 | A | G | | S | nonsynonymous_D578G |
| 23296 | T | C | | S | synonymous_D578D |
| 23298 | C | T | G | S | nonsynonymous_P579L; nonsynonymous_P579R |
| 23308 | T | C | | S | synonymous_L582L |
| 23311 | G | T | | S | nonsynonymous_E583D |
| 23315 | C | T | | S | nonsynonymous_L585F |
| 23327 | C | T | | S | nonsynonymous_P589S |

| | | | | | |
|---|---|---|---|---|---|
| 23335 | T | A | C | S | synonymous_S591S;<br>synonymous_S591S |
| 23342 | G | A | T | S | nonsynonymous_G594S;<br>nonsynonymous_G594C |
| 23343 | G | T | | S | nonsynonymous_G594V |
| 23347 | C | T | | S | synonymous_V595V |
| 23349 | G | T | | S | nonsynonymous_S596I |
| 23360 | C | T | | S | nonsynonymous_P600S |
| 23367 | C | T | | S | nonsynonymous_T602I |
| 23371 | T | C | | S | synonymous_N603N |
| 23380 | C | T | | S | synonymous_N606N |
| 23388 | C | T | | S | nonsynonymous_A609V |
| 23389 | T | C | | S | synonymous_A609A |
| 23394 | T | C | | S | nonsynonymous_L611P |
| 23397 | A | G | | S | nonsynonymous_Y612C |
| 23399 | C | T | | S | stopgain_Q613X |
| 23400 | A | G | | S | nonsynonymous_Q613R |
| 23401 | G | T | | S | nonsynonymous_Q613H |
| 23402 | G | T | | S | nonsynonymous_D614Y |
| 23403 | A | T | G | S | nonsynonymous_D614V;<br>nonsynonymous_D614G |
| 23410 | C | G | | S | nonsynonymous_N616K |
| 23411 | T | C | | S | nonsynonymous_C617R |
| 23415 | C | T | | S | nonsynonymous_T618I |
| 23416 | A | T | | S | synonymous_T618T |
| 23420 | G | T | | S | nonsynonymous_V620F |
| 23421 | T | C | | S | nonsynonymous_V620A |
| 23422 | C | T | | S | synonymous_V620V |
| 23423 | C | T | | S | nonsynonymous_P621S |
| 23424 | C | T | | S | nonsynonymous_P621L |
| 23426 | G | A | | S | nonsynonymous_V622I |

| | | | | | |
|---|---|---|---|---|---|
| 23430 | C | T | | S | nonsynonymous_A623V |
| 23435 | C | T | | S | nonsynonymous_H625Y |
| 23436 | A | G | | S | nonsynonymous_H625R |
| 23438 | G | C | | S | nonsynonymous_A626P |
| 23439 | C | T | | S | nonsynonymous_A626V |
| 23444 | C | T | | S | stopgain_Q628X |
| 23447 | C | T | | S | nonsynonymous_L629F |
| 23451 | C | T | | S | nonsynonymous_T630I |
| 23469 | A | T | | S | nonsynonymous_Y636F |
| 23481 | C | T | | S | nonsynonymous_S640F |
| 23487 | T | G | | S | nonsynonymous_V642G |
| 23490 | T | C | | S | nonsynonymous_F643S |
| 23498 | C | T | | S | nonsynonymous_R646C |
| 23515 | A | C | | S | synonymous_I651I |
| 23517 | G | T | | S | nonsynonymous_G652V |
| 23519 | G | T | | S | nonsynonymous_A653S |
| 23525 | C | T | | S | nonsynonymous_H655Y |
| 23533 | C | T | | S | synonymous_N657N |
| 23536 | C | T | | S | synonymous_N658N |
| 23538 | C | T | | S | nonsynonymous_S659L |
| 23557 | C | T | | S | synonymous_P665P |
| 23564 | G | A | | S | nonsynonymous_A668T |
| 23575 | C | T | | S | synonymous_C671C |
| 23580 | G | T | | S | nonsynonymous_S673I |
| 23587 | G | C | | S | nonsynonymous_Q675H |
| 23592 | A | T | | S | nonsynonymous_Q677L |
| 23593 | G | T | | S | nonsynonymous_Q677H |
| 23595 | C | T | | S | nonsynonymous_T678I |
| 23601 | C | T | | S | nonsynonymous_S680F |
| 23603 | C | T | | S | nonsynonymous_P681S |
| 23604 | C | T | | S | nonsynonymous_P681L |

| | | | | | |
|---|---|---|---|---|---|
| 23607 | G | A | T | S | nonsynonymous_R682Q; nonsynonymous_R682L |
| 23608 | G | T | | S | synonymous_R682R |
| 23611 | G | T | | S | synonymous_R683R |
| 23612 | G | A | T | S | nonsynonymous_A684T; nonsynonymous_A684S |
| 23625 | C | T | | S | nonsynonymous_A688V |
| 23629 | T | C | | S | synonymous_S689S |
| 23634 | C | T | | S | nonsynonymous_S691F |
| 23635 | C | T | | S | synonymous_S691S |
| 23638 | C | T | | S | synonymous_I692I |
| 23644 | C | T | | S | synonymous_A694A |
| 23655 | C | T | G | S | nonsynonymous_S698L; stopgain_S698X |
| 23660 | G | A | | S | nonsynonymous_G700S |
| 23661 | G | T | | S | nonsynonymous_G700V |
| 23662 | T | C | | S | synonymous_G700G |
| 23663 | G | T | | S | nonsynonymous_A701S |
| 23664 | C | T | | S | nonsynonymous_A701V |
| 23665 | A | T | | S | synonymous_A701A |
| 23670 | A | G | | S | nonsynonymous_N703S |
| 23673 | C | T | | S | nonsynonymous_S704L |
| 23674 | A | T | | S | synonymous_S704S |
| 23678 | G | T | | S | nonsynonymous_A706S |
| 23679 | C | T | | S | nonsynonymous_A706V |
| 23683 | C | T | | S | synonymous_Y707Y |
| 23685 | C | T | | S | nonsynonymous_S708F |
| 23692 | C | T | | S | synonymous_N710N |
| 23698 | T | C | | S | synonymous_I712I |
| 23699 | G | C | | S | nonsynonymous_A713P |
| 23701 | C | T | | S | synonymous_A713A |

| | | | | | |
|---|---|---|---|---|---|
| 23705 | C | T | | S | nonsynonymous_P715S |
| 23707 | C | T | | S | synonymous_P715P |
| 23709 | C | T | | S | nonsynonymous_T716I |
| 23712 | A | T | | S | nonsynonymous_N717I |
| 23718 | C | T | | S | nonsynonymous_T719I |
| 23724 | G | C | | S | nonsynonymous_S721T |
| 23730 | C | T | | S | nonsynonymous_T723I |
| 23731 | C | T | | S | synonymous_T723T |
| 23735 | G | C | | S | nonsynonymous_E725Q |
| 23738 | A | T | | S | nonsynonymous_I726F |
| 23741 | C | T | | S | synonymous_L727L |
| 23745 | C | T | | S | nonsynonymous_P728L |
| 23747 | G | T | | S | nonsynonymous_V729L |
| 23749 | G | T | | S | synonymous_V729V |
| 23755 | G | C | T | S | nonsynonymous_M731I; nonsynonymous_M731I |
| 23757 | C | T | | S | nonsynonymous_T732I |
| 23766 | C | T | | S | nonsynonymous_S735L |
| 23767 | A | G | | S | synonymous_S735S |
| 23768 | G | T | | S | nonsynonymous_V736L |
| 23772 | A | T | | S | nonsynonymous_D737V |
| 23773 | T | C | | S | synonymous_D737D |
| 23774 | T | A | | S | nonsynonymous_C738S |
| 23775 | G | T | | S | nonsynonymous_C738F |
| 23776 | T | C | | S | synonymous_C738C |
| 23778 | C | T | | S | nonsynonymous_T739I |
| 23781 | T | C | G | S | nonsynonymous_M740T; nonsynonymous_M740R |
| 23785 | C | T | | S | synonymous_Y741Y |
| 23786 | A | T | | S | nonsynonymous_I742F |
| 23790 | G | T | | S | nonsynonymous_C743F |

| | | | | | |
|---|---|---|---|---|---|
| 23797 | T | C | | S | synonymous_D745D |
| 23802 | C | T | | S | nonsynonymous_T747I |
| 23810 | A | T | | S | nonsynonymous_S750C |
| 23816 | C | T | | S | nonsynonymous_L752F |
| 23832 | G | T | | S | nonsynonymous_G757V |
| 23833 | C | A | | S | synonymous_G757G |
| 23841 | G | T | | S | nonsynonymous_C760F |
| 23844 | C | T | | S | nonsynonymous_T761I |
| 23849 | T | C | | S | synonymous_L763L |
| 23852 | A | T | | S | nonsynonymous_N764Y |
| 23854 | C | T | | S | synonymous_N764N |
| 23855 | C | T | | S | nonsynonymous_R765C |
| 23856 | G | T | | S | nonsynonymous_R765L |
| 23859 | C | T | | S | nonsynonymous_A766V |
| 23867 | G | T | | S | stopgain_G769X |
| 23868 | G | T | | S | nonsynonymous_G769V |
| 23873 | G | T | | S | nonsynonymous_A771S |
| 23887 | C | T | | S | synonymous_D775D |
| 23893 | C | T | | S | synonymous_N777N |
| 23895 | C | T | G | S | nonsynonymous_T778I;<br>nonsynonymous_T778S |
| 23896 | C | T | | S | synonymous_T778T |
| 23898 | A | G | | S | nonsynonymous_Q779R |
| 23912 | C | T | | S | stopgain_Q784X |
| 23917 | C | T | | S | synonymous_V785V |
| 23922 | A | G | | S | nonsynonymous_Q787R |
| 23924 | A | T | | S | nonsynonymous_I788F |
| 23929 | C | T | | S | synonymous_Y789Y |
| 23934 | C | T | | S | nonsynonymous_T791I |
| 23937 | C | T | | S | nonsynonymous_P792L |
| 23948 | G | T | | S | nonsynonymous_D796Y |

| | | | | | |
|---|---|---|---|---|---|
| 23958 | G | T | | S | nonsynonymous_G799V |
| 23972 | C | T | | S | stopgain_Q804X |
| 23981 | C | T | | S | nonsynonymous_P807S |
| 23982 | C | T | | S | nonsynonymous_P807L |
| 23984 | G | A | T | S | nonsynonymous_D808N; nonsynonymous_D808Y |
| 23985 | A | T | | S | nonsynonymous_D808V |
| 23986 | T | C | | S | synonymous_D808D |
| 23987 | C | T | | S | nonsynonymous_P809S |
| 23991 | C | T | | S | nonsynonymous_S810L |
| 23996 | C | T | | S | nonsynonymous_P812S |
| 24009 | C | T | | S | nonsynonymous_S816L |
| 24013 | T | C | | S | synonymous_F817F |
| 24015 | T | G | | S | nonsynonymous_I818S |
| 24021 | A | G | | S | nonsynonymous_D820G |
| 24023 | C | T | | S | synonymous_L821L |
| 24026 | C | T | | S | nonsynonymous_L822F |
| 24027 | T | C | | S | nonsynonymous_L822P |
| 24034 | C | T | | S | synonymous_N824N |
| 24038 | G | T | | S | nonsynonymous_V826L |
| 24040 | G | T | | S | synonymous_V826V |
| 24042 | C | T | | S | nonsynonymous_T827I |
| 24045 | T | C | | S | nonsynonymous_L828P |
| 24047 | G | A | | S | nonsynonymous_A829T |
| 24053 | G | A | T | S | nonsynonymous_A831T; nonsynonymous_A831S |
| 24055 | T | A | | S | synonymous_A831A |
| 24064 | C | A | | S | synonymous_I834I |
| 24068 | C | T | | S | stopgain_Q836X |
| 24079 | T | C | | S | synonymous_D839D |
| 24081 | G | T | | S | nonsynonymous_C840F |

| | | | | | |
|---|---|---|---|---|---|
| 24096 | C | T | | S | nonsynonymous_A845V |
| 24097 | T | A | | S | synonymous_A845A |
| 24099 | C | T | | S | nonsynonymous_A846V |
| 24104 | G | C | | S | nonsynonymous_D848H |
| 24106 | C | T | | S | synonymous_D848D |
| 24119 | C | T | | S | stopgain_Q853X |
| 24121 | A | G | | S | synonymous_Q853Q |
| 24123 | A | C | | S | nonsynonymous_K854T |
| 24124 | G | T | | S | nonsynonymous_K854N |
| 24125 | T | C | | S | nonsynonymous_F855L |
| 24131 | G | T | | S | nonsynonymous_G857C |
| 24134 | C | T | | S | nonsynonymous_L858F |
| 24138 | C | T | | S | nonsynonymous_T859I |
| 24147 | C | T | | S | nonsynonymous_P862L |
| 24152 | T | C | | S | synonymous_L864L |
| 24153 | T | C | | S | nonsynonymous_L864S |
| 24155 | C | T | | S | nonsynonymous_L865F |
| 24157 | C | T | G | S | synonymous_L865L;<br>synonymous_L865L |
| 24169 | G | T | | S | nonsynonymous_M869I |
| 24173 | G | A | T | S | nonsynonymous_A871T;<br>nonsynonymous_A871S |
| 24180 | A | T | | S | nonsynonymous_Y873F |
| 24187 | T | C | | S | synonymous_S875S |
| 24189 | C | T | | S | nonsynonymous_A876V |
| 24190 | A | C | | S | synonymous_A876A |
| 24197 | G | T | | S | nonsynonymous_A879S |
| 24198 | C | A | T | S | nonsynonymous_A879E;<br>nonsynonymous_A879V |
| 24199 | G | A | | S | synonymous_A879A |
| 24200 | G | A | | S | nonsynonymous_G880S |

| | | | | | |
|---|---|---|---|---|---|
| 24210 | C | T | | S | nonsynonymous_T883I |
| 24213 | C | T | | S | nonsynonymous_S884F |
| 24214 | T | C | | S | synonymous_S884S |
| 24220 | G | T | | S | nonsynonymous_W886C |
| 24223 | C | T | | S | synonymous_T887T |
| 24227 | G | A | | S | nonsynonymous_G889S |
| 24230 | G | A | | S | nonsynonymous_A890T |
| 24233 | G | T | | S | nonsynonymous_G891C |
| 24236 | G | T | | S | nonsynonymous_A892S |
| 24237 | C | T | | S | nonsynonymous_A892V |
| 24238 | T | C | | S | synonymous_A892A |
| 24250 | A | C | | S | synonymous_I896I |
| 24253 | A | T | | S | synonymous_P897P |
| 24278 | T | A | | S | nonsynonymous_F906I |
| 24290 | G | T | | S | stopgain_G910X |
| 24297 | C | T | | S | nonsynonymous_T912I |
| 24299 | C | T | | S | stopgain_Q913X |
| 24308 | C | T | | S | nonsynonymous_L916F |
| 24321 | A | C | | S | nonsynonymous_Q920P |
| 24325 | A | G | | S | synonymous_K921K |
| 24329 | A | T | | S | nonsynonymous_I923F |
| 24337 | C | T | | S | synonymous_N925N |
| 24340 | A | G | | S | synonymous_Q926Q |
| 24348 | G | A | | S | nonsynonymous_S929N |
| 24349 | T | A | | S | nonsynonymous_S929R |
| 24351 | C | T | | S | nonsynonymous_A930V |
| 24353 | A | T | | S | nonsynonymous_I931F |
| 24357 | G | C | | S | nonsynonymous_G932A |
| 24358 | C | A | T | S | synonymous_G932G;<br>synonymous_G932G |
| 24362 | A | T | | S | nonsynonymous_I934F |

| | | | | | |
|---|---|---|---|---|---|
| 24368 | G | T | | S | nonsynonymous_D936Y |
| 24370 | C | T | | S | synonymous_D936D |
| 24372 | C | T | | S | nonsynonymous_S937L |
| 24374 | C | T | | S | nonsynonymous_L938F |
| 24378 | C | T | | S | nonsynonymous_S939F |
| 24380 | T | C | | S | nonsynonymous_S940P |
| 24381 | C | T | | S | nonsynonymous_S940F |
| 24382 | C | T | | S | synonymous_S940S |
| 24386 | G | T | | S | nonsynonymous_A942S |
| 24389 | A | T | | S | nonsynonymous_S943C |
| 24392 | G | T | | S | nonsynonymous_A944S |
| 24393 | C | T | | S | nonsynonymous_A944V |
| 24395 | C | T | | S | nonsynonymous_L945F |
| 24406 | T | G | | S | synonymous_L948L |
| 24410 | G | C | | S | nonsynonymous_D950H |
| 24436 | T | C | | S | synonymous_A958A |
| 24438 | T | A | | S | stopgain_L959X |
| 24442 | C | T | | S | synonymous_N960N |
| 24444 | C | T | | S | nonsynonymous_T961M |
| 24453 | A | G | | S | nonsynonymous_K964R |
| 24458 | C | T | | S | nonsynonymous_L966F |
| 24463 | C | T | | S | synonymous_S967S |
| 24465 | C | T | | S | nonsynonymous_S968F |
| 24487 | T | C | | S | synonymous_S975S |
| 24493 | A | T | | S | nonsynonymous_L977F |
| 24495 | A | T | | S | nonsynonymous_N978I |
| 24497 | G | T | | S | nonsynonymous_D979Y |
| 24498 | A | T | | S | nonsynonymous_D979V |
| 24502 | C | T | | S | synonymous_I980I |
| 24517 | C | T | | S | synonymous_D985D |
| 24524 | G | T | | S | stopgain_E988X |
| 24533 | G | T | | S | nonsynonymous_V991L |

| | | | | | |
|---|---|---|---|---|---|
| 24554 | A | G | | S | nonsynonymous_T998A |
| 24556 | A | G | | S | synonymous_T998T |
| 24557 | G | T | | S | nonsynonymous_G999C |
| 24558 | G | A | | S | nonsynonymous_G999D |
| 24566 | C | T | | S | stopgain_Q1002X |
| 24579 | C | T | | S | nonsynonymous_T1006I |
| 24584 | G | A | T | S | nonsynonymous_V1008M; nonsynonymous_V1008L |
| 24588 | C | T | | S | nonsynonymous_T1009I |
| 24590 | C | T | | S | stopgain_Q1010X |
| 24605 | G | C | | S | nonsynonymous_A1015P |
| 24606 | C | A | T | S | nonsynonymous_A1015D; nonsynonymous_A1015V |
| 24607 | T | C | | S | synonymous_A1015A |
| 24616 | C | T | | S | synonymous_I1018I |
| 24621 | C | T | | S | nonsynonymous_A1020V |
| 24624 | C | T | | S | nonsynonymous_S1021F |
| 24627 | C | T | | S | nonsynonymous_A1022V |
| 24651 | C | T | | S | nonsynonymous_S1030L |
| 24654 | A | G | | S | nonsynonymous_E1031G |
| 24662 | C | T | | S | nonsynonymous_L1034F |
| 24668 | C | T | | S | stopgain_Q1036X |
| 24672 | C | T | | S | nonsynonymous_S1037L |
| 24675 | A | T | | S | nonsynonymous_K1038I |
| 24694 | A | T | | S | synonymous_G1044G |
| 24695 | A | G | | S | nonsynonymous_K1045E |
| 24704 | C | T | | S | nonsynonymous_H1048Y |
| 24716 | T | C | | S | nonsynonymous_F1052L |
| 24718 | C | T | | S | synonymous_F1052F |
| 24729 | C | T | | S | nonsynonymous_A1056V |

| | | | | | |
|---|---|---|---|---|---|
| 24731 | C | T | G | S | nonsynonymous_P1057S; nonsynonymous_P1057A |
| 24732 | C | T | | S | nonsynonymous_P1057L |
| 24734 | C | T | | S | nonsynonymous_H1058Y |
| 24740 | G | T | | S | nonsynonymous_V1060L |
| 24751 | G | T | | S | nonsynonymous_L1063F |
| 24752 | C | T | | S | nonsynonymous_H1064Y |
| 24755 | G | A | | S | nonsynonymous_V1065M |
| 24757 | G | T | | S | synonymous_V1065V |
| 24766 | C | T | | S | synonymous_V1068V |
| 24767 | C | T | | S | nonsynonymous_P1069S |
| 24771 | C | T | | S | nonsynonymous_A1070V |
| 24773 | C | T | | S | stopgain_Q1071X |
| 24776 | G | A | | S | nonsynonymous_E1072K |
| 24777 | A | G | | S | nonsynonymous_E1072G |
| 24787 | C | T | | S | synonymous_F1075F |
| 24792 | C | T | | S | nonsynonymous_T1077I |
| 24797 | C | T | | S | nonsynonymous_P1079S |
| 24802 | C | T | | S | synonymous_A1080A |
| 24816 | G | A | | S | nonsynonymous_G1085E |
| 24821 | G | C | T | S | nonsynonymous_A1087P; nonsynonymous_A1087S |
| 24826 | C | T | | S | synonymous_H1088H |
| 24844 | C | T | | S | synonymous_V1094V |
| 24845 | T | A | | S | nonsynonymous_F1095I |
| 24847 | T | A | C | S | nonsynonymous_F1095L; synonymous_F1095F |
| 24848 | G | T | | S | nonsynonymous_V1096F |
| 24856 | T | C | G | S | synonymous_N1098N; nonsynonymous_N1098K |
| 24862 | A | G | | S | synonymous_T1100T |

| | | | | | |
|---|---|---|---|---|---|
| 24863 | C | T | | S | nonsynonymous_H1101Y |
| 24865 | C | T | | S | synonymous_H1101H |
| 24868 | G | T | | S | nonsynonymous_W1102C |
| 24871 | T | C | | S | synonymous_F1103F |
| 24872 | G | T | | S | nonsynonymous_V1104L |
| 24874 | A | T | | S | synonymous_V1104V |
| 24876 | C | T | | S | nonsynonymous_T1105I |
| 24878 | C | T | | S | stopgain_Q1106X |
| 24883 | G | A | | S | synonymous_R1107R |
| 24904 | C | T | | S | synonymous_I1114I |
| 24905 | A | T | | S | nonsynonymous_I1115F |
| 24911 | A | T | | S | nonsynonymous_T1117S |
| 24912 | C | T | | S | nonsynonymous_T1117I |
| 24914 | G | C | | S | nonsynonymous_D1118H |
| 24921 | C | T | | S | nonsynonymous_T1120I |
| 24922 | A | G | | S | synonymous_T1120T |
| 24924 | T | C | | S | nonsynonymous_F1121S |
| 24926 | G | T | | S | nonsynonymous_V1122L |
| 24930 | C | T | | S | nonsynonymous_S1123F |
| 24933 | G | C | T | S | nonsynonymous_G1124A; nonsynonymous_G1124V |
| 24934 | T | C | | S | synonymous_G1124G |
| 24940 | T | C | | S | synonymous_C1126C |
| 24947 | G | C | | S | nonsynonymous_V1129L |
| 24953 | G | T | | S | stopgain_G1131X |
| 24961 | C | T | | S | synonymous_V1133V |
| 24979 | T | C | | S | synonymous_D1139D |
| 24980 | C | T | | S | nonsynonymous_P1140S |
| 24981 | C | T | | S | nonsynonymous_P1140L |
| 24982 | T | C | | S | synonymous_P1140P |
| 24986 | C | T | | S | stopgain_Q1142X |

| | | | | | |
|---|---|---|---|---|---|
| 24989 | C | T | | S | nonsynonymous_P1143S |
| 25000 | C | T | | S | synonymous_D1146D |
| 25006 | C | T | | S | synonymous_F1148F |
| 25041 | C | T | | S | nonsynonymous_T1160I |
| 25046 | C | T | | S | nonsynonymous_P1162S |
| 25047 | C | T | | S | nonsynonymous_P1162L |
| 25048 | A | T | | S | synonymous_P1162P |
| 25049 | G | T | | S | nonsynonymous_D1163Y |
| 25050 | A | G | | S | nonsynonymous_D1163G |
| 25051 | T | C | | S | synonymous_D1163D |
| 25052 | G | T | | S | nonsynonymous_V1164F |
| 25064 | G | T | | S | nonsynonymous_D1168Y |
| 25080 | A | T | | S | nonsynonymous_N1173I |
| 25086 | C | T | | S | nonsynonymous_S1175L |
| 25090 | T | C | | S | synonymous_V1176V |
| 25096 | C | T | | S | synonymous_N1178N |
| 25100 | C | T | | S | stopgain_Q1180X |
| 25104 | A | G | | S | nonsynonymous_K1181R |
| 25106 | G | A | | S | nonsynonymous_E1182K |
| 25109 | A | C | | S | nonsynonymous_I1183L |
| 25111 | T | C | | S | synonymous_I1183I |
| 25114 | C | T | | S | synonymous_D1184D |
| 25116 | G | C | | S | nonsynonymous_R1185P |
| 25135 | G | C | T | S | nonsynonymous_K1191N; nonsynonymous_K1191N |
| 25137 | A | T | | S | nonsynonymous_N1192I |
| 25148 | T | C | | S | nonsynonymous_S1196P |
| 25156 | C | T | | S | synonymous_I1198I |
| 25159 | T | C | | S | synonymous_D1199D |
| 25169 | C | T | | S | nonsynonymous_L1203F |
| 25173 | G | A | | S | nonsynonymous_G1204E |

| | | | | | |
|---|---|---|---|---|---|
| 25181 | G | T | | S | stopgain_E1207X |
| 25183 | G | T | | S | nonsynonymous_E1207D |
| 25186 | G | T | | S | nonsynonymous_Q1208H |
| 25197 | G | T | | S | nonsynonymous_W1212L |
| 25198 | G | A | T | S | stopgain_W1212X; nonsynonymous_W1212C |
| 25199 | C | T | | S | nonsynonymous_P1213S |
| 25204 | G | T | | S | nonsynonymous_W1214C |
| 25207 | C | T | | S | synonymous_Y1215Y |
| 25208 | A | T | | S | nonsynonymous_I1216F |
| 25209 | T | C | | S | nonsynonymous_I1216T |
| 25210 | T | C | | S | synonymous_I1216I |
| 25216 | A | G | | S | synonymous_L1218L |
| 25217 | G | T | | S | nonsynonymous_G1219C |
| 25219 | T | C | | S | synonymous_G1219G |
| 25227 | C | T | | S | nonsynonymous_A1222V |
| 25248 | T | C | | S | nonsynonymous_M1229T |
| 25249 | G | T | | S | nonsynonymous_M1229I |
| 25250 | G | C | T | S | nonsynonymous_V1230L; nonsynonymous_V1230L |
| 25266 | G | A | T | S | nonsynonymous_C1235Y; nonsynonymous_C1235F |
| 25268 | T | G | | S | nonsynonymous_C1236G |
| 25273 | G | T | | S | nonsynonymous_M1237I |
| 25278 | G | T | | S | nonsynonymous_S1239I |
| 25282 | C | T | | S | synonymous_C1240C |
| 25290 | G | T | | S | nonsynonymous_C1243F |
| 25294 | C | T | | S | synonymous_L1244L |
| 25297 | G | T | | S | nonsynonymous_K1245N |
| 25316 | T | C | | S | nonsynonymous_S1252P |
| 25317 | C | T | | S | nonsynonymous_S1252F |

| | | | | | |
|---|---|---|---|---|---|
| 25319 | T | A | | S | nonsynonymous_C1253S |
| 25320 | G | A | | S | nonsynonymous_C1253Y |
| 25323 | G | T | | S | nonsynonymous_C1254F |
| 25334 | G | T | | S | stopgain_E1258X |
| 25336 | A | T | | S | nonsynonymous_E1258D |
| 25337 | G | C | | S | nonsynonymous_D1259H |
| 25339 | C | T | | S | synonymous_D1259D |
| 25340 | G | A | | S | nonsynonymous_D1260N |
| 25342 | C | T | | S | synonymous_D1260D |
| 25348 | G | T | | S | nonsynonymous_E1262D |
| 25350 | C | T | | S | nonsynonymous_P1263L |
| 25352 | G | T | | S | nonsynonymous_V1264L |
| 25354 | G | T | | S | synonymous_V1264V |
| 25357 | C | T | | S | synonymous_L1265L |
| 25359 | A | G | | S | nonsynonymous_K1266R |
| 25365 | T | C | G | S | nonsynonymous_V1268A; nonsynonymous_V1268G |
| 25373 | C | T | | S | nonsynonymous_H1271Y |
| 25376 | T | C | | S | nonsynonymous_Y1272H |
| 25393 | A | G | | ORF3a | nonsynonymous_M1V |
| 25407 | G | T | | ORF3a | nonsynonymous_M5I |
| 25413 | C | T | | ORF3a | synonymous_I7I |
| 25418 | C | G | | ORF3a | nonsynonymous_T9R |
| 25421 | T | C | | ORF3a | nonsynonymous_I10T |
| 25422 | T | G | | ORF3a | nonsynonymous_I10M |
| 25429 | G | T | | ORF3a | nonsynonymous_V13L |
| 25433 | C | T | | ORF3a | nonsynonymous_T14I |
| 25441 | C | T | G | ORF3a | stopgain_Q17X; nonsynonymous_Q17E |
| 25443 | A | G | | ORF3a | synonymous_Q17Q |
| 25449 | A | G | | ORF3a | synonymous_E19E |

| | | | | | |
|---|---|---|---|---|---|
| 25452 | C | T | | ORF3a | synonymous_I20I |
| 25455 | G | T | | ORF3a | nonsynonymous_K21N |
| 25460 | C | T | | ORF3a | nonsynonymous_A23V |
| 25462 | A | G | | ORF3a | nonsynonymous_T24A |
| 25463 | C | T | | ORF3a | nonsynonymous_T24I |
| 25464 | T | C | | ORF3a | synonymous_T24T |
| 25466 | C | T | | ORF3a | nonsynonymous_P25L |
| 25469 | C | T | | ORF3a | nonsynonymous_S26L |
| 25471 | G | C | | ORF3a | nonsynonymous_D27H |
| 25472 | A | T | G | ORF3a | nonsynonymous_D27V; nonsynonymous_D27G |
| 25473 | T | C | | ORF3a | synonymous_D27D |
| 25480 | C | T | | ORF3a | nonsynonymous_R30C |
| 25483 | G | A | | ORF3a | nonsynonymous_A31T |
| 25486 | A | C | | ORF3a | nonsynonymous_T32P |
| 25487 | C | T | | ORF3a | nonsynonymous_T32I |
| 25494 | G | T | | ORF3a | synonymous_T34T |
| 25496 | T | C | | ORF3a | nonsynonymous_I35T |
| 25504 | C | T | | ORF3a | stopgain_Q38X |
| 25511 | C | T | | ORF3a | nonsynonymous_S40L |
| 25513 | C | T | | ORF3a | nonsynonymous_L41F |
| 25514 | T | C | | ORF3a | nonsynonymous_L41P |
| 25515 | C | T | | ORF3a | synonymous_L41L |
| 25516 | C | T | | ORF3a | nonsynonymous_P42S |
| 25517 | C | T | | ORF3a | nonsynonymous_P42L |
| 25520 | T | C | | ORF3a | nonsynonymous_F43S |
| 25521 | C | T | | ORF3a | synonymous_F43F |
| 25522 | G | A | | ORF3a | nonsynonymous_G44R |
| 25528 | C | T | | ORF3a | nonsynonymous_L46F |
| 25538 | G | A | | ORF3a | nonsynonymous_G49D |
| 25546 | C | A | | ORF3a | nonsynonymous_L52I |

| | | | | | |
|---|---|---|---|---|---|
| 25549 | C | T | | ORF3a | nonsynonymous_L53F |
| 25550 | T | C | | ORF3a | nonsynonymous_L53P |
| 25552 | G | T | | ORF3a | nonsynonymous_A54S |
| 25553 | C | T | | ORF3a | nonsynonymous_A54V |
| 25561 | C | T | | ORF3a | stopgain_Q57X |
| 25563 | G | C | T | ORF3a | nonsynonymous_Q57H;<br>nonsynonymous_Q57H |
| 25568 | C | T | | ORF3a | nonsynonymous_A59V |
| 25571 | C | T | | ORF3a | nonsynonymous_S60F |
| 25572 | C | T | | ORF3a | synonymous_S60S |
| 25578 | C | A | | ORF3a | synonymous_I62I |
| 25585 | C | T | | ORF3a | nonsynonymous_L65F |
| 25587 | C | T | | ORF3a | synonymous_L65L |
| 25593 | G | C | | ORF3a | nonsynonymous_K67N |
| 25598 | G | T | | ORF3a | nonsynonymous_W69L |
| 25603 | C | T | | ORF3a | synonymous_L71L |
| 25606 | G | T | | ORF3a | nonsynonymous_A72S |
| 25609 | C | T | | ORF3a | nonsynonymous_L73F |
| 25612 | T | C | | ORF3a | nonsynonymous_S74P |
| 25613 | C | T | | ORF3a | nonsynonymous_S74F |
| 25614 | C | T | | ORF3a | synonymous_S74S |
| 25615 | A | G | | ORF3a | nonsynonymous_K75E |
| 25616 | A | G | | ORF3a | nonsynonymous_K75R |
| 25621 | G | T | | ORF3a | nonsynonymous_V77F |
| 25624 | C | T | | ORF3a | nonsynonymous_H78Y |
| 25626 | C | T | | ORF3a | synonymous_H78H |
| 25627 | T | C | | ORF3a | nonsynonymous_F79L |
| 25638 | C | T | | ORF3a | synonymous_N82N |
| 25641 | G | T | | ORF3a | nonsynonymous_L83F |
| 25642 | C | T | | ORF3a | synonymous_L84L |

| | | | | | |
|---|---|---|---|---|---|
| 25644 | G | A | T | ORF3a | synonymous_L84L;<br>synonymous_L84L |
| 25654 | G | T | | ORF3a | nonsynonymous_V88L |
| 25658 | C | T | | ORF3a | nonsynonymous_T89I |
| 25663 | T | C | | ORF3a | nonsynonymous_Y91H |
| 25667 | C | T | | ORF3a | nonsynonymous_S92L |
| 25669 | C | T | | ORF3a | nonsynonymous_H93Y |
| 25676 | T | C | | ORF3a | nonsynonymous_L95S |
| 25688 | C | T | | ORF3a | nonsynonymous_A99V |
| 25689 | T | C | | ORF3a | synonymous_A99A |
| 25690 | G | T | | ORF3a | nonsynonymous_G100C |
| 25691 | G | T | | ORF3a | nonsynonymous_G100V |
| 25703 | C | T | | ORF3a | nonsynonymous_P104L |
| 25704 | T | C | | ORF3a | synonymous_P104P |
| 25710 | C | T | | ORF3a | synonymous_L106L |
| 25714 | C | T | | ORF3a | nonsynonymous_L108F |
| 25720 | G | C | T | ORF3a | nonsynonymous_A110P;<br>nonsynonymous_A110S |
| 25721 | C | T | | ORF3a | nonsynonymous_A110V |
| 25726 | G | T | | ORF3a | nonsynonymous_V112F |
| 25728 | C | T | | ORF3a | synonymous_V112V |
| 25731 | C | T | | ORF3a | synonymous_Y113Y |
| 25733 | T | C | | ORF3a | nonsynonymous_F114S |
| 25740 | G | T | | ORF3a | nonsynonymous_Q116H |
| 25750 | T | C | | ORF3a | nonsynonymous_F120L |
| 25752 | T | G | | ORF3a | nonsynonymous_F120L |
| 25753 | G | T | | ORF3a | nonsynonymous_V121L |
| 25758 | A | C | | ORF3a | nonsynonymous_R122S |
| 25763 | T | C | | ORF3a | nonsynonymous_I124T |
| 25767 | G | C | T | ORF3a | nonsynonymous_M125I;<br>nonsynonymous_M125I |

| | | | | | |
|---|---|---|---|---|---|
| 25771 | C | T | | ORF3a | nonsynonymous_L127F |
| 25777 | C | T | | ORF3a | nonsynonymous_L129F |
| 25782 | C | T | | ORF3a | synonymous_C130C |
| 25785 | G | T | | ORF3a | nonsynonymous_W131C |
| 25790 | G | T | | ORF3a | nonsynonymous_C133F |
| 25791 | C | T | | ORF3a | synonymous_C133C |
| 25796 | C | T | | ORF3a | nonsynonymous_S135F |
| 25797 | C | T | | ORF3a | synonymous_S135S |
| 25810 | C | T | | ORF3a | nonsynonymous_L140F |
| 25814 | A | T | | ORF3a | nonsynonymous_Y141F |
| 25819 | G | T | | ORF3a | nonsynonymous_A143S |
| 25831 | C | T | | ORF3a | nonsynonymous_L147F |
| 25835 | G | T | | ORF3a | nonsynonymous_C148F |
| 25836 | C | T | | ORF3a | synonymous_C148C |
| 25839 | G | A | | ORF3a | stopgain_W149X |
| 25844 | C | T | | ORF3a | nonsynonymous_T151I |
| 25855 | G | T | | ORF3a | nonsynonymous_D155Y |
| 25857 | C | T | | ORF3a | synonymous_D155D |
| 25865 | T | C | | ORF3a | nonsynonymous_I158T |
| 25867 | C | T | | ORF3a | nonsynonymous_P159S |
| 25868 | C | T | | ORF3a | nonsynonymous_P159L |
| 25879 | G | T | | ORF3a | nonsynonymous_V163L |
| 25886 | C | T | | ORF3a | nonsynonymous_S165F |
| 25904 | C | T | | ORF3a | nonsynonymous_S171L |
| 25912 | G | T | | ORF3a | nonsynonymous_G174C |
| 25913 | G | T | | ORF3a | nonsynonymous_G174V |
| 25916 | C | T | | ORF3a | nonsynonymous_T175I |
| 25931 | C | T | | ORF3a | nonsynonymous_S180F |
| 25933 | G | C | T | ORF3a | nonsynonymous_E181Q;<br>stopgain_E181X |
| 25936 | C | T | | ORF3a | nonsynonymous_H182Y |

| | | | | | |
|---|---|---|---|---|---|
| 25947 | G | T | | ORF3a | nonsynonymous_Q185H |
| 25956 | T | C | | ORF3a | synonymous_G188G |
| 25957 | T | C | | ORF3a | nonsynonymous_Y189H |
| 25968 | A | G | | ORF3a | synonymous_K192K |
| 25979 | G | T | | ORF3a | nonsynonymous_G196V |
| 25987 | G | A | T | ORF3a | nonsynonymous_D199N; nonsynonymous_D199Y |
| 26002 | C | T | | ORF3a | nonsynonymous_H204Y |
| 26020 | G | A | | ORF3a | nonsynonymous_D210N |
| 26028 | C | T | | ORF3a | synonymous_Y212Y |
| 26029 | C | A | T | ORF3a | nonsynonymous_Q213K; stopgain_Q213X |
| 26037 | C | T | | ORF3a | synonymous_Y215Y |
| 26042 | C | T | | ORF3a | nonsynonymous_T217I |
| 26044 | C | G | | ORF3a | nonsynonymous_Q218E |
| 26046 | A | G | | ORF3a | synonymous_Q218Q |
| 26049 | G | T | | ORF3a | nonsynonymous_L219F |
| 26062 | G | C | | ORF3a | nonsynonymous_G224R |
| 26066 | T | C | | ORF3a | nonsynonymous_V225A |
| 26078 | C | T | | ORF3a | nonsynonymous_T229I |
| 26079 | C | T | | ORF3a | synonymous_T229T |
| 26082 | C | T | | ORF3a | synonymous_F230F |
| 26088 | C | T | | ORF3a | synonymous_I232I |
| 26094 | T | C | | ORF3a | synonymous_N234N |
| 26099 | T | A | | ORF3a | nonsynonymous_I236N |
| 26109 | G | C | | ORF3a | nonsynonymous_E239D |
| 26110 | C | T | | ORF3a | nonsynonymous_P240S |
| 26120 | A | T | | ORF3a | nonsynonymous_H243L |
| 26121 | T | C | G | ORF3a | synonymous_H243H; nonsynonymous_H243Q |
| 26124 | C | T | | ORF3a | synonymous_V244V |

| | | | | | |
|---|---|---|---|---|---|
| 26125 | C | T | | ORF3a | stopgain_Q245X |
| 26130 | T | C | | ORF3a | synonymous_I246I |
| 26131 | C | G | | ORF3a | nonsynonymous_H247D |
| 26143 | G | T | | ORF3a | nonsynonymous_G251C |
| 26144 | G | T | | ORF3a | nonsynonymous_G251V |
| 26147 | C | T | | ORF3a | nonsynonymous_S252L |
| 26151 | C | T | | ORF3a | synonymous_S253S |
| 26158 | G | T | | ORF3a | nonsynonymous_V256F |
| 26164 | C | T | | ORF3a | nonsynonymous_P258S |
| 26167 | G | T | | ORF3a | nonsynonymous_V259L |
| 26172 | G | T | | ORF3a | nonsynonymous_M260I |
| 26176 | C | T | | ORF3a | nonsynonymous_P262S |
| 26177 | C | T | | ORF3a | nonsynonymous_P262L |
| 26185 | G | T | | ORF3a | nonsynonymous_D265Y |
| 26191 | C | G | | ORF3a | nonsynonymous_P267A |
| 26194 | A | G | | ORF3a | nonsynonymous_T268A |
| 26195 | C | T | | ORF3a | nonsynonymous_T268M |
| 26196 | G | T | | ORF3a | synonymous_T268T |
| 26198 | C | T | | ORF3a | nonsynonymous_T269M |
| 26201 | C | T | | ORF3a | nonsynonymous_T270I |
| 26212 | C | T | | ORF3a | nonsynonymous_P274S |
| 26213 | C | T | | ORF3a | nonsynonymous_P274L |
| 26217 | G | T | | ORF3a | nonsynonymous_L275F |
| 26227 | G | A | | intergenic | |
| 26228 | C | T | | intergenic | |
| 26230 | G | T | | intergenic | |
| 26233 | G | T | | intergenic | |
| 26256 | C | T | | E | synonymous_F4F |
| 26262 | G | T | | E | synonymous_S6S |
| 26263 | G | T | | E | stopgain_E7X |
| 26268 | G | T | | E | nonsynonymous_E8D |
| 26269 | A | G | | E | nonsynonymous_T9A |

| | | | | | |
|---|---|---|---|---|---|
| 26270 | C | T | | E | nonsynonymous_T9I |
| 26273 | G | T | | E | nonsynonymous_G10V |
| 26305 | C | T | | E | nonsynonymous_L21F |
| 26309 | C | T | | E | nonsynonymous_A22V |
| 26313 | C | T | | E | synonymous_F23F |
| 26316 | G | T | | E | synonymous_V24V |
| 26326 | C | T | | E | synonymous_L28L |
| 26333 | C | T | | E | nonsynonymous_T30I |
| 26335 | C | T | | E | synonymous_L31L |
| 26336 | T | C | | E | nonsynonymous_L31P |
| 26338 | G | T | | E | nonsynonymous_A32S |
| 26339 | C | T | | E | nonsynonymous_A32V |
| 26340 | C | T | | E | synonymous_A32A |
| 26348 | C | T | | E | nonsynonymous_T35I |
| 26351 | C | T | | E | nonsynonymous_A36V |
| 26367 | G | C | | E | synonymous_A41A |
| 26370 | C | T | | E | synonymous_Y42Y |
| 26373 | C | G | | E | nonsynonymous_C43W |
| 26375 | G | A | | E | nonsynonymous_C44Y |
| 26380 | A | T | | E | nonsynonymous_I46F |
| 26387 | A | T | | E | nonsynonymous_N48I |
| 26388 | C | T | | E | synonymous_N48N |
| 26389 | G | T | | E | nonsynonymous_V49L |
| 26391 | G | T | | E | synonymous_V49V |
| 26394 | T | C | | E | synonymous_S50S |
| 26398 | G | A | | E | nonsynonymous_V52I |
| 26408 | C | T | | E | nonsynonymous_S55F |
| 26418 | T | A | | E | synonymous_V58V |
| 26423 | C | T | | E | nonsynonymous_S60F |
| 26425 | C | T | | E | nonsynonymous_R61C |
| 26426 | G | T | | E | nonsynonymous_R61L |

| | | | | | |
|---|---|---|---|---|---|
| 26428 | G | A | T | E | nonsynonymous_V62I; nonsynonymous_V62F |
| 26431 | A | T | | E | stopgain_K63X |
| 26437 | C | T | | E | synonymous_L65L |
| 26439 | G | A | T | E | synonymous_L65L; synonymous_L65L |
| 26447 | C | A | T | E | nonsynonymous_S68Y; nonsynonymous_S68F |
| 26449 | A | T | | E | stopgain_R69X |
| 26455 | C | T | | E | nonsynonymous_P71S |
| 26456 | C | T | | E | nonsynonymous_P71L |
| 26457 | T | C | | E | synonymous_P71P |
| 26458 | G | C | | E | nonsynonymous_D72H |
| 26461 | C | T | | E | nonsynonymous_L73F |
| 26464 | C | T | | E | synonymous_L74L |
| 26466 | G | A | | E | synonymous_L74L |
| 26468 | T | C | | E | nonsynonymous_V75A |
| 26471 | A | T | | E | stoploss_X76L |
| 26474 | C | T | | intergenic | |
| 26483 | T | A | | intergenic | |
| 26490 | T | G | | intergenic | |
| 26497 | T | C | | intergenic | |
| 26499 | C | T | | intergenic | |
| 26518 | T | G | | intergenic | |
| 26519 | A | T | | intergenic | |
| 26520 | G | T | | intergenic | |
| 26521 | C | T | | intergenic | |
| 26522 | C | T | | intergenic | |
| 26526 | G | T | | M | nonsynonymous_A2S |
| 26529 | G | C | | M | nonsynonymous_D3H |

| | | | | | |
|---|---|---|---|---|---|
| 26530 | A | C | G | M | nonsynonymous_D3A; nonsynonymous_D3G |
| 26533 | C | T | | M | nonsynonymous_S4F |
| 26534 | C | T | | M | synonymous_S4S |
| 26542 | C | A | T | M | nonsynonymous_T7N; nonsynonymous_T7I |
| 26545 | T | G | | M | nonsynonymous_I8S |
| 26546 | T | C | | M | synonymous_I8I |
| 26548 | C | T | | M | nonsynonymous_T9I |
| 26558 | G | T | | M | nonsynonymous_E12D |
| 26561 | T | A | | M | synonymous_L13L |
| 26567 | G | T | | M | nonsynonymous_K15N |
| 26577 | C | T | | M | stopgain_Q19X |
| 26589 | G | T | | M | nonsynonymous_V23L |
| 26592 | A | C | | M | nonsynonymous_I24L |
| 26596 | G | T | | M | nonsynonymous_G25V |
| 26601 | C | T | | M | synonymous_L27L |
| 26606 | C | T | | M | synonymous_F28F |
| 26609 | T | C | | M | synonymous_L29L |
| 26615 | G | T | | M | nonsynonymous_W31C |
| 26618 | T | C | | M | synonymous_I32I |
| 26620 | G | T | | M | nonsynonymous_C33F |
| 26625 | C | T | | M | synonymous_L35L |
| 26636 | C | T | | M | synonymous_A38A |
| 26640 | G | T | | M | nonsynonymous_A40S |
| 26644 | A | T | | M | nonsynonymous_N41I |
| 26654 | G | T | | M | nonsynonymous_R44S |
| 26660 | G | T | | M | nonsynonymous_L46F |
| 26672 | G | T | | M | nonsynonymous_K50N |
| 26681 | C | T | | M | synonymous_F53F |
| 26702 | A | T | | M | synonymous_V60V |

| | | | | | |
|---|---|---|---|---|---|
| 26707 | T | C | | M | nonsynonymous_L62S |
| 26713 | G | T | | M | nonsynonymous_C64F |
| 26717 | T | C | | M | synonymous_F65F |
| 26720 | G | C | T | M | synonymous_V66V;<br>synonymous_V66V |
| 26724 | G | T | | M | nonsynonymous_A68S |
| 26727 | G | T | | M | nonsynonymous_A69S |
| 26728 | C | T | G | M | nonsynonymous_A69V;<br>nonsynonymous_A69G |
| 26729 | T | C | | M | synonymous_A69A |
| 26730 | G | A | T | M | nonsynonymous_V70I;<br>nonsynonymous_V70F |
| 26735 | C | T | | M | synonymous_Y71Y |
| 26750 | C | T | | M | synonymous_I76I |
| 26763 | G | T | | M | nonsynonymous_A81S |
| 26768 | C | T | | M | synonymous_I82I |
| 26781 | C | T | | M | nonsynonymous_L87F |
| 26792 | G | A | | M | synonymous_L90L |
| 26793 | A | G | | M | nonsynonymous_M91V |
| 26794 | T | C | | M | nonsynonymous_M91T |
| 26801 | C | T | | M | synonymous_L93L |
| 26804 | C | T | | M | synonymous_S94S |
| 26807 | C | T | | M | synonymous_Y95Y |
| 26809 | T | C | | M | nonsynonymous_F96S |
| 26812 | T | A | | M | nonsynonymous_I97N |
| 26818 | C | G | | M | nonsynonymous_S99C |
| 26819 | T | C | | M | synonymous_S99S |
| 26822 | C | T | | M | synonymous_F100F |
| 26826 | C | T | | M | synonymous_L102L |
| 26835 | C | T | | M | nonsynonymous_R105C |
| 26842 | G | T | | M | nonsynonymous_R107L |

| | | | | | |
|---|---|---|---|---|---|
| 26846 | C | T | | M | synonymous_S108S |
| 26851 | G | T | | M | nonsynonymous_W110L |
| 26854 | C | T | | M | nonsynonymous_S111L |
| 26858 | C | T | | M | synonymous_F112F |
| 26864 | A | T | G | M | synonymous_P114P;<br>synonymous_P114P |
| 26869 | C | T | | M | nonsynonymous_T116I |
| 26877 | C | T | | M | nonsynonymous_L119F |
| 26880 | C | T | | M | nonsynonymous_L120F |
| 26882 | C | T | | M | synonymous_L120L |
| 26885 | C | T | | M | synonymous_N121N |
| 26889 | C | T | | M | nonsynonymous_P123S |
| 26895 | C | T | | M | nonsynonymous_H125Y |
| 26900 | C | T | | M | synonymous_G126G |
| 26906 | T | C | | M | synonymous_I128I |
| 26910 | A | C | | M | nonsynonymous_T130P |
| 26911 | C | T | | M | nonsynonymous_T130I |
| 26912 | C | T | | M | synonymous_T130T |
| 26916 | C | T | | M | nonsynonymous_P132S |
| 26934 | C | A | | M | nonsynonymous_L138I |
| 26951 | G | A | | M | synonymous_V143V |
| 26959 | G | A | | M | nonsynonymous_R146H |
| 26964 | C | T | | M | nonsynonymous_H148Y |
| 26969 | T | C | | M | synonymous_L149L |
| 26970 | C | T | | M | nonsynonymous_R150C |
| 26971 | G | T | | M | nonsynonymous_R150L |
| 26977 | C | T | | M | nonsynonymous_A152V |
| 26988 | C | T | | M | synonymous_L156L |
| 26990 | A | G | | M | synonymous_L156L |
| 26996 | C | T | | M | synonymous_R158R |
| 27002 | C | T | | M | synonymous_D160D |

| | | | | | |
|---|---|---|---|---|---|
| 27005 | C | T | | M | synonymous_I161I |
| 27009 | G | T | | M | nonsynonymous_D163Y |
| 27011 | C | T | | M | synonymous_D163D |
| 27012 | C | A | | M | nonsynonymous_L164M |
| 27016 | C | T | | M | nonsynonymous_P165L |
| 27017 | T | A | | M | synonymous_P165P |
| 27021 | G | T | | M | stopgain_E167X |
| 27022 | A | T | | M | nonsynonymous_E167V |
| 27026 | C | T | | M | synonymous_I168I |
| 27033 | G | T | | M | nonsynonymous_A171S |
| 27034 | C | A | T | M | nonsynonymous_A171D;<br>nonsynonymous_A171V |
| 27037 | C | T | | M | nonsynonymous_T172I |
| 27040 | C | T | | M | nonsynonymous_S173L |
| 27046 | C | T | | M | nonsynonymous_T175M |
| 27052 | C | T | | M | nonsynonymous_S177F |
| 27059 | C | A | | M | stopgain_Y179X |
| 27070 | C | T | | M | nonsynonymous_A183V |
| 27075 | C | T | | M | stopgain_Q185X |
| 27077 | G | T | | M | nonsynonymous_Q185H |
| 27078 | C | T | | M | nonsynonymous_R186C |
| 27084 | G | T | | M | nonsynonymous_A188S |
| 27094 | C | T | | M | nonsynonymous_S191L |
| 27097 | G | T | | M | nonsynonymous_G192V |
| 27102 | G | T | | M | nonsynonymous_A194S |
| 27103 | C | T | | M | nonsynonymous_A194V |
| 27106 | C | T | | M | nonsynonymous_A195V |
| 27109 | A | T | | M | nonsynonymous_Y196F |
| 27112 | G | C | T | M | nonsynonymous_S197T;<br>nonsynonymous_S197I |
| 27113 | T | C | | M | synonymous_S197S |

| | | | | | |
|---|---|---|---|---|---|
| 27114 | C | T | | M | nonsynonymous_R198C |
| 27128 | C | T | | M | synonymous_G202G |
| 27129 | A | G | | M | nonsynonymous_N203D |
| 27145 | C | T | | M | nonsynonymous_T208I |
| 27163 | G | T | | M | nonsynonymous_S214I |
| 27165 | G | T | | M | nonsynonymous_D215Y |
| 27167 | C | T | | M | synonymous_D215D |
| 27196 | C | T | | intergenic | |
| 27208 | C | T | | ORF6 | nonsynonymous_H3Y |
| 27211 | C | T | | ORF6 | nonsynonymous_L4F |
| 27213 | C | T | | ORF6 | synonymous_L4L |
| 27214 | G | T | | ORF6 | nonsynonymous_V5F |
| 27217 | G | T | | ORF6 | nonsynonymous_D6Y |
| 27219 | C | T | | ORF6 | synonymous_D6D |
| 27223 | C | T | | ORF6 | stopgain_Q8X |
| 27226 | G | T | | ORF6 | nonsynonymous_V9F |
| 27230 | C | T | | ORF6 | nonsynonymous_T10I |
| 27233 | T | C | | ORF6 | nonsynonymous_I11T |
| 27247 | C | T | | ORF6 | synonymous_L16L |
| 27258 | G | T | | ORF6 | nonsynonymous_M19I |
| 27264 | T | C | | ORF6 | synonymous_T21T |
| 27267 | T | A | | ORF6 | nonsynonymous_F22L |
| 27281 | G | T | | ORF6 | nonsynonymous_W27L |
| 27282 | G | A | | ORF6 | stopgain_W27X |
| 27285 | T | C | | ORF6 | synonymous_N28N |
| 27289 | G | T | | ORF6 | nonsynonymous_D30Y |
| 27294 | C | T | | ORF6 | synonymous_Y31Y |
| 27299 | T | C | | ORF6 | nonsynonymous_I33T |
| 27304 | C | T | | ORF6 | nonsynonymous_L35F |
| 27323 | C | T | | ORF6 | nonsynonymous_S41F |
| 27329 | C | T | | ORF6 | nonsynonymous_S43L |
| 27347 | A | G | | ORF6 | nonsynonymous_Y49C |

| | | | | | |
|---|---|---|---|---|---|
| 27350 | C | T | | ORF6 | nonsynonymous_S50F |
| 27351 | T | G | | ORF6 | synonymous_S50S |
| 27355 | T | C | | ORF6 | synonymous_L52L |
| 27358 | G | T | | ORF6 | nonsynonymous_D53Y |
| 27366 | G | A | T | ORF6 | synonymous_E55E; nonsynonymous_E55D |
| 27370 | C | T | | ORF6 | nonsynonymous_P57S |
| 27371 | C | T | | ORF6 | nonsynonymous_P57L |
| 27373 | A | T | G | ORF6 | nonsynonymous_M58L; nonsynonymous_M58V |
| 27382 | G | T | | ORF6 | nonsynonymous_D61Y |
| 27383 | A | C | | ORF6 | nonsynonymous_D61A |
| 27384 | T | C | | ORF6 | synonymous_D61D |
| 27393 | C | T | | intergenic | |
| 27403 | A | T | | ORF7a | nonsynonymous_I4F |
| 27405 | T | C | | ORF7a | synonymous_I4I |
| 27406 | C | T | | ORF7a | nonsynonymous_L5F |
| 27407 | T | C | | ORF7a | nonsynonymous_L5P |
| 27408 | T | A | | ORF7a | synonymous_L5L |
| 27424 | A | G | | ORF7a | nonsynonymous_T11A |
| 27425 | C | T | | ORF7a | nonsynonymous_T11I |
| 27426 | A | T | | ORF7a | synonymous_T11T |
| 27427 | C | T | | ORF7a | nonsynonymous_L12F |
| 27429 | C | T | | ORF7a | synonymous_L12L |
| 27433 | A | T | | ORF7a | nonsynonymous_T14S |
| 27434 | C | T | | ORF7a | nonsynonymous_T14I |
| 27438 | T | C | | ORF7a | synonymous_C15C |
| 27439 | G | T | | ORF7a | stopgain_E16X |
| 27450 | C | T | | ORF7a | synonymous_H19H |
| 27459 | G | T | | ORF7a | nonsynonymous_E22D |
| 27460 | T | C | | ORF7a | nonsynonymous_C23R |

| | | | | | |
|---|---|---|---|---|---|
| 27469 | G | T | | ORF7a | nonsynonymous_G26C |
| 27476 | C | T | | ORF7a | nonsynonymous_T28I |
| 27478 | G | C | T | ORF7a | nonsynonymous_V29L; nonsynonymous_V29L |
| 27482 | T | C | | ORF7a | nonsynonymous_L30P |
| 27484 | T | C | | ORF7a | synonymous_L31L |
| 27490 | G | T | | ORF7a | stopgain_E33X |
| 27493 | C | T | | ORF7a | nonsynonymous_P34S |
| 27494 | C | T | | ORF7a | nonsynonymous_P34L |
| 27498 | C | T | | ORF7a | synonymous_C35C |
| 27500 | C | T | | ORF7a | nonsynonymous_S36F |
| 27506 | G | T | | ORF7a | nonsynonymous_G38V |
| 27509 | C | T | G | ORF7a | nonsynonymous_T39I; nonsynonymous_T39R |
| 27513 | C | T | | ORF7a | synonymous_Y40Y |
| 27514 | G | C | | ORF7a | nonsynonymous_E41Q |
| 27516 | G | T | | ORF7a | nonsynonymous_E41D |
| 27519 | C | A | | ORF7a | synonymous_G42G |
| 27524 | C | T | | ORF7a | nonsynonymous_S44L |
| 27526 | C | T | | ORF7a | nonsynonymous_P45S |
| 27527 | C | A | T | ORF7a | nonsynonymous_P45Q; nonsynonymous_P45L |
| 27532 | C | T | | ORF7a | nonsynonymous_H47Y |
| 27535 | C | T | | ORF7a | nonsynonymous_P48S |
| 27546 | T | C | | ORF7a | synonymous_D51D |
| 27549 | C | T | | ORF7a | synonymous_N52N |
| 27555 | T | C | | ORF7a | synonymous_F54F |
| 27556 | G | T | | ORF7a | nonsynonymous_A55S |
| 27559 | C | T | | ORF7a | synonymous_L56L |
| 27567 | C | T | | ORF7a | synonymous_C58C |
| 27573 | C | T | | ORF7a | synonymous_S60S |

| | | | | | |
|---|---|---|---|---|---|
| 27575 | C | T | | ORF7a | nonsynonymous_T61I |
| 27577 | C | T | | ORF7a | stopgain_Q62X |
| 27583 | G | C | | ORF7a | nonsynonymous_A64P |
| 27584 | C | T | | ORF7a | nonsynonymous_A64V |
| 27590 | C | T | | ORF7a | nonsynonymous_A66V |
| 27595 | C | T | | ORF7a | nonsynonymous_P68S |
| 27600 | C | T | | ORF7a | synonymous_D69D |
| 27603 | C | T | | ORF7a | synonymous_G70G |
| 27610 | C | T | | ORF7a | nonsynonymous_H73Y |
| 27612 | C | T | G | ORF7a | synonymous_H73H; nonsynonymous_H73Q |
| 27613 | G | C | T | ORF7a | nonsynonymous_V74L; nonsynonymous_V74F |
| 27618 | T | C | | ORF7a | synonymous_Y75Y |
| 27619 | C | T | | ORF7a | stopgain_Q76X |
| 27624 | A | T | | ORF7a | nonsynonymous_L77F |
| 27628 | G | T | | ORF7a | nonsynonymous_A79S |
| 27629 | C | T | | ORF7a | nonsynonymous_A79V |
| 27630 | C | T | | ORF7a | synonymous_A79A |
| 27632 | G | A | T | ORF7a | nonsynonymous_R80K; nonsynonymous_R80I |
| 27633 | A | C | | ORF7a | nonsynonymous_R80S |
| 27635 | C | T | | ORF7a | nonsynonymous_S81L |
| 27640 | T | C | | ORF7a | nonsynonymous_S83P |
| 27641 | C | T | | ORF7a | nonsynonymous_S83L |
| 27643 | C | T | | ORF7a | nonsynonymous_P84S |
| 27644 | C | T | | ORF7a | nonsynonymous_P84L |
| 27645 | T | A | | ORF7a | synonymous_P84P |
| 27657 | C | T | | ORF7a | synonymous_I88I |
| 27659 | G | A | T | ORF7a | nonsynonymous_R89K; nonsynonymous_R89I |

| | | | | | |
|---|---|---|---|---|---|
| 27661 | C | T | | ORF7a | stopgain_Q90X |
| 27669 | A | G | | ORF7a | synonymous_E92E |
| 27670 | G | T | | ORF7a | nonsynonymous_V93F |
| 27673 | C | A | T | ORF7a | nonsynonymous_Q94K; stopgain_Q94X |
| 27675 | A | C | | ORF7a | nonsynonymous_Q94H |
| 27679 | C | T | | ORF7a | nonsynonymous_L96F |
| 27687 | T | C | | ORF7a | synonymous_S98S |
| 27688 | C | T | | ORF7a | nonsynonymous_P99S |
| 27689 | C | T | | ORF7a | nonsynonymous_P99L |
| 27690 | A | T | | ORF7a | synonymous_P99P |
| 27691 | A | T | | ORF7a | nonsynonymous_I100F |
| 27703 | G | A | T | ORF7a | nonsynonymous_V104I; nonsynonymous_V104F |
| 27707 | C | T | | ORF7a | nonsynonymous_A105V |
| 27727 | C | T | | ORF7a | nonsynonymous_L112F |
| 27729 | T | G | | ORF7a | synonymous_L112L |
| 27737 | C | T | | ORF7a | nonsynonymous_T115I |
| 27739 | C | T | | ORF7a | nonsynonymous_L116F |
| 27750 | G | T | | ORF7a | nonsynonymous_K119N |
| 27759 | A | G | | ORF7a | stoploss_X122W |
| 27760 | T | C | | ORF7b | nonsynonymous_I2T |
| 27761 | T | C | | ORF7b | synonymous_I2I |
| 27769 | C | T | | ORF7b | nonsynonymous_S5L |
| 27770 | A | G | | ORF7b | synonymous_S5S |
| 27772 | T | C | | ORF7b | nonsynonymous_L6S |
| 27778 | A | C | | ORF7b | nonsynonymous_D8A |
| 27779 | C | G | | ORF7b | nonsynonymous_D8E |
| 27812 | C | T | | ORF7b | synonymous_F19F |
| 27816 | G | T | | ORF7b | nonsynonymous_V21F |
| 27828 | C | T | | ORF7b | nonsynonymous_L25F |

| | | | | | |
|---|---|---|---|---|---|
| 27847 | C | T | | ORF7b | nonsynonymous_S31L |
| 27849 | C | T | | ORF7b | nonsynonymous_L32F |
| 27865 | A | T | | ORF7b | nonsynonymous_H37L |
| 27872 | A | G | | ORF7b | synonymous_E39E |
| 27874 | C | T | | ORF7b | nonsynonymous_T40I |
| 27877 | G | C | T | ORF7b | nonsynonymous_C41S; nonsynonymous_C41F |
| 27879 | C | T | | ORF7b | nonsynonymous_H42Y |
| 27883 | C | T | | ORF7b | nonsynonymous_A43V |
| 27884 | C | T | | ORF7b | synonymous_A43A |
| 27887 | A | G | | ORF7b | synonymous_X44X |
| 27889 | C | T | G | intergenic | |
| 27896 | G | A | T | ORF8 | nonsynonymous_M1I; nonsynonymous_M1I |
| 27900 | T | A | | ORF8 | nonsynonymous_F3I |
| 27903 | C | T | | ORF8 | nonsynonymous_L4F |
| 27906 | G | T | | ORF8 | nonsynonymous_V5F |
| 27909 | T | C | | ORF8 | nonsynonymous_F6L |
| 27915 | G | C | T | ORF8 | nonsynonymous_G8R; stopgain_G8X |
| 27916 | G | A | T | ORF8 | nonsynonymous_G8E; nonsynonymous_G8V |
| 27919 | T | C | | ORF8 | nonsynonymous_I9T |
| 27920 | C | T | | ORF8 | synonymous_I9I |
| 27925 | C | T | | ORF8 | nonsynonymous_T11I |
| 27936 | G | T | | ORF8 | nonsynonymous_A15S |
| 27937 | C | T | | ORF8 | nonsynonymous_A15V |
| 27942 | C | T | | ORF8 | nonsynonymous_H17Y |
| 27964 | C | T | | ORF8 | nonsynonymous_S24L |
| 27970 | C | T | | ORF8 | nonsynonymous_T26I |
| 27972 | C | T | | ORF8 | stopgain_Q27X |

| | | | | | |
|---|---|---|---|---|---|
| 27975 | C | T | | ORF8 | nonsynonymous_H28Y |
| 27977 | T | G | | ORF8 | nonsynonymous_H28Q |
| 27983 | A | T | | ORF8 | synonymous_P30P |
| 27984 | T | C | | ORF8 | nonsynonymous_Y31H |
| 27996 | G | T | | ORF8 | nonsynonymous_D35Y |
| 27998 | C | T | | ORF8 | synonymous_D35D |
| 27999 | C | T | | ORF8 | nonsynonymous_P36S |
| 28000 | C | T | | ORF8 | nonsynonymous_P36L |
| 28003 | G | T | | ORF8 | nonsynonymous_C37F |
| 28005 | C | T | | ORF8 | nonsynonymous_P38S |
| 28009 | T | C | | ORF8 | nonsynonymous_I39T |
| 28010 | T | C | | ORF8 | synonymous_I39I |
| 28018 | A | T | | ORF8 | nonsynonymous_Y42F |
| 28025 | A | G | | ORF8 | synonymous_K44K |
| 28032 | A | G | | ORF8 | nonsynonymous_I47V |
| 28045 | C | T | | ORF8 | nonsynonymous_A51V |
| 28046 | T | C | | ORF8 | synonymous_A51A |
| 28048 | G | T | | ORF8 | nonsynonymous_R52I |
| 28057 | C | T | | ORF8 | nonsynonymous_A55V |
| 28064 | A | G | | ORF8 | synonymous_L57L |
| 28069 | A | T | | ORF8 | nonsynonymous_E59V |
| 28076 | C | T | | ORF8 | synonymous_C61C |
| 28077 | G | C | T | ORF8 | nonsynonymous_V62L; nonsynonymous_V62L |
| 28079 | G | T | | ORF8 | synonymous_V62V |
| 28083 | G | T | | ORF8 | stopgain_E64X |
| 28086 | G | T | | ORF8 | nonsynonymous_A65S |
| 28087 | C | G | | ORF8 | nonsynonymous_A65G |
| 28089 | G | T | | ORF8 | nonsynonymous_G66C |
| 28090 | G | T | | ORF8 | nonsynonymous_G66V |
| 28093 | C | T | | ORF8 | nonsynonymous_S67F |

| | | | | | |
|---|---|---|---|---|---|
| 28099 | C | T | | ORF8 | nonsynonymous_S69L |
| 28105 | T | C | | ORF8 | nonsynonymous_I71T |
| 28109 | G | T | | ORF8 | nonsynonymous_Q72H |
| 28115 | C | T | | ORF8 | synonymous_I74I |
| 28122 | G | A | | ORF8 | nonsynonymous_G77S |
| 28123 | G | T | | ORF8 | nonsynonymous_G77V |
| 28144 | T | C | | ORF8 | nonsynonymous_L84S |
| 28146 | C | T | | ORF8 | nonsynonymous_P85S |
| 28166 | G | T | | ORF8 | nonsynonymous_Q91H |
| 28178 | G | A | | ORF8 | synonymous_L95L |
| 28198 | G | T | | ORF8 | nonsynonymous_C102F |
| 28208 | T | C | | ORF8 | synonymous_Y105Y |
| 28214 | C | T | | ORF8 | synonymous_D107D |
| 28222 | A | G | | ORF8 | nonsynonymous_E110G |
| 28231 | A | G | | ORF8 | nonsynonymous_D113G |
| 28232 | C | T | | ORF8 | synonymous_D113D |
| 28233 | G | A | T | ORF8 | nonsynonymous_V114I; nonsynonymous_V114F |
| 28234 | T | C | | ORF8 | nonsynonymous_V114A |
| 28237 | G | T | | ORF8 | nonsynonymous_R115L |
| 28240 | T | C | | ORF8 | nonsynonymous_V116A |
| 28248 | G | T | | ORF8 | nonsynonymous_D119Y |
| 28253 | C | T | G | ORF8 | synonymous_F120F; nonsynonymous_F120L |
| 28254 | A | C | | ORF8 | nonsynonymous_I121L |
| 28259 | A | G | | ORF8 | synonymous_X122X |
| 28261 | C | T | | intergenic | |
| 28262 | G | C | | intergenic | |
| 28268 | A | G | | intergenic | |
| 28286 | G | T | | N | stopgain_G5X |
| 28287 | G | A | | N | nonsynonymous_G5E |

| | | | | | |
|---|---|---|---|---|---|
| 28289 | C | A | T | N | nonsynonymous_P6T;<br>nonsynonymous_P6S |
| 28300 | G | T | | N | nonsynonymous_Q9H |
| 28301 | C | T | | N | stopgain_R10X |
| 28302 | G | A | | N | nonsynonymous_R10Q |
| 28303 | A | G | | N | synonymous_R10R |
| 28310 | C | T | G | N | nonsynonymous_P13S;<br>nonsynonymous_P13A |
| 28311 | C | T | | N | nonsynonymous_P13L |
| 28312 | C | T | | N | synonymous_P13P |
| 28313 | C | T | | N | nonsynonymous_R14C |
| 28315 | C | T | | N | synonymous_R14R |
| 28324 | T | C | | N | synonymous_F17F |
| 28325 | G | T | | N | nonsynonymous_G18C |
| 28326 | G | T | | N | nonsynonymous_G18V |
| 28329 | G | T | | N | nonsynonymous_G19V |
| 28336 | A | T | | N | synonymous_S21S |
| 28337 | G | T | | N | nonsynonymous_D22Y |
| 28338 | A | G | | N | nonsynonymous_D22G |
| 28340 | T | C | | N | nonsynonymous_S23P |
| 28341 | C | T | | N | nonsynonymous_S23L |
| 28342 | A | T | | N | synonymous_S23S |
| 28344 | C | A | T | N | nonsynonymous_T24N;<br>nonsynonymous_T24I |
| 28346 | G | T | | N | nonsynonymous_G25C |
| 28347 | G | A | T | N | nonsynonymous_G25D;<br>nonsynonymous_G25V |
| 28353 | A | C | | N | nonsynonymous_N27T |
| 28354 | C | T | | N | synonymous_N27N |
| 28357 | G | T | | N | nonsynonymous_Q28H |
| 28360 | T | G | | N | nonsynonymous_N29K |

| | | | | | |
|---|---|---|---|---|---|
| 28361 | G | A | | N | nonsynonymous_G30R |
| 28362 | G | T | | N | nonsynonymous_G30V |
| 28366 | A | C | | N | nonsynonymous_E31D |
| 28367 | C | T | | N | nonsynonymous_R32C |
| 28368 | G | T | | N | nonsynonymous_R32L |
| 28373 | G | T | | N | nonsynonymous_G34W |
| 28374 | G | T | | N | nonsynonymous_G34V |
| 28378 | G | T | | N | synonymous_A35A |
| 28380 | G | A | | N | nonsynonymous_R36Q |
| 28390 | A | G | | N | synonymous_Q39Q |
| 28392 | G | C | T | N | nonsynonymous_R40P; nonsynonymous_R40L |
| 28396 | G | A | | N | synonymous_R41R |
| 28403 | G | T | | N | nonsynonymous_G44C |
| 28406 | T | C | | N | synonymous_L45L |
| 28409 | C | G | | N | nonsynonymous_P46A |
| 28429 | G | T | | N | nonsynonymous_W52C |
| 28430 | T | C | | N | nonsynonymous_F53L |
| 28432 | C | T | | N | synonymous_F53F |
| 28435 | C | T | | N | synonymous_T54T |
| 28451 | G | C | | N | nonsynonymous_G60R |
| 28452 | G | A | T | N | nonsynonymous_G60D; nonsynonymous_G60V |
| 28456 | G | T | | N | nonsynonymous_K61N |
| 28457 | G | A | | N | nonsynonymous_E62K |
| 28460 | G | T | | N | nonsynonymous_D63Y |
| 28472 | C | T | | N | nonsynonymous_P67S |
| 28476 | G | T | | N | nonsynonymous_R68L |
| 28478 | G | T | | N | stopgain_G69X |
| 28481 | C | T | | N | stopgain_Q70X |
| 28491 | C | A | | N | nonsynonymous_P73Q |

| | | | | | |
|---|---|---|---|---|---|
| 28509 | G | T | | N | nonsynonymous_S79I |
| 28511 | C | T | | N | nonsynonymous_P80S |
| 28517 | G | T | | N | nonsynonymous_D82Y |
| 28528 | C | T | | N | synonymous_G85G |
| 28531 | C | T | | N | synonymous_Y86Y |
| 28534 | C | T | | N | synonymous_Y87Y |
| 28537 | A | G | | N | synonymous_R88R |
| 28541 | G | A | | N | nonsynonymous_A90T |
| 28543 | T | C | | N | synonymous_A90A |
| 28546 | C | T | | N | synonymous_T91T |
| 28548 | G | T | | N | nonsynonymous_R92I |
| 28556 | C | T | | N | nonsynonymous_R95C |
| 28557 | G | C | T | N | nonsynonymous_R95P; nonsynonymous_R95L |
| 28559 | G | T | | N | nonsynonymous_G96C |
| 28569 | G | C | | N | nonsynonymous_G99A |
| 28576 | G | A | C | N | nonsynonymous_M101I; nonsynonymous_M101I |
| 28580 | G | T | | N | nonsynonymous_D103Y |
| 28583 | C | T | | N | nonsynonymous_L104F |
| 28589 | C | T | | N | nonsynonymous_P106S |
| 28594 | A | G | | N | synonymous_R107R |
| 28602 | T | C | | N | nonsynonymous_F110S |
| 28603 | C | T | | N | synonymous_F110F |
| 28606 | C | G | | N | stopgain_Y111X |
| 28614 | G | T | | N | nonsynonymous_G114V |
| 28621 | G | A | T | N | synonymous_G116G; synonymous_G116G |
| 28625 | G | T | | N | stopgain_E118X |
| 28631 | G | A | | N | nonsynonymous_G120R |
| 28632 | G | T | | N | nonsynonymous_G120V |

| | | | | | |
|---|---|---|---|---|---|
| 28634 | C | T | | N | nonsynonymous_L121F |
| 28635 | T | C | | N | nonsynonymous_L121P |
| 28640 | T | C | | N | nonsynonymous_Y123H |
| 28651 | C | T | | N | synonymous_N126N |
| 28652 | A | T | | N | stopgain_K127X |
| 28655 | G | T | | N | nonsynonymous_D128Y |
| 28657 | C | T | G | N | synonymous_D128D; nonsynonymous_D128E |
| 28660 | C | T | | N | synonymous_G129G |
| 28662 | T | C | | N | nonsynonymous_I130T |
| 28674 | C | T | | N | nonsynonymous_A134V |
| 28677 | C | T | G | N | nonsynonymous_T135I; nonsynonymous_T135S |
| 28683 | G | T | | N | nonsynonymous_G137V |
| 28686 | C | T | | N | nonsynonymous_A138V |
| 28688 | T | C | | N | synonymous_L139L |
| 28690 | G | T | | N | nonsynonymous_L139F |
| 28695 | C | T | | N | nonsynonymous_T141I |
| 28697 | C | T | | N | nonsynonymous_P142S |
| 28706 | C | T | G | N | nonsynonymous_H145Y; nonsynonymous_H145D |
| 28708 | C | T | | N | synonymous_H145H |
| 28709 | A | T | G | N | nonsynonymous_I146F; nonsynonymous_I146V |
| 28713 | G | T | | N | nonsynonymous_G147V |
| 28714 | C | T | | N | synonymous_G147G |
| 28718 | C | T | | N | nonsynonymous_R149C |
| 28724 | C | T | | N | nonsynonymous_P151S |
| 28727 | G | T | | N | nonsynonymous_A152S |
| 28728 | C | T | | N | nonsynonymous_A152V |
| 28729 | T | A | | N | synonymous_A152A |

| | | | | | |
|---|---|---|---|---|---|
| 28737 | C | T | | N | nonsynonymous_A155V |
| 28739 | G | T | | N | nonsynonymous_A156S |
| 28743 | T | C | | N | nonsynonymous_I157T |
| 28744 | C | T | | N | synonymous_I157I |
| 28748 | C | T | | N | synonymous_L159L |
| 28751 | C | T | G | N | stopgain_Q160X; nonsynonymous_Q160E |
| 28754 | C | T | | N | nonsynonymous_L161F |
| 28755 | T | C | | N | nonsynonymous_L161P |
| 28756 | T | C | | N | synonymous_L161L |
| 28760 | C | A | | N | nonsynonymous_Q163K |
| 28767 | C | T | | N | nonsynonymous_T165I |
| 28770 | C | T | | N | nonsynonymous_T166I |
| 28773 | T | C | | N | nonsynonymous_L167S |
| 28775 | C | T | | N | nonsynonymous_P168S |
| 28776 | C | A | T | N | nonsynonymous_P168Q; nonsynonymous_P168L |
| 28783 | C | T | | N | synonymous_G170G |
| 28790 | G | C | | N | nonsynonymous_A173P |
| 28793 | G | T | | N | stopgain_E174X |
| 28794 | A | C | | N | nonsynonymous_E174A |
| 28798 | G | T | | N | synonymous_G175G |
| 28808 | G | A | | N | nonsynonymous_G179S |
| 28810 | C | T | | N | synonymous_G179G |
| 28811 | A | G | | N | nonsynonymous_S180G |
| 28812 | G | T | | N | nonsynonymous_S180I |
| 28814 | C | T | G | N | stopgain_Q181X; nonsynonymous_Q181E |
| 28821 | C | A | | N | nonsynonymous_S183Y |
| 28824 | C | G | | N | nonsynonymous_S184C |
| 28826 | C | T | | N | nonsynonymous_R185C |

| | | | | | |
|---|---|---|---|---|---|
| 28827 | G | A | T | N | nonsynonymous_R185H;<br>nonsynonymous_R185L |
| 28830 | C | A | T | N | nonsynonymous_S186Y;<br>nonsynonymous_S186F |
| 28831 | C | T | | N | synonymous_S186S |
| 28833 | C | T | | N | nonsynonymous_S187L |
| 28835 | T | C | | N | nonsynonymous_S188P |
| 28836 | C | T | | N | nonsynonymous_S188L |
| 28839 | G | T | | N | nonsynonymous_R189L |
| 28841 | A | G | | N | nonsynonymous_S190G |
| 28842 | G | T | | N | nonsynonymous_S190I |
| 28844 | C | T | | N | nonsynonymous_R191C |
| 28845 | G | T | | N | nonsynonymous_R191L |
| 28849 | C | T | | N | synonymous_N192N |
| 28851 | G | T | | N | nonsynonymous_S193I |
| 28853 | T | G | | N | nonsynonymous_S194A |
| 28854 | C | T | | N | nonsynonymous_S194L |
| 28858 | A | G | | N | synonymous_R195R |
| 28859 | A | C | | N | nonsynonymous_N196H |
| 28860 | A | T | | N | nonsynonymous_N196I |
| 28861 | T | C | | N | synonymous_N196N |
| 28862 | T | C | | N | nonsynonymous_S197P |
| 28863 | C | T | | N | nonsynonymous_S197L |
| 28866 | C | T | | N | nonsynonymous_T198I |
| 28868 | C | A | T | N | nonsynonymous_P199T;<br>nonsynonymous_P199S |
| 28869 | C | T | | N | nonsynonymous_P199L |
| 28870 | A | G | | N | synonymous_P199P |
| 28873 | C | T | | N | synonymous_G200G |
| 28877 | A | G | | N | nonsynonymous_S202G |
| 28878 | G | A | | N | nonsynonymous_S202N |

| | | | | | |
|---|---|---|---|---|---|
| 28881 | G | A | | N | nonsynonymous_R203K |
| 28882 | G | A | T | N | synonymous_R203R; nonsynonymous_R203S |
| 28883 | G | C | | N | nonsynonymous_G204R |
| 28885 | A | G | | N | synonymous_G204G |
| 28887 | C | T | | N | nonsynonymous_T205I |
| 28892 | C | T | | N | nonsynonymous_P207S |
| 28893 | C | T | | N | nonsynonymous_P207L |
| 28896 | C | T | G | N | nonsynonymous_A208V; nonsynonymous_A208G |
| 28898 | A | G | | N | nonsynonymous_R209G |
| 28899 | G | A | | N | nonsynonymous_R209K |
| 28902 | T | C | | N | nonsynonymous_M210T |
| 28903 | G | C | | N | nonsynonymous_M210I |
| 28904 | G | T | | N | nonsynonymous_A211S |
| 28905 | C | T | | N | nonsynonymous_A211V |
| 28908 | G | T | | N | nonsynonymous_G212V |
| 28922 | G | T | | N | nonsynonymous_A217S |
| 28924 | T | C | | N | synonymous_A217A |
| 28928 | C | T | | N | nonsynonymous_L219F |
| 28932 | C | T | | N | nonsynonymous_A220V |
| 28933 | T | C | | N | synonymous_A220A |
| 28939 | G | T | | N | synonymous_L222L |
| 28948 | C | T | | N | synonymous_D225D |
| 28951 | A | T | | N | nonsynonymous_R226S |
| 28955 | A | T | | N | nonsynonymous_N228Y |
| 28957 | C | T | | N | synonymous_N228N |
| 28958 | C | T | | N | stopgain_Q229X |
| 28960 | G | A | | N | synonymous_Q229Q |
| 28961 | C | T | | N | nonsynonymous_L230F |
| 28963 | T | A | | N | synonymous_L230L |

| | | | | | |
|---|---|---|---|---|---|
| 28975 | G | T | | N | nonsynonymous_M234I |
| 28976 | T | C | | N | nonsynonymous_S235P |
| 28979 | G | T | | N | nonsynonymous_G236C |
| 28980 | G | T | | N | nonsynonymous_G236V |
| 28991 | C | T | | N | stopgain_Q240X |
| 28993 | A | C | | N | nonsynonymous_Q240H |
| 28994 | C | T | | N | stopgain_Q241X |
| 28995 | A | G | | N | nonsynonymous_Q241R |
| 28997 | C | T | | N | stopgain_Q242X |
| 29000 | G | T | | N | nonsynonymous_G243C |
| 29002 | C | T | | N | synonymous_G243G |
| 29003 | C | A | T | N | nonsynonymous_Q244K; stopgain_Q244X |
| 29004 | A | C | | N | nonsynonymous_Q244P |
| 29005 | A | G | | N | synonymous_Q244Q |
| 29007 | C | T | G | N | nonsynonymous_T245I; nonsynonymous_T245S |
| 29013 | C | T | | N | nonsynonymous_T247I |
| 29025 | C | T | | N | nonsynonymous_A251V |
| 29027 | G | T | | N | nonsynonymous_A252S |
| 29028 | C | T | | N | nonsynonymous_A252V |
| 29030 | G | T | | N | stopgain_E253X |
| 29032 | G | T | | N | nonsynonymous_E253D |
| 29033 | G | A | T | N | nonsynonymous_A254T; nonsynonymous_A254S |
| 29034 | C | T | G | N | nonsynonymous_A254V; nonsynonymous_A254G |
| 29035 | T | C | G | N | synonymous_A254A; synonymous_A254A |
| 29036 | T | C | | N | nonsynonymous_S255P |
| 29037 | C | T | | N | nonsynonymous_S255F |

| | | | | | |
|---|---|---|---|---|---|
| 29040 | A | G | | N | nonsynonymous_K256R |
| 29052 | A | T | | N | nonsynonymous_Q260L |
| 29061 | C | T | | N | nonsynonymous_T263I |
| 29077 | C | T | | N | synonymous_Y268Y |
| 29078 | A | T | G | N | nonsynonymous_N269Y;<br>nonsynonymous_N269D |
| 29085 | C | T | | N | nonsynonymous_T271I |
| 29087 | C | T | | N | stopgain_Q272X |
| 29095 | C | T | | N | synonymous_F274F |
| 29097 | G | T | | N | nonsynonymous_G275V |
| 29105 | G | C | | N | nonsynonymous_G278R |
| 29118 | C | T | | N | nonsynonymous_T282I |
| 29119 | C | T | | N | synonymous_T282T |
| 29134 | G | A | | N | synonymous_G287G |
| 29135 | G | T | | N | nonsynonymous_D288Y |
| 29144 | C | T | | N | synonymous_L291L |
| 29148 | T | C | | N | nonsynonymous_I292T |
| 29157 | G | T | | N | nonsynonymous_G295V |
| 29162 | G | T | | N | nonsynonymous_D297Y |
| 29167 | C | T | | N | synonymous_Y298Y |
| 29177 | C | T | | N | nonsynonymous_P302S |
| 29186 | G | T | | N | nonsynonymous_A305S |
| 29187 | C | T | | N | nonsynonymous_A305V |
| 29188 | A | G | | N | synonymous_A305A |
| 29200 | C | T | | N | synonymous_P309P |
| 29202 | G | C | T | N | nonsynonymous_S310T;<br>nonsynonymous_S310I |
| 29203 | C | T | | N | synonymous_S310S |
| 29208 | C | T | | N | nonsynonymous_S312L |
| 29211 | C | T | | N | nonsynonymous_A313V |
| 29212 | G | T | | N | synonymous_A313A |

| | | | | | |
|---|---|---|---|---|---|
| 29218 | C | T | | N | synonymous_F315F |
| 29220 | G | A | | N | nonsynonymous_G316E |
| 29227 | G | A | T | N | synonymous_S318S;<br>synonymous_S318S |
| 29228 | C | T | | N | nonsynonymous_R319C |
| 29236 | C | T | | N | synonymous_G321G |
| 29240 | G | T | | N | stopgain_E323X |
| 29242 | A | T | | N | nonsynonymous_E323D |
| 29247 | C | A | T | N | nonsynonymous_T325K;<br>nonsynonymous_T325I |
| 29253 | C | T | | N | nonsynonymous_S327L |
| 29254 | G | T | | N | synonymous_S327S |
| 29259 | C | T | | N | nonsynonymous_T329M |
| 29260 | G | A | T | N | synonymous_T329T;<br>synonymous_T329T |
| 29266 | G | T | | N | nonsynonymous_L331F |
| 29268 | C | T | | N | nonsynonymous_T332I |
| 29274 | C | T | | N | nonsynonymous_T334I |
| 29284 | C | T | | N | synonymous_I337I |
| 29296 | C | T | | N | synonymous_D341D |
| 29303 | C | T | | N | nonsynonymous_P344S |
| 29311 | C | A | | N | nonsynonymous_F346L |
| 29313 | A | G | | N | nonsynonymous_K347R |
| 29315 | G | T | | N | nonsynonymous_D348Y |
| 29321 | G | T | | N | nonsynonymous_V350F |
| 29329 | G | T | | N | nonsynonymous_L352F |
| 29330 | C | T | | N | synonymous_L353L |
| 29332 | G | T | | N | synonymous_L353L |
| 29348 | G | C | T | N | nonsynonymous_A359P;<br>nonsynonymous_A359S |
| 29349 | C | T | | N | nonsynonymous_A359V |

| | | | | | |
|---|---|---|---|---|---|
| 29353 | C | T | | N | synonymous_Y360Y |
| 29358 | C | T | | N | nonsynonymous_T362I |
| 29360 | T | C | | N | nonsynonymous_F363L |
| 29364 | C | T | | N | nonsynonymous_P364L |
| 29370 | C | T | | N | nonsynonymous_T366I |
| 29374 | G | A | T | N | synonymous_E367E;<br>nonsynonymous_E367D |
| 29386 | C | A | T | N | nonsynonymous_D371E;<br>synonymous_D371D |
| 29392 | G | T | | N | nonsynonymous_K373N |
| 29395 | G | C | T | N | nonsynonymous_K374N;<br>nonsynonymous_K374N |
| 29399 | G | T | | N | nonsynonymous_A376S |
| 29409 | C | A | T | N | nonsynonymous_T379N;<br>nonsynonymous_T379I |
| 29413 | A | T | G | N | nonsynonymous_Q380H;<br>synonymous_Q380Q |
| 29414 | G | T | | N | nonsynonymous_A381S |
| 29418 | T | C | | N | nonsynonymous_L382S |
| 29420 | C | T | | N | nonsynonymous_P383S |
| 29422 | G | A | | N | synonymous_P383P |
| 29429 | C | T | | N | stopgain_Q386X |
| 29440 | G | T | | N | nonsynonymous_Q389H |
| 29441 | C | A | T | N | nonsynonymous_Q390K;<br>stopgain_Q390X |
| 29449 | G | T | | N | synonymous_V392V |
| 29450 | A | T | | N | nonsynonymous_T393S |
| 29451 | C | T | | N | nonsynonymous_T393I |
| 29462 | G | T | | N | nonsynonymous_A397S |
| 29465 | G | C | | N | nonsynonymous_A398P |
| 29466 | C | T | | N | nonsynonymous_A398V |

| | | | | | |
|---|---|---|---|---|---|
| 29468 | G | C | T | N | nonsynonymous_D399H; nonsynonymous_D399Y |
| 29469 | A | T | | N | nonsynonymous_D399V |
| 29474 | G | T | | N | nonsynonymous_D401Y |
| 29477 | G | T | | N | nonsynonymous_D402Y |
| 29482 | C | T | | N | synonymous_F403F |
| 29483 | T | G | | N | nonsynonymous_S404A |
| 29492 | T | C | | N | synonymous_L407L |
| 29498 | C | T | | N | stopgain_Q409X |
| 29507 | A | C | | N | nonsynonymous_S412R |
| 29522 | A | G | | N | nonsynonymous_T417A |
| 29523 | C | T | | N | nonsynonymous_T417I |
| 29525 | C | T | | N | stopgain_Q418X |
| 29527 | G | A | | N | synonymous_Q418Q |
| 29528 | G | T | | N | nonsynonymous_A419S |
| 29531 | T | A | | N | stoploss_X420K |
| 29535 | C | T | | intergenic | |
| 29540 | G | A | T | intergenic | |
| 29541 | C | T | | intergenic | |
| 29554 | G | T | | intergenic | |
| 29555 | C | T | | intergenic | |
| 29563 | C | T | | ORF10 | synonymous_G2G |
| 29578 | C | T | | ORF10 | synonymous_F7F |
| 29579 | G | T | | ORF10 | nonsynonymous_A8S |
| 29580 | C | A | T | ORF10 | nonsynonymous_A8D; nonsynonymous_A8V |
| 29581 | T | A | | ORF10 | synonymous_A8A |
| 29585 | C | T | | ORF10 | nonsynonymous_P10S |
| 29586 | C | T | | ORF10 | nonsynonymous_P10L |
| 29587 | G | A | | ORF10 | synonymous_P10P |
| 29612 | T | A | | ORF10 | nonsynonymous_C19S |

| | | | | | |
|---|---|---|---|---|---|
| 29614 | C | T | | ORF10 | synonymous_C19C |
| 29618 | A | T | | ORF10 | nonsynonymous_M21L |
| 29623 | T | C | | ORF10 | synonymous_N22N |
| 29625 | C | T | | ORF10 | nonsynonymous_S23F |
| 29627 | C | T | | ORF10 | nonsynonymous_R24C |
| 29633 | T | C | | ORF10 | nonsynonymous_Y26H |
| 29635 | C | T | | ORF10 | synonymous_Y26Y |
| 29639 | G | T | | ORF10 | nonsynonymous_A28S |
| 29640 | C | T | | ORF10 | nonsynonymous_A28V |
| 29642 | C | T | | ORF10 | stopgain_Q29X |
| 29654 | G | A | T | ORF10 | nonsynonymous_V33I; nonsynonymous_V33F |
| 29666 | C | T | | ORF10 | nonsynonymous_L37F |
| 29668 | C | T | | ORF10 | synonymous_L37L |
| 29670 | C | T | | ORF10 | nonsynonymous_T38I |
| 29672 | T | C | | ORF10 | stoploss_X39Q |
| 29675 | C | T | | three_prime_UTR | |
| 29679 | C | T | | three_prime_UTR | |
| 29683 | A | G | | three_prime_UTR | |
| 29686 | C | T | | three_prime_UTR | |
| 29690 | G | T | | three_prime_UTR | |
| 29692 | G | T | | three_prime_UTR | |
| 29700 | A | G | | three_prime_UTR | |
| 29703 | G | T | | three_prime_UTR | |
| 29705 | G | T | | three_prime_UTR | |
| 29706 | G | T | | three_prime_UTR | |
| 29708 | C | T | | three_prime_UTR | |
| 29711 | G | A | T | three_prime_UTR | |
| 29715 | G | T | | three_prime_UTR | |
| 29717 | G | T | | three_prime_UTR | |
| 29718 | C | A | T | three_prime_UTR | |

| | | | | |
|---|---|---|---|---|
| 29721 | C | T | | three_prime_UTR |
| 29722 | C | T | | three_prime_UTR |
| 29724 | C | T | | three_prime_UTR |
| 29729 | T | C | | three_prime_UTR |
| 29730 | C | T | G | three_prime_UTR |
| 29731 | A | G | | three_prime_UTR |
| 29732 | C | T | | three_prime_UTR |
| 29733 | C | T | | three_prime_UTR |
| 29734 | G | C | T | three_prime_UTR |
| 29736 | G | T | | three_prime_UTR |
| 29737 | G | T | | three_prime_UTR |
| 29738 | C | T | | three_prime_UTR |
| 29739 | C | T | | three_prime_UTR |
| 29741 | C | T | | three_prime_UTR |
| 29742 | G | A | T | three_prime_UTR |
| 29743 | C | T | | three_prime_UTR |
| 29745 | G | T | | three_prime_UTR |
| 29746 | A | T | | three_prime_UTR |
| 29747 | G | T | | three_prime_UTR |
| 29750 | C | T | | three_prime_UTR |
| 29751 | G | T | | three_prime_UTR |
| 29754 | C | T | | three_prime_UTR |
| 29758 | T | A | | three_prime_UTR |
| 29761 | A | C | | three_prime_UTR |
| 29762 | C | T | | three_prime_UTR |
| 29763 | A | T | | three_prime_UTR |
| 29764 | G | T | | three_prime_UTR |
| 29769 | C | T | | three_prime_UTR |
| 29773 | G | T | | three_prime_UTR |
| 29774 | C | T | | three_prime_UTR |
| 29776 | A | G | | three_prime_UTR |
| 29778 | G | A | T | three_prime_UTR |

| | | | | |
|---|---|---|---|---|
| 29779 | G | T | | three_prime_UTR |
| 29781 | G | T | | three_prime_UTR |
| 29784 | C | T | | three_prime_UTR |
| 29786 | G | C | | three_prime_UTR |
| 29788 | C | T | | three_prime_UTR |
| 29793 | T | C | | three_prime_UTR |
| 29800 | G | C | | three_prime_UTR |
| 29802 | C | T | | three_prime_UTR |
| 29803 | C | T | | three_prime_UTR |
| 29804 | T | A | | three_prime_UTR |
| 29810 | G | T | | three_prime_UTR |
| 29825 | G | A | C | three_prime_UTR |
| 29835 | C | T | G | three_prime_UTR |
| 29836 | C | T | | three_prime_UTR |
| 29837 | C | T | | three_prime_UTR |
