## Extended Data Table for "Worldwide tracing of mutations and the evolutionary dynamics of SARS-CoV-2": Extended Data Table 4.docx

|  | **Genome Position** | **Mutation Rate** (per site per year) | **HPD** |
| --- | --- | --- | --- |
| SARS-Cov-2 | 1 – 29,903 | 1.2×10^-3^ | 6.14×10^-4^ – 1.87×10^-3^ |
| Block1 | 200 – 4,300 | 2.95×10^-3^ | 6.55×10^-4^ – 5.47×10^-3^ |
| Block2 | 5,800 – 7,500 | 3.44×10^-3^ | 4.15×10^-4^ – 7.39×10^-3^ |
| Block3 | 11,900 – 13,000 | 3.94×10^-3^ | 2.95×10^-4^ – 1 × 10^-2^ |
| Block4 | 17,700 – 22,700 | 3.23×10^-3^ | 3.75×10^-4^ – 6.91×10^-3^ |
| Block5 | 24,300 – 27,800 | 3.35×10^-3^ | 6.08×10^-4^ – 6.74×10^-3^ |

**Extended Data Table 4. Estimates of the substitution rate for the SARS-CoV-2 genome and the regions with high numbers of mutations in protein genes. HPD: 95% highest posterior density interval.**
