## Extended Data Table for "Worldwide tracing of mutations and the evolutionary dynamics of SARS-CoV-2": Extended Data Table 5.docx

| **Gene** | **π_n_** | **π_s_** | **π_n_/π_s_** |
| --- | --- | --- | --- |
| ORF1a | 0.00017 ± 0.00004 | 0.00042 ± 0.00015 | 0.410 |
| ORF1b | 0.00018 ± 0.00004 | 0.00044 ± 0.00011 | 0.420^**^ |
| S | 0.00023 ± 0.00014 | 0.00028 ± 0.00006 | 0.835 |
| ORF3a | 0.00114 ± 0.00070 | 0.00028 ± 0.00006 | 4.087 |
| E | 0.00011 ± 0.00003 | 0.00019 ± 0.00006 | 0.564 |
| M | 0.00017 ± 0.00008 | 0.00060 ± 0.00022 | 0.284^*^ |
| ORF6 | 0.00014 ± 0.00004 | 0.00040 ± 0.00022 | 0.342 |
| ORF7a | 0.00015 ± 0.00004 | 0.00020 ± 0.00005 | 0.783 |
| ORF7b | 0.00010 ± 0.00002 | 0.00037 ± 0.00015 | 0.278^*^ |
| ORF8 | 0.00119 ± 0.00080 | 0.00014 ± 0.00004 | 8.575 |
| N | 0.00098 ± 0.00048 | 0.00150 ± 0.00108 | 0.655 |
| ORF10 | 0.00019 ± 0.00005 | 0.00030 ± 0.00011 | 0.642 |
| **π_n_= π_s_**: ^*^p < 0.05, ^**^p < 0.01. | | | |

**Extended Data Table 5. Mean nonsynonymous and synonymous nucleotide diversity of each ORF.**
